# ERCC1-XPF Interacts with Topoisomerase IIβ to Facilitate the Repair of Activity-induced DNA Breaks

**DOI:** 10.1101/2020.01.03.892703

**Authors:** Georgia Chatzinikolaou, Kalliopi Stratigi, Kyriacos Agathangelou, Maria Tsekrekou, Evi Goulielmaki, Ourania Chatzidoukaki, Katerina Gkirtzimanaki, Tamara Aid-Pavlidis, Michalis Aivaliotis, Pavlos Pavlidis, Ioannis Tsamardinos, Pantelis Topalis, Britta A. M. Bouwman, Nicola Crosetto, Janine Altmüller, George A. Garinis

## Abstract

Type II DNA Topoisomerases (TOP II) generate transient double-strand DNA breaks (DSBs) to resolve topological constraints during transcription. Using genome-wide mapping of DSBs and functional genomics approaches, we show that, in the absence of exogenous genotoxic stress, transcription leads to DSB accumulation and to the recruitment of the structure-specific ERCC1-XPF endonuclease on active gene promoters. Instead, we find that the complex is released from regulatory or gene body elements in UV-irradiated cells. Abrogation of ERCC1 or re-ligation blockage of TOP II-mediated DSBs aggravates the accumulation of transcription-associated γH2Ax and 53BP1 foci, which dissolve when TOP II-mediated DNA cleavage is inhibited. An *in vivo* biotinylation tagging strategy coupled to a high-throughput proteomics approach reveals that ERCC1-XPF interacts with TOP IIβ and the CTCF/cohesin complex, which co-localize with the heterodimer on DSBs. Together; our findings provide a rational explanation for the remarkable clinical heterogeneity seen in human disorders with ERCC1-XPF defects.

## Introduction

Transcription requires the concerted action of basal transcription factors, sequence-specific DNA binding proteins, chromatin remodeling and modification enzymes to enable the synthesis of the primary transcript (Ohler and Wassarman, 2010). Besides transcription-blocking DNA insults, the process of mRNA synthesis leads to transcription-associated recombination or rearrangements that occur during robust shifts in transcription demands threatening cell viability (Gaillard and Aguilera, 2016). To ensure that genome integrity is preserved and that transcription is not compromised, cells employ a battery of partially overlapping DNA repair systems aimed at counteracting DNA damage and restore DNA to its native form (Hoeijmakers, 2001).

ERCC1-XPF is a two subunit structure-specific endonuclease where XPF contains the nuclease domain of the complex and ERCC1 is required for subsequent nuclease activity (Tripsianes et al., 2005). The complex is essential for incising DNA 5′ to the DNA lesion during nucleotide excision repair (NER) (Gregg et al., 2011), a highly conserved mechanism that removes helical distortions throughout the genome i.e. global genome NER (GG-NER) or selectively from the transcribed strand of active genes i.e. transcription-coupled NER (TC-NER) (Hanawalt, 2002; Marteijn et al., 2014; Sijbers et al., 1996; van Duin et al., 1986). Besides NER, ERCC1-XPF is required for the repair of DNA interstrand cross links (DNA ICLs) (Fisher et al., 2008; Kuraoka et al., 2000) and for removing non-homologous 3′ single-stranded tails from DNA ends during DSB repair by homologous recombination (HR) or by alternative non-homologous end joining (NHEJ), where short stretches of homology are utilized to join two broken DNA ends (Adair et al., 2000; Ahmad et al., 2008; Al-Minawi et al., 2008; Ma et al., 2003; Sargent et al., 1997). Furthermore, ERCC1-XPF is involved in telomere maintenance (Munoz et al., 2005; Zhu et al., 2003) and, recently; it was shown to play a role in a sub-pathway of long-patch base excision repair involving 5′ gap formation (Woodrick et al., 2017).

In humans, mutations in ERCC1-XPF lead to Xeroderma Pigmentosum (XP; affected proteins: XPA through XPG), Cockayne Syndrome (CS; affected proteins: CSA, CSB, UVSSA, XPB, XPD, XPF, TTDA and certain mutations in the gene encoding XPG) or Fanconi Anemia, whose clinical outcomes are exceptionally diverse (Bootsma, 1998; Bootsma, 2001; Kashiyama et al., 2013; Mori et al., 2018). For instance, patients with mutations in ERCC1 manifest a severe form of CS named cerebro-oculo-facio-skeletal syndrome (COFS) (Gregg et al., 2011; Jaspers et al., 2007). Instead the great majority of XP-F patients present with mild symptoms of XP, including sun sensitivity, freckling of the skin, and basal or squamous cell carcinomas that typically occur at later stages in life (Gregg et al., 2011). Mice carrying inborn defects in *Ercc1* and *Xpf* genes fully recapitulate the severe growth retardation and premature onset of heterogeneous pathological symptoms seen in patients with defects in the corresponding genes (Niedernhofer et al., 2006; Tian et al., 2004).

In addition to DNA repair, recent studies have shown that NER factors, including ERCC1-XPF play a role in the regulation of gene expression (Le May et al., 2010a; Le May et al., 2010c), chromatin looping (Chatzinikolaou et al., 2017; Le May et al., 2012), the transcriptional reprogramming of pluripotent stem cells (Fong et al., 2011) and the fine-tuning of growth-promoting genes during postnatal development (Kamileri et al., 2012d). However, for ERCC1-XPF, no solid evidence exists on (i.) how the endonuclease complex is functionally involved in these processes, (ii.) the associated protein factors involved, or (iii.) the *in vivo* relevance of the complex to human disorders. Using an *in vivo* biotinylation tagging strategy coupled to genome-wide mapping of DNA DSBs, functional genomics and proteomics approaches, we find that ERCC1-XPF interacts with TOP IIβ and the CTCF/cohesin complex on promoters to facilitate the repair of activity-induced DNA DSBs. The findings provide a rational basis to explain how inborn defects in ERCC1 and/or XPF lead to the remarkable heterogeneity of tissue-specific, pathological features seen in corresponding human progeroid disorders.

## Results

### Transcription initiation but not elongation triggers DNA damage response signalling (DDR)

Transcription is often so abrupt that leads to genome instability (Butuci et al., 2015). To further test this, we treated primary mouse embryonic fibroblasts (MEFs) with all-trans-retinoic acid (tRA), a pleiotropic factor known to activate transcription during cell differentiation and embryonic development (Bastien and Rochette-Egly, 2004), in the absence of exogenous genotoxic stress. We find an increase in the formation of γH2AX and 53BP1 foci, two well-established DNA damage markers, and a mild but detectable increase in FANCI protein levels in the nucleoplasm of tRA-treated cells (Niraj et al., 2019); the response was comparable to that seen when cells are treated with the potent genotoxin mitomycin (**Figure 1A** and **Supplementary Figure S1A**; as indicated). Ataxia telangiectasia mutated (ATM) and Rad3 related (ATR) kinases are central mediators of the DNA damage checkpoint (Awasthi et al., 2015). To test whether transcription triggers canonical DDR signaling, tRA-treated MEFs were cultured in the presence of KU-55933, an ATM inhibitor (Ding et al., 2006) or ATR inhibitor NU6027, which inhibits ATR kinase without interfering with irradiation-induced autophosphorylation of DNA-dependent protein kinase or ATM. We find that upon transcription induction, inhibition of ATM (tRA/ATMi cells) but not ATR (tRA/ATRi cells) abrogates the formation of γH2AX and 53BP1 foci in tRA-treated MEFs (**Figure 1B**; as indicated). The accumulation of γH2AX and 53BP1 foci was also observed in tRA-treated MEFs induced into quiescence by serum starvation (**Supplementary Figure S1B**). To test whether the pronounced formation of γH2AX and 53BP1 foci occurs predominantly during transcription initiation or elongation, we treated tRA-treated MEFs with 5,6-Dichlorobenzimidazole 1-β-D-ribofuranoside (DRB), a selective inhibitor of transcription elongation by RNA polymerase II (RNAPII) (Yankulov et al., 1995) or with triptolide (TPL), a small molecule XPB/TFIIH inhibitor that blocks transcription initiation (Chen et al., 2015). Unlike in cells treated with DRB, we find that the number of γH2AX^+^ 53BP1^+^ MEFs decreases substantially when tRA-treated cells are cultured in the presence of TPL (**Figure 1C**), indicating that transcription initiation rather than elongation is the primary instigator of the DDR in tRA-treated MEFs. Earlier studies have shown that, in addition to DNA repair, NER factors are recruited to active promoters and facilitate chromatin modification for transcription in the absence of exogenous genotoxic insults (de Boer et al., 2002; Fong et al., 2011; Laine and Egly, 2006; Le May et al., 2010b). To test the relevance of transcription-associated DDR to NER mutant cells, we exposed cells defective in TC-NER (*Csb^m/m^*), GG-NER (*Xpc^−/−^*) or in GG- and TC-NER subpathways of NER (*Xpa^−/−^* and *Ercc1^−/−^* cells) to tRA treatment. Unlike in *Csb^m/m^*, *Xpc^−/−^* or *Xpa^−/−^* cells, we find that transcription induction leads to the pronounced accumulation of γH2AX and 53BP1 foci in tRA-treated *Ercc1^−/−^* cells (**Figure 1D** and **Supplementary Figure S2A**), indicating that the observed DDR is restricted to ERCC1-defective cells and that it does not involve DNA lesions typically associated with NER. Consistent with the previously observed transcription-associated DDR in wt. cells, we detect the presence of γH2AX^+^ 53BP1^+^ cells in developing wt. mouse P15 cerebella and livers, where transcription dynamics are high. As expected, the percentage of γH2AX^+^ 53BP1^+^ cells increases substantially in P15 *Ercc1^−/−^* animals (**Figure 1E**). Taken together, our findings indicate that, in the absence of exogenous genotoxic stimuli, transcription initiation, but not elongation, triggers an ATM-dependent DDR that is further pronounced in cells defective in the ERCC1-XPF endonuclease.

**Figure 1.**
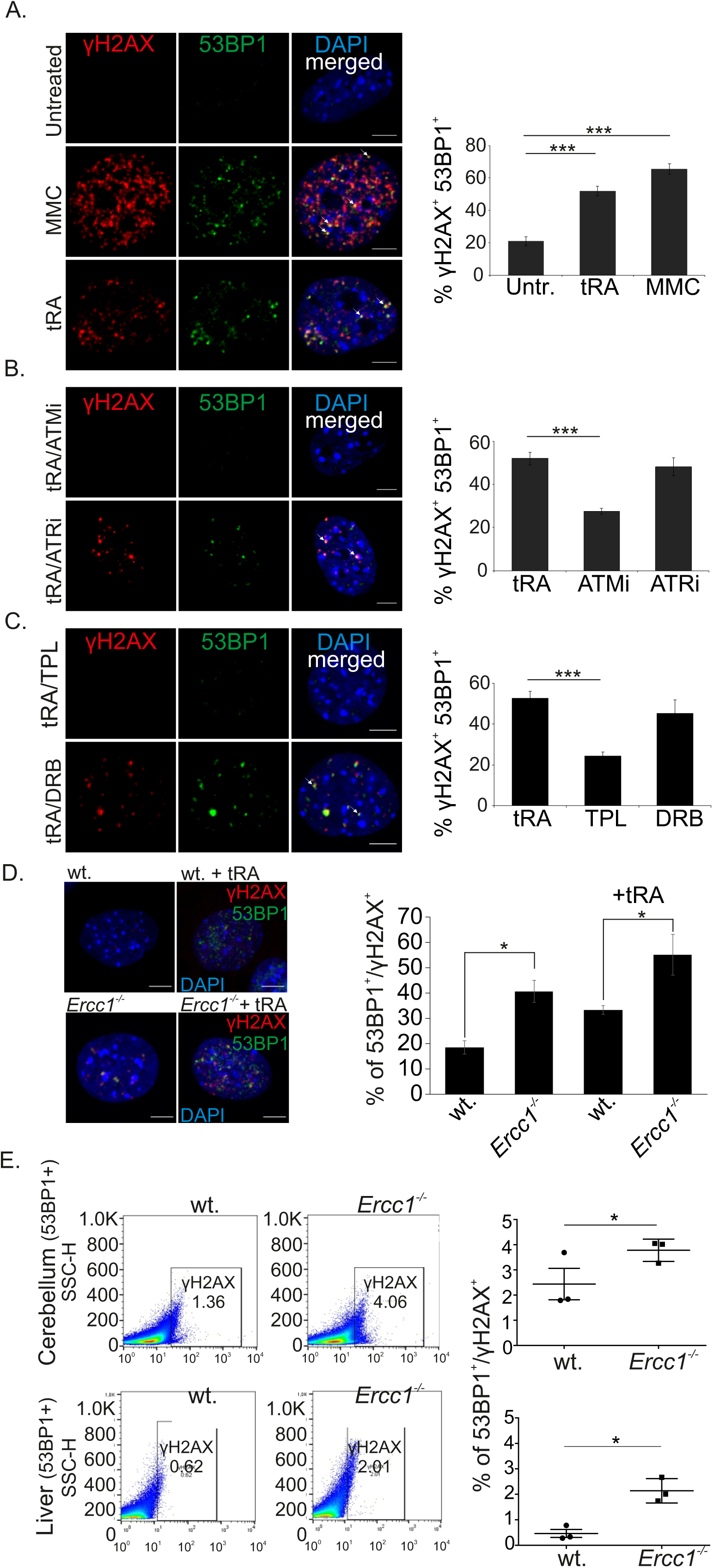
Transcription initiation triggers DDR signaling. **(A)**. Immunofluorescence detection of γH2AX and 53BP1 (white arrowheads) in primary wt. MEFs cultured upon basal conditions, exposure to MMC or treatment with tRA (as indicated). **(B)**. Immunofluorescence detection of γH2AX and 53BP1 in primary tRA-treated wt. MEFs cultured in the presence of ATM (ATMi) or ATR (ATRi) inhibitors (as indicated). **(C)**. Immunofluorescence detection of γH2AX and 53BP1 in primary tRA-treated wt. MEFs cultured in the presence of triptolide (TPL), an XPB/TFIIH inhibitor, that blocks transcription initiation or DRB, a selective inhibitor of transcription elongation by RNA polymerase II (as indicated). **(D)**. Immunofluorescence detection of γH2AX and 53BP1 in primary untreated and tRA-treated wt. and *Ercc1^−/−^* primary MEFs (see Supplementary Figure S2A). The graph represent the number of γH2AX^+^ 53BP1^+^ cells in untreated and tRA-treated wt. and *Ercc1^−/−^* primary MEFs. **(E)**. FACS representative plots and respective graphs of γH2AX^+^ 53BP1^+^ cells, in the P15 cerebellum and liver of wt. and *Ercc1*^−/−^ animals (as indicated). Error bars indicate S.E.M. among n ≥ three biological replicates. Asterisk indicates the significance set at p-value: *≤0.05, **≤ 0.01, ***≤ 0.001 (two-tailed Student’s t-test). Grey line is set at 10 μm scale.

### Transcription activation triggers the genome-wide recruitment of XPF to promoters

The relevance of transcription-associated DDR to tRA-treated *Ercc1^−/−^* cells prompted us to evaluate, genome-wide, the functional role of the ERCC1-XPF complex in transcription-associated DNA damage. To do so, we crossed homozygous av*Xpf^+/+^* knockin mice expressing the NER structure-specific endonuclease XPF fused with a 15 amino acid (aa) Tandem Affinity Purification (TAP)-tag biotinylatable sequence and a 3X FLAG tag with mice broadly expressing the HA-tagged bacterial BirA biotin ligase transgene (BirA) to generate biotin-tagged XPF (bXPF) animals (Chatzinikolaou et al., 2017) (**Figure 2A**). BirA is a bacterial ligase that specifically biotinylates the 15aa avidin within the short 15aa tag, allowing us to isolate bXPF-bound genome targets and protein complexes by binding to streptavidin. Using this approach, we performed chromatin streptavidin pulldowns followed by high-throughput sequencing (bXPF ChIP-Seq) in primary bXPF and BirA MEFs under basal conditions, upon transcription stimulation with tRA or upon exposure to ultraviolet (UV) irradiation. An Irreproducible Discovery Rate (IDR) filtering across two biological replicates (FDR≤0.05) revealed that the great majority of the 1100 identified bXPF-Seq peaks is mapped to intronic (24.1%), promoter (27%) and intergenic (31%) sequences under native conditions (**Figure 2Bi** and **D**; **Supplementary Table S1**). Treatment of MEFs with tRA led to an increase in the number of bXPF-Seq peaks by 78% (i.e. 1964) genome-wide (**Figure 2Bii** and **D**; **Supplementary Table S1**) and by 198% on promoters corresponding to 683 well-annotated genes (**Figure 2C**). Strikingly, we find a minimal number of bXPF-Seq peaks (i.e. 44) in bXPF MEFs treated with 10J/m^2^ of UV irradiation (**Figure 2Biii**) that was comparable to that seen in BirA transgenic control cells (**Figure 2Biv** and **Figure S1C**; **Supplementary Table S1**). A series of follow-up chromatin immunoprecipitation (ChIP) assays coupled to quantitative (q) PCR on peak sequences flanking the transcription start sites (TSS) of *Cfh*, *Rarb* and *Hs3st1* gene promoters selected from our RNA-Seq profiling in untreated and tRA-treated bXPF MEFs (**Supplementary Table S2**) confirmed the recruitment of bXPF on the promoters in untreated MEFs (**Figure 2D**), the significantly higher bXPF ChIP signals in tRA-treated MEFs and the substantial reduction of bXPF ChIP signals in UV-irradiated MEFs (**Figure 2E**; as indicated). In line, we find that bXPF is recruited minimally to the promoter region of the tRA non-responsive gene e.g. *Chordc1* or in non-transcribed genomic regions (**Figure 2E**; as indicated, **Supplementary table S2**). Thus, XPF is recruited predominantly to promoters under conditions that favor transcription but, in line with the random distribution of DNA damage events, it shows no selective recruitment to any DNA sites upon UV-induced DNA damage.

**Figure 2.**
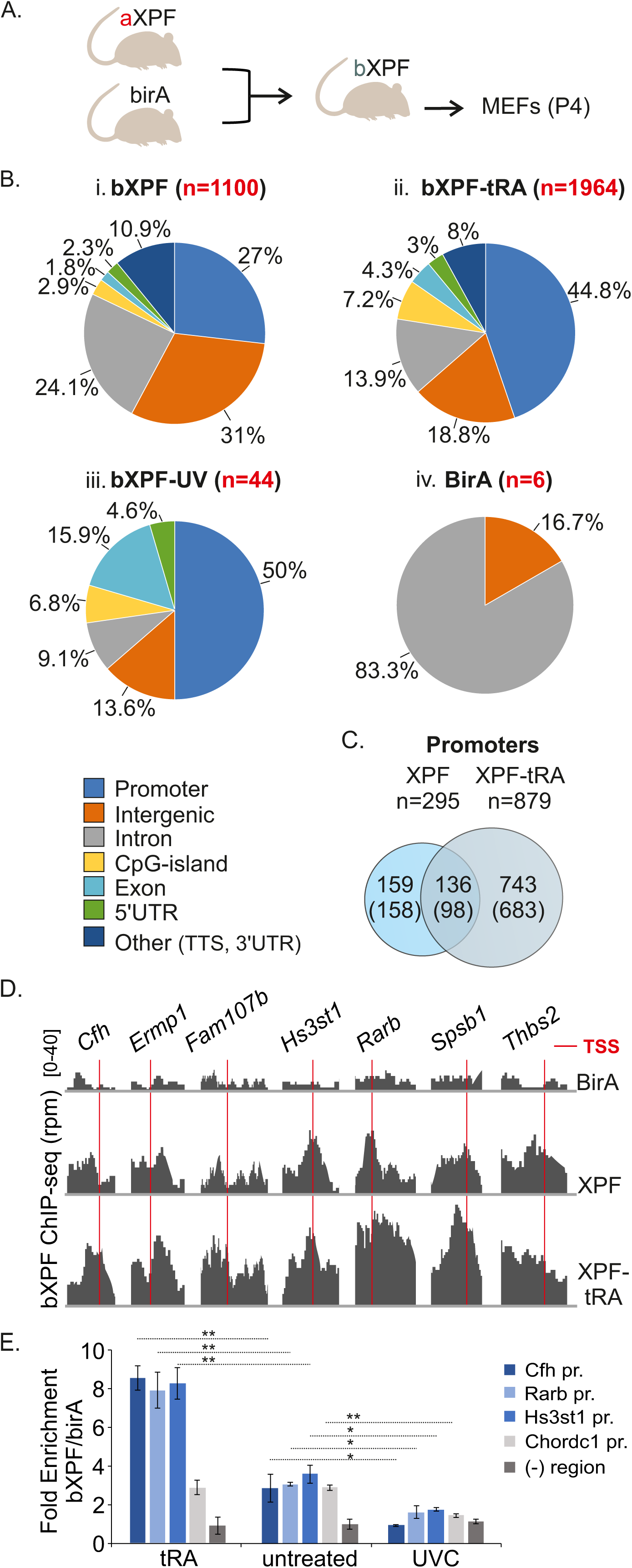
Genome-wide ChIP-Seq analysis of ERCC1-XPF occupancy in MEFs. **(A)**. Schematic representation of bXPF ChIP-Seq analysis in mouse embryonic fibroblasts (MEFs) derived from bXPF animals expressing the BirA transgene and BirA transgenic mice upon trans-retinoic acid (tRA) or UVC-irradiation. **(B)**. Pie charts illustrating the genomic distribution of bXPF binding sites in untreated, tRA- and UVC-treated bXPF and BirA (control) MEFs. Peaks occurring within ±2kb of the TSS were considered promoter. **(C)**. Venn diagram of XPF and XPF-tRA ChIP-Seq peaks mapped on promoters and corresponding number of unique genes (parenthesis). **(D)**. Genome browser views depicting bXPF ChIP-Seq signals on ±2kb genomic regions flanking the TSS of representative genes in untreated (bXPF) and tRA-treated (bXPF-tRA) MEFs. **(E)**. bXPF ChIP signals on the promoters of tRA-induced *Cfh*, *Rarb*, and *Hs3st1* genes, the tRA-non induced *Chordc1* gene and on an intergenic non-transcribed (-) region. bXPF ChIP signals are shown as fold enrichment of percentage input over percentage input BirA (for bXPF). Error bars indicate S.E.M. among n>three biological replicates. Asterisk indicates the significance set at p-value: *≤0.05, **≤0.01, ***≤0.001 (two-tailed Student’s t-test).

### bXPF recruitment on DNA coincides with RNAPII and active histone PTMs

Combinatorial ChIP-Seq profiles provide novel insights on shared or differential protein occupancies and histone marks. The preferential recruitment of bXPF to promoters in tRA-treated MEFs prompted us to contrast the genome-wide distribution of bXPF against the ChIP-Seq profiles of protein factors known to associate with active transcription i.e. RNAPII, H3K4me3, H3K27ac (Huang et al., 2019; Tie et al., 2009), gene silencing i.e. H3K4me1 (Cheng et al., 2014) and facultative or constitutive heterochromatin i.e. Lamin B (Zheng et al., 2015). Our analysis also included the CCCTC-binding factor (CTCF) factor known to interact with the ERCC1-XPF complex during postnatal murine development (Chatzinikolaou et al., 2017). Interrogation of chromosome 15 revealed that bXPF associates preferentially to gene-dense regions that closely coincide with regions bound by RNAPII, the active histone marks H3K4me3 and H3K27ac and with CTCF; in agreement, bXPF ChIP signals are excluded from Lamin B-associated heterochromatic regions reflecting low density gene regions (**Figure 3A**). Using a Pearson’s r correlation coefficient metrics, we find that, on a genome-wide level, the bXPF ChIP-Seq profiles of untreated or tRA-treated MEFs positively correlate with the RNAPII, H3K4me3 and H3K27ac ChIP-Seq profiles tested, including those of H3K4me1 that typically associates with gene repression (**Figure 3B**; left panel). Importantly, however, the Pearson’s r correlation increases significantly between bXPF (of untreated or tRA-treated MEFs) and RNAPII, H3K4me3 and H3K27ac ChIP-Seq profiles when the same analysis was carried out on promoters. Moreover, we find that the Pearson’s r correlation decreases substantially (lower correlation) or remains unaltered for H3K4me1 or CTCF ChIP-Seq signals, respectively (**Figure 3B**; right panel). To gain further insight into the recruitment of XPF to promoters, we next calculated the average coverage around the TSSs of genes bound by XPF in untreated and tRA-treated MEFs. Our analysis reveals a strong enrichment for bXPF (**Figure 3C**; green dotted line) and RNAPII, H3K4me3 and H3K27ac (**Figure 3C**; continuous lines as indicated) around TSS, which is further pronounced in tRA-treated MEFs (**Figure 3C**; red dotted line). The sharp dip of H3K27ac around the TSS represents a common feature of the TSS centered plots that likely reflects the position of the nucleosome-depleted zone (Jiang and Pugh, 2009). Instead, we find that H3K4me1 is locally depleted from TSSs, further confirming the previously observed negative correlation of bXPF with H3K4me1 on promoter regions (**Figure 3C**; as indicated). Next, to test whether the recruitment of bXPF on promoters and gene bodies associates with productive, steady-state mRNA synthesis, we performed an RNA-Seq analysis in untreated and tRA-treated bXPF MEFs. We find that 441 (441/539; 81.8%) bXPF-bound genes (including 5’UTR, promoter-TSS, exon, intron, TTS and 3’UTR region) contain ≥20 RNA-Seq counts (i.e. number of reads) in untreated MEFs (**Figure 3D**; upper pie chart; **Supplementary Table S2**). The number of bXPF-bound genes increases substantially to 1049 bXPF-bound genes (1049/1199; 87.54%) when the same analysis was carried out in tRA-treated MEFs (**Figure 3D**; lower pie chart; **Supplementary Table S2**). Thus, upon transcription activation, bXPF is recruited to the promoters of actively transcribed genes and significantly correlates with the occupancy of RNAPII and active histone PTMs on these sites.

**Figure 3.**
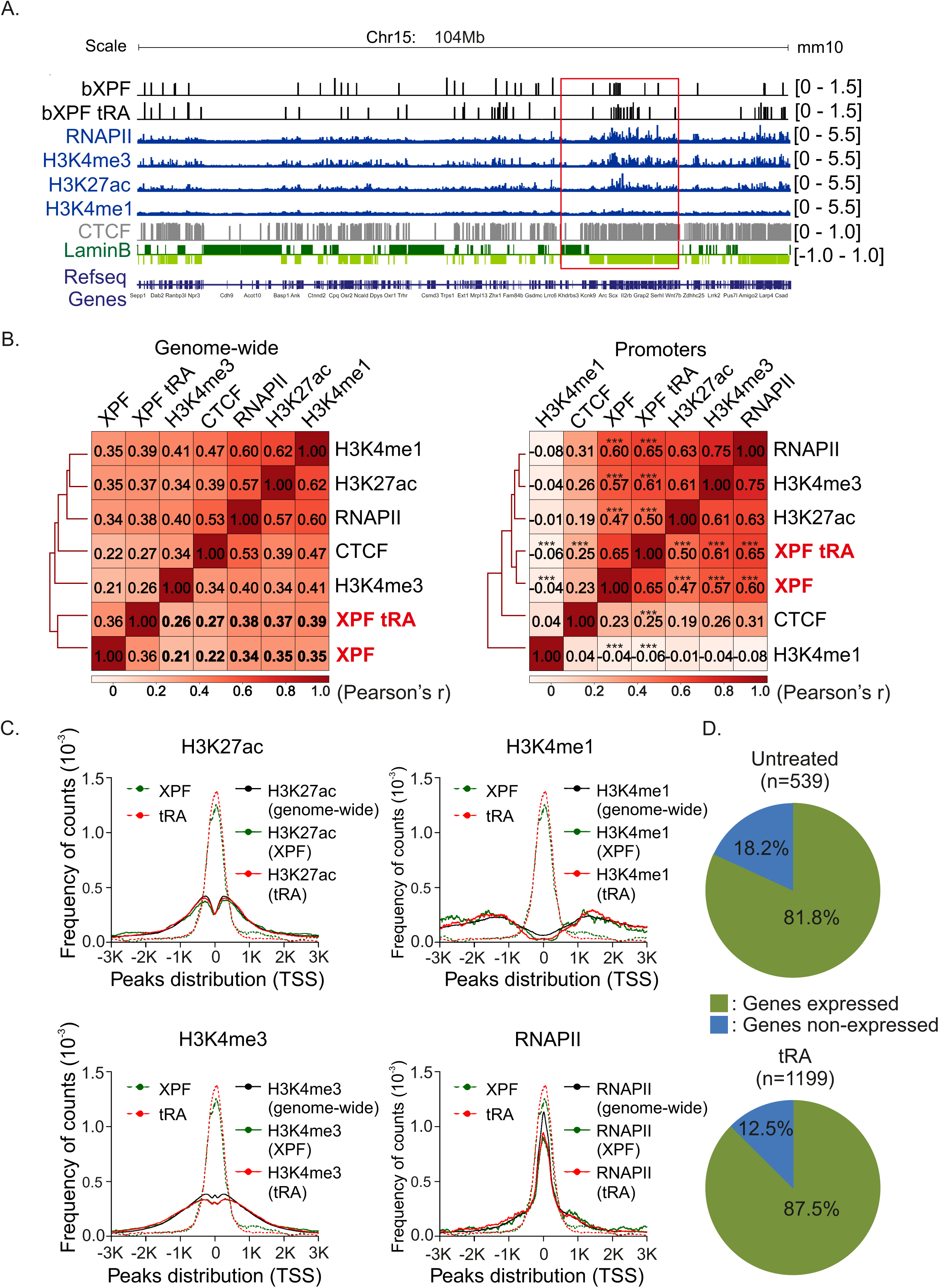
Chromatin state of ERCC1-XPF binding peaks. **(A)**. IGV overview of ChIP-Seq profiles for bXPF (untreated MEFs), bXPF-tRA (tRA-treated MEFs), RNAPII, H3K4me3, H3K27ac, H3K4me1, CTCF and Dam-ID profile of Lamin B on a representative 103 Mb (Chromosome 15) genomic region in MEFs. **(B)**. Genome- (left panel) or gene promoter-wide (right panel) heatmap representation of Pearson’s r correlation analysis of XPF (untreated bXPF MEFs), XPF-tRA (tRA-treated MEFs), RNAPII, H3K27ac, H3K4me3 and H3K4me1 ChIP-Seq profiles. For promoters, the p-value (***: P= 0.001-0.0001, **: P=0.05-0.001) is based on the comparison of Pearson’s r correlations (single sided test) from independent samples; in this case, between the correlations of genome-wide and promoter-associated ChIP-Seq signals **(C)**. Average count frequencies on +/−3Kb genomic regions flanking the TSS for RNAPII, H3K27ac and H3K4me3 activating histone marks, H3K4me1 repressive histone modification, bXPF- and XPF-tRA-bound gene targets. Dotted lines depict the genome-wide profiles of bXPF and bXPF-tRA. **(D)**. Pie charts depicting the RNA-Seq gene expression status (blue: non-expressed; green: expressed) of bXPF-bound genes (5’UTR, promoter-TSS, exon, intron, TTS, 3’UTR) in untreated (upper pie chart) and tRA-treated (lower pie chart) MEFs.

### A proteomics strategy reveals bXPF-bound protein partners involved in chromosome organization, transcription and DNA repair

We reasoned that the selective recruitment of bXPF on promoters reflects possible interactions of ERCC1-XPF with factors associated with transcription initiation and/or transcription-associated DNA damage. To test this, we combined the *in vivo* biotinylation tagging approach (Chatzinikolaou et al., 2017) with a hypothesis-free, high-throughput proteomics strategy in primary bXPF MEFs. Using high salt extraction methods, we prepared nuclear extracts from bXPF MEFs and MEFs expressing only the BirA transgene that were subsequently treated with benzonase and RNase A; the latter ensures that neither DNA nor RNA mediate the identified protein interactions (**Figure 4A**). Nuclear extracts were further incubated with streptavidin-coated beads and bound proteins were eluted and subjected to Western blot analysis, confirming that bXPF can still interact with its obligatory partner ERCC1 (**Figure 4B**). Next, the proteome was separated by 1D SDS-PAGE (∼12 fractions) followed by in-gel digestion and peptides were analyzed with high-resolution liquid chromatography-tandem mass spectrometry (nLC MS/MS) on a hybrid linear ion trap Orbitrap instrument (**Figure 4C**). From three biological replicates, which comprised a total of 72 MS runs, we identified a total of 695 proteins (**Supplementary Table S3**) with 607 proteins (87.3%) shared between all three measurements under stringent selection criteria (**Figure 4D**; **Supplementary Table S4**). To functionally characterize this dataset, we subjected the 607-shared bXPF-bound proteins to gene ontology (GO) classification. Those biological processes (**Figure 4E**) or pathways (**Figure 4F**) containing a significantly disproportionate number of proteins relative to the murine proteome were flagged as significantly over-represented (FDR<0.05). At this confidence level, the over-represented biological processes and pathways involved 77 out of the initial 607 bXPF-bound core proteins; the latter set of proteins also showed a significantly higher number of known protein interactions (i.e. 286 interactions) than expected by chance (i.e. 76 interactions; **Figure 4G**) indicating a functionally relevant and highly interconnected protein network. Using this dataset, we were able to discern four major, partially overlapping, bXPF-associated protein complexes involved in i. chromosome organization (p≤3.2×10, e.g. CTCF, HIST1h1a-e, H1F0, SMARCA5, SMC1A, SMC3, TOP1, TOP IIα, TOP IIβ) ii. transcription (p ≤ 2.8×10^−16^, e.g. TAF6, TAF10, TAF4A, KLF13, UBTF, TOP1, TOP IIα, TOP IIβ, RBM39, NUP107, NUP133, NUP153), iii. gene silencing (p ≤ 8.3×10, e.g. BMS1, GNL3, MDN1, NOP58, UTP15, WDR36, WDR43, WDR75, XRN2) and DNA replication (p ≤ 1.2×10^−12^, e.g. RCF2, RCF3, RCF4, RCF5, SSRP1, RBBP6). Together, these findings indicate that the great majority of bXPF-bound protein partners are functionally involved in genome utilization processes.

**Figure 4.**
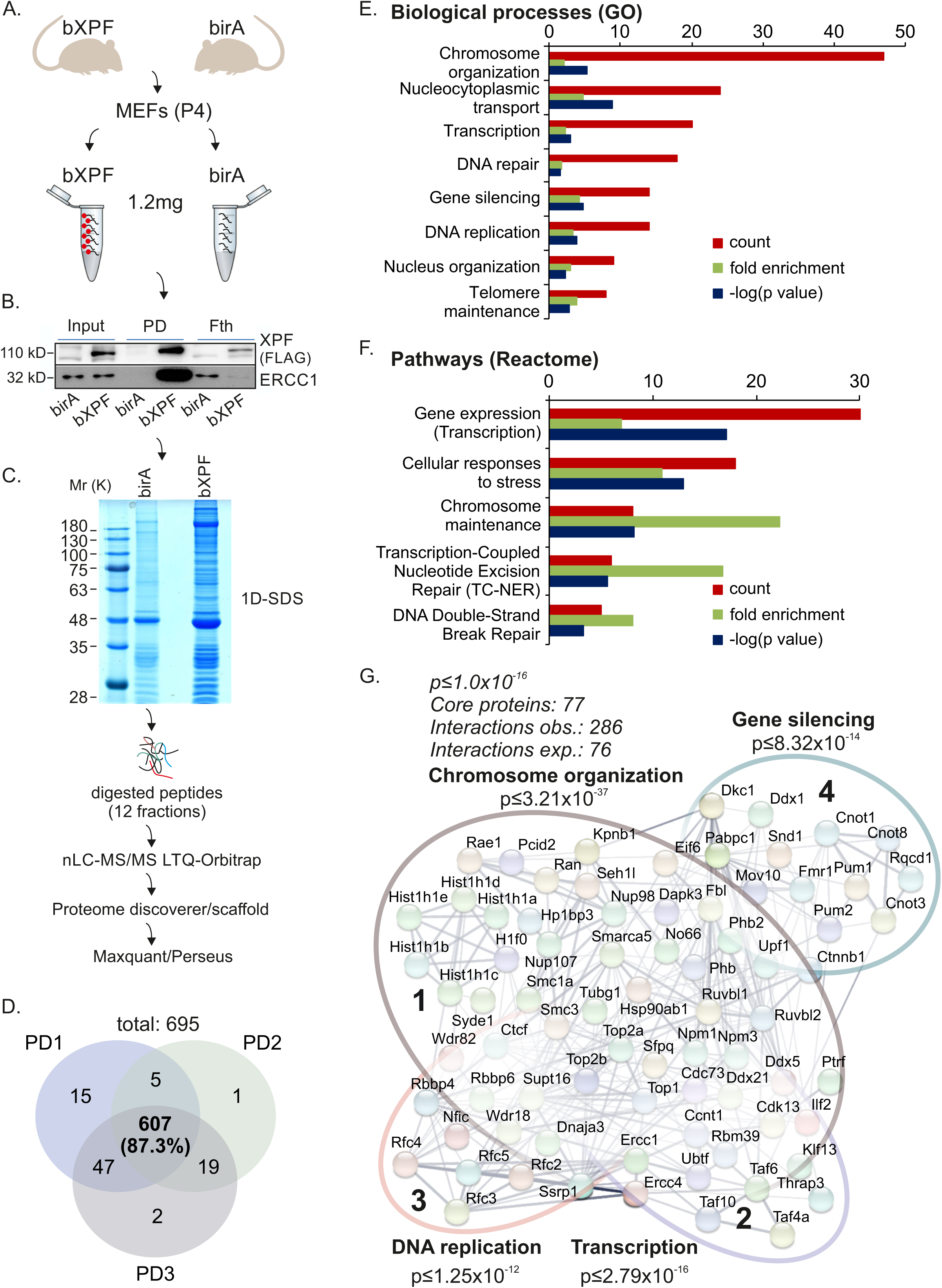
ERCC1-XPF interacts with chromatin remodeling and transcription factors. **(A)**. Schematic representation of the high-throughput MS analysis performed using nuclear extracts from bXPF and BirA MEFs. **(B).** bXPF pulldowns (fth: flow through, PD: Pulldown) and western blot with anti-FLAG and anti-ERCC1 in nuclear extracts derived from bXPF and BirA MEFs. **(C)**. A representative 2D gel of proteins extracts derived from bXPF and BirA MEFs. **(D)**. Venn diagram of bXPF-bound protein factors from three independent pulldowns (PD) and subsequent MS analyses. **(E)**. Significantly over-represented biological processes (gene ontology; GO) and **(F)**. pathways (Reactome) of the shared 607 bXPF-bound proteins **(G)**. Number of observed (obs.) and expected (exp.) known protein interactions within the core XPF-bound protein set; highlighted circles represent the four major XPF-bound protein complexes involved in chromosome organization, gene silencing, DNA replication and transcription.

### ERCC1-XPF interacts with DNA TOP IIβ on promoters

Pulldown experiments in nuclear extracts of bXPF and control BirA MEFs confirmed that the endogenous bXPF interacts with the TATA-associated factors (TAFs) TAF4, TAF6, and TAF10 of the TFIID complex (Kamileri et al., 2012c) as well as with CTCF and the cohesin subunits SMC1A and SMC3 (**Figure 5A** and **Figure 5B**) (Chatzinikolaou et al., 2017) highlighting the possible role of ERCC1-XPF complex in transcription initiation and chromatin looping (Apostolou et al., 2019; Kamileri et al., 2012a). Moreover, these findings along with the preferential recruitment of bXPF on promoters upon transcription stimulation (**Figure 2B-C**) and the identification of several topoisomerases in the 607 bXPF-bound core proteome (**Figure 4G**) fit well with the known involvement of ERCC1-XPF in DNA DSB repair (Ahmad et al., 2008; Li et al., 2019). Indeed, during the process of ongoing mRNA synthesis, TOP IIβ relieves topological constraints by triggering the formation of transient DNA DSBs on promoters of actively transcribed genes (Calderwood, 2016; Canela et al., 2019; Madabhushi et al., 2015; Pommier et al., 2016). In line, we find that the endogenous bXPF interacts with TOP IIβ but not with TOP IIα or TOP I (**Figure 5C**). Similar to bXPF, a series of immunoprecipitation experiments revealed that ERCC1 interacts specifically with TOP IIβ as well as with CTCF and the cohesin SMC1A and SMC3 subunits in primary MEFs (**Figure 5D**). Follow-up immunoprecipitation experiments with an antibody raised against TOP IIβ confirmed the reciprocity of ERCC1-XPF and TOP IIβ interaction (**Figure 5E**). TOP IIβ is known to co-localize with the evolutionarily conserved CTCF/cohesin binding sites whereas members of the cohesin complex and CTCF were recently identified as TOP II-interacting proteins in a high-throughput MS screen (Uuskula-Reimand et al., 2016). As with ERCC1-XPF, we find that TOP IIβ reciprocally interacts with CTCF and the SMC1A and SMC3 (**Figure 5F**); importantly, the interaction of TOP IIβ with SMC1A or CTCF is not abolished when ERCC1-XPF is abrogated in *Ercc1^−/−^* MEFs (**Figure 5G**). Confocal imaging in untreated and tRA-treated wt. MEFs revealed that whereas ERCC1 or bXPF are evenly scattered in the nucleoplasm, TOP IIβ localizes in clear subnuclear landmarks identified as heterochromatin by 4′,-6-diamidine-2-phenylindole (DAPI). Importantly, however, TOP IIβ is re-distributed throughout the nucleoplasm in tRA-treated MEFs (**Figure 5H** and **Supplementary Figure S2B** and **S2C**). To test for the relevance and specificity of TOP IIβ binding to bXPF-bound promoters in MEFs, we next performed a series of ChIP-qPCR assays using antibodies raised against TOP IIβ, TOP IIα and TOP I on tRA-induced *Rarb*, *Cfh* and *Hs3St1* promoters previously identified in the bXPF ChIP-Seq profiles. Our analysis reveals that TOP IIβ (**Figure 5I**), but not TOP IIα (**Supplementary Figure S3A**) or TOP I (**Supplementary Figure S3B**), is recruited preferentially to *Rarb*, *Cfh* and *Hs3St1* promoters. Similar to bXPF, we find that the TOP IIβ ChIP signals remain unchanged at the tRA non-responsive *Chordc1* gene promoter or at a non-transcribed genomic region (**Figure 5I**; as indicated). ChIP/re-ChIP analysis using antibodies against TOP IIβ (1^st^ ChIP) and ERCC1, FLAG-tagged XPF or CTCF (2^nd^ ChIP) showed that these factors co-occupy the *Rarb*, *Cfh, Hs3St1* or *Spsb3* gene promoters (**Figure 5J-K**, **Supplementary Figure S3D-E**). Unlike, however, with bXPF, we find that the affinity of TOP I, IIα or IIβ and the CTCF/cohesin complex to chromatin remains unaffected by UV-induced DNA damage as all corresponding ChIP signals remain unaltered in UV-irradiated cells (**Supplementary Figures S3C**, **S3F-H** and **S4A-B**). Taken together, our findings reveal that the ERCC1-XPF complex interacts specifically with TOP IIβ and that the endonuclease complex is recruited with TOP IIβ and the CTCF/cohesin complex to gene promoters in the absence of exogenous genotoxic stress and under conditions that favor transcription.

**Figure 5.**
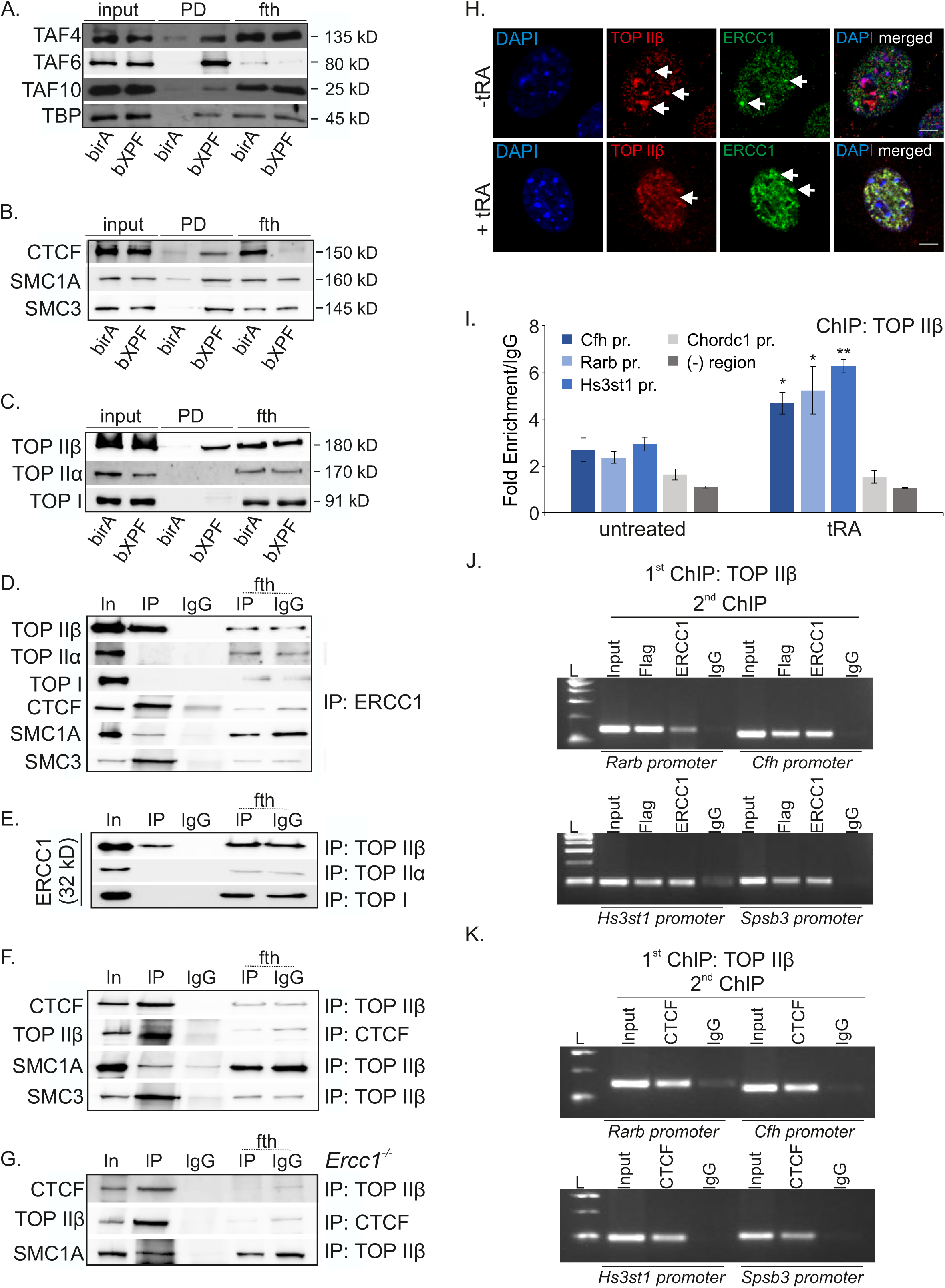
ERCC1-XPF interacts with TOP IIβ on promoters. **(A)**. bXPF pulldowns (PD) and western blot with anti-TAF4, TAF6, TAF10 and TBP. (**B**). CTCF, SMC1A and SMC3 and (**C**). TOP IIβ, TOP IIa and TOP I in nuclear extracts derived from bXPF and BirA MEFs (as indicated). **(D)**. Co-immunoprecipitation experiments using anti-ERCC1 in nuclear extracts from wt. MEFs analyzed by western blotting for TOP IIβ, TOP IIa and TOP I and CTCF, SMC1A and SMC3 (as indicated). **(E)**. Co-immunoprecipitation experiments using anti-TOP IIΒ, anti-TOP2A or anti-TOP1 in nuclear extracts from wt. MEFs analyzed by western blotting for ERCC1 (as indicated). The input and flow-through are 1/20 of the extract used. **(F)**. Co-immunoprecipitation experiments using anti-TOP IIΒ or anti-CTCF in nuclear extracts from wt. MEFs analyzed by western blotting for CTCF, TOP IIβ, SMC1A and SMC3. (**G**). Co-immunoprecipitation experiments using anti-TOP IIβ or anti-CTCF in nuclear extracts from primary *Ercc1*^−/−^ MEFs analyzed by western blotting for CTCF, TOP IIβ and SMC1A. (**H**). Immunofluorescence detection of TOP IIβ and ERCC1 (white arrowheads) in primary wt. MEFs cultured upon basal conditions (-tRA) or upon treatment with tRA (+ tRA). **(I)**. TOP IIβ ChIP signals on the promoters of tRA-induced *Cfh*, *Rarb*, and *Hs3st1* genes, the tRA-non induced *Chordc1* gene and on an intergenic non-transcribed (-) region (as indicated). (**J**). ChIP with antibodies raised against TOP IIβ and re-ChIP with antibodies raised against ERCC1 or Flag-tagged XPF on *Rarb* and *Cfh* gene promoters (upper panel) and on *Hs3st1* and Spsb3 gene promoters (lower panel). (**K**). ChIP with antibodies raised against TOP IIΒ and re-ChIP with antibodies raised against CTCF on *Rarb* and *Cfh* gene promoters (upper panel) and on *Hs3st1* and Spsb3 gene promoters (lower panel). See also Supplementary Figure S3. Error bars indicate S.E.M. among n>three biological replicates. Asterisk indicates the significance set at p-value: *≤0.05, **≤0.01, ***≤0.001 (two-tailed Student’s t-test).

### TOP IIβ-mediated DNA damage triggers the recruitment of bXPF on promoters during transcription

The interaction of ERCC1-XPF with TOP IIβ on promoters and the increased accumulation of γH2AX and 53BP1 foci in tRA-treated *Ercc1^−/−^* cells prompted us to test for the functional relevance of ERCC1-XPF to TOP IIβ-induced DNA DSB formation during transcription. To do so, we first treated tRA-treated MEFs with merbarone (tRA/merb MEFs), a DNA topoisomerase II catalytic inhibitor that acts by blocking TOP II-mediated DNA cleavage without interfering with protein-DNA binding. The latter led to the noticeable decrease of γH2AX and 53BP1 foci in tRA/merb-treated MEFs (**Figure 6A**) and bXPF ChIP signals on *Cfh*, *Rarb*, *Hs3st1* gene promoters in tRA/merb MEFs relative to tRA-treated MEFs, the non-induced *Chordc1* gene promoter or to a non-transcribed intergenic region (**Figure 6B** and **Supplementary Figure S4D**; as indicated). Confocal imaging in tRA/merb cells revealed that, unlike in tRA-treated cells (**Figure 5H**, **Supplementary Figure S2B** and **S2C**); TOP IIβ remains in the heterochromatin and does not diffuse in the nucleoplasm of primary MEFs (**Supplementary Figure S4C**). To further challenge these findings, we treated wt. MEFs with etoposide, a topoisomerase II inhibitor that prevents re-ligation of TOP II-mediated DNA DSBs. In support of our previous findings, we find a substantial accumulation of TOP IIβ-mediated γH2AX and 53BP1 foci (**Figure 6C**) and a significant increase of bXPF ChIP signals on gene promoters in etoposide-treated bXPF MEFs (**Figure 6D**). Thus, upon transcription initiation, TOP IIβ is causal to the great majority of transcription-associated DNA DSBs triggering the recruitment of bXPF on transcriptionally active promoters.

**Figure 6.**
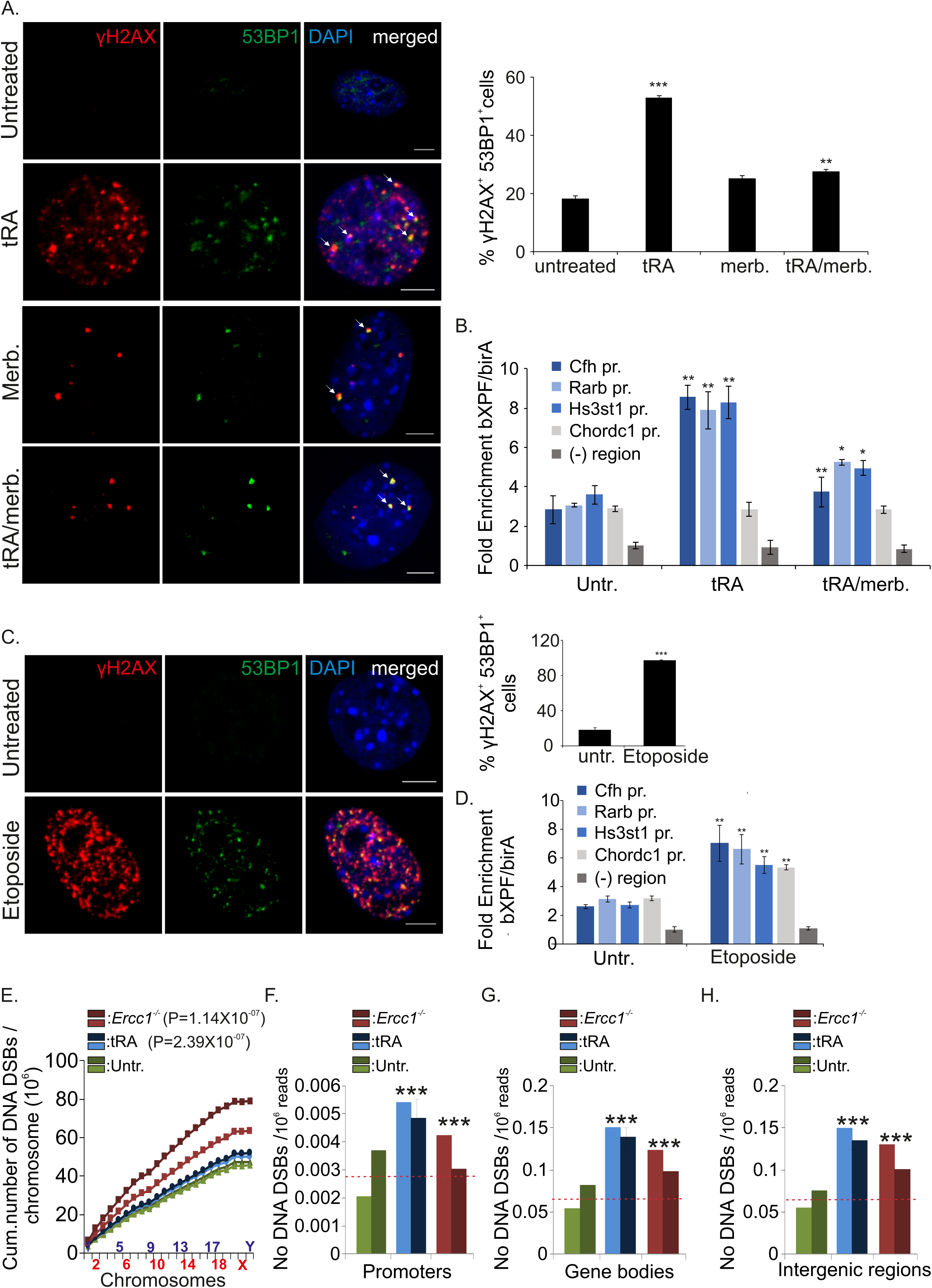
Transcription-associated DNA damage events require TOP IIβ. **(A).** Immunofluorescence detection of γH2AX and 53BP1 (white arrowheads) in wt. tRA-treated MEFs cultured in the presence of tRA, merbarone (Merb.) or tRA and merbarone (as indicated). The graph represents the number of γH2AX^+^ 53BP1^+^ cells under the conditions shown in the x-axis. **(B)**. bXPF ChIP signals on the tRA-inducible *Cfh*, *Rarb* and *Hs3st1* gene promoters, the tRA non-inducible *Chordc1* gene promoter, as well as on a non-transcribed intergenic region (-) region in MEFs treated with tRA, merbarone (merb.) or tRA and merbarone. The ChIP signals are shown as fold enrichment of the percentage of input for bXPF over the percentage of input for BirA. (**C**). Immunofluorescence detection of γH2AX and 53BP1 in wt. untreated MEFs and wt. MEFs treated with etoposide. The graph represents the number of γH2AX^+^ 53BP1^+^ cells in etoposide-treated and corresponding control cells. (**D**). bXPF ChIP signals on the tRA-inducible *Cfh*, *Rarb* and *Hs3st1* gene promoters, the tRA non-inducible *Chordc1* gene promoter, as well as on a non-transcribed intergenic region (-) region in MEFs treated with etoposide. **(E)**. Cumulative number of DNA DSBs per chromosome in untreated (Untr.), tRA-treated (tRA) or *Ercc1^−/−^* MEFs (as indicated); for each color, the light-and dark-colored lines indicate the two BLISS replicates for each experimental condition. The red dotted line represents the average of the two biological replicates in untreated (Untr.) control MEFs. **(F)**. Number of DNA DSBs per million mapped reads on gene promoters in untreated (Untr.), tRA-treated (tRA) or *Ercc1^−/−^* MEFs (as indicated); for each color, the light- and dark-colored bars indicate the two BLISS replicates for each experimental condition. The red dotted line represents the average of the two biological replicates in untreated (Untr.) control MEFs. **(G)**. Number of DNA DSBs per million mapped reads on gene bodies in untreated (Untr.), tRA-treated (tRA) or *Ercc1^−/−^* MEFs (as indicated); for each color, the light- and dark-colored bars indicate the two BLISS replicates for each experimental condition. The red dotted line represents the average of the two biological replicates in untreated (Untr.) control MEFs. (**H**). Number of DNA DSBs per million mapped reads on intergenic regions in untreated (Untr.), tRA-treated (tRA) or *Ercc1^−/−^* MEFs (as indicated); for each color, the light- and dark-colored bars indicate the two BLISS replicates for each experimental condition. Error bars indicate S.E.M. among n>three biological replicates. Asterisk indicates the significance set at p-value: *≤0.05, **≤0.01, ***≤0.001 (two-tailed Student’s t-test). For Figure 6E-H, a triple asterisk indicates the significance set at p-value ≤ 10^−15^ (Mann Whitney test). Grey line is set at 10 μm scale.

### Genome-wide association of ERCC1-XPF recruitment on promoters with activity-induced DNA DSBs

Our finding that the recruitment of ERCC1-XPF on promoters is dependent on TOP IIβ activation prompted us to compare the bXPF ChIP-Seq profiles with the genome-wide distribution and frequency of DNA DSBs in tRA-treated MEFs. To do so, we used Breaks Labeling *In Situ* and Sequencing (BLISS) (Yan et al., 2017) to identify DNA DSBs across the genome. After filtering out PCR duplications, we find that DNA DSB counts are evenly distributed across the mouse chromosomes (**Figure 6E**). Whereas *Ercc1^−/−^* MEFs have a significantly higher number of total DNA DSBs, tRA-treated MEFs have a significantly higher number of DNA DSBs in transcription-associated regions per 100ng of genomic DNA (p = 1.14×10^−7^and p = 2.39×10^−7^, respectively), including in promoters (**Figure 6F**) and gene bodies (**Figure 6G**) when compared to untreated wt. MEFs. Interestingly, we also find that, similar to *Ercc1^−/−^* MEFs, tRA treatment increases the occurrence of DNA DSBs also in intergenic regions (**Figure 6H**). In support, previous findings have shown that transcription induction requires the transient formation of DNA DSBs at intergenic cis-regulatory elements to resolve topological problems (Uuskula-Reimand et al., 2016). Further analysis of the +/−2kb genomic region flanking the TSS of all genes revealed a significant, genome-wide, enrichment of DNA DSBs on the TSS of genes in untreated and tRA-treated MEFs (**Figure 7A-B**), which was not apparent in *Ercc1^−/−^* MEFs (**Figure 7C**). A follow-up BLESS (Breaks Labeling, Enrichment on Streptavidin and next-generation Sequencing) approach coupled to qPCR on the previously identified bXPF-bound *Rarb*, *Cfh,* and *Hs3st1* gene promoters confirmed the significant increase of DNA DSBs on promoters in tRA-treated MEFs; importantly, we find no increase of DNA DSBs in a transcriptionally inactive genomic region (**Figure 7D**). Instead, DNA DSBs decreased substantially when tRA-treated MEFs were also exposed to the TOP IIβ inhibitor merbarone (**Figure 7D**; as indicated); merbarone alone was also included in our analysis as it is a known instigator of chromosomal damage (Wang and Eastmond, 2002). By using a modified BLESS approach; we next sought to confirm the significant enrichment of ERCC1, TOP IIβ and CTCF proteins on the isolated DSB genomic fragments (**Figure 7E**). Using a control sample without a ligated adapter to test for the non-specific binding of DNA fragments on beads, we find that ERCC1, TOP IIβ and CTCF are significantly enriched on DNA DSBs in MEFs. Importantly, ∼96% out of the 1100 peaks identified in bXPF ChIP-Seq profiles contained DNA DSBs (**Figure 2B, Figure 7F**; n=5803; **Supplementary Table S5**) with 26% of the identified DNA DSBs being detected on bXPF-bound promoters. Upon tRA treatment, the number of DNA DSBs increased dramatically to 9665 corresponding to 91,2% of the 1964 peaks identified in bXPF ChIP-Seq profiles with 42% of the identified DNA DSBs being detected on bXPF-bound promoters (**Figure 2B**, **Figure 7F**; n=9665; **Supplementary Table S5**). To test whether transcription activation and/or the associated DNA DSBs affect the probability of XPF to bind to gene promoters, we generated a classification model for bXPF binding (bound/unbound) using the automated machine learning (AutoML) tool JAD Bio (Lakiotaki et al., 2019) and the logarithms of RNA-Seq, BLISS and the tRA treatment as predictors (**Supplementary Table S6**). The importance of the interaction term logRNA-seq×logBLISS is visually verified in **Figure 7G** and **Supplementary Figure S4E-F** where the distributions of the bXPF-bound and -unbound sites are clearly distinguished based on this feature. In line with our previous findings, we find that recruitment of bXPF requires transcription activation and the presence of DNA DSBs, independently of the tRA treatment in MEFs. Consistent with the increased detection of γH2AX^+^ 53BP1^+^ cells in tRA-treated *Ercc1^−/−^* MEFs (**Figure 1D**) and the developing P15 cerebella and livers (**Figure 1E**), we find a higher number of DNA DSBs on the *Rarb2* promoter in tRA-treated *Ercc1^−/−^* MEFs relative to wt. controls (**Figure 7H**; as indicated) as well as on the promoters of actively transcribed genes in P15 *Ercc1^−/−^* livers (genes: *PrlR*, *Dio1*)(Kamileri et al., 2012b) and cerebella (genes: *Dhfr*, *Prnp*)(McKenzie et al., 2018; Nouspikel and Hanawalt, 2000) (**Figure 7I** and **Figure 7J**; as indicated), relative to wt. control tissues. Importantly, we find no difference in the number of DNA DSBs on the promoters of non-expressing genes in the liver i.e. *NeuN* and cerebellum i.e. *Alb* (**Figure 7I-J**; as indicated). Taken together, our findings indicate that, upon transcription induction, ERCC1-XPF is preferentially recruited on active promoters for the repair of activity-induced DNA DSBs.

**Figure 7.**
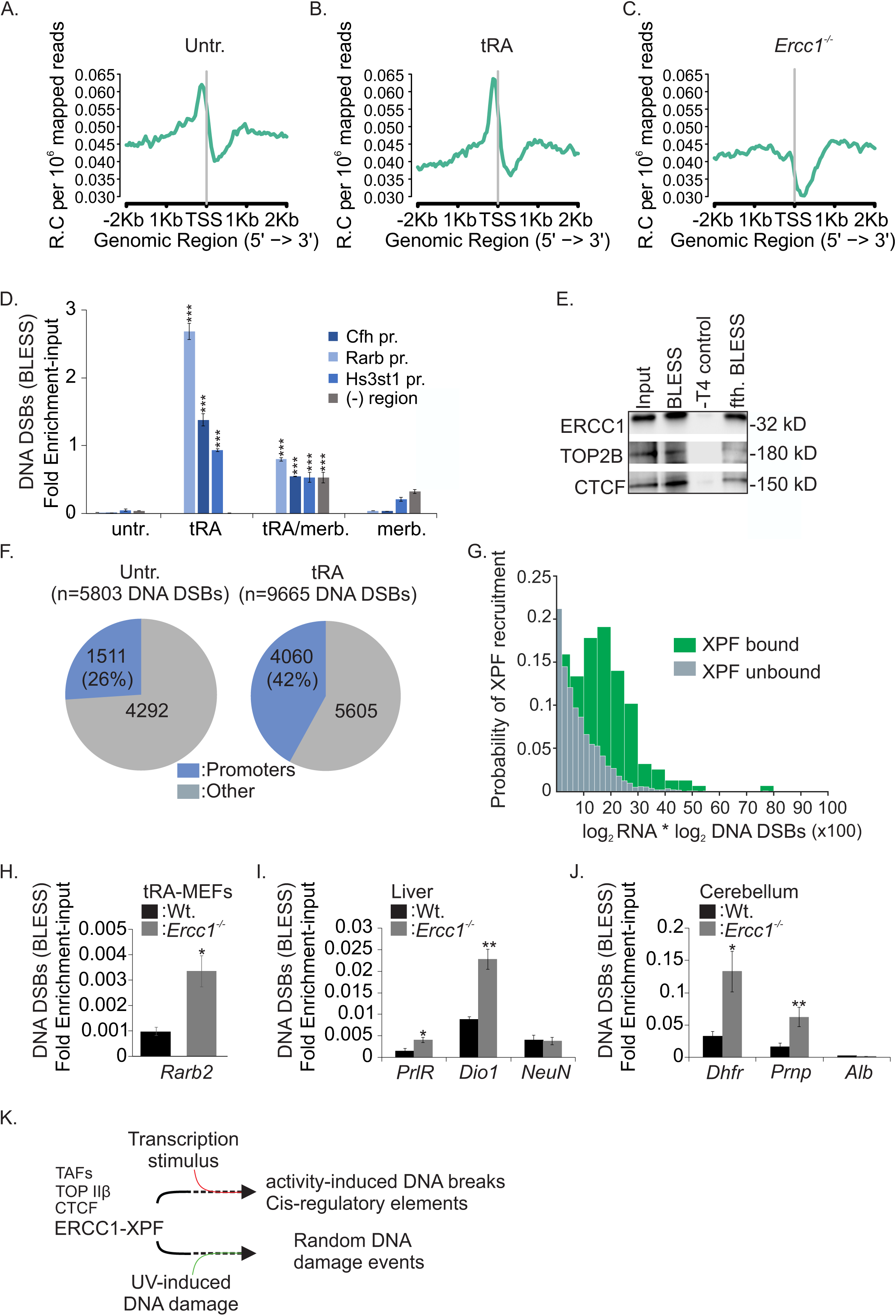
Genome-wide mapping of DSBs in tRA-treated and *Ercc1*^−/−^ MEFs. **(A)**. Genome-wide enrichment (read count; R.C.) of DNA DSBs (normalized per million mapped reads) on -/+2Kb flanking the TSS in untreated (Untr.), **(B)**. tRA-treated (tRA) and **(C)**. *Ercc1^−/−^* MEFs. **(D)**. BLESS signals quantified by qPCR on the tRA-inducible *Cfh*, *Rarb* and *Hs3st1* gene promoters, the tRA non-inducible *Chordc1* gene promoter, as well as on a non-transcribed intergenic region (-) region in untreated MEFs (Untr.) or MEFs treated with tRA (tRA) or tRA and merbarone (tRA/merb). **(E)**. Enrichment of ERCC1, TOP IIβ and CTCF proteins on DSB-containing genomic fragments derived from primary MEFs. **(F)**. Venn’s logic diagrams representing the number of transcription-associated DNA DSBs on XPF-bound genomic sites in untreated (Untr.) and tRA-treated MEFs. **(G)**. Probability of XPF recruitment on genes by means of log^2RNA*log^2Breaks variable from the RNA-Seq and BLISS data. (**H**). BLESS signals quantified by qPCR on *Rarb2* gene promoter in tRA-treated wt. and *Ercc1^−/−^* MEFs. (**I**). BLESS signals quantified by qPCR on the promoters of *PrlR* and *Dio1* genes known to be expressed in the P15 *Ercc1^−/−^* and wt. livers and on the promoter of non-expressed *Neun* gene. (**J**). BLESS signals quantified by qPCR on the promoter of *Dhfr* and *Prnp* genes known to be expressed in the P15 *Ercc1^−/−^* and wt. cerebella and on the promoter of the non-expressed *Alb* gene. Error bars indicate S.E.M. among n>three biological replicates. Asterisk indicates the significance set at p-value: *≤0.05, **≤0.01, ***≤0.001 (two-tailed Student’s t-test).

## Discussion

TOP II-mediated DSBs are essential for estradiol-stimulated activation of gene expression (Ju et al., 2006), to facilitate the expression of a subset of neuronal early-response genes (Madabhushi et al., 2015) and to regulate the expression of developmental gene expression programs (Lyu et al., 2006). Consistent with these observations, we find that, in the absence of exogenous genotoxic insults, transcription triggers the genome-wide accumulation of DNA DSBs and the formation of γH2Ax and 53BP1 foci in MEFs. The latter requires TOP II activity and a functional ATM because inhibition of the catalytic activity of TOP II or ATM leads to a substantial decrease of γH2Ax and 53BP1 foci in tRA-treated cells. In agreement, blocking the re-ligation of TOP II-mediated DNA DSBs by etoposide prompted a robust accumulation of γH2Ax and 53BP1 foci similar to that seen when cells are exposed to the potent genotoxin MMC.

We previously showed that ERCC1-XPF engages together with RNAPII and the basal transcription machinery at the promoters of growth genes during postnatal hepatic development (Apostolou et al., 2019; Kamileri et al., 2012a; Kamileri et al., 2012b). However, the structure-specific requirement of ERCC1-XPF for incision made it difficult to justify the functional role of the endonuclease complex on promoters. Using an *in vivo* biotinylation tagging approach coupled to genome-wide mapping of DNA DSBs and functional genomics approaches, we find that, upon transcription induction, XPF recruits preferentially at and upstream of the TSS of actively transcribed genes, of which a great majority bears activity-induced DNA breaks. In agreement, the XPF ChIP-Seq peak summits on promoters positively correlate with those of RNAPII, H3K4me3 and H3K27ac, associated with active transcription and are excluded from lamin-associated heterochromatin. Moreover, bXPF ChIP signals show no correlation with H3K4me1 associated with gene silencing. Importantly, we find that bXPF disassembles from promoters and shows no selective recruitment to any genomic region in UV-irradiated cells. These findings suggest that in the absence of exogenous genotoxic insults, ERCC1-XPF is recruited genome-wide to activity-induced DNA breaks, which are spatially restricted to promoters or cis acting regulatory regions. Upon UV irradiation, however, DNA lesions, and by inference the ERCC1-XPF complex itself, are expected to be randomly scattered across the mammalian genome.

Although transcriptionally active chromatin is preferentially repaired by HR (Aymard et al., 2014), recent findings indicate that the repair of activity-induced DSBs is dependent on NHEJ (Madabhushi et al., 2015) where ERCC1-XPF is required for trimming the DNA ends before resealing (Ahmad et al., 2008; Manandhar et al., 2015; McDaniel and Schultz, 2008). These data and our finding that ERCC1-XPF interacts with TFIID e.g. several TAFs and the TBP involved in transcription initiation in mouse livers (Kamileri et al., 2012b) and primary MEFs (this work) may well explain how the endonuclease complex is specifically targeted to active promoters. Furthermore, the interaction of ERCC1-XPF with TOP IIβ, CTCF and the cohesin subunits on gene promoters would facilitate the proximity of bXPF-bound promoters with enhancers (Ren et al., 2017), chromatin looping (Chatzinikolaou et al., 2017; Le May et al., 2012) and/or the positioning of TOP IIβ at topologically associating domain boundaries (Uuskula-Reimand et al., 2016) allowing the selective regulation of gene expression *in vivo*. In this respect, our finding that activity-induced DNA breaks accumulate on the promoters of actively transcribed genes in *Ercc1^−/−^* MEFs as well as in developing cerebella and livers may well explain how persistent DNA damage triggers the wide range of tissue-specific, progeroid features seen in patients and accompanying animal models with congenital ERCC1-XPF defects (**Figure 7K**).

## Methods

### Animal models and primary cells

The generation and characterization of Biotin-tagged XPF (bXPF) and NER-deficient mice has been previously described (Apostolou et al., 2019; Chatzinikolaou et al., 2017; Kamileri et al., 2012a; Kamileri et al., 2012b). Animals were kept on a regular diet and housed at the IMBB animal house, which operates in compliance with the “Animal Welfare Act” of the Greek government, using the “Guide for the Care and Use of Laboratory Animals” as its standard. As required by Greek law, formal permission to generate and use genetically modified animals was obtained from the responsible local and national authorities. All animal studies were approved by independent Animal Ethical Committees at FORTH and BSRC Al. Fleming. Primary MEFs were isolated from E13.5d animals and cultured in standard medium containing Dulbecco’s Modified Eagle Medium (DMEM) supplemented with 10% Fetal Bovine Serum (FBS), 50 μg/ml streptomycin, 50 U/ml penicillin (Sigma) and 2mM L glutamine (Gibco). Cells were rinsed with PBS, exposed to UVC irradiation (10 J/m^2^), MMC (10 μg/mL) (AppliChem), tRA (10 μM) (Sigma-Aldrich) or merbarone (2 μM) (Sigma-Aldrich) and cultured at 37°C for 1 to 16h prior to subsequent experiments. Pre-incubation with ATM inhibitor (10 μM) and ATR inhibitor (10 μM) started 1h before genotoxic treatments and lasted throughout the experiment.

### Immunofluorescence, Antibodies, Westerns blots

Immunofluorescence experiments were performed as previously described (Chatzinikolaou et al., 2017; Karakasilioti et al., 2013). Briefly, cells (primary MEFs) were fixed in 4% formaldehyde, permeabilized with 0,5% Triton-X and blocked with 1% BSA. After one-hour incubation with primary antibodies, secondary fluorescent antibodies were added and DAPI was used for nuclear counterstaining. Samples were imaged with an SP8 confocal microscope (Leica). For local DNA damage infliction, cells were UV-irradiated (10 J/m^2^) through isopore polycarbonate membranes containing 3-μm-diameter pores (Millipore) and experiments were performed 2hr post-UV irradiation. Antibodies against HA (Y-11, wb: 1:500), ERCC1 (D-10, wb: 1:500, IF: 1:50), TOP2A (C-15, wb: 1:200, IF: 1:50) were from SantaCruz Biotechnology. γH2AX (05-636, IF: 1:12000) was from Millipore. TOP1 (NBP1-30482, wb: 1:1000, IF: 1:50), TOP IIβ (NB100-40842, wb: 1:1000) and 53BP1 (NB100-304, IF: 1:300) were from Novus Biologicals. TOP IIβ (20549-I-AP, IF: 1:50) was from Proteintech. TAF-4 (TAF2B9, wb: 1:500, IF:1:50), TAF-6 (TAF2G7, wb: 1:500) and TAF-10 (6TA-2B11, wb: 1:500) were from ProteoGenix. Streptavidin-HRP (wb: 1:12,000) was from Upstate Biotechnology. pATM (wb: 1:1000, IF: 1:1000) was from Rockland. pATR (wb: 1:1000, IF: 1:500) was from Genetex. FLAGM2 (F3165, wb 1:2.000, F1804, IF: 1:1000) was from Sigma-Aldrich.

### ChIP, Co-immunoprecipitation and Chromatin Pull-Down assays

For co-immunoprecipitation assays, nuclear protein extracts from primary MEFs were prepared as previously described (Chatzinikolaou et al., 2017) using the high-salt extraction method (10mM HEPES-KOH pH 7.9, 380mM KCl, 3mM MgCl2, 0.2mM EDTA, 20% glycerol and protease inhibitors). Nuclear lysates were diluted threefold by adding ice-cold HENG buffer (10mM HEPES-KOH pH 7.9, 1.5mM MgCl2, 0.25 mM EDTA, 20% glycerol) and precipitated with antibodies overnight at 4°C followed by incubation for 3 h with protein G Sepharose beads (Millipore). Normal mouse, rabbit or goat IgG (Santa Cruz) was used as a negative control. Immunoprecipitates were washed five times (10mM HEPES-KOH pH7.9, 300mM KCl, 0.3% NP40, 1.5mM MgCl2, 0.25mM EDTA, 20% glycerol and protease inhibitors), eluted and resolved on 8-12% SDS-PAGE. Pulldowns were performed with 1.2 mg of nuclear extracts using M-280 paramagnetic streptavidin beads (Invitrogen) as previously described (Chatzinikolaou et al., 2017). For ChIP assays, primary cells (MEFs) were crosslinked at R.T. for 2.5 min with 1% formaldehyde. Chromatin was prepared and sonicated on ice 15 min using Covaris S220 Focused-ultrasonicator. Samples were immunoprecipitated with antibodies (5-8 μg) overnight at 4°C followed by incubation for 3 hours with protein G-sepharose beads (Millipore) and washed sequentially. The complexes were eluted and the crosslinking was heat reversed. Purified DNA fragments were analysed by sequencing or qPCR using sets of primers targeting different regions of tRA-responsive genes. ChIP re-ChIP experiments were performed as described above with the following modifications after the first immunoprecipitation and washing, complexes were eluted with 10 mM DTT, 1% SDS in TE buffer for 30 min. Eluted samples were diluted 1:20 with re-ChIP buffer (10 mM Tris-HCl pH 8, 1 mM EDTA, 150 mM NaCl, 0.01% SDS and 1% Triton X-100) and immunoprecipitated overnight with the second antibody.

### Mass Spectrometry studies

Proteins eluted from the beads were separated by SDS/PAGE electrophoresis on a 10% polyacrylamide gel and stained with Colloidal blue silver (ThermoFisher Scientific, USA; 70). SDS-PAGE gel lanes were cut into 2-mm slices and subjected to in-gel reduction with dithiothreitol, alkylation with iodoacetamide and digested with trypsin (sequencing grade; Promega), as described previously (Schwertman et al., 2012),(Wilm et al., 1996)). Peptide mixtures were analysed by nLC-ESI-MS/MS on a LTQ-Orbitrap XL coupled to an Easy nLC (Thermo Scientific). The sample preparation and the nLC-ESI-MS/MS analysis were performed as previously described (Rappsilber et al., 2002) with minor modifications. Briefly, the dried peptides were dissolved in 0.5% formic acid aqueous solution, and the tryptic peptide mixtures were separated on a reversed-phase column (Reprosil Pur C18 AQ, Dr. Maisch GmbH), fused silica emitters 100 mm long with a 75 μm internal diameter (ThermoFisher Scientific, USA) packed in-house using a packing bomb (Loader kit SP035, Proxeon). Tryptic peptides were separated and eluted in a linear water-acetonitrile gradient and injected into the MS.

### RNA-Seq and Quantitative PCR studies

Total RNA was isolated from cells using a Total RNA isolation kit (Qiagen) as described by the manufacturer. For RNA-Seq studies, libraries were prepared using the Illumina® TruSeq® mRNA stranded sample preparation kit. Library preparation started with 1 μg total RNA. After poly-A selection (using poly-T oligo-attached magnetic beads), mRNA was purified and fragmented using divalent cations under elevated temperature. The RNA fragments underwent reverse transcription using random primers. This is followed by second strand cDNA synthesis with DNA Polymerase I and RNase H. After end repair and A-tailing, indexing adapters were ligated. The products were then purified and amplified (14 PCR cycles) to create the final cDNA libraries. After library validation and quantification (Agilent 2100 Bioanalyzer), equimolar amounts of library were pooled. The pool was quantified by using the Peqlab KAPA Library Quantification Kit and the Applied Biosystems 7900HT Sequence Detection System. The pool was sequenced by using a S2 flowcell on the Illumina NovaSeq6000 sequencer and the 2×100nt protocol. Quantitative PCR (Q-PCR) was performed with a Biorad 1000-series thermal cycler according to the instructions of the manufacturer (Biorad) as previously described (Chatzinikolaou et al., 2017). All relevant data and primer sequences for the genes tested by qPCR are available upon request.

### sBLISS and BLESS

To map DNA DSBs genome-wide, we applied an adapted set-up of the Breaks labeling *in situ* and sequencing (BLISS) method (Yan et al., 2017). In suspension BLISS (sBLISS), DSB ends are *in situ* blunted and ligated to specialized BLISS adapters that enable selective linear amplification of the genomic sequences at the DSB ends, via T7-driven *in vitro* transcription. Briefly, after cell treatment and prior to fixation, cells were washed, trypsinized and resuspended in pre-warmed PBS supplied with 10% fetal bovine serum (FBS), ensuring single-cell suspensions. Then the cells were counted and diluted to 10^6^ cells/ml and fixed with 4% paraformaldehyde aqueous solution (Electron Microscopy Sciences #15710, Formaldehyde methanol-free) for 10 minutes at room temperature (RT). Formaldehyde was quenched with 2M glycine at a final concentration of 125 mM for 5 minutes at RT, while gently rotating, and for an additional 5 minutes on ice. Fixed cells were washed with ice-cold PBS and pelleted by centrifuging at 100-400g for 10 minutes at 4°C. For *in situ* DSB labeling, 10^6^ fixed cells were incubated in a lysis buffer (10mM Tris-HCl, 10 mM NaCl, 1 mM EDTA, and 0.2% Triton X-100 (pH 8)), for 60 min on ice and the nuclei were, thereafter permeabilized with a pre-warmed permeabilization buffer (10 mM Tris-HCl, 150 mM NaCl, 1 mM EDTA, and 0.3% SDS (pH 8)) for 60 minutes at 37°C. After pelleting, the nuclei were washed twice with pre-warmed 1x CutSmart Buffer (New England Biolabs (NEB) #B7204) supplemented with 0.1% Triton X-100 (1xCS/TX100). To prepare the DSB ends for BLISS adapter ligation, the DSB ends were blunted with NEB’s Quick Blunting Kit (NEB #E1201) according to the manufacturer’s instructions in a final volume of 100 μl for 60 minutes at RT. After blunting, the nuclei were washed twice with 1x CS/TX100 before proceeding with *in situ* ligation of BLISS adapters (see below for adapter preparation). Ligation was performed with 25 Weiss units of T4 DNA Ligase (5 U/μl, ThermoFisher Scientific #EL0011) for 20-24h at 16°C in reaction volumes of 100 μl supplemented with BSA (Thermo #AM2616) and ATP (Thermo #R0441). Per preparation of 10^6^ cells, 4 μl of the selected BLISS adapter (10 μM) was ligated. Prior to use, BLISS dsDNA adapters were prepared from two complementary HPLC-purified oligonucleotides ordered from Integrated DNA Technologies (IDT). Each dsDNA adapter contains a T7 promoter sequence for *in vitro* transcription (IVT), the RA5 Illumina RNA adapter sequence for downstream sequencing, an 8-nt Unique Molecular Identifier (UMI) sequence generated by random incorporation of the four dNTPs according to IDT’s ‘Machine mixing’ strategy, and an 8-nt sample barcode to enable multiplexing of BLISS libraries. Sense oligos diluted to 10 μM in nuclease-free water were phosphorylated with T4 PNK (NEB #M0201) supplemented with ATP, after which an equimolar amount of antisense oligo was added. Oligos were annealed in a Thermocycler (5 minutes 95°C, then ramping down to 25°C in steps of 1.5°C per minute), to generate a 10 μM phosphorylated dsDNA adapter. After overnight ligation, nuclei were washed twice with 1x CS/TX100. To reverse crosslinks and extract gDNA, nuclei were resuspended in 100 μl DNA extraction buffer (10 mM Tris-HCl, 100 mM NaCl, 50 mM EDTA, and 1% SDS (pH7.5)), supplemented with 10 μl Proteinase K (800 U/ml, NEB #P8107), and incubated at 55°C for 14-18h while shaking at 800rpm. Afterwards, Proteinase K was heat-inactivated for 10 minutes at 95°C, followed by extraction using Phenol:Chloroform:Isoamyl Alcohol 25:24:1 with 10 mM Tris, pH 8.0, 1 mM EDTA (Sigma-Aldrich/Merck #P2069) and Chloroform (Merck #1024451000), followed by ethanol precipitation. The purified gDNA was resuspended in 100 μl TE and sonicated using a BioRuptor Plus (Diagenode) with settings: 30s ON, 60s OFF, HIGH intensity, 30 cycles. Sonicated DNA was concentrated with Agencourt AMPure XP beads (Beckman Coulter) and fragment sizes were assessed using a BioAnalyzer 2100 (Agilent Technologies) to range from 300bp to 800bp, with a peak around 400-600bp. To selectively and linearly amplify BLISS adapter-tagged genomic DSB ends, 100 ng of sonicated template was used for T7-mediated IVT using the MEGAscript T7 Transcription Kit (ThermoFisher #AMB13345, supplemented with Ribosafe RNAse Inhibitor (Bioline #BIO-65028)), according to the manufacturer’s guidelines. Directly after RA3 ligation, reverse transcription was performed with Reverse Transcription Primer (RTP) (Illumina sequence, ordered via IDT) and SuperScript IV Reverse Transcriptase (ThermoFisher #18090050). The manufacturer’s protocol was followed extending the incubation time to 50 minutes at 50°C followed by 10 min heat inactivation at 80°C. Library amplification was carried out with NEBNext Ultra II Q5 Master Mix (NEB #M0544), RP1 common primer, and a selected RPIX index primer (Illumina sequences, ordered through IDT). Libraries were amplified for 8 PCR cycles, purified with a 0.8x AMPure XP bead purification, and then amplified for 4 additional PCR cycles. Then, the amplified libraries were cleaned-up according to the two-sided AMPure XP bead purification protocol, aiming at retaining library sizes from ∼300–850bp. Final library profiles were assessed and quantified on a BioAnalyzer High Sensitivity DNA chip and using the Qubit dsDNA HS Assay kit (ThermoFisher #Q32851). Sequencing was performed at the Science for Life Laboratory, Sweden, on a NextSeq 500 with NextSeq 500/550 High Output Kit v2 chemistry for SE 1×75 sequencing with an additional 6 cycles for index sequencing. Multiple indexed BLISS libraries were pooled together, aiming to retrieve at least 50 million reads per condition/library. Upon completion of the run, raw sequencing reads were demultiplexed based on index sequences by Illumina’s BaseSpace, after which the generated FASTQ files were downloaded. The Breaks Labeling, Enrichment on Streptavidin and next-generation Sequencing (BLESS) validation experiments were performed according to Crosetto et al, 2013 (Crosetto et al., 2013). The procedure resembles the sBLISS protocol and includes the *in situ* blunting of DSB ends, after mild fixation of the cells, and ligation to specialized biotinylated BLESS adapters, bearing the RA5 Illumina RNA sequence, that allow the selective affinity capture of DSBs. Upon ligation of the biotinylated adapter on DSBs, genomic DNA is purified and sonicated. Then, streptavidin beads (Dynabeads MyOne C1 #65001) are used to isolate DSB-bearing DNA fragments, followed by blunting of the other end and ligation to a second BLESS adapter containing the RA3 Illumina RNA adapter sequence. PCR amplification was performed according to Illumina’s guidelines, for 10 cycles using the RA5 and RA3 adapters, followed by purification and specific-target qPCR amplification. For the modified BLESS-Western approach, we used the above BLESS protocol, except that an incubation of C1 beads in Laemmli buffer for 10min at 95°C replaced the ligation of the second adapter. The adapter sequences were previously reported for BLISS (Yan et al., 2017) and BLESS (Crosetto et al., 2013). The RA3, RA5 adapters, RTP primer, and RP1 and RPIX primers, see the sequence information available for the Illumina smallRNA library preparation kit.

### Data analysis

Statistically significant data were extracted by means of the IBM SPSS Statistics 19 (IBM) and R-statistical package (www.r-project.org). Significant over-representation of pathways and gene networks was determined by Gene Ontology (http://geneontology.org/) and KEGG pathways (https://www.genome.jp/kegg/pathway.html). For mass spectrometry (MS), the MS/MS raw data were loaded in Proteome Discoverer 1.3.0.339 (ThermoFischer Scientific) and run using the Mascot 2.3.02 (Matrix Science) search algorithm against the Mus musculus theoretical proteome (last modified 6 July 2015) containing 46,470 entries in Uniprot. A list of common contaminants was included in the database. For protein identification, the following search parameters were used: precursor error tolerance 10 ppm, fragment ion tolerance 0.8Da, trypsin full specificity, maximum number of missed cleavages 3 and cysteine alkylation as a fixed modification. The resulting .dat and .msf files were subsequently loaded and merged in Scaffold (version 3.04.05, Proteome Software) for further processing and validation of the assigned MS/MS spectra. Thresholds for protein and peptide identification were set to 99% and 95% accordingly, for proteins with minimum 1 different peptides identified, resulting in a protein false discovery rate (FDR) of <0.1%. For single peptide identifications, we applied the same criteria in addition to manual validation of MS/MS spectra. Protein lists were constructed from the respective peptide lists through extensive manual curation based on previous knowledge. For label-free relative quantitation of proteins, we applied a label-free relative quantitation method between the different samples (control versus bait) to determine unspecific binders during the affinity purification. All .dat and .msf files created by Proteome Discoverer were merged in Scaffold where label-free relative quantification was performed using the total ion current (TIC) from each identified MS/MS spectra. The TIC is the sum of the areas under all the peaks contained in a MS/MS spectrum and total TIC value results by summing the intensity of the peaks contained in the peak list associated to a MS/MS sample. Protein lists containing the Scaffold-calculated total TIC quantitative value for each protein were exported to Microsoft Excel for further manual processing including categorization and additional curation based on previous knowledge. The fold change of protein levels was calculated by dividing the mean total TIC quantitative value in bait samples with the mean value of the control samples for each of the proteins. Proteins having ≥60% protein coverage, ≥1 peptide in each sample and a fold change ≥1.2 in all three measurements were selected as being significantly enriched in bXPF compared with BirA MEF samples. Proteins that were significantly enriched in bait samples were considered these with P value ≤0.05 and a fold change ≥2. Significant over-representation of pathways, protein-protein interactions and protein complexes were derived by STRING68 (http://string-db.org). The quality of ChIP-Seq raw reads was checked using FastQC software (https://www.bioinformatics.babraham.ac.uk/projects/fastqc/). For both transcription factors (https://www.encodeproject.org/chip-seq/transcription_factor/) and histones (https://www.encodeproject.org/chip-seq/histone/) the appropriate pipelines proposed by ENCODE were adopted. All analyses were performed using as a reference the mm10 mouse genome from UCSC using Kundaje’s lab ChIP-Seq pipeline and selecting the conservative set of peaks at the end. Peak annotation was performed using the HOMER Analysis package ((Heinz et al., 2010). Simple Combinations of Lineage-Determining Transcription Factors Prime cis-Regulatory Elements Required for Macrophage and B Cell Identities). Peak visualization around TSS was performed using ChIPSeeker R package ((Yu et al., 2015)). For sBLISS, the generated amplified RNA is sequenced using next-generation sequencing, after which the obtained reads are mapped to the reference genome to identify the genomic locations of the DSBs. As described previously ((Yan et al., 2017)), a custom-built pipeline was used to keep only those reads that contain the expected prefix of 8nt UMI and 8nt sample barcode, using SAMtools and *scan for matches*, allowing at most one mismatch in the barcode sequence. The prefixes were then clipped off and stored, and the trimmed reads per condition were aligned to the GRCm38/mm10 reference genome with BWA-MEM. Only those reads with mapping quality scores ≥ 30 were retained. Next, PCR duplicates were identified and removed, by searching for proximal reads (at most 30bp apart in the reference genome) with at most two mismatches in the UMI sequence. Finally, we generated BED files for downstream analyses, comprising a list of DSB end locations and a number of unique UMIs identified at these locations, which we refer to as ‘UMI-DSB ends’ or unique DSB ends. DSBs from all samples and all replicates have been annotated using HOMER software and a generic genome distribution (Intergenic, 3UTR, miRNA, ncRNA, TTS, pseudo, Exon, Intron, Promoter, 5UTR, snoRNA, rRNA) was created. The BLISS-ChIP-Seq comparisons were performed using bedtools. The significance of difference between correlations was tested by using the tool (https://www.psychometrica.de/correlation.html) as previously described (Eid, 2014).

### Multivariate Classification Analysis

We performed a multivariate classification analysis with a binary outcome of bXPF binding to DNA (bound / unbound). As possible predictors, we employed the log values of RNA-Seq and the BLISS measurements. The tRA treatment (yes/no) was also included as a potential predictor. The analysis determines whether the predictors correlate with the bXPF binding status in a multivariate way and included feature selection by filtering out features that are either irrelevant or redundant in predicting the outcome. Since the same biological sample was measured twice e.g. one treated with tRA and one without, these measurements are not independently and identically distributed (repeated measurements). To perform the analysis we employed the “Just Add Data Bio (JAD Bio)” tool (www.jadbio.com). JAD Bio provides conservative estimates of predictive performance and corresponding confidence intervals and included the following user preferences: enforcing feature selection, not-enforcing interpretable models, using sample ID to indicate the repeated measurements, and the extensive analysis setting, the most exhaustive in terms of models it tries. The winning model did not contain the tRA treatment in the predictors as it was thrown by the feature selection step. Out of all models tested, the winning model was a Support Vector Machine model, using the full polynomial kernel of degree 2. This is equivalent to a linear model with an intercept term and predictors logRNA-Seq, logRNA-Seq^2^, logBLISS, logBLISS^2^, and the interaction term logRNA-Seq×logBLISS. The predictive performance of the model, adjusted for trying several algorithms, is 0.726 as measured by the Area Under the Receiver operating characteristic curve (AUC), with confidence interval [0.671, 0.781]. The internal workings of JAD Bio and the methods it employs were previously described (Lagani et al., 2016).

## Acknowledgements

The Horizon 2020 ERC Consolidator grant “DeFiNER” (GA64663), the FP7 Marie Curie ITN “aDDRess” (GA316390), “CodeAge” (GA316354), “Marriage” (GA316964), the Horizon 2020 Marie Curie ITN “Chromatin3D (GA GA622934), the Santé Foundation and the ELIDEK grant 1059 supported this work. GC is supported by the IKY postdoctoral research fellowship program (MIS: 5001552), co-financed by the European Social Fund- ESF and the Greek government.

## Author Contributions

GC, KS, KA, MT, EG, OC, KG, TAP, MA, PP, IT, PT, NC, JA performed the experiments and/or analyzed data. GG interpreted data and wrote the manuscript. All relevant data are available from the authors.

## Declaration of Interests

IT is affiliated with Gnosis Data Analysis that owns JAD Bio.

## References

Adair, G.M., Rolig, R.L., Moore-Faver, D., Zabelshansky, M., Wilson, J.H., and Nairn, R.S. (2000). Role of ERCC1 in removal of long non-homologous tails during targeted homologous recombination. EMBO J 19, 5552–5561.

Ahmad, A., Robinson, A.R., Duensing, A., van Drunen, E., Beverloo, H.B., Weisberg, D.B., Hasty, P., Hoeijmakers, J.H., and Niedernhofer, L.J. (2008). ERCC1-XPF endonuclease facilitates DNA double-strand break repair. Mol Cell Biol 28, 5082–5092.

Al-Minawi, A.Z., Saleh-Gohari, N., and Helleday, T. (2008). The ERCC1/XPF endonuclease is required for efficient single-strand annealing and gene conversion in mammalian cells. Nucleic Acids Res 36, 1–9.

Apostolou, Z., Chatzinikolaou, G., Stratigi, K., and Garinis, G.A. (2019). Nucleotide Excision Repair and Transcription-Associated Genome Instability. Bioessays 41, e1800201.

Awasthi, P., Foiani, M., and Kumar, A. (2015). ATM and ATR signaling at a glance. J Cell Sci 128, 4255–4262.

Aymard, F., Bugler, B., Schmidt, C.K., Guillou, E., Caron, P., Briois, S., Iacovoni, J.S., Daburon, V., Miller, K.M., Jackson, S.P., et al. (2014). Transcriptionally active chromatin recruits homologous recombination at DNA double-strand breaks. Nat Struct Mol Biol 21, 366–374.

Bastien, J., and Rochette-Egly, C. (2004). Nuclear retinoid receptors and the transcription of retinoid-target genes. Gene 328, 1–16.

Bootsma, D., Kraemer, K.H., Cleaver, J.E., Hoeijmakers, J.H.J. (1998). Nucleotide excision repair syndromes: xeroderma pigmentosum, Cockayne syndrome and trichothiodystrophy. In The genetic basis of human cancer, B. Vogelstein, Kinzler, K.W., ed. (McGraw-Hill New-York), pp. 245–274.

Bootsma, D. K., KH. Cleaver, JE. Hoeijmakers, JHJ. (2001). The Metabolic and Molecular Basis of inherited Disease (New York: McGraw-Hill).

Butuci, M., Williams, A.B., Wong, M.M., Kramer, B., and Michael, W.M. (2015). Zygotic Genome Activation Triggers Chromosome Damage and Checkpoint Signaling in C. elegans Primordial Germ Cells. Dev Cell 34, 85–95.

Calderwood, S.K. (2016). A critical role for topoisomerase IIb and DNA double strand breaks in transcription. Transcription 7, 75–83.

Canela, A., Maman, Y., Huang, S.N., Wutz, G., Tang, W., Zagnoli-Vieira, G., Callen, E., Wong, N., Day, A., Peters, J.M., et al. (2019). Topoisomerase II-Induced Chromosome Breakage and Translocation Is Determined by Chromosome Architecture and Transcriptional Activity. Mol Cell 75, 252–266 e258.

Chatzinikolaou, G., Apostolou, Z., Aid-Pavlidis, T., Ioannidou, A., Karakasilioti, I., Papadopoulos, G.L., Aivaliotis, M., Tsekrekou, M., Strouboulis, J., Kosteas, T., et al. (2017). ERCC1-XPF cooperates with CTCF and cohesin to facilitate the developmental silencing of imprinted genes. Nat Cell Biol 19, 421–432.

Chen, F., Gao, X., and Shilatifard, A. (2015). Stably paused genes revealed through inhibition of transcription initiation by the TFIIH inhibitor triptolide. Genes Dev 29, 39–47.

Cheng, J., Blum, R., Bowman, C., Hu, D., Shilatifard, A., Shen, S., and Dynlacht, B.D. (2014). A role for H3K4 monomethylation in gene repression and partitioning of chromatin readers. Mol Cell 53, 979–992.

Crosetto, N., Mitra, A., Silva, M.J., Bienko, M., Dojer, N., Wang, Q., Karaca, E., Chiarle, R., Skrzypczak, M., Ginalski, K., et al. (2013). Nucleotide-resolution DNA double-strand break mapping by next-generation sequencing. Nat Methods 10, 361–365.

de Boer, J., Andressoo, J.O., de Wit, J., Huijmans, J., Beems, R.B., van Steeg, H., Weeda, G., van der Horst, G.T., van Leeuwen, W., Themmen, A.P., et al. (2002). Premature aging in mice deficient in DNA repair and transcription. Science 296, 1276–1279.

Ding, J., Miao, Z.H., Meng, L.H., and Geng, M.Y. (2006). Emerging cancer therapeutic opportunities target DNA-repair systems. Trends in pharmacological sciences 27, 338–344.

Eid, M., Gollwitzer, M., & Schmitt, M. (2014). Hypothesis Tests for Comparing Correlations. Bibergau (Germany): Psychometrica.

Fisher, L.A., Bessho, M., and Bessho, T. (2008). Processing of a psoralen DNA interstrand cross-link by XPF-ERCC1 complex in vitro. J Biol Chem 283, 1275–1281.

Fong, Y.W., Inouye, C., Yamaguchi, T., Cattoglio, C., Grubisic, I., and Tjian, R. (2011). A DNA repair complex functions as an oct4/sox2 coactivator in embryonic stem cells. Cell 147, 120–131.

Gaillard, H., and Aguilera, A. (2016). Transcription as a Threat to Genome Integrity. Annu Rev Biochem 85, 291–317.

Gregg, S.Q., Robinson, A.R., and Niedernhofer, L.J. (2011). Physiological consequences of defects in ERCC1-XPF DNA repair endonuclease. DNA Repair (Amst) 10, 781–791.

Hanawalt, P.C. (2002). Subpathways of nucleotide excision repair and their regulation. Oncogene 21, 8949–8956.

Heinz, S., Benner, C., Spann, N., Bertolino, E., Lin, Y.C., Laslo, P., Cheng, J.X., Murre, C., Singh, H., and Glass, C.K. (2010). Simple combinations of lineage-determining transcription factors prime cis-regulatory elements required for macrophage and B cell identities. Mol Cell 38, 576–589.

Hoeijmakers, J.H. (2001). Genome maintenance mechanisms for preventing cancer. Nature 411, 366–374.

Huang, X., Gao, X., Li, W., Jiang, S., Li, R., Hong, H., Zhao, C., Zhou, P., Chen, H., Bo, X., et al. (2019). Stable H3K4me3 is associated with transcription initiation during early embryo development. Bioinformatics 35, 3931–3936.

Jaspers, N.G., Raams, A., Silengo, M.C., Wijgers, N., Niedernhofer, L.J., Robinson, A.R., Giglia-Mari, G., Hoogstraten, D., Kleijer, W.J., Hoeijmakers, J.H., et al. (2007). First reported patient with human ERCC1 deficiency has cerebro-oculo-facio-skeletal syndrome with a mild defect in nucleotide excision repair and severe developmental failure. Am J Hum Genet 80, 457–466.

Jiang, C., and Pugh, B.F. (2009). A compiled and systematic reference map of nucleosome positions across the Saccharomyces cerevisiae genome. Genome Biol 10, R109.

Ju, B.G., Lunyak, V.V., Perissi, V., Garcia-Bassets, I., Rose, D.W., Glass, C.K., and Rosenfeld, M.G. (2006). A topoisomerase IIbeta-mediated dsDNA break required for regulated transcription. Science 312, 1798–1802.

Kamileri, I., Karakasilioti, I., and Garinis, G.A. (2012a). Nucleotide excision repair: new tricks with old bricks. Trends in genetics : TIG 28, 566–573.

Kamileri, I., Karakasilioti, I., Sideri, A., Kosteas, T., Tatarakis, A., Talianidis, I., and Garinis, G.A. (2012b). Defective transcription initiation causes postnatal growth failure in a mouse model of nucleotide excision repair (NER) progeria. Proc Natl Acad Sci U S A 109, 2995–3000.

Kamileri, I., Karakasilioti, I., Sideri, A., Kosteas, T., Tatarakis, A., Talianidis, I., and Garinis, G.A. (2012c). Defective transcription initiation causes postnatal growth failure in a mouse model of nucleotide excision repair (NER) progeria. Proc Natl Acad Sci U S A 109, 2995–3000.

Kamileri, I., Karakasilioti, I., Sideri, A., Kosteas, T., Tatarakis, A., Talianidis, I., and Garinis, G.A. (2012d). Defective transcription initiation causes postnatal growth failure in a mouse model of nucleotide excision repair (NER) progeria. Proceedings of the National Academy of Sciences 109, 2995–3000.

Karakasilioti, I., Kamileri, I., Chatzinikolaou, G., Kosteas, T., Vergadi, E., Robinson, A.R., Tsamardinos, I., Rozgaja, T.A., Siakouli, S., Tsatsanis, C., et al. (2013). DNA damage triggers a chronic autoinflammatory response, leading to fat depletion in NER progeria. Cell Metab 18, 403–415.

Kashiyama, K., Nakazawa, Y., Pilz, D.T., Guo, C., Shimada, M., Sasaki, K., Fawcett, H., Wing, J.F., Lewin, S.O., Carr, L., et al. (2013). Malfunction of nuclease ERCC1-XPF results in diverse clinical manifestations and causes Cockayne syndrome, xeroderma pigmentosum, and Fanconi anemia. Am J Hum Genet 92, 807–819.

Kuraoka, I., Kobertz, W.R., Ariza, R.R., Biggerstaff, M., Essigmann, J.M., and Wood, R.D. (2000). Repair of an interstrand DNA cross-link initiated by ERCC1-XPF repair/recombination nuclease. J Biol Chem 275, 26632–26636.

Lagani, V., Athineou, G., Farcomeni, A., Tsagris, M., and Tsamardinos, I. (2016). Journal of Statistical Software Feature Selection with the R Package MXM: Discovering Statistically-Equivalent Feature Subsets. Journal of statistical software 80.

Laine, J.P., and Egly, J.M. (2006). When transcription and repair meet: a complex system. Trends in genetics : TIG 22, 430–436.

Lakiotaki, K., Georgakopoulos, G., Castanas, E., Roe, O.D., Borboudakis, G., and Tsamardinos, I. (2019). A data driven approach reveals disease similarity on a molecular level. NPJ Syst Biol Appl 5, 39.

Le May, N., Egly, J.M., and Coin, F. (2010a). True lies: the double life of the nucleotide excision repair factors in transcription and DNA repair. Journal of nucleic acids 2010.

Le May, N., Fradin, D., Iltis, I., Bougneres, P., and Egly, J.M. (2012). XPG and XPF endonucleases trigger chromatin looping and DNA demethylation for accurate expression of activated genes. Mol Cell 47, 622–632.

Le May, N., Mota-Fernandes, D., Velez-Cruz, R., Iltis, I., Biard, D., and Egly, J.M. (2010b). NER factors are recruited to active promoters and facilitate chromatin modification for transcription in the absence of exogenous genotoxic attack. Mol Cell 38, 54–66.

Le May, N., Mota-Fernandes, D., Velez-Cruz, R., Iltis, I., Biard, D., and Egly, J.M. (2010c). NER factors are recruited to active promoters and facilitate chromatin modification for transcription in the absence of exogenous genotoxic attack. Molecular cell 38, 54–66.

Li, S., Lu, H., Wang, Z., Hu, Q., Wang, H., Xiang, R., Chiba, T., and Wu, X. (2019). ERCC1/XPF Is Important for Repair of DNA Double-Strand Breaks Containing Secondary Structures. iScience 16, 63–78.

Lyu, Y.L., Lin, C.P., Azarova, A.M., Cai, L., Wang, J.C., and Liu, L.F. (2006). Role of topoisomerase IIbeta in the expression of developmentally regulated genes. Mol Cell Biol 26, 7929–7941.

Ma, J.L., Kim, E.M., Haber, J.E., and Lee, S.E. (2003). Yeast Mre11 and Rad1 proteins define a Ku-independent mechanism to repair double-strand breaks lacking overlapping end sequences. Mol Cell Biol 23, 8820–8828.

Madabhushi, R., Gao, F., Pfenning, A.R., Pan, L., Yamakawa, S., Seo, J., Rueda, R., Phan, T.X., Yamakawa, H., Pao, P.C., et al. (2015). Activity-Induced DNA Breaks Govern the Expression of Neuronal Early-Response Genes. Cell 161, 1592–1605.

Manandhar, M., Boulware, K.S., and Wood, R.D. (2015). The ERCC1 and ERCC4 (XPF) genes and gene products. Gene 569, 153–161.

Marteijn, J.A., Lans, H., Vermeulen, W., and Hoeijmakers, J.H. (2014). Understanding nucleotide excision repair and its roles in cancer and ageing. Nat Rev Mol Cell Biol 15, 465–481.

McDaniel, L.D., and Schultz, R.A. (2008). XPF/ERCC4 and ERCC1: their products and biological roles. Adv Exp Med Biol 637, 65–82.

McKenzie, A.T., Wang, M., Hauberg, M.E., Fullard, J.F., Kozlenkov, A., Keenan, A., Hurd, Y.L., Dracheva, S., Casaccia, P., Roussos, P., et al. (2018). Brain Cell Type Specific Gene Expression and Co-expression Network Architectures. Sci Rep 8, 8868.

Mori, T., Yousefzadeh, M.J., Faridounnia, M., Chong, J.X., Hisama, F.M., Hudgins, L., Mercado, G., Wade, E.A., Barghouthy, A.S., Lee, L., et al. (2018). ERCC4 variants identified in a cohort of patients with segmental progeroid syndromes. Hum Mutat 39, 255–265.

Munoz, P., Blanco, R., Flores, J.M., and Blasco, M.A. (2005). XPF nuclease-dependent telomere loss and increased DNA damage in mice overexpressing TRF2 result in premature aging and cancer. Nat Genet 37, 1063–1071.

Niedernhofer, L.J., Garinis, G.A., Raams, A., Lalai, A.S., Robinson, A.R., Appeldoorn, E., Odijk, H., Oostendorp, R., Ahmad, A., van Leeuwen, W., et al. (2006). A new progeroid syndrome reveals that genotoxic stress suppresses the somatotroph axis. Nature 444, 1038–1043.

Niraj, J., Farkkila, A., and D’Andrea, A.D. (2019). The Fanconi Anemia Pathway in Cancer. Annu Rev Canc Biol 3, 457–478.

Nouspikel, T., and Hanawalt, P.C. (2000). Terminally differentiated human neurons repair transcribed genes but display attenuated global DNA repair and modulation of repair gene expression. Mol Cell Biol 20, 1562–1570.

Ohler, U., and Wassarman, D.A. (2010). Promoting developmental transcription. Development 137, 15–26.

Pommier, Y., Sun, Y., Huang, S.N., and Nitiss, J.L. (2016). Roles of eukaryotic topoisomerases in transcription, replication and genomic stability. Nat Rev Mol Cell Biol 17, 703–721.

Rappsilber, J., Ryder, U., Lamond, A.I., and Mann, M. (2002). Large-scale proteomic analysis of the human spliceosome. Genome Res 12, 1231–1245.

Ren, G., Jin, W., Cui, K., Rodrigez, J., Hu, G., Zhang, Z., Larson, D.R., and Zhao, K. (2017). CTCF-Mediated Enhancer-Promoter Interaction Is a Critical Regulator of Cell-to-Cell Variation of Gene Expression. Mol Cell 67, 1049–1058 e1046.

Sargent, R.G., Rolig, R.L., Kilburn, A.E., Adair, G.M., Wilson, J.H., and Nairn, R.S. (1997). Recombination-dependent deletion formation in mammalian cells deficient in the nucleotide excision repair gene ERCC1. Proc Natl Acad Sci U S A 94, 13122–13127.

Schwertman, P., Lagarou, A., Dekkers, D.H., Raams, A., van der Hoek, A.C., Laffeber, C., Hoeijmakers, J.H., Demmers, J.A., Fousteri, M., Vermeulen, W., et al. (2012). UV-sensitive syndrome protein UVSSA recruits USP7 to regulate transcription-coupled repair. Nat Genet 44, 598–602.

Sijbers, A.M., de Laat, W.L., Ariza, R.R., Biggerstaff, M., Wei, Y.F., Moggs, J.G., Carter, K.C., Shell, B.K., Evans, E., de Jong, M.C., et al. (1996). Xeroderma pigmentosum group F caused by a defect in a structure-specific DNA repair endonuclease. Cell 86, 811–822.

Tian, M., Shinkura, R., Shinkura, N., and Alt, F.W. (2004). Growth retardation, early death, and DNA repair defects in mice deficient for the nucleotide excision repair enzyme XPF. Mol Cell Biol 24, 1200–1205.

Tie, F., Banerjee, R., Stratton, C.A., Prasad-Sinha, J., Stepanik, V., Zlobin, A., Diaz, M.O., Scacheri, P.C., and Harte, P.J. (2009). CBP-mediated acetylation of histone H3 lysine 27 antagonizes Drosophila Polycomb silencing. Development 136, 3131–3141.

Tripsianes, K., Folkers, G., Ab, E., Das, D., Odijk, H., Jaspers, N.G., Hoeijmakers, J.H., Kaptein, R., and Boelens, R. (2005). The structure of the human ERCC1/XPF interaction domains reveals a complementary role for the two proteins in nucleotide excision repair. Structure 13, 1849–1858.

Uuskula-Reimand, L., Hou, H., Samavarchi-Tehrani, P., Rudan, M.V., Liang, M., Medina-Rivera, A., Mohammed, H., Schmidt, D., Schwalie, P., Young, E.J., et al. (2016). Topoisomerase II beta interacts with cohesin and CTCF at topological domain borders. Genome Biol 17, 182.

van Duin, M., de Wit, J., Odijk, H., Westerveld, A., Yasui, A., Koken, M.H.M., Hoeijmakers, J.H.J., and Bootsma, D. (1986). Molecular characterization of the human excision repair gene ERCC-1: cDNA cloning and amino acid homology with the yeast DNA repair gene RAD10. Cell 44, 913–923.

Wang, L., and Eastmond, D.A. (2002). Catalytic inhibitors of topoisomerase II are DNA-damaging agents: induction of chromosomal damage by merbarone and ICRF-187. Environ Mol Mutagen 39, 348–356.

Wilm, M., Shevchenko, A., Houthaeve, T., Breit, S., Schweigerer, L., Fotsis, T., and Mann, M. (1996). Femtomole sequencing of proteins from polyacrylamide gels by nano-electrospray mass spectrometry. Nature 379, 466–469.

Woodrick, J., Gupta, S., Camacho, S., Parvathaneni, S., Choudhury, S., Cheema, A., Bai, Y., Khatkar, P., Erkizan, H.V., Sami, F., et al. (2017). A new sub-pathway of long-patch base excision repair involving 5’ gap formation. EMBO J 36, 1605–1622.

Yan, W.X., Mirzazadeh, R., Garnerone, S., Scott, D., Schneider, M.W., Kallas, T., Custodio, J., Wernersson, E., Li, Y., Gao, L., et al. (2017). BLISS is a versatile and quantitative method for genome-wide profiling of DNA double-strand breaks. Nat Commun 8, 15058.

Yankulov, K., Yamashita, K., Roy, R., Egly, J.M., and Bentley, D.L. (1995). The transcriptional elongation inhibitor 5,6-dichloro-1-beta-D-ribofuranosylbenzimidazole inhibits transcription factor IIH-associated protein kinase. J Biol Chem 270, 23922–23925.

Yu, G., Wang, L.G., and He, Q.Y. (2015). ChIPseeker: an R/Bioconductor package for ChIP peak annotation, comparison and visualization. Bioinformatics 31, 2382–2383.

Zheng, X., Kim, Y., and Zheng, Y. (2015). Identification of lamin B-regulated chromatin regions based on chromatin landscapes. Mol Biol Cell 26, 2685–2697.

Zhu, X.D., Niedernhofer, L., Kuster, B., Mann, M., Hoeijmakers, J.H., and de Lange, T. (2003). ERCC1/XPF removes the 3’ overhang from uncapped telomeres and represses formation of telomeric DNA-containing double minute chromosomes. Mol Cell 12, 1489–1498.

