## Supplementary figures and images for "ERCC1-XPF Interacts with Topoisomerase IIβ to Facilitate the Repair of Activity-induced DNA Breaks"

### Supplementary Figure 1

A.

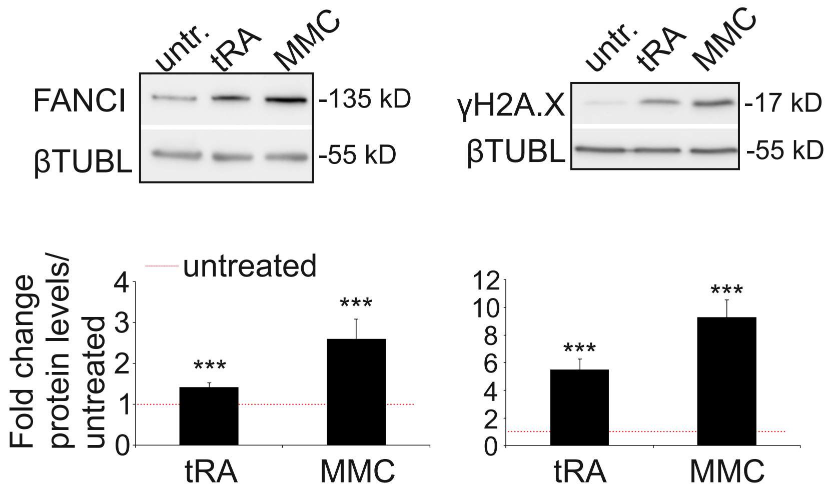


B.

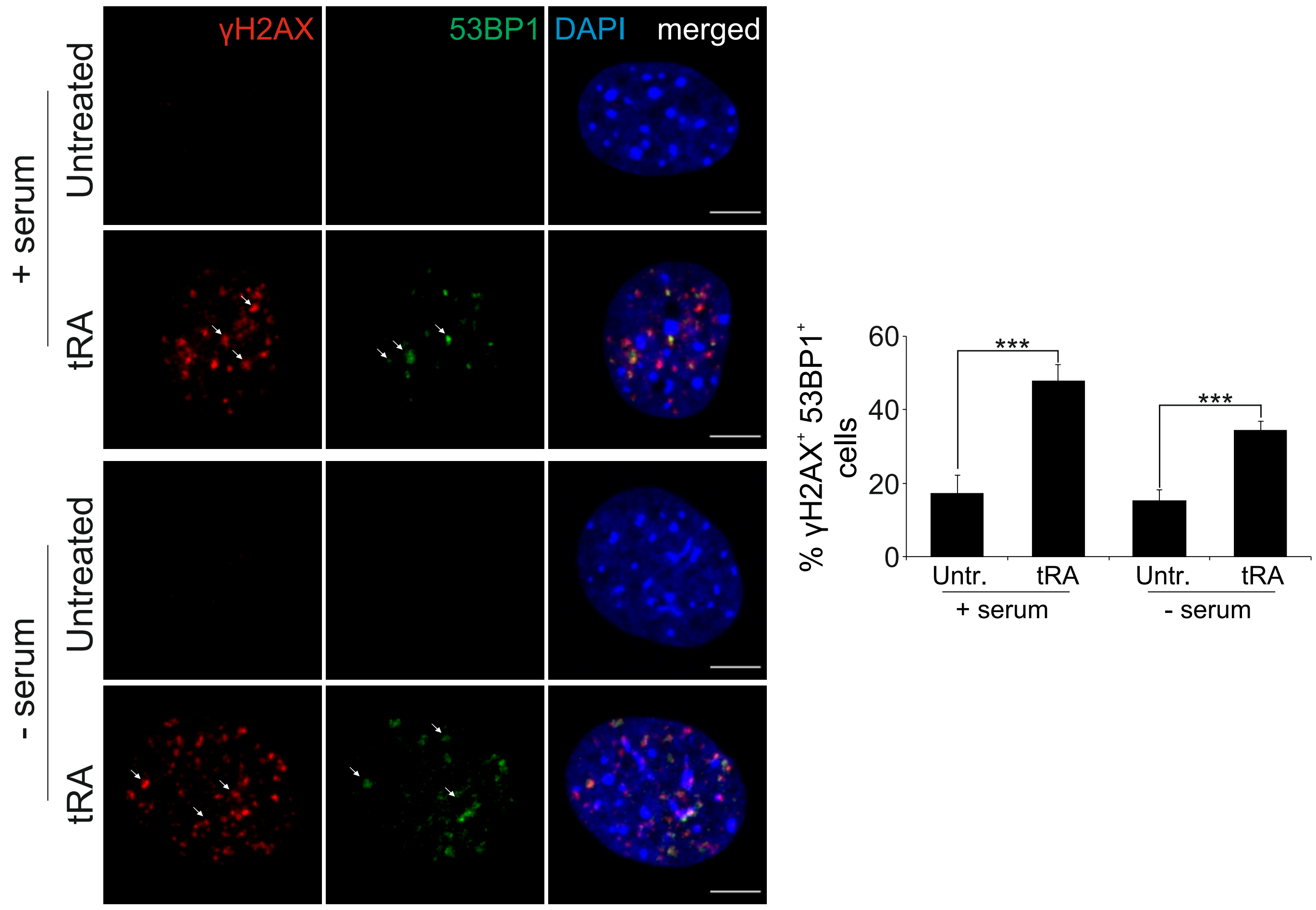


C.

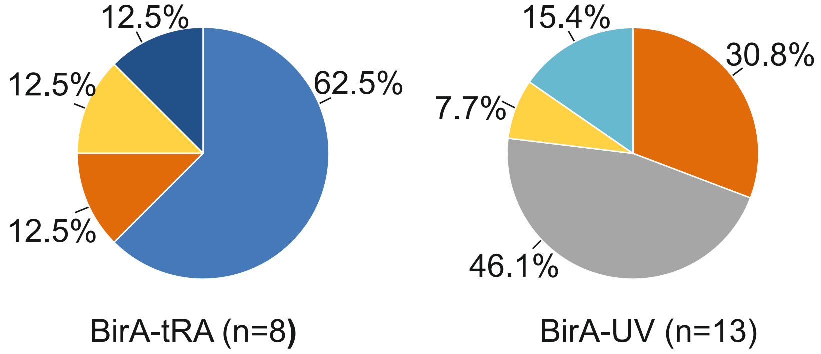


Supplementary Figure S1

### Supplementary Figure 2

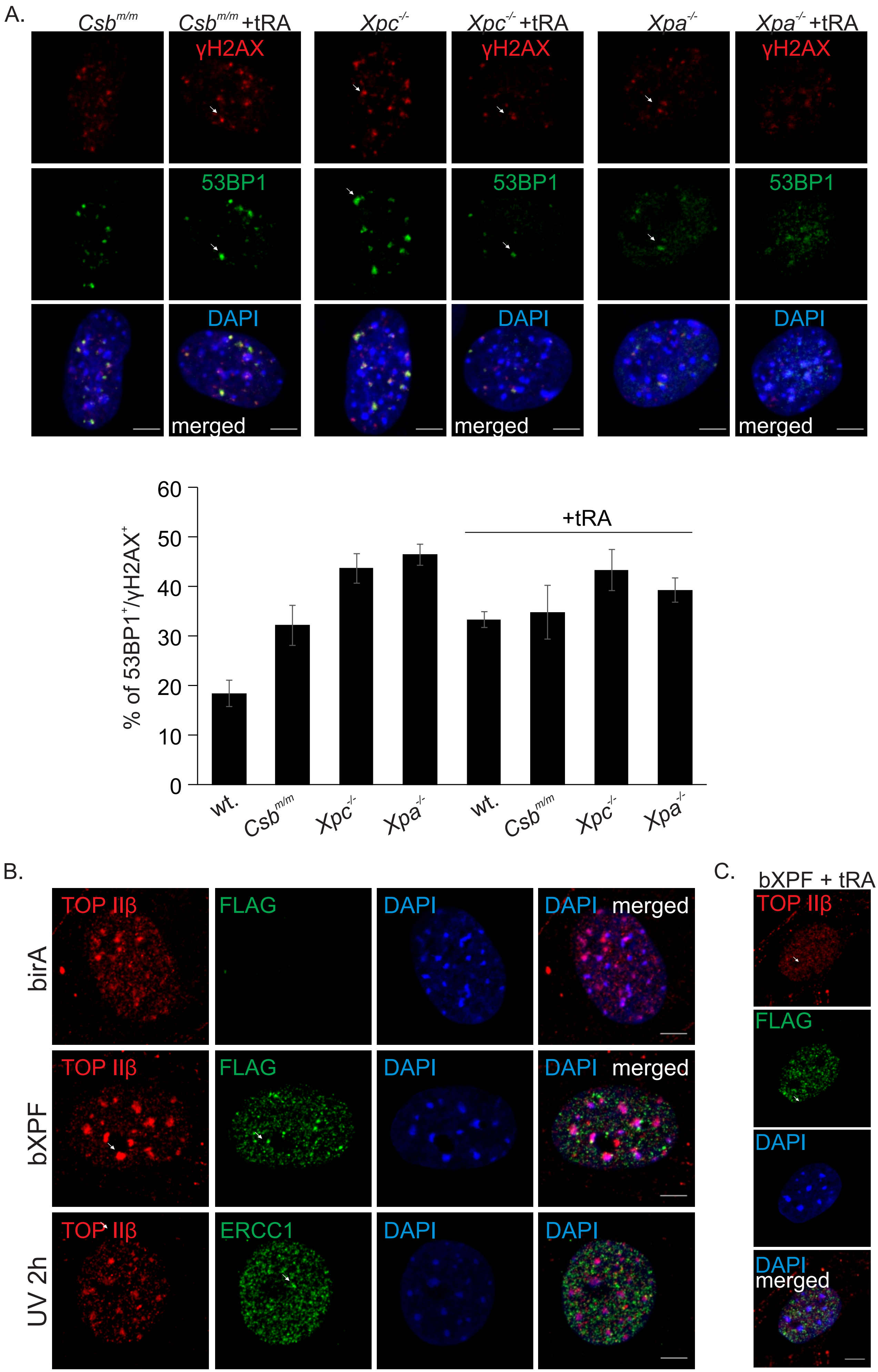


Supplementary Figure S2

### Supplementary Figure 3

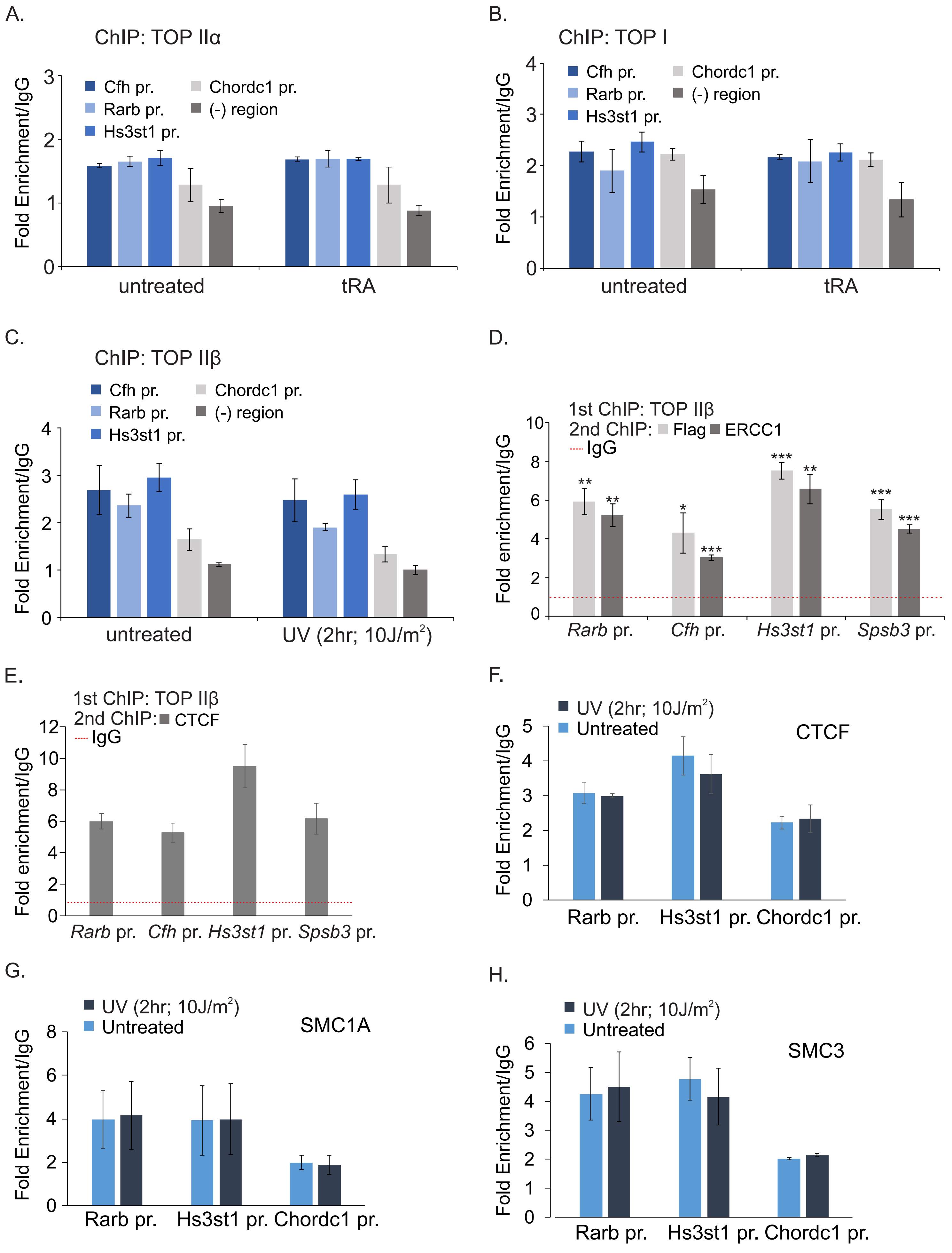


Supplementary Figure S3

### Supplementary Figure 4

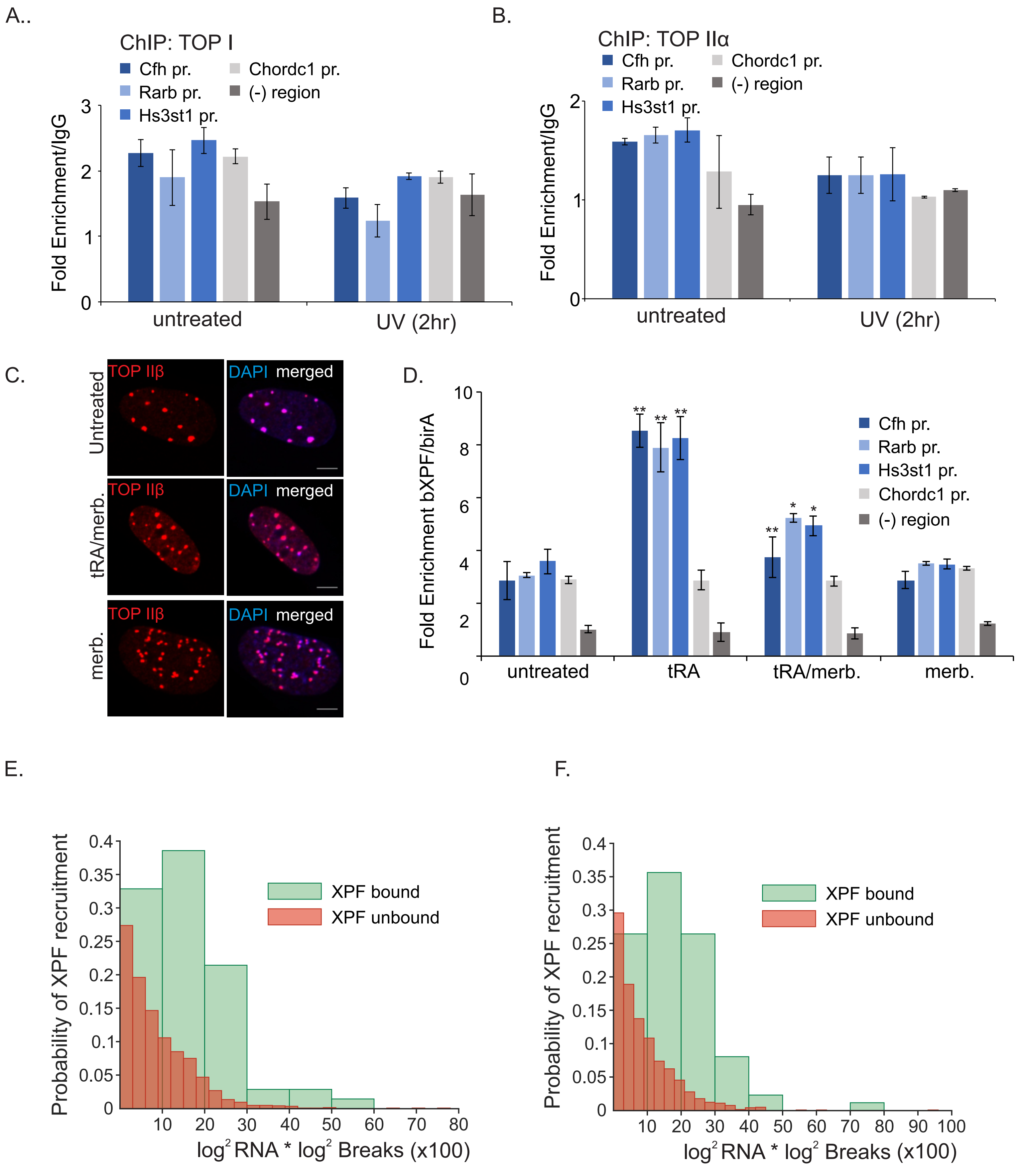


Supplementary Figure S4
