## Supplementary Table S1 for "ERCC1-XPF Interacts with Topoisomerase IIβ to Facilitate the Repair of Activity-induced DNA Breaks"

### Supplementary Table S1. bXPF and birA ChIP-Seq profiles in tRA-treated MEFs (bXPF_tRA or birA_tRA) or in UV-irradiated MEFs (bXPF_UV or birA_UV).

| PeakID | Chr | Start | End | Strand | Peak Score | Distance to TSS | Gene Name | Sample |
|---|---|---|---|---|---|---|---|---|
| 42767 | chr4 | 147421294 | 147421733 | + | 1000 | -131764 | Zfp985 | birA |
| 42887 | chr8 | 119993675 | 119994114 | + | 715 | 1456 | Crispld2 | birA |
| 42826 | chr12 | 37225308 | 37225747 | + | 639 | -16112 | Agmo | birA |
| 1 | chr11 | 82959157 | 82959596 | + | 619 | 1305 | Slfn5os | birA |
| 42856 | chr18 | 20849712 | 20850151 | + | 577 | 46147 | Trappc8 | birA |
| 42795 | chr1 | 88276317 | 88276756 | + | 558 | 1021 | A730008H23Rik | birA |
| 42917 | chr1 | 88277265 | 88277908 | + | 975 | -7 | Hjurp | birA_tRA |
| 42826 | chr5 | 151368608 | 151369251 | + | 943 | 254 | 1700028E10Rik | birA_tRA |
| 42767 | chr16 | 5049570 | 5050213 | + | 869 | 19 | Glyr1 | birA_tRA |
| 42856 | chr8 | 46551565 | 46552208 | + | 844 | -183 | Cenpu | birA_tRA |
| 42887 | chr18 | 31788768 | 31789411 | + | 720 | -70 | Polr2d | birA_tRA |
| 42795 | chr19 | 3282727 | 3283370 | + | 602 | 1 | Mrpl21 | birA_tRA |
| 1 | chr15 | 101984180 | 101984823 | + | 589 | 19841 | Krt8 | birA_tRA |
| 42948 | chr9 | 123461863 | 123462176 | + | 578 | -16682 | Limd1 | birA_tRA |
| 42826 | chr11 | 3101906 | 3102281 | + | 1000 | -21928 | Pisd-ps1 | birA_UV |
| 41275 | chr11 | 3101636 | 3102011 | + | 1000 | -22198 | Pisd-ps1 | birA_UV |
| 42767 | chr8 | 54184559 | 54184934 | + | 850 | 107214 | Vegfc | birA_UV |
| 43070 | chr1 | 88263394 | 88263769 | + | 825 | 5827 | 6430706D22Rik | birA_UV |
| 42948 | chr19 | 43689862 | 43690237 | + | 739 | 360 | Entpd7 | birA_UV |
| 43040 | chr13 | 48912795 | 48913170 | + | 682 | -42097 | Phf2 | birA_UV |
| 42856 | chr1 | 182918633 | 182919008 | + | 665 | -35968 | Tlr5 | birA_UV |
| 43009 | chr17 | 34584537 | 34584912 | + | 627 | -5082 | Gpsm3 | birA_UV |
| 42887 | chr14 | 121593566 | 121593941 | + | 609 | -88499 | Slc15a1 | birA_UV |
| 42979 | chr12 | 17881486 | 17881861 | + | 592 | 190859 | Hpcal1 | birA_UV |
| 42917 | chr12 | 69654025 | 69654400 | + | 575 | 27640 | Sos2 | birA_UV |
| 1 | chr17 | 66679990 | 66680365 | + | 559 | -85555 | Themis3 | birA_UV |
| 42795 | chr17 | 66680254 | 66680629 | + | 544 | -85819 | Themis3 | birA_UV |
| 1-1067 | chr11 | 82035278 | 82035741 | + | 1000 | -68 | Ccl2 | bXPF |
| 1-1099 | chr5 | 72855754 | 72856217 | + | 1000 | 12463 | Tec | bXPF |
| 1-1094 | chr3 | 148640466 | 148640929 | + | 1000 | 313938 | Adgrl2 | bXPF |
| 1-1097 | chr2 | 174076011 | 174076474 | + | 1000 | -809 | Stx16 | bXPF |
| 1-1095 | chr8 | 103813973 | 103814436 | + | 1000 | 210405 | 4933400L20Rik | bXPF |
| 1-1051 | chr13 | 60409789 | 60410252 | + | 1000 | -191927 | Dapk1 | bXPF |
| 1-1090 | chr16 | 21210868 | 21211331 | + | 1000 | 6304 | Ephb3 | bXPF |
| 1-199 | chr11 | 82035478 | 82035941 | + | 1000 | 132 | Ccl2 | bXPF |
| 1-1006 | chr3 | 148640665 | 148641128 | + | 1000 | 313739 | Adgrl2 | bXPF |
| 1-1098 | chr11 | 93968084 | 93968547 | + | 1000 | 206 | Nme1 | bXPF |
| 1-1093 | chr9 | 106429304 | 106429767 | + | 1000 | 124 | Rpl29 | bXPF |
| 1-1086 | chr1 | 57406394 | 57406857 | + | 1000 | 49 | Tyw5 | bXPF |
| 1-1092 | chr16 | 52067673 | 52068136 | + | 995 | 36355 | Cblb | bXPF |
| 1-1068 | chr9 | 35339462 | 35339925 | + | 989 | 58588 | 4933422A05Rik | bXPF |
| 1-168 | chr9 | 35339673 | 35340136 | + | 983 | 58799 | 4933422A05Rik | bXPF |
| 1-1035 | chr8 | 19979797 | 19980260 | + | 978 | 250452 | Gm21119 | bXPF |
| 1-1096 | chr13 | 21279566 | 21280029 | + | 972 | 12889 | Gpx5 | bXPF |
| 1-1013 | chr9 | 13633223 | 13633686 | + | 967 | 13465 | Maml2 | bXPF |
| 17899 | chr9 | 13633427 | 13633890 | + | 963 | 13669 | Maml2 | bXPF |
| 1-1078 | chr19 | 20366077 | 20366540 | + | 958 | 24363 | Anxa1 | bXPF |
| 1-1087 | chr12 | 79507170 | 79507633 | + | 954 | 210050 | Rad51b | bXPF |
| 1-1073 | chr5 | 125404070 | 125404533 | + | 950 | -14284 | Ubc | bXPF |
| 1-1046 | chr14 | 51064744 | 51065207 | + | 946 | 6467 | Olfr750 | bXPF |
| 1-1079 | chr2 | 26494328 | 26494791 | + | 942 | 9263 | Notch1 | bXPF |
| 1-1088 | chr1 | 36030525 | 36030988 | + | 938 | -37644 | Hs6st1 | bXPF |
| 1-1084 | chr8 | 46551655 | 46552118 | + | 934 | -183 | Cenpu | bXPF |
| 1-1070 | chr15 | 6934614 | 6935077 | + | 930 | -60532 | Osmr | bXPF |
| 1-1080 | chr4 | 13832945 | 13833408 | + | 926 | 48394 | Runx1t1 | bXPF |
| 1-1037 | chr4 | 143278407 | 143278870 | + | 922 | 20926 | Pdpn | bXPF |
| 1-1032 | chr15 | 76607345 | 76607808 | + | 918 | 15 | Cpsf1 | bXPF |
| 1-334 | chr10 | 83895815 | 83896278 | + | 914 | 173181 | 1500009L16Rik | bXPF |
| 1-868 | chr10 | 83895613 | 83896076 | + | 911 | 172979 | 1500009L16Rik | bXPF |

| | | | | | | | | |
|---|---|---|---|---|---|---|---|---|
| 1-614 | chr8 | 126603245 | 126603708 | + | 907 | -10040 | Irf2bp2 | bXPF |
| 1-1026 | chr13 | 28417706 | 28418169 | + | 904 | 275453 | Prl5a1 | bXPF |
| 1-1076 | chr8 | 123442959 | 123443422 | + | 901 | -69 | Def8 | bXPF |
| 1-1008 | chr3 | 36449319 | 36449782 | + | 898 | 25974 | Mir7009 | bXPF |
| 1-1074 | chr8 | 128025137 | 128025600 | + | 895 | 252857 | Mir21c | bXPF |
| 1-1047 | chr3 | 95658480 | 95658943 | + | 892 | -10 | Mcl1 | bXPF |
| 1-1058 | chr13 | 60019811 | 60020274 | + | 889 | -9669 | A530065N20Rik | bXPF |
| 1-997 | chr4 | 119539173 | 119539636 | + | 886 | -257 | Foxj3 | bXPF |
| 27395 | chr4 | 119539380 | 119539843 | + | 884 | -50 | Foxj3 | bXPF |
| 1-1066 | chr14 | 16554559 | 16555022 | + | 881 | 20686 | Rarb | bXPF |
| 1-1060 | chr19 | 32974830 | 32975293 | + | 879 | 217484 | Pten | bXPF |
| 1-1040 | chr14 | 51089321 | 51089784 | + | 876 | -1525 | Rnase4 | bXPF |
| 1-1007 | chr12 | 83631978 | 83632441 | + | 874 | -25 | Rbm25 | bXPF |
| 1-1075 | chr19 | 3282670 | 3283133 | + | 872 | 0 | Mrpl21 | bXPF |
| 1-1064 | chr2 | 20453820 | 20454283 | + | 870 | -55996 | Etl4 | bXPF |
| 1-1082 | chr9 | 43945429 | 43945892 | + | 867 | -97724 | Thy1 | bXPF |
| 1-1002 | chr13 | 15741589 | 15742052 | + | 865 | 60810 | 4933412O06Rik | bXPF |
| 1-953 | chr4 | 43492778 | 43493241 | + | 863 | 50 | Rmrp | bXPF |
| 1-1052 | chr2 | 118532986 | 118533449 | + | 861 | 11370 | Bmf | bXPF |
| 1-824 | chr1 | 82290798 | 82291261 | + | 860 | 410 | Irs1 | bXPF |
| 1-1034 | chr2 | 32549696 | 32550159 | + | 858 | 14568 | Fam102a | bXPF |
| 1-1015 | chr5 | 77080858 | 77081321 | + | 856 | 14178 | Hopx | bXPF |
| 1-139 | chr5 | 72855959 | 72856422 | + | 854 | 12258 | Tec | bXPF |
| 1-1049 | chr3 | 49642396 | 49642859 | + | 853 | 114689 | Pcdh18 | bXPF |
| 1-111 | chr3 | 49642597 | 49643060 | + | 851 | 114488 | Pcdh18 | bXPF |
| 1-1025 | chr18 | 33377868 | 33378331 | + | 849 | 85535 | Nrep | bXPF |
| 1-951 | chr14 | 76681671 | 76682134 | + | 847 | 4334 | 1700108F19Rik | bXPF |
| 35796 | chr3 | 95658692 | 95659155 | + | 845 | 202 | Mcl1 | bXPF |
| 1-1017 | chr13 | 80962848 | 80963311 | + | 844 | -995 | 9330111N05Rik | bXPF |
| 1-1024 | chr6 | 124740864 | 124741327 | + | 842 | -16 | Grcc10 | bXPF |
| 1-956 | chr9 | 21592654 | 21593117 | + | 840 | -54 | Yipf2 | bXPF |
| 1-979 | chr4 | 155943487 | 155943950 | + | 838 | -95 | Ube2j2 | bXPF |
| 1-1044 | chr13 | 51314601 | 51315064 | + | 837 | -93786 | S1pr3 | bXPF |
| 1-1085 | chr7 | 70267062 | 70267525 | + | 835 | 70529 | Gm29683 | bXPF |
| 1-1069 | chr8 | 109778337 | 109778800 | + | 833 | -15 | Ap1g1 | bXPF |
| 1-1001 | chr6 | 71632994 | 71633457 | + | 832 | -308 | Kdm3a | bXPF |
| 1-977 | chr2 | 160525934 | 160526397 | + | 830 | -119732 | Top1 | bXPF |
| 1-1023 | chr15 | 50698172 | 50698635 | + | 829 | 190646 | Trps1 | bXPF |
| 1-1077 | chr9 | 79759621 | 79760084 | + | 827 | 1 | Cox7a2 | bXPF |
| 1-235 | chr7 | 99360429 | 99360892 | + | 826 | -7421 | Serpinh1 | bXPF |
| 1-1072 | chr7 | 99360228 | 99360691 | + | 824 | -7220 | Serpinh1 | bXPF |
| 1-1089 | chr6 | 38144370 | 38144833 | + | 823 | -20015 | Atp6v0a4 | bXPF |
| 1-913 | chr16 | 24495446 | 24495909 | + | 822 | -33637 | Morf4l1-ps1 | bXPF |
| 1-514 | chr8 | 36613931 | 36614394 | + | 821 | -219 | Dlc1 | bXPF |
| 16072 | chr2 | 173351987 | 173352450 | + | 819 | -75685 | Pmepa1 | bXPF |
| 1-916 | chr14 | 31669639 | 31670102 | + | 818 | 28813 | Btd | bXPF |
| 1-236 | chr2 | 173351788 | 173352251 | + | 817 | -75486 | Pmepa1 | bXPF |
| 1-165 | chr12 | 79507371 | 79507834 | + | 815 | 210251 | Rad51b | bXPF |
| 1-1021 | chr2 | 60689025 | 60689488 | + | 814 | -15563 | Itgb6 | bXPF |
| 33604 | chr8 | 120589163 | 120589626 | + | 813 | -319 | Gins2 | bXPF |
| 1-1081 | chr8 | 120588956 | 120589419 | + | 812 | -112 | Gins2 | bXPF |
| 18994 | chr16 | 24619595 | 24620058 | + | 811 | 90512 | Morf4l1-ps1 | bXPF |
| 1-1031 | chr13 | 54750698 | 54751161 | + | 809 | -1055 | Gprin1 | bXPF |
| 1-963 | chr16 | 24619396 | 24619859 | + | 808 | 90313 | Morf4l1-ps1 | bXPF |
| 1-990 | chr17 | 83990351 | 83990814 | + | 807 | -7970 | 8430430B14Rik | bXPF |
| 1-954 | chr13 | 91165053 | 91165516 | + | 806 | 58703 | Atg10 | bXPF |
| 1-818 | chr8 | 104023028 | 104023491 | + | 805 | -78366 | Cdh5 | bXPF |
| 1-1036 | chr9 | 57831245 | 57831708 | + | 804 | 2758 | Arid3b | bXPF |
| 1-1048 | chr15 | 10890126 | 10890589 | + | 803 | 61306 | Gm19276 | bXPF |
| 1-1050 | chr1 | 20069641 | 20070104 | + | 802 | -286582 | 4930486I03Rik | bXPF |
| 1-993 | chr4 | 137048462 | 137048925 | + | 798 | 2 | Zbtb40 | bXPF |
| 1-952 | chr1 | 152556035 | 152556498 | + | 797 | 68951 | Rgl1 | bXPF |
| 1-1061 | chr4 | 58271677 | 58272140 | + | 796 | -14054 | Musk | bXPF |

| | | | | | | | | |
|---|---|---|---|---|---|---|---|---|
| 1-943 | chr2 | 83760073 | 83760536 | + | 795 | 35907 | Itgav | bXPF |
| 1-998 | chr15 | 27782596 | 27783059 | + | 793 | -101285 | Fam105a | bXPF |
| 1-1057 | chr1 | 74386787 | 74387250 | + | 792 | -4591 | Ctdsp1 | bXPF |
| 24473 | chr10 | 13553101 | 13553564 | + | 791 | -190 | Pex3 | bXPF |
| 1-960 | chr10 | 13552901 | 13553364 | + | 790 | 10 | Pex3 | bXPF |
| 1-1038 | chr5 | 136403546 | 136404009 | + | 789 | -27974 | Mir721 | bXPF |
| 1-1045 | chr3 | 130086633 | 130087096 | + | 788 | -25957 | Sec24b | bXPF |
| 1-225 | chr8 | 72136688 | 72137151 | + | 787 | 1627 | Tpm4 | bXPF |
| 1-991 | chr8 | 72136489 | 72136952 | + | 786 | 1428 | Tpm4 | bXPF |
| 1-1083 | chr3 | 148989179 | 148989642 | + | 785 | -34775 | Adgrl2 | bXPF |
| 1-936 | chr17 | 26584620 | 26585083 | + | 784 | 23339 | Ergic1 | bXPF |
| 1-927 | chr1 | 45043173 | 45043636 | + | 783 | -268134 | Col3a1 | bXPF |
| 1-924 | chr6 | 71990037 | 71990500 | + | 782 | 52260 | 9130221F21Rik | bXPF |
| 1-400 | chr14 | 54443351 | 54443814 | + | 781 | 11984 | Mmp14 | bXPF |
| 31413 | chr14 | 21985060 | 21985523 | + | 780 | 4310 | Zfp503 | bXPF |
| 1-1100 | chr8 | 19784536 | 19784787 | + | 779 | 55085 | Gm21119 | bXPF |
| 1-441 | chr14 | 16554760 | 16555223 | + | 778 | 20485 | Rarb | bXPF |
| 1-985 | chr10 | 37139160 | 37139623 | + | 777 | -465 | Marcks | bXPF |
| 18264 | chr10 | 37139371 | 37139834 | + | 776 | -676 | Marcks | bXPF |
| 1-1011 | chr18 | 75366556 | 75367019 | + | 775 | -578 | Smad7 | bXPF |
| 1-958 | chr6 | 100503310 | 100503773 | + | 774 | -23859 | 1700049E22Rik | bXPF |
| 1-934 | chr13 | 104816963 | 104817426 | + | 774 | 24 | Srek1ip1 | bXPF |
| 1-1029 | chr9 | 87021671 | 87022134 | + | 773 | -127 | Cyb5r4 | bXPF |
| 1-259 | chr1 | 45672770 | 45673233 | + | 772 | -122500 | Wdr75 | bXPF |
| 1-754 | chr1 | 45672570 | 45673033 | + | 771 | -122700 | Wdr75 | bXPF |
| 1-987 | chr4 | 84158124 | 84158587 | + | 770 | -113476 | 6030471H07Rik | bXPF |
| 1-938 | chr1 | 163362107 | 163362570 | + | 769 | 41331 | Gorab | bXPF |
| 1-876 | chr5 | 44586577 | 44587040 | + | 768 | 212938 | Ldb2 | bXPF |
| 1-146 | chr5 | 44586782 | 44587245 | + | 768 | 212733 | Ldb2 | bXPF |
| 1-965 | chr1 | 83900159 | 83900622 | + | 766 | 318571 | Mir6353 | bXPF |
| 23012 | chr5 | 125184960 | 125185423 | + | 765 | -5977 | Ncor2 | bXPF |
| 1-807 | chr3 | 51358133 | 51358596 | + | 764 | -17720 | Elf2 | bXPF |
| 1-742 | chr5 | 125184759 | 125185222 | + | 764 | -5776 | Ncor2 | bXPF |
| 1-1012 | chr2 | 127208108 | 127208571 | + | 763 | -59 | 1810024B03Rik | bXPF |
| 1-790 | chr16 | 90882521 | 90882984 | + | 762 | 51893 | Eva1c | bXPF |
| 1-140 | chr18 | 21299636 | 21300099 | + | 761 | 272 | Garem1 | bXPF |
| 1-945 | chr16 | 90453160 | 90453623 | + | 761 | 66994 | Hunk | bXPF |
| 1-966 | chr7 | 130215433 | 130215896 | + | 760 | 51144 | Fgfr2 | bXPF |
| 1-527 | chr15 | 66861654 | 66862117 | + | 759 | -29508 | Wisp1 | bXPF |
| 1-1019 | chr11 | 102556122 | 102556585 | + | 758 | -195 | Gpatch8 | bXPF |
| 1-177 | chr15 | 66861865 | 66862328 | + | 758 | -29297 | Wisp1 | bXPF |
| 1-925 | chr4 | 14273626 | 14274089 | + | 757 | 17452 | Gm2560 | bXPF |
| 1-859 | chr3 | 58168478 | 58168941 | + | 756 | -4902 | 1700007F19Rik | bXPF |
| 1-937 | chr9 | 105053053 | 105053516 | + | 755 | 16 | Mrpl3 | bXPF |
| 1-150 | chr9 | 105053258 | 105053721 | + | 755 | 221 | Mrpl3 | bXPF |
| 1-995 | chr4 | 43394535 | 43394998 | + | 754 | -11736 | Rusc2 | bXPF |
| 1-192 | chr10 | 80249191 | 80249654 | + | 753 | -30 | Ndufs7 | bXPF |
| 1-888 | chr5 | 151368632 | 151369095 | + | 752 | 188 | 1700028E10Rik | bXPF |
| 1-920 | chr9 | 62560022 | 62560485 | + | 752 | -23209 | Coro2b | bXPF |
| 1-608 | chr10 | 80248984 | 80249447 | + | 751 | -237 | Ndufs7 | bXPF |
| 1-923 | chr17 | 67241826 | 67242289 | + | 750 | 112402 | Ptprm | bXPF |
| 1-730 | chr13 | 98815289 | 98815752 | + | 750 | -71 | Fcho2 | bXPF |
| 1-978 | chr17 | 88628819 | 88629282 | + | 749 | 2495 | Ston1 | bXPF |
| 1-1016 | chr19 | 31082675 | 31083138 | + | 748 | 65 | Cstf2t | bXPF |
| 43831 | chr12 | 86800682 | 86801145 | + | 748 | 66544 | Lrrc74a | bXPF |
| 1-816 | chr4 | 84523659 | 84524122 | + | 747 | 151196 | Bnc2 | bXPF |
| 1-970 | chr18 | 52613745 | 52614208 | + | 746 | -1939 | Zfp474 | bXPF |
| 1-1033 | chr6 | 134035207 | 134035670 | + | 746 | -262 | Etv6 | bXPF |
| 1-971 | chr11 | 86380426 | 86380889 | + | 745 | -23132 | Med13 | bXPF |
| 1-872 | chr9 | 68813265 | 68813728 | + | 744 | 48150 | 9530091C08Rik | bXPF |
| 1-989 | chr9 | 21496642 | 21497105 | + | 744 | -309 | Mir199a-1 | bXPF |
| 1-1000 | chr13 | 51259016 | 51259479 | + | 743 | -56181 | Hist1h2al | bXPF |
| 1-861 | chr18 | 13620920 | 13621383 | + | 742 | 351582 | Zfp521 | bXPF |

| | | | | | | | | |
|---|---|---|---|---|---|---|---|---|
| 1-1055 | chr5 | 142821547 | 142822010 | + | 742 | -4391 | Tnrc18 | bXPF |
| 1-321 | chr3 | 142158948 | 142159411 | + | 741 | 10049 | Bmpr1b | bXPF |
| 1-836 | chr3 | 19628618 | 19629081 | + | 741 | -15611 | Trim55 | bXPF |
| 1-696 | chr5 | 31526651 | 31527114 | + | 740 | -113 | Slc4a1ap | bXPF |
| 1-899 | chr11 | 60416980 | 60417443 | + | 739 | 66 | Gid4 | bXPF |
| 1-688 | chr2 | 167569914 | 167570377 | + | 739 | 31950 | Snai1 | bXPF |
| 1-786 | chr9 | 84369074 | 84369537 | + | 738 | -405093 | 4930554C24Rik | bXPF |
| 1-488 | chr18 | 39581940 | 39582403 | + | 738 | -94926 | Nr3c1 | bXPF |
| 1-893 | chr7 | 141476445 | 141476908 | + | 737 | 296 | Tspan4 | bXPF |
| 1-967 | chr9 | 65884886 | 65885349 | + | 736 | 56 | Trip4 | bXPF |
| 1-747 | chr11 | 75046952 | 75047415 | + | 736 | 18808 | Mir684-1 | bXPF |
| 1-250 | chr1 | 174617460 | 174617923 | + | 735 | 115866 | Fmn2 | bXPF |
| 1-709 | chr1 | 174617245 | 174617708 | + | 735 | 115651 | Fmn2 | bXPF |
| 1-1009 | chr13 | 23466869 | 23467332 | + | 734 | -1199 | Btn1a1 | bXPF |
| 1-1028 | chr16 | 30187880 | 30188343 | + | 734 | 79421 | Cpn2 | bXPF |
| 1-750 | chr6 | 128089695 | 128090158 | + | 733 | 53652 | Tspan9 | bXPF |
| 1-976 | chr11 | 48818261 | 48818724 | + | 732 | -1101 | Trim41 | bXPF |
| 1-891 | chr6 | 99485041 | 99485504 | + | 732 | 35721 | Foxp1 | bXPF |
| 1-293 | chr13 | 21160625 | 21161088 | + | 731 | -17801 | Olfr1368 | bXPF |
| 1-840 | chr7 | 99352982 | 99353445 | + | 731 | 26 | Serpinh1 | bXPF |
| 1-911 | chr2 | 51145971 | 51146434 | + | 730 | 2909 | Rnd3 | bXPF |
| 1-950 | chr10 | 30810868 | 30811331 | + | 730 | -7992 | Ncoa7 | bXPF |
| 1-812 | chr10 | 86425679 | 86426142 | + | 729 | 72986 | Syn3 | bXPF |
| 1-919 | chr12 | 31218575 | 31219038 | + | 729 | -46488 | Lamb1 | bXPF |
| 1-955 | chr1 | 51478346 | 51478809 | + | 728 | -178 | Nabp1 | bXPF |
| 1-804 | chr12 | 79600165 | 79600628 | + | 728 | 303045 | Rad51b | bXPF |
| 1-982 | chr2 | 4362125 | 4362588 | + | 727 | -38620 | Frmd4a | bXPF |
| 1-757 | chr15 | 83166526 | 83166989 | + | 726 | 5451 | Cyb5r3 | bXPF |
| 1-1014 | chr19 | 43752626 | 43753089 | + | 726 | 143 | Cox15 | bXPF |
| 1-897 | chr8 | 79027873 | 79028336 | + | 725 | -333 | Zfp827 | bXPF |
| 1-932 | chr5 | 92434935 | 92435398 | + | 725 | 33 | Nup54 | bXPF |
| 1-864 | chr17 | 31982803 | 31983266 | + | 724 | 51474 | Hsf2bp | bXPF |
| 1-890 | chr13 | 40733554 | 40734017 | + | 724 | 38 | Tfap2a | bXPF |
| 1-905 | chr9 | 110790362 | 110790825 | + | 723 | -7592 | Prss42 | bXPF |
| 1-667 | chr12 | 101852043 | 101852506 | + | 723 | -33155 | Fbln5 | bXPF |
| 1-154 | chr1 | 120757512 | 120757975 | + | 722 | 155256 | En1 | bXPF |
| 1-1039 | chr7 | 75427315 | 75427778 | + | 722 | -27988 | Akap13 | bXPF |
| 1-928 | chr5 | 100798440 | 100798903 | + | 721 | -71 | Helq | bXPF |
| 1-267 | chr1 | 120757305 | 120757768 | + | 721 | 155049 | En1 | bXPF |
| 1-1030 | chr17 | 44261788 | 44262251 | + | 720 | 73447 | Clic5 | bXPF |
| 1-792 | chr4 | 153974833 | 153975296 | + | 720 | 17 | Dffb | bXPF |
| 1-1091 | chr16 | 15195279 | 15195742 | + | 719 | 288864 | Efcab1 | bXPF |
| 1-380 | chr14 | 20805096 | 20805559 | + | 719 | -11239 | Camk2g | bXPF |
| 1-795 | chr6 | 22280460 | 22280923 | + | 718 | -7536 | Wnt16 | bXPF |
| 1-806 | chr8 | 72160597 | 72161060 | + | 718 | -372 | Rab8a | bXPF |
| 1-736 | chr6 | 37617263 | 37617726 | + | 717 | 87321 | Akr1d1 | bXPF |
| 1-910 | chr1 | 45675656 | 45676119 | + | 717 | -119614 | Wdr75 | bXPF |
| 1-114 | chr13 | 80963049 | 80963512 | + | 716 | -794 | 9330111N05Rik | bXPF |
| 1-247 | chr13 | 23517507 | 23517970 | + | 716 | -13306 | Hist1h4h | bXPF |
| 1-665 | chr17 | 34121117 | 34121580 | + | 715 | 399 | Brd2 | bXPF |
| 1-862 | chr5 | 4323221 | 4323684 | + | 715 | 131085 | Mterf1b | bXPF |
| 1-195 | chr10 | 39613376 | 39613839 | + | 714 | 673 | Traf3ip2 | bXPF |
| 1-636 | chr16 | 92139319 | 92139782 | + | 713 | 81214 | Mrps6 | bXPF |
| 1-740 | chr15 | 51798568 | 51799031 | + | 713 | 66662 | Eif3h | bXPF |
| 1-719 | chr15 | 59823185 | 59823648 | + | 712 | -28457 | Gm19510 | bXPF |
| 1-801 | chr18 | 63529819 | 63530282 | + | 712 | -142867 | Piezo2 | bXPF |
| 1-629 | chr12 | 73147162 | 73147625 | + | 711 | 23676 | Mnat1 | bXPF |
| 1-895 | chr6 | 38381219 | 38381682 | + | 711 | -74 | Ttc26 | bXPF |
| 1-563 | chr5 | 134688660 | 134689123 | + | 710 | -301 | Limk1 | bXPF |
| 15342 | chr5 | 134688873 | 134689336 | + | 709 | -514 | Limk1 | bXPF |
| 1-252 | chr15 | 50698384 | 50698847 | + | 709 | 190434 | Trps1 | bXPF |
| 1-116 | chr4 | 32303402 | 32303865 | + | 708 | -113802 | Bach2 | bXPF |
| 1-729 | chr6 | 94700098 | 94700561 | + | 708 | -184 | Lrig1 | bXPF |

| | | | | | | | | |
|---|---|---|---|---|---|---|---|---|
| 1-641 | chr2 | 94010403 | 94010866 | + | 707 | 96 | Alkbh3 | bXPF |
| 1-135 | chr4 | 32303010 | 32303473 | + | 707 | -114194 | Bach2 | bXPF |
| 1-832 | chr3 | 144720293 | 144720756 | + | 706 | -189 | Sh3glb1 | bXPF |
| 14611 | chr3 | 144720495 | 144720958 | + | 706 | -391 | Sh3glb1 | bXPF |
| 1-743 | chr7 | 95914699 | 95915162 | + | 706 | -256314 | Tenm4 | bXPF |
| 1-1062 | chr5 | 149320485 | 149320948 | + | 705 | 55712 | Alox5ap | bXPF |
| 1-226 | chr2 | 94010623 | 94011086 | + | 705 | -124 | Alkbh3 | bXPF |
| 1-964 | chr6 | 145968274 | 145968737 | + | 704 | 34358 | Sspn | bXPF |
| 1-679 | chr15 | 36792565 | 36793028 | + | 703 | 1742 | Ywhaz | bXPF |
| 1-900 | chr2 | 163675151 | 163675614 | + | 703 | 16924 | Pkig | bXPF |
| 1-726 | chr9 | 66834751 | 66835214 | + | 702 | -276 | Aph1c | bXPF |
| 1-203 | chr16 | 30063751 | 30064214 | + | 702 | -1375 | Hes1 | bXPF |
| 1-856 | chr6 | 88343352 | 88343815 | + | 702 | 102956 | Eefsec | bXPF |
| 1-877 | chr2 | 44559389 | 44559852 | + | 701 | 302002 | Gtdc1 | bXPF |
| 1-983 | chr18 | 78144727 | 78145190 | + | 701 | -2839 | Slc14a1 | bXPF |
| 1-212 | chr1 | 152556243 | 152556706 | + | 700 | 68743 | Rgl1 | bXPF |
| 1-698 | chr2 | 130412034 | 130412497 | + | 700 | 5743 | Tmem239 | bXPF |
| 1-984 | chr1 | 180850585 | 180851048 | + | 699 | -335 | Sde2 | bXPF |
| 28126 | chr1 | 180850785 | 180851248 | + | 699 | -135 | Sde2 | bXPF |
| 1-701 | chr7 | 70345891 | 70346354 | + | 699 | -8270 | Gm29683 | bXPF |
| 1-556 | chr1 | 192785549 | 192786012 | + | 698 | -14561 | Hhat | bXPF |
| 1-543 | chr4 | 141790524 | 141790987 | + | 698 | -111 | Dnajc16 | bXPF |
| 1-104 | chr4 | 141790732 | 141791195 | + | 697 | -319 | Dnajc16 | bXPF |
| 1-595 | chr13 | 38410460 | 38410923 | + | 697 | 51102 | 4930579J19Rik | bXPF |
| 1-874 | chr7 | 84329086 | 84329549 | + | 697 | 80721 | Arnt2 | bXPF |
| 1-735 | chr10 | 68279917 | 68280380 | + | 696 | -1422 | Arid5b | bXPF |
| 1-141 | chr10 | 68280118 | 68280581 | + | 696 | -1623 | Arid5b | bXPF |
| 1-793 | chr11 | 106065758 | 106066221 | + | 695 | -118 | Taco1 | bXPF |
| 1-901 | chr1 | 151686662 | 151687125 | + | 695 | -68481 | Edem3 | bXPF |
| 1-962 | chr3 | 126831420 | 126831883 | + | 695 | 111735 | Ank2 | bXPF |
| 1-758 | chr11 | 118327452 | 118327915 | + | 694 | 14851 | BC100451 | bXPF |
| 1-598 | chr6 | 136468362 | 136468825 | + | 694 | -50258 | Atf7ip | bXPF |
| 1-999 | chr13 | 43171174 | 43171637 | + | 693 | 96 | Tbc1d7 | bXPF |
| 1-704 | chr10 | 7589505 | 7589968 | + | 693 | -64 | Lrp11 | bXPF |
| 1-981 | chr9 | 62828678 | 62829141 | + | 693 | -9878 | Cln6 | bXPF |
| 1-809 | chr10 | 111972478 | 111972941 | + | 692 | 14 | Krr1 | bXPF |
| 1-988 | chr1 | 192798444 | 192798907 | + | 692 | -27456 | Hhat | bXPF |
| 1-435 | chr5 | 34169461 | 34169924 | + | 692 | -166 | Poln | bXPF |
| 1-490 | chr14 | 121171972 | 121172435 | + | 691 | 136629 | Farp1 | bXPF |
| 1-322 | chr14 | 121172181 | 121172644 | + | 691 | 136838 | Farp1 | bXPF |
| 1-127 | chr11 | 86646381 | 86646844 | + | 690 | -9459 | Mir8115 | bXPF |
| 1-523 | chr11 | 86646181 | 86646644 | + | 690 | -9659 | Mir8115 | bXPF |
| 1-959 | chr15 | 64376958 | 64377421 | + | 690 | 5730 | Asap1 | bXPF |
| 1-748 | chr16 | 24490680 | 24491143 | + | 689 | -38403 | Morf4l1-ps1 | bXPF |
| 1-738 | chr4 | 59231275 | 59231738 | + | 689 | -28545 | Gm12596 | bXPF |
| 1-912 | chr9 | 59486576 | 59487039 | + | 689 | -433 | Arih1 | bXPF |
| 1-335 | chr6 | 47650639 | 47651102 | + | 688 | 4224 | Rn4.5s | bXPF |
| 1-903 | chr14 | 49493742 | 49494205 | + | 688 | 31864 | Slc35f4 | bXPF |
| 1-819 | chr2 | 145214026 | 145214489 | + | 687 | -28354 | Slc24a3 | bXPF |
| 1-360 | chr3 | 95650745 | 95651208 | + | 687 | -7745 | Mcl1 | bXPF |
| 1-206 | chr9 | 66988919 | 66989382 | + | 687 | -13666 | Lactb | bXPF |
| 1-565 | chr9 | 66988720 | 66989183 | + | 686 | -13467 | Lactb | bXPF |
| 1-463 | chr8 | 106415164 | 106415627 | + | 686 | 56 | Zfp90 | bXPF |
| 13516 | chr7 | 129591696 | 129592159 | + | 686 | 64 | Wdr11 | bXPF |
| 1-830 | chr7 | 129591494 | 129591957 | + | 685 | -138 | Wdr11 | bXPF |
| 1-917 | chr14 | 9542955 | 9543418 | + | 685 | -340909 | Mnd1-ps | bXPF |
| 1-230 | chr14 | 9543156 | 9543619 | + | 685 | -340708 | Mnd1-ps | bXPF |
| 1-509 | chr17 | 84112265 | 84112728 | + | 684 | 33836 | 4933433H22Rik | bXPF |
| 20090 | chr5 | 52834112 | 52834575 | + | 684 | 208 | Anapc4 | bXPF |
| 1-430 | chr5 | 52833912 | 52834375 | + | 684 | 8 | Anapc4 | bXPF |
| 1-703 | chr4 | 13364692 | 13365155 | + | 683 | -378379 | Runx1t1 | bXPF |
| 1-256 | chr10 | 64076232 | 64076695 | + | 683 | 13792 | Lrrtm3 | bXPF |
| 1-820 | chr2 | 129100744 | 129101207 | + | 683 | -21 | Polr1b | bXPF |

| | | | | | | | | |
|---|---|---|---|---|---|---|---|---|
| 1-777 | chr7 | 83392669 | 83393132 | + | 682 | -192031 | Tmc3 | bXPF |
| 1-373 | chr13 | 63251565 | 63252028 | + | 682 | 11767 | 2010111I01Rik | bXPF |
| 1-1063 | chr11 | 19923994 | 19924457 | + | 681 | -217 | Spred2 | bXPF |
| 1-699 | chr17 | 25792121 | 25792584 | + | 681 | 35 | Fam173a | bXPF |
| 1-969 | chr4 | 117682058 | 117682521 | + | 680 | -64 | Dmap1 | bXPF |
| 1-949 | chr17 | 84085751 | 84086214 | + | 680 | 7322 | 4933433H22Rik | bXPF |
| 1-778 | chr1 | 163899252 | 163899715 | + | 680 | -29617 | Scyl3 | bXPF |
| 1-425 | chr15 | 81159256 | 81159719 | + | 679 | 31270 | Mkl1 | bXPF |
| 45658 | chr10 | 122894781 | 122895244 | + | 679 | 90712 | Mirlet7i | bXPF |
| 1-815 | chr7 | 112499930 | 112500393 | + | 679 | 72455 | Parva | bXPF |
| 1-879 | chr7 | 110297140 | 110297603 | + | 678 | 75668 | Swap70 | bXPF |
| 1-512 | chr18 | 4518426 | 4518889 | + | 678 | -116272 | 9430020K01Rik | bXPF |
| 1-914 | chr10 | 122894577 | 122895040 | + | 678 | 90916 | Mirlet7i | bXPF |
| 1-878 | chr10 | 67287950 | 67288413 | + | 677 | -2900 | Nrbf2 | bXPF |
| 1-690 | chr6 | 117272200 | 117272663 | + | 677 | 103896 | Cxcl12 | bXPF |
| 1-902 | chr4 | 75329696 | 75330159 | + | 677 | -51641 | Tmem261 | bXPF |
| 1-208 | chr14 | 7774161 | 7774624 | + | 676 | -43565 | Flnb | bXPF |
| 1-675 | chr14 | 7773956 | 7774419 | + | 676 | -43770 | Flnb | bXPF |
| 42887 | chr10 | 67288175 | 67288638 | + | 676 | -3125 | Nrbf2 | bXPF |
| 1-972 | chr7 | 132444233 | 132444696 | + | 675 | -127309 | Chst15 | bXPF |
| 1-734 | chr17 | 83979166 | 83979629 | + | 675 | -19155 | 8430430B14Rik | bXPF |
| 22282 | chr17 | 83979365 | 83979828 | + | 675 | -18956 | 8430430B14Rik | bXPF |
| 32509 | chr7 | 132444432 | 132444895 | + | 674 | -127508 | Chst15 | bXPF |
| 1-217 | chr15 | 89195995 | 89196458 | + | 674 | 248 | Dennd6b | bXPF |
| 1-800 | chr17 | 29268649 | 29269112 | + | 674 | 92 | BC004004 | bXPF |
| 36161 | chr8 | 35591808 | 35592271 | + | 673 | -3019 | Gm16793 | bXPF |
| 1-578 | chr8 | 35591609 | 35592072 | + | 673 | -2820 | Gm16793 | bXPF |
| 1-176 | chr5 | 151368833 | 151369296 | + | 673 | 389 | 1700028E10Rik | bXPF |
| 1-368 | chr1 | 53823652 | 53824115 | + | 672 | -25541 | Mir7681 | bXPF |
| 1-477 | chr3 | 37875303 | 37875766 | + | 672 | -23279 | Gm20755 | bXPF |
| 1-1041 | chr16 | 63287636 | 63288099 | + | 672 | 433560 | Pros1 | bXPF |
| 1-149 | chr18 | 78485196 | 78485659 | + | 671 | 111523 | Slc14a2 | bXPF |
| 1-638 | chr18 | 78484949 | 78485412 | + | 671 | 111770 | Slc14a2 | bXPF |
| 1-412 | chr16 | 85173547 | 85174010 | + | 671 | -71 | App | bXPF |
| 1-845 | chr18 | 56035096 | 56035559 | + | 670 | -396805 | Gramd3 | bXPF |
| 1-863 | chr9 | 49980782 | 49981245 | + | 670 | -181620 | Ncam1 | bXPF |
| 1-1056 | chr11 | 100979911 | 100980374 | + | 670 | -9525 | Ptrf | bXPF |
| 1-418 | chr8 | 24450523 | 24450986 | + | 669 | -11808 | 1810011O10Rik | bXPF |
| 1-803 | chr1 | 79775834 | 79776297 | + | 669 | 47 | Mrpl44 | bXPF |
| 1-160 | chr1 | 79776038 | 79776501 | + | 669 | 251 | Mrpl44 | bXPF |
| 1-1010 | chr7 | 79920455 | 79920918 | + | 668 | -46 | Ap3s2 | bXPF |
| 17168 | chr14 | 51089524 | 51089987 | + | 668 | -1322 | Rnase4 | bXPF |
| 1-193 | chr3 | 21935697 | 21936160 | + | 668 | -140724 | Tbl1xr1 | bXPF |
| 1-1054 | chr9 | 64960633 | 64961096 | + | 668 | 32 | Vwa9 | bXPF |
| 1-942 | chr16 | 56852861 | 56853324 | + | 667 | 33071 | Tmem45a | bXPF |
| 29221 | chr16 | 65815158 | 65815621 | + | 667 | -244 | Vgll3 | bXPF |
| 1-658 | chr10 | 37241693 | 37242156 | + | 667 | -102998 | Marcks | bXPF |
| 1-866 | chr18 | 44472979 | 44473442 | + | 666 | 92710 | Dcp2 | bXPF |
| 1-352 | chr16 | 65814955 | 65815418 | + | 666 | -447 | Vgll3 | bXPF |
| 1-712 | chr7 | 98540796 | 98541259 | + | 666 | 46805 | Lrrc32 | bXPF |
| 1-834 | chr4 | 32179897 | 32180360 | + | 665 | 216021 | Map3k7 | bXPF |
| 1-692 | chr7 | 13034479 | 13034942 | + | 665 | 67 | Chmp2a | bXPF |
| 42736 | chr4 | 32180101 | 32180564 | + | 665 | 216225 | Map3k7 | bXPF |
| 1-702 | chr7 | 142470762 | 142471225 | + | 664 | -850 | Lsp1 | bXPF |
| 23377 | chr11 | 57962566 | 57963029 | + | 664 | -46267 | Larp1 | bXPF |
| 1-685 | chr2 | 102901130 | 102901593 | + | 664 | 304 | Cd44 | bXPF |
| 1-421 | chr11 | 57962367 | 57962830 | + | 664 | -46466 | Larp1 | bXPF |
| 1-583 | chr15 | 6874067 | 6874530 | + | 663 | 15 | Osmr | bXPF |
| 1-273 | chr13 | 23519516 | 23519979 | + | 663 | -11297 | Hist1h4h | bXPF |
| 1-618 | chr8 | 44934922 | 44935385 | + | 663 | -15055 | Fat1 | bXPF |
| 1-179 | chr14 | 70659154 | 70659617 | + | 662 | -6301 | Npm2 | bXPF |
| 1-394 | chr14 | 70658940 | 70659403 | + | 662 | -6087 | Npm2 | bXPF |
| 1-980 | chr3 | 49928887 | 49929350 | + | 662 | -171802 | Pcdh18 | bXPF |

| | | | | | | | | |
|---|---|---|---|---|---|---|---|---|
| 1-483 | chr2 | 3748617 | 3749080 | + | 661 | 35390 | Fam107b | bXPF |
| 20455 | chr5 | 125389887 | 125390350 | + | 661 | -101 | Ubc | bXPF |
| 1-787 | chr5 | 125389682 | 125390145 | + | 661 | 104 | Ubc | bXPF |
| 1-975 | chr2 | 51234372 | 51234835 | + | 661 | -7228 | Gm13497 | bXPF |
| 1-865 | chr10 | 107171898 | 107172361 | + | 660 | -48465 | Acss3 | bXPF |
| 1-713 | chr1 | 36369013 | 36369476 | + | 660 | -63 | Kansl3 | bXPF |
| 1-336 | chr19 | 8774227 | 8774690 | + | 660 | 9 | Tmem179b | bXPF |
| 24108 | chr6 | 145768931 | 145769394 | + | 659 | -39221 | Rassf8 | bXPF |
| 1-706 | chr6 | 145768726 | 145769189 | + | 659 | -39426 | Rassf8 | bXPF |
| 1-811 | chr10 | 105841148 | 105841611 | + | 659 | 1 | Mettl25 | bXPF |
| 1-848 | chr13 | 98890771 | 98891234 | + | 658 | 40 | Tnpo1 | bXPF |
| 1-680 | chr2 | 51914766 | 51915229 | + | 658 | 19647 | Rbm43 | bXPF |
| 1-839 | chr6 | 128842992 | 128843455 | + | 658 | -16908 | Klrb1b | bXPF |
| 1-582 | chr11 | 35065754 | 35066217 | + | 658 | -55471 | Slit3 | bXPF |
| 1-558 | chr13 | 15741795 | 15742258 | + | 657 | 60604 | 4933412O06Rik | bXPF |
| 1-670 | chr3 | 9839926 | 9840389 | + | 657 | -6478 | Pag1 | bXPF |
| 1-749 | chr17 | 33955652 | 33956115 | + | 657 | 1 | Vps52 | bXPF |
| 1-587 | chr6 | 100146353 | 100146816 | + | 656 | 140774 | Rybp | bXPF |
| 1-251 | chr13 | 91165262 | 91165725 | + | 656 | 58494 | Atg10 | bXPF |
| 1-886 | chr16 | 14940903 | 14941366 | + | 656 | 34488 | Efcab1 | bXPF |
| 1-789 | chr13 | 51975775 | 51976238 | + | 656 | -120735 | 4921525O09Rik | bXPF |
| 1-880 | chr19 | 42017477 | 42017940 | + | 655 | -18330 | Ankrd2 | bXPF |
| 1-480 | chr12 | 51691929 | 51692392 | + | 655 | -246 | Strn3 | bXPF |
| 1-739 | chr12 | 35246085 | 35246548 | + | 655 | 199127 | Snx13 | bXPF |
| 27760 | chr6 | 100120132 | 100120595 | + | 654 | 166995 | Rybp | bXPF |
| 1-720 | chr6 | 100119928 | 100120391 | + | 654 | 167199 | Rybp | bXPF |
| 1-659 | chr15 | 5116136 | 5116599 | + | 654 | -246 | Rpl37 | bXPF |
| 1-1053 | chr13 | 54249052 | 54249515 | + | 654 | 57154 | Hrh2 | bXPF |
| 1-639 | chr5 | 21701503 | 21701966 | + | 653 | -389 | Napepld | bXPF |
| 1-180 | chr8 | 103814177 | 103814640 | + | 653 | 210609 | 4933400L20Rik | bXPF |
| 1-825 | chr4 | 33261259 | 33261722 | + | 653 | -12703 | Pnrc1 | bXPF |
| 1-353 | chr1 | 138023445 | 138023908 | + | 652 | 57037 | Mir181b-1 | bXPF |
| 1-159 | chr1 | 138023648 | 138024111 | + | 652 | 57240 | Mir181b-1 | bXPF |
| 1-472 | chr2 | 51238666 | 51239129 | + | 652 | -2934 | Gm13497 | bXPF |
| 1-833 | chr9 | 15279813 | 15280276 | + | 652 | -177 | Med17 | bXPF |
| 1-837 | chr1 | 54438822 | 54439285 | + | 651 | -27 | Gtf3c3 | bXPF |
| 1-732 | chr13 | 21974834 | 21975297 | + | 651 | -6116 | Pom121l2 | bXPF |
| 1-403 | chr13 | 113403493 | 113403956 | + | 651 | 99120 | 4921509O07Rik | bXPF |
| 1-144 | chr7 | 109576651 | 109577114 | + | 650 | 39686 | St5 | bXPF |
| 1-858 | chr7 | 109576449 | 109576912 | + | 650 | 39888 | St5 | bXPF |
| 1-687 | chr10 | 10321005 | 10321468 | + | 650 | 151078 | Adgb | bXPF |
| 1-906 | chr18 | 78237313 | 78237776 | + | 650 | -31104 | Slc14a2 | bXPF |
| 1-887 | chr8 | 33909558 | 33910021 | + | 649 | 20074 | Rbpms | bXPF |
| 1-759 | chr1 | 12500338 | 12500801 | + | 649 | 74583 | Mir6341 | bXPF |
| 1-852 | chr12 | 73790422 | 73790885 | + | 649 | -110671 | Hif1a | bXPF |
| 1-573 | chr10 | 126144720 | 126145183 | + | 649 | 178732 | Lrig3 | bXPF |
| 1-602 | chr18 | 66140980 | 66141443 | + | 648 | 78122 | Mir694 | bXPF |
| 1-145 | chr1 | 136683310 | 136683773 | + | 648 | -88 | Gm19705 | bXPF |
| 1-599 | chr5 | 114100048 | 114100511 | + | 648 | -54 | Usp30 | bXPF |
| 1-244 | chr1 | 136683015 | 136683478 | + | 648 | -150 | Gm19705 | bXPF |
| 1-534 | chr6 | 114726825 | 114727288 | + | 647 | 83959 | Atg7 | bXPF |
| 1-973 | chr13 | 84646831 | 84647294 | + | 647 | 424766 | Tmem161b | bXPF |
| 1-277 | chr17 | 25985448 | 25985911 | + | 647 | 195 | Capn15 | bXPF |
| 1-831 | chr2 | 136955699 | 136956162 | + | 647 | 56579 | Slx4ip | bXPF |
| 1-1027 | chr16 | 90844892 | 90845355 | + | 646 | 14264 | Eva1c | bXPF |
| 1-564 | chr13 | 60417348 | 60417811 | + | 646 | -184368 | Dapk1 | bXPF |
| 1-632 | chr17 | 63660924 | 63661387 | + | 646 | -160575 | Fbxl17 | bXPF |
| 1-282 | chr4 | 150854684 | 150855147 | + | 646 | 34 | 1700045H11Rik | bXPF |
| 1-500 | chr2 | 52981067 | 52981530 | + | 645 | 123430 | Fmnl2 | bXPF |
| 1-122 | chr4 | 150854886 | 150855349 | + | 645 | 26 | Errfi1 | bXPF |
| 1-516 | chr19 | 8983240 | 8983703 | + | 645 | -5813 | Ahnak | bXPF |
| 25204 | chr7 | 70822428 | 70822891 | + | 644 | -455913 | Nr2f2 | bXPF |
| 1-684 | chr11 | 69417284 | 69417747 | + | 644 | -3840 | Kdm6b | bXPF |

| | | | | | | | | |
|---|---|---|---|---|---|---|---|---|
| 1-898 | chr9 | 120691376 | 120691839 | + | 644 | 114276 | 5830454E08Rik | bXPF |
| 1-915 | chr5 | 125273598 | 125274061 | + | 644 | 67265 | Scarb1 | bXPF |
| 1-889 | chr1 | 162015402 | 162015865 | + | 643 | 10664 | 2810442N19Rik | bXPF |
| 1-773 | chr12 | 101083468 | 101083931 | + | 643 | 3 | Smek1 | bXPF |
| 1-926 | chr8 | 64029522 | 64029985 | + | 643 | -77583 | Sgol2b | bXPF |
| 1-637 | chr10 | 39614598 | 39615061 | + | 643 | 1895 | Traf3ip2 | bXPF |
| 1-367 | chr1 | 39900265 | 39900728 | + | 643 | -417 | Map4k4 | bXPF |
| 1-473 | chr17 | 74489326 | 74489789 | + | 642 | 64 | Yipf4 | bXPF |
| 1-172 | chr17 | 74489527 | 74489990 | + | 642 | 265 | Yipf4 | bXPF |
| 1-931 | chr11 | 119933142 | 119933605 | + | 642 | -9719 | Baiap2 | bXPF |
| 1-673 | chr5 | 123141017 | 123141480 | + | 642 | -945 | Setd1b | bXPF |
| 1-821 | chr18 | 11883433 | 11883896 | + | 641 | -43226 | Mir1901 | bXPF |
| 1-841 | chr4 | 47504905 | 47505368 | + | 641 | 30475 | Sec61b | bXPF |
| 1-415 | chr12 | 116280602 | 116281065 | + | 641 | -389 | Esyt2 | bXPF |
| 1-935 | chr3 | 38283721 | 38284184 | + | 641 | -75833 | 5430434I15Rik | bXPF |
| 1-691 | chr2 | 166793473 | 166793936 | + | 640 | -63 | Trp53rkb | bXPF |
| 1-996 | chr9 | 61013099 | 61013562 | + | 640 | -3739 | Gm5122 | bXPF |
| 1-153 | chr10 | 86425881 | 86426344 | + | 640 | 72784 | Syn3 | bXPF |
| 1-853 | chr13 | 5248810 | 5249273 | + | 640 | 609051 | 1700016G22Rik | bXPF |
| 1-961 | chr11 | 86570236 | 86570699 | + | 639 | 13691 | Mir21a | bXPF |
| 1-835 | chr10 | 38908180 | 38908643 | + | 639 | -57104 | Lama4 | bXPF |
| 1-652 | chr16 | 32640038 | 32640501 | + | 639 | -4374 | Tnk2 | bXPF |
| 1-294 | chr15 | 35371260 | 35371723 | + | 639 | -55 | Vps13b | bXPF |
| 1-850 | chr9 | 63896370 | 63896833 | + | 638 | 125458 | Smad6 | bXPF |
| 1-654 | chr18 | 23989416 | 23989879 | + | 638 | 13 | Zfp35 | bXPF |
| 1-187 | chr4 | 126150473 | 126150936 | + | 638 | 2701 | Eva1b | bXPF |
| 1-870 | chr18 | 35021772 | 35022235 | + | 638 | -67652 | Hspa9 | bXPF |
| 1-142 | chr6 | 127337498 | 127337961 | + | 637 | -29762 | Gm38404 | bXPF |
| 1-617 | chr4 | 126150274 | 126150737 | + | 637 | 2502 | Eva1b | bXPF |
| 1-462 | chr6 | 127337287 | 127337750 | + | 637 | -29551 | Gm38404 | bXPF |
| 19725 | chr9 | 43945629 | 43946092 | + | 637 | -97524 | Thy1 | bXPF |
| 1-930 | chr1 | 171111218 | 171111681 | + | 636 | -10212 | Cfap126 | bXPF |
| 1-408 | chr5 | 93114418 | 93114881 | + | 636 | 21233 | Sept11 | bXPF |
| 25569 | chr5 | 93114617 | 93115080 | + | 636 | 21432 | Sept11 | bXPF |
| 1-308 | chr12 | 8913600 | 8914063 | + | 636 | -7476 | Laptm4a | bXPF |
| 1-661 | chr3 | 30508190 | 30508653 | + | 636 | 1066 | Mecom | bXPF |
| 1-781 | chr17 | 25874784 | 25875247 | + | 635 | -485 | Mettl26 | bXPF |
| 1-407 | chr12 | 83921939 | 83922402 | + | 635 | -236 | Numb | bXPF |
| 1-605 | chr11 | 97309373 | 97309836 | + | 635 | -6112 | Mrpl45 | bXPF |
| 1-810 | chr3 | 88177778 | 88178241 | + | 635 | -224 | 1700113A16Rik | bXPF |
| 1-138 | chr11 | 46073126 | 46073589 | + | 634 | 17373 | Adam19 | bXPF |
| 1-686 | chr18 | 5166535 | 5166998 | + | 634 | 120177 | Svil | bXPF |
| 1-429 | chr11 | 46072927 | 46073390 | + | 634 | 17174 | Adam19 | bXPF |
| 1-591 | chr17 | 17827064 | 17827527 | + | 634 | -2893 | Mir99b | bXPF |
| 1-725 | chr10 | 23787005 | 23787468 | + | 633 | -27 | Rps12 | bXPF |
| 1-601 | chr4 | 40853864 | 40854327 | + | 633 | 442 | B4galt1 | bXPF |
| 1-760 | chr8 | 83652258 | 83652721 | + | 633 | -189 | Gipc1 | bXPF |
| 1-132 | chr8 | 83652462 | 83652925 | + | 633 | 15 | Gipc1 | bXPF |
| 1-813 | chr8 | 92706390 | 92706853 | + | 633 | 32332 | Irx6 | bXPF |
| 1-682 | chr3 | 25145186 | 25145649 | + | 632 | -361411 | Naaladl2 | bXPF |
| 1-620 | chr7 | 98815254 | 98815717 | + | 632 | -19627 | Wnt11 | bXPF |
| 1-829 | chr7 | 96575341 | 96575804 | + | 632 | 225977 | Gm15413 | bXPF |
| 1-771 | chr8 | 23256917 | 23257380 | + | 632 | -68 | Golga7 | bXPF |
| 1-694 | chr3 | 54360785 | 54361248 | + | 631 | -91 | Postn | bXPF |
| 1-379 | chr5 | 138170940 | 138171403 | + | 631 | 691 | Mcm7 | bXPF |
| 1-485 | chr4 | 15957913 | 15958376 | + | 631 | 177 | Nbn | bXPF |
| 26665 | chr3 | 54361000 | 54361463 | + | 631 | 124 | Postn | bXPF |
| 1-434 | chr6 | 29388577 | 29389040 | + | 631 | -7767 | Ccdc136 | bXPF |
| 1-169 | chr7 | 58830813 | 58831276 | + | 630 | 172842 | Atp10a | bXPF |
| 1-554 | chr6 | 47781395 | 47781858 | + | 630 | 22026 | Mir704 | bXPF |
| 1-537 | chr1 | 62703127 | 62703590 | + | 630 | 41 | Nrp2 | bXPF |
| 1-802 | chr16 | 11909154 | 11909617 | + | 630 | 38 | Cpped1 | bXPF |
| 1-871 | chr5 | 107101821 | 107102284 | + | 629 | 137491 | Cdc7 | bXPF |

| | | | | | | | | |
|---|---|---|---|---|---|---|---|---|
| 1-498 | chr17 | 42580570 | 42581033 | + | 629 | 30512 | Opn5 | bXPF |
| 1-867 | chr6 | 91729864 | 91730327 | + | 629 | 46028 | Slc6a6 | bXPF |
| 1-304 | chr4 | 7560170 | 7560633 | + | 629 | -287 | 8430436N08Rik | bXPF |
| 1-842 | chr6 | 30549223 | 30549686 | + | 629 | 7812 | Cpa2 | bXPF |
| 1-611 | chr2 | 4286374 | 4286837 | + | 628 | -114371 | Frmd4a | bXPF |
| 1-933 | chr17 | 83861903 | 83862366 | + | 628 | -15344 | Haao | bXPF |
| 1-324 | chr1 | 93478781 | 93479244 | + | 628 | 19 | Sept2 | bXPF |
| 1-157 | chr7 | 84329313 | 84329776 | + | 628 | 80494 | Arnt2 | bXPF |
| 1-896 | chr8 | 108552529 | 108552992 | + | 628 | 31917 | Lncbate1 | bXPF |
| 1-827 | chr16 | 90533066 | 90533529 | + | 627 | 146900 | Hunk | bXPF |
| 13881 | chr16 | 90533285 | 90533748 | + | 627 | 147119 | Hunk | bXPF |
| 1-354 | chr11 | 118355127 | 118355590 | + | 627 | 53 | Timp2 | bXPF |
| 1-156 | chr11 | 118355352 | 118355815 | + | 627 | -172 | Timp2 | bXPF |
| 1-752 | chr17 | 51979067 | 51979530 | + | 626 | 96571 | Gm20098 | bXPF |
| 1-562 | chr5 | 4104322 | 4104785 | + | 626 | 144 | Cyp51 | bXPF |
| 35065 | chr5 | 134014413 | 134014876 | + | 626 | -85104 | Gatsl2 | bXPF |
| 1-233 | chr17 | 42580258 | 42580721 | + | 626 | 30824 | Opn5 | bXPF |
| 1-785 | chr16 | 84735065 | 84735528 | + | 626 | 6 | Mrpl39 | bXPF |
| 1-300 | chr5 | 25529447 | 25529910 | + | 625 | -340 | 1700096K18Rik | bXPF |
| 1-723 | chr1 | 165641433 | 165641896 | + | 625 | -7123 | Mpzl1 | bXPF |
| 1-588 | chr15 | 76327169 | 76327632 | + | 625 | 3 | Exosc4 | bXPF |
| 1-103 | chr8 | 78742012 | 78742475 | + | 625 | 78909 | Lsm6 | bXPF |
| 1-763 | chr1 | 90750099 | 90750562 | + | 625 | 93641 | Col6a3 | bXPF |
| 1-762 | chr9 | 117246172 | 117246635 | + | 624 | 6080 | Rbms3 | bXPF |
| 1-158 | chr9 | 53384203 | 53384666 | + | 624 | 411 | Kdelc2 | bXPF |
| 1-561 | chr3 | 95160076 | 95160539 | + | 624 | -113 | Sema6c | bXPF |
| 1-152 | chr5 | 111620136 | 111620599 | + | 624 | 38806 | C130026L21Rik | bXPF |
| 1-718 | chr5 | 111619907 | 111620370 | + | 624 | 38577 | C130026L21Rik | bXPF |
| 1-884 | chr6 | 116974827 | 116975290 | + | 623 | -193477 | Cxcl12 | bXPF |
| 1-764 | chr2 | 54281876 | 54282339 | + | 623 | -154280 | Galnt13 | bXPF |
| 1-633 | chr16 | 97962262 | 97962725 | + | 623 | 129 | Zbtb21 | bXPF |
| 16438 | chr2 | 54282076 | 54282539 | + | 623 | -154080 | Galnt13 | bXPF |
| 1-860 | chr4 | 63291050 | 63291513 | + | 623 | 34430 | Mir455 | bXPF |
| 1-471 | chr9 | 63123956 | 63124419 | + | 622 | 22793 | Skor1 | bXPF |
| 1-844 | chr11 | 50439807 | 50440270 | + | 622 | -8927 | Rufy1 | bXPF |
| 1-586 | chr9 | 44482435 | 44482898 | + | 622 | -16470 | Bcl9l | bXPF |
| 12420 | chr9 | 44482638 | 44483101 | + | 622 | -16267 | Bcl9l | bXPF |
| 1-746 | chr19 | 55316023 | 55316486 | + | 622 | 197 | Vti1a | bXPF |
| 1-326 | chr3 | 137552946 | 137553409 | + | 621 | -555 | Gm4861 | bXPF |
| 1-147 | chr9 | 41461807 | 41462270 | + | 621 | -12855 | 3110039I08Rik | bXPF |
| 1-454 | chr5 | 36226325 | 36226788 | + | 621 | 22535 | Psapl1 | bXPF |
| 1-904 | chr10 | 122912968 | 122913431 | + | 621 | 72525 | Mirlet7i | bXPF |
| 1-662 | chr11 | 121451708 | 121452171 | + | 621 | -10 | Tbcd | bXPF |
| 46023 | chr4 | 9668552 | 9669015 | + | 620 | 379 | Asph | bXPF |
| 1-484 | chr9 | 41461607 | 41462070 | + | 620 | -13055 | 3110039I08Rik | bXPF |
| 1-278 | chr8 | 104023229 | 104023692 | + | 620 | -78165 | Cdh5 | bXPF |
| 1-475 | chr9 | 120776644 | 120777107 | + | 620 | 138375 | 1700020M21Rik | bXPF |
| 1-770 | chr7 | 70822205 | 70822668 | + | 619 | -455690 | Nr2f2 | bXPF |
| 1-468 | chr1 | 90334898 | 90335361 | + | 619 | 131149 | Ackr3 | bXPF |
| 30317 | chr2 | 18767146 | 18767609 | + | 619 | -6515 | Gm3363 | bXPF |
| 11689 | chr1 | 90335099 | 90335562 | + | 619 | 131350 | Ackr3 | bXPF |
| 14977 | chr13 | 99391679 | 99392142 | + | 619 | 20916 | 6430562O15Rik | bXPF |
| 1-847 | chr4 | 44072472 | 44072935 | + | 618 | -30 | Gne | bXPF |
| 1-766 | chr13 | 99391477 | 99391940 | + | 618 | 21118 | 6430562O15Rik | bXPF |
| 1-769 | chr2 | 18766942 | 18767405 | + | 618 | -6311 | Gm3363 | bXPF |
| 1-947 | chr5 | 64878034 | 64878497 | + | 618 | 54434 | Tlr1 | bXPF |
| 1-506 | chr14 | 51985789 | 51986252 | + | 618 | 1187 | Arhgef40 | bXPF |
| 1-994 | chr7 | 80786734 | 80787197 | + | 617 | 16366 | Iqgap1 | bXPF |
| 1-432 | chr12 | 33193337 | 33193800 | + | 617 | 45882 | Atxn7l1 | bXPF |
| 1-238 | chr18 | 55196209 | 55196672 | + | 617 | -206260 | Zfp608 | bXPF |
| 1-431 | chr9 | 102626332 | 102626795 | + | 617 | 267 | Anapc13 | bXPF |
| 1-710 | chr1 | 74587981 | 74588444 | + | 617 | -77 | Bcs1l | bXPF |
| 1-541 | chr10 | 59539470 | 59539933 | + | 616 | 76991 | Mcu | bXPF |

| | | | | | | | | |
|---|---|---|---|---|---|---|---|---|
| 1-167 | chr14 | 21984761 | 21985224 | + | 616 | 4609 | Zfp503 | bXPF |
| 1-470 | chr17 | 80801418 | 80801881 | + | 616 | -73624 | Map4k3 | bXPF |
| 1-385 | chr4 | 116685353 | 116685816 | + | 616 | -15 | Prdx1 | bXPF |
| 1-528 | chr1 | 155233928 | 155234391 | + | 616 | 75456 | Stx6 | bXPF |
| 1-450 | chr11 | 58938712 | 58939175 | + | 615 | -37 | Rnf187 | bXPF |
| 1-869 | chr18 | 55296368 | 55296831 | + | 615 | -306419 | Zfp608 | bXPF |
| 1-1042 | chr3 | 147445868 | 147446331 | + | 615 | 593732 | Ttll7 | bXPF |
| 1-411 | chr5 | 21586626 | 21587089 | + | 615 | 43330 | Lrrc17 | bXPF |
| 1-465 | chr17 | 46277538 | 46278001 | + | 615 | 5257 | Tjap1 | bXPF |
| 1-854 | chr13 | 21215267 | 21215730 | + | 614 | 35567 | Trim27 | bXPF |
| 1-881 | chr3 | 121576887 | 121577350 | + | 614 | -44774 | Slc44a3 | bXPF |
| 1-855 | chr18 | 34751503 | 34751966 | + | 614 | -201 | Cdc25c | bXPF |
| 1-499 | chr13 | 42549008 | 42549471 | + | 614 | -131560 | Phactr1 | bXPF |
| 1-183 | chr3 | 148641733 | 148642196 | + | 614 | 312671 | Adgrl2 | bXPF |
| 1-346 | chr16 | 56853061 | 56853524 | + | 614 | 32871 | Tmem45a | bXPF |
| 1-437 | chr6 | 47835358 | 47835821 | + | 613 | -72 | Zfp398 | bXPF |
| 1-705 | chr11 | 11953308 | 11953771 | + | 613 | -55395 | Ddc | bXPF |
| 1-939 | chr12 | 69501774 | 69502237 | + | 613 | -26455 | 5830428M24Rik | bXPF |
| 1-600 | chr8 | 128685360 | 128685823 | + | 613 | -63 | Itgb1 | bXPF |
| 43101 | chr6 | 47835559 | 47836022 | + | 613 | 129 | Zfp398 | bXPF |
| 1-401 | chr11 | 120990945 | 120991408 | + | 612 | 157 | Csnk1d | bXPF |
| 1-501 | chr3 | 51553507 | 51553970 | + | 612 | -6019 | 5031434O11Rik | bXPF |
| 43040 | chr7 | 132575574 | 132576037 | + | 612 | 593 | Oat | bXPF |
| 1-922 | chr6 | 88924627 | 88925090 | + | 612 | 22607 | Tpra1 | bXPF |
| 44562 | chr15 | 99369641 | 99370104 | + | 612 | 610 | Fmnl3 | bXPF |
| 1-992 | chr12 | 100778906 | 100779369 | + | 611 | 14 | 9030617O03Rik | bXPF |
| 1-262 | chr5 | 114372629 | 114373092 | + | 611 | 7645 | Kctd10 | bXPF |
| 1-207 | chr2 | 144165523 | 144165986 | + | 611 | 23536 | Gm5535 | bXPF |
| 1-538 | chr2 | 144165320 | 144165783 | + | 611 | 23739 | Gm5535 | bXPF |
| 13150 | chr11 | 82045705 | 82046168 | + | 611 | 224 | Ccl7 | bXPF |
| 1-535 | chr11 | 82045502 | 82045965 | + | 611 | 21 | Ccl7 | bXPF |
| 1-443 | chr12 | 4233741 | 4234204 | + | 610 | -55 | Ptrhd1 | bXPF |
| 1-683 | chr3 | 18215471 | 18215934 | + | 610 | 27636 | Cyp7b1 | bXPF |
| 1-325 | chr11 | 45296757 | 45297220 | + | 610 | -180060 | Gm12159 | bXPF |
| 1-310 | chr13 | 40859491 | 40859954 | + | 610 | -46 | Gcnt2 | bXPF |
| 1-1018 | chr2 | 128967000 | 128967463 | + | 610 | -171 | Zc3h6 | bXPF |
| 1-651 | chr9 | 109953415 | 109953878 | + | 609 | 21872 | Map4 | bXPF |
| 1-968 | chr13 | 45931236 | 45931699 | + | 609 | 33524 | Atxn1 | bXPF |
| 1-761 | chr18 | 54663802 | 54664265 | + | 609 | 52289 | 9330117O12Rik | bXPF |
| 1-772 | chr2 | 145329755 | 145330218 | + | 609 | 87375 | Slc24a3 | bXPF |
| 1-536 | chr4 | 66025390 | 66025853 | + | 609 | 378862 | Astn2 | bXPF |
| 1-695 | chr3 | 63964532 | 63964995 | + | 608 | -30 | Slc33a1 | bXPF |
| 1-843 | chr17 | 80765173 | 80765636 | + | 608 | -37379 | Map4k3 | bXPF |
| 1-458 | chr1 | 163907355 | 163907818 | + | 608 | -21514 | Scyl3 | bXPF |
| 1-389 | chr2 | 28582845 | 28583308 | + | 608 | 203 | Gtf3c5 | bXPF |
| 1-129 | chr13 | 54750898 | 54751361 | + | 608 | -1255 | Gprin1 | bXPF |
| 1-521 | chr6 | 52240293 | 52240756 | + | 608 | 330 | Hoxa10 | bXPF |
| 1-351 | chr16 | 49332341 | 49332804 | + | 607 | 275086 | Myh15 | bXPF |
| 29587 | chr11 | 69063557 | 69064020 | + | 607 | 4013 | Borcs6 | bXPF |
| 1-974 | chr9 | 41877294 | 41877757 | + | 607 | 246764 | Sorl1 | bXPF |
| 1-396 | chr6 | 114921649 | 114922112 | + | 607 | -128 | Vgll4 | bXPF |
| 1-386 | chr4 | 133729479 | 133729942 | + | 607 | -12602 | Mir7227 | bXPF |
| 1-402 | chr1 | 165460878 | 165461341 | + | 606 | -99 | Mpc2 | bXPF |
| 1-625 | chr17 | 24527855 | 24528318 | + | 606 | -148 | Traf7 | bXPF |
| 1-184 | chr2 | 51238879 | 51239342 | + | 606 | -2721 | Gm13497 | bXPF |
| 1-417 | chr13 | 37312522 | 37312985 | + | 606 | -32592 | Ly86 | bXPF |
| 1-224 | chr1 | 152386525 | 152386988 | + | 606 | -74 | Tsen15 | bXPF |
| 33239 | chr9 | 41877497 | 41877960 | + | 606 | 246561 | Sorl1 | bXPF |
| 1-628 | chr3 | 101604575 | 101605038 | + | 605 | -99 | Atp1a1 | bXPF |
| 1-305 | chr7 | 35234638 | 35235101 | + | 605 | -19524 | Lrp3 | bXPF |
| 25934 | chr10 | 93177712 | 93178175 | + | 605 | 15158 | Mir1931 | bXPF |
| 16803 | chr3 | 37938441 | 37938904 | + | 605 | 39859 | Gm20755 | bXPF |
| 1-799 | chr6 | 117327377 | 117327840 | + | 605 | 159073 | Cxcl12 | bXPF |

| | | | | | | | | |
|---|---|---|---|---|---|---|---|---|
| 1-520 | chr7 | 35234437 | 35234900 | + | 605 | -19323 | Lrp3 | bXPF |
| 1-796 | chr5 | 130623226 | 130623689 | + | 604 | 174665 | Caln1 | bXPF |
| 1-341 | chr1 | 64617088 | 64617551 | + | 604 | -151 | Mettl21a | bXPF |
| 1-559 | chr10 | 93177500 | 93177963 | + | 604 | 14946 | Mir1931 | bXPF |
| 43466 | chr6 | 117327609 | 117328072 | + | 604 | 159305 | Cxcl12 | bXPF |
| 1-358 | chr11 | 79704530 | 79704993 | + | 604 | -7208 | Mir193a | bXPF |
| 1-649 | chr16 | 17809679 | 17810142 | + | 604 | 12628 | Scarf2 | bXPF |
| 1-745 | chr10 | 13782627 | 13783090 | + | 603 | 86099 | Aig1 | bXPF |
| 1-494 | chr19 | 42017679 | 42018142 | + | 603 | -18128 | Ankrd2 | bXPF |
| 1-397 | chr16 | 4963998 | 4964461 | + | 603 | 8 | Anks3 | bXPF |
| 1-724 | chr4 | 106678705 | 106679168 | + | 603 | 8 | Ttc4 | bXPF |
| 41640 | chr16 | 4880260 | 4880723 | + | 603 | -640 | Ubald1 | bXPF |
| 1-328 | chr10 | 94990704 | 94991167 | + | 602 | -46357 | Plxnc1 | bXPF |
| 1-892 | chr10 | 71012196 | 71012659 | + | 602 | 147263 | Bicc1 | bXPF |
| 45292 | chr17 | 25273844 | 25274307 | + | 602 | -49 | Gm17801 | bXPF |
| 1-753 | chr1 | 162217457 | 162217920 | + | 602 | 65 | Dnm3os | bXPF |
| 1-744 | chr3 | 59354163 | 59354626 | + | 602 | -10138 | Igsf10 | bXPF |
| 1-751 | chr3 | 81367794 | 81368257 | + | 602 | 331609 | Pdgfc | bXPF |
| 1-948 | chr17 | 25273644 | 25274107 | + | 601 | 39 | Ube2i | bXPF |
| 1-780 | chr6 | 137465243 | 137465706 | + | 601 | 54421 | Eps8 | bXPF |
| 1-647 | chr5 | 142380453 | 142380916 | + | 601 | -20813 | Foxk1 | bXPF |
| 11324 | chr5 | 142380653 | 142381116 | + | 601 | -20613 | Foxk1 | bXPF |
| 1-311 | chr9 | 84032703 | 84033166 | + | 601 | 84153 | Bckdhb | bXPF |
| 1-151 | chr5 | 107200461 | 107200924 | + | 601 | 88903 | Tgfbr3 | bXPF |
| 1-1043 | chr5 | 134313446 | 134313909 | + | 600 | 1083 | Gtf2i | bXPF |
| 1-548 | chr5 | 140024246 | 140024709 | + | 600 | 116534 | Elfn1 | bXPF |
| 1-551 | chr19 | 21794562 | 21795025 | + | 600 | 16453 | Tmem2 | bXPF |
| 1-289 | chr3 | 133091671 | 133092134 | + | 600 | 47 | Ints12 | bXPF |
| 1-113 | chr3 | 133091875 | 133092338 | + | 600 | 251 | Ints12 | bXPF |
| 1-461 | chr13 | 63245605 | 63246068 | + | 600 | 5807 | 2010111I01Rik | bXPF |
| 1-575 | chr16 | 30827984 | 30828447 | + | 599 | 228492 | Fam43a | bXPF |
| 1-545 | chr11 | 76311517 | 76311980 | + | 599 | 68012 | Rnmtl1 | bXPF |
| 1-894 | chr12 | 112236981 | 112237444 | + | 599 | 91004 | Kif26a | bXPF |
| 1-946 | chr13 | 92967350 | 92967813 | + | 599 | -105723 | Gm4814 | bXPF |
| 41275 | chr11 | 76311719 | 76312182 | + | 599 | 68214 | Rnmtl1 | bXPF |
| 29952 | chr7 | 95914899 | 95915362 | + | 599 | -256114 | Tenm4 | bXPF |
| 1-909 | chr9 | 49798788 | 49799251 | + | 598 | 50 | Ncam1 | bXPF |
| 1-496 | chr14 | 33797863 | 33798326 | + | 598 | 13189 | 4930503F20Rik | bXPF |
| 1-275 | chr15 | 33362971 | 33363434 | + | 598 | 42737 | 1700084J12Rik | bXPF |
| 1-666 | chr15 | 33362766 | 33363229 | + | 598 | 42942 | 1700084J12Rik | bXPF |
| 1-105 | chr1 | 20069841 | 20070304 | + | 598 | -286382 | 4930486I03Rik | bXPF |
| 32143 | chr14 | 33798080 | 33798543 | + | 598 | 12972 | 4930503F20Rik | bXPF |
| 1-791 | chr2 | 178454659 | 178455122 | + | 597 | 24375 | Cdh26 | bXPF |
| 1-337 | chr7 | 28919683 | 28920146 | + | 597 | 38258 | Capn12 | bXPF |
| 1-478 | chr1 | 121567655 | 121568118 | + | 597 | 94 | Ddx18 | bXPF |
| 1-359 | chr18 | 78294032 | 78294495 | + | 597 | -87823 | Slc14a2 | bXPF |
| 1-118 | chr3 | 27863393 | 27863856 | + | 597 | 32744 | Tmem212 | bXPF |
| 1-674 | chr5 | 93206466 | 93206929 | + | 597 | 179 | 2010109A12Rik | bXPF |
| 1-406 | chr18 | 58466527 | 58466990 | + | 596 | -89482 | Slc27a6 | bXPF |
| 1-447 | chr3 | 27863193 | 27863656 | + | 596 | 32944 | Tmem212 | bXPF |
| 1-944 | chr2 | 24453147 | 24453610 | + | 596 | 22221 | Pax8 | bXPF |
| 1-957 | chr10 | 68354713 | 68355176 | + | 596 | -33802 | 4930545H06Rik | bXPF |
| 1-524 | chr5 | 148994869 | 148995332 | + | 596 | 114 | 5930430L01Rik | bXPF |
| 1-716 | chr2 | 18586864 | 18587327 | + | 596 | -85367 | Commd3 | bXPF |
| 1-627 | chr16 | 13109680 | 13110143 | + | 596 | 175 | Ercc4 | bXPF |
| 42005 | chr15 | 51877454 | 51877917 | + | 595 | 244 | Utp23 | bXPF |
| 1-448 | chr11 | 75147045 | 75147508 | + | 595 | 20876 | Hic1 | bXPF |
| 1-808 | chr5 | 141303738 | 141304201 | + | 595 | 62435 | Sdk1 | bXPF |
| 1-297 | chr8 | 109693066 | 109693529 | + | 595 | -3 | Ist1 | bXPF |
| 1-279 | chr15 | 51877249 | 51877712 | + | 595 | 39 | Utp23 | bXPF |
| 1-1020 | chr11 | 33660416 | 33660879 | + | 595 | -81690 | Gabrp | bXPF |
| 1-255 | chr1 | 106400831 | 106401294 | + | 594 | 145569 | Mir3473f | bXPF |
| 34700 | chr3 | 37875546 | 37876009 | + | 594 | -23036 | Gm20755 | bXPF |

| | | | | | | | | |
|---|---|---|---|---|---|---|---|---|
| 1-689 | chr14 | 33247554 | 33248017 | + | 594 | 30962 | Arhgap22 | bXPF |
| 1-211 | chr13 | 104817163 | 104817626 | + | 594 | 224 | Srek1ip1 | bXPF |
| 1-768 | chr10 | 128458329 | 128458792 | + | 594 | -120 | Mir8105 | bXPF |
| 1-613 | chr6 | 118419174 | 118419637 | + | 594 | 12 | Bms1 | bXPF |
| 1-823 | chr2 | 52455627 | 52456090 | + | 593 | -30984 | Arl5a | bXPF |
| 1-526 | chr5 | 111890893 | 111891356 | + | 593 | -129396 | E130006D01Rik | bXPF |
| 1-1004 | chr11 | 120932685 | 120933148 | + | 593 | -44 | Ccdc57 | bXPF |
| 15707 | chr15 | 58135082 | 58135545 | + | 593 | -231 | Atad2 | bXPF |
| 1-348 | chr15 | 58134866 | 58135329 | + | 593 | -15 | Atad2 | bXPF |
| 1-316 | chr11 | 84513275 | 84513738 | + | 593 | -5 | Aatf | bXPF |
| 1-398 | chr1 | 181446386 | 181446849 | + | 592 | 64834 | Ccdc121 | bXPF |
| 1-237 | chr8 | 111033858 | 111034321 | + | 592 | 247 | Aars | bXPF |
| 1-222 | chr8 | 79208590 | 79209053 | + | 592 | 39762 | 1700011L22Rik | bXPF |
| 1-182 | chr9 | 106429512 | 106429975 | + | 592 | 332 | Rpl29 | bXPF |
| 1-882 | chr14 | 29498570 | 29499033 | + | 592 | 223063 | Cacna2d3 | bXPF |
| 1-1065 | chr16 | 95844707 | 95845170 | + | 592 | 53775 | 1600002D24Rik | bXPF |
| 1-518 | chr19 | 28591774 | 28592237 | + | 592 | -86219 | D930032P07Rik | bXPF |
| 1-650 | chr8 | 47917539 | 47918002 | + | 591 | 72781 | Wwc2 | bXPF |
| 1-239 | chr6 | 51076751 | 51077214 | + | 591 | 192928 | Mir148a | bXPF |
| 1-410 | chr3 | 96629736 | 96630199 | + | 591 | 39 | Rbm8a | bXPF |
| 1-363 | chr15 | 8806395 | 8806858 | + | 591 | -95819 | Slc1a3 | bXPF |
| 1-817 | chr7 | 25788488 | 25788951 | + | 591 | 14 | Axl | bXPF |
| 1-409 | chr5 | 122303956 | 122304419 | + | 591 | 19789 | Pptc7 | bXPF |
| 1-220 | chr6 | 15321352 | 15321815 | + | 590 | 136077 | Foxp2 | bXPF |
| 1-264 | chr1 | 127204829 | 127205292 | + | 590 | 74 | Mgat5 | bXPF |
| 1-504 | chr5 | 97953093 | 97953556 | + | 590 | 77638 | Antxr2 | bXPF |
| 1-828 | chr9 | 9124201 | 9124664 | + | 590 | -111856 | Gm16833 | bXPF |
| 1-918 | chr8 | 70504227 | 70504690 | + | 590 | 2281 | 2810428I15Rik | bXPF |
| 1-285 | chr1 | 90603180 | 90603643 | + | 590 | -14 | Cops8 | bXPF |
| 1-851 | chr5 | 143656800 | 143657263 | + | 590 | 5790 | Cyth3 | bXPF |
| 1-170 | chr17 | 5189881 | 5190344 | + | 589 | 195038 | Arid1b | bXPF |
| 1-191 | chr6 | 15321604 | 15322067 | + | 589 | 136329 | Foxp2 | bXPF |
| 1-782 | chr2 | 16451252 | 16451715 | + | 589 | 95179 | Plxdc2 | bXPF |
| 1-722 | chr11 | 29373412 | 29373875 | + | 589 | -529 | Ccdc88a | bXPF |
| 28491 | chr17 | 5190110 | 5190573 | + | 589 | 195267 | Arid1b | bXPF |
| 1-619 | chr5 | 107087077 | 107087540 | + | 589 | 122747 | Cdc7 | bXPF |
| 1-590 | chr9 | 7453189 | 7453652 | + | 588 | 7598 | Mmp3 | bXPF |
| 1-214 | chr1 | 162186926 | 162187389 | + | 588 | -30466 | Dnm3os | bXPF |
| 1-361 | chr10 | 25271676 | 25272139 | + | 588 | 27256 | Akap7 | bXPF |
| 1-814 | chr7 | 112663284 | 112663747 | + | 588 | -15805 | Tead1 | bXPF |
| 1-676 | chr2 | 44045399 | 44045862 | + | 588 | 19694 | Arhgap15os | bXPF |
| 42856 | chr2 | 16451452 | 16451915 | + | 588 | 95379 | Plxdc2 | bXPF |
| 1-517 | chr2 | 73572415 | 73572878 | + | 588 | 7692 | Chrna1 | bXPF |
| 1-119 | chr9 | 7453394 | 7453857 | + | 587 | 7803 | Mmp3 | bXPF |
| 34335 | chr2 | 51144553 | 51145016 | + | 587 | 4327 | Rnd3 | bXPF |
| 1-405 | chr11 | 89303311 | 89303774 | + | 587 | -983 | Nog | bXPF |
| 1-136 | chr18 | 75366756 | 75367219 | + | 587 | -378 | Smad7 | bXPF |
| 1-218 | chr5 | 111891093 | 111891556 | + | 587 | -129596 | E130006D01Rik | bXPF |
| 1-663 | chr13 | 21275770 | 21276233 | + | 587 | 16685 | Gpx5 | bXPF |
| 1-941 | chr6 | 124700294 | 124700757 | + | 586 | 11653 | Emg1 | bXPF |
| 1-531 | chr1 | 84302732 | 84303195 | + | 586 | -18318 | Pid1 | bXPF |
| 1-436 | chr5 | 21056127 | 21056590 | + | 586 | -561 | Ptpn12 | bXPF |
| 1-646 | chr13 | 35604050 | 35604513 | + | 586 | -55581 | Cdyl | bXPF |
| 1-197 | chr13 | 5257995 | 5258458 | + | 586 | 599866 | 1700016G22Rik | bXPF |
| 1-369 | chr11 | 22982024 | 22982487 | + | 586 | 29 | Commd1 | bXPF |
| 1-426 | chr16 | 21694434 | 21694897 | + | 586 | 0 | 2510009E07Rik | bXPF |
| 1-243 | chr11 | 93373738 | 93374201 | + | 585 | 274679 | Car10 | bXPF |
| 1-126 | chr3 | 21936019 | 21936482 | + | 585 | -140402 | Tbl1xr1 | bXPF |
| 1-330 | chr3 | 63137753 | 63138216 | + | 585 | -157451 | Mme | bXPF |
| 1-765 | chr19 | 9978508 | 9978971 | + | 585 | -1861 | Fth1 | bXPF |
| 42826 | chr11 | 93373940 | 93374403 | + | 585 | 274881 | Car10 | bXPF |
| 1-643 | chr7 | 127477646 | 127478109 | + | 585 | 6263 | Prr14 | bXPF |
| 1-331 | chr11 | 76312644 | 76313107 | + | 584 | 69139 | Rnmtl1 | bXPF |

| | | | | | | | | |
|---|---|---|---|---|---|---|---|---|
| 1-299 | chr11 | 11939481 | 11939944 | + | 584 | -41568 | Ddc | bXPF |
| 1-731 | chr3 | 66040942 | 66041405 | + | 584 | -82948 | Ccnl1 | bXPF |
| 1-542 | chr4 | 148444518 | 148444981 | + | 584 | 2 | Ubiad1 | bXPF |
| 1-424 | chr7 | 28392724 | 28393187 | + | 584 | -41 | Paf1 | bXPF |
| 1-329 | chr11 | 97699928 | 97700391 | + | 584 | 338 | Pcgf2 | bXPF |
| 46753 | chr11 | 11939692 | 11940155 | + | 584 | -41779 | Ddc | bXPF |
| 1-303 | chr17 | 29415936 | 29416399 | + | 583 | 55253 | Fgd2 | bXPF |
| 1-657 | chr12 | 4916940 | 4917403 | + | 583 | -182 | Atad2b | bXPF |
| 1-181 | chr17 | 29416173 | 29416636 | + | 583 | 55490 | Fgd2 | bXPF |
| 1-481 | chr1 | 51401983 | 51402446 | + | 583 | -68435 | 9330175M20Rik | bXPF |
| 1-826 | chr8 | 84722806 | 84723269 | + | 583 | 21580 | Lyl1 | bXPF |
| 1-161 | chr8 | 70698597 | 70699060 | + | 583 | 1089 | Jund | bXPF |
| 1-125 | chr8 | 70698854 | 70699317 | + | 583 | 1346 | Jund | bXPF |
| 1-547 | chr14 | 49171924 | 49172387 | + | 582 | -91 | Naa30 | bXPF |
| 1-568 | chr2 | 37430771 | 37431234 | + | 582 | -83 | Zbtb6 | bXPF |
| 1-737 | chr16 | 13160187 | 13160650 | + | 582 | 50682 | Ercc4 | bXPF |
| 1-270 | chr16 | 37788319 | 37788782 | + | 582 | 11495 | Fstl1 | bXPF |
| 1-131 | chr3 | 106721565 | 106722028 | + | 582 | 114 | Lrif1 | bXPF |
| 24838 | chr3 | 106721283 | 106721746 | + | 582 | -168 | Lrif1 | bXPF |
| 1-495 | chr17 | 33685020 | 33685483 | + | 582 | 207 | Hnrnpm | bXPF |
| 1-343 | chr11 | 55468838 | 55469301 | + | 581 | -683 | G3bp1 | bXPF |
| 19360 | chr10 | 68276690 | 68277153 | + | 581 | 1805 | Arid5b | bXPF |
| 1-399 | chr10 | 59702395 | 59702858 | + | 581 | 63 | Micu1 | bXPF |
| 42917 | chr4 | 76057089 | 76057552 | + | 581 | -779034 | Tmem261 | bXPF |
| 1-416 | chr4 | 76056874 | 76057337 | + | 581 | -778819 | Tmem261 | bXPF |
| 1-626 | chr6 | 13677769 | 13678232 | + | 581 | -34 | Bmt2 | bXPF |
| 1-444 | chr10 | 68276470 | 68276933 | + | 581 | 2025 | Arid5b | bXPF |
| 1-428 | chr3 | 155093124 | 155093587 | + | 580 | 23 | Fpgt | bXPF |
| 1-798 | chr12 | 9304330 | 9304793 | + | 580 | 266024 | 1700022H16Rik | bXPF |
| 1-572 | chr2 | 125290462 | 125290925 | + | 580 | 43178 | Dut | bXPF |
| 1-907 | chr1 | 163898193 | 163898656 | + | 580 | -30676 | Scyl3 | bXPF |
| 1-805 | chr5 | 57755664 | 57756127 | + | 580 | 37814 | Pcdh7 | bXPF |
| 1-741 | chr15 | 64618409 | 64618872 | + | 580 | -235721 | Asap1 | bXPF |
| 1-309 | chr12 | 79887169 | 79887632 | + | 580 | 37333 | 9430078K24Rik | bXPF |
| 1-457 | chr6 | 71440848 | 71441311 | + | 579 | -442 | Rmnd5a | bXPF |
| 1-388 | chr10 | 115315482 | 115315945 | + | 579 | -122 | Rab21 | bXPF |
| 1-202 | chr10 | 37603219 | 37603682 | + | 579 | -464524 | Marcks | bXPF |
| 1-469 | chr10 | 58810095 | 58810558 | + | 579 | -3033 | Sh3rf3 | bXPF |
| 1-130 | chr16 | 63864179 | 63864642 | + | 579 | -253 | Epha3 | bXPF |
| 1-755 | chr16 | 63863978 | 63864441 | + | 579 | -52 | Epha3 | bXPF |
| 1-553 | chr2 | 27442077 | 27442540 | + | 579 | 8322 | Mir7578 | bXPF |
| 1-940 | chr17 | 23546263 | 23546726 | + | 578 | -4305 | 6330415G19Rik | bXPF |
| 42767 | chr2 | 61354274 | 61354737 | + | 578 | -224081 | Tank | bXPF |
| 1-567 | chr16 | 91044347 | 91044810 | + | 578 | -199 | Paxbp1 | bXPF |
| 1-634 | chr2 | 61354075 | 61354538 | + | 578 | -224280 | Tank | bXPF |
| 1-668 | chr5 | 41844194 | 41844657 | + | 578 | -110 | Bod1l | bXPF |
| 1-622 | chr15 | 59828244 | 59828707 | + | 578 | -33516 | Gm19510 | bXPF |
| 1-387 | chr7 | 143013607 | 143014070 | + | 578 | 8179 | Tspan32 | bXPF |
| 1-384 | chr10 | 77211180 | 77211643 | + | 577 | -44881 | Col18a1 | bXPF |
| 1-549 | chr9 | 102645334 | 102645797 | + | 577 | 19269 | Anapc13 | bXPF |
| 1-721 | chr17 | 29093623 | 29094086 | + | 577 | 85 | Cdkn1a | bXPF |
| 1-569 | chr6 | 101374796 | 101375259 | + | 577 | 2870 | Pdzrn3 | bXPF |
| 1-487 | chr4 | 59331677 | 59332140 | + | 577 | 71857 | Gm12596 | bXPF |
| 1-393 | chr6 | 121245651 | 121246114 | + | 577 | -24 | Usp18 | bXPF |
| 1-664 | chr16 | 4273879 | 4274342 | + | 577 | 44881 | Gm5766 | bXPF |
| 1-486 | chr3 | 104220528 | 104220991 | + | 577 | -353 | Magi3 | bXPF |
| 1-603 | chr3 | 57093154 | 57093617 | + | 576 | 208534 | Tm4sf1 | bXPF |
| 1-655 | chr11 | 99024174 | 99024637 | + | 576 | -216 | Top2a | bXPF |
| 1-849 | chr10 | 39597173 | 39597636 | + | 576 | -15530 | Traf3ip2 | bXPF |
| 1-557 | chr9 | 69181302 | 69181765 | + | 576 | -108308 | Rora | bXPF |
| 1-100 | chr3 | 57093378 | 57093841 | + | 576 | 208310 | Tm4sf1 | bXPF |
| 1-438 | chr8 | 46210788 | 46211251 | + | 576 | -10 | Slc25a4 | bXPF |
| 1-451 | chr2 | 16450586 | 16451049 | + | 576 | 94513 | Plxdc2 | bXPF |

| | | | | | | | | |
|---|---|---|---|---|---|---|---|---|
| 1-714 | chr5 | 133884955 | 133885418 | + | 575 | -214562 | Gatsl2 | bXPF |
| 1-449 | chr7 | 99355674 | 99356137 | + | 575 | -2666 | Serpinh1 | bXPF |
| 1-110 | chr16 | 45968805 | 45969268 | + | 575 | 41377 | Plcxd2 | bXPF |
| 1-885 | chr8 | 103272319 | 103272782 | + | 575 | 74984 | 1600027J07Rik | bXPF |
| 1-286 | chr16 | 45968571 | 45969034 | + | 575 | 41611 | Plcxd2 | bXPF |
| 1-266 | chr9 | 72640296 | 72640759 | + | 575 | -21820 | Nedd4 | bXPF |
| 1-248 | chr10 | 122884262 | 122884725 | + | 575 | 101231 | Mirlet7i | bXPF |
| 1-166 | chr12 | 33957400 | 33957863 | + | 574 | -40 | Twist1 | bXPF |
| 1-533 | chr18 | 36411277 | 36411740 | + | 574 | 42987 | Pfdn1 | bXPF |
| 1-502 | chr3 | 79586791 | 79587254 | + | 574 | -4367 | Ppid | bXPF |
| 26299 | chr13 | 23400955 | 23401418 | + | 574 | 22680 | Abt1 | bXPF |
| 12785 | chr11 | 69662554 | 69663017 | + | 574 | -136 | Mpdu1 | bXPF |
| 1-246 | chr12 | 33957201 | 33957664 | + | 574 | -239 | Twist1 | bXPF |
| 1-213 | chr12 | 107807420 | 107807883 | + | 574 | -72163 | 4930465M20Rik | bXPF |
| 1-1022 | chr13 | 23400747 | 23401210 | + | 574 | 22888 | Abt1 | bXPF |
| 1-671 | chr15 | 31187303 | 31187766 | + | 573 | -36851 | Dap | bXPF |
| 1-550 | chr11 | 69662355 | 69662818 | + | 573 | 63 | Mpdu1 | bXPF |
| 1-476 | chr11 | 3535518 | 3535981 | + | 573 | 3543 | Smtn | bXPF |
| 1-229 | chr13 | 107890387 | 107890850 | + | 573 | -554 | Zswim6 | bXPF |
| 1-1005 | chr17 | 67988944 | 67989407 | + | 573 | 14933 | Arhgap28 | bXPF |
| 1-653 | chr3 | 18452264 | 18452727 | + | 573 | 27982 | 4930433B08Rik | bXPF |
| 1-707 | chr3 | 19953825 | 19954288 | + | 573 | -2998 | Cp | bXPF |
| 46388 | chr11 | 3535717 | 3536180 | + | 573 | 3344 | Smtn | bXPF |
| 1-700 | chr5 | 149228978 | 149229441 | + | 572 | -35795 | Alox5ap | bXPF |
| 1-1071 | chr8 | 126524515 | 126524978 | + | 572 | -49681 | Tarbp1 | bXPF |
| 1-579 | chr13 | 32471668 | 32472131 | + | 572 | -133355 | Gmds | bXPF |
| 1-838 | chr9 | 65908639 | 65909102 | + | 572 | -76 | Trip4 | bXPF |
| 1-204 | chr5 | 23214849 | 23215312 | + | 572 | 219273 | 5031425E22Rik | bXPF |
| 1-413 | chr1 | 82429087 | 82429550 | + | 572 | 90269 | Rhbdd1 | bXPF |
| 1-715 | chr1 | 45033172 | 45033635 | + | 572 | -278135 | Col3a1 | bXPF |
| 1-846 | chr2 | 53190929 | 53191392 | + | 571 | 27 | Prpf40a | bXPF |
| 1-597 | chr1 | 58445249 | 58445712 | + | 571 | 6 | Ppil3 | bXPF |
| 1-466 | chr4 | 55731970 | 55732433 | + | 571 | -199726 | Klf4 | bXPF |
| 1-420 | chr7 | 15962325 | 15962788 | + | 571 | 4979 | Ehd2 | bXPF |
| 1-365 | chr2 | 6378787 | 6379250 | + | 571 | 26282 | Usp6nl | bXPF |
| 1-571 | chr11 | 68665404 | 68665867 | + | 571 | -26280 | Myh10 | bXPF |
| 1-349 | chr2 | 103600205 | 103600668 | + | 571 | 34126 | Abtb2 | bXPF |
| 1-383 | chr19 | 53894850 | 53895313 | + | 571 | 2850 | Pdcd4 | bXPF |
| 1-580 | chr4 | 141011900 | 141012363 | + | 570 | 1494 | Mfap2 | bXPF |
| 1-143 | chr4 | 141012109 | 141012572 | + | 570 | 1703 | Mfap2 | bXPF |
| 1-493 | chr8 | 107293104 | 107293567 | + | 570 | -135 | Nfat5 | bXPF |
| 1-459 | chr19 | 17837443 | 17837906 | + | 570 | -42 | Pcsk5 | bXPF |
| 1-232 | chr6 | 65779153 | 65779616 | + | 570 | 422 | Prdm5 | bXPF |
| 1-515 | chr2 | 128127745 | 128128208 | + | 570 | 401 | Bcl2l11 | bXPF |
| 43070 | chr4 | 55732182 | 55732645 | + | 570 | -199938 | Klf4 | bXPF |
| 1-355 | chr12 | 35246299 | 35246762 | + | 570 | 199341 | Snx13 | bXPF |
| 1-697 | chr4 | 126227066 | 126227529 | + | 569 | -24587 | Thrap3 | bXPF |
| 1-616 | chr10 | 23003538 | 23004001 | + | 569 | -183641 | Tcf21 | bXPF |
| 1-455 | chr9 | 63941858 | 63942321 | + | 569 | 79970 | Smad6 | bXPF |
| 1-883 | chr6 | 47404487 | 47404950 | + | 569 | -49606 | Cul1 | bXPF |
| 1-784 | chr19 | 45058323 | 45058786 | + | 569 | 10978 | Sfxn3 | bXPF |
| 47119 | chr7 | 112023295 | 112023758 | + | 569 | 20 | Usp47 | bXPF |
| 1-318 | chr4 | 134704109 | 134704572 | + | 569 | -50 | Man1c1 | bXPF |
| 1-221 | chr7 | 112023088 | 112023551 | + | 569 | -187 | Usp47 | bXPF |
| 1-635 | chr13 | 15664790 | 15665253 | + | 568 | 137609 | 4933412O06Rik | bXPF |
| 1-610 | chr10 | 26564322 | 26564785 | + | 568 | 110098 | Gm8709 | bXPF |
| 1-489 | chr5 | 144988798 | 144989261 | + | 568 | 19022 | Kpna7 | bXPF |
| 1-314 | chr16 | 14350682 | 14351145 | + | 568 | -10645 | Abcc1 | bXPF |
| 1-482 | chr12 | 80365934 | 80366397 | + | 568 | 25512 | Mir1843a | bXPF |
| 28856 | chr12 | 69501975 | 69502438 | + | 568 | -26656 | 5830428M24Rik | bXPF |
| 21916 | chr12 | 80366133 | 80366596 | + | 568 | 25313 | Mir1843a | bXPF |
| 1-287 | chr9 | 109059521 | 109059984 | + | 568 | -29 | Trex1 | bXPF |
| 1-511 | chr18 | 13894660 | 13895123 | + | 567 | 77842 | Zfp521 | bXPF |

| | | | | | | | | |
|---|---|---|---|---|---|---|---|---|
| 1-364 | chr9 | 108201124 | 108201587 | + | 567 | -10972 | Bsn | bXPF |
| 1-585 | chr15 | 102350497 | 102350960 | + | 567 | 31 | Aaas | bXPF |
| 1-392 | chr16 | 29579185 | 29579648 | + | 567 | 82 | Opa1 | bXPF |
| 1-175 | chr3 | 21935402 | 21935865 | + | 567 | -141019 | Tbl1xr1 | bXPF |
| 1-276 | chr4 | 97575502 | 97575965 | + | 567 | -6210 | Nfia | bXPF |
| 1-117 | chr8 | 127720904 | 127721367 | + | 567 | 421483 | Pard3 | bXPF |
| 1-594 | chr2 | 166180325 | 166180788 | + | 567 | -24873 | Sulf2 | bXPF |
| 1-313 | chr13 | 79029961 | 79030424 | + | 566 | -831319 | Nr2f1 | bXPF |
| 1-209 | chr7 | 28372724 | 28373187 | + | 566 | -293 | Plekhg2 | bXPF |
| 1-642 | chr9 | 54803318 | 54803781 | + | 566 | 38801 | Crabp1 | bXPF |
| 1-433 | chr2 | 61454275 | 61454738 | + | 566 | -124080 | Tank | bXPF |
| 1-630 | chr10 | 75517726 | 75518189 | + | 566 | 15 | Gucd1 | bXPF |
| 1-607 | chr1 | 162008604 | 162009067 | + | 566 | 3866 | 2810442N19Rik | bXPF |
| 1-677 | chr6 | 4430919 | 4431382 | + | 566 | -74468 | Col1a2 | bXPF |
| 1-101 | chr7 | 28372930 | 28373393 | + | 566 | -499 | Plekhg2 | bXPF |
| 1-263 | chr5 | 36293167 | 36293630 | + | 565 | -8172 | Mir7689 | bXPF |
| 1-822 | chr9 | 84178613 | 84179076 | + | 565 | 230063 | Bckdhb | bXPF |
| 1-290 | chr11 | 103132727 | 103133190 | + | 565 | -381 | Hexim2 | bXPF |
| 1-656 | chr1 | 181144063 | 181144526 | + | 565 | -90 | Nvl | bXPF |
| 1-539 | chr1 | 186449230 | 186449693 | + | 565 | 109220 | A730004F24Rik | bXPF |
| 1-508 | chr5 | 36292964 | 36293427 | + | 565 | -8375 | Mir7689 | bXPF |
| 1-375 | chr4 | 55026094 | 55026557 | + | 565 | 78380 | Zfp462 | bXPF |
| 1-645 | chr1 | 186449011 | 186449474 | + | 565 | 109439 | A730004F24Rik | bXPF |
| 1-371 | chr1 | 36709784 | 36710247 | + | 564 | -90 | Actr1b | bXPF |
| 1-228 | chr15 | 99553688 | 99554151 | + | 564 | -25137 | Aqp2 | bXPF |
| 1-570 | chr9 | 121254032 | 121254495 | + | 564 | 22909 | Ulk4 | bXPF |
| 23743 | chr13 | 23438208 | 23438671 | + | 564 | -7422 | C230035I16Rik | bXPF |
| 1-446 | chr13 | 23438007 | 23438470 | + | 564 | -7221 | C230035I16Rik | bXPF |
| 1-332 | chr6 | 59412513 | 59412976 | + | 564 | 13546 | Gprin3 | bXPF |
| 1-298 | chr17 | 84112468 | 84112931 | + | 564 | 34039 | 4933433H22Rik | bXPF |
| 1-320 | chr13 | 44907665 | 44908128 | + | 564 | 94200 | Dtnbp1 | bXPF |
| 1-123 | chr17 | 35134908 | 35135371 | + | 563 | -39 | Bag6 | bXPF |
| 1-162 | chr1 | 106171372 | 106171835 | + | 563 | -79 | Gm20753 | bXPF |
| 1-733 | chr12 | 79346066 | 79346529 | + | 563 | 48946 | Rad51b | bXPF |
| 1-234 | chr1 | 106171116 | 106171579 | + | 563 | 177 | Gm20753 | bXPF |
| 1-120 | chr12 | 8913803 | 8914266 | + | 563 | -7273 | Laptm4a | bXPF |
| 22647 | chr14 | 76555355 | 76555818 | + | 563 | 1101 | Serp2 | bXPF |
| 1-669 | chr5 | 103620481 | 103620944 | + | 563 | 8691 | Slc10a6 | bXPF |
| 1-347 | chr14 | 76555146 | 76555609 | + | 563 | 1310 | Serp2 | bXPF |
| 1-464 | chr4 | 151411604 | 151412067 | + | 563 | 322265 | Gm13090 | bXPF |
| 1-456 | chr1 | 64226177 | 64226640 | + | 562 | -105019 | Klf7 | bXPF |
| 1-391 | chr17 | 35134703 | 35135166 | + | 562 | -244 | Bag6 | bXPF |
| 1-288 | chr8 | 25465544 | 25466007 | + | 562 | -52984 | Fgfr1 | bXPF |
| 17533 | chr7 | 83892004 | 83892467 | + | 562 | 235 | Mesdc2 | bXPF |
| 1-612 | chr7 | 83891801 | 83892264 | + | 562 | 32 | Mesdc2 | bXPF |
| 1-345 | chr1 | 59508458 | 59508921 | + | 562 | -7575 | Gm973 | bXPF |
| 1-552 | chr18 | 60628307 | 60628770 | + | 562 | -4233 | Synpo | bXPF |
| 1-419 | chr6 | 135252356 | 135252819 | + | 562 | 1749 | Gsg1 | bXPF |
| 12055 | chr6 | 4457720 | 4458183 | + | 561 | -47667 | Col1a2 | bXPF |
| 42948 | chr15 | 59823385 | 59823848 | + | 561 | -28657 | Gm19510 | bXPF |
| 1-390 | chr6 | 4457521 | 4457984 | + | 561 | -47866 | Col1a2 | bXPF |
| 1-544 | chr17 | 45359313 | 45359776 | + | 561 | 74163 | Cdc5l | bXPF |
| 1-525 | chr6 | 118883979 | 118884442 | + | 561 | 4052 | 4931430N09Rik | bXPF |
| 1-231 | chr7 | 126625617 | 126626080 | + | 561 | -378 | Nupr1 | bXPF |
| 1-223 | chr13 | 25629715 | 25630178 | + | 561 | 336370 | Gm11351 | bXPF |
| 1-258 | chr2 | 3423844 | 3424307 | + | 561 | -56 | Dclre1c | bXPF |
| 1-215 | chr13 | 25630006 | 25630469 | + | 561 | 336079 | Gm11351 | bXPF |
| 1-302 | chr7 | 118705448 | 118705911 | + | 560 | 59 | Gde1 | bXPF |
| 1-362 | chr1 | 14357701 | 14358164 | + | 560 | -47733 | Eya1 | bXPF |
| 1-440 | chr9 | 122923050 | 122923513 | + | 560 | 203 | Zfp105 | bXPF |
| 1-797 | chr8 | 45602080 | 45602543 | + | 560 | -25440 | Sorbs2 | bXPF |
| 1-510 | chr14 | 25105341 | 25105804 | + | 560 | 37669 | 4930572O13Rik | bXPF |
| 44927 | chr1 | 58029795 | 58030258 | + | 560 | 57 | Aox1 | bXPF |

| | | | | | | | | |
|---|---|---|---|---|---|---|---|---|
| 1-529 | chr7 | 130965910 | 130966373 | + | 560 | 29938 | Htra1 | bXPF |
| 1-284 | chr12 | 79481906 | 79482369 | + | 560 | 184786 | Rad51b | bXPF |
| 1-788 | chr4 | 97575301 | 97575764 | + | 559 | -6411 | Nfia | bXPF |
| 1-333 | chr1 | 58029595 | 58030058 | + | 559 | -143 | Aox1 | bXPF |
| 1-574 | chr15 | 59777262 | 59777725 | + | 559 | 17466 | Gm19510 | bXPF |
| 32874 | chr11 | 83271527 | 83271990 | + | 559 | 14968 | Slfn14 | bXPF |
| 1-857 | chr1 | 64809144 | 64809607 | + | 559 | -71625 | Fzd5 | bXPF |
| 1-216 | chr7 | 19715935 | 19716398 | + | 559 | -737 | Tomm40 | bXPF |
| 1-210 | chr10 | 39369314 | 39369777 | + | 559 | -254 | Fyn | bXPF |
| 1-395 | chr7 | 19715698 | 19716161 | + | 559 | -500 | Tomm40 | bXPF |
| 1-593 | chr16 | 17208850 | 17209313 | + | 559 | 946 | Rimbp3 | bXPF |
| 1-606 | chr13 | 24792347 | 24792810 | + | 558 | -9079 | BC005537 | bXPF |
| 1-589 | chr1 | 92934084 | 92934547 | + | 558 | -93 | Capn10 | bXPF |
| 1-581 | chr12 | 102498538 | 102499001 | + | 558 | 28859 | Golga5 | bXPF |
| 1-376 | chr10 | 62865183 | 62865646 | + | 558 | 14600 | Tet1 | bXPF |
| 1-377 | chr5 | 31048317 | 31048780 | + | 558 | 14 | Slc5a6 | bXPF |
| 1-776 | chr15 | 8684250 | 8684713 | + | 558 | 26326 | Slc1a3 | bXPF |
| 1-200 | chr3 | 109936406 | 109936869 | + | 558 | 206835 | Ntng1 | bXPF |
| 1-186 | chr8 | 105254009 | 105254472 | + | 558 | 913 | B3gnt9 | bXPF |
| 1-794 | chr15 | 77214850 | 77215313 | + | 558 | -61269 | Rbfox2 | bXPF |
| 1-249 | chr7 | 34341976 | 34342439 | + | 557 | 47333 | Lsm14a | bXPF |
| 1-344 | chr3 | 109936172 | 109936635 | + | 557 | 207069 | Ntng1 | bXPF |
| 1-693 | chr5 | 17574263 | 17574726 | + | 557 | -322 | Sema3c | bXPF |
| 1-479 | chr5 | 66549497 | 66549960 | + | 557 | 69089 | Apbb2 | bXPF |
| 1-340 | chr2 | 57182182 | 57182645 | + | 557 | 582 | 4930555B11Rik | bXPF |
| 1-201 | chr7 | 25755625 | 25756088 | + | 557 | -1136 | Hnrnpul1 | bXPF |
| 1-339 | chr4 | 65420636 | 65421099 | + | 557 | -184119 | Trim32 | bXPF |
| 1-708 | chr1 | 45864576 | 45865039 | + | 557 | 60787 | Slc40a1 | bXPF |
| 1-219 | chr2 | 32882178 | 32882641 | + | 556 | 6275 | Fam129b | bXPF |
| 1-1059 | chr19 | 57823599 | 57824062 | + | 556 | 212796 | Atrnl1 | bXPF |
| 1-272 | chr8 | 90869508 | 90869971 | + | 556 | 40904 | Chd9 | bXPF |
| 1-296 | chr16 | 4652831 | 4653294 | + | 556 | 13117 | Vasn | bXPF |
| 1-460 | chr7 | 27474341 | 27474804 | + | 556 | 732 | Sertad3 | bXPF |
| 1-631 | chr15 | 62079177 | 62079640 | + | 556 | 40151 | Pvt1 | bXPF |
| 1-621 | chr10 | 86339654 | 86340117 | + | 556 | 39473 | Timp3 | bXPF |
| 21551 | chr4 | 80912026 | 80912489 | + | 556 | 1571 | Lurap1l | bXPF |
| 1-566 | chr4 | 80911775 | 80912238 | + | 556 | 1320 | Lurap1l | bXPF |
| 1-609 | chr14 | 123532860 | 123533323 | + | 555 | 94053 | Nalcn | bXPF |
| 27030 | chr14 | 123533065 | 123533528 | + | 555 | 93848 | Nalcn | bXPF |
| 1-174 | chr8 | 123949316 | 123949779 | + | 555 | -282 | Nup133 | bXPF |
| 1-644 | chr15 | 75820833 | 75821296 | + | 555 | 20844 | Zc3h3 | bXPF |
| 1-268 | chr8 | 71881796 | 71882259 | + | 555 | -41 | Zfp709 | bXPF |
| 1-711 | chr11 | 100939367 | 100939830 | + | 555 | -58 | Stat3 | bXPF |
| 1-108 | chr8 | 123949028 | 123949491 | + | 555 | 6 | Nup133 | bXPF |
| 1-242 | chr3 | 39043532 | 39043995 | + | 555 | 156823 | Fat4 | bXPF |
| 1-245 | chr4 | 7560381 | 7560844 | + | 555 | -76 | 8430436N08Rik | bXPF |
| 1-327 | chr9 | 51419053 | 51419516 | + | 554 | -90367 | 1810046K07Rik | bXPF |
| 1-124 | chr19 | 5797467 | 5797930 | + | 554 | -1717 | Mascrna | bXPF |
| 1-592 | chr18 | 56899545 | 56900008 | + | 554 | 25772 | March3 | bXPF |
| 1-372 | chr1 | 195033970 | 195034433 | + | 554 | -2839 | Mir29b-2 | bXPF |
| 10959 | chr16 | 24490887 | 24491350 | + | 554 | -38196 | Morf4l1-ps1 | bXPF |
| 1-530 | chr4 | 131850273 | 131850736 | + | 554 | 7033 | Mecr | bXPF |
| 1-427 | chr4 | 63030144 | 63030607 | + | 554 | 64801 | Zfp618 | bXPF |
| 1-648 | chr6 | 108709563 | 108710026 | + | 554 | -48860 | 0610040F04Rik | bXPF |
| 1-323 | chr6 | 136468579 | 136469042 | + | 554 | -50041 | Atf7ip | bXPF |
| 1-240 | chr3 | 141465298 | 141465761 | + | 553 | -35 | Unc5c | bXPF |
| 33970 | chr4 | 62755467 | 62755930 | + | 553 | 81724 | Rgs3 | bXPF |
| 1-178 | chr11 | 69037597 | 69038060 | + | 553 | -7809 | Aurkb | bXPF |
| 1-194 | chr4 | 62755260 | 62755723 | + | 553 | 81517 | Rgs3 | bXPF |
| 1-442 | chr6 | 99811003 | 99811466 | + | 553 | -84842 | Prok2 | bXPF |
| 1-921 | chr11 | 32925638 | 32926101 | + | 553 | -101275 | Smim23 | bXPF |
| 1-1003 | chr6 | 129229871 | 129230334 | + | 553 | 7943 | 2310001H17Rik | bXPF |
| 1-189 | chr5 | 90750508 | 90750971 | + | 553 | -8559 | Cxcl5 | bXPF |

| | | | | | | | | |
|---|---|---|---|---|---|---|---|---|
| 1-681 | chr9 | 20385010 | 20385473 | + | 553 | -83 | Zfp560 | bXPF |
| 1-497 | chr12 | 85288045 | 85288508 | + | 552 | -315 | Zc2hc1c | bXPF |
| 1-171 | chr3 | 21953360 | 21953823 | + | 552 | -123061 | Tbl1xr1 | bXPF |
| 1-584 | chr4 | 144949735 | 144950198 | + | 552 | 56829 | Dhrs3 | bXPF |
| 1 | chr6 | 112924942 | 112925405 | + | 552 | 22093 | Srgap3 | bXPF |
| 42370 | chr9 | 20385238 | 20385701 | + | 552 | -311 | Zfp560 | bXPF |
| 1-615 | chr11 | 32226441 | 32226904 | + | 552 | 167 | Mpg | bXPF |
| 1-253 | chr7 | 145330017 | 145330480 | + | 552 | 15413 | Mrgprd | bXPF |
| 1-374 | chr6 | 112924742 | 112925205 | + | 552 | 22293 | Srgap3 | bXPF |
| 1-774 | chr17 | 17792528 | 17792991 | + | 552 | -37429 | Mir99b | bXPF |
| 1-196 | chr9 | 41950473 | 41950936 | + | 551 | 173585 | Sorl1 | bXPF |
| 1-491 | chr3 | 84666010 | 84666473 | + | 551 | 49 | Tmem154 | bXPF |
| 1-109 | chr8 | 70523090 | 70523553 | + | 551 | -141 | Kxd1 | bXPF |
| 1-265 | chr2 | 144527318 | 144527781 | + | 551 | -151 | Dzank1 | bXPF |
| 1-342 | chr5 | 130128683 | 130129146 | + | 551 | -15974 | Kctd7 | bXPF |
| 1-445 | chr4 | 97695565 | 97696028 | + | 551 | -81830 | Nfia | bXPF |
| 1-505 | chr16 | 14979172 | 14979635 | + | 551 | 72757 | Efcab1 | bXPF |
| 1-319 | chr15 | 76904026 | 76904489 | + | 551 | 186 | Rpl8 | bXPF |
| 1-274 | chr10 | 60831219 | 60831682 | + | 551 | 131 | Unc5b | bXPF |
| 1-560 | chr8 | 124643929 | 124644392 | + | 551 | 19209 | 2310022B05Rik | bXPF |
| 1-148 | chr7 | 110630318 | 110630781 | + | 550 | 2880 | Adm | bXPF |
| 1-185 | chr7 | 110630060 | 110630523 | + | 550 | 2622 | Adm | bXPF |
| 1-281 | chr8 | 70212403 | 70212866 | + | 550 | -353 | Slc25a42 | bXPF |
| 1-453 | chr19 | 5940118 | 5940581 | + | 550 | -15533 | Cdc42ep2 | bXPF |
| 1-414 | chr11 | 45274901 | 45275364 | + | 550 | -158204 | Gm12159 | bXPF |
| 1-513 | chr3 | 116129376 | 116129839 | + | 550 | 81 | Vcam1 | bXPF |
| 1-873 | chr7 | 51759701 | 51760164 | + | 550 | -102083 | Gas2 | bXPF |
| 1-604 | chr9 | 51394279 | 51394742 | + | 550 | -65593 | 1810046K07Rik | bXPF |
| 1-717 | chr14 | 30479419 | 30479882 | + | 550 | 85 | Dcp1a | bXPF |
| 1-423 | chr13 | 60364967 | 60365430 | + | 549 | 159727 | Gm5084 | bXPF |
| 1-775 | chr7 | 99381147 | 99381610 | + | 549 | -171 | Gdpd5 | bXPF |
| 1-540 | chr15 | 73154559 | 73155022 | + | 549 | 30157 | Ago2 | bXPF |
| 1-546 | chr8 | 122444348 | 122444811 | + | 549 | 923 | 9330133O14Rik | bXPF |
| 1-190 | chr8 | 126524723 | 126525186 | + | 549 | -49889 | Tarbp1 | bXPF |
| 1-155 | chr13 | 60365170 | 60365633 | + | 549 | 159930 | Gm5084 | bXPF |
| 1-624 | chr16 | 13437005 | 13437468 | + | 549 | -12287 | Mir193b | bXPF |
| 1-301 | chr6 | 49073499 | 49073962 | + | 549 | -65 | Malsu1 | bXPF |
| 1-381 | chr11 | 120800030 | 120800493 | + | 548 | -3866 | Dus1l | bXPF |
| 1-205 | chr9 | 41771246 | 41771709 | + | 548 | 189551 | Mir125b-1 | bXPF |
| 1-134 | chr15 | 34420072 | 34420535 | + | 548 | 22922 | Rpl30 | bXPF |
| 1-163 | chr11 | 120800240 | 120800703 | + | 548 | -4076 | Dus1l | bXPF |
| 1-306 | chr4 | 119422120 | 119422583 | + | 548 | 69 | Ppcs | bXPF |
| 1-492 | chr5 | 131600446 | 131600909 | + | 548 | -279592 | 4930563F08Rik | bXPF |
| 1-728 | chr11 | 48870964 | 48871427 | + | 548 | 151 | Irgm1 | bXPF |
| 1-370 | chr3 | 21937960 | 21938423 | + | 548 | -138461 | Tbl1xr1 | bXPF |
| 18629 | chr9 | 41771530 | 41771993 | + | 548 | 189835 | Mir125b-1 | bXPF |
| 42795 | chr7 | 99381393 | 99381856 | + | 548 | 75 | Gdpd5 | bXPF |
| 1-779 | chr4 | 97757375 | 97757838 | + | 547 | -20020 | Nfia | bXPF |
| 1-767 | chr15 | 43282495 | 43282958 | + | 547 | 10 | Eif3e | bXPF |
| 1-106 | chr4 | 97757608 | 97758071 | + | 547 | -19787 | Nfia | bXPF |
| 1-137 | chr19 | 8713625 | 8714088 | + | 547 | 41 | Slc3a2 | bXPF |
| 1-908 | chr11 | 103312037 | 103312500 | + | 547 | 32434 | Arhgap27 | bXPF |
| 1-241 | chr3 | 7503110 | 7503573 | + | 547 | -85 | Zc2hc1a | bXPF |
| 1-507 | chr4 | 133369467 | 133369930 | + | 547 | -74 | Slc9a1 | bXPF |
| 21186 | chr11 | 98775023 | 98775486 | + | 547 | 123 | Nr1d1 | bXPF |
| 14246 | chr15 | 4028061 | 4028524 | + | 547 | -886 | BC037032 | bXPF |
| 1-254 | chr9 | 44417849 | 44418312 | + | 546 | -73 | Ccdc84 | bXPF |
| 1-357 | chr8 | 95434716 | 95435179 | + | 546 | -78 | Cfap20 | bXPF |
| 31778 | chr11 | 63130607 | 63131070 | + | 546 | -619 | Pmp22 | bXPF |
| 1-382 | chr5 | 67306663 | 67307126 | + | 546 | -61 | Slc30a9 | bXPF |
| 1-660 | chr11 | 63130407 | 63130870 | + | 546 | -819 | Pmp22 | bXPF |
| 1-112 | chr3 | 151351714 | 151352177 | + | 546 | -85937 | Adgrl4 | bXPF |
| 1-269 | chr3 | 151351467 | 151351930 | + | 546 | -86184 | Adgrl4 | bXPF |

| | | | | | | | | |
|---|---|---|---|---|---|---|---|---|
| 1-577 | chr5 | 88553943 | 88554406 | + | 546 | -309 | Utp3 | bXPF |
| 1-875 | chr11 | 87086592 | 87087055 | + | 546 | -46 | Smg8 | bXPF |
| 1-102 | chr9 | 44418050 | 44418513 | + | 546 | -274 | Ccdc84 | bXPF |
| 1-133 | chr6 | 92941435 | 92941898 | + | 545 | 1084 | 9530026P05Rik | bXPF |
| 1-503 | chr14 | 51090364 | 51090827 | + | 545 | -482 | Rnase4 | bXPF |
| 1-576 | chr3 | 20034792 | 20035255 | + | 545 | 287 | Hps3 | bXPF |
| 1-315 | chr8 | 25518683 | 25519146 | + | 545 | 155 | Fgfr1 | bXPF |
| 1-128 | chr12 | 40222632 | 40223095 | + | 545 | -52 | Mir1938 | bXPF |
| 1-519 | chr3 | 81047237 | 81047700 | + | 545 | 11052 | Pdgfc | bXPF |
| 1-338 | chr18 | 5133927 | 5134390 | + | 545 | 87569 | Svil | bXPF |
| 1-271 | chr2 | 160366861 | 160367324 | + | 545 | -27 | Mafb | bXPF |
| 1-227 | chr6 | 92941236 | 92941699 | + | 545 | 885 | 9530026P05Rik | bXPF |
| 1-756 | chr1 | 90225648 | 90226111 | + | 545 | 21899 | Ackr3 | bXPF |
| 1-596 | chr6 | 117437693 | 117438156 | + | 544 | 269389 | Cxcl12 | bXPF |
| 1-467 | chr18 | 37806056 | 37806519 | + | 544 | -123 | Pcdhgc3 | bXPF |
| 1-173 | chr17 | 80206738 | 80207201 | + | 544 | 336 | Srsf7 | bXPF |
| 1-283 | chr7 | 96210284 | 96210747 | + | 544 | -122 | Tenm4 | bXPF |
| 1-261 | chr4 | 81309391 | 81309854 | + | 544 | 133193 | Mpdz | bXPF |
| 1-474 | chr12 | 99392981 | 99393444 | + | 544 | 56862 | Foxn3 | bXPF |
| 1-260 | chr10 | 20951966 | 20952429 | + | 544 | -350 | Ahi1 | bXPF |
| 1-422 | chr1 | 59516090 | 59516553 | + | 544 | 57 | Gm973 | bXPF |
| 1-986 | chr7 | 88200002 | 88200465 | + | 544 | -77852 | Ctsc | bXPF |
| 1-929 | chr13 | 22020969 | 22021432 | + | 543 | -11459 | Prss16 | bXPF |
| 1-366 | chr2 | 37768793 | 37769256 | + | 543 | -7225 | Crb2 | bXPF |
| 1-640 | chr14 | 47177994 | 47178457 | + | 543 | 11177 | Gch1 | bXPF |
| 1-164 | chr13 | 112800550 | 112801013 | + | 543 | 4 | Plpp1 | bXPF |
| 1-555 | chr11 | 115628088 | 115628551 | + | 543 | -24 | Slc25a19 | bXPF |
| 1-317 | chr12 | 104983358 | 104983821 | + | 543 | 15088 | Syne3 | bXPF |
| 1-378 | chr7 | 89770044 | 89770507 | + | 543 | 133354 | Ccdc81 | bXPF |
| 1-307 | chr5 | 53376921 | 53377384 | + | 543 | 110046 | Smim20 | bXPF |
| 42979 | chr13 | 112800798 | 112801261 | + | 543 | 252 | Plpp1 | bXPF |
| 20821 | chr16 | 84735268 | 84735731 | + | 543 | -197 | Mrpl39 | bXPF |
| 31048 | chr18 | 9213284 | 9213747 | + | 542 | 659 | Fzd8 | bXPF |
| 1-280 | chr19 | 4810815 | 4811278 | + | 542 | 588 | Rbm14 | bXPF |
| 1-198 | chr9 | 62811603 | 62812066 | + | 542 | -186 | Fem1b | bXPF |
| 1-532 | chr9 | 102784311 | 102784774 | + | 542 | 44894 | Gm5627 | bXPF |
| 30682 | chr9 | 102784513 | 102784976 | + | 542 | 45096 | Gm5627 | bXPF |
| 1-188 | chr8 | 74297897 | 74298360 | + | 542 | -575046 | Isx | bXPF |
| 1-107 | chr9 | 62811833 | 62812296 | + | 542 | -416 | Fem1b | bXPF |
| 1-623 | chr15 | 61189524 | 61189987 | + | 542 | -268485 | A1bg | bXPF |
| 44197 | chr19 | 4811028 | 4811491 | + | 542 | 375 | Rbm14 | bXPF |
| 1-678 | chr15 | 53134314 | 53134777 | + | 542 | 211638 | Ext1 | bXPF |
| 1-522 | chr8 | 72135265 | 72135728 | + | 541 | 204 | Tpm4 | bXPF |
| 1-115 | chr8 | 72135476 | 72135939 | + | 541 | 415 | Tpm4 | bXPF |
| 1-727 | chr6 | 72512503 | 72512966 | + | 541 | -31657 | Capg | bXPF |
| 1-672 | chr11 | 50174470 | 50174933 | + | 541 | -150 | Mrnip | bXPF |
| 1-292 | chr4 | 155837542 | 155838005 | + | 541 | -1907 | Mxra8 | bXPF |
| 1-295 | chr18 | 54987901 | 54988364 | + | 541 | 2048 | Zfp608 | bXPF |
| 1-356 | chr14 | 103346701 | 103347164 | + | 541 | -132 | Mycbp2 | bXPF |
| 1-404 | chr6 | 72655300 | 72655763 | + | 541 | 25700 | Gm15401 | bXPF |
| 1-783 | chr8 | 126566353 | 126566816 | + | 541 | 26852 | Irf2bp2 | bXPF |
| 1-257 | chr17 | 35590796 | 35591259 | + | 541 | 23542 | 2300002M23Rik | bXPF |
| 1-350 | chr2 | 154372323 | 154372786 | + | 540 | 165 | Cdk5rap1 | bXPF |
| 1-291 | chr3 | 96401970 | 96402433 | + | 540 | 12632 | Terc | bXPF |
| 35431 | chr17 | 25874984 | 25875447 | + | 540 | -285 | Mettl26 | bXPF |
| 1-452 | chr1 | 63768047 | 63768510 | + | 540 | 989 | 4933402D24Rik | bXPF |
| 43009 | chr3 | 96401632 | 96402095 | + | 540 | 12970 | Terc | bXPF |
| 1-121 | chr2 | 154372522 | 154372985 | + | 540 | -34 | Cdk5rap1 | bXPF |
| 1-312 | chr11 | 52360579 | 52361042 | + | 540 | -52 | Vdac1 | bXPF |
| 1-439 | chr13 | 41709067 | 41709530 | + | 540 | 58536 | Gm5082 | bXPF |
| 1-1534 | chr1 | 4857347 | 4857810 | + | 822 | -116 | Tcea1 | bXPF_tRA |
| 1-1623 | chr1 | 12718223 | 12718686 | + | 715 | -91 | Sulf1 | bXPF_tRA |
| 1-951 | chr1 | 24553568 | 24554031 | + | 550 | 33638 | Col19a1 | bXPF_tRA |

| | | | | | | | | |
|---|---|---|---|---|---|---|---|---|
| 1-312 | chr1 | 33814300 | 33814763 | + | 545 | 64 | Zfp451 | bXPF_tRA |
| 1-705 | chr1 | 36064734 | 36065197 | + | 680 | -3435 | Hs6st1 | bXPF_tRA |
| 1-126 | chr1 | 36471058 | 36471521 | + | 558 | -308 | Cnnm4 | bXPF_tRA |
| 1-1313 | chr1 | 37890265 | 37890728 | + | 870 | -57 | Mrpl30 | bXPF_tRA |
| 1-697 | chr1 | 39713987 | 39714450 | + | 607 | 6771 | Rfx8 | bXPF_tRA |
| 1-915 | chr1 | 40444776 | 40445239 | + | 633 | 5378 | Il1rl1 | bXPF_tRA |
| 1-379 | chr1 | 40444994 | 40445457 | + | 633 | 5596 | Il1rl1 | bXPF_tRA |
| 1-872 | chr1 | 44147418 | 44147881 | + | 616 | -95 | Ercc5 | bXPF_tRA |
| 1-993 | chr1 | 45043199 | 45043662 | + | 675 | -268108 | Col3a1 | bXPF_tRA |
| 41640 | chr1 | 45043411 | 45043874 | + | 675 | -267896 | Col3a1 | bXPF_tRA |
| 1-562 | chr1 | 45502792 | 45503255 | + | 596 | 259 | Col5a2 | bXPF_tRA |
| 1-1065 | chr1 | 45672792 | 45673255 | + | 595 | -122478 | Wdr75 | bXPF_tRA |
| 1-1055 | chr1 | 45837637 | 45838100 | + | 589 | 42367 | Wdr75 | bXPF_tRA |
| 1-1148 | chr1 | 51689750 | 51690213 | + | 602 | -211582 | Nabp1 | bXPF_tRA |
| 1-1867 | chr1 | 54438791 | 54439254 | + | 732 | 4 | Gtf3c3 | bXPF_tRA |
| 1-1500 | chr1 | 55087813 | 55088276 | + | 555 | -104 | Hspe1 | bXPF_tRA |
| 1-1750 | chr1 | 57406355 | 57406818 | + | 843 | 38 | Maip1 | bXPF_tRA |
| 1-608 | chr1 | 58445197 | 58445660 | + | 582 | 58 | Ppil3 | bXPF_tRA |
| 1-129 | chr1 | 58445396 | 58445859 | + | 582 | -141 | Ppil3 | bXPF_tRA |
| 42887 | chr1 | 60180269 | 60180732 | + | 559 | -99 | Nbeal1 | bXPF_tRA |
| 18629 | chr1 | 60180655 | 60181118 | + | 559 | 287 | Nbeal1 | bXPF_tRA |
| 1-497 | chr1 | 60825809 | 60826272 | + | 547 | 79652 | Cd28 | bXPF_tRA |
| 1-378 | chr1 | 62703269 | 62703732 | + | 567 | 183 | Nrp2 | bXPF_tRA |
| 1-460 | chr1 | 63176564 | 63177027 | + | 650 | 27 | Ndufs1 | bXPF_tRA |
| 1-402 | chr1 | 63176775 | 63177238 | + | 650 | 175 | Eef1b2 | bXPF_tRA |
| 1-1333 | chr1 | 63180976 | 63181439 | + | 561 | 2184 | Snora41 | bXPF_tRA |
| 1-1813 | chr1 | 64226046 | 64226509 | + | 663 | -104888 | Klf7 | bXPF_tRA |
| 1-1354 | chr1 | 64305482 | 64305945 | + | 699 | -184324 | Klf7 | bXPF_tRA |
| 1-1607 | chr1 | 71556971 | 71557434 | + | 697 | 46 | Atic | bXPF_tRA |
| 12055 | chr1 | 74386748 | 74387211 | + | 981 | -4630 | Ctdsp1 | bXPF_tRA |
| 1-1494 | chr1 | 74587851 | 74588314 | + | 680 | -54 | Zfp142 | bXPF_tRA |
| 1-1400 | chr1 | 78326337 | 78326800 | + | 556 | 16222 | Sgpp2 | bXPF_tRA |
| 11324 | chr1 | 79775884 | 79776347 | + | 1000 | 97 | Mrpl44 | bXPF_tRA |
| 1-1327 | chr1 | 82283177 | 82283640 | + | 549 | 8031 | Irs1 | bXPF_tRA |
| 1-1850 | chr1 | 82290828 | 82291291 | + | 1000 | 380 | Irs1 | bXPF_tRA |
| 1-552 | chr1 | 82839158 | 82839621 | + | 574 | -71 | Agfg1 | bXPF_tRA |
| 1-1201 | chr1 | 84218552 | 84219015 | + | 563 | 178 | Mir6353 | bXPF_tRA |
| 1-633 | chr1 | 84839265 | 84839728 | + | 639 | -192 | Trip12 | bXPF_tRA |
| 1-245 | chr1 | 84839500 | 84839963 | + | 639 | -110 | Fbxo36 | bXPF_tRA |
| 1-1893 | chr1 | 86487320 | 86487783 | + | 725 | -39185 | Ptma | bXPF_tRA |
| 1-208 | chr1 | 86503796 | 86504259 | + | 686 | -22709 | Ptma | bXPF_tRA |
| 1-1639 | chr1 | 86504113 | 86504576 | + | 685 | -22392 | Ptma | bXPF_tRA |
| 1-287 | chr1 | 86525050 | 86525513 | + | 556 | -1455 | Ptma | bXPF_tRA |
| 1-171 | chr1 | 86525258 | 86525721 | + | 556 | -1247 | Ptma | bXPF_tRA |
| 8402 | chr1 | 86586873 | 86587336 | + | 880 | 4 | Cops7b | bXPF_tRA |
| 1-1283 | chr1 | 87213709 | 87214172 | + | 724 | 1 | Eif4e2 | bXPF_tRA |
| 1-1728 | chr1 | 90100560 | 90101023 | + | 749 | 52610 | Iqca | bXPF_tRA |
| 1-548 | chr1 | 90100760 | 90101223 | + | 749 | 52410 | Iqca | bXPF_tRA |
| 1-1412 | chr1 | 90334951 | 90335414 | + | 752 | 131202 | Ackr3 | bXPF_tRA |
| 1-672 | chr1 | 90603204 | 90603667 | + | 581 | 10 | Cops8 | bXPF_tRA |
| 1-991 | chr1 | 90974980 | 90975443 | + | 583 | -5591 | Rab17 | bXPF_tRA |
| 1-103 | chr1 | 90975181 | 90975644 | + | 583 | -5792 | Rab17 | bXPF_tRA |
| 1-1364 | chr1 | 91250102 | 91250565 | + | 602 | 14 | Ube2f | bXPF_tRA |
| 1-1382 | chr1 | 93305737 | 93306200 | + | 648 | -89 | Mterf4 | bXPF_tRA |
| 1-464 | chr1 | 93478239 | 93478702 | + | 551 | 447 | Hdlbp | bXPF_tRA |
| 1-1100 | chr1 | 97769888 | 97770351 | + | 609 | -27 | Ppip5k2 | bXPF_tRA |
| 1-188 | chr1 | 97770213 | 97770676 | + | 609 | 272 | Gin1 | bXPF_tRA |
| 1-973 | chr1 | 97976932 | 97977395 | + | 614 | 70630 | B230216N24Rik | bXPF_tRA |
| 13150 | chr1 | 97977131 | 97977594 | + | 614 | 70431 | B230216N24Rik | bXPF_tRA |
| 1-320 | chr1 | 106759270 | 106759733 | + | 673 | 241 | Kdsr | bXPF_tRA |
| 1-976 | chr1 | 111864664 | 111865127 | + | 768 | 23 | Dsel | bXPF_tRA |
| 1-1345 | chr1 | 118311137 | 118311600 | + | 670 | -236 | Tsn | bXPF_tRA |
| 1-537 | chr1 | 118976429 | 118976892 | + | 582 | 22440 | Mir6346 | bXPF_tRA |

| | | | | | | | | |
|---|---|---|---|---|---|---|---|---|
| 1-776 | chr1 | 118993632 | 118994095 | + | 570 | 39643 | Mir6346 | bXPF_tRA |
| 1-1188 | chr1 | 119053657 | 119054120 | + | 563 | -269 | Gli2 | bXPF_tRA |
| 1-1591 | chr1 | 119526067 | 119526530 | + | 798 | 144 | Tmem185b | bXPF_tRA |
| 1-239 | chr1 | 119526271 | 119526734 | + | 797 | 348 | Tmem185b | bXPF_tRA |
| 1-1270 | chr1 | 120757219 | 120757682 | + | 549 | 154963 | En1 | bXPF_tRA |
| 1-710 | chr1 | 121567833 | 121568296 | + | 623 | -84 | Ddx18 | bXPF_tRA |
| 1-945 | chr1 | 126830625 | 126831088 | + | 568 | -224 | Nckap5 | bXPF_tRA |
| 1-1199 | chr1 | 128416827 | 128417290 | + | 609 | 358 | Dars | bXPF_tRA |
| 1-1890 | chr1 | 132417191 | 132417654 | + | 902 | -31 | Dstyk | bXPF_tRA |
| 1-1562 | chr1 | 133033892 | 133034355 | + | 691 | -3536 | Mdm4 | bXPF_tRA |
| 1-1828 | chr1 | 134556906 | 134557369 | + | 724 | -3041 | Kdm5b | bXPF_tRA |
| 1-1172 | chr1 | 135636029 | 135636492 | + | 544 | -50905 | Nav1 | bXPF_tRA |
| 43009 | chr1 | 135636232 | 135636695 | + | 544 | -51108 | Nav1 | bXPF_tRA |
| 1-371 | chr1 | 135770977 | 135771440 | + | 580 | 5123 | Phlda3 | bXPF_tRA |
| 1-897 | chr1 | 140183250 | 140183713 | + | 581 | -70 | Cfh | bXPF_tRA |
| 1-1567 | chr1 | 152556160 | 152556623 | + | 794 | 68826 | Rgl1 | bXPF_tRA |
| 1-572 | chr1 | 153130022 | 153130485 | + | 553 | 56194 | Lamc2 | bXPF_tRA |
| 1-255 | chr1 | 153130225 | 153130688 | + | 554 | 55991 | Lamc2 | bXPF_tRA |
| 1-1203 | chr1 | 153487362 | 153487825 | + | 677 | 67 | Dhx9 | bXPF_tRA |
| 1-1596 | chr1 | 153572193 | 153572656 | + | 821 | -22710 | Npl | bXPF_tRA |
| 1-1075 | chr1 | 155050097 | 155050560 | + | 608 | 49308 | Ier5 | bXPF_tRA |
| 25569 | chr1 | 155050302 | 155050765 | + | 608 | 49103 | Ier5 | bXPF_tRA |
| 1-923 | chr1 | 155091689 | 155092152 | + | 587 | 7716 | Ier5 | bXPF_tRA |
| 1-1661 | chr1 | 155093477 | 155093940 | + | 640 | 5928 | Ier5 | bXPF_tRA |
| 1-329 | chr1 | 155972853 | 155973316 | + | 612 | 171 | Cep350 | bXPF_tRA |
| 1-132 | chr1 | 156035716 | 156036179 | + | 577 | -89 | Tor1aip2 | bXPF_tRA |
| 1-799 | chr1 | 156036355 | 156036818 | + | 634 | -106 | Tor1aip1 | bXPF_tRA |
| 1-1170 | chr1 | 158732882 | 158733345 | + | 574 | 224325 | Pappa2 | bXPF_tRA |
| 1-346 | chr1 | 162003883 | 162004346 | + | 550 | -855 | 2810442N19Rik | bXPF_tRA |
| 1-1231 | chr1 | 163994590 | 163995053 | + | 700 | -40 | BC055324 | bXPF_tRA |
| 35065 | chr1 | 163994798 | 163995261 | + | 700 | 84 | Mettl18 | bXPF_tRA |
| 1-918 | chr1 | 165642173 | 165642636 | + | 607 | -7863 | Mpzl1 | bXPF_tRA |
| 1-1817 | chr1 | 169165888 | 169166351 | + | 698 | 365345 | Nuf2 | bXPF_tRA |
| 1-571 | chr1 | 169166089 | 169166552 | + | 599 | 365144 | Nuf2 | bXPF_tRA |
| 1-1862 | chr1 | 169531205 | 169531668 | + | 875 | 28 | Nuf2 | bXPF_tRA |
| 1-353 | chr1 | 170110713 | 170111176 | + | 620 | -22000 | Ddr2 | bXPF_tRA |
| 1-101 | chr1 | 170110965 | 170111428 | + | 620 | -22252 | Ddr2 | bXPF_tRA |
| 1-1548 | chr1 | 171033670 | 171034133 | + | 811 | 14975 | Fcgr4 | bXPF_tRA |
| 1-1549 | chr1 | 171102713 | 171103176 | + | 773 | -18717 | Cfap126 | bXPF_tRA |
| 1-219 | chr1 | 171102915 | 171103378 | + | 773 | -18515 | Cfap126 | bXPF_tRA |
| 1-729 | chr1 | 171247020 | 171247483 | + | 593 | -129 | Ndufs2 | bXPF_tRA |
| 1-659 | chr1 | 172206271 | 172206734 | + | 666 | 302 | Pea15a | bXPF_tRA |
| 1-221 | chr1 | 172206472 | 172206935 | + | 666 | 101 | Pea15a | bXPF_tRA |
| 1-1643 | chr1 | 177648988 | 177649451 | + | 718 | -6276 | 2310043L19Rik | bXPF_tRA |
| 1-1418 | chr1 | 177708485 | 177708948 | + | 759 | -21098 | 1700016C15Rik | bXPF_tRA |
| 1-666 | chr1 | 177708684 | 177709147 | + | 759 | -20899 | 1700016C15Rik | bXPF_tRA |
| 1-1435 | chr1 | 180387448 | 180387911 | + | 701 | -57130 | Gm5069 | bXPF_tRA |
| 1-178 | chr1 | 180387648 | 180388111 | + | 700 | -57330 | Gm5069 | bXPF_tRA |
| 1-1894 | chr1 | 180850708 | 180851171 | + | 942 | -212 | Sde2 | bXPF_tRA |
| 1-1669 | chr1 | 182124834 | 182125297 | + | 770 | 328 | Srp9 | bXPF_tRA |
| 1-1078 | chr1 | 182908307 | 182908770 | + | 697 | -46250 | Tlr5 | bXPF_tRA |
| 1-770 | chr1 | 186749050 | 186749513 | + | 555 | 77 | Rrp15 | bXPF_tRA |
| 1-1035 | chr1 | 187487908 | 187488371 | + | 564 | -120867 | Esrrg | bXPF_tRA |
| 1-1167 | chr1 | 191025944 | 191026407 | + | 643 | 15 | Mfsd7b | bXPF_tRA |
| 1-102 | chr1 | 191026143 | 191026606 | + | 642 | -184 | Mfsd7b | bXPF_tRA |
| 1-1834 | chr1 | 191062728 | 191063191 | + | 929 | -27 | Tatdn3 | bXPF_tRA |
| 1-735 | chr1 | 191317876 | 191318339 | + | 562 | 11 | Nenf | bXPF_tRA |
| 1-1790 | chr1 | 191575799 | 191576262 | + | 708 | 394 | Ints7 | bXPF_tRA |
| 1-1018 | chr1 | 192855795 | 192856258 | + | 584 | -274 | Sertad4 | bXPF_tRA |
| 1-1048 | chr10 | 3573086 | 3573549 | + | 615 | 33038 | Iyd | bXPF_tRA |
| 1-719 | chr10 | 3740010 | 3740473 | + | 543 | -136 | Plekhg1 | bXPF_tRA |
| 1-961 | chr10 | 4775985 | 4776448 | + | 660 | 66058 | Esr1 | bXPF_tRA |
| 1-360 | chr10 | 5823992 | 5824455 | + | 638 | -280 | Mtrf1l | bXPF_tRA |

| | | | | | | | | |
|---|---|---|---|---|---|---|---|---|
| 16803 | chr10 | 5824208 | 5824671 | + | 638 | -496 | Mtrf1l | bXPF_tRA |
| 1-599 | chr10 | 12922391 | 12922854 | + | 729 | 505 | B230208H11Rik | bXPF_tRA |
| 1-1762 | chr10 | 13552908 | 13553371 | + | 938 | 3 | Pex3 | bXPF_tRA |
| 1-1211 | chr10 | 16990285 | 16990748 | + | 608 | 105266 | Gm20125 | bXPF_tRA |
| 1-1300 | chr10 | 17012326 | 17012789 | + | 819 | 83225 | Gm20125 | bXPF_tRA |
| 1-544 | chr10 | 17724425 | 17724888 | + | 606 | 1428 | Cited2 | bXPF_tRA |
| 1-1344 | chr10 | 19203846 | 19204309 | + | 612 | 36839 | Gm20139 | bXPF_tRA |
| 43040 | chr10 | 19204046 | 19204509 | + | 612 | 37039 | Gm20139 | bXPF_tRA |
| 1-1246 | chr10 | 24765326 | 24765789 | + | 665 | -53398 | Enpp1 | bXPF_tRA |
| 12785 | chr10 | 24869764 | 24870227 | + | 654 | 9 | Med23 | bXPF_tRA |
| 1-509 | chr10 | 25271703 | 25272166 | + | 549 | 27229 | Akap7 | bXPF_tRA |
| 1-179 | chr10 | 25271905 | 25272368 | + | 549 | 27027 | Akap7 | bXPF_tRA |
| 1-747 | chr10 | 25296893 | 25297356 | + | 548 | 2039 | Akap7 | bXPF_tRA |
| 1-1337 | chr10 | 25520417 | 25520880 | + | 796 | 15624 | Smlr1 | bXPF_tRA |
| 1-601 | chr10 | 26892909 | 26893372 | + | 668 | 120628 | Arhgap18 | bXPF_tRA |
| 1-451 | chr10 | 26893124 | 26893587 | + | 668 | 120843 | Arhgap18 | bXPF_tRA |
| 1-946 | chr10 | 34206963 | 34207426 | + | 565 | 357 | Dse | bXPF_tRA |
| 1-835 | chr10 | 34418173 | 34418636 | + | 591 | 147 | Nt5dc1 | bXPF_tRA |
| 1-276 | chr10 | 37136100 | 37136563 | + | 643 | 2595 | Marcks | bXPF_tRA |
| 1-300 | chr10 | 37136380 | 37136843 | + | 686 | 2315 | Marcks | bXPF_tRA |
| 1-262 | chr10 | 37291722 | 37292185 | + | 681 | -153027 | Marcks | bXPF_tRA |
| 1-1667 | chr10 | 37495370 | 37495833 | + | 594 | -356675 | Marcks | bXPF_tRA |
| 1-1410 | chr10 | 38908183 | 38908646 | + | 854 | -57101 | Lama4 | bXPF_tRA |
| 1-498 | chr10 | 38973050 | 38973513 | + | 629 | 7766 | Lama4 | bXPF_tRA |
| 19360 | chr10 | 38973293 | 38973756 | + | 629 | 8009 | Lama4 | bXPF_tRA |
| 1-1202 | chr10 | 39659271 | 39659734 | + | 718 | 46568 | Traf3ip2 | bXPF_tRA |
| 1-273 | chr10 | 39659472 | 39659935 | + | 718 | 46769 | Traf3ip2 | bXPF_tRA |
| 1-614 | chr10 | 41292925 | 41293388 | + | 635 | 10085 | Fig4 | bXPF_tRA |
| 1-351 | chr10 | 41293124 | 41293587 | + | 635 | 9886 | Fig4 | bXPF_tRA |
| 1-1297 | chr10 | 42277835 | 42278298 | + | 738 | -1324 | Foxo3 | bXPF_tRA |
| 1-609 | chr10 | 43578504 | 43578967 | + | 604 | -434 | Cd24a | bXPF_tRA |
| 1-1774 | chr10 | 52417208 | 52417671 | + | 707 | -108 | Nus1 | bXPF_tRA |
| 1-971 | chr10 | 54075588 | 54076051 | + | 657 | -23 | Man1a | bXPF_tRA |
| 1-365 | chr10 | 56921002 | 56921465 | + | 553 | 543933 | Gja1 | bXPF_tRA |
| 1-204 | chr10 | 56921258 | 56921721 | + | 546 | 544189 | Gja1 | bXPF_tRA |
| 1-1464 | chr10 | 59951564 | 59952027 | + | 711 | -25 | Ddit4 | bXPF_tRA |
| 1-683 | chr10 | 61147258 | 61147721 | + | 661 | 195 | Sgpl1 | bXPF_tRA |
| 1-1527 | chr10 | 61782700 | 61783163 | + | 681 | 933 | H2afy2 | bXPF_tRA |
| 1-1727 | chr10 | 62650957 | 62651420 | + | 621 | 10 | Ddx50 | bXPF_tRA |
| 1-410 | chr10 | 62651198 | 62651661 | + | 617 | -231 | Ddx50 | bXPF_tRA |
| 1-542 | chr10 | 63023773 | 63024236 | + | 840 | -155 | Hnrnph3 | bXPF_tRA |
| 1-780 | chr10 | 63023983 | 63024446 | + | 841 | -298 | Pbld2 | bXPF_tRA |
| 1-798 | chr10 | 63429322 | 63429785 | + | 698 | -545 | Ctnna3 | bXPF_tRA |
| 1-1261 | chr10 | 64343061 | 64343524 | + | 594 | -253037 | Lrrtm3 | bXPF_tRA |
| 1-120 | chr10 | 67002880 | 67003343 | + | 734 | -1964 | Gm31763 | bXPF_tRA |
| 1-1650 | chr10 | 68136404 | 68136867 | + | 750 | 142091 | Arid5b | bXPF_tRA |
| 1-1827 | chr10 | 68354742 | 68355205 | + | 716 | -33831 | 4930545H06Rik | bXPF_tRA |
| 1-762 | chr10 | 68558413 | 68558876 | + | 570 | -16769 | 1700040L02Rik | bXPF_tRA |
| 1-1763 | chr10 | 75517687 | 75518150 | + | 836 | 54 | Gucd1 | bXPF_tRA |
| 1-634 | chr10 | 76903080 | 76903543 | + | 548 | 58636 | Pcbp3 | bXPF_tRA |
| 1-393 | chr10 | 80167634 | 80168097 | + | 584 | 24 | Cirbp | bXPF_tRA |
| 19725 | chr10 | 80249200 | 80249663 | + | 712 | -21 | Ndufs7 | bXPF_tRA |
| 1-1139 | chr10 | 80433699 | 80434162 | + | 690 | -277 | Tcf3 | bXPF_tRA |
| 1-479 | chr10 | 80701775 | 80702238 | + | 600 | -186 | Mob3a | bXPF_tRA |
| 1-1729 | chr10 | 80751415 | 80751878 | + | 739 | -3560 | Dot1l | bXPF_tRA |
| 1-364 | chr10 | 80751614 | 80752077 | + | 603 | -3361 | Dot1l | bXPF_tRA |
| 1-1131 | chr10 | 80774624 | 80775087 | + | 578 | 19649 | Dot1l | bXPF_tRA |
| 1-1034 | chr10 | 80855088 | 80855551 | + | 644 | 44 | Sppl2b | bXPF_tRA |
| 1-1278 | chr10 | 81037226 | 81037689 | + | 674 | -3545 | Slc39a3 | bXPF_tRA |
| 1-1895 | chr10 | 81285697 | 81286160 | + | 940 | -147 | Tjp3 | bXPF_tRA |
| 1-488 | chr10 | 81292505 | 81292968 | + | 575 | -227 | Pip5k1c | bXPF_tRA |
| 35796 | chr10 | 81292852 | 81293315 | + | 575 | 120 | Pip5k1c | bXPF_tRA |
| 1-1772 | chr10 | 82178142 | 82178605 | + | 608 | 50360 | AU041133 | bXPF_tRA |

| | | | | | | | | |
|---|---|---|---|---|---|---|---|---|
| 1-740 | chr10 | 82763823 | 82764286 | + | 639 | 87 | Nfyb | bXPF_tRA |
| 1-457 | chr10 | 82764026 | 82764489 | + | 639 | -116 | Nfyb | bXPF_tRA |
| 1-1662 | chr10 | 83648400 | 83648863 | + | 557 | 33 | Appl2 | bXPF_tRA |
| 1-227 | chr10 | 83882088 | 83882551 | + | 696 | 159454 | 1500009L16Rik | bXPF_tRA |
| 42856 | chr10 | 83882356 | 83882819 | + | 696 | 159722 | 1500009L16Rik | bXPF_tRA |
| 1-790 | chr10 | 84388452 | 84388915 | + | 672 | 51788 | Nuak1 | bXPF_tRA |
| 1-275 | chr10 | 84388712 | 84389175 | + | 671 | 51528 | Nuak1 | bXPF_tRA |
| 1-1355 | chr10 | 84622132 | 84622595 | + | 737 | -74 | Polr3b | bXPF_tRA |
| 1-1040 | chr10 | 85916939 | 85917402 | + | 657 | -184 | Prdm4 | bXPF_tRA |
| 1-627 | chr10 | 86705126 | 86705589 | + | 565 | 87 | Hsp90b1 | bXPF_tRA |
| 1-1398 | chr10 | 91181799 | 91182262 | + | 661 | -10411 | Tmpo | bXPF_tRA |
| 1-511 | chr10 | 94147750 | 94148213 | + | 556 | 50 | Nr2c1 | bXPF_tRA |
| 1-285 | chr10 | 94147982 | 94148445 | + | 556 | 282 | Nr2c1 | bXPF_tRA |
| 1-706 | chr10 | 94824327 | 94824790 | + | 676 | 120020 | Plxnc1 | bXPF_tRA |
| 1-454 | chr10 | 94943609 | 94944072 | + | 587 | 738 | Plxnc1 | bXPF_tRA |
| 1-409 | chr10 | 94943828 | 94944291 | + | 587 | 519 | Plxnc1 | bXPF_tRA |
| 1-1692 | chr10 | 96213897 | 96214360 | + | 833 | -12079 | 4930459C07Rik | bXPF_tRA |
| 1-1787 | chr10 | 96616613 | 96617076 | + | 781 | -157 | Btg1 | bXPF_tRA |
| 11324 | chr10 | 96616815 | 96617278 | + | 569 | 45 | Btg1 | bXPF_tRA |
| 1-989 | chr10 | 100645653 | 100646116 | + | 688 | 53498 | 1700017N19Rik | bXPF_tRA |
| 1-1823 | chr10 | 103650089 | 103650552 | + | 1000 | 282512 | Slc6a15 | bXPF_tRA |
| 1-800 | chr10 | 107171878 | 107172341 | + | 587 | -48445 | Acss3 | bXPF_tRA |
| 1-1679 | chr10 | 111389831 | 111390294 | + | 779 | -83130 | Nap1l1 | bXPF_tRA |
| 1-1092 | chr10 | 111506298 | 111506761 | + | 544 | 243 | Phlda1 | bXPF_tRA |
| 1-658 | chr10 | 111506501 | 111506964 | + | 544 | 446 | Phlda1 | bXPF_tRA |
| 1-1874 | chr10 | 111972530 | 111972993 | + | 877 | 66 | Krr1 | bXPF_tRA |
| 1-210 | chr10 | 111972735 | 111973198 | + | 662 | 271 | Krr1 | bXPF_tRA |
| 1-559 | chr10 | 117846111 | 117846574 | + | 553 | -368 | Rap1b | bXPF_tRA |
| 1-756 | chr10 | 119615253 | 119615716 | + | 617 | 146215 | Grip1os2 | bXPF_tRA |
| 42917 | chr10 | 120476160 | 120476623 | + | 590 | 78 | Hmga2 | bXPF_tRA |
| 46023 | chr10 | 122096542 | 122097005 | + | 689 | 329 | Tmem5 | bXPF_tRA |
| 1-1616 | chr10 | 122912964 | 122913427 | + | 755 | 72529 | Mirlet7i | bXPF_tRA |
| 1-933 | chr10 | 127063329 | 127063792 | + | 568 | -43 | Cdk4 | bXPF_tRA |
| 1-482 | chr10 | 127290395 | 127290858 | + | 558 | -167 | Ddit3 | bXPF_tRA |
| 1-187 | chr10 | 127290657 | 127291120 | + | 557 | 95 | Ddit3 | bXPF_tRA |
| 1-1845 | chr10 | 128096524 | 128096987 | + | 823 | 3972 | Baz2a | bXPF_tRA |
| 1-225 | chr10 | 128096740 | 128097203 | + | 591 | 4188 | Baz2a | bXPF_tRA |
| 1-1143 | chr10 | 128411456 | 128411919 | + | 581 | 71 | Rnf41 | bXPF_tRA |
| 43466 | chr10 | 128411668 | 128412131 | + | 581 | 283 | Rnf41 | bXPF_tRA |
| 1-1428 | chr10 | 128458341 | 128458804 | + | 674 | -108 | Mir8105 | bXPF_tRA |
| 1-1168 | chr10 | 128909446 | 128909909 | + | 597 | -189 | Cd63 | bXPF_tRA |
| 1-1266 | chr10 | 128922978 | 128923441 | + | 657 | 315 | Bloc1s1 | bXPF_tRA |
| 30317 | chr10 | 128923179 | 128923642 | + | 657 | 114 | Bloc1s1 | bXPF_tRA |
| 1-1600 | chr11 | 3266301 | 3266764 | + | 745 | -146 | Drg1 | bXPF_tRA |
| 1-1488 | chr11 | 3963559 | 3964022 | + | 646 | -185 | Pes1 | bXPF_tRA |
| 1-620 | chr11 | 5788379 | 5788842 | + | 550 | 127 | Dbnl | bXPF_tRA |
| 12420 | chr11 | 5788580 | 5789043 | + | 550 | 328 | Dbnl | bXPF_tRA |
| 1-1404 | chr11 | 5878193 | 5878656 | + | 616 | -168 | Pold2 | bXPF_tRA |
| 1-1531 | chr11 | 6200014 | 6200477 | + | 605 | 206 | Nudcd3 | bXPF_tRA |
| 1-1853 | chr11 | 6267593 | 6268056 | + | 915 | -95 | Ddx56 | bXPF_tRA |
| 1-327 | chr11 | 16546534 | 16546997 | + | 582 | -38281 | Sec61g | bXPF_tRA |
| 1-268 | chr11 | 16546744 | 16547207 | + | 583 | -38491 | Sec61g | bXPF_tRA |
| 1-757 | chr11 | 16591401 | 16591864 | + | 702 | -83148 | Sec61g | bXPF_tRA |
| 1-570 | chr11 | 17083887 | 17084350 | + | 580 | 32184 | Cnrip1 | bXPF_tRA |
| 1-912 | chr11 | 17386719 | 17387182 | + | 635 | 129262 | C1d | bXPF_tRA |
| 1-731 | chr11 | 19920217 | 19920680 | + | 597 | -3994 | Spred2 | bXPF_tRA |
| 1-1097 | chr11 | 19923982 | 19924445 | + | 778 | -229 | Spred2 | bXPF_tRA |
| 1-911 | chr11 | 20112777 | 20113240 | + | 576 | -57 | Actr2 | bXPF_tRA |
| 1-1331 | chr11 | 20859076 | 20859539 | + | 579 | -28199 | Lgalsl | bXPF_tRA |
| 1-838 | chr11 | 21091145 | 21091608 | + | 865 | 52 | Peli1 | bXPF_tRA |
| 20090 | chr11 | 21091363 | 21091826 | + | 866 | 270 | Peli1 | bXPF_tRA |
| 1-1050 | chr11 | 22990582 | 22991045 | + | 637 | 220 | Cct4 | bXPF_tRA |
| 1-110 | chr11 | 22990784 | 22991247 | + | 637 | 422 | Cct4 | bXPF_tRA |

| | | | | | | | | |
|---|---|---|---|---|---|---|---|---|
| 1-603 | chr11 | 23257180 | 23257643 | + | 555 | 1370 | Xpo1 | bXPF_tRA |
| 1-858 | chr11 | 23666090 | 23666553 | + | 685 | 345 | Pus10 | bXPF_tRA |
| 1-112 | chr11 | 23666295 | 23666758 | + | 685 | 550 | Pus10 | bXPF_tRA |
| 1-1051 | chr11 | 30649028 | 30649491 | + | 592 | 137 | Acyp2 | bXPF_tRA |
| 1-826 | chr11 | 32435698 | 32436161 | + | 631 | -19443 | Ubtd2 | bXPF_tRA |
| 1-1326 | chr11 | 33660372 | 33660835 | + | 542 | -81646 | Gabrp | bXPF_tRA |
| 1-1779 | chr11 | 33861059 | 33861522 | + | 893 | -17705 | Kcnip1 | bXPF_tRA |
| 1-1744 | chr11 | 35065828 | 35066291 | + | 989 | -55397 | Slit3 | bXPF_tRA |
| 1-391 | chr11 | 35066035 | 35066498 | + | 991 | -55190 | Slit3 | bXPF_tRA |
| 1-308 | chr11 | 35769228 | 35769691 | + | 560 | -36 | Pank3 | bXPF_tRA |
| 1-758 | chr11 | 40694585 | 40695048 | + | 611 | 387 | Mat2b | bXPF_tRA |
| 1-1039 | chr11 | 44616940 | 44617403 | + | 624 | -929 | Ebf1 | bXPF_tRA |
| 1-1262 | chr11 | 45296759 | 45297222 | + | 682 | -180062 | Gm12159 | bXPF_tRA |
| 1-879 | chr11 | 45926653 | 45927116 | + | 584 | 18051 | Lsm11 | bXPF_tRA |
| 1-1542 | chr11 | 45955261 | 45955724 | + | 702 | 11 | Thg1l | bXPF_tRA |
| 1-1516 | chr11 | 48817124 | 48817587 | + | 653 | 36 | Trim41 | bXPF_tRA |
| 1-1587 | chr11 | 48818304 | 48818767 | + | 797 | -1144 | Trim41 | bXPF_tRA |
| 1-191 | chr11 | 48818526 | 48818989 | + | 560 | -1366 | Trim41 | bXPF_tRA |
| 1-1695 | chr11 | 48853439 | 48853902 | + | 735 | 15665 | Trim7 | bXPF_tRA |
| 23377 | chr11 | 48853654 | 48854117 | + | 580 | 15880 | Trim7 | bXPF_tRA |
| 1-1611 | chr11 | 48856094 | 48856557 | + | 594 | 15021 | Irgm1 | bXPF_tRA |
| 1-1193 | chr11 | 48856385 | 48856848 | + | 595 | 14730 | Irgm1 | bXPF_tRA |
| 1-939 | chr11 | 48871191 | 48871654 | + | 582 | -76 | Irgm1 | bXPF_tRA |
| 1-1460 | chr11 | 49250412 | 49250875 | + | 608 | 91 | Mgat1 | bXPF_tRA |
| 1-1376 | chr11 | 50174598 | 50175061 | + | 705 | -22 | Mrnip | bXPF_tRA |
| 1-1380 | chr11 | 50292202 | 50292665 | + | 564 | -97 | Maml1 | bXPF_tRA |
| 1-437 | chr11 | 51605850 | 51606313 | + | 633 | 800 | Hnrnpab | bXPF_tRA |
| 1-1130 | chr11 | 51688656 | 51689119 | + | 614 | -253 | 0610009B22Rik | bXPF_tRA |
| 1-941 | chr11 | 51967411 | 51967874 | + | 727 | 11 | Cdkn2aipnl | bXPF_tRA |
| 1-1627 | chr11 | 54060226 | 54060689 | + | 636 | 8560 | Pdlim4 | bXPF_tRA |
| 1-404 | chr11 | 55469080 | 55469543 | + | 605 | -441 | G3bp1 | bXPF_tRA |
| 27760 | chr11 | 55469393 | 55469856 | + | 605 | -128 | G3bp1 | bXPF_tRA |
| 1-726 | chr11 | 57719670 | 57720133 | + | 752 | 74459 | Galnt10 | bXPF_tRA |
| 1-194 | chr11 | 57719869 | 57720332 | + | 752 | 74658 | Galnt10 | bXPF_tRA |
| 1-1595 | chr11 | 58168217 | 58168680 | + | 671 | 91 | Gemin5 | bXPF_tRA |
| 1-1205 | chr11 | 58302309 | 58302772 | + | 578 | -4529 | Zfp692 | bXPF_tRA |
| 1-1106 | chr11 | 58954847 | 58955310 | + | 699 | 393 | Hist3h2a | bXPF_tRA |
| 1-1151 | chr11 | 59458935 | 59459398 | + | 763 | 9121 | Jmjd4 | bXPF_tRA |
| 1-202 | chr11 | 59459153 | 59459616 | + | 762 | 9339 | Jmjd4 | bXPF_tRA |
| 1-859 | chr11 | 60046175 | 60046638 | + | 583 | 83 | Pemt | bXPF_tRA |
| 1-1019 | chr11 | 60417025 | 60417488 | + | 596 | 111 | Gid4 | bXPF_tRA |
| 17899 | chr11 | 60454291 | 60454754 | + | 1000 | -95 | Drg2 | bXPF_tRA |
| 1-1854 | chr11 | 61407701 | 61408164 | + | 851 | -29857 | Slc47a1 | bXPF_tRA |
| 1-1174 | chr11 | 61930215 | 61930678 | + | 846 | -189 | Akap10 | bXPF_tRA |
| 44562 | chr11 | 62281455 | 62281918 | + | 547 | 213 | Ttc19 | bXPF_tRA |
| 1-678 | chr11 | 62551183 | 62551646 | + | 667 | -90 | Gm1821 | bXPF_tRA |
| 1-190 | chr11 | 62551394 | 62551857 | + | 667 | -106 | Ubb | bXPF_tRA |
| 1-1212 | chr11 | 62602422 | 62602885 | + | 601 | -224 | 2410006H16Rik | bXPF_tRA |
| 1-691 | chr11 | 62756772 | 62757235 | + | 567 | -27910 | Zfp287 | bXPF_tRA |
| 1-1743 | chr11 | 62921077 | 62921540 | + | 926 | -29885 | Cdrt4 | bXPF_tRA |
| 1-1356 | chr11 | 67515546 | 67516009 | + | 576 | -17221 | Gas7 | bXPF_tRA |
| 1-1560 | chr11 | 69040117 | 69040580 | + | 780 | -5289 | Aurkb | bXPF_tRA |
| 1-630 | chr11 | 69040837 | 69041300 | + | 582 | -4569 | Aurkb | bXPF_tRA |
| 1-1496 | chr11 | 69062751 | 69063214 | + | 888 | 3207 | Borcs6 | bXPF_tRA |
| 1-1882 | chr11 | 69073309 | 69073772 | + | 908 | 2731 | Tmem107 | bXPF_tRA |
| 1-115 | chr11 | 69073509 | 69073972 | + | 906 | 2931 | Tmem107 | bXPF_tRA |
| 1-1565 | chr11 | 69118616 | 69119079 | + | 793 | -1606 | Hes7 | bXPF_tRA |
| 1-1284 | chr11 | 69124231 | 69124694 | + | 859 | -1915 | Aloxe3 | bXPF_tRA |
| 1-1133 | chr11 | 69125028 | 69125491 | + | 563 | -1118 | Aloxe3 | bXPF_tRA |
| 1-1163 | chr11 | 69332118 | 69332581 | + | 650 | 6091 | Kcnab3 | bXPF_tRA |
| 1-1222 | chr11 | 69417415 | 69417878 | + | 666 | -3971 | Kdm6b | bXPF_tRA |
| 1-1671 | chr11 | 69838207 | 69838670 | + | 895 | -87 | Tmem256 | bXPF_tRA |
| 1-567 | chr11 | 70982616 | 70983079 | + | 580 | 179 | C1qbp | bXPF_tRA |

| | | | | | | | | |
|---|---|---|---|---|---|---|---|---|
| 1-789 | chr11 | 72689820 | 72690283 | + | 624 | 49 | Ankfy1 | bXPF_tRA |
| 1-148 | chr11 | 72690034 | 72690497 | + | 624 | 263 | Ankfy1 | bXPF_tRA |
| 1-1647 | chr11 | 74649127 | 74649590 | + | 852 | -137 | Cluh | bXPF_tRA |
| 1-1066 | chr11 | 75012932 | 75013395 | + | 630 | 52828 | Mir684-1 | bXPF_tRA |
| 1-240 | chr11 | 75013137 | 75013600 | + | 630 | 52623 | Mir684-1 | bXPF_tRA |
| 1-850 | chr11 | 75170172 | 75170635 | + | 722 | -148 | Hic1 | bXPF_tRA |
| 47119 | chr11 | 75170386 | 75170849 | + | 722 | -362 | Hic1 | bXPF_tRA |
| 1-1580 | chr11 | 75454519 | 75454982 | + | 781 | -33 | Wdr81 | bXPF_tRA |
| 1-1389 | chr11 | 75630580 | 75631043 | + | 736 | -209 | Inpp5k | bXPF_tRA |
| 1-1123 | chr11 | 76292278 | 76292741 | + | 581 | 48773 | Rnmtl1 | bXPF_tRA |
| 1-1135 | chr11 | 76763220 | 76763683 | + | 613 | 104 | Gosr1 | bXPF_tRA |
| 1-1309 | chr11 | 77685850 | 77686313 | + | 601 | -58 | Nufip2 | bXPF_tRA |
| 1-794 | chr11 | 78751366 | 78751829 | + | 561 | 132 | Fam58b | bXPF_tRA |
| 1-881 | chr11 | 79213991 | 79214454 | + | 560 | 28802 | Wsb1 | bXPF_tRA |
| 1-1339 | chr11 | 79805986 | 79806449 | + | 678 | 79817 | Mir365-2 | bXPF_tRA |
| 1-1379 | chr11 | 80080898 | 80081361 | + | 723 | -138 | Crlf3 | bXPF_tRA |
| 1-1664 | chr11 | 80141965 | 80142428 | + | 714 | -43 | Tefm | bXPF_tRA |
| 1-1622 | chr11 | 80377811 | 80378274 | + | 730 | -27 | 5730455P16Rik | bXPF_tRA |
| 1-728 | chr11 | 83298765 | 83299228 | + | 553 | -19 | Pex12 | bXPF_tRA |
| 1-1083 | chr11 | 84819367 | 84819830 | + | 598 | -83 | Mrm1 | bXPF_tRA |
| 1-623 | chr11 | 85430833 | 85431296 | + | 550 | 77900 | Bcas3 | bXPF_tRA |
| 1-1109 | chr11 | 85925348 | 85925811 | + | 599 | 28814 | Tbx4 | bXPF_tRA |
| 1-342 | chr11 | 86580091 | 86580554 | + | 628 | 3836 | Mir21a | bXPF_tRA |
| 1-175 | chr11 | 86580352 | 86580815 | + | 628 | 3575 | Mir21a | bXPF_tRA |
| 1-315 | chr11 | 86580720 | 86581183 | + | 778 | 3207 | Mir21a | bXPF_tRA |
| 1-420 | chr11 | 86581022 | 86581485 | + | 555 | 2905 | Mir21a | bXPF_tRA |
| 1-967 | chr11 | 86603659 | 86604122 | + | 545 | -19732 | Mir21a | bXPF_tRA |
| 1-1240 | chr11 | 86650620 | 86651083 | + | 659 | -5220 | Mir8115 | bXPF_tRA |
| 1-1103 | chr11 | 87086692 | 87087155 | + | 590 | -146 | Smg8 | bXPF_tRA |
| 1-1521 | chr11 | 89186763 | 89187226 | + | 628 | 113153 | Gm525 | bXPF_tRA |
| 1-1802 | chr11 | 93968101 | 93968564 | + | 643 | 189 | Nme1 | bXPF_tRA |
| 1-823 | chr11 | 94276903 | 94277366 | + | 556 | -34555 | Wfikkn2 | bXPF_tRA |
| 1-1230 | chr11 | 94327628 | 94328091 | + | 688 | -142 | Ankrd40 | bXPF_tRA |
| 1-1871 | chr11 | 94548903 | 94549366 | + | 836 | 73 | Rsad1 | bXPF_tRA |
| 1-195 | chr11 | 94549102 | 94549565 | + | 835 | -126 | Rsad1 | bXPF_tRA |
| 1-514 | chr11 | 94880856 | 94881319 | + | 545 | -31494 | Gm11544 | bXPF_tRA |
| 1-333 | chr11 | 94881073 | 94881536 | + | 545 | -31711 | Gm11544 | bXPF_tRA |
| 1-564 | chr11 | 94948367 | 94948830 | + | 661 | 12374 | Col1a1 | bXPF_tRA |
| 1-1469 | chr11 | 94989573 | 94990036 | + | 711 | -1408 | Ppp1r9b | bXPF_tRA |
| 1-1137 | chr11 | 95413618 | 95414081 | + | 627 | -234 | Spop | bXPF_tRA |
| 1-1586 | chr11 | 96065206 | 96065669 | + | 548 | -73 | Ube2z | bXPF_tRA |
| 1-720 | chr11 | 96747282 | 96747745 | + | 569 | 30042 | Snx11 | bXPF_tRA |
| 1-968 | chr11 | 96977446 | 96977909 | + | 579 | 11 | Sp2 | bXPF_tRA |
| 1-739 | chr11 | 97187779 | 97188242 | + | 616 | -118 | Kpnb1 | bXPF_tRA |
| 1-1220 | chr11 | 97361942 | 97362405 | + | 543 | -378 | Socs7 | bXPF_tRA |
| 1-1510 | chr11 | 97988033 | 97988496 | + | 734 | -1818 | Plxdc1 | bXPF_tRA |
| 1-742 | chr11 | 100738068 | 100738531 | + | 578 | -84 | Rab5c | bXPF_tRA |
| 1-277 | chr11 | 100738267 | 100738730 | + | 579 | -283 | Rab5c | bXPF_tRA |
| 1-1274 | chr11 | 100939484 | 100939947 | + | 868 | -175 | Stat3 | bXPF_tRA |
| 1-936 | chr11 | 101442056 | 101442519 | + | 585 | 42 | Rpl27 | bXPF_tRA |
| 20455 | chr11 | 102156186 | 102156649 | + | 1000 | -10 | Tmem101 | bXPF_tRA |
| 1-1792 | chr11 | 102245015 | 102245478 | + | 832 | -3636 | BC030867 | bXPF_tRA |
| 1-232 | chr11 | 102245220 | 102245683 | + | 588 | -3431 | BC030867 | bXPF_tRA |
| 1-466 | chr11 | 102437476 | 102437939 | + | 566 | 7385 | Grn | bXPF_tRA |
| 1-560 | chr11 | 102466943 | 102467406 | + | 585 | 2709 | Itga2b | bXPF_tRA |
| 1-1425 | chr11 | 102556142 | 102556605 | + | 692 | -215 | Gpatch8 | bXPF_tRA |
| 1-837 | chr11 | 103003160 | 103003623 | + | 552 | -3470 | Mir6931 | bXPF_tRA |
| 1-952 | chr11 | 103051243 | 103051706 | + | 553 | 22912 | Nmt1 | bXPF_tRA |
| 1-1708 | chr11 | 106065873 | 106066336 | + | 824 | -3 | Taco1 | bXPF_tRA |
| 17899 | chr11 | 106066080 | 106066543 | + | 573 | 204 | Taco1 | bXPF_tRA |
| 1-1734 | chr11 | 106084131 | 106084594 | + | 730 | -540 | Map3k3 | bXPF_tRA |
| 12785 | chr11 | 106084331 | 106084794 | + | 729 | -340 | Map3k3 | bXPF_tRA |
| 1-1052 | chr11 | 106272688 | 106273151 | + | 632 | 53 | Smarcd2 | bXPF_tRA |

| | | | | | | | | |
|---|---|---|---|---|---|---|---|---|
| 1-1471 | chr11 | 106500748 | 106501211 | + | 667 | -12 | Snord104 | bXPF_tRA |
| 1-1080 | chr11 | 109439845 | 109440308 | + | 564 | -2 | LOC100503496 | bXPF_tRA |
| 1-474 | chr11 | 112473192 | 112473655 | + | 585 | 237933 | BC006965 | bXPF_tRA |
| 1-1077 | chr11 | 113565512 | 113565975 | + | 551 | 72 | Slc39a11 | bXPF_tRA |
| 1-1899 | chr11 | 115277023 | 115277486 | + | 932 | -285 | Fdxr | bXPF_tRA |
| 1-521 | chr11 | 115413546 | 115414009 | + | 551 | 6142 | Atp5h | bXPF_tRA |
| 1-1592 | chr11 | 116024042 | 116024505 | + | 695 | 231 | H3f3b | bXPF_tRA |
| 1-1105 | chr11 | 116852781 | 116853244 | + | 629 | 82 | Srsf2 | bXPF_tRA |
| 1-698 | chr11 | 118289923 | 118290386 | + | 726 | 90 | Usp36 | bXPF_tRA |
| 1-1761 | chr11 | 120237649 | 120238112 | + | 719 | 4933 | Bahcc1 | bXPF_tRA |
| 1-1589 | chr11 | 120796172 | 120796635 | + | 693 | -8 | Dus1l | bXPF_tRA |
| 1-1552 | chr11 | 120932683 | 120933146 | + | 561 | -42 | Ccdc57 | bXPF_tRA |
| 9498 | chr11 | 120990965 | 120991428 | + | 1000 | 137 | Csnk1d | bXPF_tRA |
| 1-1036 | chr11 | 121672868 | 121673331 | + | 550 | 52 | B3gntl1 | bXPF_tRA |
| 1-1551 | chr11 | 121702052 | 121702515 | + | 726 | -144 | Metrnl | bXPF_tRA |
| 1-1374 | chr12 | 4558586 | 4559049 | + | 633 | -34191 | Itsn2 | bXPF_tRA |
| 1-1547 | chr12 | 4907091 | 4907554 | + | 603 | 198 | Ubxn2a | bXPF_tRA |
| 1-1707 | chr12 | 4917096 | 4917559 | + | 798 | -26 | Atad2b | bXPF_tRA |
| 1-1740 | chr12 | 8313243 | 8313706 | + | 745 | 41 | Hs1bp3 | bXPF_tRA |
| 1-423 | chr12 | 8313455 | 8313918 | + | 594 | 253 | Hs1bp3 | bXPF_tRA |
| 1-801 | chr12 | 9304176 | 9304639 | + | 549 | 266178 | 1700022H16Rik | bXPF_tRA |
| 1-1651 | chr12 | 10390700 | 10391163 | + | 609 | 151 | Rdh14 | bXPF_tRA |
| 1-338 | chr12 | 32379348 | 32379811 | + | 564 | 790 | Ccdc71l | bXPF_tRA |
| 1-1532 | chr12 | 32546332 | 32546795 | + | 766 | 167774 | Ccdc71l | bXPF_tRA |
| 1-1060 | chr12 | 32819805 | 32820268 | + | 552 | -299 | Nampt | bXPF_tRA |
| 1-1581 | chr12 | 34040043 | 34040506 | + | 652 | 82603 | Twist1 | bXPF_tRA |
| 1-655 | chr12 | 52493413 | 52493876 | + | 623 | -13035 | Gm35135 | bXPF_tRA |
| 1-1535 | chr12 | 52503399 | 52503862 | + | 921 | -12447 | Arhgap5 | bXPF_tRA |
| 1-663 | chr12 | 55492168 | 55492631 | + | 561 | 248 | Nfkbia | bXPF_tRA |
| 1-1108 | chr12 | 57230205 | 57230668 | + | 586 | 11 | Mipol1 | bXPF_tRA |
| 1-557 | chr12 | 65172663 | 65173126 | + | 561 | -314 | Mis18bp1 | bXPF_tRA |
| 1-331 | chr12 | 69158772 | 69159235 | + | 545 | 183 | Rps29 | bXPF_tRA |
| 1-138 | chr12 | 69158998 | 69159461 | + | 545 | -43 | Rps29 | bXPF_tRA |
| 1-1024 | chr12 | 69183811 | 69184274 | + | 633 | 25 | Rpl36al | bXPF_tRA |
| 1-1816 | chr12 | 69296372 | 69296835 | + | 835 | -78 | Klhdc2 | bXPF_tRA |
| 1-818 | chr12 | 70794981 | 70795444 | + | 588 | -30302 | Frmd6 | bXPF_tRA |
| 1-1559 | chr12 | 76837221 | 76837684 | + | 610 | -15 | Fntb | bXPF_tRA |
| 1-411 | chr12 | 80109973 | 80110436 | + | 622 | 2809 | Zfp36l1 | bXPF_tRA |
| 1-959 | chr12 | 80230005 | 80230468 | + | 958 | 30135 | Actn1 | bXPF_tRA |
| 1-357 | chr12 | 80230214 | 80230677 | + | 957 | 29926 | Actn1 | bXPF_tRA |
| 1-679 | chr12 | 80365986 | 80366449 | + | 620 | 25460 | Mir1843a | bXPF_tRA |
| 1-1524 | chr12 | 80504677 | 80505140 | + | 661 | -2298 | 2310002D06Rik | bXPF_tRA |
| 1-197 | chr12 | 82257933 | 82258396 | + | 549 | -53185 | Sipa1l1 | bXPF_tRA |
| 1-1058 | chr12 | 83921836 | 83922299 | + | 641 | -133 | Numb | bXPF_tRA |
| 1-1234 | chr12 | 84772934 | 84773397 | + | 611 | -53 | Npc2 | bXPF_tRA |
| 1-377 | chr12 | 86801010 | 86801473 | + | 733 | 66872 | Lrrc74a | bXPF_tRA |
| 1-1002 | chr12 | 86824006 | 86824469 | + | 716 | 60577 | Irf2bpl | bXPF_tRA |
| 1-783 | chr12 | 86888058 | 86888521 | + | 542 | -3475 | Irf2bpl | bXPF_tRA |
| 1-304 | chr12 | 86888286 | 86888749 | + | 542 | -3703 | Irf2bpl | bXPF_tRA |
| 1-1856 | chr12 | 86947030 | 86947493 | + | 795 | -82 | Cipc | bXPF_tRA |
| 1-1440 | chr12 | 94509735 | 94510198 | + | 611 | 582628 | Mir8099-2 | bXPF_tRA |
| 1-1411 | chr12 | 100199349 | 100199812 | + | 605 | 145 | Calm1 | bXPF_tRA |
| 1-547 | chr12 | 100724981 | 100725444 | + | 708 | -92 | Rps6ka5 | bXPF_tRA |
| 1-334 | chr12 | 100725192 | 100725655 | + | 708 | -303 | Rps6ka5 | bXPF_tRA |
| 1462 | chr12 | 104751977 | 104752440 | + | 802 | -256 | Dicer1 | bXPF_tRA |
| 1-1312 | chr12 | 105685065 | 105685528 | + | 559 | -55 | Atg2b | bXPF_tRA |
| 1-752 | chr12 | 107950659 | 107951122 | + | 591 | 52524 | Bcl11b | bXPF_tRA |
| 1-1795 | chr12 | 107952243 | 107952706 | + | 766 | 50940 | Bcl11b | bXPF_tRA |
| 1-1206 | chr12 | 108835876 | 108836339 | + | 655 | -231 | Slc25a29 | bXPF_tRA |
| 14246 | chr12 | 108894035 | 108894498 | + | 872 | -6 | Wdr25 | bXPF_tRA |
| 1-988 | chr12 | 110122845 | 110123308 | + | 632 | 34997 | 3110009F21Rik | bXPF_tRA |
| 1-1045 | chr12 | 110687968 | 110688431 | + | 684 | -5580 | 1700001K19Rik | bXPF_tRA |
| 1-1865 | chr12 | 110703080 | 110703543 | + | 951 | -6916 | Hsp90aa1 | bXPF_tRA |

| | | | | | | | | |
|---|---|---|---|---|---|---|---|---|
| 1-445 | chr12 | 110703288 | 110703751 | + | 585 | -7124 | Hsp90aa1 | bXPF_tRA |
| 1-843 | chr12 | 111537699 | 111538162 | + | 595 | -171 | Eif5 | bXPF_tRA |
| 1-935 | chr12 | 117843466 | 117843929 | + | 599 | -164 | Cdca7l | bXPF_tRA |
| 1-1504 | chr13 | 3893247 | 3893710 | + | 632 | 103 | Net1 | bXPF_tRA |
| 1-983 | chr13 | 5248746 | 5249209 | + | 578 | 609115 | 1700016G22Rik | bXPF_tRA |
| 1-1785 | chr13 | 15741735 | 15742198 | + | 815 | 60664 | 4933412O06Rik | bXPF_tRA |
| 1-694 | chr13 | 21161206 | 21161669 | + | 631 | -18382 | Olfr1368 | bXPF_tRA |
| 1-612 | chr13 | 21172278 | 21172741 | + | 609 | -7422 | Trim27 | bXPF_tRA |
| 1-596 | chr13 | 21233904 | 21234367 | + | 642 | 54204 | Trim27 | bXPF_tRA |
| 1-580 | chr13 | 21259604 | 21260067 | + | 557 | 32851 | Gpx5 | bXPF_tRA |
| 1-1472 | chr13 | 21279449 | 21279912 | + | 850 | 13006 | Gpx5 | bXPF_tRA |
| 1-1397 | chr13 | 21530808 | 21531271 | + | 628 | 75 | Zkscan8 | bXPF_tRA |
| 1-1448 | chr13 | 21881728 | 21882191 | + | 588 | 15090 | Mir1983 | bXPF_tRA |
| 1-1332 | chr13 | 21918982 | 21919445 | + | 656 | -22164 | Mir1983 | bXPF_tRA |
| 1-366 | chr13 | 21919185 | 21919648 | + | 656 | -22367 | Mir1983 | bXPF_tRA |
| 1-1748 | chr13 | 21940165 | 21940628 | + | 828 | -4698 | Zfp184 | bXPF_tRA |
| 23012 | chr13 | 21940789 | 21941252 | + | 603 | -4074 | Zfp184 | bXPF_tRA |
| 1-244 | chr13 | 21944848 | 21945311 | + | 541 | -15 | Zfp184 | bXPF_tRA |
| 1-1873 | chr13 | 21974824 | 21975287 | + | 890 | -6126 | Pom121l2 | bXPF_tRA |
| 1-1514 | chr13 | 22015591 | 22016054 | + | 606 | -6081 | Prss16 | bXPF_tRA |
| 1-1219 | chr13 | 22020964 | 22021427 | + | 616 | -11454 | Prss16 | bXPF_tRA |
| 1-954 | chr13 | 22029021 | 22029484 | + | 610 | 6391 | Hist1h2ah | bXPF_tRA |
| 1-1839 | chr13 | 23299933 | 23300396 | + | 663 | 13243 | 4930557F10Rik | bXPF_tRA |
| 11689 | chr13 | 23317257 | 23317720 | + | 843 | 3963 | 4930586N03Rik | bXPF_tRA |
| 1-1620 | chr13 | 23400869 | 23401332 | + | 549 | 22766 | Abt1 | bXPF_tRA |
| 1-1898 | chr13 | 23425405 | 23425868 | + | 983 | -1770 | Abt1 | bXPF_tRA |
| 1-577 | chr13 | 23426342 | 23426805 | + | 561 | -2707 | Abt1 | bXPF_tRA |
| 1-842 | chr13 | 23432502 | 23432965 | + | 581 | -1716 | C230035I16Rik | bXPF_tRA |
| 1-1673 | chr13 | 23436795 | 23437258 | + | 651 | -6009 | C230035I16Rik | bXPF_tRA |
| 1-625 | chr13 | 23437015 | 23437478 | + | 651 | -6229 | C230035I16Rik | bXPF_tRA |
| 1-348 | chr13 | 23445692 | 23446155 | + | 552 | -14906 | C230035I16Rik | bXPF_tRA |
| 1-354 | chr13 | 23446446 | 23446909 | + | 575 | -15660 | C230035I16Rik | bXPF_tRA |
| 1-716 | chr13 | 23450409 | 23450872 | + | 547 | 15261 | Btn1a1 | bXPF_tRA |
| 1-805 | chr13 | 23453367 | 23453830 | + | 596 | 12303 | Btn1a1 | bXPF_tRA |
| 16438 | chr13 | 23466830 | 23467293 | + | 994 | -1160 | Btn1a1 | bXPF_tRA |
| 1-1550 | chr13 | 23503772 | 23504235 | + | 769 | -15146 | Btn2a2 | bXPF_tRA |
| 17168 | chr13 | 25312134 | 25312597 | + | 1000 | -42369 | Nrsn1 | bXPF_tRA |
| 1-139 | chr13 | 25312366 | 25312829 | + | 1000 | -42601 | Nrsn1 | bXPF_tRA |
| 3654 | chr13 | 28417608 | 28418071 | + | 919 | 275355 | Prl5a1 | bXPF_tRA |
| 1-431 | chr13 | 28417812 | 28418275 | + | 917 | 275559 | Prl5a1 | bXPF_tRA |
| 1-1138 | chr13 | 32802410 | 32802873 | + | 704 | 611 | Wrnip1 | bXPF_tRA |
| 1-581 | chr13 | 33004257 | 33004720 | + | 546 | -53 | Serpinb9 | bXPF_tRA |
| 1-458 | chr13 | 34002302 | 34002765 | + | 614 | 261 | Serpinb6a | bXPF_tRA |
| 1-1519 | chr13 | 34726094 | 34726557 | + | 747 | -13317 | Fam50b | bXPF_tRA |
| 1-518 | chr13 | 37827551 | 37828014 | + | 560 | 389 | Rreb1 | bXPF_tRA |
| 1-270 | chr13 | 37827755 | 37828218 | + | 560 | 593 | Rreb1 | bXPF_tRA |
| 1-868 | chr13 | 38204731 | 38205194 | + | 603 | 23 | Snrnp48 | bXPF_tRA |
| 1-1515 | chr13 | 39306727 | 39307190 | + | 611 | -346083 | Slc35b3 | bXPF_tRA |
| 1-631 | chr13 | 42548984 | 42549447 | + | 568 | -131584 | Phactr1 | bXPF_tRA |
| 1-1353 | chr13 | 42575755 | 42576218 | + | 723 | -104813 | Phactr1 | bXPF_tRA |
| 1-1846 | chr13 | 43171244 | 43171707 | + | 747 | 26 | Tbc1d7 | bXPF_tRA |
| 1-172 | chr13 | 43171444 | 43171907 | + | 747 | -174 | Tbc1d7 | bXPF_tRA |
| 1-1887 | chr13 | 45389459 | 45389922 | + | 808 | -52 | Mylip | bXPF_tRA |
| 1-812 | chr13 | 45751222 | 45751685 | + | 624 | 213538 | Atxn1 | bXPF_tRA |
| 1-1145 | chr13 | 46930027 | 46930490 | + | 562 | -540 | Kif13a | bXPF_tRA |
| 1-797 | chr13 | 48227746 | 48228209 | + | 678 | -33450 | Id4 | bXPF_tRA |
| 1-1717 | chr13 | 50417723 | 50418186 | + | 684 | 77 | Fbxw17 | bXPF_tRA |
| 1-1277 | chr13 | 51847019 | 51847482 | + | 603 | 575 | Gadd45g | bXPF_tRA |
| 1-1474 | chr13 | 51968763 | 51969226 | + | 748 | 122319 | Gadd45g | bXPF_tRA |
| 1-893 | chr13 | 51975813 | 51976276 | + | 674 | -120697 | 4921525O09Rik | bXPF_tRA |
| 1-206 | chr13 | 51976014 | 51976477 | + | 675 | -120496 | 4921525O09Rik | bXPF_tRA |
| 1-930 | chr13 | 54693819 | 54694282 | + | 672 | -90 | Rnf44 | bXPF_tRA |
| 45292 | chr13 | 54694021 | 54694484 | + | 671 | -292 | Rnf44 | bXPF_tRA |

| | | | | | | | | |
|---|---|---|---|---|---|---|---|---|
| 1-1753 | chr13 | 54750681 | 54751144 | + | 594 | -1038 | Gprin1 | bXPF_tRA |
| 1-738 | chr13 | 55510719 | 55511182 | + | 647 | 2496 | Pdlim7 | bXPF_tRA |
| 27030 | chr13 | 55510922 | 55511385 | + | 647 | 2293 | Pdlim7 | bXPF_tRA |
| 1-540 | chr13 | 55536451 | 55536914 | + | 604 | -24 | Ddx41 | bXPF_tRA |
| 1-1886 | chr13 | 55599765 | 55600228 | + | 716 | 100 | B4galt7 | bXPF_tRA |
| 42948 | chr13 | 58128390 | 58128853 | + | 806 | -65 | Hnrnpa0 | bXPF_tRA |
| 1-234 | chr13 | 58128737 | 58129200 | + | 646 | -412 | Hnrnpa0 | bXPF_tRA |
| 41275 | chr13 | 58129125 | 58129588 | + | 645 | -800 | Hnrnpa0 | bXPF_tRA |
| 1-1688 | chr13 | 58402487 | 58402950 | + | 871 | 121 | Rmi1 | bXPF_tRA |
| 1-621 | chr13 | 59769740 | 59770203 | + | 600 | 5 | Etohd2 | bXPF_tRA |
| 1-1758 | chr13 | 60019814 | 60020277 | + | 764 | -9666 | A530065N20Rik | bXPF_tRA |
| 1-121 | chr13 | 60020014 | 60020477 | + | 548 | -9466 | A530065N20Rik | bXPF_tRA |
| 13516 | chr13 | 60176314 | 60176777 | + | 673 | 990 | Gas1 | bXPF_tRA |
| 1-917 | chr13 | 60177354 | 60177817 | + | 585 | -50 | Gas1 | bXPF_tRA |
| 15707 | chr13 | 60409788 | 60410251 | + | 1000 | -191928 | Dapk1 | bXPF_tRA |
| 1-824 | chr13 | 60427630 | 60428093 | + | 616 | -174086 | Dapk1 | bXPF_tRA |
| 1-1415 | chr13 | 63045553 | 63046016 | + | 774 | 80891 | 2010111I01Rik | bXPF_tRA |
| 1-845 | chr13 | 63258227 | 63258690 | + | 607 | 18429 | 2010111I01Rik | bXPF_tRA |
| 1-182 | chr13 | 63567899 | 63568362 | + | 686 | -2395 | Ptch1 | bXPF_tRA |
| 22647 | chr13 | 63568103 | 63568566 | + | 686 | -2599 | Ptch1 | bXPF_tRA |
| 1-515 | chr13 | 63814694 | 63815157 | + | 583 | -395 | Ercc6l2 | bXPF_tRA |
| 1-453 | chr13 | 65240847 | 65241310 | + | 549 | -35362 | Nlrp4f | bXPF_tRA |
| 1-699 | chr13 | 70637819 | 70638282 | + | 570 | -416 | Ice1 | bXPF_tRA |
| 1-1104 | chr13 | 71959271 | 71959734 | + | 559 | 4221 | Irx1 | bXPF_tRA |
| 1-1573 | chr13 | 72631506 | 72631969 | + | 619 | 2940 | Irx2 | bXPF_tRA |
| 1-1245 | chr13 | 72639514 | 72639977 | + | 653 | 10948 | Irx2 | bXPF_tRA |
| 1-1142 | chr13 | 73603706 | 73604169 | + | 552 | -65 | Clptm1l | bXPF_tRA |
| 1-1306 | chr13 | 74493759 | 74494222 | + | 639 | -40 | Zfp825 | bXPF_tRA |
| 1-309 | chr13 | 75707343 | 75707806 | + | 628 | 90 | Ell2 | bXPF_tRA |
| 17168 | chr13 | 75707546 | 75708009 | + | 628 | 293 | Ell2 | bXPF_tRA |
| 1-862 | chr13 | 76018393 | 76018856 | + | 577 | -18 | Rfesd | bXPF_tRA |
| 43831 | chr13 | 76018597 | 76019060 | + | 577 | -116 | Rfesd | bXPF_tRA |
| 1-1129 | chr13 | 80557133 | 80557596 | + | 645 | -326058 | Arrdc3 | bXPF_tRA |
| 1-519 | chr13 | 80883046 | 80883509 | + | 772 | -145 | Arrdc3 | bXPF_tRA |
| 1-150 | chr13 | 80883277 | 80883740 | + | 773 | 86 | Arrdc3 | bXPF_tRA |
| 1-1714 | chr13 | 80963001 | 80963464 | + | 721 | -842 | 9330111N05Rik | bXPF_tRA |
| 1-715 | chr13 | 90089491 | 90089954 | + | 576 | 55 | Tmem167 | bXPF_tRA |
| 20090 | chr13 | 91165017 | 91165480 | + | 1000 | 58739 | Atg10 | bXPF_tRA |
| 1-1017 | chr13 | 95618348 | 95618811 | + | 596 | -146 | F2r | bXPF_tRA |
| 1-759 | chr13 | 96670731 | 96671194 | + | 583 | -26 | Hmgcr | bXPF_tRA |
| 1-1085 | chr13 | 97137707 | 97138170 | + | 564 | 1 | Gfm2 | bXPF_tRA |
| 1-1889 | chr13 | 98890842 | 98891305 | + | 847 | -31 | Tnpo1 | bXPF_tRA |
| 1-1561 | chr13 | 99391397 | 99391860 | + | 734 | 21198 | 6430562O15Rik | bXPF_tRA |
| 1-1443 | chr13 | 102232479 | 102232942 | + | 549 | -460848 | Cd180 | bXPF_tRA |
| 1-1768 | chr13 | 104024906 | 104025369 | + | 768 | 84477 | Nln | bXPF_tRA |
| 1-166 | chr13 | 104025105 | 104025568 | + | 636 | 84278 | Nln | bXPF_tRA |
| 1-648 | chr13 | 107831162 | 107831625 | + | 692 | 58671 | Zswim6 | bXPF_tRA |
| 1-433 | chr13 | 107890408 | 107890871 | + | 588 | -575 | Zswim6 | bXPF_tRA |
| 1-1112 | chr13 | 108622586 | 108623049 | + | 598 | -31360 | Pde4d | bXPF_tRA |
| 1-1014 | chr13 | 110907540 | 110908003 | + | 546 | -3925 | 4930526H09Rik | bXPF_tRA |
| 1-512 | chr13 | 111489807 | 111490270 | + | 683 | 3 | Gpbp1 | bXPF_tRA |
| 1-1235 | chr13 | 113202291 | 113202754 | + | 573 | -7137 | Esm1 | bXPF_tRA |
| 1-361 | chr13 | 113794291 | 113794754 | + | 558 | 14 | Arl15 | bXPF_tRA |
| 1-135 | chr13 | 113794492 | 113794955 | + | 558 | 215 | Arl15 | bXPF_tRA |
| 1-1320 | chr13 | 116829060 | 116829523 | + | 558 | 196225 | Parp8 | bXPF_tRA |
| 1-1574 | chr14 | 7773832 | 7774295 | + | 565 | -43894 | Flnb | bXPF_tRA |
| 1-667 | chr14 | 7774031 | 7774494 | + | 565 | -43695 | Flnb | bXPF_tRA |
| 1-607 | chr14 | 7822654 | 7823117 | + | 592 | 4928 | Flnb | bXPF_tRA |
| 1-751 | chr14 | 8097900 | 8098363 | + | 542 | -82 | Pxk | bXPF_tRA |
| 1-1526 | chr14 | 14832875 | 14833338 | + | 665 | 12291 | Nek10 | bXPF_tRA |
| 1-216 | chr14 | 16554933 | 16555396 | + | 558 | 20312 | Rarb | bXPF_tRA |
| 1-1585 | chr14 | 20082831 | 20083294 | + | 565 | 110 | Saysd1 | bXPF_tRA |
| 1 | chr14 | 20347937 | 20348400 | + | 1000 | 6 | Fam149b | bXPF_tRA |

| | | | | | | | | |
|---|---|---|---|---|---|---|---|---|
| 1-1053 | chr14 | 20388644 | 20389107 | + | 547 | 35 | Dnajc9 | bXPF_tRA |
| 1-969 | chr14 | 20734263 | 20734726 | + | 584 | 68 | Ndst2 | bXPF_tRA |
| 44927 | chr14 | 20734462 | 20734925 | + | 584 | -131 | Ndst2 | bXPF_tRA |
| 1-532 | chr14 | 21985796 | 21986259 | + | 574 | 3574 | Zfp503 | bXPF_tRA |
| 1-517 | chr14 | 24486932 | 24487395 | + | 557 | -117 | Polr3a | bXPF_tRA |
| 1-712 | chr14 | 25457448 | 25457911 | + | 559 | -1506 | Zmiz1 | bXPF_tRA |
| 1-686 | chr14 | 29968407 | 29968870 | + | 635 | 258 | Selenok | bXPF_tRA |
| 1-1417 | chr14 | 30479342 | 30479805 | + | 595 | 8 | Dcp1a | bXPF_tRA |
| 1-725 | chr14 | 31001176 | 31001639 | + | 658 | 14 | Glt8d1 | bXPF_tRA |
| 1-330 | chr14 | 31001409 | 31001872 | + | 658 | 26 | Spcs1 | bXPF_tRA |
| 1-1282 | chr14 | 31336619 | 31337082 | + | 543 | 126 | Capn7 | bXPF_tRA |
| 1-765 | chr14 | 31661992 | 31662455 | + | 635 | 21166 | Btd | bXPF_tRA |
| 1-1076 | chr14 | 34673522 | 34673985 | + | 642 | -175 | Wapl | bXPF_tRA |
| 1-449 | chr14 | 34673725 | 34674188 | + | 642 | 28 | Wapl | bXPF_tRA |
| 1-656 | chr14 | 45388629 | 45389092 | + | 587 | -64 | Gnpnat1 | bXPF_tRA |
| 1-1478 | chr14 | 46436443 | 46436906 | + | 731 | -46005 | Bmp4 | bXPF_tRA |
| 1-305 | chr14 | 46436644 | 46437107 | + | 732 | -46206 | Bmp4 | bXPF_tRA |
| 1-1368 | chr14 | 46831816 | 46832279 | + | 575 | -197 | Cgrrf1 | bXPF_tRA |
| 1-1633 | chr14 | 47472294 | 47472757 | + | 791 | -36 | Fbxo34 | bXPF_tRA |
| 1-588 | chr14 | 48446057 | 48446520 | + | 591 | -64 | Tmem260 | bXPF_tRA |
| 1 | chr14 | 48446258 | 48446721 | + | 591 | 137 | Tmem260 | bXPF_tRA |
| 1-1626 | chr14 | 49066313 | 49066776 | + | 554 | 49 | Ap5m1 | bXPF_tRA |
| 1-863 | chr14 | 49171751 | 49172214 | + | 597 | -264 | Naa30 | bXPF_tRA |
| 1097 | chr14 | 51064716 | 51065179 | + | 999 | 6495 | Olfr750 | bXPF_tRA |
| 1-1578 | chr14 | 51088461 | 51088924 | + | 632 | -2385 | Rnase4 | bXPF_tRA |
| 1-1782 | chr14 | 51089265 | 51089728 | + | 825 | -1581 | Rnase4 | bXPF_tRA |
| 1-1860 | chr14 | 51090476 | 51090939 | + | 923 | -370 | Rnase4 | bXPF_tRA |
| 1-704 | chr14 | 54253924 | 54254387 | + | 547 | 26 | Abhd4 | bXPF_tRA |
| 1-874 | chr14 | 54443693 | 54444156 | + | 578 | 12326 | Mmp14 | bXPF_tRA |
| 1-876 | chr14 | 54517474 | 54517937 | + | 638 | -235 | Prmt5 | bXPF_tRA |
| 1-889 | chr14 | 54554084 | 54554547 | + | 541 | 46 | Haus4 | bXPF_tRA |
| 1-549 | chr14 | 54577352 | 54577815 | + | 588 | 78 | Ajuba | bXPF_tRA |
| 1-223 | chr14 | 54577556 | 54578019 | + | 588 | -126 | Ajuba | bXPF_tRA |
| 1-1132 | chr14 | 55823311 | 55823774 | + | 613 | -1253 | Nfatc4 | bXPF_tRA |
| 1-996 | chr14 | 57826262 | 57826725 | + | 547 | 254 | Mrpl57 | bXPF_tRA |
| 1-447 | chr14 | 60264836 | 60265299 | + | 608 | -138 | Mtmr6 | bXPF_tRA |
| 1-830 | chr14 | 61681001 | 61681464 | + | 601 | 1141 | Dleu2 | bXPF_tRA |
| 1-768 | chr14 | 63132191 | 63132654 | + | 720 | 9960 | Ctsb | bXPF_tRA |
| 25204 | chr14 | 63132395 | 63132858 | + | 720 | 10164 | Ctsb | bXPF_tRA |
| 1-986 | chr14 | 65900406 | 65900869 | + | 544 | 53107 | Scara3 | bXPF_tRA |
| 1-1788 | chr14 | 67008863 | 67009326 | + | 713 | -217 | Bnip3l | bXPF_tRA |
| 1-325 | chr14 | 67715752 | 67716215 | + | 568 | -115 | Kctd9 | bXPF_tRA |
| 1-100 | chr14 | 67716030 | 67716493 | + | 568 | 163 | Kctd9 | bXPF_tRA |
| 1-966 | chr14 | 70659046 | 70659509 | + | 541 | -6193 | Npm2 | bXPF_tRA |
| 1-1171 | chr14 | 72983391 | 72983854 | + | 659 | -41981 | Gm9199 | bXPF_tRA |
| 1-901 | chr14 | 73143360 | 73143823 | + | 715 | 806 | Rcbtb2 | bXPF_tRA |
| 1-677 | chr14 | 73293574 | 73294037 | + | 587 | 31986 | Rb1 | bXPF_tRA |
| 1-1522 | chr14 | 73524780 | 73525243 | + | 712 | 14962 | Med4 | bXPF_tRA |
| 1-1285 | chr14 | 74309471 | 74309934 | + | 637 | -45825 | 4933402J15Rik | bXPF_tRA |
| 1-1454 | chr14 | 76110855 | 76111318 | + | 643 | 195 | Nufip1 | bXPF_tRA |
| 1-593 | chr14 | 76504383 | 76504846 | + | 633 | 104 | Tsc22d1 | bXPF_tRA |
| 1-1760 | chr14 | 76681673 | 76682136 | + | 779 | 4336 | 1700108F19Rik | bXPF_tRA |
| 1-1069 | chr14 | 78849069 | 78849532 | + | 602 | 122 | Vwa8 | bXPF_tRA |
| 1-1593 | chr14 | 78900778 | 78901241 | + | 654 | 51831 | Vwa8 | bXPF_tRA |
| 1-985 | chr14 | 79426413 | 79426876 | + | 590 | 133 | Kbtbd7 | bXPF_tRA |
| 1-1429 | chr14 | 101784357 | 101784820 | + | 636 | 54660 | Lmo7 | bXPF_tRA |
| 1-1648 | chr14 | 101804757 | 101805220 | + | 637 | -35635 | Lmo7 | bXPF_tRA |
| 1-520 | chr14 | 101804976 | 101805439 | + | 554 | -35416 | Lmo7 | bXPF_tRA |
| 1-1064 | chr14 | 102983014 | 102983477 | + | 548 | -608 | Kctd12 | bXPF_tRA |
| 1-1244 | chr14 | 106012127 | 106012590 | + | 690 | -93840 | Trim52 | bXPF_tRA |
| 1-583 | chr14 | 118566398 | 118566861 | + | 599 | 25843 | Mir6391 | bXPF_tRA |
| 1-1601 | chr14 | 120137881 | 120138344 | + | 645 | -95677 | Oxgr1 | bXPF_tRA |
| 33239 | chr14 | 120138080 | 120138543 | + | 644 | -95876 | Oxgr1 | bXPF_tRA |

| | | | | | | | | |
|---|---|---|---|---|---|---|---|---|
| 1-1571 | chr14 | 121878194 | 121878657 | + | 802 | 79 | 1810041H14Rik | bXPF_tRA |
| 1-1185 | chr14 | 122534052 | 122534515 | + | 597 | -45 | Pcca | bXPF_tRA |
| 34700 | chr14 | 123236435 | 123236898 | + | 546 | 67480 | AA536875 | bXPF_tRA |
| 1-1628 | chr15 | 3995691 | 3996154 | + | 813 | -117 | A630020A06 | bXPF_tRA |
| 1-205 | chr15 | 3995902 | 3996365 | + | 812 | 94 | A630020A06 | bXPF_tRA |
| 1-1479 | chr15 | 5116308 | 5116771 | + | 579 | -74 | Rpl37 | bXPF_tRA |
| 1-1357 | chr15 | 6386343 | 6386806 | + | 666 | -174 | Dab2 | bXPF_tRA |
| 1-128 | chr15 | 6386583 | 6387046 | + | 666 | 66 | Dab2 | bXPF_tRA |
| 1-1310 | chr15 | 8684322 | 8684785 | + | 718 | 26254 | Slc1a3 | bXPF_tRA |
| 1-565 | chr15 | 10759982 | 10760445 | + | 652 | 45313 | 4930556M19Rik | bXPF_tRA |
| 44197 | chr15 | 10760194 | 10760657 | + | 652 | 45525 | 4930556M19Rik | bXPF_tRA |
| 1-1458 | chr15 | 10890129 | 10890592 | + | 641 | 61309 | Gm19276 | bXPF_tRA |
| 1-258 | chr15 | 10890339 | 10890802 | + | 641 | 61519 | Gm19276 | bXPF_tRA |
| 1-668 | chr15 | 12739802 | 12740265 | + | 578 | 84624 | 6030458C11Rik | bXPF_tRA |
| 1-781 | chr15 | 25767626 | 25768089 | + | 542 | -75441 | Fam134b | bXPF_tRA |
| 1-1752 | chr15 | 27923703 | 27924166 | + | 869 | 101914 | Trio | bXPF_tRA |
| 1-834 | chr15 | 34443123 | 34443586 | + | 613 | -78 | Rpl30 | bXPF_tRA |
| 1-1824 | chr15 | 35371247 | 35371710 | + | 703 | -68 | Vps13b | bXPF_tRA |
| 1-181 | chr15 | 38519458 | 38519921 | + | 596 | -423 | Azin1 | bXPF_tRA |
| 1-1122 | chr15 | 43170660 | 43171123 | + | 612 | -73 | Rspo2 | bXPF_tRA |
| 1-131 | chr15 | 44427597 | 44428060 | + | 543 | -283 | Eny2 | bXPF_tRA |
| 1-256 | chr15 | 44427937 | 44428400 | + | 543 | 57 | Eny2 | bXPF_tRA |
| 1-1838 | chr15 | 51798682 | 51799145 | + | 944 | 66548 | Eif3h | bXPF_tRA |
| 1-1301 | chr15 | 53134082 | 53134545 | + | 582 | 211870 | Ext1 | bXPF_tRA |
| 1-336 | chr15 | 53342882 | 53343345 | + | 540 | 3070 | Ext1 | bXPF_tRA |
| 31778 | chr15 | 53343082 | 53343545 | + | 540 | 2870 | Ext1 | bXPF_tRA |
| 1-1271 | chr15 | 55557109 | 55557572 | + | 547 | -28 | Mrpl13 | bXPF_tRA |
| 1-895 | chr15 | 57659965 | 57660428 | + | 619 | -34471 | Zhx2 | bXPF_tRA |
| 1-1450 | chr15 | 57975748 | 57976211 | + | 644 | -9924 | Fam83a | bXPF_tRA |
| 1-1687 | chr15 | 58480340 | 58480803 | + | 751 | 65103 | Fam91a1 | bXPF_tRA |
| 1-1292 | chr15 | 58823330 | 58823793 | + | 699 | -134 | Tmem65 | bXPF_tRA |
| 1-286 | chr15 | 58823539 | 58824002 | + | 699 | -343 | Tmem65 | bXPF_tRA |
| 1-690 | chr15 | 59374015 | 59374478 | + | 599 | 48 | Nsmce2 | bXPF_tRA |
| 1-1316 | chr15 | 59648394 | 59648857 | + | 642 | -29 | Trib1 | bXPF_tRA |
| 1-248 | chr15 | 59680727 | 59681190 | + | 587 | 32304 | Trib1 | bXPF_tRA |
| 1-592 | chr15 | 59777401 | 59777864 | + | 593 | 17327 | Gm19510 | bXPF_tRA |
| 1-207 | chr15 | 59777601 | 59778064 | + | 593 | 17127 | Gm19510 | bXPF_tRA |
| 1-852 | chr15 | 60824574 | 60825037 | + | 553 | 275 | Fam84b | bXPF_tRA |
| 1-1764 | chr15 | 63340263 | 63340726 | + | 885 | 139114 | Gm20740 | bXPF_tRA |
| 1-265 | chr15 | 63340464 | 63340927 | + | 795 | 138913 | Gm20740 | bXPF_tRA |
| 1-711 | chr15 | 67176882 | 67177345 | + | 607 | -231 | St3gal1 | bXPF_tRA |
| 1-1038 | chr15 | 73060912 | 73061375 | + | 742 | 61 | Trappc9 | bXPF_tRA |
| 1-1880 | chr15 | 75940798 | 75941261 | + | 777 | 77 | Zfp623 | bXPF_tRA |
| 1-1789 | chr15 | 76327126 | 76327589 | + | 572 | -40 | Exosc4 | bXPF_tRA |
| 10228 | chr15 | 76607330 | 76607793 | + | 855 | 30 | Cpsf1 | bXPF_tRA |
| 1-855 | chr15 | 76903898 | 76904361 | + | 577 | 58 | Rpl8 | bXPF_tRA |
| 1-1825 | chr15 | 76948312 | 76948775 | + | 867 | -1 | 1110038F14Rik | bXPF_tRA |
| 1-340 | chr15 | 76948513 | 76948976 | + | 695 | 200 | 1110038F14Rik | bXPF_tRA |
| 1-636 | chr15 | 77076490 | 77076953 | + | 578 | -7677 | 1700109K24Rik | bXPF_tRA |
| 1-652 | chr15 | 77153577 | 77154040 | + | 572 | 4 | Rbfox2 | bXPF_tRA |
| 1-587 | chr15 | 77336377 | 77336840 | + | 548 | -29555 | Rbfox2 | bXPF_tRA |
| 1-792 | chr15 | 77358501 | 77358964 | + | 553 | 40378 | Apol7a | bXPF_tRA |
| 1-528 | chr15 | 77366729 | 77367192 | + | 655 | 32150 | Apol7a | bXPF_tRA |
| 1-1697 | chr15 | 77798630 | 77799093 | + | 659 | 43314 | Myh9 | bXPF_tRA |
| 1-772 | chr15 | 77882767 | 77883230 | + | 604 | -40823 | Myh9 | bXPF_tRA |
| 1-1156 | chr15 | 78450004 | 78450467 | + | 572 | 18399 | Tmprss6 | bXPF_tRA |
| 1-395 | chr15 | 78926472 | 78926935 | + | 642 | -22 | Lgals1 | bXPF_tRA |
| 1-651 | chr15 | 78934672 | 78935135 | + | 550 | -30 | Nol12 | bXPF_tRA |
| 1-702 | chr15 | 79028234 | 79028697 | + | 552 | 253 | H1f0 | bXPF_tRA |
| 1-1776 | chr15 | 79328087 | 79328550 | + | 785 | 53 | Pla2g6 | bXPF_tRA |
| 35431 | chr15 | 79328288 | 79328751 | + | 784 | -148 | Pla2g6 | bXPF_tRA |
| 1-641 | chr15 | 80133010 | 80133473 | + | 746 | 87 | Tab1 | bXPF_tRA |
| 1-1726 | chr15 | 80264122 | 80264585 | + | 692 | -47 | Rps19bp1 | bXPF_tRA |

| | | | | | | | | |
|---|---|---|---|---|---|---|---|---|
| 1-1722 | chr15 | 81858131 | 81858594 | + | 636 | -36 | Tob2 | bXPF_tRA |
| 1-1360 | chr15 | 85859268 | 85859731 | + | 627 | -208 | Gtse1 | bXPF_tRA |
| 1-821 | chr15 | 95931263 | 95931726 | + | 549 | 140651 | Ano6 | bXPF_tRA |
| 1-832 | chr15 | 98108329 | 98108792 | + | 573 | 89 | Pfkm | bXPF_tRA |
| 1-1153 | chr15 | 98662632 | 98663095 | + | 547 | 26 | Ddx23 | bXPF_tRA |
| 1-1456 | chr15 | 98953255 | 98953718 | + | 708 | 15 | Tuba1a | bXPF_tRA |
| 16438 | chr15 | 98953473 | 98953936 | + | 708 | -203 | Tuba1a | bXPF_tRA |
| 1-154 | chr15 | 99369826 | 99370289 | + | 564 | 425 | Fmnl3 | bXPF_tRA |
| 1-684 | chr15 | 99614768 | 99615231 | + | 602 | 13599 | Aqp6 | bXPF_tRA |
| 1-1041 | chr15 | 99701931 | 99702394 | + | 725 | -125 | Smarcd1 | bXPF_tRA |
| 1-1393 | chr15 | 99972681 | 99973144 | + | 726 | 132 | Larp4 | bXPF_tRA |
| 1-1826 | chr15 | 100551632 | 100552095 | + | 896 | 145 | Tfcp2 | bXPF_tRA |
| 1-363 | chr15 | 102247519 | 102247982 | + | 542 | -1250 | Rarg | bXPF_tRA |
| 1-1422 | chr15 | 102470151 | 102470614 | + | 676 | -250 | Pcbp2 | bXPF_tRA |
| 1-1042 | chr15 | 102670873 | 102671336 | + | 667 | -57 | Atp5g2 | bXPF_tRA |
| 1-782 | chr16 | 4003831 | 4004294 | + | 607 | -2384 | Slx4 | bXPF_tRA |
| 1-302 | chr16 | 4004030 | 4004493 | + | 607 | -2583 | Slx4 | bXPF_tRA |
| 1-730 | chr16 | 4077645 | 4078108 | + | 558 | -66 | Trap1 | bXPF_tRA |
| 1-877 | chr16 | 4273840 | 4274303 | + | 588 | 44842 | Gm5766 | bXPF_tRA |
| 1-1603 | chr16 | 5255746 | 5256209 | + | 691 | -21 | Eef2kmt | bXPF_tRA |
| 1-516 | chr16 | 8671753 | 8672216 | + | 592 | 169 | Carhsp1 | bXPF_tRA |
| 1-359 | chr16 | 10542855 | 10543318 | + | 590 | -32 | Dexi | bXPF_tRA |
| 1-491 | chr16 | 11066165 | 11066628 | + | 768 | 98 | Snn | bXPF_tRA |
| 1-452 | chr16 | 15084855 | 15085318 | + | 701 | 178440 | Efcab1 | bXPF_tRA |
| 1-907 | chr16 | 17759390 | 17759853 | + | 631 | 0 | Klhl22 | bXPF_tRA |
| 4384 | chr16 | 17807147 | 17807610 | + | 807 | 10096 | Scarf2 | bXPF_tRA |
| 1-337 | chr16 | 20516621 | 20517084 | + | 574 | -134 | Dvl3 | bXPF_tRA |
| 1-224 | chr16 | 20516823 | 20517286 | + | 574 | -10 | Dvl3 | bXPF_tRA |
| 1-754 | chr16 | 20716094 | 20716557 | + | 542 | 311 | Clcn2 | bXPF_tRA |
| 1-628 | chr16 | 21318889 | 21319352 | + | 555 | 14236 | Magef1 | bXPF_tRA |
| 1-767 | chr16 | 21353203 | 21353666 | + | 570 | -20078 | Magef1 | bXPF_tRA |
| 1-481 | chr16 | 21377520 | 21377983 | + | 606 | -44395 | Magef1 | bXPF_tRA |
| 1-1858 | chr16 | 23107330 | 23107793 | + | 613 | 93 | Eif4a2 | bXPF_tRA |
| 1-1869 | chr16 | 23988547 | 23989010 | + | 653 | -166 | Bcl6 | bXPF_tRA |
| 1-598 | chr16 | 24494560 | 24495023 | + | 549 | -34523 | Morf4l1-ps1 | bXPF_tRA |
| 1-833 | chr16 | 25531977 | 25532440 | + | 543 | -151557 | Trp63 | bXPF_tRA |
| 1-1709 | chr16 | 29579175 | 29579638 | + | 863 | 72 | Opa1 | bXPF_tRA |
| 1-1463 | chr16 | 29946001 | 29946464 | + | 651 | 32892 | Gm1968 | bXPF_tRA |
| 1-827 | chr16 | 30008862 | 30009325 | + | 543 | 426 | 4632428C04Rik | bXPF_tRA |
| 1-1645 | chr16 | 30066953 | 30067416 | + | 627 | 1827 | Hes1 | bXPF_tRA |
| 1-137 | chr16 | 30067220 | 30067683 | + | 627 | 2094 | Hes1 | bXPF_tRA |
| 1-344 | chr16 | 30068325 | 30068788 | + | 673 | 3199 | Hes1 | bXPF_tRA |
| 1-324 | chr16 | 30068577 | 30069040 | + | 673 | 3451 | Hes1 | bXPF_tRA |
| 1-1498 | chr16 | 30187983 | 30188446 | + | 616 | 79318 | Cpn2 | bXPF_tRA |
| 1-682 | chr16 | 30550291 | 30550754 | + | 722 | 56 | Tmem44 | bXPF_tRA |
| 1-1218 | chr16 | 30797799 | 30798262 | + | 814 | 198307 | Fam43a | bXPF_tRA |
| 1-645 | chr16 | 30827791 | 30828254 | + | 602 | 228299 | Fam43a | bXPF_tRA |
| 1-1436 | chr16 | 31233846 | 31234309 | + | 592 | -32839 | Acap2 | bXPF_tRA |
| 1-1844 | chr16 | 31598547 | 31599010 | + | 801 | -64665 | Dlg1 | bXPF_tRA |
| 1-388 | chr16 | 31663386 | 31663849 | + | 791 | 174 | Dlg1 | bXPF_tRA |
| 32874 | chr16 | 31663601 | 31664064 | + | 790 | -103 | Dlg1 | bXPF_tRA |
| 20821 | chr16 | 32639940 | 32640403 | + | 1000 | -4472 | Tnk2 | bXPF_tRA |
| 1-1254 | chr16 | 32859110 | 32859573 | + | 642 | 9025 | Rubcn | bXPF_tRA |
| 1-1572 | chr16 | 33380419 | 33380882 | + | 720 | 86 | 1700007L15Rik | bXPF_tRA |
| 18264 | chr16 | 33380618 | 33381081 | + | 721 | 74 | Zfp148 | bXPF_tRA |
| 1-854 | chr16 | 33922584 | 33923047 | + | 580 | 44188 | Umps | bXPF_tRA |
| 1-123 | chr16 | 33922813 | 33923276 | + | 580 | 43959 | Umps | bXPF_tRA |
| 1-1608 | chr16 | 34958630 | 34959093 | + | 737 | 3 | E130310I04Rik | bXPF_tRA |
| 1-1227 | chr16 | 35490681 | 35491144 | + | 610 | -39 | Pdia5 | bXPF_tRA |
| 1-591 | chr16 | 35501070 | 35501533 | + | 640 | 8118 | 1600019K03Rik | bXPF_tRA |
| 1-1256 | chr16 | 36071736 | 36072199 | + | 626 | 307 | Ccdc58 | bXPF_tRA |
| 1-653 | chr16 | 37788402 | 37788865 | + | 566 | 11578 | Fstl1 | bXPF_tRA |
| 1-953 | chr16 | 37867892 | 37868355 | + | 554 | -277 | Lrrc58 | bXPF_tRA |

| | | | | | | | | |
|---|---|---|---|---|---|---|---|---|
| 1-597 | chr16 | 42190083 | 42190546 | + | 664 | 113155 | Mir6540 | bXPF_tRA |
| 23743 | chr16 | 42190292 | 42190755 | + | 664 | 112946 | Mir6540 | bXPF_tRA |
| 1-777 | chr16 | 44745923 | 44746386 | + | 541 | 209 | Gtpbp8 | bXPF_tRA |
| 1-1340 | chr16 | 45844149 | 45844612 | + | 567 | -2 | Phldb2 | bXPF_tRA |
| 32143 | chr16 | 45892408 | 45892871 | + | 577 | -48261 | Phldb2 | bXPF_tRA |
| 1-288 | chr16 | 45892722 | 45893185 | + | 577 | -48575 | Phldb2 | bXPF_tRA |
| 1-192 | chr16 | 46497507 | 46497970 | + | 605 | -771 | Nectin3 | bXPF_tRA |
| 1-1737 | chr16 | 49855073 | 49855536 | + | 910 | -350 | Cd47 | bXPF_tRA |
| 1-1612 | chr16 | 50432223 | 50432686 | + | 647 | 12 | Bbx | bXPF_tRA |
| 1-980 | chr16 | 55731054 | 55731517 | + | 623 | 90853 | Nfkbiz | bXPF_tRA |
| 1-317 | chr16 | 55731262 | 55731725 | + | 622 | 90645 | Nfkbiz | bXPF_tRA |
| 1-1465 | chr16 | 56037486 | 56037949 | + | 626 | 57 | Trmt10c | bXPF_tRA |
| 1-1660 | chr16 | 57231175 | 57231638 | + | 645 | 60 | Tbc1d23 | bXPF_tRA |
| 1-1461 | chr16 | 57427473 | 57427936 | + | 784 | 74427 | Filip1l | bXPF_tRA |
| 1-793 | chr16 | 57606683 | 57607146 | + | 605 | -47 | Cmss1 | bXPF_tRA |
| 1-1150 | chr16 | 63857060 | 63857523 | + | 602 | 6866 | Epha3 | bXPF_tRA |
| 1-873 | chr16 | 64851598 | 64852061 | + | 617 | -106 | Zfp654 | bXPF_tRA |
| 1-561 | chr16 | 70314237 | 70314700 | + | 947 | 519 | Gbe1 | bXPF_tRA |
| 1-476 | chr16 | 72663017 | 72663480 | + | 544 | 99 | Robo1 | bXPF_tRA |
| 1-582 | chr16 | 72663262 | 72663725 | + | 542 | 344 | Robo1 | bXPF_tRA |
| 1-1663 | chr16 | 84735100 | 84735563 | + | 808 | -29 | Mrpl39 | bXPF_tRA |
| 1-717 | chr16 | 90143808 | 90144271 | + | 594 | 1802 | Gm10789 | bXPF_tRA |
| 1-695 | chr16 | 90220444 | 90220907 | + | 548 | -67 | Sod1 | bXPF_tRA |
| 1-721 | chr16 | 90934629 | 90935092 | + | 568 | -11 | 1110004E09Rik | bXPF_tRA |
| 1-1724 | chr16 | 91044406 | 91044869 | + | 670 | -258 | Paxbp1 | bXPF_tRA |
| 1-1118 | chr16 | 93603582 | 93604045 | + | 678 | 2 | Setd4 | bXPF_tRA |
| 1-553 | chr16 | 95702364 | 95702827 | + | 595 | 188 | Ets2 | bXPF_tRA |
| 1-846 | chr16 | 96144932 | 96145395 | + | 595 | -256 | Wrb | bXPF_tRA |
| 1-1276 | chr16 | 97962275 | 97962738 | + | 683 | 116 | Zbtb21 | bXPF_tRA |
| 1-1021 | chr17 | 4173880 | 4174343 | + | 830 | -52009 | 4930548J01Rik | bXPF_tRA |
| 4019 | chr17 | 6429767 | 6430230 | + | 1000 | -114 | Dynlt1b | bXPF_tRA |
| 10959 | chr17 | 6429966 | 6430429 | + | 1000 | 85 | Dynlt1b | bXPF_tRA |
| 1-673 | chr17 | 6948402 | 6948865 | + | 595 | -277 | Rsph3b | bXPF_tRA |
| 1-654 | chr17 | 12782818 | 12783281 | + | 597 | -13343 | Igf2r | bXPF_tRA |
| 1-489 | chr17 | 12992398 | 12992861 | + | 546 | -90 | Wtap | bXPF_tRA |
| 1-1506 | chr17 | 14546050 | 14546513 | + | 620 | -147339 | 4930474M22Rik | bXPF_tRA |
| 1-1182 | chr17 | 14693912 | 14694375 | + | 630 | 119 | Thbs2 | bXPF_tRA |
| 1-490 | chr17 | 14694111 | 14694574 | + | 630 | -80 | Thbs2 | bXPF_tRA |
| 1-1159 | chr17 | 15499670 | 15500133 | + | 557 | 13 | Tbp | bXPF_tRA |
| 1-1003 | chr17 | 15564162 | 15564625 | + | 654 | -1070 | Prdm9 | bXPF_tRA |
| 1-1674 | chr17 | 17837742 | 17838205 | + | 727 | 6998 | Spaca6 | bXPF_tRA |
| 1-1690 | chr17 | 20945274 | 20945737 | + | 754 | 51 | Ppp2r1a | bXPF_tRA |
| 1-1746 | chr17 | 23546259 | 23546722 | + | 818 | -4309 | 6330415G19Rik | bXPF_tRA |
| 1-1843 | chr17 | 24073317 | 24073780 | + | 874 | -63 | Kctd5 | bXPF_tRA |
| 1-1840 | chr17 | 24470025 | 24470488 | + | 949 | -217 | Pgp | bXPF_tRA |
| 1-538 | chr17 | 24478716 | 24479179 | + | 564 | 131 | Mlst8 | bXPF_tRA |
| 1-1870 | chr17 | 25083300 | 25083763 | + | 671 | -2417 | Tmem204 | bXPF_tRA |
| 1-496 | chr17 | 25273641 | 25274104 | + | 615 | 42 | Ube2i | bXPF_tRA |
| 1-875 | chr17 | 25773601 | 25774064 | + | 580 | 56 | Narfl | bXPF_tRA |
| 1-1191 | chr17 | 25792227 | 25792690 | + | 750 | -64 | Fam173a | bXPF_tRA |
| 1-218 | chr17 | 27133366 | 27133829 | + | 561 | 294 | Uqcc2 | bXPF_tRA |
| 1-1215 | chr17 | 28084898 | 28085361 | + | 588 | -4490 | Tcp11 | bXPF_tRA |
| 46388 | chr17 | 28085098 | 28085561 | + | 588 | -4690 | Tcp11 | bXPF_tRA |
| 1-909 | chr17 | 28232342 | 28232805 | + | 578 | -181 | Ppard | bXPF_tRA |
| 1-1255 | chr17 | 28313366 | 28313829 | + | 579 | 67 | Fance | bXPF_tRA |
| 1-1781 | chr17 | 28691068 | 28691531 | + | 831 | -43 | Mapk14 | bXPF_tRA |
| 1-1808 | chr17 | 29456453 | 29456916 | + | 864 | -34361 | Pim1 | bXPF_tRA |
| 1-1302 | chr17 | 29660515 | 29660978 | + | 763 | 151 | Cmtr1 | bXPF_tRA |
| 1-1553 | chr17 | 30465325 | 30465788 | + | 560 | -69376 | Gm6402 | bXPF_tRA |
| 1-1642 | chr17 | 33760245 | 33760708 | + | 593 | 10 | Rab11b | bXPF_tRA |
| 1-1786 | chr17 | 33890636 | 33891099 | + | 813 | -234 | Kifc1 | bXPF_tRA |
| 1-1711 | chr17 | 33929232 | 33929695 | + | 765 | -431 | Rgl2 | bXPF_tRA |
| 1-173 | chr17 | 33929433 | 33929896 | + | 765 | -230 | Rgl2 | bXPF_tRA |

| | | | | | | | | |
|---|---|---|---|---|---|---|---|---|
| 1-857 | chr17 | 33955638 | 33956101 | + | 712 | -13 | Vps52 | bXPF_tRA |
| 29587 | chr17 | 34118990 | 34119453 | + | 680 | 679 | Brd2 | bXPF_tRA |
| 1-209 | chr17 | 35134729 | 35135192 | + | 598 | -218 | Bag6 | bXPF_tRA |
| 1-355 | chr17 | 35821156 | 35821619 | + | 612 | -326 | Ier3 | bXPF_tRA |
| 1-386 | chr17 | 35821477 | 35821940 | + | 611 | -5 | Ier3 | bXPF_tRA |
| 1-894 | chr17 | 35841076 | 35841539 | + | 584 | -191 | Mdc1 | bXPF_tRA |
| 1-1851 | chr17 | 35879484 | 35879947 | + | 883 | -63 | Dhx16 | bXPF_tRA |
| 1-950 | chr17 | 36951513 | 36951976 | + | 812 | 48 | Ppp1r11 | bXPF_tRA |
| 1-1730 | chr17 | 44105522 | 44105985 | + | 567 | 55 | Enpp4 | bXPF_tRA |
| 1-992 | chr17 | 44261621 | 44262084 | + | 682 | 73280 | Clic5 | bXPF_tRA |
| 1-156 | chr17 | 44261834 | 44262297 | + | 681 | 73493 | Clic5 | bXPF_tRA |
| 1-1347 | chr17 | 44500174 | 44500637 | + | 650 | 235771 | Runx2 | bXPF_tRA |
| 1-1384 | chr17 | 45359511 | 45359974 | + | 594 | 73965 | Cdc5l | bXPF_tRA |
| 1-948 | chr17 | 45506617 | 45507080 | + | 626 | 7 | Aars2 | bXPF_tRA |
| 1-1694 | chr17 | 47736781 | 47737244 | + | 884 | -25 | Tfeb | bXPF_tRA |
| 1-155 | chr17 | 47736983 | 47737446 | + | 563 | 177 | Tfeb | bXPF_tRA |
| 1-1473 | chr17 | 53689219 | 53689682 | + | 559 | -135 | Sgol1 | bXPF_tRA |
| 1-750 | chr17 | 55891978 | 55892441 | + | 606 | 116 | Zfp959 | bXPF_tRA |
| 1-1317 | chr17 | 56036466 | 56036929 | + | 703 | -60 | Sh3gl1 | bXPF_tRA |
| 1-994 | chr17 | 56079085 | 56079548 | + | 670 | -315 | Hdgfrp2 | bXPF_tRA |
| 1-1149 | chr17 | 56242017 | 56242480 | + | 582 | -740 | Mir7b | bXPF_tRA |
| 1-1029 | chr17 | 56507192 | 56507655 | + | 677 | 5060 | Znrf4 | bXPF_tRA |
| 1-839 | chr17 | 56613196 | 56613659 | + | 647 | 32 | Rpl36 | bXPF_tRA |
| 1-1117 | chr17 | 63678224 | 63678687 | + | 677 | -177875 | Fbxl17 | bXPF_tRA |
| 1-693 | chr17 | 63712807 | 63713270 | + | 621 | -150943 | Fer | bXPF_tRA |
| 1-1810 | chr17 | 64332064 | 64332527 | + | 953 | -412 | Pja2 | bXPF_tRA |
| 1-1287 | chr17 | 65613673 | 65614136 | + | 552 | -349 | Vapa | bXPF_tRA |
| 1-1087 | chr17 | 69417165 | 69417628 | + | 599 | 949 | C030034I22Rik | bXPF_tRA |
| 1-629 | chr17 | 70683790 | 70684253 | + | 550 | 122274 | Dlgap1 | bXPF_tRA |
| 1-1638 | chr17 | 70719668 | 70720131 | + | 881 | 126878 | Tgif1 | bXPF_tRA |
| 1-744 | chr17 | 70719869 | 70720332 | + | 655 | 126677 | Tgif1 | bXPF_tRA |
| 1-1842 | chr17 | 73970374 | 73970837 | + | 771 | -20409 | Xdh | bXPF_tRA |
| 1-127 | chr17 | 73970573 | 73971036 | + | 769 | -20608 | Xdh | bXPF_tRA |
| 1-428 | chr17 | 74527714 | 74528177 | + | 617 | -350 | Birc6 | bXPF_tRA |
| 1-463 | chr17 | 75110454 | 75110917 | + | 565 | -68096 | Ltbp1 | bXPF_tRA |
| 1-323 | chr17 | 75110746 | 75111209 | + | 566 | -67804 | Ltbp1 | bXPF_tRA |
| 1-1459 | chr17 | 75178495 | 75178958 | + | 572 | -55 | Ltbp1 | bXPF_tRA |
| 1-1525 | chr17 | 78322544 | 78323007 | + | 548 | 95377 | Fez2 | bXPF_tRA |
| 1-743 | chr17 | 79421535 | 79421998 | + | 550 | 56119 | 4930429F11Rik | bXPF_tRA |
| 1-1299 | chr17 | 79895979 | 79896442 | + | 597 | -87 | Atl2 | bXPF_tRA |
| 1-1237 | chr17 | 80373180 | 80373643 | + | 604 | 131 | Gm10190 | bXPF_tRA |
| 1-1160 | chr17 | 80482783 | 80483246 | + | 635 | -2561 | Sos1 | bXPF_tRA |
| 1-1576 | chr17 | 80801606 | 80802069 | + | 669 | -73812 | Map4k3 | bXPF_tRA |
| 1-1365 | chr17 | 81605304 | 81605767 | + | 623 | 132852 | Slc8a1 | bXPF_tRA |
| 1-866 | chr17 | 83903410 | 83903873 | + | 576 | -56851 | Haao | bXPF_tRA |
| 22647 | chr17 | 84186556 | 84186768 | + | 1000 | 1285 | Zfp36l2 | bXPF_tRA |
| 1-1751 | chr17 | 84186664 | 84187127 | + | 1000 | 1052 | Zfp36l2 | bXPF_tRA |
| 1-1257 | chr17 | 84510278 | 84510741 | + | 668 | -1386 | Plekhh2 | bXPF_tRA |
| 1-385 | chr17 | 87170058 | 87170521 | + | 556 | 24653 | C330024C12Rik | bXPF_tRA |
| 1-233 | chr17 | 87202256 | 87202719 | + | 573 | -7545 | C330024C12Rik | bXPF_tRA |
| 1-201 | chr17 | 87202645 | 87203108 | + | 573 | -7934 | C330024C12Rik | bXPF_tRA |
| 1-1427 | chr17 | 87314926 | 87315389 | + | 684 | 32271 | Ttc7 | bXPF_tRA |
| 1-471 | chr17 | 87446844 | 87447307 | + | 582 | -140 | Calm2 | bXPF_tRA |
| 1-1503 | chr18 | 3337351 | 3337814 | + | 748 | 7 | Crem | bXPF_tRA |
| 1-405 | chr18 | 5163386 | 5163849 | + | 574 | 117028 | Svil | bXPF_tRA |
| 1-822 | chr18 | 6515905 | 6516368 | + | 554 | -28 | Epc1 | bXPF_tRA |
| 1-1507 | chr18 | 7004591 | 7005054 | + | 575 | -43 | Mkx | bXPF_tRA |
| 46753 | chr18 | 7004798 | 7005261 | + | 575 | -250 | Mkx | bXPF_tRA |
| 1-1311 | chr18 | 10725401 | 10725864 | + | 679 | 7 | Mib1 | bXPF_tRA |
| 1-732 | chr18 | 16713565 | 16714028 | + | 547 | 95450 | Cdh2 | bXPF_tRA |
| 1-1513 | chr18 | 21300076 | 21300539 | + | 566 | -168 | Garem1 | bXPF_tRA |
| 1-1375 | chr18 | 23803664 | 23804127 | + | 562 | -75 | Mapre2 | bXPF_tRA |
| 1-1765 | chr18 | 32677209 | 32677672 | + | 596 | -117406 | Gypc | bXPF_tRA |

| | | | | | | | | |
|---|---|---|---|---|---|---|---|---|
| 1-1068 | chr18 | 32836934 | 32837397 | + | 543 | -60 | Wdr36 | bXPF_tRA |
| 1-1699 | chr18 | 33463670 | 33464133 | + | 792 | 128 | Nrep | bXPF_tRA |
| 1-1272 | chr18 | 33794713 | 33795176 | + | 617 | 52 | Epb41l4aos | bXPF_tRA |
| 1-1528 | chr18 | 34330852 | 34331315 | + | 723 | -62 | Srp19 | bXPF_tRA |
| 13150 | chr18 | 34751490 | 34751953 | + | 1000 | -188 | Cdc25c | bXPF_tRA |
| 1-1325 | chr18 | 34859771 | 34860234 | + | 600 | -1205 | Egr1 | bXPF_tRA |
| 1-675 | chr18 | 36679126 | 36679589 | + | 710 | 9 | Apbb3 | bXPF_tRA |
| 1-1754 | chr18 | 36783048 | 36783511 | + | 706 | 70 | Hars2 | bXPF_tRA |
| 1-662 | chr18 | 36801616 | 36802079 | + | 567 | 84 | Vaultrc5 | bXPF_tRA |
| 1-1152 | chr18 | 38296482 | 38296945 | + | 601 | 78 | Rnf14 | bXPF_tRA |
| 1-501 | chr18 | 38804542 | 38805005 | + | 570 | -87683 | Gm5820 | bXPF_tRA |
| 1-688 | chr18 | 42511053 | 42511516 | + | 664 | -203 | Tcerg1 | bXPF_tRA |
| 1-1700 | chr18 | 52639457 | 52639920 | + | 656 | -6532 | 1700034E13Rik | bXPF_tRA |
| 1-1061 | chr18 | 53744416 | 53744879 | + | 688 | -100 | Cep120 | bXPF_tRA |
| 1-1635 | chr18 | 54374096 | 54374559 | + | 576 | -47968 | Redrum | bXPF_tRA |
| 33604 | chr18 | 54374299 | 54374762 | + | 575 | -47765 | Redrum | bXPF_tRA |
| 1-1315 | chr18 | 54378054 | 54378517 | + | 619 | -44010 | Redrum | bXPF_tRA |
| 1-1658 | chr18 | 54902047 | 54902510 | + | 542 | 87902 | Zfp608 | bXPF_tRA |
| 1-1733 | chr18 | 54985545 | 54986008 | + | 777 | 4404 | Zfp608 | bXPF_tRA |
| 1-116 | chr18 | 54985745 | 54986208 | + | 676 | 4204 | Zfp608 | bXPF_tRA |
| 4750 | chr18 | 55133806 | 55134269 | + | 904 | -143857 | Zfp608 | bXPF_tRA |
| 1-1408 | chr18 | 55134007 | 55134470 | + | 751 | -144058 | Zfp608 | bXPF_tRA |
| 1-1556 | chr18 | 55344006 | 55344469 | + | 788 | -354057 | Zfp608 | bXPF_tRA |
| 1-1569 | chr18 | 60648832 | 60649295 | + | 754 | -14347 | G630071F17Rik | bXPF_tRA |
| 1-649 | chr18 | 64685062 | 64685525 | + | 562 | -24293 | Atp8b1 | bXPF_tRA |
| 1-1775 | chr18 | 65934829 | 65935292 | + | 909 | 4029 | Rax | bXPF_tRA |
| 6941 | chr18 | 66162126 | 66162589 | + | 620 | 56976 | Mir694 | bXPF_tRA |
| 1-1178 | chr18 | 66162325 | 66162788 | + | 573 | 56777 | Mir694 | bXPF_tRA |
| 1-709 | chr18 | 67641324 | 67641787 | + | 569 | -44 | Psmg2 | bXPF_tRA |
| 1-949 | chr18 | 73703693 | 73704156 | + | 618 | -183 | Smad4 | bXPF_tRA |
| 1-1086 | chr18 | 75366562 | 75367025 | + | 551 | -572 | Smad7 | bXPF_tRA |
| 1-1402 | chr18 | 78546956 | 78547419 | + | 648 | 49763 | Slc14a2 | bXPF_tRA |
| 12055 | chr18 | 78547161 | 78547624 | + | 648 | 49558 | Slc14a2 | bXPF_tRA |
| 1-914 | chr18 | 78554199 | 78554662 | + | 634 | 42520 | Slc14a2 | bXPF_tRA |
| 1-473 | chr18 | 79109737 | 79110200 | + | 706 | -577 | Setbp1 | bXPF_tRA |
| 1-1307 | chr18 | 79145430 | 79145893 | + | 574 | -36270 | Setbp1 | bXPF_tRA |
| 28856 | chr18 | 79145633 | 79146096 | + | 574 | -36473 | Setbp1 | bXPF_tRA |
| 1-1815 | chr18 | 79466947 | 79467410 | + | 838 | -357787 | Setbp1 | bXPF_tRA |
| 1-624 | chr18 | 82462091 | 82462554 | + | 562 | -12801 | Mbp | bXPF_tRA |
| 1-1049 | chr18 | 82487231 | 82487694 | + | 584 | 12339 | Mbp | bXPF_tRA |
| 1-1484 | chr18 | 84851155 | 84851618 | + | 644 | -28 | Cyb5a | bXPF_tRA |
| 1-392 | chr18 | 84851372 | 84851835 | + | 547 | 189 | Cyb5a | bXPF_tRA |
| 1-1896 | chr19 | 3282683 | 3283146 | + | 826 | 13 | Mrpl21 | bXPF_tRA |
| 1-1383 | chr19 | 4125605 | 4126068 | + | 643 | 22 | Aip | bXPF_tRA |
| 1-1545 | chr19 | 4169438 | 4169901 | + | 776 | 5810 | Carns1 | bXPF_tRA |
| 1-1800 | chr19 | 4333901 | 4334364 | + | 690 | -27910 | Grk2 | bXPF_tRA |
| 1-1127 | chr19 | 4928146 | 4928609 | + | 604 | -90 | Dpp3 | bXPF_tRA |
| 1-637 | chr19 | 4988947 | 4989410 | + | 546 | 793 | Npas4 | bXPF_tRA |
| 1-1395 | chr19 | 5366532 | 5366995 | + | 696 | -50 | Eif1ad | bXPF_tRA |
| 1-1442 | chr19 | 5817935 | 5818398 | + | 692 | -15494 | Malat1 | bXPF_tRA |
| 1-251 | chr19 | 5818149 | 5818612 | + | 583 | -15708 | Malat1 | bXPF_tRA |
| 1-107 | chr19 | 5843800 | 5844263 | + | 570 | 1447 | Neat1 | bXPF_tRA |
| 1-1540 | chr19 | 5850039 | 5850502 | + | 740 | -4790 | Neat1 | bXPF_tRA |
| 42736 | chr19 | 5850244 | 5850707 | + | 740 | -4995 | Neat1 | bXPF_tRA |
| 1-1482 | chr19 | 6076987 | 6077450 | + | 634 | -31 | Vps51 | bXPF_tRA |
| 1-1369 | chr19 | 6128055 | 6128518 | + | 744 | -71 | Snx15 | bXPF_tRA |
| 1-617 | chr19 | 6278301 | 6278764 | + | 607 | 1636 | Ehd1 | bXPF_tRA |
| 1-524 | chr19 | 6980218 | 6980681 | + | 585 | -9 | Fkbp2 | bXPF_tRA |
| 23012 | chr19 | 7422818 | 7423281 | + | 1000 | 476 | Mir6991 | bXPF_tRA |
| 14977 | chr19 | 7423023 | 7423486 | + | 882 | 681 | Mir6991 | bXPF_tRA |
| 1-1321 | chr19 | 8741240 | 8741703 | + | 666 | 47 | Stx5a | bXPF_tRA |
| 1-504 | chr19 | 8871171 | 8871634 | + | 595 | -157 | Ubxn1 | bXPF_tRA |
| 1-1431 | chr19 | 8966807 | 8967270 | + | 679 | -3 | Eef1g | bXPF_tRA |

| | | | | | | | | |
|---|---|---|---|---|---|---|---|---|
| 1-616 | chr19 | 8989118 | 8989581 | + | 578 | 65 | Ahnak | bXPF_tRA |
| 1-529 | chr19 | 10100599 | 10101062 | + | 629 | 673 | Fads2 | bXPF_tRA |
| 1-236 | chr19 | 10100799 | 10101262 | + | 629 | 473 | Fads2 | bXPF_tRA |
| 1-955 | chr19 | 10101445 | 10101908 | + | 621 | -173 | Fads2 | bXPF_tRA |
| 1-1721 | chr19 | 10577236 | 10577699 | + | 930 | -81 | Tmem138 | bXPF_tRA |
| 1-1866 | chr19 | 10605254 | 10605717 | + | 726 | -140 | Ddb1 | bXPF_tRA |
| 1-1636 | chr19 | 17837445 | 17837908 | + | 587 | -44 | Pcsk5 | bXPF_tRA |
| 1-1883 | chr19 | 20366022 | 20366485 | + | 955 | 24418 | Anxa1 | bXPF_tRA |
| 1-813 | chr19 | 23273588 | 23274051 | + | 566 | 78 | Smc5 | bXPF_tRA |
| 1-1632 | chr19 | 29648135 | 29648598 | + | 913 | 54 | Ermp1 | bXPF_tRA |
| 1-1755 | chr19 | 31082766 | 31083229 | + | 800 | 156 | Cstf2t | bXPF_tRA |
| 1-1179 | chr19 | 32756253 | 32756716 | + | 605 | -1093 | Pten | bXPF_tRA |
| 1-185 | chr19 | 32756477 | 32756940 | + | 605 | -869 | Pten | bXPF_tRA |
| 1-1771 | chr19 | 34435140 | 34435603 | + | 622 | 39764 | Ch25h | bXPF_tRA |
| 1-350 | chr19 | 36688040 | 36688503 | + | 731 | 1208 | 1500017E21Rik | bXPF_tRA |
| 1-141 | chr19 | 36688247 | 36688710 | + | 731 | 1001 | 1500017E21Rik | bXPF_tRA |
| 1-1116 | chr19 | 36736524 | 36736987 | + | 609 | -151 | Ppp1r3c | bXPF_tRA |
| 1-345 | chr19 | 36736725 | 36737188 | + | 618 | -352 | Ppp1r3c | bXPF_tRA |
| 1-1485 | chr19 | 36918891 | 36919354 | + | 761 | 477 | Fgfbp3 | bXPF_tRA |
| 12420 | chr19 | 38223979 | 38224442 | + | 1000 | -78 | Fra10ac1 | bXPF_tRA |
| 1-1541 | chr19 | 38818998 | 38819461 | + | 754 | 8 | Noc3l | bXPF_tRA |
| 1-771 | chr19 | 40513727 | 40514190 | + | 556 | -146 | Sorbs1 | bXPF_tRA |
| 1-1434 | chr19 | 44293671 | 44294134 | + | 698 | 226 | Scd2 | bXPF_tRA |
| 1-1606 | chr19 | 45363544 | 45364007 | + | 626 | 41 | Btrc | bXPF_tRA |
| 1-1043 | chr19 | 46003378 | 46003841 | + | 638 | 131 | Hps6 | bXPF_tRA |
| 1-1543 | chr19 | 46598808 | 46599271 | + | 648 | -67 | Wbp1l | bXPF_tRA |
| 1-1452 | chr19 | 46689841 | 46690304 | + | 691 | 166 | Borcs7 | bXPF_tRA |
| 1-1221 | chr19 | 47177047 | 47177510 | + | 646 | -1542 | Neurl1a | bXPF_tRA |
| 1-610 | chr19 | 47330711 | 47331174 | + | 623 | -57353 | Mir6995 | bXPF_tRA |
| 1-108 | chr19 | 47330922 | 47331385 | + | 623 | -57564 | Mir6995 | bXPF_tRA |
| 1-483 | chr19 | 53528778 | 53529241 | + | 583 | -309 | Dusp5 | bXPF_tRA |
| 1-558 | chr19 | 53555056 | 53555519 | + | 571 | 25969 | Dusp5 | bXPF_tRA |
| 1-189 | chr19 | 53555316 | 53555779 | + | 571 | 26229 | Dusp5 | bXPF_tRA |
| 1-922 | chr19 | 53731755 | 53732218 | + | 674 | 54680 | Rbm20 | bXPF_tRA |
| 1-352 | chr19 | 53943740 | 53944203 | + | 571 | -335 | Shoc2 | bXPF_tRA |
| 1-283 | chr19 | 53944070 | 53944533 | + | 571 | -5 | Shoc2 | bXPF_tRA |
| 1-1759 | chr19 | 55315991 | 55316454 | + | 853 | 165 | Vti1a | bXPF_tRA |
| 1-1091 | chr2 | 3283765 | 3284228 | + | 589 | -216 | Nmt2 | bXPF_tRA |
| 1-1350 | chr2 | 3712320 | 3712783 | + | 707 | -907 | Fam107b | bXPF_tRA |
| 1-266 | chr2 | 4445595 | 4446058 | + | 561 | 44850 | Frmd4a | bXPF_tRA |
| 1-657 | chr2 | 4445915 | 4446378 | + | 562 | 45170 | Frmd4a | bXPF_tRA |
| 1-865 | chr2 | 6352590 | 6353053 | + | 565 | 85 | Usp6nl | bXPF_tRA |
| 1-687 | chr2 | 7056297 | 7056760 | + | 566 | 24774 | Celf2 | bXPF_tRA |
| 1-1046 | chr2 | 7088987 | 7089450 | + | 620 | -7916 | Celf2 | bXPF_tRA |
| 1-1367 | chr2 | 8781159 | 8781622 | + | 671 | -408125 | 1700061F12Rik | bXPF_tRA |
| 1-904 | chr2 | 18027453 | 18027916 | + | 570 | 435 | Gm17762 | bXPF_tRA |
| 732 | chr2 | 18586865 | 18587328 | + | 646 | -85366 | Commd3 | bXPF_tRA |
| 1-111 | chr2 | 18587072 | 18587535 | + | 646 | -85159 | Commd3 | bXPF_tRA |
| 1-806 | chr2 | 18693954 | 18694417 | + | 570 | -4837 | Spag6 | bXPF_tRA |
| 1-1455 | chr2 | 18767036 | 18767499 | + | 824 | -6405 | Gm3363 | bXPF_tRA |
| 1-526 | chr2 | 19657463 | 19657926 | + | 571 | 203 | Gm3230 | bXPF_tRA |
| 1-713 | chr2 | 22813778 | 22814241 | + | 714 | 39682 | Apbb1ip | bXPF_tRA |
| 1-347 | chr2 | 22814002 | 22814465 | + | 714 | 39906 | Apbb1ip | bXPF_tRA |
| 1-981 | chr2 | 22895274 | 22895737 | + | 609 | -17 | Pdss1 | bXPF_tRA |
| 1-555 | chr2 | 23571513 | 23571976 | + | 636 | 360 | Spopl | bXPF_tRA |
| 1-1681 | chr2 | 24453160 | 24453623 | + | 856 | 22208 | Pax8 | bXPF_tRA |
| 1-282 | chr2 | 24453439 | 24453902 | + | 600 | 21929 | Pax8 | bXPF_tRA |
| 1-563 | chr2 | 25196559 | 25197022 | + | 548 | 23 | Tor4a | bXPF_tRA |
| 1-1071 | chr2 | 25224513 | 25224976 | + | 560 | -42 | Tubb4b | bXPF_tRA |
| 29952 | chr2 | 25224726 | 25225189 | + | 560 | -255 | Tubb4b | bXPF_tRA |
| 5845 | chr2 | 26964290 | 26964753 | + | 648 | -135 | Rexo4 | bXPF_tRA |
| 1-1533 | chr2 | 27442185 | 27442648 | + | 828 | 8214 | Mir7578 | bXPF_tRA |
| 1-212 | chr2 | 27442385 | 27442848 | + | 829 | 8014 | Mir7578 | bXPF_tRA |

| | | | | | | | | |
|---|---|---|---|---|---|---|---|---|
| 1-769 | chr2 | 27539720 | 27540183 | + | 619 | 24804 | Wdr5 | bXPF_tRA |
| 1-925 | chr2 | 28467813 | 28468276 | + | 583 | -22 | Mrps2 | bXPF_tRA |
| 1-795 | chr2 | 30077449 | 30077912 | + | 589 | -1086 | Pkn3 | bXPF_tRA |
| 1-142 | chr2 | 30077649 | 30078112 | + | 589 | -886 | Pkn3 | bXPF_tRA |
| 1-1649 | chr2 | 30133554 | 30134017 | + | 658 | 39 | Tbc1d13 | bXPF_tRA |
| 1-193 | chr2 | 30266508 | 30266971 | + | 649 | 18 | Phyhd1 | bXPF_tRA |
| 1-902 | chr2 | 30415271 | 30415734 | + | 545 | 246 | Crat | bXPF_tRA |
| 1-819 | chr2 | 30870819 | 30871282 | + | 549 | 25683 | Prrx2 | bXPF_tRA |
| 1-1793 | chr2 | 30981953 | 30982416 | + | 743 | -95 | Usp20 | bXPF_tRA |
| 1-802 | chr2 | 31757436 | 31757899 | + | 558 | -2272 | Abl1 | bXPF_tRA |
| 1-1146 | chr2 | 32914875 | 32915338 | + | 573 | 38972 | Fam129b | bXPF_tRA |
| 1-646 | chr2 | 35461981 | 35462444 | + | 544 | -763 | Ggta1 | bXPF_tRA |
| 1-1120 | chr2 | 36136450 | 36136913 | + | 738 | 23 | Rbm18 | bXPF_tRA |
| 1-118 | chr2 | 36136650 | 36137113 | + | 739 | -177 | Rbm18 | bXPF_tRA |
| 1-692 | chr2 | 44145916 | 44146379 | + | 553 | -80823 | Arhgap15os | bXPF_tRA |
| 1-1162 | chr2 | 50296510 | 50296973 | + | 588 | -64 | Mmadhc | bXPF_tRA |
| 1-1279 | chr2 | 51288552 | 51289015 | + | 568 | 46952 | Gm13497 | bXPF_tRA |
| 1-784 | chr2 | 57181887 | 57182350 | + | 561 | 287 | 4930555B11Rik | bXPF_tRA |
| 1-888 | chr2 | 57182124 | 57182587 | + | 545 | 524 | 4930555B11Rik | bXPF_tRA |
| 1-763 | chr2 | 57238199 | 57238662 | + | 555 | 48 | Gpd2 | bXPF_tRA |
| 1-1423 | chr2 | 58160943 | 58161406 | + | 618 | -1052 | Cytip | bXPF_tRA |
| 1-1715 | chr2 | 60689002 | 60689465 | + | 842 | -15540 | Itgb6 | bXPF_tRA |
| 1-1696 | chr2 | 68983492 | 68983955 | + | 810 | 122282 | Cers6 | bXPF_tRA |
| 1-871 | chr2 | 69885351 | 69885814 | + | 656 | 33 | Mettl5 | bXPF_tRA |
| 1-1706 | chr2 | 70661210 | 70661673 | + | 775 | -68 | Gorasp2 | bXPF_tRA |
| 19360 | chr2 | 71644034 | 71644497 | + | 1000 | -75152 | Platr26 | bXPF_tRA |
| 1-1183 | chr2 | 72980617 | 72981080 | + | 617 | -402 | Sp3 | bXPF_tRA |
| 1-1194 | chr2 | 73047101 | 73047564 | + | 613 | -66886 | Sp3 | bXPF_tRA |
| 1-579 | chr2 | 73386469 | 73386932 | + | 655 | -220 | Gpr155 | bXPF_tRA |
| 1-786 | chr2 | 74672732 | 74673195 | + | 556 | -2050 | Hoxd12 | bXPF_tRA |
| 1-183 | chr2 | 74672932 | 74673395 | + | 556 | -1850 | Hoxd12 | bXPF_tRA |
| 1-884 | chr2 | 75685371 | 75685834 | + | 707 | 19061 | Nfe2l2 | bXPF_tRA |
| 1-387 | chr2 | 75685577 | 75686040 | + | 707 | 18855 | Nfe2l2 | bXPF_tRA |
| 1-788 | chr2 | 76647967 | 76648430 | + | 580 | -204 | Prkra | bXPF_tRA |
| 1-840 | chr2 | 80086782 | 80087245 | + | 554 | 41958 | Pde1a | bXPF_tRA |
| 1-534 | chr2 | 80638527 | 80638990 | + | 579 | -54 | Nup35 | bXPF_tRA |
| 1-271 | chr2 | 80638727 | 80639190 | + | 579 | 146 | Nup35 | bXPF_tRA |
| 1-825 | chr2 | 84650535 | 84650998 | + | 617 | -26 | Ctnnd1 | bXPF_tRA |
| 1-220 | chr2 | 84743521 | 84743984 | + | 580 | -2574 | Mir130a | bXPF_tRA |
| 1-328 | chr2 | 84961155 | 84961618 | + | 639 | -19075 | Prg2 | bXPF_tRA |
| 1-1801 | chr2 | 90904383 | 90904846 | + | 680 | 107 | Ndufs3 | bXPF_tRA |
| 1-1093 | chr2 | 91061864 | 91062327 | + | 643 | 8079 | Psmc3 | bXPF_tRA |
| 1-1778 | chr2 | 91202591 | 91203054 | + | 822 | -90 | Acp2 | bXPF_tRA |
| 1-1610 | chr2 | 94010590 | 94011053 | + | 626 | -91 | Alkbh3 | bXPF_tRA |
| 1-1323 | chr2 | 94437889 | 94438352 | + | 721 | 66 | Api5 | bXPF_tRA |
| 1-1568 | chr2 | 102400350 | 102400813 | + | 753 | 319 | Trim44 | bXPF_tRA |
| 1-1200 | chr2 | 102943998 | 102944461 | + | 703 | -42564 | Cd44 | bXPF_tRA |
| 1-1238 | chr2 | 103761101 | 103761564 | + | 641 | -82 | Nat10 | bXPF_tRA |
| 1-1371 | chr2 | 105398790 | 105399253 | + | 645 | 298 | Rcn1 | bXPF_tRA |
| 1-1495 | chr2 | 106003027 | 106003490 | + | 577 | 291 | Dnajc24 | bXPF_tRA |
| 1-380 | chr2 | 109917352 | 109917815 | + | 599 | -64 | Lgr4 | bXPF_tRA |
| 1-198 | chr2 | 109917553 | 109918016 | + | 600 | 137 | Lgr4 | bXPF_tRA |
| 1-1140 | chr2 | 110362920 | 110363383 | + | 604 | -158 | Fibin | bXPF_tRA |
| 1-1124 | chr2 | 114175085 | 114175548 | + | 579 | 23 | Aqr | bXPF_tRA |
| 1-1184 | chr2 | 114201097 | 114201560 | + | 608 | 104 | Zfp770 | bXPF_tRA |
| 1-1676 | chr2 | 117249538 | 117250001 | + | 730 | 30 | Fam98b | bXPF_tRA |
| 1-369 | chr2 | 117249739 | 117250202 | + | 566 | 231 | Fam98b | bXPF_tRA |
| 1-568 | chr2 | 117881978 | 117882441 | + | 563 | 60281 | 4930412B13Rik | bXPF_tRA |
| 1-546 | chr2 | 118533083 | 118533546 | + | 543 | 11273 | Bmf | bXPF_tRA |
| 1-1832 | chr2 | 119046559 | 119047022 | + | 901 | -329 | Knl1 | bXPF_tRA |
| 1-477 | chr2 | 119478080 | 119478543 | + | 627 | -682 | Ino80 | bXPF_tRA |
| 1-1630 | chr2 | 119546992 | 119547455 | + | 618 | 404 | Exd1 | bXPF_tRA |
| 2558 | chr2 | 119662612 | 119663075 | + | 905 | -45 | Ndufaf1 | bXPF_tRA |

| | | | | | | | | |
|---|---|---|---|---|---|---|---|---|
| 1-1804 | chr2 | 119872835 | 119873298 | + | 892 | -24162 | Mga | bXPF_tRA |
| 1-1430 | chr2 | 119897229 | 119897692 | + | 786 | 232 | Mga | bXPF_tRA |
| 1-733 | chr2 | 119972204 | 119972667 | + | 603 | -264 | Mapkbp1 | bXPF_tRA |
| 1-1809 | chr2 | 121413462 | 121413925 | + | 789 | 99 | Catsper2 | bXPF_tRA |
| 10594 | chr2 | 122377117 | 122377580 | + | 1000 | -8430 | Shf | bXPF_tRA |
| 42795 | chr2 | 122377321 | 122377784 | + | 1000 | -8634 | Shf | bXPF_tRA |
| 1-503 | chr2 | 125278505 | 125278968 | + | 603 | 31221 | Dut | bXPF_tRA |
| 1-1026 | chr2 | 125480154 | 125480617 | + | 556 | 26053 | Fbn1 | bXPF_tRA |
| 1-774 | chr2 | 125624868 | 125625331 | + | 541 | 14 | Cep152 | bXPF_tRA |
| 1-1820 | chr2 | 126151982 | 126152445 | + | 845 | 72 | Dtwd1 | bXPF_tRA |
| 1-167 | chr2 | 126152198 | 126152661 | + | 621 | 288 | Dtwd1 | bXPF_tRA |
| 1-615 | chr2 | 126675182 | 126675645 | + | 560 | 179 | Gabpb1 | bXPF_tRA |
| 13516 | chr2 | 126932776 | 126933239 | + | 1000 | 228 | Sppl2a | bXPF_tRA |
| 1-1236 | chr2 | 127247795 | 127248258 | + | 677 | 51 | Tmem127 | bXPF_tRA |
| 1-1209 | chr2 | 127722387 | 127722850 | + | 668 | 7279 | Mall | bXPF_tRA |
| 16072 | chr2 | 128967025 | 128967488 | + | 742 | -146 | Zc3h6 | bXPF_tRA |
| 1-905 | chr2 | 129100735 | 129101198 | + | 640 | -30 | Polr1b | bXPF_tRA |
| 1-160 | chr2 | 129100968 | 129101431 | + | 640 | 203 | Polr1b | bXPF_tRA |
| 1-1348 | chr2 | 130179056 | 130179519 | + | 684 | 77 | Snrpb | bXPF_tRA |
| 1-1391 | chr2 | 130667605 | 130668068 | + | 753 | -5 | Itpa | bXPF_tRA |
| 1-1684 | chr2 | 131392751 | 131393214 | + | 868 | -40090 | Rnf24 | bXPF_tRA |
| 1-943 | chr2 | 131548223 | 131548686 | + | 593 | 13831 | Adra1d | bXPF_tRA |
| 1-370 | chr2 | 132006349 | 132006812 | + | 614 | 23408 | Rassf2 | bXPF_tRA |
| 1-400 | chr2 | 132252997 | 132253460 | + | 568 | -48 | Pcna | bXPF_tRA |
| 1-122 | chr2 | 132253196 | 132253659 | + | 569 | -247 | Pcna | bXPF_tRA |
| 1-508 | chr2 | 134643800 | 134644263 | + | 546 | 90 | Tmx4 | bXPF_tRA |
| 1-415 | chr2 | 134786046 | 134786509 | + | 567 | 113 | Plcb1 | bXPF_tRA |
| 1-1322 | chr2 | 137067630 | 137068093 | + | 548 | 48659 | Jag1 | bXPF_tRA |
| 1-1204 | chr2 | 137114329 | 137114792 | + | 694 | 1960 | Jag1 | bXPF_tRA |
| 1-1712 | chr2 | 144165275 | 144165738 | + | 857 | 23784 | Gm5535 | bXPF_tRA |
| 1-755 | chr2 | 144165479 | 144165942 | + | 799 | 23580 | Gm5535 | bXPF_tRA |
| 1-506 | chr2 | 145759881 | 145760344 | + | 625 | -26004 | Rin2 | bXPF_tRA |
| 1-439 | chr2 | 145760094 | 145760557 | + | 625 | -25791 | Rin2 | bXPF_tRA |
| 1-158 | chr2 | 145887981 | 145888444 | + | 554 | -15029 | Naa20 | bXPF_tRA |
| 1-978 | chr2 | 146959295 | 146959758 | + | 566 | -53534 | Xrn2 | bXPF_tRA |
| 1-1154 | chr2 | 150748932 | 150749395 | + | 647 | 82 | Entpd6 | bXPF_tRA |
| 1-1833 | chr2 | 151493952 | 151494415 | + | 912 | 1 | Nsfl1c | bXPF_tRA |
| 1-929 | chr2 | 152105371 | 152105834 | + | 587 | 78 | Srxn1 | bXPF_tRA |
| 1-432 | chr2 | 152332149 | 152332612 | + | 608 | 45 | Rbck1 | bXPF_tRA |
| 1-1196 | chr2 | 152847716 | 152848179 | + | 601 | -17 | Tpx2 | bXPF_tRA |
| 1-1134 | chr2 | 154613087 | 154613550 | + | 687 | -50 | Zfp341 | bXPF_tRA |
| 1-525 | chr2 | 155692127 | 155692590 | + | 608 | 26 | Trpc4ap | bXPF_tRA |
| 1-1505 | chr2 | 156180104 | 156180567 | + | 787 | -97 | Rbm39 | bXPF_tRA |
| 42767 | chr2 | 156180305 | 156180768 | + | 787 | -298 | Rbm39 | bXPF_tRA |
| 1-1529 | chr2 | 157204366 | 157204829 | + | 659 | -63 | Rbl1 | bXPF_tRA |
| 1-284 | chr2 | 157204568 | 157205031 | + | 660 | -265 | Rbl1 | bXPF_tRA |
| 1-814 | chr2 | 157278992 | 157279455 | + | 556 | 125 | Rpn2 | bXPF_tRA |
| 1-1678 | chr2 | 158064549 | 158065012 | + | 792 | 27017 | 2010009K17Rik | bXPF_tRA |
| 1-1180 | chr2 | 158409467 | 158409930 | + | 625 | -155 | Ralgapb | bXPF_tRA |
| 1-1289 | chr2 | 164432664 | 164433127 | + | 615 | 10293 | Sdc4 | bXPF_tRA |
| 1-708 | chr2 | 164460775 | 164461238 | + | 576 | 35 | Sys1 | bXPF_tRA |
| 1-296 | chr2 | 164461024 | 164461487 | + | 594 | 284 | Sys1 | bXPF_tRA |
| 1-670 | chr2 | 164497267 | 164497730 | + | 559 | -27 | Pigt | bXPF_tRA |
| 1-1136 | chr2 | 164785575 | 164786038 | + | 596 | -215 | Snx21 | bXPF_tRA |
| 1-1280 | chr2 | 165914279 | 165914742 | + | 644 | -29672 | Zmynd8 | bXPF_tRA |
| 1-297 | chr2 | 165914482 | 165914945 | + | 644 | -29875 | Zmynd8 | bXPF_tRA |
| 1-618 | chr2 | 166426829 | 166427292 | + | 603 | -20391 | 5031425F14Rik | bXPF_tRA |
| 1-311 | chr2 | 166427028 | 166427491 | + | 602 | -20192 | 5031425F14Rik | bXPF_tRA |
| 1-1407 | chr2 | 166463417 | 166463880 | + | 764 | 16197 | 5031425F14Rik | bXPF_tRA |
| 1-1857 | chr2 | 166793495 | 166793958 | + | 1000 | -41 | Trp53rkb | bXPF_tRA |
| 1-306 | chr2 | 166793696 | 166794159 | + | 1000 | 160 | Trp53rkb | bXPF_tRA |
| 1-831 | chr2 | 167188854 | 167189317 | + | 649 | -267 | Kcnb1 | bXPF_tRA |
| 1-640 | chr2 | 167765070 | 167765533 | + | 563 | 74124 | A530013C23Rik | bXPF_tRA |

| | | | | | | | | |
|---|---|---|---|---|---|---|---|---|
| 1-1157 | chr2 | 168114913 | 168115376 | + | 743 | 34140 | Pard6b | bXPF_tRA |
| 1-1682 | chr2 | 170319073 | 170319536 | + | 757 | -37057 | Bcas1os2 | bXPF_tRA |
| 1-987 | chr2 | 173335801 | 173336264 | + | 545 | -59499 | Pmepa1 | bXPF_tRA |
| 1-430 | chr2 | 173337285 | 173337748 | + | 685 | -60983 | Pmepa1 | bXPF_tRA |
| 1-186 | chr2 | 173337591 | 173338054 | + | 685 | -61289 | Pmepa1 | bXPF_tRA |
| 22282 | chr2 | 174076193 | 174076324 | + | 1000 | -793 | Stx16 | bXPF_tRA |
| 1-290 | chr2 | 174110604 | 174111067 | + | 545 | 484 | Npepl1 | bXPF_tRA |
| 1-820 | chr2 | 174932707 | 174933170 | + | 592 | 134831 | Gm14393 | bXPF_tRA |
| 1-1492 | chr2 | 181520205 | 181520668 | + | 787 | -49 | Dnajc5 | bXPF_tRA |
| 1-253 | chr3 | 5218337 | 5218800 | + | 565 | 14 | Zfhx4 | bXPF_tRA |
| 1-1072 | chr3 | 5576007 | 5576470 | + | 1000 | 10 | Pex2 | bXPF_tRA |
| 1-373 | chr3 | 5576208 | 5576671 | + | 602 | -191 | Pex2 | bXPF_tRA |
| 1-1030 | chr3 | 8923358 | 8923821 | + | 560 | 268 | Mrps28 | bXPF_tRA |
| 1-1010 | chr3 | 9609950 | 9610413 | + | 570 | -96 | Zfp704 | bXPF_tRA |
| 1-847 | chr3 | 9715542 | 9716005 | + | 541 | -105688 | Zfp704 | bXPF_tRA |
| 1-1830 | chr3 | 18452172 | 18452635 | + | 860 | 28074 | 4930433B08Rik | bXPF_tRA |
| 1-313 | chr3 | 18452373 | 18452836 | + | 638 | 27873 | 4930433B08Rik | bXPF_tRA |
| 1-1264 | chr3 | 19628586 | 19629049 | + | 831 | -15643 | Trim55 | bXPF_tRA |
| 1-1836 | chr3 | 21935863 | 21936326 | + | 1000 | -140558 | Tbl1xr1 | bXPF_tRA |
| 1-291 | chr3 | 21936068 | 21936531 | + | 562 | -140353 | Tbl1xr1 | bXPF_tRA |
| 1-1438 | chr3 | 27182701 | 27183164 | + | 559 | -72 | Nceh1 | bXPF_tRA |
| 8037 | chr3 | 30514960 | 30515423 | + | 961 | -5704 | Mecom | bXPF_tRA |
| 1-1037 | chr3 | 30995226 | 30995689 | + | 563 | -314 | Prkci | bXPF_tRA |
| 1-375 | chr3 | 32510420 | 32510883 | + | 585 | 59 | Zfp639 | bXPF_tRA |
| 1-1486 | chr3 | 33799805 | 33800268 | + | 709 | -147 | Ttc14 | bXPF_tRA |
| 1-163 | chr3 | 33800004 | 33800467 | + | 566 | 52 | Ttc14 | bXPF_tRA |
| 1-1652 | chr3 | 34081079 | 34081542 | + | 784 | 44 | Dnajc19 | bXPF_tRA |
| 1-899 | chr3 | 36449547 | 36450010 | + | 578 | 25746 | Mir7009 | bXPF_tRA |
| 1-1275 | chr3 | 41185418 | 41185881 | + | 540 | -102603 | Pgrmc2 | bXPF_tRA |
| 1-1698 | chr3 | 49642331 | 49642794 | + | 838 | 114754 | Pcdh18 | bXPF_tRA |
| 1-117 | chr3 | 49642535 | 49642998 | + | 839 | 114550 | Pcdh18 | bXPF_tRA |
| 14611 | chr3 | 51358114 | 51358577 | + | 729 | -17701 | Elf2 | bXPF_tRA |
| 1-1829 | chr3 | 51359166 | 51359629 | + | 631 | -18753 | Elf2 | bXPF_tRA |
| 1-1481 | chr3 | 51483863 | 51484326 | + | 711 | 128 | Rab33b | bXPF_tRA |
| 1-696 | chr3 | 53075489 | 53075952 | + | 541 | 34173 | Lhfp | bXPF_tRA |
| 27395 | chr3 | 53075703 | 53076166 | + | 541 | 34387 | Lhfp | bXPF_tRA |
| 1-1446 | chr3 | 58372301 | 58372764 | + | 634 | -42188 | Tsc22d2 | bXPF_tRA |
| 1-1449 | chr3 | 58416653 | 58417116 | + | 601 | 2164 | Tsc22d2 | bXPF_tRA |
| 1-1005 | chr3 | 58576668 | 58577131 | + | 546 | 241 | Selenot | bXPF_tRA |
| 1-749 | chr3 | 65958131 | 65958594 | + | 637 | -137 | Ccnl1 | bXPF_tRA |
| 1-1378 | chr3 | 66373854 | 66374317 | + | 569 | -77248 | Veph1 | bXPF_tRA |
| 1-1226 | chr3 | 66985525 | 66985988 | + | 646 | 84 | Rsrc1 | bXPF_tRA |
| 1-1216 | chr3 | 75868867 | 75869330 | + | 604 | 87851 | Golim4 | bXPF_tRA |
| 1-414 | chr3 | 79531867 | 79532330 | + | 573 | 35581 | Fnip2 | bXPF_tRA |
| 1-1749 | chr3 | 79977571 | 79978034 | + | 690 | 91872 | Fam198b | bXPF_tRA |
| 1-779 | chr3 | 79989288 | 79989751 | + | 630 | 103589 | Fam198b | bXPF_tRA |
| 1-604 | chr3 | 81058356 | 81058819 | + | 663 | 22171 | Pdgfc | bXPF_tRA |
| 1-1067 | chr3 | 84814223 | 84814686 | + | 581 | -1123 | Fbxw7 | bXPF_tRA |
| 1-1480 | chr3 | 84952101 | 84952564 | + | 702 | 121 | Fbxw7 | bXPF_tRA |
| 1-796 | chr3 | 87885260 | 87885723 | + | 556 | 71 | Prcc | bXPF_tRA |
| 1-1467 | chr3 | 89419424 | 89419887 | + | 651 | 1104 | Shc1 | bXPF_tRA |
| 14611 | chr3 | 89419626 | 89420089 | + | 651 | 1306 | Shc1 | bXPF_tRA |
| 1-1114 | chr3 | 90612864 | 90613327 | + | 654 | 201 | S100a6 | bXPF_tRA |
| 1-1613 | chr3 | 93568518 | 93568981 | + | 676 | 13632 | S100a10 | bXPF_tRA |
| 1-1653 | chr3 | 94309933 | 94310396 | + | 849 | 32 | Them4 | bXPF_tRA |
| 1-203 | chr3 | 94310141 | 94310604 | + | 851 | 240 | Them4 | bXPF_tRA |
| 1-1483 | chr3 | 95110841 | 95111304 | + | 727 | 30 | Vps72 | bXPF_tRA |
| 1-1693 | chr3 | 95217646 | 95218109 | + | 598 | 65 | Gabpb2 | bXPF_tRA |
| 1-500 | chr3 | 95228186 | 95228649 | + | 552 | -7 | Cdc42se1 | bXPF_tRA |
| 1-1084 | chr3 | 95658501 | 95658964 | + | 649 | 11 | Mcl1 | bXPF_tRA |
| 1-998 | chr3 | 95929046 | 95929509 | + | 551 | 20 | Anp32e | bXPF_tRA |
| 1-1158 | chr3 | 96635332 | 96635795 | + | 634 | 133 | Pex11b | bXPF_tRA |
| 1-1624 | chr3 | 97900966 | 97901429 | + | 717 | -30 | Sec22b | bXPF_tRA |

| | | | | | | | | |
|---|---|---|---|---|---|---|---|---|
| 1-499 | chr3 | 98990715 | 98991178 | + | 669 | -58876 | Gm12440 | bXPF_tRA |
| 1-585 | chr3 | 104864109 | 104864572 | + | 564 | 165 | Capza1 | bXPF_tRA |
| 1-896 | chr3 | 105687290 | 105687753 | + | 583 | 50 | Ddx20 | bXPF_tRA |
| 1-1705 | chr3 | 106547424 | 106547887 | + | 1000 | -143 | Dram2 | bXPF_tRA |
| 1-714 | chr3 | 107631310 | 107631773 | + | 625 | 169 | Strip1 | bXPF_tRA |
| 1-1241 | chr3 | 108209968 | 108210431 | + | 728 | 328 | Atxn7l2 | bXPF_tRA |
| 1-426 | chr3 | 108210170 | 108210633 | + | 679 | 126 | Atxn7l2 | bXPF_tRA |
| 1-1381 | chr3 | 108389164 | 108389627 | + | 641 | 5557 | Psrc1 | bXPF_tRA |
| 1-1295 | chr3 | 108562193 | 108562656 | + | 572 | 42 | Tmem167b | bXPF_tRA |
| 1-151 | chr3 | 108562393 | 108562856 | + | 572 | -158 | Tmem167b | bXPF_tRA |
| 1-920 | chr3 | 108770643 | 108771106 | + | 553 | 31216 | Aknad1 | bXPF_tRA |
| 1-136 | chr3 | 109122972 | 109123435 | + | 550 | 54 | Slc25a24 | bXPF_tRA |
| 42826 | chr3 | 109123188 | 109123651 | + | 550 | 270 | Slc25a24 | bXPF_tRA |
| 1-1267 | chr3 | 113629776 | 113630239 | + | 848 | 142 | Rnpc3 | bXPF_tRA |
| 33970 | chr3 | 113629980 | 113630443 | + | 848 | -62 | Rnpc3 | bXPF_tRA |
| 1-418 | chr3 | 115028237 | 115028700 | + | 583 | -52497 | Olfm3 | bXPF_tRA |
| 1-1619 | chr3 | 116712120 | 116712583 | + | 575 | -71 | Slc35a3 | bXPF_tRA |
| 1-1597 | chr3 | 118431337 | 118431800 | + | 682 | -2289 | Mir137 | bXPF_tRA |
| 1-1523 | chr3 | 121577023 | 121577486 | + | 758 | -44910 | Slc44a3 | bXPF_tRA |
| 1-1641 | chr3 | 122122071 | 122122534 | + | 610 | 77842 | Abca4 | bXPF_tRA |
| 1-1875 | chr3 | 122292763 | 122293226 | + | 662 | 709 | Mir760 | bXPF_tRA |
| 1-701 | chr3 | 123508094 | 123508557 | + | 584 | 11 | Snhg8 | bXPF_tRA |
| 1-1797 | chr3 | 124320956 | 124321419 | + | 858 | 150 | Tram1l1 | bXPF_tRA |
| 1-1096 | chr3 | 126505712 | 126506175 | + | 545 | -91008 | Camk2d | bXPF_tRA |
| 1-295 | chr3 | 126597112 | 126597575 | + | 564 | 392 | Camk2d | bXPF_tRA |
| 1-1767 | chr3 | 126831366 | 126831829 | + | 771 | 111789 | Ank2 | bXPF_tRA |
| 1-1298 | chr3 | 129532127 | 129532590 | + | 577 | -28 | Elovl6 | bXPF_tRA |
| 29221 | chr3 | 129532336 | 129532799 | + | 577 | 181 | Elovl6 | bXPF_tRA |
| 1-650 | chr3 | 131490223 | 131490686 | + | 591 | -74314 | Papss1 | bXPF_tRA |
| 1-1831 | chr3 | 132683762 | 132684225 | + | 855 | -114 | Aimp1 | bXPF_tRA |
| 1-1631 | chr3 | 134033843 | 134034306 | + | 646 | -202421 | Cxxc4 | bXPF_tRA |
| 1-531 | chr3 | 135437844 | 135438307 | + | 622 | 590 | 4930539J05Rik | bXPF_tRA |
| 1-510 | chr3 | 135438043 | 135438506 | + | 623 | 391 | 4930539J05Rik | bXPF_tRA |
| 1-1025 | chr3 | 136150767 | 136151230 | + | 557 | 175068 | Bank1 | bXPF_tRA |
| 1-1057 | chr3 | 136725157 | 136725620 | + | 593 | 55322 | Ppp3ca | bXPF_tRA |
| 1-685 | chr3 | 138864265 | 138864728 | + | 553 | 122288 | Tspan5 | bXPF_tRA |
| 11689 | chr3 | 138864465 | 138864928 | + | 553 | 122488 | Tspan5 | bXPF_tRA |
| 1-1794 | chr3 | 142169080 | 142169543 | + | 873 | -83 | Bmpr1b | bXPF_tRA |
| 1-1855 | chr3 | 142395423 | 142395886 | + | 568 | 42 | Pdlim5 | bXPF_tRA |
| 1-1723 | chr3 | 142764987 | 142765450 | + | 971 | -29 | Gtf2b | bXPF_tRA |
| 1-298 | chr3 | 142765188 | 142765651 | + | 617 | 172 | Gtf2b | bXPF_tRA |
| 1-1366 | chr3 | 144570417 | 144570880 | + | 573 | 221 | Selenof | bXPF_tRA |
| 1-1401 | chr3 | 144720402 | 144720865 | + | 557 | -298 | Sh3glb1 | bXPF_tRA |
| 1-1675 | chr3 | 145649770 | 145650233 | + | 1000 | -16 | Cyr61 | bXPF_tRA |
| 1-997 | chr3 | 145758756 | 145759219 | + | 750 | 295 | Ddah1 | bXPF_tRA |
| 1-1291 | chr3 | 145886025 | 145886488 | + | 575 | -38006 | Bcl10 | bXPF_tRA |
| 1-1125 | chr3 | 145923945 | 145924408 | + | 552 | -86 | Bcl10 | bXPF_tRA |
| 1-643 | chr3 | 145937820 | 145938283 | + | 555 | 19 | 2410004B18Rik | bXPF_tRA |
| 1-133 | chr3 | 145988260 | 145988723 | + | 665 | 621 | Syde2 | bXPF_tRA |
| 1-153 | chr3 | 145988474 | 145988937 | + | 665 | 835 | Syde2 | bXPF_tRA |
| 1-440 | chr3 | 147445792 | 147446255 | + | 807 | 593656 | Ttll7 | bXPF_tRA |
| 1-1735 | chr3 | 147853313 | 147853776 | + | 924 | 1001177 | Ttll7 | bXPF_tRA |
| 1-1352 | chr3 | 150610521 | 150610984 | + | 611 | -827130 | Adgrl4 | bXPF_tRA |
| 1-382 | chr3 | 150610721 | 150611184 | + | 611 | -826930 | Adgrl4 | bXPF_tRA |
| 1-1059 | chr3 | 153944212 | 153944675 | + | 674 | 200 | Acadm | bXPF_tRA |
| 1-1570 | chr3 | 154692269 | 154692732 | + | 758 | -18633 | Erich3 | bXPF_tRA |
| 1-944 | chr3 | 158036335 | 158036798 | + | 565 | 73 | Srsf11 | bXPF_tRA |
| 1-869 | chr3 | 159865789 | 159866252 | + | 586 | 26325 | Wls | bXPF_tRA |
| 1-990 | chr4 | 3835306 | 3835769 | + | 696 | 118 | Rps20 | bXPF_tRA |
| 1-238 | chr4 | 3835520 | 3835983 | + | 697 | -96 | Rps20 | bXPF_tRA |
| 1-1520 | chr4 | 4792970 | 4793433 | + | 637 | 105 | Impad1 | bXPF_tRA |
| 1-1164 | chr4 | 6190791 | 6191254 | + | 732 | -83 | Ubxn2b | bXPF_tRA |
| 21186 | chr4 | 6190996 | 6191459 | + | 732 | 122 | Ubxn2b | bXPF_tRA |

| | | | | | | | | |
|---|---|---|---|---|---|---|---|---|
| 1-1564 | chr4 | 7560274 | 7560737 | + | 776 | -183 | 8430436N08Rik | bXPF_tRA |
| 1-1773 | chr4 | 9668906 | 9669369 | + | 827 | 25 | Asph | bXPF_tRA |
| 1-436 | chr4 | 11175926 | 11176389 | + | 557 | -15194 | Ccne2 | bXPF_tRA |
| 16803 | chr4 | 11321997 | 11322460 | + | 1000 | -97 | Dpy19l4 | bXPF_tRA |
| 1-665 | chr4 | 12906760 | 12907223 | + | 581 | 154 | Triqk | bXPF_tRA |
| 1-671 | chr4 | 14826255 | 14826718 | + | 560 | 101 | Otud6b | bXPF_tRA |
| 1-1328 | chr4 | 15957975 | 15958438 | + | 619 | 239 | Nbn | bXPF_tRA |
| 1-1269 | chr4 | 19574876 | 19575339 | + | 579 | 41 | Rmdn1 | bXPF_tRA |
| 1-1013 | chr4 | 21879557 | 21880020 | + | 729 | 113 | Coq3 | bXPF_tRA |
| 1-1373 | chr4 | 25281586 | 25282049 | + | 669 | 4 | Ufl1 | bXPF_tRA |
| 1-785 | chr4 | 26259475 | 26259938 | + | 579 | 86946 | Manea | bXPF_tRA |
| 1-791 | chr4 | 31963703 | 31964166 | + | 572 | -173 | Map3k7 | bXPF_tRA |
| 15342 | chr4 | 31963917 | 31964380 | + | 572 | 41 | Map3k7 | bXPF_tRA |
| 1-1891 | chr4 | 32179881 | 32180344 | + | 986 | 216005 | Map3k7 | bXPF_tRA |
| 1-882 | chr4 | 32250849 | 32251312 | + | 561 | -166355 | Bach2 | bXPF_tRA |
| 1-1101 | chr4 | 34614667 | 34615130 | + | 660 | 44 | Orc3 | bXPF_tRA |
| 1-152 | chr4 | 34614869 | 34615332 | + | 661 | 142 | Rars2 | bXPF_tRA |
| 5480 | chr4 | 34687117 | 34687580 | + | 1000 | 90 | Slc35a1 | bXPF_tRA |
| 1-942 | chr4 | 34886505 | 34886968 | + | 748 | -3788 | Zfp292 | bXPF_tRA |
| 1-475 | chr4 | 34886762 | 34887225 | + | 749 | -4045 | Zfp292 | bXPF_tRA |
| 1-1358 | chr4 | 40757434 | 40757897 | + | 565 | 220 | Smu1 | bXPF_tRA |
| 1-589 | chr4 | 40948185 | 40948648 | + | 615 | -122 | Bag1 | bXPF_tRA |
| 1-427 | chr4 | 41314583 | 41315046 | + | 659 | 87 | Dcaf12 | bXPF_tRA |
| 1-600 | chr4 | 43381611 | 43382074 | + | 565 | -140 | Rusc2 | bXPF_tRA |
| 1-816 | chr4 | 43398695 | 43399158 | + | 611 | -7576 | Rusc2 | bXPF_tRA |
| 1-140 | chr4 | 43398925 | 43399388 | + | 611 | -7346 | Rusc2 | bXPF_tRA |
| 1-487 | chr4 | 43406346 | 43406809 | + | 576 | 75 | Rusc2 | bXPF_tRA |
| 1-1008 | chr4 | 43492684 | 43493147 | + | 702 | 144 | Rmrp | bXPF_tRA |
| 1-505 | chr4 | 43492908 | 43493371 | + | 702 | -80 | Rmrp | bXPF_tRA |
| 1-1646 | chr4 | 44072454 | 44072917 | + | 745 | -12 | Gne | bXPF_tRA |
| 15342 | chr4 | 45341770 | 45342233 | + | 1000 | -100 | Dcaf10 | bXPF_tRA |
| 1-1634 | chr4 | 47504901 | 47505364 | + | 789 | 30471 | Sec61b | bXPF_tRA |
| 1-1419 | chr4 | 53158594 | 53159057 | + | 636 | 1070 | Abca1 | bXPF_tRA |
| 1-775 | chr4 | 55349599 | 55350062 | + | 597 | -212 | Rad23b | bXPF_tRA |
| 1-1476 | chr4 | 55718418 | 55718881 | + | 653 | -186174 | Klf4 | bXPF_tRA |
| 1-1377 | chr4 | 58206615 | 58207078 | + | 585 | -250 | Svep1 | bXPF_tRA |
| 1-416 | chr4 | 59323283 | 59323746 | + | 766 | 63463 | Gm12596 | bXPF_tRA |
| 1-681 | chr4 | 59783776 | 59784239 | + | 574 | -152 | Inip | bXPF_tRA |
| 1-1501 | chr4 | 62470719 | 62471182 | + | 756 | -55 | Wdr31 | bXPF_tRA |
| 1-1677 | chr4 | 62519702 | 62520165 | + | 746 | -24 | Alad | bXPF_tRA |
| 1-1324 | chr4 | 62525075 | 62525538 | + | 540 | -63 | 4933430I17Rik | bXPF_tRA |
| 1-486 | chr4 | 64901355 | 64901818 | + | 568 | -222588 | Pappa | bXPF_tRA |
| 1-848 | chr4 | 65420578 | 65421041 | + | 540 | -184177 | Trim32 | bXPF_tRA |
| 1-1468 | chr4 | 70410199 | 70410662 | + | 621 | 5 | Cdk5rap2 | bXPF_tRA |
| 43070 | chr4 | 70410399 | 70410862 | + | 622 | -195 | Cdk5rap2 | bXPF_tRA |
| 1-556 | chr4 | 72946434 | 72946897 | + | 546 | -94031 | Aldoart1 | bXPF_tRA |
| 1-495 | chr4 | 76449795 | 76450258 | + | 624 | -1171740 | Tmem261 | bXPF_tRA |
| 1-211 | chr4 | 76449996 | 76450459 | + | 624 | -1171941 | Tmem261 | bXPF_tRA |
| 1-1032 | chr4 | 80911821 | 80912284 | + | 586 | 1366 | Lurap1l | bXPF_tRA |
| 1-1121 | chr4 | 83417310 | 83417773 | + | 568 | -203 | Snapc3 | bXPF_tRA |
| 1-1747 | chr4 | 88033055 | 88033518 | + | 694 | 121 | Mllt3 | bXPF_tRA |
| 31048 | chr4 | 88033256 | 88033719 | + | 693 | -80 | Mllt3 | bXPF_tRA |
| 1-1020 | chr4 | 89128174 | 89128637 | + | 669 | -8965 | Mtap | bXPF_tRA |
| 1-1499 | chr4 | 89294413 | 89294876 | + | 649 | -25 | Cdkn2a | bXPF_tRA |
| 1-289 | chr4 | 89294625 | 89295088 | + | 649 | -237 | Cdkn2a | bXPF_tRA |
| 1-1811 | chr4 | 89310705 | 89311168 | + | 799 | 96 | Cdkn2b | bXPF_tRA |
| 1-1214 | chr4 | 98511292 | 98511755 | + | 561 | -35111 | Patj | bXPF_tRA |
| 1-279 | chr4 | 98511491 | 98511954 | + | 561 | -34912 | Patj | bXPF_tRA |
| 1-1577 | chr4 | 99829009 | 99829472 | + | 705 | -122 | Itgb3bp | bXPF_tRA |
| 1-1798 | chr4 | 101202685 | 101203148 | + | 837 | 62366 | Jak1 | bXPF_tRA |
| 1-1223 | chr4 | 105157142 | 105157605 | + | 604 | 26 | Plpp3 | bXPF_tRA |
| 1-1260 | chr4 | 105898370 | 105898833 | + | 622 | -417612 | Usp24 | bXPF_tRA |
| 1-1082 | chr4 | 106678767 | 106679230 | + | 635 | -54 | Ttc4 | bXPF_tRA |

| | | | | | | | | |
|---|---|---|---|---|---|---|---|---|
| 1-867 | chr4 | 108619789 | 108620252 | + | 544 | 64 | Cc2d1b | bXPF_tRA |
| 1-247 | chr4 | 109666466 | 109666929 | + | 606 | 59 | Cdkn2c | bXPF_tRA |
| 36161 | chr4 | 109666756 | 109667219 | + | 606 | -231 | Cdkn2c | bXPF_tRA |
| 1-1095 | chr4 | 109676260 | 109676723 | + | 565 | -136 | Faf1 | bXPF_tRA |
| 1-1001 | chr4 | 111719886 | 111720349 | + | 549 | 107 | Spata6 | bXPF_tRA |
| 1-1414 | chr4 | 115784748 | 115785211 | + | 683 | 165 | Atpaf1 | bXPF_tRA |
| 1-106 | chr4 | 115784950 | 115785413 | + | 683 | 367 | Atpaf1 | bXPF_tRA |
| 1-1361 | chr4 | 116053484 | 116053947 | + | 783 | 161 | Nsun4 | bXPF_tRA |
| 28491 | chr4 | 116053690 | 116054153 | + | 782 | -45 | Nsun4 | bXPF_tRA |
| 1-632 | chr4 | 116555725 | 116556188 | + | 545 | -13 | Tmem69 | bXPF_tRA |
| 1-1432 | chr4 | 116685424 | 116685887 | + | 612 | 56 | Prdx1 | bXPF_tRA |
| 1-1362 | chr4 | 117682123 | 117682586 | + | 563 | -129 | Dmap1 | bXPF_tRA |
| 1-766 | chr4 | 117997595 | 117998058 | + | 548 | -68063 | Artn | bXPF_tRA |
| 1-1181 | chr4 | 118134625 | 118135088 | + | 555 | 90 | St3gal3 | bXPF_tRA |
| 1-200 | chr4 | 118134852 | 118135315 | + | 555 | -137 | St3gal3 | bXPF_tRA |
| 1-1147 | chr4 | 119105729 | 119106192 | + | 612 | -2785 | Slc2a1 | bXPF_tRA |
| 1-1847 | chr4 | 119539114 | 119539577 | + | 965 | -316 | Foxj3 | bXPF_tRA |
| 1-412 | chr4 | 120380568 | 120381031 | + | 557 | -24482 | Scmh1 | bXPF_tRA |
| 1-1766 | chr4 | 121097933 | 121098396 | + | 805 | 79 | Zmpste24 | bXPF_tRA |
| 1-177 | chr4 | 121098133 | 121098596 | + | 804 | -121 | Zmpste24 | bXPF_tRA |
| 1-900 | chr4 | 122859266 | 122859729 | + | 577 | 23270 | Ppt1 | bXPF_tRA |
| 1-157 | chr4 | 122859476 | 122859939 | + | 577 | 23480 | Ppt1 | bXPF_tRA |
| 1-622 | chr4 | 122885714 | 122886177 | + | 598 | 112 | Cap1 | bXPF_tRA |
| 1-1115 | chr4 | 123535135 | 123535598 | + | 584 | -81054 | Mir6398 | bXPF_tRA |
| 1-982 | chr4 | 123917054 | 123917517 | + | 638 | -148 | Rragc | bXPF_tRA |
| 1-1770 | chr4 | 124708690 | 124709153 | + | 819 | -5940 | Sf3a3 | bXPF_tRA |
| 1-1701 | chr4 | 124714574 | 124715037 | + | 688 | -56 | Sf3a3 | bXPF_tRA |
| 1-1258 | chr4 | 125066332 | 125066795 | + | 709 | -130 | Snip1 | bXPF_tRA |
| 1-829 | chr4 | 126046755 | 126047218 | + | 581 | 58 | Mrps15 | bXPF_tRA |
| 1-734 | chr4 | 126150143 | 126150606 | + | 547 | 2371 | Eva1b | bXPF_tRA |
| 1-836 | chr4 | 126321484 | 126321947 | + | 710 | -12 | Adprhl2 | bXPF_tRA |
| 1-727 | chr4 | 126677257 | 126677720 | + | 591 | -155 | Psmb2 | bXPF_tRA |
| 1-1372 | chr4 | 127243595 | 127244058 | + | 679 | 42 | Smim12 | bXPF_tRA |
| 1-522 | chr4 | 130273612 | 130274075 | + | 541 | 1377 | Serinc2 | bXPF_tRA |
| 1-1228 | chr4 | 132270054 | 132270517 | + | 679 | -99 | Rnu11 | bXPF_tRA |
| 1-1342 | chr4 | 132509623 | 132510086 | + | 599 | 602 | Sesn2 | bXPF_tRA |
| 1-1819 | chr4 | 133277610 | 133278073 | + | 920 | -51 | Tmem222 | bXPF_tRA |
| 1-808 | chr4 | 133545743 | 133546206 | + | 547 | 53 | Nudc | bXPF_tRA |
| 1-1007 | chr4 | 133729543 | 133730006 | + | 559 | -12666 | Mir7227 | bXPF_tRA |
| 1-492 | chr4 | 133754236 | 133754699 | + | 589 | -856 | Arid1a | bXPF_tRA |
| 19725 | chr4 | 133754440 | 133754903 | + | 590 | -1060 | Arid1a | bXPF_tRA |
| 1-114 | chr4 | 134186420 | 134186883 | + | 624 | 434 | Cep85 | bXPF_tRA |
| 1-278 | chr4 | 135272569 | 135273032 | + | 616 | -40 | Clic4 | bXPF_tRA |
| 1-1198 | chr4 | 136021508 | 136021971 | + | 562 | -90 | Eloa | bXPF_tRA |
| 1-1370 | chr4 | 136053188 | 136053651 | + | 649 | -48 | Rpl11 | bXPF_tRA |
| 1-441 | chr4 | 136142955 | 136143418 | + | 552 | -636 | Id3 | bXPF_tRA |
| 1-1111 | chr4 | 136781704 | 136782167 | + | 604 | 54077 | Ephb2 | bXPF_tRA |
| 1-1555 | chr4 | 138304538 | 138305001 | + | 664 | 31 | Ddost | bXPF_tRA |
| 1-1225 | chr4 | 140699500 | 140699963 | + | 742 | -1742 | Rcc2 | bXPF_tRA |
| 1-924 | chr4 | 141060508 | 141060971 | + | 648 | -194 | Crocc | bXPF_tRA |
| 1-1232 | chr4 | 141425695 | 141426158 | + | 638 | 5147 | Hspb7 | bXPF_tRA |
| 1-1119 | chr4 | 143278334 | 143278797 | + | 684 | 20999 | Pdpn | bXPF_tRA |
| 1-1637 | chr4 | 144949719 | 144950182 | + | 746 | 56813 | Dhrs3 | bXPF_tRA |
| 1-184 | chr4 | 144949918 | 144950381 | + | 600 | 57012 | Dhrs3 | bXPF_tRA |
| 1-407 | chr4 | 144983694 | 144984157 | + | 562 | 90788 | Dhrs3 | bXPF_tRA |
| 2193 | chr4 | 144999763 | 145000226 | + | 1000 | 106857 | Dhrs3 | bXPF_tRA |
| 1-1640 | chr4 | 144999999 | 145000462 | + | 935 | 107093 | Dhrs3 | bXPF_tRA |
| 1-1665 | chr4 | 145194748 | 145195211 | + | 554 | 26 | Vps13d | bXPF_tRA |
| 1-1837 | chr4 | 145239421 | 145239884 | + | 691 | 7218 | Tnfrsf1b | bXPF_tRA |
| 1-880 | chr4 | 148444413 | 148444876 | + | 569 | 107 | Ubiad1 | bXPF_tRA |
| 1-1680 | chr4 | 149954894 | 149955357 | + | 586 | -119 | Spsb1 | bXPF_tRA |
| 1-196 | chr4 | 149955097 | 149955560 | + | 586 | -322 | Spsb1 | bXPF_tRA |
| 1-530 | chr4 | 150008480 | 150008943 | + | 662 | 312 | H6pd | bXPF_tRA |

| | | | | | | | | |
|---|---|---|---|---|---|---|---|---|
| 17533 | chr4 | 150008680 | 150009143 | + | 662 | 112 | H6pd | bXPF_tRA |
| 1-887 | chr4 | 150371507 | 150371970 | + | 586 | 89822 | Rere | bXPF_tRA |
| 1-261 | chr4 | 150371714 | 150372177 | + | 586 | 90029 | Rere | bXPF_tRA |
| 1-1304 | chr4 | 151411644 | 151412107 | + | 614 | 322305 | Gm13090 | bXPF_tRA |
| 1-1741 | chr4 | 152283078 | 152283541 | + | 567 | 8353 | Hes3 | bXPF_tRA |
| 1-398 | chr4 | 153956873 | 153957336 | + | 563 | -133 | A430005L14Rik | bXPF_tRA |
| 1-1803 | chr4 | 153974878 | 153975341 | + | 826 | -28 | Dffb | bXPF_tRA |
| 1-1814 | chr4 | 155803307 | 155803770 | + | 814 | -80 | Mrpl20 | bXPF_tRA |
| 1-1780 | chr4 | 155943432 | 155943895 | + | 825 | -150 | Ube2j2 | bXPF_tRA |
| 1-861 | chr4 | 156109624 | 156110087 | + | 585 | -143 | 9430015G10Rik | bXPF_tRA |
| 34335 | chr4 | 156109829 | 156110292 | + | 585 | 62 | 9430015G10Rik | bXPF_tRA |
| 1-462 | chr5 | 3543554 | 3544017 | + | 563 | -48 | Fam133b | bXPF_tRA |
| 1-170 | chr5 | 3543766 | 3544229 | + | 563 | 164 | Fam133b | bXPF_tRA |
| 1-1821 | chr5 | 5749135 | 5749598 | + | 656 | -49 | Steap1 | bXPF_tRA |
| 2923 | chr5 | 12910765 | 12911228 | + | 690 | -485788 | Sema3a | bXPF_tRA |
| 1-1615 | chr5 | 17556145 | 17556608 | + | 756 | -18440 | Sema3c | bXPF_tRA |
| 1-1318 | chr5 | 21168887 | 21169350 | + | 655 | -17149 | Gsap | bXPF_tRA |
| 1-1497 | chr5 | 21701366 | 21701829 | + | 643 | -252 | Napepld | bXPF_tRA |
| 1-1462 | chr5 | 21786449 | 21786912 | + | 704 | 1397 | Psmc2 | bXPF_tRA |
| 1-332 | chr5 | 23432880 | 23433343 | + | 676 | 1242 | 5031425E22Rik | bXPF_tRA |
| 1-397 | chr5 | 24444733 | 24445196 | + | 619 | 271 | Fastk | bXPF_tRA |
| 1-828 | chr5 | 24604442 | 24604905 | + | 688 | -2671 | Smarcd3 | bXPF_tRA |
| 1-1349 | chr5 | 24685357 | 24685820 | + | 782 | -227 | Nub1 | bXPF_tRA |
| 1-478 | chr5 | 24685573 | 24686036 | + | 782 | -11 | Nub1 | bXPF_tRA |
| 1-419 | chr5 | 28071608 | 28072071 | + | 647 | 427 | Insig1 | bXPF_tRA |
| 1-724 | chr5 | 30888193 | 30888656 | + | 613 | -428 | Agbl5 | bXPF_tRA |
| 1-164 | chr5 | 30888415 | 30888878 | + | 613 | -206 | Agbl5 | bXPF_tRA |
| 1-1685 | chr5 | 30912959 | 30913422 | + | 801 | -596 | Emilin1 | bXPF_tRA |
| 1-1892 | chr5 | 31526673 | 31527136 | + | 997 | -91 | Slc4a1ap | bXPF_tRA |
| 1-162 | chr5 | 31526880 | 31527343 | + | 719 | 116 | Slc4a1ap | bXPF_tRA |
| 1-1413 | chr5 | 31602322 | 31602785 | + | 714 | -11398 | Mrpl33 | bXPF_tRA |
| 1-605 | chr5 | 31876475 | 31876938 | + | 599 | 178656 | Bre | bXPF_tRA |
| 1-326 | chr5 | 31876674 | 31877137 | + | 599 | 178855 | Bre | bXPF_tRA |
| 1-446 | chr5 | 33782297 | 33782760 | + | 551 | 176 | Letm1 | bXPF_tRA |
| 1-1710 | chr5 | 34761424 | 34761887 | + | 771 | -85 | Htt | bXPF_tRA |
| 1-1233 | chr5 | 35921568 | 35922031 | + | 571 | 28480 | Afap1 | bXPF_tRA |
| 1-748 | chr5 | 37824253 | 37824716 | + | 563 | 101 | Msx1 | bXPF_tRA |
| 17533 | chr5 | 38039001 | 38039464 | + | 1000 | 2 | Stx18 | bXPF_tRA |
| 1-113 | chr5 | 38039201 | 38039664 | + | 618 | 202 | Stx18 | bXPF_tRA |
| 1-539 | chr5 | 39644424 | 39644887 | + | 672 | -24 | Hs3st1 | bXPF_tRA |
| 1-849 | chr5 | 43948859 | 43949322 | + | 575 | 32709 | Fgfbp1 | bXPF_tRA |
| 1-1518 | chr5 | 46790492 | 46790955 | + | 804 | -858314 | 4930405L22Rik | bXPF_tRA |
| 1-1308 | chr5 | 48402396 | 48402859 | + | 641 | 7054 | 5730480H06Rik | bXPF_tRA |
| 1-1056 | chr5 | 49755536 | 49755999 | + | 610 | 303229 | Adgra3 | bXPF_tRA |
| 1-1208 | chr5 | 52782801 | 52783264 | + | 677 | -23 | Zcchc4 | bXPF_tRA |
| 1-1248 | chr5 | 52833837 | 52834300 | + | 710 | -67 | Anapc4 | bXPF_tRA |
| 1-1818 | chr5 | 53809326 | 53809789 | + | 774 | -70 | Tbc1d19 | bXPF_tRA |
| 1-928 | chr5 | 57755640 | 57756103 | + | 610 | 37790 | Pcdh7 | bXPF_tRA |
| 1-1000 | chr5 | 64802779 | 64803242 | + | 689 | -513 | Klf3 | bXPF_tRA |
| 1-134 | chr5 | 64803101 | 64803564 | + | 689 | -191 | Klf3 | bXPF_tRA |
| 1-1190 | chr5 | 66199944 | 66200407 | + | 587 | -8840 | 1700126H18Rik | bXPF_tRA |
| 1-1062 | chr5 | 74792236 | 74792699 | + | 624 | -89564 | Lnx1 | bXPF_tRA |
| 1-1618 | chr5 | 74944038 | 74944501 | + | 817 | -4925 | Gm6116 | bXPF_tRA |
| 1-381 | chr5 | 74944238 | 74944701 | + | 634 | -4725 | Gm6116 | bXPF_tRA |
| 1-1716 | chr5 | 76140345 | 76140808 | + | 800 | 303 | Srd5a3 | bXPF_tRA |
| 1-425 | chr5 | 81364189 | 81364652 | + | 593 | 342827 | Adgrl3 | bXPF_tRA |
| 1-773 | chr5 | 92277804 | 92278267 | + | 569 | 146 | Naaa | bXPF_tRA |
| 1-957 | chr5 | 92505314 | 92505777 | + | 650 | 112 | Scarb2 | bXPF_tRA |
| 1-1668 | chr5 | 93206451 | 93206914 | + | 785 | 164 | 2010109A12Rik | bXPF_tRA |
| 3289 | chr5 | 96209846 | 96210309 | + | 933 | -38 | Mrpl1 | bXPF_tRA |
| 1-422 | chr5 | 96996985 | 96997448 | + | 544 | -473 | Bmp2k | bXPF_tRA |
| 1-999 | chr5 | 98329235 | 98329698 | + | 554 | 162 | 1700007G11Rik | bXPF_tRA |
| 1-1433 | chr5 | 100798443 | 100798906 | + | 675 | -74 | Helq | bXPF_tRA |

| | | | | | | | | |
|---|---|---|---|---|---|---|---|---|
| 1-958 | chr5 | 105779384 | 105779847 | + | 567 | 78805 | Lrrc8d | bXPF_tRA |
| 1-1421 | chr5 | 107087226 | 107087689 | + | 665 | 122896 | Cdc7 | bXPF_tRA |
| 42005 | chr5 | 107087427 | 107087890 | + | 665 | 123097 | Cdc7 | bXPF_tRA |
| 1-1253 | chr5 | 107088126 | 107088589 | + | 594 | 123796 | Cdc7 | bXPF_tRA |
| 1-1683 | chr5 | 107289344 | 107289807 | + | 894 | 20 | Tgfbr3 | bXPF_tRA |
| 15707 | chr5 | 107289578 | 107290041 | + | 551 | -214 | Tgfbr3 | bXPF_tRA |
| 1-1841 | chr5 | 107437702 | 107438165 | + | 709 | -64 | Btbd8 | bXPF_tRA |
| 1-613 | chr5 | 107900530 | 107900993 | + | 609 | 233 | Rpl5 | bXPF_tRA |
| 18994 | chr5 | 108065407 | 108065870 | + | 1000 | -36 | Mtf2 | bXPF_tRA |
| 1-853 | chr5 | 108629388 | 108629851 | + | 624 | 158 | Gak | bXPF_tRA |
| 1-676 | chr5 | 108693810 | 108694273 | + | 555 | -188 | Fgfrl1 | bXPF_tRA |
| 1-723 | chr5 | 110107545 | 110108008 | + | 614 | 421 | Gtpbp6 | bXPF_tRA |
| 1-401 | chr5 | 110107849 | 110108312 | + | 614 | 117 | Gtpbp6 | bXPF_tRA |
| 1-977 | chr5 | 112326165 | 112326628 | + | 550 | 27 | Tfip11 | bXPF_tRA |
| 1-472 | chr5 | 114370437 | 114370900 | + | 579 | 9837 | Kctd10 | bXPF_tRA |
| 1-1538 | chr5 | 115158114 | 115158577 | + | 741 | -169 | Mlec | bXPF_tRA |
| 1-443 | chr5 | 115489498 | 115489961 | + | 548 | -5245 | Sirt4 | bXPF_tRA |
| 1-257 | chr5 | 115489709 | 115490172 | + | 548 | -5456 | Sirt4 | bXPF_tRA |
| 1-764 | chr5 | 115559316 | 115559779 | + | 727 | 80 | Rplp0 | bXPF_tRA |
| 1-1033 | chr5 | 118458912 | 118459375 | + | 631 | -101576 | Med13l | bXPF_tRA |
| 1-237 | chr5 | 118459115 | 118459578 | + | 632 | -101373 | Med13l | bXPF_tRA |
| 1-921 | chr5 | 118465286 | 118465749 | + | 793 | -95202 | Med13l | bXPF_tRA |
| 1-1703 | chr5 | 122621406 | 122621869 | + | 833 | -7119 | Ift81 | bXPF_tRA |
| 8767 | chr5 | 122821207 | 122821670 | + | 1000 | -96 | Anapc5 | bXPF_tRA |
| 1-226 | chr5 | 123749327 | 123749790 | + | 547 | -144 | Rsrc2 | bXPF_tRA |
| 1-263 | chr5 | 124439615 | 124440078 | + | 631 | -84 | Kmt5a | bXPF_tRA |
| 1-940 | chr5 | 124578888 | 124579351 | + | 562 | 12 | Eif2b1 | bXPF_tRA |
| 1-533 | chr5 | 125389465 | 125389928 | + | 756 | 321 | Ubc | bXPF_tRA |
| 1-280 | chr5 | 125389722 | 125390185 | + | 757 | 64 | Ubc | bXPF_tRA |
| 1-1457 | chr5 | 125404078 | 125404541 | + | 613 | -14292 | Ubc | bXPF_tRA |
| 1-222 | chr5 | 125404277 | 125404740 | + | 543 | -14491 | Ubc | bXPF_tRA |
| 1-293 | chr5 | 125405454 | 125405917 | + | 790 | -15668 | Ubc | bXPF_tRA |
| 1-574 | chr5 | 125405816 | 125406279 | + | 567 | -16030 | Ubc | bXPF_tRA |
| 31413 | chr5 | 125406019 | 125406482 | + | 567 | -16233 | Ubc | bXPF_tRA |
| 1-1011 | chr5 | 127617277 | 127617740 | + | 626 | -116 | Slc15a4 | bXPF_tRA |
| 1-1784 | chr5 | 129669895 | 129670358 | + | 937 | -38 | Zfp11 | bXPF_tRA |
| 1-761 | chr5 | 129724753 | 129725216 | + | 598 | -91 | Gbas | bXPF_tRA |
| 1-1250 | chr5 | 131711227 | 131711690 | + | 564 | -168811 | 4930563F08Rik | bXPF_tRA |
| 1-1141 | chr5 | 133173257 | 133173720 | + | 736 | -631145 | Auts2 | bXPF_tRA |
| 1-810 | chr5 | 134313457 | 134313920 | + | 695 | 1072 | Gtf2i | bXPF_tRA |
| 1-235 | chr5 | 134313657 | 134314120 | + | 695 | 872 | Gtf2i | bXPF_tRA |
| 1-1852 | chr5 | 134688649 | 134689112 | + | 628 | -290 | Limk1 | bXPF_tRA |
| 1-147 | chr5 | 134688848 | 134689311 | + | 628 | -489 | Limk1 | bXPF_tRA |
| 1-367 | chr5 | 135168295 | 135168758 | + | 571 | 154 | Bcl7b | bXPF_tRA |
| 1-242 | chr5 | 135168498 | 135168961 | + | 571 | 357 | Bcl7b | bXPF_tRA |
| 1-745 | chr5 | 136116410 | 136116873 | + | 630 | -50 | Polr2j | bXPF_tRA |
| 32509 | chr5 | 136116609 | 136117072 | + | 630 | 149 | Polr2j | bXPF_tRA |
| 1-1689 | chr5 | 136403461 | 136403924 | + | 701 | -27889 | Mir721 | bXPF_tRA |
| 1-374 | chr5 | 136403679 | 136404142 | + | 603 | -28107 | Mir721 | bXPF_tRA |
| 1-1359 | chr5 | 137533016 | 137533479 | + | 728 | -18 | Gnb2 | bXPF_tRA |
| 1-272 | chr5 | 137533215 | 137533678 | + | 728 | -217 | Gnb2 | bXPF_tRA |
| 1-1490 | chr5 | 137611125 | 137611588 | + | 571 | 48 | Pcolce | bXPF_tRA |
| 1-1536 | chr5 | 138116712 | 138117175 | + | 743 | 38 | Zscan21 | bXPF_tRA |
| 1-1807 | chr5 | 138170877 | 138171340 | + | 783 | 754 | Mcm7 | bXPF_tRA |
| 1-1704 | chr5 | 138280150 | 138280613 | + | 777 | -128 | Stag3 | bXPF_tRA |
| 1-807 | chr5 | 138754388 | 138754851 | + | 587 | -462 | Fam20c | bXPF_tRA |
| 1-1015 | chr5 | 139344975 | 139345438 | + | 620 | -40 | Cox19 | bXPF_tRA |
| 1-554 | chr5 | 139775364 | 139775827 | + | 542 | 83 | Ints1 | bXPF_tRA |
| 1-870 | chr5 | 141241175 | 141241638 | + | 625 | -128 | Sdk1 | bXPF_tRA |
| 1-1165 | chr5 | 143642270 | 143642733 | + | 682 | -8740 | Cyth3 | bXPF_tRA |
| 1-124 | chr5 | 143642470 | 143642933 | + | 682 | -8540 | Cyth3 | bXPF_tRA |
| 1-1617 | chr5 | 144014635 | 144015098 | + | 619 | -13 | Ccz1 | bXPF_tRA |
| 1-1602 | chr5 | 145140007 | 145140470 | + | 644 | -138 | Bud31 | bXPF_tRA |

| | | | | | | | | |
|---|---|---|---|---|---|---|---|---|
| 1-1144 | chr5 | 146221255 | 146221718 | + | 697 | -29 | Rnf6 | bXPF_tRA |
| 1-938 | chr5 | 147076395 | 147076858 | + | 572 | -54 | Lnx2 | bXPF_tRA |
| 1-1512 | chr5 | 148552518 | 148552981 | + | 900 | 39 | Ubl3 | bXPF_tRA |
| 1-199 | chr5 | 148552718 | 148553181 | + | 899 | -161 | Ubl3 | bXPF_tRA |
| 9133 | chr5 | 149184385 | 149184848 | + | 916 | 56 | Uspl1 | bXPF_tRA |
| 6211 | chr5 | 149228958 | 149229421 | + | 780 | -35815 | Alox5ap | bXPF_tRA |
| 1-1073 | chr5 | 150665493 | 150665956 | + | 605 | -112 | N4bp2l2 | bXPF_tRA |
| 1-1451 | chr5 | 151368707 | 151369170 | + | 712 | 263 | 1700028E10Rik | bXPF_tRA |
| 1-741 | chr6 | 8172054 | 8172517 | + | 551 | 20332 | Col28a1 | bXPF_tRA |
| 1-231 | chr6 | 8172254 | 8172717 | + | 551 | 20132 | Col28a1 | bXPF_tRA |
| 1-1110 | chr6 | 17771564 | 17772027 | + | 598 | 22625 | St7 | bXPF_tRA |
| 1-1563 | chr6 | 22356059 | 22356522 | + | 717 | -209 | Fam3c | bXPF_tRA |
| 1-1169 | chr6 | 24527339 | 24527802 | + | 744 | 115 | Ndufa5 | bXPF_tRA |
| 10959 | chr6 | 29388568 | 29389031 | + | 1000 | -7776 | Ccdc136 | bXPF_tRA |
| 1-1582 | chr6 | 31082036 | 31082499 | + | 653 | -19174 | Mir29b-1 | bXPF_tRA |
| 1-576 | chr6 | 31088801 | 31089264 | + | 571 | -25939 | Mir29b-1 | bXPF_tRA |
| 1-1351 | chr6 | 34317262 | 34317725 | + | 684 | -4 | Akr1b3 | bXPF_tRA |
| 1-937 | chr6 | 34721731 | 34722194 | + | 639 | 12518 | Cald1 | bXPF_tRA |
| 1-1390 | chr6 | 37617315 | 37617778 | + | 687 | 87373 | Akr1d1 | bXPF_tRA |
| 1-1016 | chr6 | 37617572 | 37618035 | + | 687 | 87630 | Akr1d1 | bXPF_tRA |
| 1-1796 | chr6 | 38381187 | 38381650 | + | 633 | -106 | Ttc26 | bXPF_tRA |
| 1-1566 | chr6 | 38533557 | 38534020 | + | 692 | -1073 | Fmc1 | bXPF_tRA |
| 1-406 | chr6 | 38551887 | 38552350 | + | 568 | 784 | Luc7l2 | bXPF_tRA |
| 1-149 | chr6 | 39118300 | 39118763 | + | 542 | 58 | 4930599N23Rik | bXPF_tRA |
| 1-1031 | chr6 | 40325288 | 40325751 | + | 683 | 41 | Agk | bXPF_tRA |
| 1-1388 | chr6 | 40471165 | 40471628 | + | 794 | -19 | Ssbp1 | bXPF_tRA |
| 1-1644 | chr6 | 42349480 | 42349943 | + | 969 | -117 | Zyx | bXPF_tRA |
| 1-1897 | chr6 | 47650691 | 47651154 | + | 689 | 4172 | Rn4.5s | bXPF_tRA |
| 1-1387 | chr6 | 47781355 | 47781818 | + | 566 | 22066 | Mir704 | bXPF_tRA |
| 1-886 | chr6 | 47877351 | 47877814 | + | 658 | 27 | Zfp282 | bXPF_tRA |
| 1-541 | chr6 | 47920489 | 47920952 | + | 545 | 152 | Zfp212 | bXPF_tRA |
| 1-1557 | chr6 | 49319090 | 49319553 | + | 563 | 47 | Ccdc126 | bXPF_tRA |
| 1-1405 | chr6 | 50379164 | 50379627 | + | 652 | 3442 | Osbpl3 | bXPF_tRA |
| 24838 | chr6 | 50379364 | 50379827 | + | 610 | 3242 | Osbpl3 | bXPF_tRA |
| 1-417 | chr6 | 52221628 | 52222091 | + | 597 | -3286 | Hoxa7 | bXPF_tRA |
| 1-1575 | chr6 | 53573019 | 53573482 | + | 667 | -124 | Creb5 | bXPF_tRA |
| 1-1594 | chr6 | 54992622 | 54993085 | + | 737 | 14 | Ggct | bXPF_tRA |
| 13881 | chr6 | 54992822 | 54993285 | + | 738 | -186 | Ggct | bXPF_tRA |
| 1-1012 | chr6 | 59208744 | 59209207 | + | 595 | 105 | Tigd2 | bXPF_tRA |
| 1-1341 | chr6 | 65778899 | 65779362 | + | 821 | 168 | Prdm5 | bXPF_tRA |
| 1-611 | chr6 | 66534997 | 66535460 | + | 569 | -240 | Mad2l1 | bXPF_tRA |
| 1-1655 | chr6 | 67037124 | 67037587 | + | 569 | 52 | Gadd45a | bXPF_tRA |
| 1-1079 | chr6 | 67102428 | 67102891 | + | 598 | -65252 | Gadd45a | bXPF_tRA |
| 1-1192 | chr6 | 71632984 | 71633447 | + | 687 | -298 | Kdm3a | bXPF_tRA |
| 1-1251 | chr6 | 72344826 | 72345289 | + | 593 | 118 | Usp39 | bXPF_tRA |
| 1-523 | chr6 | 72439564 | 72440027 | + | 595 | -237 | Mat2a | bXPF_tRA |
| 1-1195 | chr6 | 72980925 | 72981388 | + | 555 | -22408 | Tmsb10 | bXPF_tRA |
| 1-1363 | chr6 | 82996788 | 82997251 | + | 622 | 36452 | Dok1 | bXPF_tRA |
| 1-1739 | chr6 | 83054386 | 83054849 | + | 755 | -46 | Htra2 | bXPF_tRA |
| 1-1054 | chr6 | 83325835 | 83326298 | + | 566 | 27 | Mob1a | bXPF_tRA |
| 1-1604 | chr6 | 83506664 | 83507127 | + | 670 | 74 | Dguok | bXPF_tRA |
| 1-841 | chr6 | 84181182 | 84181645 | + | 619 | 162027 | Dysf | bXPF_tRA |
| 1-975 | chr6 | 86403698 | 86404161 | + | 704 | -290 | Tia1 | bXPF_tRA |
| 1-578 | chr6 | 86472047 | 86472510 | + | 650 | 8552 | A430078I02Rik | bXPF_tRA |
| 1-168 | chr6 | 86472253 | 86472716 | + | 650 | 8346 | A430078I02Rik | bXPF_tRA |
| 1-891 | chr6 | 86584907 | 86585370 | + | 554 | -43036 | Asprv1 | bXPF_tRA |
| 1-573 | chr6 | 87042385 | 87042848 | + | 658 | -230 | Gfpt1 | bXPF_tRA |
| 30682 | chr6 | 87042608 | 87043071 | + | 657 | -7 | Gfpt1 | bXPF_tRA |
| 1-1822 | chr6 | 87838602 | 87839065 | + | 806 | -74 | Isy1 | bXPF_tRA |
| 1-1088 | chr6 | 88446356 | 88446819 | + | 573 | -48 | Eefsec | bXPF_tRA |
| 1-1396 | chr6 | 88465208 | 88465671 | + | 645 | 30 | Ruvbl1 | bXPF_tRA |
| 1-442 | chr6 | 88842099 | 88842562 | + | 591 | -395 | Abtb1 | bXPF_tRA |
| 1-429 | chr6 | 88935835 | 88936298 | + | 576 | 33815 | Tpra1 | bXPF_tRA |

| | | | | | | | | |
|---|---|---|---|---|---|---|---|---|
| 1-660 | chr6 | 88936088 | 88936551 | + | 576 | 34068 | Tpra1 | bXPF_tRA |
| 1-1517 | chr6 | 92827345 | 92827808 | + | 735 | 73865 | Adamts9 | bXPF_tRA |
| 1-1081 | chr6 | 99523186 | 99523649 | + | 632 | -2424 | Foxp1 | bXPF_tRA |
| 1-144 | chr6 | 99523385 | 99523848 | + | 632 | -2623 | Foxp1 | bXPF_tRA |
| 1-753 | chr6 | 101251123 | 101251586 | + | 715 | 126543 | Pdzrn3 | bXPF_tRA |
| 14246 | chr6 | 101251325 | 101251788 | + | 715 | 126341 | Pdzrn3 | bXPF_tRA |
| 1-1437 | chr6 | 108668170 | 108668633 | + | 796 | -7467 | 0610040F04Rik | bXPF_tRA |
| 1-804 | chr6 | 112888883 | 112889346 | + | 553 | 58152 | Srgap3 | bXPF_tRA |
| 1-972 | chr6 | 112946769 | 112947232 | + | 741 | 266 | Srgap3 | bXPF_tRA |
| 1-736 | chr6 | 113076069 | 113076532 | + | 579 | 944 | Gt(ROSA)26Sor | bXPF_tRA |
| 1-318 | chr6 | 113077071 | 113077534 | + | 817 | -58 | Gt(ROSA)26Sor | bXPF_tRA |
| 1-180 | chr6 | 113077272 | 113077735 | + | 818 | -136 | Setd5 | bXPF_tRA |
| 1-1439 | chr6 | 113099600 | 113100063 | + | 564 | -5104 | Setd5 | bXPF_tRA |
| 1-281 | chr6 | 113099816 | 113100279 | + | 555 | -4888 | Setd5 | bXPF_tRA |
| 1-1394 | chr6 | 113237521 | 113237984 | + | 723 | -91 | Mtmr14 | bXPF_tRA |
| 1-246 | chr6 | 113306355 | 113306818 | + | 580 | -551 | Brpf1 | bXPF_tRA |
| 1-700 | chr6 | 113377523 | 113377986 | + | 565 | -234 | Tada3 | bXPF_tRA |
| 1-1713 | chr6 | 114888900 | 114889363 | + | 809 | 32621 | Vgll4 | bXPF_tRA |
| 1-1881 | chr6 | 115676347 | 115676810 | + | 775 | 57 | Raf1 | bXPF_tRA |
| 1-970 | chr6 | 117168479 | 117168942 | + | 753 | 175 | Cxcl12 | bXPF_tRA |
| 1-1475 | chr6 | 117879342 | 117879805 | + | 669 | 16496 | Zfp239 | bXPF_tRA |
| 1-860 | chr6 | 118419245 | 118419708 | + | 552 | -59 | Bms1 | bXPF_tRA |
| 1-165 | chr6 | 118419444 | 118419907 | + | 552 | -258 | Bms1 | bXPF_tRA |
| 1-1859 | chr6 | 119848178 | 119848641 | + | 898 | 216 | 3110021A11Rik | bXPF_tRA |
| 1-545 | chr6 | 119902573 | 119903036 | + | 546 | 106 | Rad52 | bXPF_tRA |
| 1-947 | chr6 | 122308533 | 122308996 | + | 580 | -246 | M6pr | bXPF_tRA |
| 1-638 | chr6 | 122669360 | 122669823 | + | 559 | -37898 | Nanog | bXPF_tRA |
| 1-1334 | chr6 | 122869831 | 122870294 | + | 615 | -4495 | Necap1 | bXPF_tRA |
| 1-1063 | chr6 | 124711944 | 124712407 | + | 560 | 3 | Emg1 | bXPF_tRA |
| 1-963 | chr6 | 124740798 | 124741261 | + | 647 | 50 | Grcc10 | bXPF_tRA |
| 1-674 | chr6 | 124756674 | 124757137 | + | 591 | -418 | Atn1 | bXPF_tRA |
| 1-376 | chr6 | 124931173 | 124931636 | + | 637 | 16 | Mlf2 | bXPF_tRA |
| 1-1265 | chr6 | 125008805 | 125009268 | + | 596 | -202 | Zfp384 | bXPF_tRA |
| 24473 | chr6 | 125009011 | 125009474 | + | 596 | 4 | Zfp384 | bXPF_tRA |
| 1-1738 | chr6 | 125215394 | 125215857 | + | 779 | 44 | Vamp1 | bXPF_tRA |
| 1-1441 | chr6 | 127266067 | 127266530 | + | 562 | 41669 | Gm38404 | bXPF_tRA |
| 1-1691 | chr6 | 127453438 | 127453901 | + | 772 | 42 | Parp11 | bXPF_tRA |
| 1-358 | chr6 | 128300273 | 128300736 | + | 562 | 309 | Tead4 | bXPF_tRA |
| 1-267 | chr6 | 128438386 | 128438849 | + | 590 | 14 | Fkbp4 | bXPF_tRA |
| 1-844 | chr6 | 128763862 | 128764325 | + | 601 | 24548 | Klrb1c | bXPF_tRA |
| 1-1217 | chr6 | 128803056 | 128803519 | + | 636 | -14646 | Klrb1c | bXPF_tRA |
| 1-1252 | chr6 | 134035188 | 134035651 | + | 676 | -281 | Etv6 | bXPF_tRA |
| 1-307 | chr6 | 134035389 | 134035852 | + | 622 | -80 | Etv6 | bXPF_tRA |
| 1-1281 | chr6 | 136518752 | 136519215 | + | 652 | 132 | Atf7ip | bXPF_tRA |
| 1-292 | chr6 | 136519014 | 136519477 | + | 616 | 394 | Atf7ip | bXPF_tRA |
| 1-1489 | chr6 | 137465178 | 137465641 | + | 566 | 54486 | Eps8 | bXPF_tRA |
| 1-746 | chr6 | 137560299 | 137560762 | + | 541 | 10479 | Eps8 | bXPF_tRA |
| 1-1508 | chr6 | 143166851 | 143167314 | + | 886 | -148 | Etnk1 | bXPF_tRA |
| 1-1268 | chr6 | 145793096 | 145793559 | + | 562 | -15056 | Rassf8 | bXPF_tRA |
| 1-851 | chr6 | 145933877 | 145934340 | + | 540 | -39 | Sspn | bXPF_tRA |
| 1-1229 | chr6 | 147476058 | 147476521 | + | 622 | 418 | Ccdc91 | bXPF_tRA |
| 1-1509 | chr6 | 149188433 | 149188896 | + | 713 | 48 | Amn1 | bXPF_tRA |
| 1828 | chr7 | 6410025 | 6410488 | + | 878 | -4919 | Smim17 | bXPF_tRA |
| 1-1343 | chr7 | 12977844 | 12978307 | + | 740 | 227 | Zfp446 | bXPF_tRA |
| 1-1672 | chr7 | 13009550 | 13010013 | + | 786 | 19 | Zbtb45 | bXPF_tRA |
| 1-910 | chr7 | 16296675 | 16297138 | + | 541 | -10111 | Ccdc9 | bXPF_tRA |
| 1-965 | chr7 | 16400878 | 16401341 | + | 713 | -87 | Zc3h4 | bXPF_tRA |
| 1-1330 | chr7 | 16891031 | 16891494 | + | 667 | -524 | Gng8 | bXPF_tRA |
| 21186 | chr7 | 16923991 | 16924454 | + | 1000 | -190 | Calm3 | bXPF_tRA |
| 1-1876 | chr7 | 19344902 | 19345365 | + | 811 | 62 | Ercc1 | bXPF_tRA |
| 1-1239 | chr7 | 25686968 | 25687431 | + | 673 | 197 | Tgfb1 | bXPF_tRA |
| 1-1047 | chr7 | 25711695 | 25712158 | + | 554 | 7127 | Ccdc97 | bXPF_tRA |
| 1-956 | chr7 | 25754610 | 25755073 | + | 627 | -121 | Hnrnpul1 | bXPF_tRA |

| | | | | | | | | |
|---|---|---|---|---|---|---|---|---|
| 1-1329 | chr7 | 27474555 | 27475018 | + | 607 | 946 | Sertad3 | bXPF_tRA |
| 1-1783 | chr7 | 28372605 | 28373068 | + | 887 | -174 | Plekhg2 | bXPF_tRA |
| 1-1128 | chr7 | 28372805 | 28373268 | + | 889 | -374 | Plekhg2 | bXPF_tRA |
| 1-335 | chr7 | 28380876 | 28381339 | + | 709 | -1879 | Zfp36 | bXPF_tRA |
| 1-903 | chr7 | 34233750 | 34234213 | + | 627 | -3645 | Gpi1 | bXPF_tRA |
| 1-159 | chr7 | 34233949 | 34234412 | + | 627 | -3844 | Gpi1 | bXPF_tRA |
| 1-1406 | chr7 | 34318991 | 34319454 | + | 627 | -33613 | 4931406P16Rik | bXPF_tRA |
| 1-1385 | chr7 | 34342104 | 34342567 | + | 660 | 47205 | Lsm14a | bXPF_tRA |
| 1-1732 | chr7 | 34655411 | 34655874 | + | 767 | -2801 | Kctd15 | bXPF_tRA |
| 1-979 | chr7 | 38084777 | 38085240 | + | 543 | 22482 | Ccne1 | bXPF_tRA |
| 1-1070 | chr7 | 43671629 | 43672092 | + | 546 | -171 | Ctu1 | bXPF_tRA |
| 1-424 | chr7 | 44246533 | 44246996 | + | 550 | 42 | 2410002F23Rik | bXPF_tRA |
| 1-470 | chr7 | 44816361 | 44816824 | + | 546 | 66 | Atf5 | bXPF_tRA |
| 1-960 | chr7 | 45062429 | 45062892 | + | 592 | 231 | Nosip | bXPF_tRA |
| 1-718 | chr7 | 45434551 | 45435014 | + | 590 | -57 | Gys1 | bXPF_tRA |
| 1-1207 | chr7 | 45717941 | 45718404 | + | 652 | 101 | Rpl18 | bXPF_tRA |
| 1-1885 | chr7 | 47008166 | 47008629 | + | 963 | 17 | Spty2d1 | bXPF_tRA |
| 9863 | chr7 | 70267074 | 70267537 | + | 1000 | 70517 | Gm29683 | bXPF_tRA |
| 1-1848 | chr7 | 70346031 | 70346494 | + | 641 | -8410 | Gm29683 | bXPF_tRA |
| 1-213 | chr7 | 70359707 | 70360170 | + | 551 | 655 | Nr2f2 | bXPF_tRA |
| 1-109 | chr7 | 70359914 | 70360377 | + | 551 | 448 | Nr2f2 | bXPF_tRA |
| 1-241 | chr7 | 70361348 | 70361811 | + | 648 | -986 | Nr2f2 | bXPF_tRA |
| 1-703 | chr7 | 70633552 | 70634015 | + | 680 | -267037 | Nr2f2 | bXPF_tRA |
| 1-1849 | chr7 | 73537651 | 73538114 | + | 834 | 3864 | Chd2 | bXPF_tRA |
| 1-455 | chr7 | 73617728 | 73618191 | + | 558 | -59564 | 1810026B05Rik | bXPF_tRA |
| 1-362 | chr7 | 73618024 | 73618487 | + | 558 | -59860 | 1810026B05Rik | bXPF_tRA |
| 1-815 | chr7 | 74107160 | 74107623 | + | 582 | -93709 | St8sia2 | bXPF_tRA |
| 1-1868 | chr7 | 79466218 | 79466681 | + | 967 | -176 | Polg | bXPF_tRA |
| 1-130 | chr7 | 79466440 | 79466903 | + | 621 | -398 | Polg | bXPF_tRA |
| 13881 | chr7 | 79920468 | 79920931 | + | 1000 | -59 | Ap3s2 | bXPF_tRA |
| 1-469 | chr7 | 80324062 | 80324525 | + | 551 | 161 | Rccd1 | bXPF_tRA |
| 26299 | chr7 | 80324268 | 80324731 | + | 552 | -45 | Rccd1 | bXPF_tRA |
| 1-1090 | chr7 | 81455212 | 81455675 | + | 558 | -685 | Cpeb1 | bXPF_tRA |
| 1-174 | chr7 | 81455417 | 81455880 | + | 558 | -890 | Cpeb1 | bXPF_tRA |
| 1-1530 | chr7 | 83884160 | 83884623 | + | 585 | -50 | Mesdc1 | bXPF_tRA |
| 1-493 | chr7 | 84151227 | 84151690 | + | 598 | 435 | Abhd17c | bXPF_tRA |
| 1-569 | chr7 | 84332579 | 84333042 | + | 551 | 77228 | Arnt2 | bXPF_tRA |
| 1-1769 | chr7 | 88200125 | 88200588 | + | 1000 | -77729 | Ctsc | bXPF_tRA |
| 1-680 | chr7 | 89563520 | 89563983 | + | 606 | -46165 | Prss23 | bXPF_tRA |
| 1-760 | chr7 | 89769287 | 89769750 | + | 554 | 134111 | Ccdc81 | bXPF_tRA |
| 1-1293 | chr7 | 98116864 | 98117327 | + | 672 | 2398 | Myo7a | bXPF_tRA |
| 1-1579 | chr7 | 98361203 | 98361666 | + | 614 | -106 | Tsku | bXPF_tRA |
| 1-635 | chr7 | 98489622 | 98490085 | + | 564 | -4369 | Lrrc32 | bXPF_tRA |
| 1-626 | chr7 | 98524032 | 98524495 | + | 593 | 30041 | Lrrc32 | bXPF_tRA |
| 1-301 | chr7 | 98524280 | 98524743 | + | 543 | 30289 | Lrrc32 | bXPF_tRA |
| 1-1539 | chr7 | 98815254 | 98815717 | + | 770 | -19627 | Wnt11 | bXPF_tRA |
| 1-260 | chr7 | 98815466 | 98815929 | + | 769 | -19415 | Wnt11 | bXPF_tRA |
| 1-1599 | chr7 | 99353116 | 99353579 | + | 741 | -108 | Serpinh1 | bXPF_tRA |
| 1-259 | chr7 | 99359887 | 99360350 | + | 725 | -6879 | Serpinh1 | bXPF_tRA |
| 1-669 | chr7 | 99360236 | 99360699 | + | 725 | -7228 | Serpinh1 | bXPF_tRA |
| 1-1718 | chr7 | 101108154 | 101108617 | + | 861 | -390 | Fchsd2 | bXPF_tRA |
| 1-1544 | chr7 | 105810796 | 105811259 | + | 762 | 60 | Mrpl17 | bXPF_tRA |
| 1-550 | chr7 | 105811001 | 105811464 | + | 762 | -145 | Mrpl17 | bXPF_tRA |
| 1-1835 | chr7 | 108934118 | 108934581 | + | 840 | -66 | Eif3f | bXPF_tRA |
| 1-1614 | chr7 | 108993141 | 108993604 | + | 805 | -17508 | Tub | bXPF_tRA |
| 1-105 | chr7 | 109671340 | 109671803 | + | 640 | -32166 | Akip1 | bXPF_tRA |
| 1-243 | chr7 | 109671581 | 109672044 | + | 640 | -31925 | Akip1 | bXPF_tRA |
| 1-926 | chr7 | 110164145 | 110164608 | + | 596 | 42317 | Wee1 | bXPF_tRA |
| 1-299 | chr7 | 110164421 | 110164884 | + | 546 | 42593 | Wee1 | bXPF_tRA |
| 1-1742 | chr7 | 110297149 | 110297612 | + | 742 | 75677 | Swap70 | bXPF_tRA |
| 1-1686 | chr7 | 110701993 | 110702456 | + | 653 | -65982 | Ampd3 | bXPF_tRA |
| 1-461 | chr7 | 110702227 | 110702690 | + | 602 | -65748 | Ampd3 | bXPF_tRA |
| 1-1806 | chr7 | 110706663 | 110707126 | + | 717 | -61312 | Ampd3 | bXPF_tRA |

| | | | | | | | | |
|---|---|---|---|---|---|---|---|---|
| 1-1812 | chr7 | 111083021 | 111083484 | + | 862 | -222 | Eif4g2 | bXPF_tRA |
| 1-1027 | chr7 | 111122151 | 111122614 | + | 561 | 293 | 1700012D14Rik | bXPF_tRA |
| 1-252 | chr7 | 111122353 | 111122816 | + | 559 | 91 | 1700012D14Rik | bXPF_tRA |
| 1-1176 | chr7 | 112023130 | 112023593 | + | 674 | -145 | Usp47 | bXPF_tRA |
| 1-394 | chr7 | 112378302 | 112378765 | + | 733 | 10225 | Micalcl | bXPF_tRA |
| 1-310 | chr7 | 112378508 | 112378971 | + | 733 | 10431 | Micalcl | bXPF_tRA |
| 1-1502 | chr7 | 112453785 | 112454248 | + | 788 | 26310 | Parva | bXPF_tRA |
| 1-803 | chr7 | 112697133 | 112697596 | + | 663 | 18044 | Tead1 | bXPF_tRA |
| 1-964 | chr7 | 113140065 | 113140528 | + | 598 | 1719 | 4930543E12Rik | bXPF_tRA |
| 1-1187 | chr7 | 114275955 | 114276418 | + | 557 | -70 | Psma1 | bXPF_tRA |
| 42979 | chr7 | 114276157 | 114276620 | + | 557 | -272 | Psma1 | bXPF_tRA |
| 1-1757 | chr7 | 114562578 | 114563041 | + | 816 | 163 | Cyp2r1 | bXPF_tRA |
| 20455 | chr7 | 116093276 | 116093739 | + | 557 | 127 | 1110004F10Rik | bXPF_tRA |
| 1-1336 | chr7 | 120917424 | 120917887 | + | 600 | -89 | Polr3e | bXPF_tRA |
| 1-1288 | chr7 | 120982360 | 120982823 | + | 586 | 82 | Mfsd13b | bXPF_tRA |
| 1-1022 | chr7 | 125491231 | 125491694 | + | 592 | 80 | Nsmce1 | bXPF_tRA |
| 1-215 | chr7 | 125491442 | 125491905 | + | 592 | -131 | Nsmce1 | bXPF_tRA |
| 1-1731 | chr7 | 126503478 | 126503941 | + | 803 | -407 | Atxn2l | bXPF_tRA |
| 1-1554 | chr7 | 126695680 | 126696143 | + | 717 | -89 | Bola2 | bXPF_tRA |
| 1-1294 | chr7 | 127876609 | 127877072 | + | 581 | -17 | Zfp668 | bXPF_tRA |
| 1-1863 | chr7 | 129591628 | 129592091 | + | 651 | -4 | Wdr11 | bXPF_tRA |
| 1-250 | chr7 | 129591829 | 129592292 | + | 651 | 197 | Wdr11 | bXPF_tRA |
| 1-1224 | chr7 | 130519510 | 130519973 | + | 615 | -161 | Ate1 | bXPF_tRA |
| 1-161 | chr7 | 130546533 | 130546996 | + | 586 | 617 | Nsmce4a | bXPF_tRA |
| 1-1099 | chr7 | 130965765 | 130966228 | + | 544 | 29793 | Htra1 | bXPF_tRA |
| 43101 | chr7 | 130965983 | 130966446 | + | 544 | 30011 | Htra1 | bXPF_tRA |
| 1-1242 | chr7 | 131362509 | 131362972 | + | 809 | -42 | 2310057M21Rik | bXPF_tRA |
| 1-1319 | chr7 | 131370974 | 131371437 | + | 540 | 59 | Pstk | bXPF_tRA |
| 1-1392 | chr7 | 132444268 | 132444731 | + | 738 | -127344 | Chst15 | bXPF_tRA |
| 1-1303 | chr7 | 134381414 | 134381877 | + | 540 | -4817 | D7Ertd443e | bXPF_tRA |
| 1-892 | chr7 | 137410381 | 137410844 | + | 549 | 144 | 9430038I01Rik | bXPF_tRA |
| 1-543 | chr7 | 137437503 | 137437966 | + | 620 | 60 | Gm12669 | bXPF_tRA |
| 21916 | chr7 | 137437703 | 137438166 | + | 620 | 260 | Gm12669 | bXPF_tRA |
| 1-230 | chr7 | 139248083 | 139248546 | + | 541 | -168 | Pwwp2b | bXPF_tRA |
| 1-575 | chr7 | 139971450 | 139971913 | + | 597 | 7074 | 6430531B16Rik | bXPF_tRA |
| 1-551 | chr7 | 140036143 | 140036606 | + | 550 | -17 | Zfp511 | bXPF_tRA |
| 1-1493 | chr7 | 140116203 | 140116666 | + | 695 | -11 | Echs1 | bXPF_tRA |
| 1-535 | chr7 | 140137425 | 140137888 | + | 544 | 92 | Mtg1 | bXPF_tRA |
| 1-1173 | chr7 | 141131083 | 141131546 | + | 541 | 28 | Ptdss2 | bXPF_tRA |
| 1-434 | chr7 | 143685481 | 143685944 | + | 631 | 163 | Tnfrsf23 | bXPF_tRA |
| 1-1537 | chr7 | 143740087 | 143740550 | + | 803 | 42 | Osbpl5 | bXPF_tRA |
| 1-778 | chr7 | 144941520 | 144941983 | + | 591 | -1826 | Ccnd1 | bXPF_tRA |
| 1-1420 | chr7 | 145222135 | 145222598 | + | 678 | -14247 | 1810010D01Rik | bXPF_tRA |
| 1-1161 | chr8 | 3515222 | 3515685 | + | 704 | 69 | Pnpla6 | bXPF_tRA |
| 1-995 | chr8 | 9977583 | 9978046 | + | 673 | 97 | Abhd13 | bXPF_tRA |
| 1-1657 | chr8 | 11497309 | 11497772 | + | 569 | -28 | Naxd | bXPF_tRA |
| 1-856 | chr8 | 11758026 | 11758489 | + | 541 | -73 | Arhgef7 | bXPF_tRA |
| 1-916 | chr8 | 13644228 | 13644691 | + | 578 | 33128 | Rasa3 | bXPF_tRA |
| 1-1243 | chr8 | 18619119 | 18619582 | + | 601 | 24177 | Mcph1 | bXPF_tRA |
| 23377 | chr8 | 19784538 | 19784850 | + | 1000 | 55118 | Gm21119 | bXPF_tRA |
| 21916 | chr8 | 19979865 | 19980328 | + | 1000 | 250520 | Gm21119 | bXPF_tRA |
| 6576 | chr8 | 20424671 | 20425134 | + | 694 | -88 | 2610005L07Rik | bXPF_tRA |
| 1-1799 | chr8 | 22411186 | 22411649 | + | 724 | 77 | Mrps31 | bXPF_tRA |
| 1-228 | chr8 | 25465330 | 25465793 | + | 592 | -53198 | Fgfr1 | bXPF_tRA |
| 1-906 | chr8 | 25754120 | 25754583 | + | 560 | -71 | Ddhd2 | bXPF_tRA |
| 1-1756 | chr8 | 26978706 | 26979169 | + | 976 | 1601 | Zfp703 | bXPF_tRA |
| 1-125 | chr8 | 26978905 | 26979368 | + | 978 | 1800 | Zfp703 | bXPF_tRA |
| 1-502 | chr8 | 33909328 | 33909791 | + | 641 | 20304 | Rbpms | bXPF_tRA |
| 1-403 | chr8 | 36613642 | 36614105 | + | 609 | 70 | Dlc1 | bXPF_tRA |
| 1-1409 | chr8 | 41374556 | 41375019 | + | 613 | -90 | Asah1 | bXPF_tRA |
| 1-1213 | chr8 | 45999925 | 46000388 | + | 711 | -306 | Ankrd37 | bXPF_tRA |
| 1-527 | chr8 | 46210854 | 46211317 | + | 551 | -76 | Slc25a4 | bXPF_tRA |
| 1-1745 | chr8 | 46429790 | 46430253 | + | 739 | -41016 | Acsl1 | bXPF_tRA |

| | | | | | | | | |
|---|---|---|---|---|---|---|---|---|
| 1-1583 | chr8 | 46551630 | 46552093 | + | 619 | -208 | Cenpu | bXPF_tRA |
| 1-962 | chr8 | 47082457 | 47082920 | + | 618 | 95801 | Enpp6 | bXPF_tRA |
| 1-1098 | chr8 | 47932235 | 47932698 | + | 590 | 58085 | Wwc2 | bXPF_tRA |
| 1-1605 | chr8 | 66486301 | 66486764 | + | 546 | -25 | Tma16 | bXPF_tRA |
| 1-1719 | chr8 | 68431206 | 68431669 | + | 758 | 62978 | 1700125H03Rik | bXPF_tRA |
| 1-104 | chr8 | 68431407 | 68431870 | + | 759 | 63179 | 1700125H03Rik | bXPF_tRA |
| 1-1107 | chr8 | 68707335 | 68707798 | + | 590 | 27580 | Csgalnact1 | bXPF_tRA |
| 1-484 | chr8 | 68707536 | 68707999 | + | 590 | 27379 | Csgalnact1 | bXPF_tRA |
| 1-1177 | chr8 | 68734888 | 68735351 | + | 612 | 27 | Csgalnact1 | bXPF_tRA |
| 1-642 | chr8 | 69716459 | 69716922 | + | 607 | -204 | Zfp869 | bXPF_tRA |
| 24108 | chr8 | 69716665 | 69717128 | + | 607 | 6 | Zfp869 | bXPF_tRA |
| 1-1584 | chr8 | 69749668 | 69750131 | + | 670 | 63 | Zfp963 | bXPF_tRA |
| 1-513 | chr8 | 70698387 | 70698850 | + | 592 | 879 | Jund | bXPF_tRA |
| 1-647 | chr8 | 71272379 | 71272842 | + | 655 | -20 | Haus8 | bXPF_tRA |
| 1-1491 | chr8 | 72160674 | 72161137 | + | 761 | -295 | Rab8a | bXPF_tRA |
| 1-408 | chr8 | 72160899 | 72161362 | + | 547 | -70 | Rab8a | bXPF_tRA |
| 1-1777 | chr8 | 72318858 | 72319321 | + | 703 | 27 | Klf2 | bXPF_tRA |
| 1-1386 | chr8 | 74993470 | 74993933 | + | 693 | 51 | Hmgxb4 | bXPF_tRA |
| 1-341 | chr8 | 74993683 | 74994146 | + | 694 | 264 | Hmgxb4 | bXPF_tRA |
| 1-595 | chr8 | 75083616 | 75084079 | + | 761 | -9771 | Hmox1 | bXPF_tRA |
| 1-661 | chr8 | 77723780 | 77724243 | + | 663 | 441 | Ednra | bXPF_tRA |
| 1-1888 | chr8 | 79027793 | 79028256 | + | 1000 | -413 | Zfp827 | bXPF_tRA |
| 1-217 | chr8 | 79028001 | 79028464 | + | 1000 | -205 | Zfp827 | bXPF_tRA |
| 1-1447 | chr8 | 79294470 | 79294933 | + | 681 | 255 | Mmaa | bXPF_tRA |
| 1-1666 | chr8 | 82943184 | 82943647 | + | 675 | 80059 | Rnf150 | bXPF_tRA |
| 1-467 | chr8 | 82943426 | 82943889 | + | 671 | 80301 | Rnf150 | bXPF_tRA |
| 1-1453 | chr8 | 83165354 | 83165817 | + | 577 | 233 | Tbc1d9 | bXPF_tRA |
| 1-584 | chr8 | 84723058 | 84723521 | + | 584 | 21832 | Lyl1 | bXPF_tRA |
| 1-264 | chr8 | 84743703 | 84744166 | + | 660 | 30435 | Nfix | bXPF_tRA |
| 21551 | chr8 | 84743914 | 84744377 | + | 659 | 30224 | Nfix | bXPF_tRA |
| 1-1621 | chr8 | 84799182 | 84799645 | + | 698 | 925 | Nfix | bXPF_tRA |
| 1-1126 | chr8 | 84976232 | 84976695 | + | 661 | 2285 | Junb | bXPF_tRA |
| 1-811 | chr8 | 85840733 | 85841196 | + | 542 | -15 | Itfg1 | bXPF_tRA |
| 1-1296 | chr8 | 90908087 | 90908550 | + | 668 | -23931 | Chd9 | bXPF_tRA |
| 1-343 | chr8 | 91069823 | 91070286 | + | 573 | -3 | Rbl2 | bXPF_tRA |
| 1-269 | chr8 | 91070067 | 91070530 | + | 573 | 241 | Rbl2 | bXPF_tRA |
| 1-1102 | chr8 | 92721480 | 92721943 | + | 576 | 47422 | Irx6 | bXPF_tRA |
| 1-448 | chr8 | 94838289 | 94838752 | + | 601 | 103 | Coq9 | bXPF_tRA |
| 1-1403 | chr8 | 94917383 | 94917846 | + | 689 | 484 | Ccdc102a | bXPF_tRA |
| 1-1338 | chr8 | 95434774 | 95435237 | + | 645 | -136 | Cfap20 | bXPF_tRA |
| 1-590 | chr8 | 95633290 | 95633753 | + | 567 | -38 | Gins3 | bXPF_tRA |
| 1-1335 | chr8 | 95888196 | 95888659 | + | 635 | -62 | Got2 | bXPF_tRA |
| 1-1074 | chr8 | 103414148 | 103414611 | + | 612 | -66845 | 1600027J07Rik | bXPF_tRA |
| 1-1625 | chr8 | 103913918 | 103914381 | + | 706 | -187476 | Cdh5 | bXPF_tRA |
| 1-1864 | chr8 | 104022986 | 104023449 | + | 540 | -78408 | Cdh5 | bXPF_tRA |
| 1-664 | chr8 | 104442566 | 104443029 | + | 604 | 250 | Dync1li2 | bXPF_tRA |
| 21551 | chr8 | 109778385 | 109778848 | + | 1000 | 33 | Ap1g1 | bXPF_tRA |
| 1-1346 | chr8 | 110941453 | 110941916 | + | 736 | 21819 | St3gal2 | bXPF_tRA |
| 1-1872 | chr8 | 111449833 | 111450296 | + | 820 | -136 | Mir1957b | bXPF_tRA |
| 1-707 | chr8 | 113635397 | 113635860 | + | 567 | 42 | Mon1b | bXPF_tRA |
| 1-1466 | chr8 | 113804145 | 113804608 | + | 593 | 44463 | Adamts18 | bXPF_tRA |
| 1-1175 | chr8 | 114439433 | 114439896 | + | 586 | 12 | Wwox | bXPF_tRA |
| 1-356 | chr8 | 119778191 | 119778654 | + | 625 | -6 | Tldc1 | bXPF_tRA |
| 1-249 | chr8 | 119778463 | 119778926 | + | 625 | -278 | Tldc1 | bXPF_tRA |
| 18629 | chr8 | 120588990 | 120589453 | + | 1000 | -146 | Gins2 | bXPF_tRA |
| 1-1166 | chr8 | 121829430 | 121829893 | + | 558 | -92 | Klhdc4 | bXPF_tRA |
| 1-389 | chr8 | 122281375 | 122281838 | + | 571 | -535 | Zfpm1 | bXPF_tRA |
| 1-1210 | chr8 | 122444404 | 122444867 | + | 637 | 979 | 9330133O14Rik | bXPF_tRA |
| 367 | chr8 | 122661102 | 122661565 | + | 1000 | 16842 | Cbfa2t3 | bXPF_tRA |
| 1-396 | chr8 | 122661304 | 122661767 | + | 653 | 16640 | Cbfa2t3 | bXPF_tRA |
| 1-1273 | chr8 | 124652510 | 124652973 | + | 594 | 10628 | 2310022B05Rik | bXPF_tRA |
| 1-421 | chr8 | 126593867 | 126594330 | + | 606 | -662 | Irf2bp2 | bXPF_tRA |
| 1-1023 | chr8 | 126600877 | 126601340 | + | 544 | -7672 | Irf2bp2 | bXPF_tRA |

| | | | | | | | | |
|---|---|---|---|---|---|---|---|---|
| 1-1670 | chr8 | 126603197 | 126603660 | + | 706 | -9992 | Irf2bp2 | bXPF_tRA |
| 18264 | chr8 | 127720737 | 127721200 | + | 1000 | 421316 | Pard3 | bXPF_tRA |
| 1-644 | chr8 | 128024965 | 128025428 | + | 601 | 253029 | Mir21c | bXPF_tRA |
| 1-459 | chr8 | 128025197 | 128025660 | + | 678 | 252797 | Mir21c | bXPF_tRA |
| 1-602 | chr9 | 7960292 | 7960755 | + | 570 | 44073 | Yap1 | bXPF_tRA |
| 25934 | chr9 | 7960495 | 7960958 | + | 570 | 43870 | Yap1 | bXPF_tRA |
| 1-1736 | chr9 | 13296523 | 13296986 | + | 763 | 49798 | Ccdc82 | bXPF_tRA |
| 1-1189 | chr9 | 14585729 | 14586192 | + | 720 | 29040 | Amotl1 | bXPF_tRA |
| 1-984 | chr9 | 15279623 | 15280086 | + | 589 | 13 | Med17 | bXPF_tRA |
| 1-294 | chr9 | 15279829 | 15280292 | + | 589 | -193 | Med17 | bXPF_tRA |
| 1-913 | chr9 | 15306179 | 15306642 | + | 621 | 38 | 4931406C07Rik | bXPF_tRA |
| 18994 | chr9 | 15306378 | 15306841 | + | 621 | -161 | 4931406C07Rik | bXPF_tRA |
| 1-1884 | chr9 | 18291915 | 18292378 | + | 760 | -121 | Chordc1 | bXPF_tRA |
| 1-145 | chr9 | 18292115 | 18292578 | + | 760 | 79 | Chordc1 | bXPF_tRA |
| 1-1629 | chr9 | 20384895 | 20385358 | + | 687 | 32 | Zfp560 | bXPF_tRA |
| 1-1249 | chr9 | 20607137 | 20607600 | + | 654 | -107 | Fbxl12os | bXPF_tRA |
| 1-456 | chr9 | 20607356 | 20607819 | + | 654 | 112 | Fbxl12os | bXPF_tRA |
| 1-619 | chr9 | 20877913 | 20878376 | + | 617 | 1566 | Angptl6 | bXPF_tRA |
| 1-1044 | chr9 | 21168741 | 21169204 | + | 600 | 3258 | Pde4a | bXPF_tRA |
| 22282 | chr9 | 21168997 | 21169460 | + | 600 | 3514 | Pde4a | bXPF_tRA |
| 1-1477 | chr9 | 21496675 | 21497138 | + | 735 | -342 | Mir199a-1 | bXPF_tRA |
| 28126 | chr9 | 21496877 | 21497340 | + | 735 | -544 | Mir199a-1 | bXPF_tRA |
| 1-737 | chr9 | 23728790 | 23729253 | + | 625 | -368997 | Npsr1 | bXPF_tRA |
| 1-438 | chr9 | 25252247 | 25252710 | + | 681 | 39 | Sept7 | bXPF_tRA |
| 1-507 | chr9 | 31280540 | 31281003 | + | 705 | 9370 | Gm7244 | bXPF_tRA |
| 20821 | chr9 | 31280854 | 31281317 | + | 705 | 9684 | Gm7244 | bXPF_tRA |
| 1-890 | chr9 | 32695686 | 32696149 | + | 544 | -125 | Ets1 | bXPF_tRA |
| 1-1186 | chr9 | 35116777 | 35117240 | + | 757 | -198 | St3gal4 | bXPF_tRA |
| 1-1511 | chr9 | 35339498 | 35339961 | + | 657 | 58624 | 4933422A05Rik | bXPF_tRA |
| 1-435 | chr9 | 37146705 | 37147168 | + | 606 | 386 | Pknox2 | bXPF_tRA |
| 1-908 | chr9 | 40730682 | 40731145 | + | 615 | 44949 | Clmp | bXPF_tRA |
| 1-1656 | chr9 | 41515233 | 41515696 | + | 791 | -15961 | Mir100 | bXPF_tRA |
| 1-1877 | chr9 | 41705453 | 41705916 | + | 691 | 123758 | Mir125b-1 | bXPF_tRA |
| 1-303 | chr9 | 41705657 | 41706120 | + | 571 | 123962 | Mir125b-1 | bXPF_tRA |
| 1-566 | chr9 | 41877319 | 41877782 | + | 574 | 246739 | Sorl1 | bXPF_tRA |
| 1-919 | chr9 | 41924183 | 41924646 | + | 640 | 199875 | Sorl1 | bXPF_tRA |
| 7306 | chr9 | 41950385 | 41950848 | + | 1000 | 173673 | Sorl1 | bXPF_tRA |
| 1-384 | chr9 | 41951204 | 41951667 | + | 633 | 172854 | Sorl1 | bXPF_tRA |
| 1-322 | chr9 | 44498495 | 44498958 | + | 572 | -410 | Bcl9l | bXPF_tRA |
| 1-176 | chr9 | 44498730 | 44499193 | + | 572 | -175 | Bcl9l | bXPF_tRA |
| 5115 | chr9 | 44881242 | 44881705 | + | 1000 | -199 | Kmt2a | bXPF_tRA |
| 1-169 | chr9 | 44881449 | 44881912 | + | 765 | -406 | Kmt2a | bXPF_tRA |
| 1-606 | chr9 | 49571709 | 49572172 | + | 589 | 53652 | Gm11149 | bXPF_tRA |
| 1-316 | chr9 | 49571913 | 49572376 | + | 589 | 53856 | Gm11149 | bXPF_tRA |
| 1-536 | chr9 | 49798930 | 49799393 | + | 543 | -92 | Ncam1 | bXPF_tRA |
| 1-1546 | chr9 | 49816094 | 49816557 | + | 581 | -16932 | Ncam1 | bXPF_tRA |
| 1-1725 | chr9 | 49980722 | 49981185 | + | 815 | -181560 | Ncam1 | bXPF_tRA |
| 1-1399 | chr9 | 50617011 | 50617474 | + | 700 | -79 | Pih1d2 | bXPF_tRA |
| 1-1155 | chr9 | 50775184 | 50775647 | + | 582 | 190 | Alg9 | bXPF_tRA |
| 1-927 | chr9 | 56041722 | 56042185 | + | 610 | 108 | Rcn2 | bXPF_tRA |
| 1-314 | chr9 | 56041992 | 56042455 | + | 610 | 378 | Rcn2 | bXPF_tRA |
| 1-934 | chr9 | 56417794 | 56418257 | + | 629 | 25 | Peak1 | bXPF_tRA |
| 1-368 | chr9 | 56417998 | 56418461 | + | 629 | -179 | Peak1 | bXPF_tRA |
| 1-1089 | chr9 | 56463639 | 56464102 | + | 616 | 45024 | Hmg20a | bXPF_tRA |
| 1-932 | chr9 | 57605992 | 57606455 | + | 552 | 58 | Cplx3 | bXPF_tRA |
| 1-1878 | chr9 | 57831422 | 57831885 | + | 566 | 2581 | Arid3b | bXPF_tRA |
| 1-931 | chr9 | 58214620 | 58215083 | + | 575 | -6299 | Mir6385 | bXPF_tRA |
| 1-1113 | chr9 | 59291157 | 59291620 | + | 615 | -184 | Adpgk | bXPF_tRA |
| 1-1444 | chr9 | 59486640 | 59487103 | + | 654 | -497 | Arih1 | bXPF_tRA |
| 1-586 | chr9 | 61914333 | 61914796 | + | 568 | -54 | Rplp1 | bXPF_tRA |
| 1-1558 | chr9 | 62559972 | 62560435 | + | 656 | -23159 | Coro2b | bXPF_tRA |
| 1-878 | chr9 | 62811658 | 62812121 | + | 697 | -241 | Fem1b | bXPF_tRA |
| 1-1009 | chr9 | 63109134 | 63109597 | + | 664 | 37615 | Skor1 | bXPF_tRA |

| | | | | | | | | |
|---|---|---|---|---|---|---|---|---|
| 1-722 | chr9 | 63123918 | 63124381 | + | 559 | 22831 | Skor1 | bXPF_tRA |
| 1-372 | chr9 | 63132755 | 63133218 | + | 572 | 13994 | Skor1 | bXPF_tRA |
| 1-1263 | chr9 | 63716101 | 63716564 | + | 548 | 41662 | Smad3 | bXPF_tRA |
| 1-494 | chr9 | 63757914 | 63758377 | + | 557 | -151 | Smad3 | bXPF_tRA |
| 1-1659 | chr9 | 63896261 | 63896724 | + | 618 | 125567 | Smad6 | bXPF_tRA |
| 1-146 | chr9 | 63896463 | 63896926 | + | 618 | 125365 | Smad6 | bXPF_tRA |
| 1-898 | chr9 | 65032108 | 65032571 | + | 844 | -119 | Dpp8 | bXPF_tRA |
| 1-390 | chr9 | 65032322 | 65032785 | + | 845 | 95 | Dpp8 | bXPF_tRA |
| 1-864 | chr9 | 65196099 | 65196562 | + | 658 | -18360 | Parp16 | bXPF_tRA |
| 1-485 | chr9 | 65196318 | 65196781 | + | 658 | -18141 | Parp16 | bXPF_tRA |
| 1-1305 | chr9 | 65827570 | 65828033 | + | 629 | -237 | Zfp609 | bXPF_tRA |
| 1-1805 | chr9 | 65884887 | 65885350 | + | 974 | 55 | Trip4 | bXPF_tRA |
| 1-1590 | chr9 | 66124579 | 66125042 | + | 876 | 76 | Snx1 | bXPF_tRA |
| 1-1720 | chr9 | 66919323 | 66919786 | + | 679 | 151 | Rab8b | bXPF_tRA |
| 1-1426 | chr9 | 66945758 | 66946221 | + | 722 | -87 | Rps27l | bXPF_tRA |
| 1-119 | chr9 | 66945973 | 66946436 | + | 721 | 128 | Rps27l | bXPF_tRA |
| 1-413 | chr9 | 67043524 | 67043987 | + | 575 | 116 | Tpm1 | bXPF_tRA |
| 42370 | chr9 | 67043837 | 67044300 | + | 626 | -197 | Tpm1 | bXPF_tRA |
| 1-1654 | chr9 | 67228941 | 67229404 | + | 693 | 7554 | Mir190a | bXPF_tRA |
| 1-383 | chr9 | 67229152 | 67229615 | + | 694 | 7343 | Mir190a | bXPF_tRA |
| 1-1791 | chr9 | 69989329 | 69989792 | + | 751 | 94 | Bnip2 | bXPF_tRA |
| 1-349 | chr9 | 72111815 | 72112278 | + | 554 | -227 | Tcf12 | bXPF_tRA |
| 1-319 | chr9 | 72133053 | 72133516 | + | 554 | -21465 | Tcf12 | bXPF_tRA |
| 1-1445 | chr9 | 72274583 | 72275046 | + | 755 | -74 | Zfp280d | bXPF_tRA |
| 1-1290 | chr9 | 73905778 | 73906241 | + | 589 | 27558 | Unc13c | bXPF_tRA |
| 45658 | chr9 | 73905990 | 73906453 | + | 589 | 27346 | Unc13c | bXPF_tRA |
| 1-321 | chr9 | 77655424 | 77655887 | + | 542 | 18923 | Klhl31 | bXPF_tRA |
| 1-1004 | chr9 | 77828443 | 77828906 | + | 670 | 74139 | Gclc | bXPF_tRA |
| 1-1470 | chr9 | 77917145 | 77917608 | + | 631 | 11 | Elovl5 | bXPF_tRA |
| 1-885 | chr9 | 79759445 | 79759908 | + | 543 | 177 | Cox7a2 | bXPF_tRA |
| 1-1598 | chr9 | 83548150 | 83548613 | + | 574 | 43 | Sh3bgrl2 | bXPF_tRA |
| 1-143 | chr9 | 83548356 | 83548819 | + | 574 | 249 | Sh3bgrl2 | bXPF_tRA |
| 1-1006 | chr9 | 84369205 | 84369668 | + | 542 | -404962 | 4930554C24Rik | bXPF_tRA |
| 1-229 | chr9 | 85748702 | 85749165 | + | 570 | 401 | Ibtk | bXPF_tRA |
| 1-883 | chr9 | 92250448 | 92250911 | + | 615 | 485 | Plscr1 | bXPF_tRA |
| 1-1879 | chr9 | 95511834 | 95512297 | + | 639 | -69 | U2surp | bXPF_tRA |
| 1-1416 | chr9 | 97110678 | 97111141 | + | 710 | 132 | Slc25a36 | bXPF_tRA |
| 1-1487 | chr9 | 100440059 | 100440522 | + | 713 | 46183 | Il20rb | bXPF_tRA |
| 1-1094 | chr9 | 100545980 | 100546443 | + | 719 | -77 | Nck1 | bXPF_tRA |
| 1-465 | chr9 | 100643285 | 100643748 | + | 630 | -107 | Stag1 | bXPF_tRA |
| 1-274 | chr9 | 102626025 | 102626488 | + | 634 | -40 | Anapc13 | bXPF_tRA |
| 16072 | chr9 | 102626321 | 102626784 | + | 634 | 256 | Anapc13 | bXPF_tRA |
| 1-594 | chr9 | 102726044 | 102726507 | + | 927 | 8471 | Amotl2 | bXPF_tRA |
| 1-817 | chr9 | 102779431 | 102779894 | + | 656 | 40014 | Gm5627 | bXPF_tRA |
| 14977 | chr9 | 104001941 | 104002404 | + | 569 | -372 | Nphp3 | bXPF_tRA |
| 1-787 | chr9 | 104002288 | 104002751 | + | 569 | -25 | Nphp3 | bXPF_tRA |
| 1-639 | chr9 | 105052967 | 105053430 | + | 553 | -70 | Mrpl3 | bXPF_tRA |
| 1-809 | chr9 | 106418309 | 106418772 | + | 559 | -10871 | Rpl29 | bXPF_tRA |
| 7672 | chr9 | 106429230 | 106429693 | + | 860 | 50 | Rpl29 | bXPF_tRA |
| 1-1197 | chr9 | 107296545 | 107297008 | + | 663 | 117 | Cish | bXPF_tRA |
| 1-1314 | chr9 | 108566733 | 108567196 | + | 623 | 378 | Ndufaf3 | bXPF_tRA |
| 1-480 | chr9 | 109059084 | 109059547 | + | 662 | -64 | Trex1 | bXPF_tRA |
| 1-254 | chr9 | 109059363 | 109059826 | + | 662 | 129 | Trex1 | bXPF_tRA |
| 1-689 | chr9 | 109832604 | 109833067 | + | 672 | 41 | Nme6 | bXPF_tRA |
| 1-1286 | chr9 | 109931290 | 109931753 | + | 541 | -253 | Map4 | bXPF_tRA |
| 1-450 | chr9 | 109931494 | 109931957 | + | 545 | -49 | Map4 | bXPF_tRA |
| 1-1702 | chr9 | 110448903 | 110449366 | + | 775 | 27560 | Klhl18 | bXPF_tRA |
| 1-1861 | chr9 | 110476421 | 110476884 | + | 733 | 42 | Klhl18 | bXPF_tRA |
| 1-339 | chr9 | 110781388 | 110781851 | + | 829 | -16566 | Prss42 | bXPF_tRA |
| 1-1259 | chr9 | 110790442 | 110790905 | + | 730 | -7512 | Prss42 | bXPF_tRA |
| 1-1588 | chr9 | 113018996 | 113019459 | + | 699 | 567925 | AU023762 | bXPF_tRA |
| 1-1028 | chr9 | 114640053 | 114640516 | + | 553 | -84 | Cnot10 | bXPF_tRA |
| 1-1424 | chr9 | 116375957 | 116376420 | + | 668 | -200825 | Tgfbr2 | bXPF_tRA |

| | | | | | | | | |
|---|---|---|---|---|---|---|---|---|
| 1-214 | chr9 | 117252263 | 117252726 | + | 603 | -11 | Rbms3 | bXPF_tRA |
| 1-399 | chr9 | 117252618 | 117253081 | + | 540 | -366 | Rbms3 | bXPF_tRA |
| 26665 | chr9 | 117252837 | 117253300 | + | 540 | -585 | Rbms3 | bXPF_tRA |
| 1-468 | chr9 | 118040118 | 118040581 | + | 767 | -173 | Azi2 | bXPF_tRA |
| 1-974 | chr9 | 119339598 | 119340061 | + | 649 | 211 | Myd88 | bXPF_tRA |
| 1-444 | chr9 | 120691306 | 120691769 | + | 626 | 114206 | 5830454E08Rik | bXPF_tRA |
| 1-1247 | chr9 | 123020990 | 123021453 | + | 719 | -105 | Tmem42 | bXPF_tRA |
| 1-1609 | chr9 | 123529224 | 123529687 | + | 701 | -427 | Sacm1l | bXPF_tRA |
| 13516 | chr7 | 16924056 | 16924495 | + | 1000 | -243 | Calm3 | bXPF_UV |
| 14977 | chr8 | 19784486 | 19784925 | + | 1000 | 55129 | Gm21119 | bXPF_UV |
| 42795 | chr1 | 88277453 | 88277892 | + | 1000 | -93 | Hjurp | bXPF_UV |
| 15707 | chr1 | 88277247 | 88277686 | + | 1000 | 91 | A730008H23Rik | bXPF_UV |
| 16072 | chr17 | 84186325 | 84186526 | + | 1000 | 1522 | Zfp36l2 | bXPF_UV |
| 11689 | chr17 | 84186435 | 84186874 | + | 1000 | 1293 | Zfp36l2 | bXPF_UV |
| 10959 | chr6 | 137558987 | 137559426 | + | 995 | 11803 | Eps8 | bXPF_UV |
| 12420 | chr18 | 32836992 | 32837431 | + | 963 | -14 | Wdr36 | bXPF_UV |
| 13150 | chr7 | 79466114 | 79466553 | + | 935 | -60 | Polg | bXPF_UV |
| 12785 | chr8 | 120588973 | 120589412 | + | 910 | -117 | Gins2 | bXPF_UV |
| 14611 | chr4 | 45341777 | 45342216 | + | 890 | -105 | Dcaf10 | bXPF_UV |
| 42856 | chr11 | 120256677 | 120257116 | + | 872 | 23949 | Bahcc1 | bXPF_UV |
| 15342 | chr9 | 123461936 | 123462075 | + | 855 | -16696 | Limd1 | bXPF_UV |
| 14246 | chr6 | 22280303 | 22280742 | + | 841 | -7705 | Wnt16 | bXPF_UV |
| 42736 | chr11 | 116843444 | 116843883 | + | 827 | 148 | Mettl23 | bXPF_UV |
| 46388 | chr5 | 108065366 | 108065805 | + | 814 | -89 | Mtf2 | bXPF_UV |
| 44197 | chr2 | 117249572 | 117250011 | + | 800 | 52 | Fam98b | bXPF_UV |
| 43070 | chr10 | 80249118 | 80249557 | + | 787 | -115 | Ndufs7 | bXPF_UV |
| 46023 | chr10 | 80248905 | 80249344 | + | 776 | -328 | Ndufs7 | bXPF_UV |
| 42979 | chr5 | 30888123 | 30888562 | + | 742 | -510 | Agbl5 | bXPF_UV |
| 45292 | chr5 | 30887910 | 30888349 | + | 731 | -723 | Agbl5 | bXPF_UV |
| 43831 | chr1 | 139195186 | 139195625 | + | 719 | 181671 | Crb1 | bXPF_UV |
| 42948 | chr6 | 71633033 | 71633472 | + | 708 | -335 | Kdm3a | bXPF_UV |
| 45658 | chr5 | 149228890 | 149229329 | + | 699 | -35895 | Alox5ap | bXPF_UV |
| 42826 | chr7 | 79466318 | 79466757 | + | 688 | -264 | Polg | bXPF_UV |
| 42917 | chr7 | 99360450 | 99360889 | + | 678 | -7430 | Serpinh1 | bXPF_UV |
| 42370 | chr7 | 99360251 | 99360690 | + | 669 | -7231 | Serpinh1 | bXPF_UV |
| 43009 | chr14 | 31669183 | 31669622 | + | 661 | 28345 | Btd | bXPF_UV |
| 43101 | chr11 | 75166687 | 75167126 | + | 653 | 1246 | Hic1 | bXPF_UV |
| 11324 | chr5 | 88553995 | 88554434 | + | 645 | -269 | Utp3 | bXPF_UV |
| 43466 | chr13 | 44569375 | 44569814 | + | 637 | 129867 | 1700029N11Rik | bXPF_UV |
| 41275 | chr1 | 161968988 | 161969427 | + | 628 | 19 | Pigc | bXPF_UV |
| 1 | chr17 | 35051394 | 35051833 | + | 619 | 1370 | Clic1 | bXPF_UV |
| 42767 | chr17 | 35051195 | 35051634 | + | 611 | 1171 | Clic1 | bXPF_UV |
| 42887 | chr13 | 60176151 | 60176590 | + | 604 | 1165 | Gas1 | bXPF_UV |
| 46753 | chr18 | 79467065 | 79467504 | + | 597 | -357893 | Setbp1 | bXPF_UV |
| 13881 | chr11 | 93968096 | 93968535 | + | 590 | 206 | Nme1 | bXPF_UV |
| 42005 | chr2 | 26964318 | 26964757 | + | 583 | -151 | Rexo4 | bXPF_UV |
| 47119 | chr14 | 54641283 | 54641722 | + | 576 | -138 | Cdh24 | bXPF_UV |
| 12055 | chr14 | 47863006 | 47863445 | + | 570 | -35639 | 4930447J18Rik | bXPF_UV |
| 41640 | chr8 | 22411120 | 22411559 | + | 563 | -1 | Mrps31 | bXPF_UV |
| 43040 | chr18 | 47368699 | 47369138 | + | 556 | -48 | Sema6a | bXPF_UV |
| 44562 | chr16 | 17276290 | 17276729 | + | 550 | 209 | Tmem191c | bXPF_UV |
| 44927 | chr9 | 67043499 | 67043938 | + | 543 | 153 | Tpm1 | bXPF_UV |
