## Supplementary Table S2 for "ERCC1-XPF Interacts with Topoisomerase IIβ to Facilitate the Repair of Activity-induced DNA Breaks"

### Supplementary Table S2. bXPF-bound genes (ChIP-Seq) with ≥ 20 counts (RNA-Seq) in untreated and tRA-treated MEFs

| bXPF-bound genes with ≥ 20 counts (RNA-Seq) | tRA_bXPF-bound genes with ≥ 20 counts (RNA-Seq) |
|---|---|
| Gm21119 | Mfsd7b |
| Pdpn | Tmem5 |
| Def8 | Rnmtl1 |
| Mir7009 | Fam58b |
| Rmrp | LOC100503496 |
| Grcc10 | A630020A06 |
| Morf4l1-ps1 | Fam134b |
| 4930486I03Rik | Morf4l1-ps1 |
| Fam105a | E130310I04Rik |
| Mir721 | Gm10789 |
| Gm2560 | 1110004E09Rik |
| 9530091C08Rik | Gm6402 |
| 4930579J19Rik | Sgol1 |
| BC100451 | Hdgfrp2 |
| Lrp11 | Apbb3 |
| Mir8115 | 1500017E21Rik |
| Slc35f4 | Gm17762 |
| 4933433H22Rik | Gm3230 |
| Mirlet7i | Tmem261 |
| Mir7681 | Bre |
| Vwa9 | 1700007G11Rik |
| Tmem179b | Gbas |
| Mir694 | 3110021A11Rik |
| 2810442N19Rik | Mesdc1 |
| Smek1 | Micalcl |
| Sgol2b | 9330133O14Rik |
| Mir1901 | Snora41 |
| Mettl26 | Mir6353 |
| Zbtb21 | Iqca |
| Mir455 | B230216N24Rik |
| Psapl1 | Mir6346 |
| Oat | 2810442N19Rik |
| 9030617O03Rik | Smlr1 |
| Mir7227 | Grip1os2 |
| Mir1931 | Mirlet7i |
| Gm17801 | Hist3h2a |
| Tmem2 | Mir21a |
| Rnmtl1 | Mir8115 |
| 1700084J12Rik | Mir6931 |
| 5930430L01Rik | Nmt1 |
| Mir8105 | Mir1843a |
| Atad2 | 4930557F10Rik |
| Ccdc121 | Mir6391 |
| 1600002D24Rik | Zbtb21 |
| Gm16833 | Dynlt1b |
| 2810428I15Rik | Mir7b |
| Arhgap15os | Gm10190 |

| | |
|---|---|
| Tmem261 | Vaultrc5 |
| Mir1843a | 1700034E13Rik |
| Trex1 | Mir694 |
| Mir7689 | Aip |
| Gm13090 | Carns1 |
| Mesdc2 | Vps51 |
| 4931430N09Rik | Ehd1 |
| 4930555B11Rik | Mir6991 |
| Vasn | Hps6 |
| Mascrna | Mir6995 |
| Mir29b-2 | Phyhd1 |
| 9330133O14Rik | 4930555B11Rik |
| 4930563F08Rik | Platr26 |
| Mir1938 | Mall |
| Pcdhgc3 | 2010009K17Rik |
| Rbm14 | Mir7009 |
| 4933402D24Rik | Mir760 |
| Serpinh1 | Rmrp |
| Postn | Mir6398 |
| Fstl1 | Rnu11 |
| Tpm4 | Mir7227 |
| Timp3 | Pdpn |
| Itgb1 | Gm13090 |
| Timp2 | 4930563F08Rik |
| Axl | Mir721 |
| App | Grcc10 |
| Rpl8 | 2410002F23Rik |
| Sept11 | Ate1 |
| Fbln5 | Nsmce4a |
| Cd44 | 2310057M21Rik |
| Iqgap1 | 1810010D01Rik |
| Atp1a1 | Gm21119 |
| Unc5b | Mir1957b |
| Fam129b | Zfpm1 |
| Parva | Mir100 |
| Marcks | Mir190a |
| Slc25a4 | Klhl31 |
| Ehd2 | Trex1 |
| Kctd10 | Myd88 |
| Cyb5r3 | Col1a1 |
| Map4 | Tpm1 |
| Slc6a6 | Col5a2 |
| Aars | Myh9 |
| Adam19 | Serpinh1 |
| Itgav | Actn1 |
| Ergic1 | Cald1 |
| Rps12 | Lgals1 |
| Cdkn1a | Scd2 |
| Fgfr1 | Rplp0 |
| Vdac1 | Flnb |

| | |
|---|---|
| Ubc | Cyr61 |
| Ago2 | Fstl1 |
| Ywhaz | Thbs2 |
| Map4k4 | F2r |
| G3bp1 | Eif4g2 |
| Brd2 | Fbn1 |
| Top2a | Zyx |
| 2310022B05Rik | S100a6 |
| Stat3 | Ahnak |
| Prdx1 | Ddb1 |
| Wwc2 | Rplp1 |
| Prpf40a | Hsp90b1 |
| Farp1 | Eef1g |
| Hic1 | Kpnb1 |
| Vgll3 | Hdlbp |
| Bag6 | Rpl8 |
| Esyt2 | Amotl2 |
| Plekhg2 | Tead1 |
| Cyth3 | Hic1 |
| Hnrnpul1 | Cmss1 |
| Slc3a2 | Ctsb |
| Rpl37 | Pcolce |
| Eva1b | Hnrnpab |
| Tspan9 | Sept7 |
| Tnpo1 | Calm1 |
| Ncam1 | Parva |
| Nrp2 | Mlec |
| Csnk1d | Rpl36 |
| Asap1 | Kctd10 |
| Rbfox2 | Igf2r |
| Mcl1 | Slc25a4 |
| Smtn | Calm3 |
| Usp47 | Amotl1 |
| Actr1b | Tubb4b |
| Eif3e | Rpl18 |
| Ston1 | Pdlim7 |
| Gtf2i | Rpn2 |
| Cyp51 | Eef1b2 |
| Dlc1 | Pdlim5 |
| Plpp1 | Cap1 |
| Ube2i | Phldb2 |
| Arhgef40 | Rpl5 |
| B4galt1 | Clic4 |
| Hnrnpm | Map4 |
| Arl5a | Id3 |
| Arid5b | Rpl11 |
| Tspan4 | Hmga2 |
| Lsp1 | Ltbp1 |
| Sh3glb1 | Wls |
| Rnd3 | Htra1 |

| | |
|---|---|
| Frmd4a | Cdk4 |
| Rpl30 | Marcks |
| Pcdh7 | Ubc |
| Mcm7 | Cd63 |
| Traf7 | H3f3b |
| Errfi1 | Fam129b |
| Arih1 | Gas1 |
| Ext1 | Cxcl12 |
| Ctdsp1 | Hnrnpa0 |
| Rnf187 | Atf5 |
| Pmp22 | Ubb |
| Lrrc17 | Srsf2 |
| Bicc1 | Trip12 |
| Rbm25 | Rps29 |
| Pdzrn3 | G3bp1 |
| Htra1 | Grn |
| Eif3h | 2310022B05Rik |
| Nfat5 | Brd2 |
| Tomm40 | Atxn2l |
| Osmr | Jak1 |
| Rpl29 | Nuak1 |
| Zfp503 | Mlf2 |
| Fgfr2 | Dhx9 |
| Strn3 | Tsc22d2 |
| Rusc2 | Rps20 |
| Fmnl3 | Wwc2 |
| Mycbp2 | Nfix |
| Asph | Ddost |
| Epha3 | Prdx1 |
| Plxdc2 | Pea15a |
| Notch1 | Gadd45g |
| Foxp1 | Rad23b |
| Mkl1 | Sdc4 |
| Srsf7 | Pcbp2 |
| St5 | Sipa1l1 |
| Etv6 | Anapc5 |
| Cops8 | Eif5 |
| Cpsf1 | Tsc22d1 |
| Apbb2 | Bag6 |
| Lrig1 | Rcn1 |
| Smad7 | Calm2 |
| Rab21 | Abl1 |
| Rbms3 | Filip1l |
| Fyn | Yap1 |
| Rab8a | Npc2 |
| Lurap1l | Ano6 |
| Ddx18 | Rbm39 |
| Mpdz | Plekhg2 |
| Adm | Sgpl1 |
| Bod1l | Rpl27 |

| | |
|---|---|
| Ccdc88a | Zfp36l2 |
| Fam107b | Rpl37 |
| Ephb3 | Cct4 |
| Bms1 | Ppp2r1a |
| Vcam1 | Hnrnpul1 |
| Gipc1 | Peak1 |
| Eps8 | Tnpo1 |
| 2010111I01Rik | Asah1 |
| Twist1 | Actr2 |
| Sema3c | Dhrs3 |
| Irs1 | Trio |
| Slc9a1 | Rarg |
| Man1c1 | Stat3 |
| Ap1g1 | Csnk1d |
| Aaas | Usp47 |
| Tjap1 | Eif3f |
| Pdgfc | Sulf1 |
| Golga7 | Emilin1 |
| Ptpn12 | Ephb2 |
| Tbcd | Ddah1 |
| Anapc4 | Psmc3 |
| Ist1 | Rbfox2 |
| Arid1b | Metrnl |
| Rgl1 | Scarf2 |
| Utp3 | Selenof |
| Tgfbr3 | Rusc2 |
| Ubald1 | Hmgcr |
| Rnase4 | Fads2 |
| Micu1 | Arid1a |
| Edem3 | Frmd4a |
| Nup133 | Ncam1 |
| Foxn3 | Snrpb |
| Paf1 | M6pr |
| Kansl3 | Gtf2i |
| Mfap2 | Fam107b |
| BC004004 | Ube2i |
| Ube2j2 | Fkbp4 |
| Fem1b | Tpx2 |
| Setd1b | Pcdh7 |
| Chmp2a | Mob1a |
| Vps52 | Dlc1 |
| Stx16 | Tcf3 |
| Atrnl1 | Dlg1 |
| Zfp827 | Smad3 |
| Emg1 | Ext1 |
| Opa1 | Il1rl1 |
| Atxn1 | Gorasp2 |
| Zfp462 | Bcl9l |
| Syne3 | Pip5k1c |
| Spred2 | Got2 |

| | |
|---|---|
| Rbm8a | St3gal2 |
| Ptprm | Ints1 |
| Chd9 | Rab5c |
| Ccl2 | Mcl1 |
| Etl4 | Ube2z |
| Antxr2 | Clptm1l |
| Kdelc2 | Arih1 |
| Zfp521 | Xpo1 |
| Cstf2t | Luc7l2 |
| Nvl | Eva1b |
| Slc30a9 | Cyth3 |
| Nupr1 | Klf3 |
| Ccl7 | Lhfp |
| Pfdn1 | Nrp2 |
| Mrpl3 | Lama4 |
| Trim32 | Rpl30 |
| Gde1 | Afap1 |
| Maml2 | Tcf12 |
| Gpatch8 | Mcm7 |
| Limk1 | Shc1 |
| Polr1b | Setd5 |
| Gins2 | Sh3gl1 |
| Arhgap28 | Dync1li2 |
| Dtnbp1 | Birc6 |
| Wdr11 | Sh3glb1 |
| Nr1d1 | Atic |
| Fzd5 | Ndufs2 |
| Sde2 | C1qbp |
| Cox7a2 | Srsf11 |
| Paxbp1 | Mgat1 |
| Vgll4 | Prkci |
| Pcgf2 | Elovl5 |
| Foxj3 | Impad1 |
| Rbpms | Eif4a2 |
| Dcp2 | Cluh |
| Capn15 | H6pd |
| Dnm3os | Ets1 |
| Unc5c | Svep1 |
| Cpped1 | Dot1l |
| Ndufs7 | Tor1aip2 |
| Zswim6 | Ggta1 |
| Fmnl2 | Nudc |
| Pptc7 | Fez2 |
| Cblb | Hspe1 |
| Ttc4 | St3gal1 |
| Mcu | Junb |
| B3gnt9 | Trpc4ap |
| Ap3s2 | Ctdsp1 |
| Fzd8 | Ajuba |
| Aatf | Dok1 |

| | |
|---|---|
| Trps1 | Zfp703 |
| Krr1 | Pja2 |
| Mecom | Anp32e |
| Gtf3c5 | Carhsp1 |
| Fcho2 | Rgl1 |
| Med17 | Zfp36l1 |
| Tmem45a | Kmt5a |
| Gne | Ddx23 |
| Golga5 | Sod1 |
| Anks3 | Xrn2 |
| Rmnd5a | Kctd5 |
| Ank2 | Rap1b |
| Yipf4 | Osbpl5 |
| Bmt2 | Pten |
| Dcp1a | Slc25a24 |
| Cfap20 | Akr1b3 |
| Rhbdd1 | H1f0 |
| Smg8 | Bnip2 |
| Nup54 | Wbp1l |
| Mrpl39 | Mib1 |
| Magi3 | Eif3h |
| Ubiad1 | Plpp3 |
| Kdm3a | Sorbs1 |
| Gtf3c3 | Selenot |
| Mrpl21 | Pdia5 |
| Gid4 | Zc3h4 |
| Cox15 | Rpl29 |
| Usp30 | Pigt |
| Nbn | Jag1 |
| Mgat5 | Ermp1 |
| Trip4 | Trim44 |
| Naa30 | Tmem127 |
| Zc2hc1a | Serpinb6a |
| Vti1a | Nfe2l2 |
| Gucd1 | Rnf44 |
| Mpdu1 | Tcerg1 |
| Slc33a1 | Pes1 |
| Arnt2 | Gtse1 |
| Gcnt2 | Grk2 |
| Exosc4 | Insig1 |
| Hoxa10 | Mapk14 |
| Nabp1 | Azin1 |
| Usp6nl | Lmo7 |
| Dnajc16 | Arid5b |
| Zfp618 | Rcn2 |
| Vps13b | Bag1 |
| Fpgt | Galnt10 |
| Mecr | Spop |
| Numb | Api5 |
| Plcxd2 | Asph |

| | |
|---|---|
| Mnat1 | Smad7 |
| Atad2b | Pcna |
| Scyl3 | Lurap1l |
| Kxd1 | Cdh2 |
| Pvt1 | Dnajc5 |
| Eefsec | Ice1 |
| Slc4a1ap | Rps27l |
| Atg7 | Eif4e2 |
| Traf3ip2 | Rere |
| Srgap3 | Zfp503 |
| Cyb5r4 | Cited2 |
| Ercc4 | Wapl |
| 2510009E07Rik | Zfhx4 |
| Mpg | Dpp8 |
| Aig1 | Cpsf1 |
| Bcl2l11 | Ankrd40 |
| Cdc42ep2 | Dvl3 |
| Rora | Rab8a |
| Zfp608 | Tnfrsf1b |
| Sorbs2 | Rab11b |
| Slc1a3 | Psmc2 |
| Hunk | Larp4 |
| Pkig | Pdzrn3 |
| Dmap1 | Scarb2 |
| Cdc25c | Irf2bp2 |
| Gdpd5 | Pank3 |
| Capn10 | Tsku |
| Ppil3 | Adamts9 |
| Tenm4 | Prmt5 |
| Tsen15 | Gak |
| Zbtb40 | U2surp |
| Nog | Htra2 |
| Mpc2 | Kmt2a |
| Prdm5 | Srxn1 |
| Sdk1 | Eloa |
| Arhgap22 | Rragc |
| Bnc2 | Mob3a |
| Zfp90 | Smc5 |
| Ints12 | Polg |
| Runx1t1 | Prrx2 |
| Bcs1l | Dpp3 |
| Anapc13 | Nus1 |
| Pex3 | Map3k7 |
| Garem1 | Elovl6 |
| Ahi1 | Ankfy1 |
| Fam173a | Cers6 |
| Helq | Rasa3 |
| Cenpu | Wdr36 |
| Foxp2 | Zmpste24 |
| Tec | Dysf |

| | |
|---|---|
| Srek1ip1 | Maml1 |
| Alkbh3 | Mdc1 |
| Mafb | Eps8 |
| Mettl21a | Cops8 |
| Atxn7l1 | Agfg1 |
| Nfia | Dbnl |
| Abtb2 | Zfp827 |
| Zfp35 | Nudcd3 |
| Mmp3 | Fibin |
| Slc5a6 | Epha3 |
| Hps3 | Camk2d |
| Zbtb6 | Dse |
| Slc24a3 | Spcs1 |
| Pcsk5 | Trap1 |
| Mrpl44 | Rbck1 |
| Elfn1 | Etv6 |
| Zc3h3 | Phlda1 |
| Bmpr1b | Vapa |
| Tbc1d7 | Gfpt1 |
| Tyw5 | Raf1 |
| Malsu1 | Fmnl3 |
| Pdcd4 | Gys1 |
| Utp23 | Ddx18 |
| Lrif1 | Sesn2 |
| Zfp398 | Dab2 |
| Taco1 | Dicer1 |
| Fam102a | Sf3a3 |
| 5031425E22Rik | Kifc1 |
| Slc25a19 | Bms1 |
| Ldb2 | Ap1g1 |
| Slc25a42 | Lrrc58 |
| Bckdhb | Fastk |
| Sertad3 | Etnk1 |
| Ptrhd1 | Bahcc1 |
| Kctd7 | Ubxn1 |
| Slx4ip | Npepl1 |
| Ppcs | Klhl22 |
| Zfp105 | Tmem167 |
| Tfap2a | Gemin5 |
| March3 | Pdgfc |
| Ccdc84 | Baz2a |
| Cdk5rap1 | Rbpms |
| Sema6c | Gypc |
| Akap7 | Arhgef7 |
| Yipf2 | Nsfl1c |
| Dennd6b | Strip1 |
| Jund | Ccdc102a |
| Dclre1c | Psma1 |
| Mettl25 | Ralgapb |
| Dffb | Tsn |

| | |
|---|---|
| Astn2 | Neat1 |
| Tmem154 | Pgp |
| Ccdc57 | Capza1 |
| Tet1 | 2010111I01Rik |
| Arid3b | Rbms3 |
| Irgm1 | Dusp5 |
| Zfp560 | Faf1 |
| Ttc26 | Sp3 |
| Lrrtm3 | Acap2 |
| Commd1 | Kctd12 |
| Serp2 | Lrrc8d |
| Zfp709 | Psmb2 |
| Aox1 | Cdkn2b |
| Mrnip | Drg2 |
| Trp53rkb | Adpgk |
| Ulk4 | Usp36 |
| Rad51b | Cdc42se1 |
| Napepld | Zfp148 |
| Cacna2d3 | Tfip11 |
| Atg10 | Shoc2 |
| Aph1c | Dars |
| Cyp7b1 | Chordc1 |
| Ddc | Rnf14 |
| Gch1 | Nub1 |
| Zc3h6 | Ier3 |
| Car10 | Atp5g2 |
| Gm20755 | Celf2 |
| Gprin3 | Stx16 |
| 9330111N05Rik | 1110004F10Rik |
| Cdh26 | Anapc4 |
| Bmf | Bud31 |
| Fmn2 | Ndufs1 |
| Hexim2 | Mga |
| Syn3 | Stag1 |
| Eva1c | Vps13d |
| Poln | Itfg1 |
| Usp18 | Rarb |
| Rarb | Map3k3 |
| Lyl1 | Tgif1 |
| 3110039I08Rik | Smu1 |
| Ntng1 | Smad4 |
| Gm19705 | Ddx50 |
| 9530026P05Rik | Abca1 |
| 1810024B03Rik | Ppp3ca |
| Clic5 | Hes1 |
| Sept2 | Umps |
| Hsf2bp | Abhd4 |
| BC037032 | Ddit4 |
| Zc2hc1c | Sertad4 |
| 8430436N08Rik | Rnf6 |

| | |
|---|---|
| 5031434O11Rik | Atxn1 |
| Atp6v0a4 | Nufip2 |
| Gm15413 | Klhdc2 |
| Slc14a2 | Tcea1 |
| Gsg1 | Atg2b |
| Chrna1 | Ddx56 |
| Ccdc81 | Nat10 |
| Gm16793 | Mapre2 |
| Pax8 | Irs1 |
| Gm20753 | Mmadhc |
| 1700096K18Rik | Sacm1l |
| Nalcn | H2afy2 |
| Ankrd2 | Tmem65 |
| 1700113A16Rik | Spred2 |
| Tspan32 | Letm1 |
| Gm973 | Ruvbl1 |
| Slc10a6 | Drg1 |
| Itgb6 | Smarcd2 |
| Mir684-1 | Tor1aip1 |
| Gm29683 | Fgfrl1 |
| 6430562O15Rik | Crat |
| Opn5 | Klf2 |
| Npm2 | Opa1 |
| Nme1 | Aqr |
| Dzank1 | Ets2 |
| 1700028E10Rik | Tgfb1 |
| Cpa2 | Cd24a |
| 4933412O06Rik | Mad2l1 |
| Mir199a-1 | Fem1b |
| Gabrp | Emg1 |
| Rimbp3 | Irx1 |
| 1700108F19Rik | Cdk5rap2 |
| Capn12 | Mtmr6 |
| 1700045H11Rik | Vps52 |
| Kpna7 | Ube2j2 |
| Gm4861 | Pde4a |
| Caln1 | Dcaf12 |
| 2010109A12Rik | Zfp609 |
| D930032P07Rik | Ppt1 |
| Gm12596 | Sec22b |
| Gm15401 | Zfp384 |
| Lncbate1 | Mrpl17 |
| | Otud6b |
| | Rexo4 |
| | Nln |
| | Tspan5 |
| | Wrb |
| | Chd9 |
| | Mrpl20 |
| | Wdr81 |

| |
|---|
| Ppip5k2 |
| Slc8a1 |
| Zswim6 |
| Rab8b |
| Mapkbp1 |
| Bcl10 |
| Dnajc9 |
| Spaca6 |
| Atf7ip |
| Htt |
| Brpf1 |
| Rnf41 |
| Rnase4 |
| Kctd9 |
| Ppp1r11 |
| Dhx16 |
| Ccz1 |
| Srgap3 |
| Tob2 |
| Nr2f2 |
| Cc2d1b |
| Sppl2b |
| Polr3e |
| Ccnl1 |
| Tbc1d13 |
| Snx1 |
| Nbeal1 |
| Syde2 |
| Cnot10 |
| Ccdc97 |
| Pxk |
| Ibtk |
| Kif13a |
| Eif2b1 |
| Rsrc2 |
| Osbpl3 |
| Trim32 |
| Ppard |
| Btg1 |
| Cd47 |
| Irx2 |
| Setbp1 |
| Prdm4 |
| Tgfbr3 |
| Mrpl3 |
| Nuf2 |
| Ttc14 |
| Azi2 |
| Ube2f |
| Cipc |

| |
|---|
| Knl1 |
| Acp2 |
| Ttc19 |
| Eif1ad |
| Uqcc2 |
| Sppl2a |
| Limk1 |
| Echs1 |
| Gpatch8 |
| Spty2d1 |
| Chd2 |
| Gabpb2 |
| Adra1d |
| Rubcn |
| Slc25a36 |
| Pold2 |
| Rbm18 |
| Ercc1 |
| Eny2 |
| Atl2 |
| Ercc6l2 |
| Gins2 |
| Zfp280d |
| Mrps2 |
| Paxbp1 |
| Wdr11 |
| Ddx41 |
| Rsrc1 |
| Selenok |
| Gpd2 |
| Gosr1 |
| Ndufs7 |
| Smarcd1 |
| Srp9 |
| Cep350 |
| Nosip |
| Cstf2t |
| Polr1b |
| Aimp1 |
| 2410006H16Rik |
| Mrps15 |
| Tmem167b |
| Fam3c |
| Bnip3l |
| Cox7a2 |
| Dus1l |
| Zfp451 |
| Vps72 |
| Ints7 |
| Wtap |

| |
|---|
| Wrnip1 |
| Foxj3 |
| Itpa |
| Ndst2 |
| Stx5a |
| Vwa8 |
| Fkbp2 |
| Tnfrsf23 |
| Polr3a |
| Capn7 |
| Rcbtb2 |
| 9430015G10Rik |
| Nmt2 |
| Rnpc3 |
| Dpy19l4 |
| Bbx |
| Fam84b |
| Clmp |
| Nsmce1 |
| Man1a |
| Fam20c |
| Fig4 |
| Ctnnd1 |
| Gtpbp6 |
| St3gal3 |
| Naxd |
| Spsb1 |
| Uspl1 |
| Trim41 |
| Snn |
| Id4 |
| Ap3s2 |
| Gpbp1 |
| Rreb1 |
| Trappc9 |
| Trib1 |
| Ttc4 |
| Mat2b |
| Naa20 |
| Cep120 |
| Runx2 |
| Rrp15 |
| Klhdc4 |
| Rgl2 |
| Ino80 |
| Adgrl3 |
| Alad |
| Fam98b |
| Net1 |
| Sde2 |

| |
|---|
| Nrep |
| Vgll4 |
| Nenf |
| Prcc |
| Robo1 |
| Apbb1ip |
| Med23 |
| Mcph1 |
| Tada3 |
| Gne |
| Klhl18 |
| Zfp142 |
| Nup35 |
| Abhd17c |
| Ptdss2 |
| Gm1821 |
| Pfkm |
| Orc3 |
| Abhd13 |
| Mis18bp1 |
| Tbc1d23 |
| Tmem222 |
| Nol12 |
| Srp19 |
| Cyb5a |
| N4bp2l2 |
| Mrpl30 |
| Polr3b |
| Nectin3 |
| Gli2 |
| Hmg20a |
| Gtf3c3 |
| Cep85 |
| Slc39a11 |
| Mrpl39 |
| Mrpl13 |
| Ank2 |
| Zfp704 |
| Dsel |
| Hs1bp3 |
| Smg8 |
| Dcp1a |
| Med17 |
| Noc3l |
| Snx11 |
| Lgr4 |
| Krr1 |
| Rbl1 |
| Bcl7b |
| Naa30 |

| |
|---|
| Cnnm4 |
| Rnf24 |
| Mrpl21 |
| Socs7 |
| Ebf1 |
| Cdkn2c |
| Mlst8 |
| Cfap20 |
| Haus8 |
| Entpd6 |
| Mtf2 |
| B4galt7 |
| Jmjd4 |
| Dcaf10 |
| Dstyk |
| Alg9 |
| Kdsr |
| Adprhl2 |
| Cfh |
| Nfyb |
| Arnt2 |
| Bola2 |
| Hmgxb4 |
| Sp2 |
| Fance |
| Coq9 |
| Ubiad1 |
| Zfp282 |
| Kdm3a |
| Fntb |
| Pex2 |
| Myo7a |
| Gucd1 |
| Slc35a1 |
| Eefsec |
| Arhgap18 |
| Nampt |
| Trip4 |
| Ercc5 |
| Atad2b |
| Rassf2 |
| Haus4 |
| Tor4a |
| Rnf150 |
| Ddhd2 |
| Gid4 |
| Cdkn2aipnl |
| Polr2j |
| Ell2 |
| Tmx4 |

| |
|---|
| Nfkbia |
| Fnip2 |
| Rps19bp1 |
| Vti1a |
| Gabpb1 |
| Ddit3 |
| Cops7b |
| Snip1 |
| Mtmr14 |
| Ubl3 |
| Fam149b |
| Acadm |
| Crlf3 |
| Numb |
| Exosc4 |
| Vps13b |
| Pnpla6 |
| Pla2g6 |
| Ttc7 |
| Med4 |
| Bmp2k |
| Fchsd2 |
| Zfp639 |
| Nbn |
| Slc15a4 |
| Cdkn2a |
| Epc1 |
| Ptch1 |
| Glt8d1 |
| Ufl1 |
| Fbxo34 |
| Hnrnph3 |
| 5730455P16Rik |
| Tead4 |
| Gadd45a |
| Mipol1 |
| Nck1 |
| Rab33b |
| Cirbp |
| Ddx20 |
| Gtf2b |
| Usp6nl |
| Mylip |
| Spopl |
| Traf3ip2 |
| Akap10 |
| Tmem185b |
| Tmem256 |
| Zfp608 |
| Snx15 |

| |
|---|
| Slc4a1ap |
| Fbxw17 |
| Sspn |
| Snrnp48 |
| Nufip1 |
| Peli1 |
| Btd |
| Gnpnat1 |
| Aars2 |
| Dmap1 |
| Zbtb45 |
| Psmg2 |
| Naaa |
| Tab1 |
| Gfm2 |
| Spata6 |
| Ednra |
| Snx21 |
| Dram2 |
| Hars2 |
| Rbl2 |
| Zfp654 |
| Cdc25c |
| Ccdc58 |
| 4931406C07Rik |
| Lnx2 |
| 2410004B18Rik |
| Zkscan8 |
| Srd5a3 |
| Tia1 |
| Stx18 |
| Arrdc3 |
| Cox19 |
| Smim12 |
| 1110038F14Rik |
| Usp20 |
| Prkra |
| Ccdc71l |
| Ubxn2a |
| Narfl |
| Hs3st1 |
| Tmem260 |
| Crocc |
| Mrpl57 |
| Ssbp1 |
| Pus10 |
| BC055324 |
| Fbxw7 |
| Prdm5 |
| St3gal4 |

| |
|---|
| Bcl11b |
| Ndufa5 |
| Steap1 |
| Inip |
| Nsmce2 |
| Plscr1 |
| Ppil3 |
| Fam173a |
| Ccdc91 |
| Fam133b |
| Trmt10c |
| Zfp668 |
| Mllt3 |
| Arl15 |
| Dexi |
| Tbp |
| Plekhg1 |
| Tbc1d19 |
| Cep152 |
| Csgalnact1 |
| Tfcp2 |
| Zfp623 |
| Nceh1 |
| Helq |
| Mon1b |
| Nsun4 |
| Dnajc19 |
| Btrc |
| Sdk1 |
| Tmem101 |
| Tbc1d9 |
| Cpeb1 |
| Inpp5k |
| Eef2kmt |
| Zfp212 |
| Nr2c1 |
| Appl2 |
| Pex3 |
| Pcca |
| Zfp770 |
| Alkbh3 |
| Rars2 |
| Dguok |
| Zfp511 |
| Garem1 |
| Snapc3 |
| Tma16 |
| Cish |
| Maip1 |
| Ap5m1 |

| |
|---|
| Cgrrf1 |
| Thg1l |
| Zscan21 |
| Tldc1 |
| St7 |
| Mrps31 |
| Pstk |
| Anapc13 |
| A430005L14Rik |
| Ctu1 |
| Cenpu |
| Snhg8 |
| Mbp |
| Rb1 |
| Mrpl44 |
| Coq3 |
| Bcas3 |
| Pdss1 |
| Msx1 |
| Mrpl1 |
| Tmem138 |
| Mtg1 |
| Sh3bgrl2 |
| Wwox |
| Zcchc4 |
| Pcsk5 |
| Mmaa |
| Tbc1d7 |
| Mrps28 |
| Agk |
| Bcl6 |
| Creb5 |
| Clcn2 |
| Patj |
| Enpp4 |
| Rmi1 |
| Atxn7l2 |
| Pcbp3 |
| Ubxn2b |
| 2610005L07Rik |
| Pappa2 |
| 5730480H06Rik |
| Nme6 |
| Rsad1 |
| Akap7 |
| Serpinb9 |
| Pkn3 |
| Ccdc126 |
| Taco1 |
| Gng8 |

| |
|---|
| Borcs7 |
| Rdh14 |
| Bmpr1b |
| 5031425E22Rik |
| Fdxr |
| Gbe1 |
| Agbl5 |
| Pdlim4 |
| Mterf4 |
| Cdca7l |
| B3gntl1 |
| Zfp869 |
| Nt5dc1 |
| Rad52 |
| Fra10ac1 |
| Nphp3 |
| Parp11 |
| Ndufaf3 |
| Gtpbp8 |
| Tigd2 |
| Rccd1 |
| Atpaf1 |
| Pknox2 |
| Epb41l4aos |
| Gins3 |
| Ndufaf1 |
| Dleu2 |
| Tmem69 |
| Plxnc1 |
| Abtb1 |
| Mrm1 |
| Lamc2 |
| Npas4 |
| Sertad3 |
| Serinc2 |
| Kbtbd7 |
| Tfeb |
| Tefm |
| Jund |
| 0610009B22Rik |
| Dnajc24 |
| Rps6ka5 |
| Saysd1 |
| Setd4 |
| Dtwd1 |
| Plcb1 |
| Amn1 |
| Crem |
| Zfp341 |
| Slc35a3 |

| |
|---|
| Rsph3b |
| Gt(ROSA)26Sor |
| Mtrf1l |
| Cmtr1 |
| Mettl5 |
| Tmem44 |
| Wdr25 |
| Irgm1 |
| Zfp11 |
| Dffb |
| Bloc1s1 |
| Rfesd |
| Tmem107 |
| Rmdn1 |
| Arid3b |
| Tatdn3 |
| Tram1l1 |
| Slc1a3 |
| Vamp1 |
| Ccdc57 |
| Zfp560 |
| Mkx |
| Them4 |
| Acyp2 |
| Hspb7 |
| Rspo2 |
| Triqk |
| Pde1a |
| 9430038I01Rik |
| Zfp825 |
| Mrnip |
| Trp53rkb |
| Gin1 |
| Zfp446 |
| Rbm20 |
| Atg10 |
| Angptl6 |
| Pemt |
| Stag3 |
| Ttc26 |
| Zfp959 |
| Col28a1 |
| Slc25a29 |
| Napepld |
| Artn |
| Kcnab3 |
| Itga2b |
| 4930556M19Rik |
| Tmem204 |
| Zc3h6 |

| |
|---|
| Btbd8 |
| Zfp184 |
| 9330111N05Rik |
| Mettl18 |
| Snord104 |
| Pex12 |
| Fgfbp3 |
| Mat2a |
| Unc13c |
| Ccdc81 |
| Bmf |
| Lrrtm3 |
| Col19a1 |
| Fbxo36 |
| Exd1 |
| Kcnip1 |
| Ggct |
| Nckap5 |
| Gpr155 |
| Atn1 |
| Catsper2 |
| Esr1 |
| Zfp963 |
| Cyp2r1 |
| Lyl1 |
| Wdr31 |
| Gm12669 |
| Rpl36al |
| 1700007L15Rik |
| Isy1 |
| Ankrd37 |
| Dlgap1 |
| Pbld2 |
| Pih1d2 |
| Abca4 |
| 8430436N08Rik |
| C030034I22Rik |
| Adamts18 |
| Usp39 |
| Fbxl12os |
| 1700012D14Rik |
| Clic5 |
| Pax8 |
| 4930599N23Rik |
| Gm20125 |
| Itgb6 |
| Kcnb1 |
| Gm5069 |
| 4930539J05Rik |
| Npm2 |

| |
|---|
| Gm20139 |
| Enpp6 |
| Bank1 |
| Slc14a2 |
| 4930412B13Rik |
| 6430562O15Rik |
| Ppp1r3c |
| 4933430I17Rik |
| Nek10 |
| Cbfa2t3 |
| Nme1 |
| Hes3 |
| Gm29683 |
| 1700028E10Rik |
| 4632428C04Rik |
| Tuba1a |
| Gm11149 |
| Aknad1 |
| Etohd2 |
| Sys1 |
| Rfx8 |
| Mfsd13b |
| B230208H11Rik |
| 4930586N03Rik |
| Gnb2 |
| Tjp3 |
| 1810041H14Rik |
| Olfm3 |
| A430078I02Rik |
| Ctnna3 |
| 1700108F19Rik |
| Mir684-1 |
| Mir199a-1 |
| Rax |
| Pwwp2b |
| 4930526H09Rik |
| Sgpp2 |
| Mir8105 |
| 2010109A12Rik |
| Tmem42 |
| Gm13497 |
| Gm12596 |
| 4930543E12Rik |
| 1700125H03Rik |
| Gabrp |
| Tmprss6 |
| Ndufs3 |
| Pex11b |
| 6430531B16Rik |
| Cplx3 |

| | |
|---|---|
| | Arhgap15os |
| | Itgb3bp |
