## Supplementary Table S3 for "ERCC1-XPF Interacts with Topoisomerase IIβ to Facilitate the Repair of Activity-induced DNA Breaks"

### Supplementary Table S3. A list of 695 bXPF-bound proteins identified in the three high-throughput MS runs.

| | | peptide count | | | | | | probability | | | | | |
|---|---|---|---|---|---|---|---|---|---|---|---|---|---|
| Uniprot ID | Uniprot name | bXpf1 | bXpf2 | bXpf3 | birA1 | birA2 | birA3 | bXPF1 | bXPF2 | bXPF3 | birA1 | birA2 | birA3 |
| Q9QZQ1 | AFAD_MOUSE | 1 | 1 | 2 | 1 | 0 | 0 | 0.99 | 0.8 | 1 | 1 | 0.72 | 0 |
| Q9Z204 | HNRPC_MOUSE | 8 | 7 | 6 | 4 | 1 | 0 | 1 | 1 | 1 | 1 | 1 | 0 |
| E9QN37 | E9QN37_MOUSE | 1 | 3 | 3 | 1 | 1 | 0 | 0.83 | 1 | 1 | 0.89 | 0.91 | 0 |
| Q9R087 | GPC6_MOUSE | 2 | 2 | 6 | 0 | 0 | 0 | 1 | 1 | 1 | 0.49 | 0.88 | 0 |
| Q6A068 | CDC5L_MOUSE | 3 | 1 | 1 | 1 | 0 | 0 | 1 | 1 | 0.95 | 0.53 | 0.4 | 0.29 |
| P01831 | THY1_MOUSE | 6 | 6 | 5 | 1 | 2 | 4 | 1 | 1 | 1 | 1 | 1 | 1 |
| Q9CY27 | TECR_MOUSE | 5 | 5 | 7 | 1 | 2 | 2 | 1 | 1 | 1 | 1 | 0.98 | 1 |
| Q8BH97 | RCN3_MOUSE | 4 | 1 | 3 | 4 | 0 | 2 | 1 | 0.8 | 1 | 1 | 0 | 1 |
| Q9EPU4 | CPSF1_MOUSE | 7 | 4 | 6 | 2 | 1 | 1 | 1 | 1 | 1 | 1 | 0.95 | 0.84 |
| P63087 | PP1G_MOUSE | 3 | 4 | 2 | 1 | 1 | 1 | 1 | 1 | 1 | 1 | 1 | 0.99 |
| P21981 | TGM2_MOUSE | 5 | 9 | 3 | 3 | 1 | 0 | 1 | 1 | 1 | 1 | 0.99 | 0 |
| Q91YU6 | LZTS2_MOUSE | 1 | 5 | 1 | 0 | 1 | 0 | 0.92 | 1 | 1 | 1 | 0.91 | 0 |
| Q91VY9 | ZN622_MOUSE | 4 | 2 | 2 | 1 | 0 | 1 | 1 | 1 | 1 | 1 | 0.92 | 0.87 |
| Q8R323 | RFC3_MOUSE | 5 | 3 | 6 | 0 | 1 | 1 | 1 | 1 | 1 | 1 | 1 | 0.9 |
| Q8CFC7 | CLASR_MOUSE | 4 | 4 | 1 | 0 | 1 | 1 | 1 | 1 | 0.95 | 0.97 | 0.95 | 0.75 |
| O88967 | YMEL1_MOUSE | 3 | 4 | 3 | 1 | 0 | 0 | 1 | 1 | 1 | 1 | 0.95 | 0 |
| Q3TUH1 | TAM41_MOUSE | 0 | 3 | 2 | 1 | 1 | 0 | 0.94 | 1 | 1 | 0.99 | 1 | 0 |
| G5E896 | G5E896_MOUSE | 3 | 0 | 3 | 1 | 0 | 0 | 1 | 0.97 | 1 | 0.99 | 1 | 0 |
| Q60932 | VDAC1_MOUSE | 3 | 4 | 5 | 1 | 2 | 0 | 1 | 1 | 1 | 1 | 1 | 0 |
| G5E8R1 | G5E8R1_MOUSE | 4 | 0 | 0 | 4 | 0 | 0 | 1 | 1 | 1 | 1 | 1 | 0.82 |
| Q922Q8 | LRC59_MOUSE | 5 | 10 | 6 | 4 | 2 | 1 | 1 | 1 | 1 | 1 | 1 | 1 |
| Q64012 | RALY_MOUSE | 6 | 1 | 9 | 1 | 1 | 0 | 1 | 1 | 1 | 1 | 0.97 | 0 |
| Q8R2U0 | SEH1_MOUSE | 2 | 1 | 1 | 1 | 0 | 0 | 1 | 0.99 | 0.93 | 0.97 | 0.39 | 0 |
| Q8BVA4 | LMOD1_MOUSE | 11 | 16 | 3 | 4 | 2 | 1 | 1 | 1 | 1 | 1 | 1 | 0.87 |
| Q9DCT2 | NDUS3_MOUSE | 3 | 2 | 2 | 0 | 0 | 1 | 1 | 1 | 1 | 0 | 0.96 | 0.98 |
| P62137 | PP1A_MOUSE | 2 | 3 | 5 | 1 | 3 | 1 | 1 | 1 | 1 | 1 | 1 | 0.97 |
| Q3V1V3 | ESF1_MOUSE | 6 | 1 | 2 | 1 | 0 | 0 | 1 | 1 | 1 | 1 | 0 | 0 |
| P62996 | TRA2B_MOUSE | 5 | 3 | 7 | 3 | 1 | 0 | 1 | 1 | 1 | 1 | 1 | 0.3 |
| Q9D0I8 | MRT4_MOUSE | 2 | 3 | 4 | 0 | 1 | 0 | 1 | 1 | 1 | 0.98 | 0.98 | 0 |
| E9PUK6 | E9PUK6_MOUSE | 38 | 0 | 0 | 18 | 0 | 2 | 1 | 1 | 1 | 1 | 1 | 1 |
| P09103 | PDIA1_MOUSE | 8 | 13 | 7 | 5 | 5 | 3 | 1 | 1 | 1 | 1 | 1 | 1 |
| Q99LH1 | NOG2_MOUSE | 12 | 3 | 11 | 2 | 0 | 1 | 1 | 1 | 1 | 1 | 0.17 | 1 |
| Q9CQ19 | MYL9_MOUSE | 2 | 3 | 2 | 1 | 1 | 1 | 1 | 1 | 1 | 1 | 1 | 0.99 |
| O54931 | AKAP2_MOUSE | 3 | 3 | 1 | 1 | 0 | 0 | 1 | 1 | 0.95 | 0.94 | 0.75 | 0 |
| Q2KN98 | CYTSA_MOUSE | 15 | 19 | 17 | 7 | 4 | 6 | 1 | 1 | 1 | 1 | 1 | 1 |
| A1BN54 | A1BN54_MOUSE | 12 | 22 | 20 | 2 | 1 | 5 | 1 | 1 | 1 | 1 | 0.96 | 1 |
| G3X9J0 | G3X9J0_MOUSE | 6 | 6 | 0 | 1 | 1 | 0 | 1 | 1 | 1 | 1 | 0.95 | 0 |
| Q9Z0X1 | AIFM1_MOUSE | 2 | 5 | 8 | 2 | 1 | 0 | 1 | 1 | 1 | 1 | 1 | 0 |
| P62141 | PP1B_MOUSE | 12 | 12 | 14 | 7 | 6 | 2 | 1 | 1 | 1 | 1 | 1 | 1 |
| O35682 | MYADM_MOUSE | 1 | 3 | 2 | 1 | 1 | 1 | 1 | 1 | 1 | 1 | 1 | 0.99 |
| Q9D2V8 | MFS10_MOUSE | 0 | 1 | 4 | 1 | 0 | 0 | 0.96 | 1 | 1 | 1 | 0 | 0 |
| Q60930 | VDAC2_MOUSE | 5 | 5 | 7 | 1 | 0 | 2 | 1 | 1 | 1 | 1 | 0.98 | 1 |
| Q61712 | DNJC1_MOUSE | 1 | 4 | 2 | 0 | 1 | 0 | 0.99 | 1 | 0.95 | 0.23 | 0.73 | 0 |
| O88874 | CCNK_MOUSE | 3 | 2 | 3 | 0 | 1 | 0 | 1 | 1 | 1 | 0 | 1 | 0 |
| Q80UM7 | MOGS_MOUSE | 1 | 1 | 1 | 0 | 0 | 1 | 0.99 | 1 | 0.91 | 0.33 | 0.25 | 0.99 |
| Q9R099 | TBL2_MOUSE | 5 | 2 | 3 | 2 | 1 | 0 | 1 | 1 | 1 | 1 | 0.98 | 0 |
| Q9JMB0 | GKAP1_MOUSE | 5 | 2 | 1 | 1 | 0 | 0 | 1 | 1 | 0.98 | 1 | 0.85 | 0 |
| P97855 | G3BP1_MOUSE | 4 | 2 | 3 | 1 | 0 | 0 | 1 | 1 | 1 | 0.98 | 0.23 | 0 |
| Q01853 | TERA_MOUSE | 11 | 14 | 27 | 2 | 0 | 7 | 1 | 1 | 1 | 1 | 1 | 1 |
| Q69ZA1 | CDK13_MOUSE | 8 | 2 | 2 | 1 | 1 | 0 | 1 | 0.88 | 0.97 | 1 | 0.99 | 0 |
| Q61136 | PRP4B_MOUSE | 15 | 19 | 25 | 8 | 4 | 2 | 1 | 1 | 1 | 1 | 1 | 0.58 |
| Q8BG81 | PDIP3_MOUSE | 8 | 6 | 3 | 3 | 1 | 0 | 1 | 1 | 1 | 1 | 1 | 0 |
| Q8VDJ3 | VIGLN_MOUSE | 9 | 12 | 8 | 3 | 0 | 2 | 1 | 1 | 1 | 1 | 1 | 1 |
| Q80UZ2 | SDA1_MOUSE | 5 | 2 | 4 | 1 | 0 | 1 | 1 | 1 | 1 | 1 | 0.52 | 0.97 |
| Q9JJF3 | NO66_MOUSE | 6 | 7 | 7 | 5 | 1 | 2 | 1 | 1 | 1 | 1 | 1 | 0.77 |
| P57780 | ACTN4_MOUSE | 15 | 35 | 26 | 5 | 2 | 13 | 1 | 1 | 1 | 1 | 1 | 1 |
| Q9CY57 | CHTOP_MOUSE | 2 | 1 | 1 | 1 | 0 | 0 | 1 | 0.99 | 0.98 | 1 | 0.53 | 0 |
| Q3THE2 | ML12B_MOUSE | 9 | 9 | 7 | 9 | 4 | 5 | 1 | 1 | 1 | 1 | 1 | 1 |

| | | | | | | | | | | | | | |
|---|---|---|---|---|---|---|---|---|---|---|---|---|---|
| Q9DBE9 | SPB1_MOUSE | 5 | 5 | 9 | 3 | 1 | 1 | 1 | 1 | 1 | 1 | 0.96 | 0.99 |
| Q91YE7 | RBM5_MOUSE | 4 | 1 | 3 | 1 | 0 | 1 | 1 | 1 | 1 | 0.98 | 0 | 0.47 |
| Q03265 | ATPA_MOUSE | 11 | 25 | 6 | 9 | 14 | 2 | 1 | 1 | 1 | 1 | 1 | 1 |
| Q5SUA5 | MYO1G_MOUSE | 2 | 4 | 14 | 1 | 0 | 3 | 1 | 1 | 1 | 0.98 | 0 | 1 |
| Q9WVB4 | SLIT3_MOUSE | 8 | 22 | 20 | 2 | 8 | 1 | 1 | 1 | 1 | 1 | 1 | 0.98 |
| Q8BH59 | CMC1_MOUSE | 11 | 10 | 9 | 4 | 6 | 2 | 1 | 1 | 1 | 1 | 1 | 1 |
| Q8BFZ9 | ERLN2_MOUSE | 2 | 1 | 10 | 1 | 0 | 3 | 1 | 1 | 1 | 0.99 | 0 | 1 |
| Q99M87 | DNJA3_MOUSE | 4 | 3 | 2 | 1 | 1 | 0 | 1 | 1 | 1 | 1 | 0.98 | 0.21 |
| Q8BP67 | RL24_MOUSE | 29 | 13 | 33 | 20 | 16 | 23 | 1 | 1 | 1 | 1 | 1 | 1 |
| Q0VBL3 | Q0VBL3_MOUSE | 4 | 0 | 0 | 0 | 0 | 1 | 1 | 1 | 0.98 | 1 | 0.98 | 0.37 |
| Q8CAQ8 | IMMT_MOUSE | 7 | 14 | 8 | 3 | 2 | 0 | 1 | 1 | 1 | 1 | 1 | 0.16 |
| Q3UIX4 | Q3UIX4_MOUSE | 4 | 3 | 2 | 1 | 0 | 1 | 1 | 1 | 1 | 0.97 | 0 | 0.79 |
| Q8R1B4 | EIF3C_MOUSE | 4 | 10 | 4 | 2 | 2 | 0 | 1 | 1 | 1 | 1 | 1 | 0.96 |
| P41105 | RL28_MOUSE | 2 | 12 | 1 | 2 | 11 | 9 | 1 | 1 | 0.99 | 1 | 1 | 1 |
| E9PZ16 | E9PZ16_MOUSE | 44 | 62 | 0 | 24 | 12 | 11 | 1 | 1 | 1 | 1 | 1 | 1 |
| Q9DBS1 | TMM43_MOUSE | 2 | 5 | 10 | 2 | 2 | 3 | 1 | 1 | 1 | 1 | 1 | 1 |
| Q6PHQ9 | Q6PHQ9_MOUSE | 8 | 0 | 3 | 3 | 0 | 0 | 1 | 1 | 1 | 1 | 1 | 0.85 |
| P23116 | EIF3A_MOUSE | 8 | 8 | 4 | 6 | 2 | 1 | 1 | 1 | 1 | 1 | 1 | 0.48 |
| O55135 | IF6_MOUSE | 4 | 4 | 4 | 2 | 2 | 3 | 1 | 1 | 1 | 1 | 1 | 1 |
| Q9DC69 | NDUA9_MOUSE | 4 | 5 | 5 | 3 | 1 | 1 | 1 | 1 | 1 | 1 | 1 | 0.99 |
| Q9CPP0 | NPM3_MOUSE | 1 | 3 | 3 | 2 | 1 | 1 | 1 | 1 | 1 | 1 | 0.99 | 0.99 |
| Q9CZ52 | ANTR1_MOUSE | 0 | 0 | 3 | 1 | 1 | 0 | 0.95 | 0.94 | 1 | 1 | 0.75 | 0 |
| Q9CY50 | SSRA_MOUSE | 3 | 2 | 3 | 1 | 0 | 0 | 1 | 1 | 1 | 1 | 0.31 | 0.55 |
| Q6PFR5 | TRA2A_MOUSE | 6 | 6 | 6 | 2 | 2 | 1 | 1 | 1 | 1 | 1 | 1 | 0.82 |
| Q8BGJ5 | Q8BGJ5_MOUSE | 3 | 2 | 0 | 1 | 1 | 0 | 1 | 1 | 1 | 1 | 0.81 | 0 |
| Q9R0L6 | PCM1_MOUSE | 3 | 0 | 5 | 1 | 0 | 2 | 1 | 0.98 | 1 | 0.95 | 0 | 1 |
| F8VQ70 | F8VQ70_MOUSE | 5 | 1 | 3 | 2 | 0 | 0 | 1 | 0.84 | 1 | 1 | 1 | 0 |
| P35550 | FBRL_MOUSE | 10 | 9 | 17 | 7 | 6 | 6 | 1 | 1 | 1 | 1 | 1 | 1 |
| P58771 | TPM1_MOUSE | 35 | 28 | 21 | 20 | 12 | 9 | 1 | 1 | 1 | 1 | 1 | 1 |
| Q8R4U7 | LUZP1_MOUSE | 17 | 17 | 55 | 8 | 2 | 16 | 1 | 1 | 1 | 1 | 1 | 1 |
| Q91VD9 | NDUS1_MOUSE | 7 | 8 | 4 | 2 | 1 | 0 | 1 | 1 | 1 | 1 | 1 | 0.39 |
| Q8K4Q8 | COL12_MOUSE | 2 | 2 | 5 | 1 | 0 | 0 | 1 | 1 | 1 | 0.38 | 0 | 0 |
| Q99J62 | RFC4_MOUSE | 7 | 9 | 12 | 4 | 6 | 2 | 1 | 1 | 1 | 1 | 1 | 1 |
| Q9D8X5 | CNOT8_MOUSE | 2 | 1 | 1 | 1 | 0 | 0 | 1 | 1 | 0.97 | 1 | 0.97 | 0 |
| Q6NWW9 | FND3B_MOUSE | 12 | 16 | 15 | 5 | 3 | 5 | 1 | 1 | 1 | 1 | 1 | 1 |
| P51655 | GPC4_MOUSE | 3 | 14 | 13 | 3 | 3 | 4 | 1 | 1 | 1 | 1 | 1 | 1 |
| Q91VR2 | ATPG_MOUSE | 0 | 6 | 2 | 0 | 2 | 0 | 0.99 | 1 | 1 | 0.98 | 1 | 0 |
| D3Z5I1 | D3Z5I1_MOUSE | 4 | 4 | 2 | 3 | 1 | 0 | 1 | 1 | 1 | 1 | 1 | 0.05 |
| F8VQC7 | F8VQC7_MOUSE | 13 | 19 | 0 | 5 | 0 | 3 | 1 | 1 | 1 | 1 | 0.28 | 1 |
| O08992 | SDCB1_MOUSE | 2 | 5 | 1 | 0 | 1 | 0 | 1 | 1 | 1 | 0.94 | 1 | 0 |
| Q9R112 | SQRD_MOUSE | 1 | 3 | 1 | 3 | 1 | 0 | 1 | 1 | 1 | 1 | 1 | 0 |
| Q920B9 | SP16H_MOUSE | 1 | 7 | 19 | 2 | 2 | 7 | 1 | 1 | 1 | 1 | 1 | 1 |
| Q6P9Q6 | FKB15_MOUSE | 2 | 4 | 3 | 1 | 1 | 1 | 1 | 1 | 1 | 1 | 1 | 0.95 |
| P58774 | TPM2_MOUSE | 7 | 6 | 2 | 3 | 2 | 1 | 1 | 1 | 1 | 1 | 1 | 0.97 |
| D3YYT1 | D3YYT1_MOUSE | 8 | 0 | 3 | 1 | 0 | 0 | 1 | 1 | 1 | 1 | 1 | 0 |
| Q9JIK5 | DDX21_MOUSE | 15 | 24 | 10 | 9 | 0 | 7 | 1 | 1 | 1 | 1 | 0.98 | 1 |
| B9EJ54 | B9EJ54_MOUSE | 10 | 0 | 2 | 1 | 0 | 0 | 1 | 1 | 1 | 1 | 0.15 | 0 |
| Q9EP71 | RAI14_MOUSE | 36 | 38 | 54 | 22 | 4 | 18 | 1 | 1 | 1 | 1 | 1 | 1 |
| Q78PY7 | SND1_MOUSE | 5 | 8 | 9 | 4 | 0 | 2 | 1 | 1 | 1 | 1 | 0 | 1 |
| Q8K297 | GT251_MOUSE | 6 | 8 | 10 | 0 | 2 | 3 | 1 | 1 | 1 | 0.81 | 1 | 1 |
| F8VPM7 | F8VPM7_MOUSE | 2 | 4 | 6 | 1 | 0 | 0 | 1 | 1 | 1 | 1 | 0 | 0 |
| Q8VHZ7 | IMP4_MOUSE | 5 | 6 | 3 | 2 | 1 | 0 | 1 | 1 | 1 | 1 | 0.88 | 0 |
| O88532 | ZFR_MOUSE | 17 | 12 | 29 | 8 | 1 | 10 | 1 | 1 | 1 | 1 | 1 | 1 |
| Q68FD5 | CLH1_MOUSE | 12 | 13 | 27 | 4 | 5 | 7 | 1 | 1 | 1 | 1 | 1 | 1 |
| O35286 | DHX15_MOUSE | 13 | 10 | 10 | 8 | 0 | 5 | 1 | 1 | 1 | 1 | 0.96 | 1 |
| Q9CVB6 | ARPC2_MOUSE | 16 | 16 | 21 | 15 | 10 | 10 | 1 | 1 | 1 | 1 | 1 | 1 |
| Q04857 | CO6A1_MOUSE | 6 | 4 | 2 | 2 | 0 | 1 | 1 | 1 | 1 | 1 | 0 | 0.75 |
| G3XA10 | G3XA10_MOUSE | 3 | 13 | 11 | 4 | 1 | 6 | 1 | 1 | 1 | 1 | 1 | 1 |
| Q8K019 | BCLF1_MOUSE | 25 | 23 | 36 | 10 | 3 | 10 | 1 | 1 | 1 | 1 | 1 | 1 |
| Q569Z6 | TR150_MOUSE | 38 | 33 | 48 | 23 | 2 | 26 | 1 | 1 | 1 | 1 | 1 | 1 |
| P84099 | RL19_MOUSE | 28 | 19 | 28 | 25 | 14 | 15 | 1 | 1 | 1 | 1 | 1 | 1 |
| P26231 | CTNA1_MOUSE | 4 | 10 | 8 | 2 | 0 | 3 | 1 | 1 | 1 | 1 | 0.36 | 1 |
| Q8BSY0 | ASPH_MOUSE | 11 | 19 | 25 | 4 | 0 | 9 | 1 | 1 | 1 | 1 | 0 | 1 |

| | | | | | | | | | | | | | |
|---|---|---|---|---|---|---|---|---|---|---|---|---|---|
| Q9CRB9 | CHCH3_MOUSE | 3 | 2 | 6 | 2 | 0 | 1 | 1 | 1 | 1 | 1 | 0.45 | 0.98 |
| Q8BMK4 | CKAP4_MOUSE | 37 | 31 | 42 | 20 | 7 | 16 | 1 | 1 | 1 | 1 | 1 | 1 |
| Q91Z92 | B3GT6_MOUSE | 3 | 2 | 2 | 1 | 0 | 0 | 1 | 1 | 1 | 1 | 0.96 | 0 |
| Q91VR5 | DDX1_MOUSE | 31 | 22 | 30 | 20 | 13 | 20 | 1 | 1 | 1 | 1 | 1 | 1 |
| Q5RJG1 | NOL10_MOUSE | 10 | 5 | 2 | 5 | 1 | 0 | 1 | 1 | 1 | 1 | 1 | 0 |
| Q99ME9 | NOG1_MOUSE | 11 | 12 | 21 | 10 | 4 | 5 | 1 | 1 | 1 | 1 | 1 | 1 |
| P56480 | ATPB_MOUSE | 8 | 23 | 6 | 7 | 5 | 0 | 1 | 1 | 1 | 1 | 1 | 0.31 |
| A2AJT5 | A2AJT5_MOUSE | 1 | 2 | 1 | 1 | 0 | 0 | 1 | 1 | 0.95 | 0.99 | 0 | 0 |
| Q9R0U0 | SRS10_MOUSE | 5 | 3 | 5 | 2 | 2 | 2 | 1 | 1 | 1 | 1 | 1 | 1 |
| Q7TPW1 | NEXN_MOUSE | 4 | 24 | 9 | 1 | 7 | 4 | 1 | 1 | 1 | 1 | 1 | 1 |
| Q7TNC4 | LC7L2_MOUSE | 15 | 11 | 11 | 13 | 5 | 3 | 1 | 1 | 1 | 1 | 1 | 1 |
| Q3U9G9 | LBR_MOUSE | 8 | 3 | 9 | 4 | 0 | 3 | 1 | 1 | 1 | 1 | 0.97 | 1 |
| Q9WUK4 | RFC2_MOUSE | 3 | 2 | 9 | 3 | 4 | 0 | 1 | 1 | 1 | 1 | 1 | 0 |
| B2RY56 | RBM25_MOUSE | 23 | 17 | 32 | 13 | 2 | 14 | 1 | 1 | 1 | 1 | 1 | 1 |
| Q8K1N2 | PHLB2_MOUSE | 19 | 14 | 25 | 8 | 2 | 8 | 1 | 1 | 1 | 1 | 1 | 1 |
| Q99KV1 | DJB11_MOUSE | 1 | 0 | 3 | 1 | 2 | 0 | 1 | 1 | 1 | 0.97 | 1 | 0 |
| P62874 | GBB1_MOUSE | 6 | 7 | 7 | 1 | 2 | 2 | 1 | 1 | 1 | 1 | 1 | 1 |
| P20152 | VIME_MOUSE | 53 | 37 | 39 | 27 | 27 | 27 | 1 | 1 | 1 | 1 | 1 | 1 |
| Q5SU73 | COIL_MOUSE | 1 | 1 | 1 | 1 | 1 | 0 | 0.9 | 1 | 0.99 | 0.59 | 0.98 | 0 |
| Q99LE6 | ABCF2_MOUSE | 10 | 10 | 21 | 5 | 3 | 7 | 1 | 1 | 1 | 1 | 1 | 1 |
| Q80VB6 | Q80VB6_MOUSE | 8 | 0 | 17 | 5 | 0 | 4 | 1 | 1 | 1 | 1 | 1 | 1 |
| O08579 | EMD_MOUSE | 9 | 7 | 8 | 4 | 5 | 2 | 1 | 1 | 1 | 1 | 1 | 1 |
| Q9JJZ6 | KLF13_MOUSE | 3 | 3 | 2 | 2 | 0 | 0 | 1 | 1 | 1 | 1 | 0 | 0 |
| Q9D883 | U2AF1_MOUSE | 7 | 6 | 10 | 2 | 6 | 1 | 1 | 1 | 1 | 1 | 1 | 1 |
| Q8K310 | MATR3_MOUSE | 20 | 17 | 31 | 14 | 1 | 9 | 1 | 1 | 1 | 1 | 0.99 | 1 |
| O08810 | U5S1_MOUSE | 21 | 22 | 25 | 6 | 0 | 7 | 1 | 1 | 1 | 1 | 0.61 | 1 |
| A3KGQ6 | A3KGQ6_MOUSE | 3 | 3 | 2 | 0 | 3 | 0 | 1 | 1 | 1 | 0 | 1 | 0 |
| Q9WV32 | ARC1B_MOUSE | 7 | 15 | 24 | 8 | 4 | 10 | 1 | 1 | 1 | 1 | 1 | 1 |
| G3X9V2 | G3X9V2_MOUSE | 21 | 20 | 16 | 6 | 3 | 7 | 1 | 1 | 1 | 1 | 1 | 1 |
| P55096 | ABCD3_MOUSE | 4 | 0 | 6 | 0 | 0 | 1 | 1 | 0.97 | 1 | 0.93 | 0.65 | 1 |
| Q99KG3 | RBM10_MOUSE | 8 | 11 | 16 | 5 | 0 | 2 | 1 | 1 | 1 | 1 | 0 | 1 |
| Q5SFM8 | RBM27_MOUSE | 2 | 7 | 4 | 2 | 1 | 1 | 1 | 1 | 1 | 1 | 0.99 | 0.61 |
| Q8VE97 | SRSF4_MOUSE | 7 | 0 | 2 | 2 | 2 | 0 | 1 | 0.94 | 1 | 1 | 0.96 | 0 |
| Q9DB96 | NGDN_MOUSE | 3 | 6 | 1 | 2 | 2 | 0 | 1 | 1 | 1 | 1 | 1 | 0 |
| P10126 | EF1A1_MOUSE | 10 | 10 | 7 | 4 | 4 | 3 | 1 | 1 | 1 | 1 | 1 | 1 |
| Q8K224 | NAT10_MOUSE | 9 | 16 | 13 | 4 | 0 | 7 | 1 | 1 | 1 | 1 | 0.58 | 1 |
| P09405 | NUCL_MOUSE | 9 | 16 | 10 | 5 | 1 | 3 | 1 | 1 | 1 | 1 | 1 | 1 |
| Q99KW3 | TARA_MOUSE | 7 | 11 | 3 | 4 | 3 | 0 | 1 | 1 | 1 | 1 | 1 | 0 |
| P46061 | RAGP1_MOUSE | 10 | 6 | 5 | 4 | 2 | 1 | 1 | 1 | 1 | 1 | 1 | 0.82 |
| Q91VE6 | MK67I_MOUSE | 2 | 2 | 2 | 0 | 1 | 1 | 1 | 1 | 1 | 0.95 | 0.99 | 0.95 |
| Q9CXW4 | RL11_MOUSE | 16 | 16 | 16 | 7 | 11 | 10 | 1 | 1 | 1 | 1 | 1 | 1 |
| P14115 | RL27A_MOUSE | 0 | 8 | 3 | 0 | 6 | 2 | 0.97 | 1 | 1 | 0.98 | 1 | 1 |
| Q9D8N0 | EF1G_MOUSE | 5 | 7 | 4 | 2 | 2 | 0 | 1 | 1 | 1 | 1 | 1 | 0 |
| Q8VHE0 | SEC63_MOUSE | 3 | 7 | 5 | 2 | 0 | 3 | 1 | 1 | 1 | 1 | 0.24 | 1 |
| Q8BG67 | EFR3A_MOUSE | 6 | 5 | 7 | 0 | 3 | 2 | 1 | 1 | 1 | 0.95 | 1 | 1 |
| Q9DBZ9 | SYDE1_MOUSE | 3 | 1 | 2 | 1 | 0 | 0 | 1 | 1 | 1 | 0.86 | 0.36 | 0 |
| P62880 | GBB2_MOUSE | 5 | 11 | 12 | 6 | 4 | 4 | 1 | 1 | 1 | 1 | 1 | 1 |
| P55937 | GOGA3_MOUSE | 14 | 18 | 41 | 7 | 4 | 11 | 1 | 1 | 1 | 1 | 1 | 1 |
| Q8JZQ9 | EIF3B_MOUSE | 5 | 9 | 4 | 3 | 0 | 2 | 1 | 1 | 1 | 1 | 0.63 | 1 |
| P58252 | EF2_MOUSE | 0 | 10 | 6 | 0 | 0 | 5 | 0.88 | 1 | 1 | 0 | 0.92 | 1 |
| Q9ESV0 | DDX24_MOUSE | 3 | 5 | 2 | 2 | 0 | 2 | 1 | 1 | 1 | 1 | 0 | 1 |
| Q9JJT0 | RCL1_MOUSE | 3 | 2 | 5 | 1 | 2 | 0 | 1 | 1 | 1 | 0.99 | 1 | 0 |
| Q6ZQ58 | J3QNB1_MOUSE | 10 | 9 | 12 | 6 | 6 | 3 | 1 | 1 | 1 | 1 | 1 | 1 |
| Q6ZWN5 | RS9_MOUSE | 28 | 18 | 4 | 16 | 13 | 13 | 1 | 1 | 1 | 1 | 1 | 1 |
| P20029 | GRP78_MOUSE | 22 | 30 | 32 | 12 | 7 | 9 | 1 | 1 | 1 | 1 | 1 | 1 |
| Q61398 | PCOC1_MOUSE | 7 | 4 | 9 | 2 | 2 | 3 | 1 | 1 | 1 | 1 | 1 | 1 |
| Q3URQ0 | TEX10_MOUSE | 11 | 10 | 17 | 4 | 2 | 6 | 1 | 1 | 1 | 1 | 1 | 1 |
| Q8BFZ3 | ACTBL_MOUSE | 4 | 2 | 2 | 2 | 1 | 2 | 1 | 1 | 1 | 1 | 0.9 | 1 |
| P14869 | RLA0_MOUSE | 29 | 27 | 31 | 22 | 15 | 10 | 1 | 1 | 1 | 1 | 1 | 1 |
| Q69ZR9 | F208A_MOUSE | 0 | 1 | 3 | 1 | 0 | 0 | 1 | 0.99 | 1 | 1 | 0 | 0 |
| Q6A0A9 | F120A_MOUSE | 19 | 16 | 36 | 14 | 4 | 12 | 1 | 1 | 1 | 1 | 1 | 1 |
| Q08093 | CNN2_MOUSE | 3 | 5 | 2 | 4 | 1 | 0 | 1 | 1 | 1 | 1 | 1 | 0 |
| Q9WV92 | E41L3_MOUSE | 2 | 3 | 4 | 0 | 0 | 1 | 1 | 1 | 1 | 0.25 | 0 | 0.9 |

| | | | | | | | | | | | | | |
|---|---|---|---|---|---|---|---|---|---|---|---|---|---|
| Q3USZ8 | DIA1_MOUSE | 1 | 1 | 4 | 2 | 3 | 0 | 0.97 | 0.82 | 1 | 1 | 1 | 0.7 |
| E9PZ21 | E9PZ21_MOUSE | 7 | 0 | 2 | 2 | 0 | 0 | 1 | 0.9 | 1 | 1 | 0 | 0 |
| P13020 | GELS_MOUSE | 41 | 40 | 47 | 24 | 24 | 22 | 1 | 1 | 1 | 1 | 1 | 1 |
| Q80YS6 | AFAP1_MOUSE | 9 | 26 | 18 | 8 | 3 | 11 | 1 | 1 | 1 | 1 | 1 | 1 |
| B2RRE7 | OTUD4_MOUSE | 8 | 2 | 2 | 3 | 0 | 1 | 1 | 0.94 | 1 | 1 | 0 | 0.95 |
| O70305 | ATX2_MOUSE | 6 | 3 | 7 | 2 | 1 | 5 | 1 | 1 | 1 | 1 | 0.99 | 1 |
| Q8VIJ6 | SFPQ_MOUSE | 3 | 10 | 2 | 2 | 2 | 2 | 1 | 1 | 1 | 0.99 | 0.86 | 1 |
| Q80WJ7 | LYRIC_MOUSE | 8 | 9 | 10 | 6 | 3 | 4 | 1 | 1 | 1 | 1 | 1 | 1 |
| Q99J47 | DRS7B_MOUSE | 7 | 6 | 5 | 2 | 3 | 0 | 1 | 1 | 1 | 1 | 1 | 0 |
| Q8BFV2 | PCID2_MOUSE | 5 | 4 | 3 | 1 | 0 | 0 | 1 | 1 | 1 | 1 | 0.92 | 0 |
| Q8JZU2 | Q8JZU2_MOUSE | 8 | 6 | 5 | 7 | 0 | 1 | 1 | 1 | 1 | 1 | 1 | 1 |
| Q9EP97 | SENP3_MOUSE | 6 | 1 | 3 | 2 | 1 | 2 | 1 | 1 | 1 | 1 | 0.69 | 1 |
| G3UYG6 | G3UYG6_MOUSE | 31 | 14 | 13 | 20 | 2 | 7 | 1 | 1 | 1 | 1 | 1 | 1 |
| Q8BL97 | SRSF7_MOUSE | 12 | 10 | 11 | 10 | 6 | 5 | 1 | 1 | 1 | 1 | 1 | 1 |
| Q91VS8 | FARP2_MOUSE | 5 | 6 | 11 | 0 | 0 | 3 | 1 | 1 | 1 | 0.99 | 0.7 | 1 |
| Q9QX47 | SON_MOUSE | 5 | 2 | 1 | 1 | 0 | 1 | 1 | 1 | 1 | 1 | 0.42 | 0.95 |
| Q8K0V4 | CNOT3_MOUSE | 2 | 1 | 1 | 1 | 0 | 1 | 1 | 1 | 1 | 1 | 0 | 1 |
| F8VPU2 | FARP1_MOUSE | 11 | 16 | 31 | 5 | 5 | 11 | 1 | 1 | 1 | 1 | 1 | 1 |
| P61161 | ARP2_MOUSE | 23 | 22 | 36 | 21 | 9 | 20 | 1 | 1 | 1 | 1 | 1 | 1 |
| Q6P9R1 | DDX51_MOUSE | 7 | 11 | 6 | 7 | 6 | 1 | 1 | 1 | 1 | 1 | 1 | 0.8 |
| Q9D8S3 | ARFG3_MOUSE | 3 | 5 | 6 | 2 | 2 | 3 | 1 | 1 | 1 | 1 | 1 | 1 |
| Q9DBR7 | MYPT1_MOUSE | 23 | 29 | 30 | 10 | 6 | 14 | 1 | 1 | 1 | 1 | 1 | 1 |
| E9QA15 | E9QA15_MOUSE | 29 | 47 | 17 | 14 | 21 | 0 | 1 | 1 | 1 | 1 | 1 | 1 |
| Q8R422 | CD109_MOUSE | 7 | 7 | 20 | 7 | 6 | 2 | 1 | 1 | 1 | 1 | 1 | 1 |
| P29341 | PABP1_MOUSE | 22 | 19 | 22 | 14 | 9 | 8 | 1 | 1 | 1 | 1 | 1 | 1 |
| Q8BH74 | NU107_MOUSE | 3 | 1 | 2 | 1 | 0 | 1 | 1 | 0.95 | 1 | 0.55 | 0.87 | 0.72 |
| Q3UFM5 | NOM1_MOUSE | 7 | 4 | 1 | 4 | 1 | 0 | 1 | 1 | 1 | 1 | 0.84 | 0.24 |
| P97434 | MPRIP_MOUSE | 5 | 3 | 5 | 2 | 0 | 2 | 1 | 1 | 1 | 1 | 0.5 | 1 |
| Q91VN6 | DDX41_MOUSE | 9 | 4 | 8 | 6 | 4 | 2 | 1 | 1 | 1 | 1 | 1 | 1 |
| Q7TMK9 | HNRPQ_MOUSE | 3 | 6 | 4 | 3 | 2 | 0 | 1 | 1 | 1 | 1 | 1 | 0 |
| Q8CIE6 | COPA_MOUSE | 4 | 6 | 12 | 2 | 1 | 1 | 1 | 1 | 1 | 1 | 1 | 1 |
| Q99LB0 | TDIF1_MOUSE | 4 | 5 | 1 | 2 | 1 | 0 | 1 | 1 | 0.92 | 1 | 0.99 | 0 |
| P61255 | RL26_MOUSE | 27 | 17 | 31 | 20 | 13 | 21 | 1 | 1 | 1 | 1 | 1 | 1 |
| P62242 | RS8_MOUSE | 25 | 14 | 18 | 16 | 7 | 13 | 1 | 1 | 1 | 1 | 1 | 1 |
| E9QA16 | E9QA16_MOUSE | 2 | 0 | 2 | 1 | 2 | 0 | 1 | 1 | 1 | 1 | 1 | 0.95 |
| Q9CZU3 | SK2L2_MOUSE | 10 | 10 | 18 | 5 | 0 | 6 | 1 | 1 | 1 | 1 | 0.62 | 1 |
| P11276 | FINC_MOUSE | 66 | 85 | 56 | 35 | 25 | 29 | 1 | 1 | 1 | 1 | 1 | 1 |
| D3YXK2 | SAFB1_MOUSE | 2 | 3 | 4 | 1 | 0 | 0 | 1 | 1 | 1 | 1 | 0 | 0 |
| Q9CR62 | M2OM_MOUSE | 5 | 11 | 8 | 6 | 7 | 0 | 1 | 1 | 1 | 1 | 1 | 0 |
| J3QK52 | J3QK52_MOUSE | 1 | 0 | 2 | 1 | 1 | 1 | 0.85 | 1 | 1 | 0.96 | 0.48 | 1 |
| P07356 | ANXA2_MOUSE | 4 | 11 | 7 | 5 | 3 | 4 | 1 | 1 | 1 | 1 | 1 | 1 |
| P21107 | TPM3_MOUSE | 1 | 12 | 10 | 0 | 9 | 5 | 0.97 | 1 | 1 | 0 | 1 | 1 |
| Q91YK2 | RRP1B_MOUSE | 5 | 5 | 5 | 3 | 1 | 4 | 1 | 1 | 1 | 1 | 1 | 1 |
| Q8BMA6 | SRP68_MOUSE | 14 | 16 | 10 | 6 | 9 | 1 | 1 | 1 | 1 | 1 | 1 | 0.97 |
| Q3TJZ6 | FA98A_MOUSE | 13 | 6 | 9 | 6 | 4 | 5 | 1 | 1 | 1 | 1 | 1 | 1 |
| Q8BYK6 | YTHD3_MOUSE | 4 | 2 | 2 | 2 | 0 | 0 | 1 | 1 | 1 | 1 | 0.23 | 0 |
| Q80X90 | FLNB_MOUSE | 45 | 46 | 28 | 32 | 10 | 8 | 1 | 1 | 1 | 1 | 1 | 1 |
| P62717 | RL18A_MOUSE | 32 | 19 | 5 | 25 | 11 | 20 | 1 | 1 | 1 | 1 | 1 | 1 |
| Q6ZWZ7 | Q6ZWZ7_MOUSE | 30 | 10 | 3 | 20 | 0 | 0 | 1 | 1 | 1 | 1 | 1 | 1 |
| Q3TDQ1 | STT3B_MOUSE | 4 | 7 | 7 | 2 | 4 | 1 | 1 | 1 | 1 | 1 | 1 | 0.97 |
| P62754 | RS6_MOUSE | 31 | 29 | 43 | 35 | 24 | 23 | 1 | 1 | 1 | 1 | 1 | 1 |
| P70168 | IMB1_MOUSE | 4 | 5 | 2 | 1 | 2 | 1 | 1 | 1 | 1 | 1 | 1 | 0.77 |
| P25444 | RS2_MOUSE | 21 | 23 | 29 | 16 | 15 | 13 | 1 | 1 | 1 | 1 | 1 | 1 |
| P15864 | H12_MOUSE | 4 | 3 | 11 | 5 | 0 | 4 | 1 | 1 | 1 | 1 | 0.94 | 1 |
| Q80Y44 | DDX10_MOUSE | 2 | 3 | 6 | 0 | 0 | 2 | 1 | 1 | 1 | 0.94 | 0 | 1 |
| E9PWQ3 | E9PWQ3_MOUSE | 23 | 1 | 3 | 3 | 0 | 0 | 1 | 1 | 1 | 1 | 1 | 0.22 |
| E9QN52 | E9QN52_MOUSE | 1 | 0 | 1 | 1 | 0 | 0 | 0.87 | 0.91 | 0.92 | 0.77 | 0.18 | 0 |
| Q921U8 | SMTN_MOUSE | 7 | 5 | 17 | 6 | 1 | 4 | 1 | 1 | 1 | 1 | 0.78 | 1 |
| Q8K4L0 | DDX54_MOUSE | 14 | 16 | 20 | 7 | 0 | 6 | 1 | 1 | 1 | 1 | 0.71 | 1 |
| Q99JY9 | ARP3_MOUSE | 19 | 14 | 19 | 15 | 9 | 9 | 1 | 1 | 1 | 1 | 1 | 1 |
| Q9D0E1 | HNRPM_MOUSE | 50 | 39 | 32 | 35 | 24 | 14 | 1 | 1 | 1 | 1 | 1 | 1 |
| Q569Z5 | DDX46_MOUSE | 28 | 17 | 44 | 17 | 0 | 10 | 1 | 1 | 1 | 1 | 0.6 | 1 |
| Q60865 | CAPR1_MOUSE | 11 | 9 | 8 | 4 | 0 | 5 | 1 | 1 | 1 | 1 | 0 | 1 |

| | | | | | | | | | | | | | |
|---|---|---|---|---|---|---|---|---|---|---|---|---|---|
| H7BX95 | H7BX95_MOUSE | 14 | 16 | 10 | 13 | 7 | 0 | 1 | 1 | 1 | 1 | 1 | 0.98 |
| A2BE28 | LAS1L_MOUSE | 12 | 2 | 13 | 6 | 1 | 8 | 1 | 1 | 1 | 1 | 0.76 | 1 |
| Q99MJ9 | DDX50_MOUSE | 18 | 16 | 15 | 8 | 6 | 11 | 1 | 1 | 1 | 1 | 1 | 1 |
| A2AMY5 | A2AMY5_MOUSE | 9 | 9 | 12 | 6 | 1 | 5 | 1 | 1 | 1 | 1 | 1 | 1 |
| P32067 | LA_MOUSE | 4 | 1 | 3 | 3 | 0 | 0 | 1 | 1 | 1 | 1 | 0.55 | 0 |
| P43276 | H15_MOUSE | 7 | 10 | 19 | 12 | 8 | 6 | 1 | 1 | 1 | 1 | 1 | 1 |
| P97868 | RBBP6_MOUSE | 10 | 5 | 2 | 4 | 0 | 0 | 1 | 1 | 1 | 1 | 0.19 | 0 |
| Q8R4R6 | NUP53_MOUSE | 3 | 3 | 2 | 1 | 0 | 0 | 1 | 1 | 1 | 0.99 | 0.58 | 0 |
| P35979 | RL12_MOUSE | 14 | 15 | 16 | 11 | 12 | 9 | 1 | 1 | 1 | 1 | 1 | 1 |
| E9QLA5 | E9QLA5_MOUSE | 21 | 6 | 10 | 13 | 1 | 0 | 1 | 1 | 1 | 1 | 0.62 | 0 |
| Q6DVA0 | LEMD2_MOUSE | 1 | 1 | 2 | 1 | 0 | 0 | 1 | 1 | 1 | 0.99 | 0.47 | 0 |
| Q9QXS6 | DREB_MOUSE | 19 | 22 | 23 | 12 | 0 | 10 | 1 | 1 | 1 | 1 | 0 | 1 |
| Q9JKY0 | RCD1_MOUSE | 3 | 3 | 3 | 1 | 0 | 1 | 1 | 1 | 1 | 1 | 0.65 | 0.95 |
| P37889 | FBLN2_MOUSE | 2 | 7 | 2 | 2 | 0 | 2 | 1 | 1 | 1 | 1 | 0 | 1 |
| Q60605 | MYL6_MOUSE | 6 | 12 | 2 | 4 | 9 | 1 | 1 | 1 | 1 | 1 | 1 | 0.84 |
| Q8BTI8 | SRRM2_MOUSE | 52 | 27 | 38 | 31 | 8 | 23 | 1 | 1 | 1 | 1 | 1 | 1 |
| Q8VDM6 | HNRL1_MOUSE | 1 | 1 | 5 | 1 | 0 | 1 | 1 | 0.99 | 1 | 0.96 | 0.41 | 0.95 |
| P62281 | RS11_MOUSE | 10 | 12 | 31 | 4 | 13 | 17 | 1 | 1 | 1 | 1 | 1 | 1 |
| Q9CR57 | RL14_MOUSE | 28 | 19 | 39 | 25 | 21 | 27 | 1 | 1 | 1 | 1 | 1 | 1 |
| Q9D6Z1 | NOP56_MOUSE | 34 | 35 | 39 | 24 | 16 | 18 | 1 | 1 | 1 | 1 | 1 | 1 |
| Q8BZX4 | SREK1_MOUSE | 2 | 2 | 2 | 1 | 0 | 0 | 1 | 1 | 1 | 0.98 | 0.92 | 0 |
| O08532 | CA2D1_MOUSE | 5 | 6 | 13 | 2 | 3 | 4 | 1 | 1 | 1 | 1 | 1 | 1 |
| O09044 | SNP23_MOUSE | 2 | 3 | 2 | 3 | 0 | 1 | 1 | 1 | 1 | 1 | 0.85 | 0.95 |
| P47915 | RL29_MOUSE | 19 | 9 | 13 | 18 | 10 | 12 | 1 | 1 | 1 | 1 | 1 | 1 |
| Q91YQ5 | RPN1_MOUSE | 19 | 26 | 32 | 21 | 11 | 15 | 1 | 1 | 1 | 1 | 1 | 1 |
| B2RVL6 | ZCH24_MOUSE | 1 | 3 | 1 | 1 | 1 | 0 | 1 | 1 | 1 | 1 | 0.99 | 0 |
| P21278 | GNA11_MOUSE | 1 | 1 | 4 | 2 | 3 | 1 | 1 | 0.94 | 1 | 1 | 1 | 0.99 |
| Q9DCA5 | BRX1_MOUSE | 8 | 1 | 9 | 7 | 1 | 0 | 1 | 1 | 1 | 1 | 0.99 | 0 |
| Q6IRU2 | TPM4_MOUSE | 3 | 13 | 3 | 3 | 3 | 1 | 1 | 1 | 1 | 1 | 1 | 0.98 |
| Q811S7 | UBIP1_MOUSE | 2 | 2 | 2 | 1 | 0 | 1 | 1 | 1 | 1 | 1 | 0.91 | 1 |
| Q9WTI7 | MYO1C_MOUSE | 50 | 61 | 97 | 42 | 17 | 47 | 1 | 1 | 1 | 1 | 1 | 1 |
| D3YWX2 | D3YWX2_MOUSE | 19 | 0 | 0 | 14 | 0 | 0 | 1 | 1 | 1 | 1 | 1 | 1 |
| E9Q6L0 | E9Q6L0_MOUSE | 3 | 0 | 0 | 3 | 0 | 0 | 1 | 1 | 1 | 1 | 0.44 | 1 |
| P47955 | RLA1_MOUSE | 4 | 7 | 1 | 3 | 5 | 0 | 1 | 1 | 0.95 | 1 | 1 | 0 |
| P12970 | RL7A_MOUSE | 42 | 40 | 49 | 37 | 36 | 36 | 1 | 1 | 1 | 1 | 1 | 1 |
| P29268 | CTGF_MOUSE | 8 | 13 | 1 | 6 | 5 | 0 | 1 | 1 | 0.95 | 1 | 1 | 0 |
| Q6DFW4 | NOP58_MOUSE | 10 | 8 | 10 | 5 | 3 | 5 | 1 | 1 | 1 | 1 | 1 | 1 |
| P51410 | RL9_MOUSE | 18 | 20 | 5 | 13 | 16 | 10 | 1 | 1 | 1 | 1 | 1 | 1 |
| Q80X50 | UBP2L_MOUSE | 22 | 15 | 25 | 12 | 6 | 10 | 1 | 1 | 1 | 1 | 1 | 1 |
| P63017 | HSP7C_MOUSE | 19 | 22 | 24 | 14 | 13 | 11 | 1 | 1 | 1 | 1 | 1 | 1 |
| Q6ZWV3 | RL10_MOUSE | 24 | 19 | 12 | 20 | 15 | 21 | 1 | 1 | 1 | 1 | 1 | 1 |
| Q6PDH0 | PHLB1_MOUSE | 10 | 15 | 17 | 6 | 0 | 3 | 1 | 1 | 1 | 1 | 0.99 | 1 |
| Q8C5N3 | CWC22_MOUSE | 5 | 4 | 6 | 3 | 0 | 4 | 1 | 1 | 1 | 1 | 1 | 1 |
| Q61543 | GSLG1_MOUSE | 7 | 12 | 17 | 4 | 0 | 7 | 1 | 1 | 1 | 1 | 0.85 | 1 |
| G3X8T2 | G3X8T2_MOUSE | 16 | 14 | 17 | 13 | 6 | 8 | 1 | 1 | 1 | 1 | 1 | 1 |
| O35326 | SRSF5_MOUSE | 23 | 15 | 17 | 18 | 11 | 5 | 1 | 1 | 1 | 1 | 1 | 1 |
| Q5SWZ5 | Q5SWZ5_MOUSE | 32 | 36 | 33 | 26 | 7 | 10 | 1 | 1 | 1 | 1 | 1 | 1 |
| P68040 | GBLP_MOUSE | 24 | 20 | 15 | 18 | 11 | 11 | 1 | 1 | 1 | 1 | 1 | 1 |
| Q8BTV1 | TUSC3_MOUSE | 1 | 2 | 3 | 2 | 1 | 1 | 1 | 1 | 1 | 1 | 0.97 | 0.62 |
| Q80YR9 | R12BA_MOUSE | 2 | 2 | 3 | 0 | 0 | 1 | 1 | 1 | 1 | 0.45 | 0.29 | 0.82 |
| Q8R1A4 | DOCK7_MOUSE | 18 | 6 | 6 | 18 | 5 | 1 | 1 | 1 | 1 | 1 | 1 | 0.95 |
| Q921L3 | TMCO1_MOUSE | 6 | 2 | 4 | 3 | 3 | 2 | 1 | 1 | 1 | 1 | 1 | 1 |
| Q0KL02 | TRIO_MOUSE | 3 | 9 | 2 | 4 | 2 | 0 | 1 | 1 | 1 | 1 | 1 | 0 |
| Q91YN9 | BAG2_MOUSE | 1 | 1 | 1 | 1 | 1 | 0 | 0.99 | 0.82 | 0.84 | 0.99 | 0.97 | 0 |
| Q8JZM7 | CDC73_MOUSE | 2 | 1 | 2 | 0 | 1 | 0 | 1 | 1 | 1 | 0.93 | 0.99 | 0 |
| Q7TQH0 | ATX2L_MOUSE | 15 | 8 | 17 | 7 | 3 | 7 | 1 | 1 | 1 | 1 | 1 | 1 |
| Q9DBE7 | Q9DBE7_MOUSE | 11 | 8 | 3 | 4 | 0 | 0 | 1 | 1 | 1 | 1 | 1 | 0.19 |
| Q9D6M3 | GHC1_MOUSE | 5 | 5 | 1 | 4 | 2 | 0 | 1 | 1 | 1 | 1 | 1 | 0 |
| P51881 | ADT2_MOUSE | 29 | 19 | 23 | 25 | 7 | 14 | 1 | 1 | 1 | 1 | 1 | 1 |
| O70551 | SRPK1_MOUSE | 2 | 2 | 2 | 0 | 0 | 1 | 1 | 1 | 1 | 0.26 | 0 | 0.87 |
| P61620 | S61A1_MOUSE | 11 | 13 | 12 | 6 | 6 | 7 | 1 | 1 | 1 | 1 | 1 | 1 |
| Q99LF4 | RTCB_MOUSE | 18 | 10 | 14 | 10 | 5 | 5 | 1 | 1 | 1 | 1 | 1 | 1 |
| P61358 | RL27_MOUSE | 0 | 10 | 1 | 0 | 8 | 1 | 0.94 | 1 | 0.99 | 0.37 | 1 | 0.89 |

| | | | | | | | | | | | | | |
|---|---|---|---|---|---|---|---|---|---|---|---|---|---|
| E9QNN1 | E9QNN1_MOUSE | 17 | 7 | 13 | 8 | 2 | 8 | 1 | 1 | 1 | 1 | 1 | 1 |
| Q8K205 | Q8K205_MOUSE | 4 | 0 | 0 | 1 | 0 | 0 | 1 | 1 | 1 | 0.99 | 0 | 0.98 |
| Q60598 | SRC8_MOUSE | 29 | 39 | 34 | 22 | 16 | 10 | 1 | 1 | 1 | 1 | 1 | 1 |
| P68033 | ACTC_MOUSE | 30 | 25 | 38 | 21 | 17 | 22 | 1 | 1 | 1 | 1 | 1 | 1 |
| Q8VEJ4 | NLE1_MOUSE | 2 | 1 | 1 | 0 | 1 | 1 | 1 | 0.92 | 0.98 | 0.97 | 0.91 | 0.99 |
| Q6P5B0 | RRP12_MOUSE | 11 | 5 | 13 | 6 | 2 | 5 | 1 | 1 | 1 | 1 | 1 | 1 |
| O54784 | DAPK3_MOUSE | 8 | 7 | 7 | 4 | 3 | 1 | 1 | 1 | 1 | 1 | 1 | 0.98 |
| Q01149 | CO1A2_MOUSE | 23 | 25 | 37 | 16 | 8 | 19 | 1 | 1 | 1 | 1 | 1 | 1 |
| Q91YR7 | PRP6_MOUSE | 1 | 19 | 16 | 0 | 3 | 13 | 0.93 | 1 | 1 | 0 | 1 | 1 |
| Q8VDD5 | MYH9_MOUSE | 335 | 283 | 439 | 281 | 189 | 298 | 1 | 1 | 1 | 1 | 1 | 1 |
| P47757 | CAPZB_MOUSE | 9 | 11 | 17 | 7 | 6 | 6 | 1 | 1 | 1 | 1 | 1 | 1 |
| Q921F2 | TADBP_MOUSE | 3 | 1 | 2 | 1 | 0 | 0 | 1 | 1 | 1 | 1 | 0.65 | 0 |
| P38647 | GRP75_MOUSE | 1 | 3 | 1 | 1 | 3 | 0 | 0.99 | 1 | 1 | 1 | 1 | 0 |
| P62702 | RS4X_MOUSE | 27 | 28 | 29 | 25 | 19 | 15 | 1 | 1 | 1 | 1 | 1 | 1 |
| Q04736 | YES_MOUSE | 3 | 6 | 4 | 2 | 3 | 3 | 1 | 1 | 1 | 1 | 1 | 1 |
| P08121 | CO3A1_MOUSE | 7 | 4 | 7 | 4 | 0 | 2 | 1 | 1 | 1 | 1 | 0.88 | 1 |
| Q60716 | P4HA2_MOUSE | 5 | 5 | 3 | 2 | 2 | 0 | 1 | 1 | 1 | 0.99 | 1 | 0 |
| P35980 | RL18_MOUSE | 34 | 19 | 7 | 28 | 17 | 21 | 1 | 1 | 1 | 1 | 1 | 1 |
| E9Q634 | MYO1E_MOUSE | 8 | 13 | 29 | 8 | 0 | 9 | 1 | 1 | 1 | 1 | 1 | 1 |
| Q62167 | DDX3X_MOUSE | 40 | 27 | 41 | 36 | 21 | 22 | 1 | 1 | 1 | 1 | 1 | 1 |
| Q80YR5 | SAFB2_MOUSE | 1 | 0 | 2 | 0 | 0 | 0 | 1 | 0.96 | 1 | 0.27 | 0.24 | 0 |
| Q6ZQ11 | CHSS1_MOUSE | 2 | 2 | 2 | 1 | 0 | 0 | 0.98 | 1 | 1 | 0.98 | 0 | 0 |
| Q80YQ1 | Q80YQ1_MOUSE | 37 | 0 | 8 | 21 | 9 | 2 | 1 | 1 | 1 | 1 | 1 | 1 |
| Q99104 | MYO5A_MOUSE | 46 | 36 | 83 | 26 | 15 | 19 | 1 | 1 | 1 | 1 | 1 | 1 |
| E1U8D0 | SOGA1_MOUSE | 4 | 2 | 3 | 2 | 0 | 0 | 1 | 1 | 1 | 1 | 0.43 | 0 |
| P47962 | RL5_MOUSE | 44 | 34 | 54 | 39 | 29 | 28 | 1 | 1 | 1 | 1 | 1 | 1 |
| Q91V55 | Q91V55_MOUSE | 22 | 0 | 1 | 15 | 0 | 15 | 1 | 1 | 0.87 | 1 | 1 | 1 |
| Q3TIX9 | SNUT2_MOUSE | 11 | 4 | 4 | 4 | 4 | 2 | 1 | 1 | 1 | 1 | 1 | 1 |
| P19253 | RL13A_MOUSE | 38 | 22 | 6 | 26 | 18 | 30 | 1 | 1 | 1 | 1 | 1 | 1 |
| O70194 | EIF3D_MOUSE | 7 | 8 | 7 | 4 | 4 | 1 | 1 | 1 | 1 | 1 | 1 | 0.97 |
| F7B5B5 | F7B5B5_MOUSE | 1 | 4 | 1 | 1 | 2 | 0 | 1 | 1 | 0.95 | 0.87 | 1 | 0 |
| Q9CX11 | UTP23_MOUSE | 0 | 2 | 2 | 0 | 1 | 0 | 0.93 | 1 | 1 | 0.44 | 0.93 | 0 |
| Q6ZQI3 | MLEC_MOUSE | 2 | 9 | 2 | 3 | 3 | 1 | 1 | 1 | 1 | 1 | 1 | 0.99 |
| Q9D903 | EBP2_MOUSE | 8 | 3 | 7 | 3 | 1 | 2 | 1 | 1 | 1 | 1 | 0.89 | 1 |
| P14206 | RSSA_MOUSE | 23 | 21 | 22 | 21 | 19 | 23 | 1 | 1 | 1 | 1 | 1 | 1 |
| Q9QZD4 | XPF_MOUSE | 4 | 4 | 25 | 0 | 0 | 0 | 1 | 1 | 1 | 0 | 0 | 0 |
| Q8VI84 | NOC3L_MOUSE | 1 | 0 | 2 | 0 | 0 | 0 | 0.99 | 1 | 1 | 0.27 | 0.55 | 0 |
| Q6PE01 | SNR40_MOUSE | 3 | 2 | 1 | 0 | 0 | 0 | 1 | 1 | 0.99 | 0.93 | 0.36 | 0 |
| Q61263 | SOAT1_MOUSE | 5 | 1 | 6 | 1 | 0 | 0 | 1 | 1 | 1 | 1 | 0 | 0 |
| P07903 | ERCC1_MOUSE | 1 | 3 | 7 | 0 | 0 | 0 | 1 | 1 | 1 | 0.57 | 0.9 | 0 |
| Q922K7 | NOP2_MOUSE | 1 | 4 | 1 | 0 | 0 | 0 | 0.95 | 1 | 0.94 | 0.33 | 0.66 | 0 |
| O35218 | CPSF2_MOUSE | 4 | 5 | 3 | 0 | 0 | 0 | 1 | 1 | 1 | 0.97 | 0.31 | 0 |
| P25911 | LYN_MOUSE | 4 | 3 | 2 | 0 | 0 | 0 | 1 | 1 | 1 | 0.53 | 0.72 | 0 |
| E9Q3L2 | E9Q3L2_MOUSE | 1 | 0 | 1 | 4 | 0 | 1 | 1 | 0.99 | 1 | 1 | 0.99 | 0.87 |
| Q8CH18 | CCAR1_MOUSE | 0 | 2 | 2 | 0 | 0 | 0 | 0.86 | 1 | 1 | 0.61 | 0 | 0.16 |
| Q9QYJ0 | DNJA2_MOUSE | 1 | 2 | 1 | 0 | 0 | 0 | 0.99 | 1 | 0.97 | 0.95 | 0.44 | 0 |
| P67778 | PHB_MOUSE | 1 | 2 | 1 | 0 | 0 | 0 | 1 | 1 | 0.97 | 0.33 | 0.94 | 0 |
| Q6ZQL4 | WDR43_MOUSE | 2 | 1 | 2 | 0 | 0 | 0 | 1 | 0.81 | 1 | 0.43 | 0 | 0 |
| Q31125 | S39A7_MOUSE | 1 | 1 | 4 | 0 | 0 | 0 | 1 | 0.98 | 1 | 0 | 0.71 | 0 |
| Q9WV55 | VAPA_MOUSE | 1 | 2 | 1 | 0 | 0 | 0 | 1 | 1 | 1 | 0.46 | 0.98 | 0 |
| Q922V4 | PLRG1_MOUSE | 2 | 2 | 1 | 0 | 0 | 0 | 1 | 1 | 0.98 | 0.25 | 0.92 | 0.92 |
| E9PVZ8 | E9PVZ8_MOUSE | 4 | 0 | 2 | 0 | 0 | 0 | 1 | 1 | 1 | 0.1 | 0 | 0 |
| Q3UU43 | Q3UU43_MOUSE | 3 | 0 | 2 | 0 | 0 | 0 | 1 | 0.99 | 1 | 0.28 | 0.96 | 0.39 |
| E9PWW6 | E9PWW6_MOUSE | 4 | 0 | 0 | 0 | 0 | 0 | 1 | 1 | 1 | 0.3 | 0 | 0.27 |
| Q921M4 | GOGA2_MOUSE | 2 | 3 | 3 | 0 | 0 | 0 | 1 | 1 | 1 | 0.17 | 0 | 0 |
| P47758 | SRPRB_MOUSE | 2 | 5 | 2 | 1 | 0 | 0 | 1 | 1 | 1 | 0.98 | 0.62 | 0 |
| O88455 | DHCR7_MOUSE | 2 | 0 | 3 | 4 | 0 | 1 | 1 | 0.93 | 1 | 1 | 0 | 0.99 |
| P58854 | GCP3_MOUSE | 1 | 1 | 2 | 0 | 0 | 0 | 0.99 | 1 | 1 | 0.23 | 0 | 0 |
| Q61749 | EI2BD_MOUSE | 1 | 2 | 1 | 0 | 0 | 0 | 1 | 1 | 0.89 | 0.98 | 0.65 | 0 |
| P04925 | PRIO_MOUSE | 1 | 1 | 1 | 0 | 0 | 0 | 0.99 | 0.99 | 0.99 | 0 | 0.94 | 0 |
| Q9CYA6 | ZCHC8_MOUSE | 3 | 0 | 1 | 0 | 0 | 0 | 1 | 1 | 1 | 0.23 | 0 | 0 |
| Q9Z2A7 | DGAT1_MOUSE | 1 | 1 | 1 | 0 | 0 | 0 | 1 | 0.94 | 0.98 | 0.27 | 0 | 0 |
| Q8BJS4 | SUN2_MOUSE | 3 | 2 | 3 | 0 | 0 | 0 | 1 | 1 | 1 | 0 | 0.51 | 0 |

| | | | | | | | | | | | | | |
|---|---|---|---|---|---|---|---|---|---|---|---|---|---|
| Q8R480 | NUP85_MOUSE | 1 | 1 | 1 | 0 | 0 | 0 | 0.99 | 1 | 0.94 | 0 | 0 | 0 |
| Q6ZPE2 | MTMR5_MOUSE | 1 | 1 | 2 | 0 | 0 | 0 | 0.98 | 0.95 | 0.85 | 0.38 | 0.35 | 0 |
| Q9R233 | TPSN_MOUSE | 0 | 1 | 1 | 0 | 0 | 0 | 0.99 | 0.98 | 0.96 | 0 | 0.49 | 0 |
| Q69Z99 | ZN512_MOUSE | 1 | 0 | 2 | 0 | 0 | 0 | 0.95 | 0.86 | 1 | 0.18 | 0.21 | 0 |
| Q6ZPF4 | FMNL3_MOUSE | 2 | 2 | 1 | 0 | 0 | 0 | 1 | 0.99 | 0.95 | 0 | 0.48 | 0 |
| Q1W617 | SHRM4_MOUSE | 3 | 0 | 1 | 0 | 0 | 0 | 1 | 0.94 | 0.95 | 0 | 0 | 0 |
| Q05186 | RCN1_MOUSE | 2 | 1 | 1 | 0 | 0 | 0 | 1 | 0.88 | 0.99 | 0 | 0 | 0 |
| Q3TDN2 | FAF2_MOUSE | 0 | 1 | 1 | 0 | 0 | 0 | 0.97 | 1 | 0.98 | 0 | 0 | 0 |
| Q91YI6 | GLD2_MOUSE | 3 | 2 | 1 | 0 | 0 | 0 | 1 | 1 | 1 | 0 | 0 | 0 |
| P70298 | CUX2_MOUSE | 1 | 1 | 1 | 0 | 0 | 0 | 0.84 | 0.8 | 0.97 | 0 | 0.37 | 0 |
| Q62158 | TRI27_MOUSE | 2 | 2 | 1 | 0 | 0 | 0 | 1 | 0.98 | 0.97 | 0 | 0 | 0 |
| P13011 | ACOD2_MOUSE | 1 | 1 | 1 | 0 | 0 | 0 | 0.94 | 0.9 | 0.89 | 0 | 0 | 0 |
| Q8CH25 | SLTM_MOUSE | 12 | 7 | 29 | 8 | 6 | 8 | 1 | 1 | 1 | 1 | 1 | 1 |
| Q9JKF1 | IQGA1_MOUSE | 24 | 19 | 18 | 30 | 7 | 3 | 1 | 1 | 1 | 1 | 1 | 1 |
| Q9D0R4 | DDX56_MOUSE | 1 | 2 | 1 | 1 | 1 | 0 | 1 | 1 | 1 | 0.93 | 0.99 | 0 |
| E9Q7L1 | E9Q7L1_MOUSE | 5 | 4 | 0 | 2 | 0 | 0 | 1 | 1 | 1 | 1 | 0.67 | 1 |
| Q61656 | DDX5_MOUSE | 54 | 36 | 48 | 41 | 19 | 24 | 1 | 1 | 1 | 1 | 1 | 1 |
| P62960 | YBOX1_MOUSE | 12 | 3 | 6 | 7 | 1 | 2 | 1 | 1 | 1 | 1 | 1 | 1 |
| P54823 | DDX6_MOUSE | 9 | 4 | 5 | 6 | 2 | 0 | 1 | 1 | 1 | 1 | 1 | 0 |
| Q9D0R8 | LSM12_MOUSE | 8 | 6 | 3 | 7 | 2 | 4 | 1 | 1 | 1 | 1 | 1 | 1 |
| Q9Z315 | SNUT1_MOUSE | 14 | 11 | 18 | 10 | 0 | 9 | 1 | 1 | 1 | 1 | 0.37 | 1 |
| Q9WTM5 | RUVB2_MOUSE | 11 | 4 | 5 | 6 | 1 | 1 | 1 | 1 | 1 | 1 | 1 | 0.99 |
| P14148 | RL7_MOUSE | 48 | 40 | 57 | 43 | 41 | 41 | 1 | 1 | 1 | 1 | 1 | 1 |
| Q64511 | TOP2B_MOUSE | 10 | 21 | 13 | 6 | 9 | 3 | 1 | 1 | 1 | 1 | 1 | 1 |
| P62737 | ACTA_MOUSE | 1 | 1 | 1 | 2 | 0 | 1 | 0.99 | 0.99 | 1 | 1 | 0.92 | 0.99 |
| K3W4R2 | K3W4R2_MOUSE | 2 | 1 | 2 | 1 | 2 | 0 | 1 | 0.98 | 1 | 0.92 | 1 | 0 |
| Q8VEM8 | MPCP_MOUSE | 4 | 4 | 6 | 4 | 1 | 2 | 1 | 1 | 1 | 1 | 0.97 | 1 |
| Q62093 | SRSF2_MOUSE | 9 | 8 | 3 | 5 | 3 | 1 | 1 | 1 | 1 | 1 | 1 | 0.97 |
| E9PYF4 | E9PYF4_MOUSE | 28 | 0 | 0 | 16 | 0 | 0 | 1 | 1 | 1 | 1 | 1 | 1 |
| Q5XJF6 | Q5XJF6_MOUSE | 14 | 0 | 9 | 12 | 0 | 0 | 1 | 1 | 1 | 1 | 1 | 1 |
| Q9DBG6 | RPN2_MOUSE | 14 | 16 | 10 | 9 | 6 | 5 | 1 | 1 | 1 | 1 | 1 | 1 |
| P59222 | SREC2_MOUSE | 2 | 1 | 4 | 0 | 0 | 1 | 1 | 1 | 1 | 0.09 | 0 | 1 |
| Q8C570 | RAE1L_MOUSE | 1 | 2 | 3 | 2 | 1 | 0 | 1 | 1 | 1 | 1 | 1 | 0 |
| Q9CY58 | PAIRB_MOUSE | 23 | 9 | 7 | 15 | 6 | 3 | 1 | 1 | 1 | 1 | 1 | 1 |
| P48962 | ADT1_MOUSE | 13 | 7 | 8 | 5 | 5 | 3 | 1 | 1 | 1 | 1 | 1 | 1 |
| Q99PL5 | RRBP1_MOUSE | 68 | 0 | 0 | 50 | 48 | 48 | 1 | 1 | 1 | 1 | 1 | 1 |
| P27659 | RL3_MOUSE | 56 | 42 | 51 | 50 | 42 | 24 | 1 | 1 | 1 | 1 | 1 | 1 |
| Q3TEA8 | HP1B3_MOUSE | 12 | 9 | 18 | 7 | 8 | 6 | 1 | 1 | 1 | 1 | 1 | 1 |
| Q9JHJ0 | TMOD3_MOUSE | 13 | 9 | 15 | 15 | 8 | 8 | 1 | 1 | 1 | 1 | 1 | 1 |
| P43277 | H13_MOUSE | 12 | 12 | 32 | 9 | 12 | 17 | 1 | 1 | 1 | 1 | 1 | 1 |
| Q9ERU9 | RBP2_MOUSE | 48 | 15 | 27 | 35 | 5 | 3 | 1 | 1 | 1 | 1 | 1 | 1 |
| D3Z0M9 | D3Z0M9_MOUSE | 12 | 0 | 0 | 11 | 6 | 0 | 1 | 1 | 1 | 1 | 1 | 1 |
| P43274 | H14_MOUSE | 3 | 4 | 10 | 2 | 4 | 6 | 1 | 1 | 1 | 1 | 1 | 1 |
| Q99JX4 | EIF3M_MOUSE | 3 | 3 | 1 | 1 | 3 | 0 | 1 | 1 | 1 | 1 | 1 | 0 |
| Q7TPV4 | MBB1A_MOUSE | 81 | 76 | 111 | 72 | 27 | 49 | 1 | 1 | 1 | 1 | 1 | 1 |
| Q3TMW1 | C102A_MOUSE | 5 | 11 | 8 | 3 | 10 | 1 | 1 | 1 | 1 | 1 | 1 | 0.76 |
| Q3V1Z5 | Q3V1Z5_MOUSE | 4 | 0 | 2 | 2 | 1 | 0 | 1 | 0.96 | 1 | 1 | 0.94 | 1 |
| Q9EPU0 | RENT1_MOUSE | 19 | 16 | 22 | 7 | 0 | 8 | 1 | 1 | 1 | 1 | 0 | 1 |
| Q04750 | TOP1_MOUSE | 2 | 9 | 4 | 4 | 0 | 2 | 1 | 1 | 1 | 1 | 0.57 | 1 |
| Q6NV83 | SR140_MOUSE | 10 | 7 | 9 | 9 | 0 | 1 | 1 | 1 | 1 | 1 | 0.64 | 1 |
| Q9D8E6 | RL4_MOUSE | 70 | 60 | 78 | 58 | 45 | 44 | 1 | 1 | 1 | 1 | 1 | 1 |
| Q922U1 | PRPF3_MOUSE | 9 | 10 | 7 | 3 | 7 | 2 | 1 | 1 | 1 | 1 | 1 | 1 |
| Q6NS46 | RRP5_MOUSE | 5 | 3 | 2 | 5 | 1 | 0 | 1 | 1 | 1 | 1 | 0.99 | 0 |
| P70279 | SURF6_MOUSE | 3 | 4 | 2 | 3 | 3 | 0 | 1 | 1 | 1 | 1 | 0.99 | 0 |
| Q91VJ2 | PRDBP_MOUSE | 5 | 2 | 7 | 5 | 3 | 0 | 1 | 1 | 1 | 1 | 1 | 0 |
| P62911 | RL32_MOUSE | 2 | 9 | 1 | 1 | 9 | 2 | 0.99 | 1 | 1 | 0.88 | 1 | 1 |
| P63260 | ACTG_MOUSE | 63 | 57 | 72 | 55 | 50 | 52 | 1 | 1 | 1 | 1 | 1 | 1 |
| Q3UM18 | LSG1_MOUSE | 3 | 1 | 5 | 2 | 3 | 1 | 1 | 1 | 1 | 1 | 1 | 1 |
| Q9Z1X4 | ILF3_MOUSE | 15 | 13 | 26 | 10 | 0 | 13 | 1 | 1 | 1 | 1 | 1 | 1 |
| Q9DAW6 | PRP4_MOUSE | 6 | 8 | 3 | 3 | 4 | 2 | 1 | 1 | 1 | 1 | 1 | 1 |
| O88207 | CO5A1_MOUSE | 14 | 16 | 16 | 9 | 7 | 4 | 1 | 1 | 1 | 1 | 1 | 1 |
| Q7TQD7 | Q7TQD7_MOUSE | 17 | 2 | 43 | 12 | 0 | 13 | 1 | 1 | 1 | 1 | 0 | 1 |
| Q61584 | FXR1_MOUSE | 18 | 26 | 17 | 18 | 13 | 9 | 1 | 1 | 1 | 1 | 1 | 1 |

| | | | | | | | | | | | | | |
|---|---|---|---|---|---|---|---|---|---|---|---|---|---|
| B7ZNY3 | B7ZNY3_MOUSE | 2 | 0 | 0 | 0 | 0 | 1 | 1 | 1 | 1 | 0.98 | 0 | 1 |
| P68372 | TBB4B_MOUSE | 25 | 9 | 13 | 11 | 6 | 4 | 1 | 1 | 1 | 1 | 1 | 1 |
| Q99NB9 | SF3B1_MOUSE | 19 | 15 | 19 | 15 | 1 | 6 | 1 | 1 | 1 | 1 | 1 | 1 |
| Q9D0B0 | SRSF9_MOUSE | 4 | 4 | 2 | 3 | 1 | 0 | 1 | 1 | 0.99 | 1 | 1 | 0 |
| E9QML5 | E9QML5_MOUSE | 18 | 5 | 9 | 14 | 3 | 0 | 1 | 1 | 1 | 1 | 1 | 0.18 |
| Q07113 | MPRI_MOUSE | 5 | 0 | 3 | 4 | 2 | 3 | 1 | 1 | 1 | 1 | 1 | 1 |
| Q9CXY6 | ILF2_MOUSE | 15 | 7 | 18 | 16 | 12 | 8 | 1 | 1 | 1 | 1 | 1 | 1 |
| Q9ERA6 | TFP11_MOUSE | 3 | 6 | 7 | 0 | 1 | 5 | 1 | 1 | 1 | 0.97 | 0.93 | 1 |
| P70255 | NFIC_MOUSE | 7 | 3 | 2 | 2 | 3 | 1 | 1 | 1 | 1 | 1 | 1 | 1 |
| Q9CS00 | CATIN_MOUSE | 2 | 4 | 3 | 2 | 0 | 0 | 1 | 1 | 1 | 1 | 0 | 0 |
| Q6ZPR5 | NSMA3_MOUSE | 1 | 3 | 1 | 1 | 0 | 1 | 0.8 | 1 | 0.95 | 0.86 | 0.42 | 0.95 |
| Q9CWK3 | CD2B2_MOUSE | 2 | 2 | 1 | 1 | 0 | 0 | 1 | 1 | 0.94 | 1 | 0 | 0 |
| Q8VH51 | RBM39_MOUSE | 19 | 16 | 14 | 16 | 11 | 10 | 1 | 1 | 1 | 1 | 1 | 1 |
| Q80VI1 | TRI56_MOUSE | 6 | 5 | 3 | 5 | 3 | 0 | 1 | 1 | 1 | 1 | 1 | 0 |
| Q9QYB5 | ADDG_MOUSE | 0 | 3 | 4 | 0 | 2 | 4 | 1 | 1 | 1 | 0.11 | 1 | 1 |
| P62852 | RS25_MOUSE | 3 | 10 | 3 | 1 | 8 | 1 | 1 | 1 | 1 | 1 | 1 | 0.99 |
| Q9Z1M8 | RED_MOUSE | 2 | 4 | 1 | 1 | 1 | 0 | 1 | 1 | 0.95 | 1 | 0.98 | 0 |
| P62918 | RL8_MOUSE | 28 | 27 | 37 | 26 | 18 | 22 | 1 | 1 | 1 | 1 | 1 | 1 |
| P47911 | RL6_MOUSE | 40 | 38 | 47 | 41 | 32 | 33 | 1 | 1 | 1 | 1 | 1 | 1 |
| O35646 | CAN6_MOUSE | 1 | 2 | 8 | 3 | 5 | 3 | 1 | 1 | 1 | 1 | 1 | 1 |
| G3UWX1 | G3UWX1_MOUSE | 6 | 0 | 8 | 4 | 0 | 3 | 1 | 1 | 1 | 1 | 0 | 1 |
| E9QPE7 | E9QPE7_MOUSE | 31 | 16 | 29 | 4 | 0 | 1 | 1 | 1 | 1 | 1 | 1 | 0.99 |
| Q922Q4 | P5CR2_MOUSE | 1 | 2 | 1 | 0 | 1 | 1 | 0.99 | 1 | 0.94 | 0 | 0.98 | 0.89 |
| Q8C7X2 | EMC1_MOUSE | 4 | 4 | 2 | 2 | 2 | 0 | 1 | 1 | 0.95 | 0.99 | 0.93 | 0 |
| Q3U962 | CO5A2_MOUSE | 13 | 3 | 14 | 14 | 0 | 5 | 1 | 1 | 1 | 1 | 1 | 1 |
| Q80U78 | PUM1_MOUSE | 3 | 2 | 5 | 1 | 0 | 1 | 1 | 1 | 1 | 0.99 | 0.79 | 0.99 |
| P84104 | SRSF3_MOUSE | 8 | 4 | 4 | 7 | 4 | 1 | 1 | 1 | 1 | 1 | 1 | 0.8 |
| Q8BU14 | SEC62_MOUSE | 1 | 2 | 2 | 1 | 0 | 1 | 1 | 1 | 1 | 0.89 | 0.98 | 0.97 |
| P26369 | U2AF2_MOUSE | 18 | 17 | 12 | 15 | 12 | 7 | 1 | 1 | 1 | 1 | 1 | 1 |
| Q9Z0U1 | ZO2_MOUSE | 19 | 12 | 18 | 15 | 2 | 8 | 1 | 1 | 1 | 1 | 0.99 | 1 |
| E9QAZ2 | E9QAZ2_MOUSE | 23 | 15 | 7 | 17 | 9 | 21 | 1 | 1 | 1 | 1 | 1 | 1 |
| Q8BK35 | Q8BK35_MOUSE | 5 | 0 | 1 | 2 | 0 | 0 | 1 | 1 | 0.95 | 1 | 0 | 0 |
| D3YWS8 | D3YWS8_MOUSE | 3 | 0 | 0 | 3 | 0 | 1 | 1 | 1 | 1 | 1 | 0.71 | 1 |
| Q9ESX5 | DKC1_MOUSE | 8 | 7 | 4 | 6 | 4 | 2 | 1 | 1 | 1 | 1 | 1 | 1 |
| Q62261 | SPTB2_MOUSE | 35 | 54 | 21 | 28 | 21 | 3 | 1 | 1 | 1 | 1 | 1 | 1 |
| B2RSH2 | GNAI1_MOUSE | 3 | 1 | 2 | 1 | 0 | 0 | 1 | 0.9 | 1 | 0.99 | 0 | 0 |
| Q8BWW4 | LARP4_MOUSE | 8 | 1 | 4 | 3 | 0 | 1 | 1 | 1 | 1 | 1 | 0 | 1 |
| P62908 | RS3_MOUSE | 37 | 32 | 39 | 32 | 29 | 23 | 1 | 1 | 1 | 1 | 1 | 1 |
| Q9EPR5 | SORC2_MOUSE | 2 | 4 | 5 | 0 | 0 | 3 | 1 | 1 | 1 | 0.24 | 0 | 1 |
| B7FAU9 | B7FAU9_MOUSE | 56 | 41 | 44 | 38 | 21 | 20 | 1 | 1 | 1 | 1 | 1 | 1 |
| F7DBB3 | F7DBB3_MOUSE | 3 | 0 | 4 | 2 | 0 | 0 | 1 | 1 | 1 | 1 | 1 | 0 |
| Q80XI3 | IF4G3_MOUSE | 16 | 3 | 7 | 8 | 3 | 2 | 1 | 1 | 1 | 1 | 1 | 1 |
| P62082 | RS7_MOUSE | 26 | 19 | 3 | 16 | 12 | 15 | 1 | 1 | 1 | 1 | 1 | 1 |
| Q4VBF2 | R3HD4_MOUSE | 1 | 2 | 2 | 1 | 2 | 0 | 0.99 | 1 | 1 | 0.93 | 1 | 0 |
| Q3UJB0 | Q3UJB0_MOUSE | 6 | 0 | 2 | 3 | 0 | 0 | 1 | 1 | 1 | 1 | 0 | 0 |
| P61222 | ABCE1_MOUSE | 2 | 3 | 2 | 1 | 0 | 0 | 1 | 1 | 1 | 1 | 0 | 0 |
| Q99PU8 | DHX30_MOUSE | 4 | 7 | 6 | 2 | 0 | 0 | 1 | 1 | 1 | 1 | 0.38 | 0 |
| Q9JJ28 | FLII_MOUSE | 8 | 13 | 12 | 8 | 4 | 8 | 1 | 1 | 1 | 1 | 1 | 1 |
| P83887 | TBG1_MOUSE | 1 | 1 | 3 | 1 | 1 | 1 | 1 | 0.92 | 1 | 1 | 0.89 | 0.88 |
| Q8BJ05 | ZC3HE_MOUSE | 1 | 1 | 2 | 0 | 0 | 2 | 1 | 1 | 1 | 0 | 1 | 1 |
| O70589 | CSKP_MOUSE | 0 | 5 | 5 | 0 | 0 | 2 | 0.88 | 1 | 1 | 0.89 | 0.62 | 1 |
| Q3TWW8 | Q3TWW8_MOUSE | 15 | 3 | 9 | 16 | 2 | 2 | 1 | 1 | 1 | 1 | 1 | 1 |
| Q91ZW3 | SMCA5_MOUSE | 3 | 6 | 11 | 2 | 0 | 8 | 1 | 1 | 1 | 1 | 0.36 | 1 |
| H7BX23 | H7BX23_MOUSE | 5 | 0 | 6 | 0 | 0 | 2 | 1 | 1 | 1 | 0.96 | 1 | 1 |
| O35129 | PHB2_MOUSE | 1 | 7 | 3 | 0 | 3 | 0 | 0.92 | 1 | 1 | 0.63 | 1 | 0 |
| Q6PHZ5 | RB15B_MOUSE | 5 | 7 | 4 | 0 | 0 | 3 | 1 | 1 | 1 | 0.97 | 0 | 1 |
| Q9D0F6 | RFC5_MOUSE | 3 | 6 | 6 | 4 | 4 | 1 | 1 | 1 | 1 | 1 | 1 | 0.95 |
| Q9R0Q6 | ARC1A_MOUSE | 1 | 4 | 4 | 2 | 1 | 0 | 0.91 | 1 | 1 | 1 | 1 | 0.51 |
| P27048 | RSMB_MOUSE | 5 | 0 | 3 | 1 | 1 | 1 | 1 | 1 | 1 | 0.95 | 0.99 | 0.96 |
| Q9R1C7 | PR40A_MOUSE | 12 | 18 | 24 | 11 | 0 | 8 | 1 | 1 | 1 | 1 | 0.95 | 1 |
| Q8R3H7 | HS2ST_MOUSE | 3 | 1 | 2 | 3 | 2 | 0 | 1 | 1 | 1 | 1 | 0.86 | 0 |
| Q9CQT2 | RBM7_MOUSE | 4 | 3 | 3 | 3 | 1 | 0 | 1 | 1 | 1 | 1 | 0.86 | 0 |
| Q8BFQ4 | WDR82_MOUSE | 7 | 2 | 2 | 2 | 5 | 0 | 1 | 1 | 1 | 1 | 1 | 0 |

| | | | | | | | | | | | | | |
|---|---|---|---|---|---|---|---|---|---|---|---|---|---|
| Q5SUH6 | Q5SUH6_MOUSE | 4 | 3 | 1 | 1 | 0 | 0 | 1 | 1 | 1 | 0.95 | 0 | 0 |
| D3YYI8 | D3YYI8_MOUSE | 0 | 1 | 2 | 0 | 1 | 1 | 0.97 | 0.81 | 1 | 0.99 | 0.72 | 0.75 |
| O55029 | COPB2_MOUSE | 0 | 2 | 2 | 1 | 0 | 3 | 0.94 | 1 | 1 | 0.99 | 0.37 | 1 |
| Q00PI9 | HNRL2_MOUSE | 7 | 8 | 21 | 4 | 0 | 9 | 1 | 1 | 1 | 1 | 1 | 1 |
| Q3U741 | Q3U741_MOUSE | 33 | 16 | 14 | 20 | 0 | 14 | 1 | 1 | 1 | 1 | 1 | 1 |
| E9Q7G0 | E9Q7G0_MOUSE | 48 | 0 | 36 | 41 | 0 | 9 | 1 | 1 | 1 | 1 | 1 | 1 |
| P99024 | TBB5_MOUSE | 4 | 3 | 3 | 3 | 2 | 2 | 1 | 1 | 1 | 1 | 1 | 1 |
| P63101 | 1433Z_MOUSE | 1 | 3 | 1 | 1 | 2 | 0 | 0.99 | 1 | 0.86 | 1 | 1 | 0.58 |
| P97449 | AMPN_MOUSE | 3 | 5 | 19 | 2 | 3 | 6 | 1 | 1 | 1 | 1 | 1 | 1 |
| P25976 | UBF1_MOUSE | 2 | 5 | 5 | 0 | 2 | 3 | 1 | 1 | 1 | 0 | 1 | 1 |
| A2ADH1 | A2ADH1_MOUSE | 3 | 5 | 3 | 4 | 4 | 3 | 1 | 1 | 1 | 1 | 1 | 1 |
| Q9CX86 | ROA0_MOUSE | 5 | 6 | 2 | 4 | 2 | 0 | 1 | 1 | 1 | 1 | 1 | 0 |
| Q61937 | NPM_MOUSE | 19 | 21 | 15 | 13 | 16 | 6 | 1 | 1 | 1 | 1 | 1 | 1 |
| P23242 | CXA1_MOUSE | 3 | 9 | 8 | 5 | 4 | 0 | 1 | 1 | 1 | 1 | 1 | 0 |
| Q8K4L3 | SVIL_MOUSE | 43 | 13 | 1 | 37 | 1 | 23 | 1 | 1 | 1 | 1 | 1 | 1 |
| Q5EBP8 | Q5EBP8_MOUSE | 1 | 3 | 1 | 1 | 1 | 0 | 1 | 1 | 0.85 | 0.99 | 0.97 | 0 |
| Q80U58 | PUM2_MOUSE | 6 | 4 | 12 | 3 | 0 | 5 | 1 | 1 | 1 | 1 | 0.35 | 1 |
| Q3UMC0 | SPAT5_MOUSE | 3 | 1 | 8 | 0 | 0 | 4 | 1 | 1 | 1 | 0.87 | 0 | 1 |
| P08752 | GNAI2_MOUSE | 15 | 23 | 26 | 13 | 18 | 9 | 1 | 1 | 1 | 1 | 1 | 1 |
| P47754 | CAZA2_MOUSE | 7 | 2 | 9 | 5 | 3 | 5 | 1 | 1 | 1 | 1 | 1 | 1 |
| Q99PV0 | PRP8_MOUSE | 41 | 28 | 49 | 32 | 10 | 9 | 1 | 1 | 1 | 1 | 1 | 1 |
| Q61879 | MYH10_MOUSE | 226 | 178 | 361 | 201 | 134 | 193 | 1 | 1 | 1 | 1 | 1 | 1 |
| Q5RKN9 | Q5RKN9_MOUSE | 7 | 8 | 9 | 4 | 4 | 2 | 1 | 1 | 1 | 1 | 1 | 1 |
| Q9DBD5 | PELP1_MOUSE | 6 | 7 | 10 | 5 | 6 | 6 | 1 | 1 | 1 | 1 | 1 | 1 |
| E9PU96 | E9PU96_MOUSE | 8 | 8 | 4 | 9 | 1 | 0 | 1 | 1 | 1 | 1 | 1 | 0 |
| P60335 | PCBP1_MOUSE | 6 | 4 | 8 | 7 | 4 | 3 | 1 | 1 | 1 | 1 | 1 | 1 |
| P16546 | SPTN1_MOUSE | 30 | 45 | 24 | 27 | 16 | 4 | 1 | 1 | 1 | 1 | 1 | 1 |
| O08573 | LEG9_MOUSE | 2 | 4 | 1 | 0 | 3 | 0 | 1 | 1 | 0.97 | 0 | 1 | 0 |
| Q9QWV9 | CCNT1_MOUSE | 4 | 3 | 2 | 3 | 1 | 0 | 1 | 1 | 1 | 1 | 0.99 | 0 |
| Q80YX1 | TENA_MOUSE | 23 | 29 | 4 | 13 | 11 | 4 | 1 | 1 | 1 | 1 | 1 | 1 |
| Q9R0E1 | PLOD3_MOUSE | 9 | 10 | 7 | 8 | 4 | 3 | 1 | 1 | 1 | 1 | 1 | 1 |
| P23249 | MOV10_MOUSE | 2 | 3 | 2 | 2 | 0 | 3 | 1 | 1 | 1 | 1 | 0 | 1 |
| Q60749 | KHDR1_MOUSE | 8 | 4 | 6 | 10 | 4 | 1 | 1 | 1 | 1 | 1 | 1 | 0.97 |
| Q811D0 | DLG1_MOUSE | 2 | 3 | 3 | 1 | 0 | 3 | 1 | 1 | 1 | 0.99 | 0.84 | 1 |
| P97351 | RS3A_MOUSE | 33 | 30 | 41 | 29 | 28 | 23 | 1 | 1 | 1 | 1 | 1 | 1 |
| P35922 | FMR1_MOUSE | 17 | 14 | 18 | 11 | 8 | 9 | 1 | 1 | 1 | 1 | 1 | 1 |
| O35737 | HNRH1_MOUSE | 8 | 3 | 10 | 7 | 4 | 2 | 1 | 1 | 1 | 1 | 1 | 1 |
| G3X922 | G3X922_MOUSE | 33 | 0 | 32 | 21 | 0 | 0 | 1 | 1 | 1 | 1 | 1 | 1 |
| Q60760 | GRB10_MOUSE | 2 | 3 | 2 | 3 | 1 | 0 | 1 | 1 | 1 | 1 | 1 | 0.93 |
| F8VQC1 | F8VQC1_MOUSE | 9 | 0 | 4 | 5 | 0 | 0 | 1 | 1 | 1 | 1 | 1 | 0 |
| O55143 | AT2A2_MOUSE | 5 | 11 | 9 | 4 | 0 | 1 | 1 | 1 | 1 | 1 | 0.71 | 1 |
| P52479 | UBP10_MOUSE | 3 | 1 | 3 | 2 | 0 | 1 | 1 | 1 | 1 | 1 | 0 | 0.91 |
| Q9DCF9 | SSRG_MOUSE | 6 | 2 | 3 | 1 | 1 | 5 | 1 | 1 | 1 | 1 | 0.99 | 1 |
| P46978 | STT3A_MOUSE | 6 | 6 | 12 | 9 | 4 | 8 | 1 | 1 | 1 | 1 | 1 | 1 |
| Q9DBY8 | NVL_MOUSE | 5 | 1 | 8 | 1 | 0 | 5 | 1 | 1 | 1 | 1 | 0 | 1 |
| Q02248 | CTNB1_MOUSE | 1 | 5 | 7 | 1 | 0 | 5 | 1 | 1 | 1 | 1 | 0.99 | 1 |
| Q9JM76 | ARPC3_MOUSE | 2 | 8 | 5 | 0 | 6 | 4 | 1 | 1 | 1 | 0.87 | 1 | 1 |
| Q91VC3 | IF4A3_MOUSE | 6 | 4 | 5 | 7 | 0 | 0 | 1 | 1 | 1 | 1 | 0.91 | 0 |
| Q8R5K4 | NOL6_MOUSE | 2 | 2 | 3 | 2 | 0 | 2 | 1 | 1 | 1 | 1 | 0 | 1 |
| Q8JZX4 | SPF45_MOUSE | 9 | 2 | 4 | 5 | 1 | 0 | 1 | 1 | 1 | 1 | 1 | 0 |
| Q9DC51 | GNAI3_MOUSE | 2 | 7 | 4 | 4 | 4 | 2 | 1 | 1 | 1 | 1 | 1 | 1 |
| Q3TLH4 | PRC2C_MOUSE | 24 | 5 | 14 | 14 | 4 | 7 | 1 | 1 | 1 | 1 | 1 | 1 |
| E9PVC5 | E9PVC5_MOUSE | 28 | 12 | 12 | 17 | 9 | 3 | 1 | 1 | 1 | 1 | 1 | 1 |
| F7CVJ5 | F7CVJ5_MOUSE | 2 | 0 | 0 | 1 | 0 | 0 | 1 | 1 | 0.88 | 0.99 | 1 | 0 |
| Q3UEB3 | PUF60_MOUSE | 19 | 9 | 14 | 7 | 8 | 8 | 1 | 1 | 1 | 1 | 1 | 1 |
| P62827 | RAN_MOUSE | 1 | 1 | 2 | 0 | 1 | 0 | 1 | 0.98 | 1 | 0.81 | 0.98 | 0 |
| Q6PGG6 | GNL3L_MOUSE | 4 | 2 | 3 | 4 | 0 | 0 | 1 | 1 | 1 | 1 | 0.57 | 0 |
| Q9QXS1 | PLEC_MOUSE | 432 | 338 | 603 | 294 | 253 | 320 | 1 | 1 | 1 | 1 | 1 | 1 |
| Q9CQM8 | Q9CQM8_MOUSE | 24 | 13 | 24 | 21 | 0 | 14 | 1 | 1 | 1 | 1 | 1 | 1 |
| P46935 | NEDD4_MOUSE | 1 | 2 | 7 | 4 | 0 | 1 | 1 | 1 | 1 | 1 | 0.41 | 0.85 |
| P67984 | RL22_MOUSE | 2 | 8 | 1 | 2 | 8 | 0 | 1 | 1 | 0.91 | 0.99 | 1 | 0 |
| D3YZC9 | D3YZC9_MOUSE | 2 | 2 | 1 | 2 | 0 | 0 | 1 | 1 | 0.94 | 1 | 0 | 0 |
| Q9EPA7 | NMNA1_MOUSE | 3 | 2 | 1 | 1 | 0 | 2 | 1 | 1 | 1 | 0.99 | 0.36 | 1 |

| | | | | | | | | | | | | | |
|---|---|---|---|---|---|---|---|---|---|---|---|---|---|
| Q5XG71 | UTP20_MOUSE | 27 | 6 | 8 | 20 | 1 | 3 | 1 | 1 | 1 | 1 | 0.92 | 1 |
| P62267 | RS23_MOUSE | 1 | 10 | 1 | 1 | 8 | 0 | 1 | 1 | 0.99 | 1 | 1 | 0 |
| Q9EP89 | LACTB_MOUSE | 5 | 6 | 5 | 3 | 1 | 2 | 1 | 1 | 1 | 1 | 1 | 1 |
| Q9JHU4 | DYHC1_MOUSE | 2 | 0 | 2 | 0 | 1 | 0 | 1 | 1 | 1 | 0.35 | 0.8 | 0 |
| Q9DBG7 | SRPR_MOUSE | 3 | 1 | 1 | 0 | 1 | 0 | 1 | 0.98 | 0.95 | 0.09 | 0.99 | 0 |
| O54724 | PTRF_MOUSE | 17 | 12 | 10 | 12 | 10 | 4 | 1 | 1 | 1 | 1 | 1 | 1 |
| Q6P4T2 | U520_MOUSE | 50 | 29 | 49 | 40 | 16 | 15 | 1 | 1 | 1 | 1 | 1 | 1 |
| Q8BH64 | EHD2_MOUSE | 1 | 3 | 1 | 0 | 1 | 0 | 1 | 1 | 0.95 | 0.16 | 0.85 | 0 |
| Q9EQ61 | PESC_MOUSE | 3 | 0 | 1 | 4 | 0 | 0 | 1 | 0.97 | 0.89 | 1 | 0 | 0 |
| Q8CGZ0 | CHERP_MOUSE | 6 | 2 | 2 | 3 | 0 | 0 | 1 | 1 | 1 | 1 | 0 | 0 |
| E9QKG6 | E9QKG6_MOUSE | 18 | 2 | 5 | 11 | 1 | 0 | 1 | 1 | 1 | 1 | 0.58 | 0 |
| H3BKH9 | H3BKH9_MOUSE | 4 | 3 | 10 | 0 | 0 | 3 | 1 | 1 | 1 | 0.99 | 0 | 1 |
| Q924Z5 | TRAM2_MOUSE | 1 | 2 | 1 | 0 | 1 | 1 | 0.93 | 1 | 1 | 0.27 | 0.99 | 0.97 |
| P32233 | DRG1_MOUSE | 6 | 3 | 2 | 7 | 1 | 1 | 1 | 1 | 1 | 1 | 1 | 0.97 |
| Q61235 | SNTB2_MOUSE | 2 | 6 | 1 | 3 | 3 | 1 | 1 | 1 | 0.83 | 1 | 1 | 0.56 |
| P47963 | RL13_MOUSE | 29 | 21 | 24 | 30 | 16 | 23 | 1 | 1 | 1 | 1 | 1 | 1 |
| Q99MR8 | MCCA_MOUSE | 9 | 24 | 46 | 14 | 15 | 35 | 1 | 1 | 1 | 1 | 1 | 1 |
| Q9QZD9 | EIF3I_MOUSE | 4 | 6 | 1 | 4 | 5 | 1 | 1 | 1 | 0.97 | 1 | 1 | 0.83 |
| Q8K301 | DDX52_MOUSE | 6 | 6 | 2 | 6 | 3 | 0 | 1 | 1 | 1 | 1 | 1 | 0 |
| Q61545 | EWS_MOUSE | 4 | 2 | 3 | 3 | 1 | 0 | 1 | 1 | 1 | 1 | 0.99 | 0 |
| O70318 | E41L2_MOUSE | 2 | 5 | 9 | 3 | 1 | 4 | 1 | 1 | 1 | 1 | 1 | 1 |
| Q6ZQ08 | CNOT1_MOUSE | 15 | 5 | 8 | 17 | 2 | 4 | 1 | 1 | 1 | 1 | 1 | 1 |
| E9Q616 | E9Q616_MOUSE | 10 | 2 | 2 | 2 | 0 | 2 | 1 | 1 | 1 | 1 | 0.77 | 1 |
| Q9R0E2 | PLOD1_MOUSE | 3 | 3 | 2 | 4 | 0 | 1 | 1 | 1 | 1 | 1 | 0.96 | 0.97 |
| Q9QZE5 | COPG1_MOUSE | 2 | 4 | 6 | 2 | 0 | 0 | 1 | 1 | 1 | 1 | 0.39 | 0.32 |
| Q9D8L3 | Q9D8L3_MOUSE | 1 | 3 | 2 | 0 | 1 | 0 | 0.92 | 1 | 1 | 0.36 | 1 | 0.64 |
| O89079 | COPE_MOUSE | 1 | 2 | 1 | 1 | 1 | 0 | 0.86 | 1 | 0.89 | 1 | 0.99 | 0 |
| H7BX26 | H7BX26_MOUSE | 5 | 3 | 2 | 0 | 2 | 0 | 1 | 1 | 1 | 0 | 0.99 | 0.17 |
| Q91ZX7 | LRP1_MOUSE | 0 | 2 | 4 | 1 | 1 | 0 | 0.91 | 1 | 1 | 1 | 1 | 0 |
| P28301 | LYOX_MOUSE | 3 | 2 | 3 | 3 | 4 | 0 | 1 | 1 | 1 | 1 | 1 | 0 |
| P62900 | RL31_MOUSE | 2 | 10 | 3 | 1 | 9 | 1 | 0.95 | 1 | 1 | 1 | 1 | 0.99 |
| P68369 | TBA1A_MOUSE | 15 | 14 | 13 | 12 | 12 | 5 | 1 | 1 | 1 | 1 | 1 | 1 |
| P58871 | TB182_MOUSE | 32 | 5 | 9 | 28 | 8 | 1 | 1 | 1 | 1 | 1 | 1 | 0.93 |
| P97479 | MYO7A_MOUSE | 16 | 7 | 39 | 17 | 9 | 18 | 1 | 1 | 1 | 1 | 1 | 1 |
| O35465 | FKBP8_MOUSE | 1 | 2 | 2 | 3 | 2 | 0 | 1 | 1 | 1 | 1 | 1 | 0 |
| Q9D1C9 | RRP7A_MOUSE | 0 | 2 | 1 | 1 | 0 | 0 | 0.99 | 1 | 0.89 | 1 | 0.56 | 0 |
| Q922F4 | TBB6_MOUSE | 6 | 2 | 3 | 2 | 0 | 0 | 1 | 1 | 1 | 0.98 | 0 | 0 |
| P39061 | COIA1_MOUSE | 7 | 2 | 3 | 4 | 2 | 0 | 1 | 1 | 1 | 1 | 1 | 0 |
| P10922 | H10_MOUSE | 7 | 4 | 4 | 5 | 3 | 0 | 1 | 1 | 1 | 1 | 1 | 0 |
| Q8QZY1 | EIF3L_MOUSE | 3 | 4 | 4 | 6 | 3 | 0 | 1 | 1 | 1 | 1 | 1 | 0 |
| Q922Q1 | MOSC2_MOUSE | 1 | 1 | 4 | 2 | 0 | 0 | 1 | 1 | 1 | 1 | 0.87 | 0 |
| Q810A7 | DDX42_MOUSE | 5 | 1 | 3 | 5 | 1 | 2 | 1 | 1 | 1 | 1 | 0.62 | 1 |
| P21956 | MFGM_MOUSE | 9 | 14 | 22 | 8 | 14 | 11 | 1 | 1 | 1 | 1 | 1 | 1 |
| Q9CQE8 | CN166_MOUSE | 8 | 6 | 6 | 11 | 6 | 3 | 1 | 1 | 1 | 1 | 1 | 1 |
| Q9CWF2 | TBB2B_MOUSE | 3 | 0 | 1 | 0 | 0 | 0 | 1 | 0.98 | 0.94 | 0.59 | 0 | 0 |
| P62751 | RL23A_MOUSE | 13 | 14 | 16 | 7 | 14 | 12 | 1 | 1 | 1 | 1 | 1 | 1 |
| P11499 | HS90B_MOUSE | 1 | 5 | 6 | 3 | 9 | 6 | 1 | 1 | 1 | 1 | 1 | 1 |
| Q91ZN5 | S35B2_MOUSE | 1 | 1 | 3 | 1 | 1 | 1 | 0.96 | 0.99 | 1 | 0.99 | 0.98 | 1 |
| Q9WVJ9 | FBLN4_MOUSE | 2 | 0 | 1 | 1 | 1 | 0 | 1 | 0.98 | 0.91 | 0.95 | 0.93 | 0 |
| P19973 | LSP1_MOUSE | 7 | 3 | 7 | 6 | 0 | 2 | 1 | 1 | 1 | 1 | 0.54 | 1 |
| Q8R3N1 | NOP14_MOUSE | 4 | 2 | 2 | 2 | 0 | 0 | 1 | 1 | 0.9 | 1 | 0 | 0 |
| Q9QXK7 | CPSF3_MOUSE | 0 | 2 | 1 | 2 | 0 | 0 | 0.98 | 1 | 0.8 | 1 | 0 | 0 |
| Q5XJY5 | COPD_MOUSE | 3 | 1 | 1 | 2 | 1 | 0 | 1 | 0.97 | 0.95 | 1 | 0.99 | 0 |
| Q4VBE8 | WDR18_MOUSE | 4 | 2 | 6 | 4 | 2 | 5 | 1 | 1 | 1 | 1 | 1 | 1 |
| P63325 | RS10_MOUSE | 2 | 7 | 6 | 0 | 8 | 1 | 1 | 1 | 1 | 1 | 1 | 1 |
| P16858 | G3P_MOUSE | 1 | 4 | 2 | 0 | 3 | 1 | 0.99 | 1 | 1 | 0.84 | 1 | 0.72 |
| P62259 | 1433E_MOUSE | 0 | 3 | 3 | 0 | 0 | 2 | 0.97 | 1 | 1 | 0 | 0.97 | 1 |
| Q61554 | A2AQ53_MOUSE | 3 | 0 | 1 | 3 | 0 | 0 | 1 | 1 | 0.99 | 1 | 0 | 1 |
| Q8BTH8 | KC1G1_MOUSE | 1 | 2 | 2 | 1 | 0 | 0 | 1 | 1 | 1 | 1 | 0.82 | 0 |
| Q9R0G7 | ZEB2_MOUSE | 7 | 1 | 2 | 1 | 0 | 0 | 1 | 1 | 0.95 | 1 | 0 | 0 |
| Q8CBY8 | DCTN4_MOUSE | 1 | 3 | 3 | 0 | 1 | 0 | 0.99 | 1 | 1 | 0.86 | 0.98 | 0 |
| P43275 | H11_MOUSE | 6 | 4 | 3 | 6 | 1 | 3 | 1 | 1 | 1 | 1 | 0.71 | 1 |
| Q922J9 | FACR1_MOUSE | 6 | 5 | 4 | 7 | 0 | 1 | 1 | 1 | 1 | 1 | 0.97 | 0.99 |

| | | | | | | | | | | | | | |
|---|---|---|---|---|---|---|---|---|---|---|---|---|---|
| Q8CCF0 | PRP31_MOUSE | 6 | 7 | 4 | 6 | 3 | 2 | 1 | 1 | 1 | 1 | 1 | 1 |
| A2AWN8 | A2AWN8_MOUSE | 5 | 3 | 7 | 8 | 3 | 0 | 1 | 1 | 1 | 1 | 1 | 0.96 |
| B9EHJ3 | B9EHJ3_MOUSE | 38 | 1 | 22 | 31 | 10 | 11 | 1 | 1 | 1 | 1 | 1 | 1 |
| P60122 | RUVB1_MOUSE | 4 | 4 | 2 | 3 | 4 | 0 | 1 | 1 | 1 | 1 | 1 | 0 |
| P62983 | RS27A_MOUSE | 2 | 6 | 2 | 0 | 5 | 0 | 1 | 1 | 1 | 0.68 | 1 | 0.51 |
| Q8VHX6 | FLNC_MOUSE | 31 | 12 | 6 | 23 | 4 | 1 | 1 | 1 | 1 | 1 | 1 | 1 |
| Q8C4U8 | Q8C4U8_MOUSE | 1 | 1 | 3 | 3 | 2 | 0 | 1 | 0.99 | 1 | 1 | 1 | 0 |
| Q8VE73 | CUL7_MOUSE | 6 | 0 | 5 | 3 | 3 | 0 | 1 | 0.94 | 1 | 1 | 1 | 0 |
| Q8BPM0 | DAAM1_MOUSE | 1 | 3 | 2 | 2 | 0 | 0 | 1 | 1 | 1 | 1 | 0 | 0.13 |
| E9PYQ1 | E9PYQ1_MOUSE | 3 | 1 | 1 | 1 | 0 | 0 | 1 | 0.98 | 0.96 | 0.99 | 0 | 0 |
| O54734 | OST48_MOUSE | 10 | 4 | 12 | 7 | 6 | 3 | 1 | 1 | 1 | 1 | 1 | 1 |
| Q9Z110 | P5CS_MOUSE | 1 | 3 | 1 | 2 | 0 | 1 | 1 | 1 | 1 | 1 | 0.98 | 0.94 |
| E9Q6H8 | E9Q6H8_MOUSE | 2 | 0 | 4 | 2 | 0 | 0 | 1 | 0.99 | 1 | 1 | 0 | 0 |
| Q01320 | TOP2A_MOUSE | 6 | 1 | 1 | 10 | 1 | 0 | 1 | 1 | 0.86 | 1 | 0.91 | 0 |
| P63037 | DNJA1_MOUSE | 2 | 2 | 2 | 3 | 0 | 0 | 1 | 1 | 1 | 1 | 0.98 | 0 |
| P19324 | SERPH_MOUSE | 9 | 8 | 12 | 7 | 8 | 5 | 1 | 1 | 1 | 1 | 1 | 1 |
| P60229 | EIF3E_MOUSE | 4 | 2 | 1 | 3 | 3 | 0 | 1 | 1 | 0.91 | 1 | 1 | 0 |
| Q91V04 | TRAM1_MOUSE | 1 | 4 | 3 | 2 | 0 | 0 | 1 | 1 | 1 | 1 | 0.96 | 0.56 |
| Q07646 | MEST_MOUSE | 1 | 1 | 1 | 1 | 0 | 1 | 0.98 | 0.99 | 1 | 0.78 | 0.43 | 0.86 |
| Q60847 | COCA1_MOUSE | 85 | 83 | 157 | 76 | 72 | 100 | 1 | 1 | 1 | 1 | 1 | 1 |
| P54116 | STOM_MOUSE | 2 | 2 | 1 | 0 | 3 | 0 | 1 | 1 | 0.95 | 0.95 | 1 | 0 |
| Q8BXQ2 | PIGT_MOUSE | 1 | 2 | 1 | 2 | 3 | 1 | 0.98 | 1 | 0.95 | 1 | 1 | 0.96 |
| Q0P688 | Q0P688_MOUSE | 2 | 1 | 2 | 2 | 1 | 0 | 1 | 1 | 1 | 0.99 | 0.99 | 0 |
| P61982 | 1433G_MOUSE | 1 | 2 | 3 | 0 | 1 | 1 | 1 | 1 | 1 | 0.3 | 1 | 0.99 |
| Q8CGB3 | UACA_MOUSE | 2 | 2 | 2 | 1 | 0 | 0 | 1 | 1 | 1 | 0.86 | 0.38 | 0 |
| Q9D880 | TIM50_MOUSE | 1 | 2 | 2 | 3 | 2 | 0 | 1 | 1 | 1 | 1 | 1 | 0 |
| Q99MN9 | PCCB_MOUSE | 1 | 5 | 3 | 1 | 6 | 5 | 0.99 | 1 | 1 | 1 | 1 | 1 |
| Q08943 | SSRP1_MOUSE | 4 | 9 | 8 | 5 | 6 | 1 | 1 | 1 | 1 | 1 | 1 | 1 |
| Q99K28 | ARFG2_MOUSE | 1 | 1 | 2 | 2 | 4 | 1 | 1 | 0.92 | 1 | 1 | 1 | 0.97 |
| Q62036 | AZI1_MOUSE | 2 | 3 | 11 | 4 | 0 | 3 | 1 | 1 | 1 | 1 | 0 | 1 |
| Q60972 | RBBP4_MOUSE | 1 | 1 | 1 | 2 | 0 | 0 | 1 | 0.94 | 0.98 | 1 | 0 | 0 |
| O35382 | EXOC4_MOUSE | 1 | 2 | 4 | 1 | 0 | 2 | 0.9 | 1 | 1 | 0.5 | 0.31 | 1 |
| A2APB8 | TPX2_MOUSE | 2 | 1 | 1 | 3 | 0 | 0 | 1 | 1 | 0.8 | 1 | 0.44 | 0 |
| Q9CR68 | UCRI_MOUSE | 2 | 1 | 2 | 2 | 0 | 1 | 1 | 0.99 | 1 | 1 | 0 | 0.82 |
| Q61245 | COBA1_MOUSE | 2 | 2 | 9 | 2 | 7 | 1 | 1 | 1 | 1 | 1 | 1 | 0.99 |
| P70333 | HNRH2_MOUSE | 3 | 1 | 2 | 3 | 0 | 0 | 1 | 0.91 | 1 | 1 | 0 | 0 |
| Q8C0T5 | SI1L1_MOUSE | 11 | 2 | 9 | 13 | 0 | 0 | 1 | 1 | 1 | 1 | 0 | 0 |
| Q8BGA9 | OXA1L_MOUSE | 2 | 1 | 1 | 2 | 0 | 0 | 1 | 1 | 1 | 1 | 0 | 0 |
| Q91WB7 | UBTD1_MOUSE | 1 | 1 | 2 | 1 | 1 | 0 | 1 | 1 | 1 | 0.99 | 0.99 | 0 |
| Q3ULD5 | MCCB_MOUSE | 10 | 19 | 26 | 8 | 16 | 36 | 1 | 1 | 1 | 1 | 1 | 1 |
| P53690 | MMP14_MOUSE | 1 | 3 | 7 | 3 | 7 | 2 | 1 | 1 | 1 | 1 | 1 | 1 |
| E9PZY8 | E9PZY8_MOUSE | 8 | 3 | 5 | 6 | 2 | 0 | 1 | 1 | 1 | 1 | 1 | 0 |
| P28653 | PGS1_MOUSE | 8 | 8 | 9 | 5 | 7 | 4 | 1 | 1 | 1 | 1 | 1 | 1 |
| E9Q784 | E9Q784_MOUSE | 5 | 0 | 2 | 4 | 0 | 0 | 1 | 1 | 1 | 1 | 0 | 0 |
| A2ANY6 | A2ANY6_MOUSE | 15 | 0 | 2 | 14 | 0 | 0 | 1 | 1 | 1 | 1 | 0.53 | 0 |
| Q60634 | FLOT2_MOUSE | 3 | 1 | 1 | 1 | 2 | 0 | 1 | 1 | 1 | 0.97 | 1 | 0 |
| Q6PFD9 | NUP98_MOUSE | 1 | 1 | 3 | 2 | 0 | 0 | 1 | 1 | 1 | 1 | 0.28 | 0 |
| Q91W39 | NCOA5_MOUSE | 3 | 1 | 4 | 4 | 0 | 0 | 1 | 1 | 1 | 1 | 0.93 | 0 |
| P42208 | SEPT2_MOUSE | 2 | 5 | 3 | 2 | 0 | 1 | 1 | 1 | 1 | 1 | 0.76 | 0.83 |
| Q5SWU9 | ACACA_MOUSE | 13 | 4 | 42 | 9 | 10 | 47 | 1 | 1 | 1 | 1 | 1 | 1 |
| Q3UV17 | K22O_MOUSE | 1 | 1 | 1 | 1 | 0 | 2 | 0.98 | 0.99 | 0.98 | 0.95 | 0.95 | 1 |
| Q8BTW3 | EXOS6_MOUSE | 1 | 1 | 1 | 1 | 2 | 0 | 0.98 | 1 | 0.98 | 0.95 | 1 | 0 |
| Q9CQU1 | MFAP1_MOUSE | 3 | 2 | 2 | 0 | 1 | 0 | 1 | 1 | 1 | 1 | 0.99 | 0 |
| P27546 | MAP4_MOUSE | 3 | 1 | 2 | 2 | 2 | 1 | 1 | 1 | 0.98 | 0.99 | 1 | 0.96 |
| E9Q2I4 | E9Q2I4_MOUSE | 1 | 0 | 0 | 2 | 0 | 0 | 1 | 1 | 1 | 1 | 0 | 0 |
| Q02257 | PLAK_MOUSE | 1 | 1 | 3 | 1 | 0 | 6 | 1 | 1 | 1 | 0.99 | 0.55 | 1 |
| O08917 | FLOT1_MOUSE | 2 | 3 | 3 | 0 | 3 | 0 | 1 | 1 | 1 | 0 | 1 | 0 |
| E9Q6R7 | E9Q6R7_MOUSE | 34 | 1 | 1 | 30 | 28 | 1 | 1 | 1 | 0.99 | 1 | 1 | 1 |
| Q91VM5 | RMXL1_MOUSE | 12 | 5 | 7 | 9 | 1 | 3 | 1 | 1 | 1 | 1 | 1 | 1 |
| A2BDX3 | MOCS3_MOUSE | 1 | 0 | 1 | 0 | 0 | 1 | 0.9 | 0.88 | 0.92 | 0.36 | 0 | 0.9 |
| Q5U405 | TMPSD_MOUSE | 1 | 1 | 1 | 0 | 1 | 1 | 0.9 | 0.8 | 0.88 | 0.64 | 0.99 | 0.99 |
| E9Q557 | DESP_MOUSE | 1 | 0 | 1 | 0 | 1 | 6 | 1 | 0.92 | 0.95 | 0.89 | 0.79 | 1 |
| Q9CU62 | SMC1A_MOUSE | 1 | 5 | 2 | 2 | 3 | 0 | 0.05 | 0.93 | 1 | 0.98 | 0.55 | 0 |

| | | | | | | | | | | | | | |
|---|---|---|---|---|---|---|---|---|---|---|---|---|---|
| Q61164 | CTCF_MOUSE | 1 | 1 | 4 | 0 | 0 | 0 | 0 | 0.45 | 1 | 0 | 0 | 0 |
| Q04750 | TOP1_MOUSE | 7 | 18 | 9 | 7 | 0 | 7 | 1 | 1 | 1 | 1 | 0.57 | 1 |
| Q8K0H5 | TAF10_MOUSE | 1 | 1 | 1 | 0 | 1 | 0 | 0.92 | 0.99 | 0 | 0 | 0.86 | 0 |
| Q01320 | TOP2A_MOUSE | 17 | 4 | 1 | 14 | 2 | 0 | 1 | 1 | 0.86 | 1 | 0.91 | 0 |
| Q9CW03 | SMC3_MOUSE | 3 | 6 | 1 | 3 | 3 | 0 | 1 | 0.96 | 0 | 0.99 | 0.97 | 0 |
| A0A2I3BPN6 | TAF4_MOUSE | 1 | 1 | 1 | 0 | 0 | 0 | 0.6 | 0.75 | 0 | 0 | 0 | 0 |
| Q62311 | TAF6_MOUSE | 1 | 1 | 1 | 1 | 0 | 0 | 0 | 0.15 | 0 | 0.17 | 0 | 0 |
