## Supplementary Table S5 for "ERCC1-XPF Interacts with Topoisomerase IIβ to Facilitate the Repair of Activity-induced DNA Breaks"

**Supplementary Table S5. List of bXPF ChIP-Seq peaks with DSBs in untreated and tRA treated MEFs.**

| PeakID | Strand | Peak Score | Annotation | Gene Name | Count (DSBs) | Sample ID |
|---|---|---|---|---|---|---|
| 1-506 | + | 554 | non-coding (NR_002847, exon 1 of 1) | Mascrna | 50 | Untreated (DSB peaks) |
| 1-479 | + | 614 | promoter-TSS (NM_009860) | Cdc25c | 32 | Untreated (DSB peaks) |
| 1-430 | + | 606 | promoter-TSS (NM_001172114) | Traf7 | 31 | Untreated (DSB peaks) |
| 1-673 | + | 863 | promoter-TSS (NR_001460) | Rmrp | 27 | Untreated (DSB peaks) |
| 1-260 | + | 558 | Intergenic | BC005537 | 23 | Untreated (DSB peaks) |
| 1-918 | + | 674 | Intergenic | Chst15 | 23 | Untreated (DSB peaks) |
| 1-940 | + | 550 | promoter-TSS (NM_001007570) | Slc25a42 | 21 | Untreated (DSB peaks) |
| 1-196 | + | 543 | promoter-TSS (NM_026071) | Slc25a19 | 20 | Untreated (DSB peaks) |
| 1-630 | + | 635 | promoter-TSS (NR_045997) | 1700113A16Rik | 18 | Untreated (DSB peaks) |
| 1-723 | + | 563 | intron (NM_001081557, intron 5 of 23) | Gm13090 | 18 | Untreated (DSB peaks) |
| 1-809 | + | 688 | Intergenic | Rn4.5s | 18 | Untreated (DSB peaks) |
| 1-881 | + | 722 | Intergenic | Akap13 | 18 | Untreated (DSB peaks) |
| 1-232 | + | 748 | Intergenic | Lrrc74a | 18 | Untreated (DSB peaks) |
| 1-917 | + | 675 | Intergenic | Chst15 | 18 | Untreated (DSB peaks) |
| 1-524 | + | 561 | promoter-TSS (NM_146114) | Dclre1c | 17 | Untreated (DSB peaks) |
| 1-189 | + | 576 | promoter-TSS (NM_011623) | Top2a | 17 | Untreated (DSB peaks) |
| 1-520 | + | 569 | Intergenic | Sfxn3 | 17 | Untreated (DSB peaks) |
| 1-883 | + | 617 | intron (NM_016721, intron 2 of 37) | Iqgap1 | 16 | Untreated (DSB peaks) |
| 1-810 | + | 630 | Intergenic | Mir704 | 16 | Untreated (DSB peaks) |
| 1-613 | + | 851 | Intergenic | Pcdh18 | 16 | Untreated (DSB peaks) |
| 1-1010 | + | 670 | Intergenic | Ncam1 | 16 | Untreated (DSB peaks) |
| 1-152 | + | 581 | promoter-TSS (NM_013716) | G3bp1 | 15 | Untreated (DSB peaks) |
| 43466 | + | 559 | promoter-TSS (NM_009676) | Aox1 | 15 | Untreated (DSB peaks) |
| 1-1027 | + | 572 | promoter-TSS (NM_019797) | Trip4 | 15 | Untreated (DSB peaks) |
| 1-190 | + | 555 | promoter-TSS (NM_011486) | Stat3 | 15 | Untreated (DSB peaks) |
| 1-912 | + | 585 | Intergenic | Prr14 | 15 | Untreated (DSB peaks) |
| 1-535 | + | 596 | intron (NM_011040, intron 2 of 11) | Pax8 | 14 | Untreated (DSB peaks) |
| 1-870 | + | 584 | promoter-TSS (NM_026042) | Paf1 | 14 | Untreated (DSB peaks) |
| 10959 | + | 617 | promoter-TSS (NM_001310668) | Bcs1l | 14 | Untreated (DSB peaks) |
| 1-230 | + | 635 | promoter-TSS (NR_073563) | Numb | 14 | Untreated (DSB peaks) |
| 1-565 | + | 664 | 5' UTR (NM_001177785, exon 1 of 16) | Cd44 | 14 | Untreated (DSB peaks) |
| 21916 | + | 602 | promoter-TSS (NR_002870) | Dnm3os | 14 | Untreated (DSB peaks) |
| 1-675 | + | 641 | Intergenic | Sec61b | 14 | Untreated (DSB peaks) |
| 14611 | + | 625 | Intergenic | Col6a3 | 14 | Untreated (DSB peaks) |
| 1-123 | + | 602 | Intergenic | Plxnc1 | 14 | Untreated (DSB peaks) |
| 1-764 | + | 593 | Intergenic | E130006D01Rik | 14 | Untreated (DSB peaks) |
| 1-356 | + | 593 | promoter-TSS (NM_027435) | Atad2 | 13 | Untreated (DSB peaks) |
| 1-374 | + | 726 | intron (NM_029787, intron 1 of 8) | Cyb5r3 | 13 | Untreated (DSB peaks) |
| 1-188 | + | 547 | 5' UTR (NM_145434, exon 1 of 8) | Nr1d1 | 13 | Untreated (DSB peaks) |
| 1-767 | + | 611 | intron (NM_001159941, intron 2 of 6) | Kctd10 | 13 | Untreated (DSB peaks) |
| 1-769 | + | 642 | promoter-TSS (NM_001040398) | Setd1b | 13 | Untreated (DSB peaks) |
| 1-571 | + | 610 | promoter-TSS (NM_178404) | Zc3h6 | 13 | Untreated (DSB peaks) |
| 1-385 | + | 582 | Intergenic | Ercc4 | 13 | Untreated (DSB peaks) |
| 1-600 | + | 548 | Intergenic | Tbl1xr1 | 13 | Untreated (DSB peaks) |

| | | | | | | |
|---|---|---|---|---|---|---|
| 1-252 | + | 543 | Intergenic | Prss16 | 13 | Untreated (DSB peaks) |
| 1-928 | + | 562 | Intergenic | Fgfr1 | 13 | Untreated (DSB peaks) |
| 1-671 | + | 653 | Intergenic | Pnrc1 | 13 | Untreated (DSB peaks) |
| 1-1030 | + | 687 | Intergenic | Lactb | 13 | Untreated (DSB peaks) |
| 1-897 | + | 824 | Intergenic | Serpinh1 | 13 | Untreated (DSB peaks) |
| 1-472 | + | 634 | Intergenic | Gm26682 | 12 | Untreated (DSB peaks) |
| 1-504 | + | 542 | intron (NM_019869, intron 1 of 2) | Rbm14 | 12 | Untreated (DSB peaks) |
| 1-863 | + | 559 | promoter-TSS (NM_016871) | Tomm40 | 12 | Untreated (DSB peaks) |
| 1-482 | + | 544 | promoter-TSS (NM_033581) | Pcdhgc3 | 12 | Untreated (DSB peaks) |
| 1-183 | + | 585 | intron (NM_001361708, intron 4 of 9) | Car10 | 12 | Untreated (DSB peaks) |
| 1-510 | + | 645 | Intergenic | Ahnak | 12 | Untreated (DSB peaks) |
| 1-185 | + | 1000 | 5' UTR (NM_008704, exon 1 of 5) | Nme1 | 12 | Untreated (DSB peaks) |
| 1-638 | + | 582 | 5' UTR (NM_001039488, exon 1 of 4) | Lrif1 | 12 | Untreated (DSB peaks) |
| 1-387 | + | 568 | Intergenic | Abcc1 | 12 | Untreated (DSB peaks) |
| 1-595 | + | 741 | Intergenic | Trim55 | 12 | Untreated (DSB peaks) |
| 1-447 | + | 541 | Intergenic | 2300002M23Rik | 12 | Untreated (DSB peaks) |
| 1-451 | + | 561 | Intergenic | Cdc5l | 12 | Untreated (DSB peaks) |
| 1-612 | + | 853 | Intergenic | Pcdh18 | 12 | Untreated (DSB peaks) |
| 1-357 | + | 559 | Intergenic | Gm19510 | 12 | Untreated (DSB peaks) |
| 1-954 | + | 592 | Intergenic | 1700011L22Rik | 12 | Untreated (DSB peaks) |
| 1-822 | + | 612 | Intergenic | Tpra1 | 12 | Untreated (DSB peaks) |
| 1-395 | + | 822 | intron (NM_178665, intron 3 of 12) | Morf4l1-ps1 | 11 | Untreated (DSB peaks) |
| 1-456 | + | 573 | intron (NM_172964, intron 1 of 17) | Arhgap28 | 11 | Untreated (DSB peaks) |
| 1-179 | + | 690 | intron (NM_001356531, intron 5 of 11) | Vmp1 | 11 | Untreated (DSB peaks) |
| 1-197 | + | 694 | intron (NM_011594, intron 1 of 4) | Cep295nl | 11 | Untreated (DSB peaks) |
| 1-974 | + | 813 | promoter-TSS (NM_178856) | Gins2 | 11 | Untreated (DSB peaks) |
| 1-850 | + | 733 | intron (NM_175414, intron 1 of 8) | Tspan9 | 11 | Untreated (DSB peaks) |
| 1-783 | + | 709 | promoter-TSS (NM_010717) | Limk1 | 11 | Untreated (DSB peaks) |
| 19725 | + | 797 | intron (NM_016846, intron 5 of 17) | Rgl1 | 11 | Untreated (DSB peaks) |
| 1-386 | + | 549 | Intergenic | Mir193b | 11 | Untreated (DSB peaks) |
| 1-583 | + | 703 | intron (NM_001039390, intron 2 of 4) | Pkig | 11 | Untreated (DSB peaks) |
| 1-259 | + | 663 | Intergenic | Hist1h4h | 11 | Untreated (DSB peaks) |
| 1-1002 | + | 551 | Intergenic | Sorl1 | 11 | Untreated (DSB peaks) |
| 1-320 | + | 876 | Intergenic | Rnase4 | 11 | Untreated (DSB peaks) |
| 1-1023 | + | 638 | Intergenic | Smad6 | 11 | Untreated (DSB peaks) |
| 1-1029 | + | 686 | Intergenic | Lactb | 11 | Untreated (DSB peaks) |
| 1-708 | + | 569 | Intergenic | Thrap3 | 11 | Untreated (DSB peaks) |
| 1-787 | + | 601 | Intergenic | Foxk1 | 11 | Untreated (DSB peaks) |
| 1-341 | + | 803 | Intergenic | Gm19276 | 10 | Untreated (DSB peaks) |
| 1-381 | + | 603 | promoter-TSS (NM_145359) | Ubald1 | 10 | Untreated (DSB peaks) |
| 1-990 | + | 553 | promoter-TSS (NM_001004190) | Zfp560 | 10 | Untreated (DSB peaks) |
| 1-433 | + | 681 | promoter-TSS (NM_001285982) | Fam173a | 10 | Untreated (DSB peaks) |
| 1-867 | + | 556 | intron (NM_133210, intron 1 of 1) | Sertad3 | 10 | Untreated (DSB peaks) |
| 1-438 | + | 577 | promoter-TSS (NM_007669) | Cdkn1a | 10 | Untreated (DSB peaks) |
| 1-351 | + | 713 | intron (NM_080635, intron 3 of 7) | Eif3h | 10 | Untreated (DSB peaks) |
| 1-408 | + | 614 | intron (NM_019631, intron 1 of 5) | Tmem45a | 10 | Untreated (DSB peaks) |
| 1-744 | + | 580 | intron (NM_001310608, intron 1 of 3) | Pcdh7 | 10 | Untreated (DSB peaks) |

| | | | | | | |
|---|---|---|---|---|---|---|
| 1-355 | + | 593 | promoter-TSS (NM_027435) | Atad2 | 10 | Untreated (DSB peaks) |
| 1-747 | + | 1000 | intron (NM_013689, intron 1 of 17) | Tec | 10 | Untreated (DSB peaks) |
| 1-372 | + | 558 | intron (NM_001358772, intron 2 of 14) | Rbfox2 | 10 | Untreated (DSB peaks) |
| 1-171 | + | 1000 | promoter-TSS (NM_011333) | Ccl2 | 10 | Untreated (DSB peaks) |
| 1-172 | + | 1000 | exon (NM_011333, exon 1 of 3) | Ccl2 | 10 | Untreated (DSB peaks) |
| 1-825 | + | 545 | intron (NR_015530, intron 1 of 2) | 9530026P05Rik | 10 | Untreated (DSB peaks) |
| 1-635 | + | 591 | promoter-TSS (NM_025875) | Rbm8a | 10 | Untreated (DSB peaks) |
| 1-238 | + | 543 | intron (NM_172500, intron 1 of 16) | Syne3 | 10 | Untreated (DSB peaks) |
| 1-1045 | + | 1000 | 5' UTR (NM_001324534, exon 1 of 3) | Rpl29 | 10 | Untreated (DSB peaks) |
| 1-390 | + | 604 | Intergenic | Scarf2 | 10 | Untreated (DSB peaks) |
| 1-1049 | + | 609 | intron (NM_001205332, intron 1 of 17) | Map4 | 10 | Untreated (DSB peaks) |
| 1-973 | + | 812 | promoter-TSS (NM_178856) | Gins2 | 10 | Untreated (DSB peaks) |
| 1-978 | + | 555 | promoter-TSS (NM_172288) | Nup133 | 10 | Untreated (DSB peaks) |
| 1-706 | + | 637 | TTS (NM_172145) | Eva1b | 10 | Untreated (DSB peaks) |
| 1-711 | + | 607 | intron (NM_001080819, intron 1 of 19) | Mir7227 | 10 | Untreated (DSB peaks) |
| 1-921 | + | 664 | promoter-TSS (NM_019391) | Lsp1 | 10 | Untreated (DSB peaks) |
| 1-792 | + | 596 | intron (NR_102383, intron 1 of 1) | 5930430L01Rik | 10 | Untreated (DSB peaks) |
| 20090 | + | 700 | intron (NM_016846, intron 5 of 17) | Rgl1 | 10 | Untreated (DSB peaks) |
| 1-429 | + | 578 | Intergenic | 6330415G19Rik | 10 | Untreated (DSB peaks) |
| 1-401 | + | 639 | Intergenic | Tnk2 | 10 | Untreated (DSB peaks) |
| 1-263 | + | 586 | Intergenic | Cdyl | 10 | Untreated (DSB peaks) |
| 1-550 | + | 658 | Intergenic | Rbm43 | 10 | Untreated (DSB peaks) |
| 1-679 | + | 796 | Intergenic | Musk | 10 | Untreated (DSB peaks) |
| 1-162 | + | 644 | Intergenic | Kdm6b | 10 | Untreated (DSB peaks) |
| 1-1033 | + | 575 | Intergenic | Nedd4 | 10 | Untreated (DSB peaks) |
| 1-894 | + | 632 | Intergenic | Wnt11 | 10 | Untreated (DSB peaks) |
| 1-1042 | + | 542 | Intergenic | Gm5627 | 10 | Untreated (DSB peaks) |
| 1-194 | + | 547 | Intergenic | Arhgap27 | 10 | Untreated (DSB peaks) |
| 1-1047 | + | 567 | Intergenic | Bsn | 10 | Untreated (DSB peaks) |
| 1-765 | + | 587 | Intergenic | E130006D01Rik | 10 | Untreated (DSB peaks) |
| 1-200 | + | 548 | Intergenic | Dus1l | 10 | Untreated (DSB peaks) |
| 1-775 | + | 950 | Intergenic | Ubc | 10 | Untreated (DSB peaks) |
| 1-983 | + | 907 | Intergenic | Irf2bp2 | 10 | Untreated (DSB peaks) |
| 24473 | + | 636 | Intergenic | Cfap126 | 10 | Untreated (DSB peaks) |
| 1-796 | + | 566 | Intergenic | Col1a2 | 9 | Untreated (DSB peaks) |
| 1-471 | + | 545 | Intergenic | Gm26682 | 9 | Untreated (DSB peaks) |
| 1-242 | + | 640 | Intergenic | 1700016G22Rik | 9 | Untreated (DSB peaks) |
| 1-511 | + | 585 | Intergenic | Fth1 | 9 | Untreated (DSB peaks) |
| 1-389 | + | 719 | Intergenic | Efcab1 | 9 | Untreated (DSB peaks) |
| 1-427 | + | 552 | Intergenic | Mir99b | 9 | Untreated (DSB peaks) |
| 1-594 | + | 573 | Intergenic | 4930433B08Rik | 9 | Untreated (DSB peaks) |
| 1-665 | + | 757 | intron (NR_131155, intron 2 of 2) | Gm2560 | 9 | Untreated (DSB peaks) |
| 1-868 | + | 566 | promoter-TSS (NM_001083912) | Plekhg2 | 9 | Untreated (DSB peaks) |
| 1-515 | + | 748 | promoter-TSS (NM_031249) | Cstf2t | 9 | Untreated (DSB peaks) |
| 1-307 | + | 719 | Intergenic | Gm30108 | 9 | Untreated (DSB peaks) |
| 1-539 | + | 858 | intron (NM_153560, intron 1 of 10) | Fam102a | 9 | Untreated (DSB peaks) |
| 1-443 | + | 657 | promoter-TSS (NM_011296) | Vps52 | 9 | Untreated (DSB peaks) |

| | | | | | | |
|---|---|---|---|---|---|---|
| 1-736 | + | 621 | intron (NM_030889, intron 2 of 26) | Psapl1 | 9 | Untreated (DSB peaks) |
| 1-518 | + | 603 | intron (NM_145500, intron 1 of 2) | Ankrd2 | 9 | Untreated (DSB peaks) |
| 1-1005 | + | 546 | promoter-TSS (NM_201372) | Ccdc84 | 9 | Untreated (DSB peaks) |
| 1-484 | + | 666 | intron (NM_001085373, intron 11 of 18) | Dcp2 | 9 | Untreated (DSB peaks) |
| 1-937 | + | 934 | promoter-TSS (NM_027973) | Cenpu | 9 | Untreated (DSB peaks) |
| 1-1015 | + | 804 | intron (NM_019689, intron 1 of 7) | Arid3b | 9 | Untreated (DSB peaks) |
| 1-363 | + | 690 | intron (NM_001276461, intron 1 of 28) | Asap1 | 9 | Untreated (DSB peaks) |
| 1-369 | + | 625 | promoter-TSS (NM_175399) | Exosc4 | 9 | Untreated (DSB peaks) |
| 1-802 | + | 718 | Intergenic | Wnt16 | 9 | Untreated (DSB peaks) |
| 1-114 | + | 751 | promoter-TSS (NM_029272) | Ndufs7 | 9 | Untreated (DSB peaks) |
| 1-115 | + | 753 | promoter-TSS (NM_029272) | Ndufs7 | 9 | Untreated (DSB peaks) |
| 1-636 | + | 577 | promoter-TSS (NM_001159354) | Magi3 | 9 | Untreated (DSB peaks) |
| 15707 | + | 563 | promoter-TSS (NM_133821) | Gm20753 | 9 | Untreated (DSB peaks) |
| 1-969 | + | 628 | TTS (NR_077224) | Lncbate1 | 9 | Untreated (DSB peaks) |
| 1-125 | + | 692 | promoter-TSS (NM_178610) | Krr1 | 9 | Untreated (DSB peaks) |
| 1-908 | + | 679 | intron (NM_020606, intron 1 of 12) | Parvaos | 9 | Untreated (DSB peaks) |
| 1-643 | + | 695 | intron (NM_001025438, intron 14 of 18) | Ank2 | 9 | Untreated (DSB peaks) |
| 1-649 | + | 741 | intron (NM_001277216, intron 1 of 11) | Bmpr1b | 9 | Untreated (DSB peaks) |
| 1-726 | + | 838 | promoter-TSS (NM_001039157) | Ube2j2 | 9 | Untreated (DSB peaks) |
| 1-610 | + | 641 | Intergenic | 5430434I15Rik | 9 | Untreated (DSB peaks) |
| 34335 | + | 639 | Intergenic | Lama4 | 9 | Untreated (DSB peaks) |
| 1-1003 | + | 867 | Intergenic | Thy1 | 9 | Untreated (DSB peaks) |
| 1-104 | + | 677 | Intergenic | Nrbf2 | 9 | Untreated (DSB peaks) |
| 1-113 | + | 577 | Intergenic | Col18a1 | 9 | Untreated (DSB peaks) |
| 1-460 | + | 608 | Intergenic | Map4k3 | 9 | Untreated (DSB peaks) |
| 1-465 | + | 807 | Intergenic | 8430430B14Rik | 9 | Untreated (DSB peaks) |
| 1-1041 | + | 542 | Intergenic | Gm5627 | 9 | Untreated (DSB peaks) |
| 1-759 | + | 589 | Intergenic | Cdc7 | 9 | Untreated (DSB peaks) |
| 1-909 | + | 588 | Intergenic | Tead1 | 9 | Untreated (DSB peaks) |
| 1-1053 | + | 620 | Intergenic | 1700020M21Rik | 9 | Untreated (DSB peaks) |
| 1-858 | + | 659 | Intergenic | Rassf8 | 9 | Untreated (DSB peaks) |
| 1-654 | + | 1000 | Intergenic | Adgrl2 | 9 | Untreated (DSB peaks) |
| 1-475 | + | 742 | Intergenic | Zfp521 | 8 | Untreated (DSB peaks) |
| 1-205 | + | 610 | promoter-TSS (NM_001204912) | Cenpo | 8 | Untreated (DSB peaks) |
| 1-526 | + | 628 | intron (NM_172475, intron 1 of 22) | Frmd4a | 8 | Untreated (DSB peaks) |
| 1-336 | + | 654 | promoter-TSS (NM_026069) | Rpl37 | 8 | Untreated (DSB peaks) |
| 1-508 | + | 547 | promoter-TSS (NM_008577) | Slc3a2 | 8 | Untreated (DSB peaks) |
| 1-861 | + | 665 | promoter-TSS (NM_026885) | Chmp2a | 8 | Untreated (DSB peaks) |
| 1-664 | + | 926 | intron (NM_009822, intron 1 of 10) | Runx1t1 | 8 | Untreated (DSB peaks) |
| 1-244 | + | 568 | intron (NM_008130, intron 10 of 14) | 4933412O06Rik | 8 | Untreated (DSB peaks) |
| 1-864 | + | 559 | promoter-TSS (NM_016871) | Tomm40 | 8 | Untreated (DSB peaks) |
| 43191 | + | 598 | intron (NM_153179, intron 66 of 66) | 4930486I03Rik | 8 | Untreated (DSB peaks) |
| 1-992 | + | 744 | promoter-TSS (NR_029585) | Mir199a-1 | 8 | Untreated (DSB peaks) |
| 1-513 | + | 600 | intron (NM_001033759, intron 2 of 23) | Tmem2 | 8 | Untreated (DSB peaks) |
| 1-865 | + | 557 | TTS (NM_001190974) | Hnrnpul1 | 8 | Untreated (DSB peaks) |
| 1-342 | + | 793 | intron (NM_001081302, intron 34 of 56) | Fam105a | 8 | Untreated (DSB peaks) |
| 1-869 | + | 566 | promoter-TSS (NM_001083912) | Plekhg2 | 8 | Untreated (DSB peaks) |

| | | | | | | |
|---|---|---|---|---|---|---|
| 1-606 | + | 898 | intron (NM_009673, intron 12 of 12) | Mir7009 | 8 | Untreated (DSB peaks) |
| 1-265 | + | 697 | intron (NM_007556, intron 1 of 6) | 4930579J19Rik | 8 | Untreated (DSB peaks) |
| 1-517 | + | 655 | intron (NM_145500, intron 1 of 2) | Ankrd2 | 8 | Untreated (DSB peaks) |
| 1-146 | + | 634 | intron (NM_001291890, intron 3 of 22) | Adam19 | 8 | Untreated (DSB peaks) |
| 42005 | + | 728 | promoter-TSS (NM_001310549) | Nabp1 | 8 | Untreated (DSB peaks) |
| 1-352 | + | 595 | promoter-TSS (NM_030132) | Utp23 | 8 | Untreated (DSB peaks) |
| 44927 | + | 544 | promoter-TSS (NM_001013771) | Gm973 | 8 | Untreated (DSB peaks) |
| 1-1031 | + | 744 | intron (NM_013646, intron 1 of 10) | 9530091C08Rik | 8 | Untreated (DSB peaks) |
| 1-948 | + | 786 | intron (NM_001001491, intron 1 of 7) | Tpm4 | 8 | Untreated (DSB peaks) |
| 1-949 | + | 787 | intron (NM_001001491, intron 1 of 7) | Tpm4 | 8 | Untreated (DSB peaks) |
| 1-367 | + | 549 | intron (NM_153178, intron 1 of 18) | Ago2 | 8 | Untreated (DSB peaks) |
| 47119 | + | 792 | TTS (NM_013612) | Ctdsp1 | 8 | Untreated (DSB peaks) |
| 1-112 | + | 566 | promoter-TSS (NM_001347201) | Gucd1 | 8 | Untreated (DSB peaks) |
| 1-501 | + | 671 | intron (NM_207651, intron 2 of 21) | Slc14a2 | 8 | Untreated (DSB peaks) |
| 1-1034 | + | 827 | promoter-TSS (NM_009945) | Cox7a2 | 8 | Untreated (DSB peaks) |
| 1-755 | + | 597 | promoter-TSS (NM_017367) | 2010109A12Rik | 8 | Untreated (DSB peaks) |
| 1-826 | + | 708 | promoter-TSS (NM_008377) | Lrig1 | 8 | Untreated (DSB peaks) |
| 1-631 | + | 624 | promoter-TSS (NM_001272024) | Sema6c | 8 | Untreated (DSB peaks) |
| 16072 | + | 563 | promoter-TSS (NR_040630) | Gm20753 | 8 | Untreated (DSB peaks) |
| 1-1048 | + | 568 | promoter-TSS (NM_011637) | Trex1 | 8 | Untreated (DSB peaks) |
| 1-979 | + | 551 | intron (NM_175149, intron 2 of 3) | 2310022B05Rik | 8 | Untreated (DSB peaks) |
| 1-710 | + | 547 | promoter-TSS (NM_001358455) | Slc9a1 | 8 | Untreated (DSB peaks) |
| 1-140 | + | 553 | Intergenic | Smim23 | 8 | Untreated (DSB peaks) |
| 1-920 | + | 737 | intron (NM_001252588, intron 1 of 7) | Tspan4 | 8 | Untreated (DSB peaks) |
| 1-717 | + | 697 | promoter-TSS (NM_172338) | Dnajc16 | 8 | Untreated (DSB peaks) |
| 1-718 | + | 922 | intron (NM_010329, intron 1 of 5) | Pdpn | 8 | Untreated (DSB peaks) |
| 21551 | + | 588 | intron (NM_001038619, intron 14 of 20) | Dnm3os | 8 | Untreated (DSB peaks) |
| 1-590 | + | 597 | intron (NM_198656, intron 3 of 14) | Cdh26 | 8 | Untreated (DSB peaks) |
| 1-142 | + | 658 | Intergenic | Slit3 | 8 | Untreated (DSB peaks) |
| 1-213 | + | 655 | Intergenic | Mir680-3 | 8 | Untreated (DSB peaks) |
| 1-483 | + | 738 | Intergenic | Nr3c1 | 8 | Untreated (DSB peaks) |
| 1-998 | + | 548 | Intergenic | Mir125b-1 | 8 | Untreated (DSB peaks) |
| 41275 | + | 557 | Intergenic | Slc40a1 | 8 | Untreated (DSB peaks) |
| 1-321 | + | 668 | Intergenic | Rnase4 | 8 | Untreated (DSB peaks) |
| 1-615 | + | 764 | Intergenic | Elf2 | 8 | Untreated (DSB peaks) |
| 1-743 | + | 543 | Intergenic | Smim20 | 8 | Untreated (DSB peaks) |
| 1-493 | + | 562 | Intergenic | Synpo | 8 | Untreated (DSB peaks) |
| 1-1018 | + | 752 | Intergenic | Coro2b | 8 | Untreated (DSB peaks) |
| 1-1022 | + | 622 | Intergenic | Skor1 | 8 | Untreated (DSB peaks) |
| 1-454 | + | 646 | Intergenic | Fbxl17 | 8 | Untreated (DSB peaks) |
| 12785 | + | 766 | Intergenic | Mir6353 | 8 | Untreated (DSB peaks) |
| 1-466 | + | 680 | non-coding (NR_045458, exon 4 of 6) | 4933433H22Rik | 8 | Untreated (DSB peaks) |
| 1-888 | + | 544 | Intergenic | Ctsc | 8 | Untreated (DSB peaks) |
| 1-896 | + | 575 | Intergenic | Serpinh1 | 8 | Untreated (DSB peaks) |
| 1-377 | + | 564 | Intergenic | Aqp2 | 8 | Untreated (DSB peaks) |
| 1-828 | + | 553 | Intergenic | Prok2 | 8 | Untreated (DSB peaks) |
| 1-830 | + | 654 | Intergenic | Rybp | 8 | Untreated (DSB peaks) |

| | | | | | | |
|---|---|---|---|---|---|---|
| 1-191 | + | 670 | Intergenic | Cavin1 | 8 | Untreated (DSB peaks) |
| 1-201 | + | 548 | Intergenic | Dus1l | 8 | Untreated (DSB peaks) |
| 1-568 | + | 580 | Intergenic | Dut | 8 | Untreated (DSB peaks) |
| 1-789 | + | 742 | Intergenic | Tnrc18 | 8 | Untreated (DSB peaks) |
| 22282 | + | 769 | Intergenic | Gorab | 8 | Untreated (DSB peaks) |
| 1-209 | + | 580 | Intergenic | 1700022H16Rik | 7 | Untreated (DSB peaks) |
| 1-245 | + | 865 | Intergenic | 4933412O06Rik | 7 | Untreated (DSB peaks) |
| 1-251 | + | 651 | Intergenic | Pom121l2 | 7 | Untreated (DSB peaks) |
| 32143 | + | 568 | Intergenic | Gm8709 | 7 | Untreated (DSB peaks) |
| 1-803 | + | 631 | Intergenic | Ccdc136 | 7 | Untreated (DSB peaks) |
| 1-210 | + | 729 | Intergenic | Lamb1 | 7 | Untreated (DSB peaks) |
| 1-312 | + | 818 | Intergenic | Btd | 7 | Untreated (DSB peaks) |
| 1-660 | + | 629 | promoter-TSS (NR_040645) | 8430436N08Rik | 7 | Untreated (DSB peaks) |
| 1-134 | + | 584 | intron (NM_001177629, intron 10 of 15) | Ddc | 7 | Untreated (DSB peaks) |
| 1-730 | + | 615 | intron (NM_001199632, intron 7 of 20) | Lrrc17 | 7 | Untreated (DSB peaks) |
| 1-138 | + | 586 | promoter-TSS (NM_144514) | Commd1 | 7 | Untreated (DSB peaks) |
| 1-926 | + | 632 | promoter-TSS (NM_001042484) | Golga7 | 7 | Untreated (DSB peaks) |
| 1-393 | + | 689 | intron (NM_178665, intron 3 of 12) | Morf4l1-ps1 | 7 | Untreated (DSB peaks) |
| 1-432 | + | 602 | promoter-TSS (NR_027452) | Gm17801 | 7 | Untreated (DSB peaks) |
| 1-866 | + | 591 | promoter-TSS (NM_001190974) | Axl | 7 | Untreated (DSB peaks) |
| 1-437 | + | 784 | intron (NM_026170, intron 1 of 9) | Ergic1 | 7 | Untreated (DSB peaks) |
| 1-804 | + | 629 | intron (NM_001024698, intron 6 of 10) | Cpa2 | 7 | Untreated (DSB peaks) |
| 1-994 | + | 989 | Intergenic | 4933422A05Rik | 7 | Untreated (DSB peaks) |
| 1-930 | + | 649 | intron (NM_001042675, intron 1 of 6) | Rbpms | 7 | Untreated (DSB peaks) |
| 1-446 | + | 563 | promoter-TSS (NM_001252468) | Bag6 | 7 | Untreated (DSB peaks) |
| 1-932 | + | 673 | intron (NM_001081279, intron 1 of 2) | Gm16793 | 7 | Untreated (DSB peaks) |
| 43221 | + | 938 | Intergenic | Hs6st1 | 7 | Untreated (DSB peaks) |
| 43282 | + | 564 | promoter-TSS (NM_146107) | Actr1b | 7 | Untreated (DSB peaks) |
| 1-935 | + | 560 | intron (NM_001205219, intron 1 of 23) | Sorbs2 | 7 | Untreated (DSB peaks) |
| 1-406 | + | 995 | intron (NM_001033238, intron 3 of 17) | Cblb | 7 | Untreated (DSB peaks) |
| 1-521 | + | 571 | intron (NM_011050, intron 1 of 11) | Pdcd4 | 7 | Untreated (DSB peaks) |
| 1-618 | + | 631 | exon (NM_001198766, exon 1 of 21) | Postn | 7 | Untreated (DSB peaks) |
| 1-407 | + | 667 | intron (NM_019631, intron 1 of 5) | Tmem45a | 7 | Untreated (DSB peaks) |
| 1-557 | + | 814 | intron (NM_001159564, intron 1 of 17) | Itgb6 | 7 | Untreated (DSB peaks) |
| 1-157 | + | 546 | promoter-TSS (NM_008885) | Pmp22 | 7 | Untreated (DSB peaks) |
| 1-106 | + | 581 | intron (NM_023598, intron 2 of 9) | Arid5b | 7 | Untreated (DSB peaks) |
| 1-877 | + | 835 | intron (NR_131145, intron 1 of 3) | Gm29683 | 7 | Untreated (DSB peaks) |
| 1-942 | + | 551 | promoter-TSS (NM_029366) | Kxd1 | 7 | Untreated (DSB peaks) |
| 1-219 | + | 711 | intron (NM_008612, intron 1 of 7) | Mnat1 | 7 | Untreated (DSB peaks) |
| 1-327 | + | 563 | intron (NM_001360046, intron 2 of 3) | Serp2 | 7 | Untreated (DSB peaks) |
| 1-608 | + | 594 | Intergenic | Gm20755 | 7 | Untreated (DSB peaks) |
| 1-328 | + | 563 | intron (NM_001360046, intron 2 of 3) | Serp2 | 7 | Untreated (DSB peaks) |
| 1-231 | + | 552 | promoter-TSS (NM_172414) | Zc2hc1c | 7 | Untreated (DSB peaks) |
| 1-824 | + | 545 | intron (NR_015530, intron 1 of 2) | 9530026P05Rik | 7 | Untreated (DSB peaks) |
| 1-754 | + | 636 | intron (NM_001009818, intron 1 of 9) | Sept11 | 7 | Untreated (DSB peaks) |
| 1-121 | + | 604 | intron (NM_146239, intron 1 of 16) | Mir1931 | 7 | Untreated (DSB peaks) |
| 1-423 | + | 592 | intron (NR_040484, intron 1 of 3) | 1600002D24Rik | 7 | Untreated (DSB peaks) |

| | | | | | | |
|---|---|---|---|---|---|---|
| 1-195 | + | 695 | promoter-TSS (NM_027346) | Taco1 | 7 | Untreated (DSB peaks) |
| 1-567 | + | 861 | intron (NM_001331221, intron 3 of 3) | Bmf | 7 | Untreated (DSB peaks) |
| 1-130 | + | 621 | intron (NM_001110218, intron 8 of 9) | Mirlet7i | 7 | Untreated (DSB peaks) |
| 1-707 | + | 638 | TTS (NM_172145) | Eva1b | 7 | Untreated (DSB peaks) |
| 1-777 | + | 604 | intron (NM_021371, intron 3 of 5) | Caln1 | 7 | Untreated (DSB peaks) |
| 1-919 | + | 612 | intron (NM_016978, intron 1 of 9) | Oat | 7 | Untreated (DSB peaks) |
| 1-785 | + | 631 | promoter-TSS (NM_021392) | Mcm7 | 7 | Untreated (DSB peaks) |
| 1-577 | + | 551 | promoter-TSS (NM_029763) | Dzank1 | 7 | Untreated (DSB peaks) |
| 1-795 | + | 673 | intron (NR_045700, intron 1 of 3) | 1700028E10Rik | 7 | Untreated (DSB peaks) |
| 19360 | + | 695 | intron (NM_022018, intron 5 of 13) | 2810414N06Rik | 7 | Untreated (DSB peaks) |
| 23743 | + | 606 | promoter-TSS (NM_027430) | Mpc2 | 7 | Untreated (DSB peaks) |
| 1-544 | + | 701 | Intergenic | Gtdc1 | 7 | Untreated (DSB peaks) |
| 1-279 | + | 549 | Intergenic | Gm5084 | 7 | Untreated (DSB peaks) |
| 1-560 | + | 566 | Intergenic | Tank | 7 | Untreated (DSB peaks) |
| 1-625 | + | 584 | Intergenic | Ccnl1 | 7 | Untreated (DSB peaks) |
| 1-220 | + | 649 | Intergenic | Hif1a | 7 | Untreated (DSB peaks) |
| 1-688 | + | 677 | Intergenic | Dmac1 | 7 | Untreated (DSB peaks) |
| 1-749 | + | 856 | Intergenic | Hopx | 7 | Untreated (DSB peaks) |
| 1-498 | + | 701 | Intergenic | Slc14a1 | 7 | Untreated (DSB peaks) |
| 1-226 | + | 580 | Intergenic | 9430078K24Rik | 7 | Untreated (DSB peaks) |
| 1-462 | + | 628 | Intergenic | C430042M11Rik | 7 | Untreated (DSB peaks) |
| 1-467 | + | 684 | Intergenic | 4933433H22Rik | 7 | Untreated (DSB peaks) |
| 1-178 | + | 639 | Intergenic | Mir21a | 7 | Untreated (DSB peaks) |
| 13881 | + | 619 | Intergenic | Ackr3 | 7 | Untreated (DSB peaks) |
| 1-186 | + | 635 | Intergenic | Mrpl45 | 7 | Untreated (DSB peaks) |
| 1-841 | + | 605 | Intergenic | Cxcl12 | 7 | Untreated (DSB peaks) |
| 16803 | + | 721 | Intergenic | En1 | 7 | Untreated (DSB peaks) |
| 1-642 | + | 614 | Intergenic | Slc44a3 | 7 | Untreated (DSB peaks) |
| 1-770 | + | 764 | Intergenic | Ncor2 | 7 | Untreated (DSB peaks) |
| 1-981 | + | 549 | Intergenic | Tarbp1 | 7 | Untreated (DSB peaks) |
| 1-848 | + | 637 | Intergenic | Gm38404 | 7 | Untreated (DSB peaks) |
| 1-644 | + | 788 | Intergenic | Sec24b | 7 | Untreated (DSB peaks) |
| 1-923 | + | 552 | Intergenic | Mrgprd | 7 | Untreated (DSB peaks) |
| 1-860 | + | 704 | Intergenic | Sspn | 7 | Untreated (DSB peaks) |
| 1-588 | + | 819 | Intergenic | Pmepa1 | 7 | Untreated (DSB peaks) |
| 28126 | + | 692 | Intergenic | Hhat | 7 | Untreated (DSB peaks) |
| 1-340 | + | 591 | Intergenic | Slc1a3 | 6 | Untreated (DSB peaks) |
| 1-208 | + | 563 | Intergenic | Laptm4a | 6 | Untreated (DSB peaks) |
| 1-246 | + | 657 | Intergenic | 4933412O06Rik | 6 | Untreated (DSB peaks) |
| 1-532 | + | 596 | Intergenic | Commd3 | 6 | Untreated (DSB peaks) |
| 1-249 | + | 587 | Intergenic | Gpx5 | 6 | Untreated (DSB peaks) |
| 1-597 | + | 567 | Intergenic | Tbl1xr1 | 6 | Untreated (DSB peaks) |
| 1-309 | + | 560 | Intergenic | Gm32123 | 6 | Untreated (DSB peaks) |
| 1-602 | + | 632 | Intergenic | Naaladl2 | 6 | Untreated (DSB peaks) |
| 1-604 | + | 597 | Intergenic | Tmem212 | 6 | Untreated (DSB peaks) |
| 32509 | + | 730 | Intergenic | Ncoa7 | 6 | Untreated (DSB peaks) |
| 1-343 | + | 573 | Intergenic | Dap | 6 | Untreated (DSB peaks) |

| | | | | | | |
|---|---|---|---|---|---|---|
| 1-516 | + | 879 | Intergenic | Mir3970 | 6 | Untreated (DSB peaks) |
| 1-873 | + | 605 | Intergenic | Lrp3 | 6 | Untreated (DSB peaks) |
| 1-874 | + | 605 | Intergenic | Lrp3 | 6 | Untreated (DSB peaks) |
| 1-214 | + | 570 | Intergenic | Mir680-3 | 6 | Untreated (DSB peaks) |
| 1-1001 | + | 606 | Intergenic | Sorl1 | 6 | Untreated (DSB peaks) |
| 1-507 | + | 550 | TTS (NM_008893) | Cdc42ep2 | 6 | Untreated (DSB peaks) |
| 29587 | + | 790 | promoter-TSS (NM_001347361) | Pex3 | 6 | Untreated (DSB peaks) |
| 30317 | + | 603 | intron (NM_025446, intron 3 of 5) | Aig1 | 6 | Untreated (DSB peaks) |
| 1-801 | + | 589 | intron (NM_212435, intron 3 of 16) | Foxp2 | 6 | Untreated (DSB peaks) |
| 1-531 | + | 588 | intron (NM_026162, intron 1 of 13) | Plxdc2 | 6 | Untreated (DSB peaks) |
| 43160 | + | 802 | intron (NM_153179, intron 66 of 66) | 4930486I03Rik | 6 | Untreated (DSB peaks) |
| 1-871 | + | 597 | intron (NM_021895, intron 1 of 20) | Capn12 | 6 | Untreated (DSB peaks) |
| 1-310 | + | 592 | intron (NM_009785, intron 3 of 37) | Cacna2d3 | 6 | Untreated (DSB peaks) |
| 1-345 | + | 598 | intron (NM_176073, intron 4 of 7) | 1700084J12Rik | 6 | Untreated (DSB peaks) |
| 1-735 | + | 692 | promoter-TSS (NM_001289804) | Poln | 6 | Untreated (DSB peaks) |
| 32874 | + | 777 | promoter-TSS (NM_008538) | Marcks | 6 | Untreated (DSB peaks) |
| 1-402 | + | 582 | intron (NM_008047, intron 2 of 10) | Fstl1 | 6 | Untreated (DSB peaks) |
| 35065 | + | 714 | promoter-TSS (NR_152260) | Gm16364 | 6 | Untreated (DSB peaks) |
| 1-215 | + | 545 | promoter-TSS (NR_035459) | Mir1938 | 6 | Untreated (DSB peaks) |
| 1-449 | + | 629 | intron (NM_181753, intron 4 of 6) | Opn5 | 6 | Untreated (DSB peaks) |
| 1-672 | + | 754 | intron (NM_001037709, intron 1 of 11) | Rusc2 | 6 | Untreated (DSB peaks) |
| 1-1006 | + | 546 | promoter-TSS (NM_201372) | Ccdc84 | 6 | Untreated (DSB peaks) |
| 43374 | + | 771 | Intergenic | Wdr75 | 6 | Untreated (DSB peaks) |
| 1-145 | + | 634 | intron (NM_001291890, intron 3 of 22) | Adam19 | 6 | Untreated (DSB peaks) |
| 1-936 | + | 576 | promoter-TSS (NM_007450) | Slc25a4 | 6 | Untreated (DSB peaks) |
| 1-323 | + | 618 | TTS (NR_045728) | Arhgef40 | 6 | Untreated (DSB peaks) |
| 1-617 | + | 631 | promoter-TSS (NM_001198766) | Postn | 6 | Untreated (DSB peaks) |
| 1-556 | + | 557 | promoter-TSS (NR_040356) | 4930555B11Rik | 6 | Untreated (DSB peaks) |
| 44197 | + | 571 | promoter-TSS (NM_027351) | Ppil3 | 6 | Untreated (DSB peaks) |
| 1-815 | + | 564 | intron (NM_183183, intron 1 of 1) | Gprin3 | 6 | Untreated (DSB peaks) |
| 1-808 | + | 569 | Intergenic | Cul1 | 6 | Untreated (DSB peaks) |
| 1-156 | + | 739 | promoter-TSS (NM_145427) | Gid4 | 6 | Untreated (DSB peaks) |
| 1-284 | + | 682 | intron (NM_001289926, intron 13 of 15) | 2010111I01Rik | 6 | Untreated (DSB peaks) |
| 1-939 | + | 643 | intron (NM_009390, intron 18 of 20) | Sgo2b | 6 | Untreated (DSB peaks) |
| 46388 | + | 604 | promoter-TSS (NM_025964) | Mettl21a | 6 | Untreated (DSB peaks) |
| 1-1028 | + | 702 | promoter-TSS (NM_026674) | Aph1c | 6 | Untreated (DSB peaks) |
| 1-1032 | + | 576 | intron (NM_013646, intron 1 of 10) | Rora | 6 | Untreated (DSB peaks) |
| 1-817 | + | 832 | promoter-TSS (NM_173001) | Kdm3a | 6 | Untreated (DSB peaks) |
| 1-165 | + | 736 | intron (NM_001002764, intron 12 of 18) | Mir684-1 | 6 | Untreated (DSB peaks) |
| 1-166 | + | 595 | intron (NM_001002764, intron 15 of 18) | Hic1 | 6 | Untreated (DSB peaks) |
| 1-150 | + | 622 | Intergenic | Rufy1 | 6 | Untreated (DSB peaks) |
| 1-329 | + | 847 | intron (NR_015485, intron 4 of 4) | 1700108F19Rik | 6 | Untreated (DSB peaks) |
| 1-223 | + | 954 | intron (NM_009014, intron 7 of 10) | Rad51b | 6 | Untreated (DSB peaks) |
| 12420 | + | 572 | intron (NM_029777, intron 6 of 6) | Rhbdd1 | 6 | Untreated (DSB peaks) |
| 1-887 | + | 628 | intron (NM_007488, intron 5 of 18) | Arnt2 | 6 | Untreated (DSB peaks) |
| 1-180 | + | 690 | intron (NM_001356531, intron 5 of 11) | Vmp1 | 6 | Untreated (DSB peaks) |
| 1-821 | + | 702 | intron (NR_149744, intron 3 of 5) | Eefsec | 6 | Untreated (DSB peaks) |

| | | | | | | |
|---|---|---|---|---|---|---|
| 1-416 | + | 761 | intron (NM_015755, intron 3 of 10) | Hunk | 6 | Untreated (DSB peaks) |
| 1-289 | + | 806 | intron (NM_025770, intron 2 of 7) | Atg10 | 6 | Untreated (DSB peaks) |
| 1-290 | + | 656 | intron (NM_025770, intron 2 of 7) | Atg10 | 6 | Untreated (DSB peaks) |
| 1-563 | + | 707 | promoter-TSS (NM_026944) | Alkbh3 | 6 | Untreated (DSB peaks) |
| 1-892 | + | 632 | intron (NM_011858, intron 5 of 32) | Gm15413 | 6 | Untreated (DSB peaks) |
| 1-294 | + | 618 | intron (NM_173757, intron 4 of 10) | 6430562O15Rik | 6 | Untreated (DSB peaks) |
| 1-234 | + | 611 | promoter-TSS (NM_001361020) | Dglucy | 6 | Untreated (DSB peaks) |
| 1-758 | + | 563 | intron (NM_029415, intron 1 of 5) | Slc10a6 | 6 | Untreated (DSB peaks) |
| 1-967 | + | 686 | promoter-TSS (NM_011764) | Zfp90 | 6 | Untreated (DSB peaks) |
| 1-485 | + | 746 | Intergenic | Zfp474 | 6 | Untreated (DSB peaks) |
| 1-637 | + | 582 | promoter-TSS (NM_001039488) | Lrif1 | 6 | Untreated (DSB peaks) |
| 1-970 | + | 595 | promoter-TSS (NM_028018) | Ist1 | 6 | Untreated (DSB peaks) |
| 1-639 | + | 557 | intron (NM_001163348, intron 2 of 6) | Ntng1 | 6 | Untreated (DSB peaks) |
| 1-905 | + | 550 | TTS (NM_009627) | Adm | 6 | Untreated (DSB peaks) |
| 1-972 | + | 592 | 5' UTR (NM_146217, exon 1 of 21) | Aars | 6 | Untreated (DSB peaks) |
| 1-1051 | + | 624 | intron (NM_001172124, intron 1 of 13) | Rbms3 | 6 | Untreated (DSB peaks) |
| 1-204 | + | 621 | promoter-TSS (NM_029878) | Tbcd | 6 | Untreated (DSB peaks) |
| 1-768 | + | 591 | intron (NM_177242, intron 1 of 5) | Pptc7 | 6 | Untreated (DSB peaks) |
| 1-129 | + | 679 | intron (NM_001110218, intron 6 of 9) | Mirlet7i | 6 | Untreated (DSB peaks) |
| 1-774 | + | 661 | promoter-TSS (NM_019639) | Ubc | 6 | Untreated (DSB peaks) |
| 1-569 | + | 763 | promoter-TSS (NM_198630) | 1810024B03Rik | 6 | Untreated (DSB peaks) |
| 1-782 | + | 710 | promoter-TSS (NM_010717) | Limk1 | 6 | Untreated (DSB peaks) |
| 1-854 | + | 562 | intron (NM_001080553, intron 1 of 5) | Gsg1 | 6 | Untreated (DSB peaks) |
| 18264 | + | 648 | promoter-TSS (NR_045324) | Gm19705 | 6 | Untreated (DSB peaks) |
| 1-857 | + | 601 | TTS (NM_001164401) | Eps8 | 6 | Untreated (DSB peaks) |
| 1-677 | + | 571 | Intergenic | Klf4 | 6 | Untreated (DSB peaks) |
| 1-678 | + | 570 | Intergenic | Klf4 | 6 | Untreated (DSB peaks) |
| 1-490 | + | 670 | Intergenic | Gramd3 | 6 | Untreated (DSB peaks) |
| 1-922 | + | 578 | intron (NM_001128082, intron 3 of 5) | Tspan32 | 6 | Untreated (DSB peaks) |
| 20821 | + | 566 | intron (NR_040562, intron 2 of 2) | 2810442N19Rik | 6 | Untreated (DSB peaks) |
| 26299 | + | 565 | promoter-TSS (NM_026171) | Nvl | 6 | Untreated (DSB peaks) |
| 1-153 | + | 664 | Intergenic | Larp1 | 6 | Untreated (DSB peaks) |
| 1-154 | + | 664 | Intergenic | Larp1 | 6 | Untreated (DSB peaks) |
| 1-358 | + | 712 | Intergenic | Gm19510 | 6 | Untreated (DSB peaks) |
| 1-278 | + | 889 | Intergenic | A530065N20Rik | 6 | Untreated (DSB peaks) |
| 1-282 | + | 646 | Intergenic | Dapk1 | 6 | Untreated (DSB peaks) |
| 1-1017 | + | 640 | Intergenic | Gm5122 | 6 | Untreated (DSB peaks) |
| 1-361 | + | 542 | Intergenic | A1bg | 6 | Untreated (DSB peaks) |
| 1-623 | + | 585 | Intergenic | Mme | 6 | Untreated (DSB peaks) |
| 1-409 | + | 672 | Intergenic | Pros1 | 6 | Untreated (DSB peaks) |
| 1-745 | + | 618 | Intergenic | Tlr1 | 6 | Untreated (DSB peaks) |
| 1-365 | + | 759 | Intergenic | Wisp1 | 6 | Untreated (DSB peaks) |
| 1-105 | + | 676 | Intergenic | Nrbf2 | 6 | Untreated (DSB peaks) |
| 1-109 | + | 696 | Intergenic | Arid5b | 6 | Untreated (DSB peaks) |
| 1-110 | + | 596 | Intergenic | 4930545H06Rik | 6 | Untreated (DSB peaks) |
| 1-218 | + | 568 | Intergenic | 5830428M24Rik | 6 | Untreated (DSB peaks) |
| 1-463 | + | 675 | Intergenic | 8430430B14Rik | 6 | Untreated (DSB peaks) |

| | | | | | | |
|---|---|---|---|---|---|---|
| 1-1037 | + | 738 | Intergenic | 4930554C24Rik | 6 | Untreated (DSB peaks) |
| 1-177 | + | 745 | Intergenic | Med13 | 6 | Untreated (DSB peaks) |
| 13150 | + | 545 | Intergenic | Ackr3 | 6 | Untreated (DSB peaks) |
| 13516 | + | 619 | Intergenic | Ackr3 | 6 | Untreated (DSB peaks) |
| 1-751 | + | 553 | Intergenic | Cxcl5 | 6 | Untreated (DSB peaks) |
| 1-632 | + | 687 | Intergenic | Mcl1 | 6 | Untreated (DSB peaks) |
| 1-890 | + | 706 | Intergenic | Tenm4 | 6 | Untreated (DSB peaks) |
| 1-891 | + | 599 | Intergenic | Tenm4 | 6 | Untreated (DSB peaks) |
| 1-893 | + | 666 | Intergenic | Lrrc32 | 6 | Untreated (DSB peaks) |
| 1-829 | + | 654 | Intergenic | Rybp | 6 | Untreated (DSB peaks) |
| 1-964 | + | 805 | Intergenic | Cdh5 | 6 | Untreated (DSB peaks) |
| 1-965 | + | 620 | Intergenic | Cdh5 | 6 | Untreated (DSB peaks) |
| 1-843 | + | 544 | Intergenic | 6820426E19Rik | 6 | Untreated (DSB peaks) |
| 1-199 | + | 642 | Intergenic | Baiap2 | 6 | Untreated (DSB peaks) |
| 1-1052 | + | 644 | Intergenic | 5830454E08Rik | 6 | Untreated (DSB peaks) |
| 1-772 | + | 644 | Intergenic | Scarb1 | 6 | Untreated (DSB peaks) |
| 1-849 | + | 637 | Intergenic | Gm38404 | 6 | Untreated (DSB peaks) |
| 1-851 | + | 658 | Intergenic | Klrb1b | 6 | Untreated (DSB peaks) |
| 1-719 | + | 552 | Intergenic | Dhrs3 | 6 | Untreated (DSB peaks) |
| 1-859 | + | 659 | Intergenic | Rassf8 | 6 | Untreated (DSB peaks) |
| 1-652 | + | 615 | Intergenic | Ttll7 | 6 | Untreated (DSB peaks) |
| 1-653 | + | 1000 | Intergenic | Adgrl2 | 6 | Untreated (DSB peaks) |
| 24108 | + | 625 | Intergenic | Mpzl1 | 6 | Untreated (DSB peaks) |
| 1-584 | + | 567 | Intergenic | Sulf2 | 6 | Untreated (DSB peaks) |
| 1-586 | + | 739 | Intergenic | Snai1 | 6 | Untreated (DSB peaks) |
| 1-728 | + | 715 | Intergenic | Mterf1b | 5 | Untreated (DSB peaks) |
| 1-207 | + | 636 | Intergenic | Laptm4a | 5 | Untreated (DSB peaks) |
| 1-663 | + | 683 | Intergenic | Runx1t1 | 5 | Untreated (DSB peaks) |
| 1-596 | + | 573 | Intergenic | Cp | 5 | Untreated (DSB peaks) |
| 1-248 | + | 614 | Intergenic | Trim27 | 5 | Untreated (DSB peaks) |
| 1-603 | + | 596 | Intergenic | Tmem212 | 5 | Untreated (DSB peaks) |
| 1-400 | + | 599 | Intergenic | Fam43a | 5 | Untreated (DSB peaks) |
| 1-668 | + | 665 | Intergenic | Map3k7 | 5 | Untreated (DSB peaks) |
| 1-1000 | + | 607 | Intergenic | Sorl1 | 5 | Untreated (DSB peaks) |
| 1-269 | + | 614 | Intergenic | Phactr1 | 5 | Untreated (DSB peaks) |
| 1-1004 | + | 637 | Intergenic | Thy1 | 5 | Untreated (DSB peaks) |
| 1-934 | + | 663 | Intergenic | Fat1 | 5 | Untreated (DSB peaks) |
| 43344 | + | 783 | Intergenic | Col3a1 | 5 | Untreated (DSB peaks) |
| 1-143 | + | 550 | Intergenic | Gm12159 | 5 | Untreated (DSB peaks) |
| 43405 | + | 772 | Intergenic | Wdr75 | 5 | Untreated (DSB peaks) |
| 1-147 | + | 732 | Intergenic | Trim41 | 5 | Untreated (DSB peaks) |
| 1-453 | + | 626 | Intergenic | Gm20098 | 5 | Untreated (DSB peaks) |
| 1-276 | + | 654 | Intergenic | Hrh2 | 5 | Untreated (DSB peaks) |
| 1-554 | + | 623 | Intergenic | Galnt13 | 5 | Untreated (DSB peaks) |
| 1-555 | + | 623 | Intergenic | Galnt13 | 5 | Untreated (DSB peaks) |
| 1-1014 | + | 566 | Intergenic | Crabp1 | 5 | Untreated (DSB peaks) |
| 1-620 | + | 576 | Intergenic | Tm4sf1 | 5 | Untreated (DSB peaks) |

| | | | | | | |
|---|---|---|---|---|---|---|
| 1-503 | + | 872 | promoter-TSS (NM_001347420) | Ighmbp2 | 5 | Untreated (DSB peaks) |
| 1-206 | + | 583 | promoter-TSS (NM_001099628) | Atad2b | 5 | Untreated (DSB peaks) |
| 1-986 | + | 588 | intron (NM_010809, intron 8 of 9) | Mmp3 | 5 | Untreated (DSB peaks) |
| 1-591 | + | 547 | promoter-TSS (NM_173181) | Zc2hc1a | 5 | Untreated (DSB peaks) |
| 1-384 | + | 596 | 5' UTR (NM_015769, exon 1 of 11) | Ercc4 | 5 | Untreated (DSB peaks) |
| 29952 | + | 791 | promoter-TSS (NM_001347361) | Pex3 | 5 | Untreated (DSB peaks) |
| 1-799 | + | 581 | promoter-TSS (NM_175312) | Bmt2 | 5 | Untreated (DSB peaks) |
| 1-306 | + | 778 | intron (NM_001289761, intron 2 of 8) | Rarb | 5 | Untreated (DSB peaks) |
| 1-593 | + | 610 | intron (NM_007825, intron 1 of 5) | Cyp7b1 | 5 | Untreated (DSB peaks) |
| 1-137 | + | 681 | promoter-TSS (NM_033523) | Spred2 | 5 | Untreated (DSB peaks) |
| 35796 | + | 579 | Intergenic | Sh3rf3 | 5 | Untreated (DSB peaks) |
| 1-534 | + | 870 | intron (NM_001081006, intron 2 of 21) | Etl4 | 5 | Untreated (DSB peaks) |
| 30682 | + | 544 | promoter-TSS (NM_026203) | Ahi1 | 5 | Untreated (DSB peaks) |
| 1-732 | + | 572 | intron (NM_029990, intron 2 of 2) | 5031425E22Rik | 5 | Untreated (DSB peaks) |
| 1-431 | + | 601 | promoter-TSS (NR_027452) | Ube2i | 5 | Untreated (DSB peaks) |
| 1-929 | + | 545 | 5' UTR (NM_010206, exon 1 of 18) | Fgfr1 | 5 | Untreated (DSB peaks) |
| 1-311 | + | 550 | promoter-TSS (NM_133761) | Dcp1a | 5 | Untreated (DSB peaks) |
| 1-540 | + | 556 | intron (NM_146119, intron 1 of 13) | Fam129b | 5 | Untreated (DSB peaks) |
| 1-359 | + | 561 | Intergenic | Gm19510 | 5 | Untreated (DSB peaks) |
| 1-280 | + | 549 | Intergenic | Gm5084 | 5 | Untreated (DSB peaks) |
| 1-346 | + | 548 | intron (NM_001358780, intron 10 of 18) | Rpl30 | 5 | Untreated (DSB peaks) |
| 1-445 | + | 562 | promoter-TSS (NM_001252468) | Bag6 | 5 | Untreated (DSB peaks) |
| 1-931 | + | 673 | intron (NM_001081279, intron 1 of 2) | Gm16793 | 5 | Untreated (DSB peaks) |
| 1-609 | + | 605 | intron (NR_040559, intron 2 of 3) | Gm20755 | 5 | Untreated (DSB peaks) |
| 34700 | + | 559 | promoter-TSS (NM_001122892) | Fyn | 5 | Untreated (DSB peaks) |
| 1-682 | + | 553 | Intergenic | Rgs3 | 5 | Untreated (DSB peaks) |
| 1-1007 | + | 622 | promoter-TSS (NM_001357490) | Bcl9l | 5 | Untreated (DSB peaks) |
| 1-740 | + | 768 | intron (NM_001359071, intron 2 of 7) | Ldb2 | 5 | Untreated (DSB peaks) |
| 1-272 | + | 609 | intron (NM_001199304, intron 2 of 6) | Atxn1 | 5 | Untreated (DSB peaks) |
| 1-938 | + | 591 | intron (NM_133791, intron 2 of 22) | Wwc2 | 5 | Untreated (DSB peaks) |
| 1-148 | + | 548 | 5' UTR (NM_008326, exon 1 of 3) | Irgm1 | 5 | Untreated (DSB peaks) |
| 1-813 | + | 549 | promoter-TSS (NM_029353) | Malsu1 | 5 | Untreated (DSB peaks) |
| 1-318 | + | 688 | intron (NM_029238, intron 1 of 7) | Slc35f4 | 5 | Untreated (DSB peaks) |
| 1-149 | + | 541 | promoter-TSS (NM_026543) | Mrnip | 5 | Untreated (DSB peaks) |
| 1-322 | + | 545 | promoter-TSS (NM_021472) | Rnase4 | 5 | Untreated (DSB peaks) |
| 1-616 | + | 612 | intron (NM_080793, intron 1 of 7) | 5031434O11Rik | 5 | Untreated (DSB peaks) |
| 1-216 | + | 655 | promoter-TSS (NM_052973) | Strn3 | 5 | Untreated (DSB peaks) |
| 1-354 | + | 542 | intron (NM_010162, intron 1 of 10) | Ext1 | 5 | Untreated (DSB peaks) |
| 1-1013 | + | 624 | intron (NM_212445, intron 1 of 7) | Kdelc2 | 5 | Untreated (DSB peaks) |
| 1-158 | + | 546 | promoter-TSS (NM_008885) | Pmp22 | 5 | Untreated (DSB peaks) |
| 1-685 | + | 623 | intron (NM_025685, intron 24 of 60) | Mir455 | 5 | Untreated (DSB peaks) |
| 46023 | + | 562 | Intergenic | Klf7 | 5 | Untreated (DSB peaks) |
| 45658 | + | 540 | intron (NM_001256158, intron 2 of 2) | 4933402D24Rik | 5 | Untreated (DSB peaks) |
| 1-748 | + | 854 | intron (NM_013689, intron 1 of 17) | Tec | 5 | Untreated (DSB peaks) |
| 1-364 | + | 580 | Intergenic | Asap1 | 5 | Untreated (DSB peaks) |
| 1-169 | + | 584 | intron (NM_008750, intron 1 of 7) | Mrm3 | 5 | Untreated (DSB peaks) |
| 1-499 | + | 650 | intron (NM_207651, intron 3 of 21) | Slc14a2 | 5 | Untreated (DSB peaks) |

| | | | | | | |
|---|---|---|---|---|---|---|
| 1-500 | + | 597 | intron (NM_207651, intron 3 of 21) | Slc14a2 | 5 | Untreated (DSB peaks) |
| 1-459 | + | 544 | intron (NM_001360435, intron 1 of 6) | Srsf7 | 5 | Untreated (DSB peaks) |
| 1-956 | + | 633 | promoter-TSS (NM_018771) | Gipc1 | 5 | Untreated (DSB peaks) |
| 1-884 | + | 562 | promoter-TSS (NM_023403) | Mesd | 5 | Untreated (DSB peaks) |
| 1-886 | + | 697 | intron (NM_007488, intron 5 of 18) | Arnt2 | 5 | Untreated (DSB peaks) |
| 1-957 | + | 583 | intron (NM_001297601, intron 6 of 8) | Lyl1 | 5 | Untreated (DSB peaks) |
| 1-414 | + | 543 | promoter-TSS (NM_017404) | Mrpl39 | 5 | Untreated (DSB peaks) |
| 1-415 | + | 671 | promoter-TSS (NM_001198824) | App | 5 | Untreated (DSB peaks) |
| 1-119 | + | 729 | intron (NM_013722, intron 3 of 13) | Syn3 | 5 | Untreated (DSB peaks) |
| 1-421 | + | 578 | promoter-TSS (NM_026110) | Paxbp1 | 5 | Untreated (DSB peaks) |
| 1-895 | + | 731 | promoter-TSS (NM_009825) | Serpinh1 | 5 | Untreated (DSB peaks) |
| 1-899 | + | 549 | promoter-TSS (NM_001362166) | Gdpd5 | 5 | Untreated (DSB peaks) |
| 1-159 | + | 571 | Intergenic | Myh10 | 5 | Untreated (DSB peaks) |
| 1-1039 | + | 617 | promoter-TSS (NM_001301689) | Anapc13 | 5 | Untreated (DSB peaks) |
| 1-1046 | + | 592 | 5' UTR (NM_001324534, exon 1 of 3) | Rpl29 | 5 | Untreated (DSB peaks) |
| 1-761 | + | 601 | intron (NM_011578, intron 3 of 16) | Tgfbr3 | 5 | Untreated (DSB peaks) |
| 1-901 | + | 650 | intron (NM_001001326, intron 1 of 19) | St5 | 5 | Untreated (DSB peaks) |
| 1-904 | + | 550 | TTS (NM_009627) | Adm | 5 | Untreated (DSB peaks) |
| 1-126 | + | 579 | promoter-TSS (NM_024454) | Rab21 | 5 | Untreated (DSB peaks) |
| 1-878 | + | 699 | Intergenic | Gm29683 | 5 | Untreated (DSB peaks) |
| 1-203 | + | 612 | 5' UTR (NM_139059, exon 1 of 9) | Csnk1d | 5 | Untreated (DSB peaks) |
| 1-911 | + | 561 | promoter-TSS (NM_019738) | Nupr1 | 5 | Untreated (DSB peaks) |
| 1-776 | + | 551 | intron (NM_001130476, intron 3 of 5) | Kctd7 | 5 | Untreated (DSB peaks) |
| 1-784 | + | 789 | intron (NM_001291238, intron 3 of 22) | Mir721 | 5 | Untreated (DSB peaks) |
| 1-574 | + | 647 | intron (NR_149746, intron 4 of 8) | Slx4ip | 5 | Untreated (DSB peaks) |
| 1-716 | + | 698 | promoter-TSS (NM_172338) | Dnajc16 | 5 | Untreated (DSB peaks) |
| 1-790 | + | 590 | intron (NM_001163548, intron 1 of 12) | Cyth3 | 5 | Untreated (DSB peaks) |
| 1-721 | + | 646 | promoter-TSS (NR_040649) | 1700045H11Rik | 5 | Untreated (DSB peaks) |
| 1-461 | + | 616 | Intergenic | Map4k3 | 5 | Untreated (DSB peaks) |
| 1-117 | + | 914 | Intergenic | 1500009L16Rik | 5 | Untreated (DSB peaks) |
| 1-468 | + | 564 | Intergenic | 4933433H22Rik | 5 | Untreated (DSB peaks) |
| 1-898 | + | 826 | Intergenic | Serpinh1 | 5 | Untreated (DSB peaks) |
| 1-832 | + | 774 | Intergenic | 1700049E22Rik | 5 | Untreated (DSB peaks) |
| 1-962 | + | 1000 | Intergenic | 4933400L20Rik | 5 | Untreated (DSB peaks) |
| 16438 | + | 594 | Intergenic | Mir3473f | 5 | Untreated (DSB peaks) |
| 1-239 | + | 574 | Intergenic | 4930465M20Rik | 5 | Untreated (DSB peaks) |
| 1-1050 | + | 723 | Intergenic | Prss42 | 5 | Untreated (DSB peaks) |
| 1-763 | + | 624 | Intergenic | C130026L21Rik | 5 | Untreated (DSB peaks) |
| 17168 | + | 722 | Intergenic | En1 | 5 | Untreated (DSB peaks) |
| 1-980 | + | 572 | Intergenic | Tarbp1 | 5 | Untreated (DSB peaks) |
| 1-793 | + | 705 | Intergenic | Alox5ap | 5 | Untreated (DSB peaks) |
| 1-658 | + | 546 | Intergenic | Adgrl4 | 5 | Untreated (DSB peaks) |
| 1-582 | + | 830 | Intergenic | Top1 | 5 | Untreated (DSB peaks) |
| 1-587 | + | 817 | Intergenic | Pmepa1 | 5 | Untreated (DSB peaks) |
| 1-379 | + | 577 | Intergenic | Gm5766 | 4 | Untreated (DSB peaks) |
| 1-797 | + | 561 | Intergenic | Col1a2 | 4 | Untreated (DSB peaks) |
| 1-470 | + | 678 | Intergenic | Jcad | 4 | Untreated (DSB peaks) |

| | | | | | | |
|---|---|---|---|---|---|---|
| 1-243 | + | 586 | Intergenic | 1700016G22Rik | 4 | Untreated (DSB peaks) |
| 1-301 | + | 676 | Intergenic | Flnb | 4 | Untreated (DSB peaks) |
| 1-388 | + | 656 | Intergenic | Efcab1 | 4 | Untreated (DSB peaks) |
| 1-512 | + | 958 | Intergenic | Anxa1 | 4 | Untreated (DSB peaks) |
| 1-247 | + | 731 | Intergenic | Olfr1368 | 4 | Untreated (DSB peaks) |
| 1-250 | + | 972 | Intergenic | Gpx5 | 4 | Untreated (DSB peaks) |
| 1-601 | + | 552 | Intergenic | Tbl1xr1 | 4 | Untreated (DSB peaks) |
| 31048 | + | 569 | Intergenic | Tcf21 | 4 | Untreated (DSB peaks) |
| 1-258 | + | 716 | Intergenic | Hist1h4h | 4 | Untreated (DSB peaks) |
| 1-927 | + | 669 | Intergenic | Tcim | 4 | Untreated (DSB peaks) |
| 1-399 | + | 734 | Intergenic | Cpn2 | 4 | Untreated (DSB peaks) |
| 1-667 | + | 665 | Intergenic | Map3k7 | 4 | Untreated (DSB peaks) |
| 1-872 | + | 557 | Intergenic | Lsm14a | 4 | Untreated (DSB peaks) |
| 1-995 | + | 983 | Intergenic | 4933422A05Rik | 4 | Untreated (DSB peaks) |
| 33604 | + | 667 | Intergenic | Marcks | 4 | Untreated (DSB peaks) |
| 33970 | + | 579 | Intergenic | Marcks | 4 | Untreated (DSB peaks) |
| 1-607 | + | 672 | Intergenic | Gm20755 | 4 | Untreated (DSB peaks) |
| 43435 | + | 717 | Intergenic | Wdr75 | 4 | Untreated (DSB peaks) |
| 1-614 | + | 662 | Intergenic | Pcdh18 | 4 | Untreated (DSB peaks) |
| 1-319 | + | 946 | Intergenic | Olfr750 | 4 | Untreated (DSB peaks) |
| 1-814 | + | 591 | Intergenic | Mir148a | 4 | Untreated (DSB peaks) |
| 1-274 | + | 837 | Intergenic | S1pr3 | 4 | Untreated (DSB peaks) |
| 1-324 | + | 781 | Intergenic | Mmp14 | 4 | Untreated (DSB peaks) |
| 1-488 | + | 617 | Intergenic | Zfp608 | 4 | Untreated (DSB peaks) |
| 1-366 | + | 758 | Intergenic | Wisp1 | 4 | Untreated (DSB peaks) |
| 1-108 | + | 696 | Intergenic | Arid5b | 4 | Untreated (DSB peaks) |
| 1-879 | + | 619 | Intergenic | Nr2f2 | 4 | Untreated (DSB peaks) |
| 1-818 | + | 782 | Intergenic | 9130221F21Rik | 4 | Untreated (DSB peaks) |
| 1-505 | + | 542 | exon (NM_019869, exon 1 of 3) | Rbm14 | 4 | Untreated (DSB peaks) |
| 1-987 | + | 587 | exon (NM_010809, exon 9 of 10) | Mmp3 | 4 | Untreated (DSB peaks) |
| 1-661 | + | 555 | promoter-TSS (NR_040645) | 8430436N08Rik | 4 | Untreated (DSB peaks) |
| 1-339 | + | 558 | intron (NM_148938, intron 3 of 9) | Slc1a3 | 4 | Untreated (DSB peaks) |
| 1-988 | + | 590 | intron (NM_027823, intron 3 of 22) | Gm16833 | 4 | Untreated (DSB peaks) |
| 1-476 | + | 567 | intron (NM_001360804, intron 3 of 7) | Zfp521 | 4 | Untreated (DSB peaks) |
| 1-800 | + | 590 | intron (NM_212435, intron 3 of 16) | Foxp2 | 4 | Untreated (DSB peaks) |
| 1-862 | + | 571 | intron (NM_153068, intron 3 of 5) | Ehd2 | 4 | Untreated (DSB peaks) |
| 1-530 | + | 589 | intron (NM_026162, intron 1 of 13) | Plxdc2 | 4 | Untreated (DSB peaks) |
| 1-993 | + | 840 | promoter-TSS (NM_138303) | Yipf2 | 4 | Untreated (DSB peaks) |
| 31413 | + | 633 | promoter-TSS (NM_011295) | Rps12 | 4 | Untreated (DSB peaks) |
| 1-394 | + | 554 | intron (NM_178665, intron 3 of 12) | Morf4l1-ps1 | 4 | Untreated (DSB peaks) |
| 31778 | + | 588 | intron (NM_018747, intron 3 of 7) | Akap7 | 4 | Untreated (DSB peaks) |
| 1-514 | + | 592 | intron (NM_175459, intron 2 of 10) | D930032P07Rik | 4 | Untreated (DSB peaks) |
| 1-439 | + | 674 | promoter-TSS (NM_030561) | BC004004 | 4 | Untreated (DSB peaks) |
| 1-398 | + | 567 | promoter-TSS (NM_001199177) | Opa1 | 4 | Untreated (DSB peaks) |
| 1-441 | + | 724 | intron (NM_028902, intron 7 of 7) | Hsf2bp | 4 | Untreated (DSB peaks) |
| 1-313 | + | 594 | intron (NM_153800, intron 2 of 10) | Arhgap22 | 4 | Untreated (DSB peaks) |
| 1-141 | + | 595 | intron (NM_001190885, intron 1 of 8) | Gabrp | 4 | Untreated (DSB peaks) |

| | | | | | | |
|---|---|---|---|---|---|---|
| 1-444 | + | 715 | intron (NM_001204973, intron 1 of 12) | Brd2 | 4 | Untreated (DSB peaks) |
| 1-347 | + | 639 | promoter-TSS (NM_177151) | Vps13b | 4 | Untreated (DSB peaks) |
| 1-481 | + | 574 | intron (NM_026027, intron 3 of 3) | Pfdn1 | 4 | Untreated (DSB peaks) |
| 1-541 | + | 582 | promoter-TSS (NM_146253) | Zbtb6 | 4 | Untreated (DSB peaks) |
| 35431 | + | 643 | intron (NM_134000, intron 1 of 8) | Gm16364 | 4 | Untreated (DSB peaks) |
| 1-448 | + | 626 | intron (NM_181753, intron 5 of 6) | Opn5 | 4 | Untreated (DSB peaks) |
| 1-674 | + | 618 | promoter-TSS (NM_001190414) | Gne | 4 | Untreated (DSB peaks) |
| 1-1008 | + | 622 | promoter-TSS (NM_001357490) | Bcl9l | 4 | Untreated (DSB peaks) |
| 1-739 | + | 768 | intron (NM_001359071, intron 2 of 7) | Ldb2 | 4 | Untreated (DSB peaks) |
| 1-271 | + | 564 | intron (NM_001205043, intron 9 of 17) | Dtnbp1 | 4 | Untreated (DSB peaks) |
| 1-546 | + | 730 | intron (NM_028810, intron 2 of 4) | Rnd3 | 4 | Untreated (DSB peaks) |
| 1-551 | + | 593 | intron (NM_001037099, intron 9 of 13) | Arl5a | 4 | Untreated (DSB peaks) |
| 43101 | + | 1000 | promoter-TSS (NM_001302963) | Tyw5 | 4 | Untreated (DSB peaks) |
| 43831 | + | 560 | promoter-TSS (NM_009676) | Aox1 | 4 | Untreated (DSB peaks) |
| 1-681 | + | 577 | intron (NM_001163288, intron 13 of 16) | Gm12596 | 4 | Untreated (DSB peaks) |
| 1-1020 | + | 542 | promoter-TSS (NM_010193) | Fem1b | 4 | Untreated (DSB peaks) |
| 1-102 | + | 558 | intron (NM_001253857, intron 2 of 12) | Tet1 | 4 | Untreated (DSB peaks) |
| 1-684 | + | 554 | intron (NM_028326, intron 1 of 14) | Zfp618 | 4 | Untreated (DSB peaks) |
| 1-952 | + | 625 | Intergenic | Lsm6 | 4 | Untreated (DSB peaks) |
| 1-624 | + | 608 | promoter-TSS (NM_001331067) | Slc33a1 | 4 | Untreated (DSB peaks) |
| 1-285 | + | 566 | Intergenic | Nr2f1 | 4 | Untreated (DSB peaks) |
| 1-1025 | + | 668 | promoter-TSS (NM_175153) | Ints14 | 4 | Untreated (DSB peaks) |
| 1-495 | + | 648 | intron (NM_178793, intron 2 of 10) | Mir694 | 4 | Untreated (DSB peaks) |
| 1-107 | + | 581 | intron (NM_023598, intron 2 of 9) | Arid5b | 4 | Untreated (DSB peaks) |
| 1-164 | + | 574 | promoter-TSS (NM_001301710) | Mpdu1 | 4 | Untreated (DSB peaks) |
| 1-941 | + | 590 | TTS (NM_018827) | Rex1bd | 4 | Untreated (DSB peaks) |
| 1-816 | + | 579 | promoter-TSS (NM_024288) | Rmnd5a | 4 | Untreated (DSB peaks) |
| 1-950 | + | 718 | promoter-TSS (NM_023126) | Rab8a | 4 | Untreated (DSB peaks) |
| 1-561 | + | 588 | intron (NM_007389, intron 4 of 8) | Chrna1 | 4 | Untreated (DSB peaks) |
| 1-168 | + | 599 | intron (NM_008750, intron 1 of 7) | Mrm3 | 4 | Untreated (DSB peaks) |
| 1-371 | + | 551 | intron (NM_012053, intron 1 of 5) | Rpl8 | 4 | Untreated (DSB peaks) |
| 1-228 | + | 568 | intron (NM_177267, intron 6 of 8) | Mir1843a | 4 | Untreated (DSB peaks) |
| 1-691 | + | 556 | intron (NM_026821, intron 1 of 1) | Lurap1l | 4 | Untreated (DSB peaks) |
| 1-173 | + | 611 | promoter-TSS (NM_013654) | Ccl7 | 4 | Untreated (DSB peaks) |
| 1-174 | + | 611 | intron (NM_013654, intron 1 of 2) | Ccl7 | 4 | Untreated (DSB peaks) |
| 1-955 | + | 633 | promoter-TSS (NM_018771) | Gipc1 | 4 | Untreated (DSB peaks) |
| 1-562 | + | 795 | intron (NM_008402, intron 4 of 29) | Itgav | 4 | Untreated (DSB peaks) |
| 1-885 | + | 562 | 5' UTR (NM_023403, exon 1 of 3) | Mesd | 4 | Untreated (DSB peaks) |
| 1-118 | + | 556 | intron (NM_011595, intron 2 of 4) | Timp3 | 4 | Untreated (DSB peaks) |
| 1-120 | + | 640 | intron (NM_013722, intron 3 of 13) | Syn3 | 4 | Untreated (DSB peaks) |
| 1-181 | + | 546 | promoter-TSS (NM_024262) | Smg8 | 4 | Untreated (DSB peaks) |
| 1-958 | + | 556 | intron (NM_177224, intron 2 of 39) | Chd9 | 4 | Untreated (DSB peaks) |
| 1-752 | + | 725 | promoter-TSS (NM_183392) | Nup54 | 4 | Untreated (DSB peaks) |
| 14977 | + | 558 | promoter-TSS (NM_011796) | Capn10 | 4 | Untreated (DSB peaks) |
| 1-753 | + | 636 | intron (NM_001009818, intron 1 of 9) | Sept11 | 4 | Untreated (DSB peaks) |
| 1-122 | + | 605 | intron (NM_146239, intron 1 of 16) | Mir1931 | 4 | Untreated (DSB peaks) |
| 1-184 | + | 585 | intron (NM_001361708, intron 4 of 9) | Car10 | 4 | Untreated (DSB peaks) |

| | | | | | | |
|---|---|---|---|---|---|---|
| 1-564 | + | 705 | promoter-TSS (NM_026944) | Alkbh3 | 4 | Untreated (DSB peaks) |
| 1-697 | + | 567 | intron (NR_015513, intron 1 of 2) | Nfia | 4 | Untreated (DSB peaks) |
| 1-698 | + | 551 | intron (NM_001122952, intron 1 of 10) | Nfia | 4 | Untreated (DSB peaks) |
| 1-699 | + | 547 | intron (NM_001122952, intron 1 of 10) | Nfia | 4 | Untreated (DSB peaks) |
| 1-700 | + | 547 | intron (NM_001122952, intron 1 of 10) | Nfia | 4 | Untreated (DSB peaks) |
| 1-295 | + | 619 | intron (NM_173757, intron 4 of 10) | 6430562O15Rik | 4 | Untreated (DSB peaks) |
| 1-827 | + | 732 | intron (NM_001347345, intron 1 of 19) | Foxp1 | 4 | Untreated (DSB peaks) |
| 1-833 | + | 577 | intron (NM_018884, intron 1 of 9) | Pdzrn3 | 4 | Untreated (DSB peaks) |
| 1-378 | + | 567 | promoter-TSS (NM_153416) | Aaas | 4 | Untreated (DSB peaks) |
| 1-116 | + | 911 | Intergenic | 1500009L16Rik | 4 | Untreated (DSB peaks) |
| 1-902 | + | 650 | intron (NM_001001326, intron 1 of 19) | St5 | 4 | Untreated (DSB peaks) |
| 1-464 | + | 675 | Intergenic | 8430430B14Rik | 4 | Untreated (DSB peaks) |
| 1-971 | + | 833 | promoter-TSS (NM_009677) | Ap1g1 | 4 | Untreated (DSB peaks) |
| 1-694 | + | 770 | Intergenic | 6030471H07Rik | 4 | Untreated (DSB peaks) |
| 1-835 | + | 552 | intron (NM_080448, intron 1 of 21) | Srgap3 | 4 | Untreated (DSB peaks) |
| 1-702 | + | 616 | promoter-TSS (NM_011034) | Prdx1 | 4 | Untreated (DSB peaks) |
| 1-331 | + | 691 | intron (NM_134082, intron 2 of 26) | B930095G15Rik | 4 | Untreated (DSB peaks) |
| 1-1054 | + | 564 | intron (NM_177589, intron 13 of 36) | Ulk4 | 4 | Untreated (DSB peaks) |
| 17533 | + | 597 | promoter-TSS (NM_025860) | Ddx18 | 4 | Untreated (DSB peaks) |
| 1-127 | + | 575 | intron (NM_001110218, intron 6 of 9) | Mirlet7i | 4 | Untreated (DSB peaks) |
| 1-1055 | + | 560 | intron (NM_009544, intron 1 of 3) | Zfp105 | 4 | Untreated (DSB peaks) |
| 1-976 | + | 901 | promoter-TSS (NM_001281803) | Def8 | 4 | Untreated (DSB peaks) |
| 1-977 | + | 555 | promoter-TSS (NM_172288) | Nup133 | 4 | Untreated (DSB peaks) |
| 1-846 | + | 586 | intron (NR_149320, intron 4 of 11) | Emg1 | 4 | Untreated (DSB peaks) |
| 17899 | + | 590 | promoter-TSS (NM_145128) | Mgat5 | 4 | Untreated (DSB peaks) |
| 1-913 | + | 685 | promoter-TSS (NM_172255) | Wdr11 | 4 | Untreated (DSB peaks) |
| 1-915 | + | 760 | intron (NM_201601, intron 4 of 15) | Fgfr2 | 4 | Untreated (DSB peaks) |
| 1-646 | + | 600 | promoter-TSS (NM_026231) | Gstcd | 4 | Untreated (DSB peaks) |
| 1-712 | + | 569 | promoter-TSS (NM_207237) | Man1c1 | 4 | Untreated (DSB peaks) |
| 18629 | + | 648 | promoter-TSS (NR_045325) | Gm19705 | 4 | Untreated (DSB peaks) |
| 1-791 | + | 568 | intron (NM_001013774, intron 11 of 12) | Kpna7 | 4 | Untreated (DSB peaks) |
| 1-722 | + | 645 | promoter-TSS (NR_040649) | Errfi1 | 4 | Untreated (DSB peaks) |
| 1-794 | + | 752 | intron (NR_045700, intron 1 of 3) | 1700028E10Rik | 4 | Untreated (DSB peaks) |
| 1-580 | + | 540 | intron (NM_025876, intron 1 of 13) | Cdk5rap1 | 4 | Untreated (DSB peaks) |
| 1-581 | + | 540 | promoter-TSS (NM_025876) | Cdk5rap1 | 4 | Untreated (DSB peaks) |
| 1-659 | + | 580 | promoter-TSS (NM_028938) | Fpgt | 4 | Untreated (DSB peaks) |
| 1-589 | + | 1000 | promoter-TSS (NM_172675) | Stx16 | 4 | Untreated (DSB peaks) |
| 25569 | + | 699 | promoter-TSS (NM_145943) | Sde2 | 4 | Untreated (DSB peaks) |
| 26665 | + | 592 | intron (NM_028408, intron 3 of 5) | Ccdc121 | 4 | Untreated (DSB peaks) |
| 1-422 | + | 713 | Intergenic | Mrps6 | 4 | Untreated (DSB peaks) |
| 1-291 | + | 599 | Intergenic | Gm4814 | 4 | Untreated (DSB peaks) |
| 1-831 | + | 656 | Intergenic | Rybp | 4 | Untreated (DSB peaks) |
| 1-240 | + | 599 | Intergenic | Kif26a | 4 | Untreated (DSB peaks) |
| 1-839 | + | 623 | Intergenic | Cxcl12 | 4 | Untreated (DSB peaks) |
| 1-840 | + | 677 | Intergenic | Cxcl12 | 4 | Untreated (DSB peaks) |
| 1-842 | + | 604 | Intergenic | Cxcl12 | 4 | Untreated (DSB peaks) |
| 1-771 | + | 765 | Intergenic | Ncor2 | 4 | Untreated (DSB peaks) |

| | | | | | | |
|---|---|---|---|---|---|---|
| 1-982 | + | 541 | Intergenic | Irf2bp2 | 4 | Untreated (DSB peaks) |
| 1-985 | + | 895 | Intergenic | Mir21c | 4 | Untreated (DSB peaks) |
| 1-573 | + | 700 | Intergenic | Tmem239 | 4 | Untreated (DSB peaks) |
| 1-780 | + | 626 | Intergenic | Gatsl2 | 4 | Untreated (DSB peaks) |
| 1-788 | + | 601 | Intergenic | Foxk1 | 4 | Untreated (DSB peaks) |
| 1-576 | + | 611 | Intergenic | Gm5535 | 4 | Untreated (DSB peaks) |
| 1-656 | + | 785 | Intergenic | Adgrl2 | 4 | Untreated (DSB peaks) |
| 27030 | + | 565 | Intergenic | A730004F24Rik | 4 | Untreated (DSB peaks) |
| 27760 | + | 698 | Intergenic | Hhat | 4 | Untreated (DSB peaks) |
| 1-338 | + | 930 | Intergenic | Osmr | 3 | Untreated (DSB peaks) |
| 1-592 | + | 657 | Intergenic | Pag1 | 3 | Untreated (DSB peaks) |
| 29221 | + | 650 | Intergenic | Adgb | 3 | Untreated (DSB peaks) |
| 1-598 | + | 668 | Intergenic | Tbl1xr1 | 3 | Untreated (DSB peaks) |
| 1-253 | + | 574 | Intergenic | Abt1 | 3 | Untreated (DSB peaks) |
| 1-261 | + | 904 | Intergenic | Prl5a1 | 3 | Untreated (DSB peaks) |
| 1-669 | + | 707 | Intergenic | Bach2 | 3 | Untreated (DSB peaks) |
| 1-262 | + | 572 | Intergenic | Gmds | 3 | Untreated (DSB peaks) |
| 1-478 | + | 849 | Intergenic | Nrep | 3 | Untreated (DSB peaks) |
| 1-314 | + | 598 | Intergenic | 4930503F20Rik | 3 | Untreated (DSB peaks) |
| 1-264 | + | 606 | Intergenic | Ly86 | 3 | Untreated (DSB peaks) |
| 1-611 | + | 555 | Intergenic | Fat4 | 3 | Untreated (DSB peaks) |
| 1-268 | + | 540 | Intergenic | Tmem170b | 3 | Untreated (DSB peaks) |
| 1-405 | + | 607 | Intergenic | Myh15 | 3 | Untreated (DSB peaks) |
| 1-549 | + | 606 | Intergenic | Gm13497 | 3 | Untreated (DSB peaks) |
| 1-1011 | + | 550 | Intergenic | 1810046K07Rik | 3 | Untreated (DSB peaks) |
| 1-1012 | + | 554 | Intergenic | 1810046K07Rik | 3 | Untreated (DSB peaks) |
| 1-275 | + | 656 | Intergenic | 4921525O09Rik | 3 | Untreated (DSB peaks) |
| 1-486 | + | 609 | Intergenic | 9330117O12Rik | 3 | Untreated (DSB peaks) |
| 1-619 | + | 576 | Intergenic | Tm4sf1 | 3 | Untreated (DSB peaks) |
| 1-621 | + | 756 | Intergenic | 1700007F19Rik | 3 | Untreated (DSB peaks) |
| 1-622 | + | 602 | Intergenic | Igsf10 | 3 | Untreated (DSB peaks) |
| 1-683 | + | 553 | Intergenic | Rgs3 | 3 | Untreated (DSB peaks) |
| 1-160 | + | 553 | Intergenic | Aurkb | 3 | Untreated (DSB peaks) |
| 1-217 | + | 613 | Intergenic | 5830428M24Rik | 3 | Untreated (DSB peaks) |
| 1-880 | + | 644 | Intergenic | Nr2f2 | 3 | Untreated (DSB peaks) |
| 1-626 | + | 574 | Intergenic | Ppid | 3 | Untreated (DSB peaks) |
| 1-628 | + | 602 | Intergenic | Pdgfc | 3 | Untreated (DSB peaks) |
| 1-418 | + | 627 | Intergenic | Hunk | 3 | Untreated (DSB peaks) |
| 1-525 | + | 661 | intron (NM_025626, intron 1 of 3) | Fam107b | 3 | Untreated (DSB peaks) |
| 1-425 | + | 589 | intron (NM_001085355, intron 4 of 19) | Arid1b | 3 | Untreated (DSB peaks) |
| 1-528 | + | 571 | intron (NM_181399, intron 1 of 13) | Usp6nl | 3 | Untreated (DSB peaks) |
| 1-383 | + | 630 | promoter-TSS (NM_146067) | Cpped1 | 3 | Untreated (DSB peaks) |
| 1-135 | + | 584 | intron (NM_001177629, intron 10 of 15) | Ddc | 3 | Untreated (DSB peaks) |
| 1-529 | + | 576 | intron (NM_026162, intron 1 of 13) | Plxdc2 | 3 | Untreated (DSB peaks) |
| 1-305 | + | 881 | intron (NM_001289761, intron 2 of 8) | Rarb | 3 | Untreated (DSB peaks) |
| 1-729 | + | 586 | promoter-TSS (NM_001356591) | Ptpn12 | 3 | Untreated (DSB peaks) |
| 1-434 | + | 635 | promoter-TSS (NM_026686) | Mettl26 | 3 | Untreated (DSB peaks) |

| | | | | | | |
|---|---|---|---|---|---|---|
| 1-435 | + | 540 | promoter-TSS (NM_026686) | Mettl26 | 3 | Untreated (DSB peaks) |
| 1-538 | + | 608 | 5' UTR (NM_001290484, exon 1 of 12) | Gtf3c5 | 3 | Untreated (DSB peaks) |
| 1-211 | + | 617 | intron (NM_028139, intron 2 of 3) | Atxn7l1 | 3 | Untreated (DSB peaks) |
| 1-806 | + | 823 | intron (NM_194061, intron 15 of 19) | Atp6v0a4 | 3 | Untreated (DSB peaks) |
| 1-807 | + | 711 | promoter-TSS (NM_153600) | Ttc26 | 3 | Untreated (DSB peaks) |
| 1-266 | + | 724 | promoter-TSS (NM_011547) | Tfap2a | 3 | Untreated (DSB peaks) |
| 1-267 | + | 610 | promoter-TSS (NM_023887) | Gcnt2 | 3 | Untreated (DSB peaks) |
| 1-519 | + | 726 | promoter-TSS (NM_025530) | Cox15 | 3 | Untreated (DSB peaks) |
| 1-452 | + | 615 | intron (NM_028751, intron 2 of 10) | Tjap1 | 3 | Untreated (DSB peaks) |
| 1-811 | + | 613 | promoter-TSS (NM_027477) | Zfp398 | 3 | Untreated (DSB peaks) |
| 1-1009 | + | 598 | promoter-TSS (NM_001113204) | Ncam1 | 3 | Untreated (DSB peaks) |
| 1-349 | + | 829 | intron (NM_001310481, intron 4 of 5) | Trps1 | 3 | Untreated (DSB peaks) |
| 1-350 | + | 709 | intron (NM_001310481, intron 4 of 5) | Trps1 | 3 | Untreated (DSB peaks) |
| 1-742 | + | 684 | exon (NM_024213, exon 1 of 28) | Anapc4 | 3 | Untreated (DSB peaks) |
| 1-553 | + | 571 | promoter-TSS (NM_022989) | Prpf40a | 3 | Untreated (DSB peaks) |
| 42370 | + | 672 | intron (NM_001001883, intron 28 of 28) | Mir7681 | 3 | Untreated (DSB peaks) |
| 36161 | + | 616 | intron (NM_001033259, intron 1 of 7) | Mcu | 3 | Untreated (DSB peaks) |
| 1-100 | + | 581 | promoter-TSS (NM_001291443) | Micu1 | 3 | Untreated (DSB peaks) |
| 1-634 | + | 540 | Intergenic | Terc | 3 | Untreated (DSB peaks) |
| 1-362 | + | 556 | intron (NR_132746, intron 1 of 6) | Pvt1 | 3 | Untreated (DSB peaks) |
| 45292 | + | 630 | promoter-TSS (NM_010939) | Nrp2 | 3 | Untreated (DSB peaks) |
| 1-410 | + | 579 | promoter-TSS (NM_010140) | Epha3 | 3 | Untreated (DSB peaks) |
| 46753 | + | 559 | intron (NM_001039493, intron 7 of 7) | Fzd5 | 3 | Untreated (DSB peaks) |
| 1-687 | + | 609 | intron (NM_207109, intron 5 of 22) | Astn2 | 3 | Untreated (DSB peaks) |
| 1-746 | + | 557 | intron (NM_001310626, intron 1 of 15) | Apbb2 | 3 | Untreated (DSB peaks) |
| 1-455 | + | 750 | intron (NM_001357625, intron 1 of 32) | Ptprm | 3 | Untreated (DSB peaks) |
| 1-943 | + | 583 | 5' UTR (NM_010592, exon 1 of 1) | Jund | 3 | Untreated (DSB peaks) |
| 1-111 | + | 602 | intron (NM_031397, intron 3 of 20) | Bicc1 | 3 | Untreated (DSB peaks) |
| 1-947 | + | 541 | intron (NM_001001491, intron 1 of 7) | Tpm4 | 3 | Untreated (DSB peaks) |
| 1-368 | + | 555 | intron (NM_172121, intron 3 of 11) | Zc3h3 | 3 | Untreated (DSB peaks) |
| 1-370 | + | 918 | promoter-TSS (NM_001164173) | Cpsf1 | 3 | Untreated (DSB peaks) |
| 1-953 | + | 725 | promoter-TSS (NM_178267) | Zfp827 | 3 | Untreated (DSB peaks) |
| 1-225 | + | 728 | intron (NM_009014, intron 7 of 10) | Rad51b | 3 | Untreated (DSB peaks) |
| 1-882 | + | 668 | promoter-TSS (NM_009682) | Ap3s2 | 3 | Untreated (DSB peaks) |
| 1-287 | + | 716 | promoter-TSS (NR_015587) | 9330111N05Rik | 3 | Untreated (DSB peaks) |
| 12055 | + | 860 | 5' UTR (NM_010570, exon 1 of 2) | Irs1 | 3 | Untreated (DSB peaks) |
| 1-229 | + | 874 | promoter-TSS (NM_027349) | Rbm25 | 3 | Untreated (DSB peaks) |
| 1-176 | + | 593 | promoter-TSS (NM_019816) | Aatf | 3 | Untreated (DSB peaks) |
| 1-695 | + | 747 | intron (NM_172870, intron 2 of 5) | Bnc2 | 3 | Untreated (DSB peaks) |
| 1-469 | + | 749 | intron (NM_029858, intron 1 of 3) | Ston1 | 3 | Untreated (DSB peaks) |
| 1-419 | + | 646 | intron (NM_001316761, intron 1 of 5) | Eva1c | 3 | Untreated (DSB peaks) |
| 1-823 | + | 629 | intron (NM_009320, intron 5 of 14) | Slc6a6 | 3 | Untreated (DSB peaks) |
| 1-960 | + | 546 | promoter-TSS (NM_008187) | Cfap20 | 3 | Untreated (DSB peaks) |
| 1-696 | + | 559 | intron (NR_015513, intron 1 of 2) | Nfia | 3 | Untreated (DSB peaks) |
| 1-187 | + | 584 | promoter-TSS (NM_001163307) | Pcgf2 | 3 | Untreated (DSB peaks) |
| 1-756 | + | 590 | intron (NM_133738, intron 12 of 16) | Antxr2 | 3 | Untreated (DSB peaks) |
| 1-292 | + | 750 | promoter-TSS (NM_172591) | Fcho2 | 3 | Untreated (DSB peaks) |

| | | | | | | |
|---|---|---|---|---|---|---|
| 1-330 | + | 541 | promoter-TSS (NM_207215) | Mycbp2 | 3 | Untreated (DSB peaks) |
| 1-968 | + | 570 | promoter-TSS (NM_133957) | Nfat5 | 3 | Untreated (DSB peaks) |
| 1-640 | + | 558 | intron (NM_001163348, intron 2 of 6) | Ntng1 | 3 | Untreated (DSB peaks) |
| 1-907 | + | 569 | promoter-TSS (NM_177249) | Usp47 | 3 | Untreated (DSB peaks) |
| 1-1040 | + | 577 | Intergenic | Anapc13 | 3 | Untreated (DSB peaks) |
| 1-298 | + | 543 | promoter-TSS (NM_008247) | Plpp1 | 3 | Untreated (DSB peaks) |
| 1-299 | + | 543 | 5' UTR (NM_008247, exon 1 of 6) | Plpp1 | 3 | Untreated (DSB peaks) |
| 1-961 | + | 575 | Intergenic | 1600027J07Rik | 3 | Untreated (DSB peaks) |
| 1-838 | + | 607 | promoter-TSS (NM_177683) | Vgll4 | 3 | Untreated (DSB peaks) |
| 1-844 | + | 594 | promoter-TSS (NM_194339) | Bms1 | 3 | Untreated (DSB peaks) |
| 1-332 | + | 691 | intron (NM_134082, intron 2 of 26) | B930095G15Rik | 3 | Untreated (DSB peaks) |
| 1-963 | + | 653 | Intergenic | 4933400L20Rik | 3 | Untreated (DSB peaks) |
| 1-845 | + | 577 | promoter-TSS (NM_011909) | Usp18 | 3 | Untreated (DSB peaks) |
| 1-975 | + | 549 | TTS (NR_015497) | 9330133O14Rik | 3 | Untreated (DSB peaks) |
| 1-128 | + | 678 | intron (NM_001110218, intron 6 of 9) | Mirlet7i | 3 | Untreated (DSB peaks) |
| 1-334 | + | 555 | intron (NM_177393, intron 7 of 43) | Nalcn | 3 | Untreated (DSB peaks) |
| 1-131 | + | 594 | promoter-TSS (NR_106183) | Mir8105 | 3 | Untreated (DSB peaks) |
| 1-572 | + | 683 | promoter-TSS (NM_009086) | Polr1b | 3 | Untreated (DSB peaks) |
| 1-781 | + | 600 | intron (NM_001359066, intron 1 of 32) | Gtf2i | 3 | Untreated (DSB peaks) |
| 1-647 | + | 621 | promoter-TSS (NM_177665) | Gm4861 | 3 | Untreated (DSB peaks) |
| 1-579 | + | 609 | intron (NM_053195, intron 2 of 16) | Slc24a3 | 3 | Untreated (DSB peaks) |
| 22647 | + | 580 | intron (NM_001305643, intron 19 of 19) | Scyl3 | 3 | Untreated (DSB peaks) |
| 1-585 | + | 640 | promoter-TSS (NM_023815) | Trp53rkb | 3 | Untreated (DSB peaks) |
| 1-762 | + | 624 | Intergenic | C130026L21Rik | 3 | Untreated (DSB peaks) |
| 1-984 | + | 567 | Intergenic | Pard3 | 3 | Untreated (DSB peaks) |
| 1-852 | + | 553 | Intergenic | 2310001H17Rik | 3 | Untreated (DSB peaks) |
| 1-779 | + | 575 | Intergenic | Gatsl2 | 3 | Untreated (DSB peaks) |
| 1-855 | + | 694 | Intergenic | Atf7ip | 3 | Untreated (DSB peaks) |
| 1-856 | + | 554 | Intergenic | Atf7ip | 3 | Untreated (DSB peaks) |
| 1-575 | + | 611 | Intergenic | Gm5535 | 3 | Untreated (DSB peaks) |
| 1-657 | + | 546 | Intergenic | Adgrl4 | 3 | Untreated (DSB peaks) |
| 1-725 | + | 541 | Intergenic | Mxra8 | 3 | Untreated (DSB peaks) |
| 28491 | + | 554 | non-coding (NR_015566, exon 8 of 8) | Mir29b-2 | 3 | Untreated (DSB peaks) |
| 1-798 | + | 561 | Intergenic | Col1a2 | 2 | Untreated (DSB peaks) |
| 1-302 | + | 676 | Intergenic | Flnb | 2 | Untreated (DSB peaks) |
| 1-303 | + | 685 | Intergenic | Mnd1-ps | 2 | Untreated (DSB peaks) |
| 43132 | + | 560 | Intergenic | Eya1 | 2 | Untreated (DSB peaks) |
| 1-254 | + | 574 | Intergenic | Abt1 | 2 | Untreated (DSB peaks) |
| 1-255 | + | 564 | Intergenic | C230035I16Rik | 2 | Untreated (DSB peaks) |
| 1-670 | + | 708 | Intergenic | Bach2 | 2 | Untreated (DSB peaks) |
| 1-315 | + | 598 | Intergenic | 4930503F20Rik | 2 | Untreated (DSB peaks) |
| 1-480 | + | 638 | Intergenic | Hspa9 | 2 | Untreated (DSB peaks) |
| 1-805 | + | 717 | Intergenic | Akr1d1 | 2 | Untreated (DSB peaks) |
| 1-542 | + | 543 | Intergenic | Crb2 | 2 | Untreated (DSB peaks) |
| 1-999 | + | 548 | Intergenic | Mir125b-1 | 2 | Untreated (DSB peaks) |
| 1-144 | + | 610 | Intergenic | Gm12159 | 2 | Untreated (DSB peaks) |
| 1-547 | + | 661 | Intergenic | Gm13497 | 2 | Untreated (DSB peaks) |

| | | | | | | |
|---|---|---|---|---|---|---|
| 1-548 | + | 652 | Intergenic | Gm13497 | 2 | Untreated (DSB peaks) |
| 1-273 | + | 743 | Intergenic | Hist1h2al | 2 | Untreated (DSB peaks) |
| 1-875 | + | 550 | Intergenic | Gas2 | 2 | Untreated (DSB peaks) |
| 1-876 | + | 630 | Intergenic | Atp10a | 2 | Untreated (DSB peaks) |
| 1-680 | + | 689 | Intergenic | Gm12596 | 2 | Untreated (DSB peaks) |
| 44562 | + | 562 | Intergenic | Gm973 | 2 | Untreated (DSB peaks) |
| 1-360 | + | 578 | Intergenic | Gm19510 | 2 | Untreated (DSB peaks) |
| 1-558 | + | 578 | Intergenic | Tank | 2 | Untreated (DSB peaks) |
| 1-559 | + | 578 | Intergenic | Tank | 2 | Untreated (DSB peaks) |
| 1-1021 | + | 693 | Intergenic | Cln6 | 2 | Untreated (DSB peaks) |
| 1-494 | + | 712 | Intergenic | Piezo2 | 2 | Untreated (DSB peaks) |
| 1-325 | + | 662 | Intergenic | Npm2 | 2 | Untreated (DSB peaks) |
| 1-326 | + | 662 | Intergenic | Npm2 | 2 | Untreated (DSB peaks) |
| 1-170 | + | 604 | Intergenic | Mir193a | 2 | Untreated (DSB peaks) |
| 1-175 | + | 559 | Intergenic | Slfn14 | 2 | Untreated (DSB peaks) |
| 1-1036 | + | 565 | Intergenic | Bckdhb | 2 | Untreated (DSB peaks) |
| 1-288 | + | 647 | Intergenic | Tmem161b | 2 | Untreated (DSB peaks) |
| 1-417 | + | 627 | Intergenic | Hunk | 2 | Untreated (DSB peaks) |
| 1-959 | + | 633 | Intergenic | Irx6 | 2 | Untreated (DSB peaks) |
| 1-633 | + | 540 | Intergenic | Terc | 2 | Untreated (DSB peaks) |
| 1-335 | + | 547 | promoter-TSS (NR_028266) | BC037032 | 2 | Untreated (DSB peaks) |
| 1-727 | + | 626 | 5' UTR (NM_020010, exon 1 of 10) | Cyp51 | 2 | Untreated (DSB peaks) |
| 1-527 | + | 727 | intron (NM_172475, intron 1 of 22) | Frmd4a | 2 | Untreated (DSB peaks) |
| 1-426 | + | 589 | intron (NM_001085355, intron 4 of 19) | Arid1b | 2 | Untreated (DSB peaks) |
| 1-662 | + | 620 | exon (NM_023066, exon 1 of 24) | Asph | 2 | Untreated (DSB peaks) |
| 1-124 | + | 660 | Intergenic | Acss3 | 2 | Untreated (DSB peaks) |
| 1-136 | + | 613 | intron (NM_001177629, intron 4 of 15) | Ddc | 2 | Untreated (DSB peaks) |
| 1-989 | + | 652 | promoter-TSS (NM_144933) | Med17 | 2 | Untreated (DSB peaks) |
| 1-666 | + | 631 | intron (NM_013752, intron 1 of 15) | Nbn | 2 | Untreated (DSB peaks) |
| 1-834 | + | 554 | Intergenic | 0610040F04Rik | 2 | Untreated (DSB peaks) |
| 1-925 | + | 978 | intron (NM_001270553, intron 6 of 9) | Gm21119 | 2 | Untreated (DSB peaks) |
| 1-991 | + | 552 | promoter-TSS (NM_001004190) | Zfp560 | 2 | Untreated (DSB peaks) |
| 1-308 | + | 616 | exon (NM_145459, exon 3 of 3) | Zfp503 | 2 | Untreated (DSB peaks) |
| 1-477 | + | 638 | promoter-TSS (NM_011755) | Zfp35 | 2 | Untreated (DSB peaks) |
| 1-396 | + | 808 | intron (NM_178665, intron 5 of 12) | Morf4l1-ps1 | 2 | Untreated (DSB peaks) |
| 1-733 | + | 625 | promoter-TSS (NR_027388) | 1700096K18Rik | 2 | Untreated (DSB peaks) |
| 1-536 | + | 942 | intron (NM_008714, intron 2 of 33) | Notch1 | 2 | Untreated (DSB peaks) |
| 1-537 | + | 579 | promoter-TSS (NM_001360790) | Brd3os | 2 | Untreated (DSB peaks) |
| 1-903 | + | 678 | Intergenic | Swap70 | 2 | Untreated (DSB peaks) |
| 1-344 | + | 598 | intron (NM_176073, intron 4 of 7) | 1700084J12Rik | 2 | Untreated (DSB peaks) |
| 1-442 | + | 582 | TTS (NM_145486) | Hnrnpm | 2 | Untreated (DSB peaks) |
| 1-737 | + | 565 | intron (NM_030889, intron 1 of 26) | Mir7689 | 2 | Untreated (DSB peaks) |
| 1-738 | + | 565 | intron (NM_030889, intron 1 of 26) | Mir7689 | 2 | Untreated (DSB peaks) |
| 1-933 | + | 821 | promoter-TSS (NM_015802) | Dlc1 | 2 | Untreated (DSB peaks) |
| 33239 | + | 776 | promoter-TSS (NM_008538) | Marcks | 2 | Untreated (DSB peaks) |
| 1-348 | + | 547 | promoter-TSS (NM_008388) | Eif3e | 2 | Untreated (DSB peaks) |
| 1-317 | + | 582 | promoter-TSS (NM_027492) | Naa30 | 2 | Untreated (DSB peaks) |

| | | | | | | |
|---|---|---|---|---|---|---|
| 1-545 | + | 587 | intron (NM_028810, intron 2 of 4) | Rnd3 | 2 | Untreated (DSB peaks) |
| 1-353 | + | 595 | exon (NM_030132, exon 1 of 3) | Utp23 | 2 | Untreated (DSB peaks) |
| 1-741 | + | 684 | promoter-TSS (NM_024213) | Anapc4 | 2 | Untreated (DSB peaks) |
| 1-552 | + | 645 | intron (NM_172409, intron 1 of 26) | Fmnl2 | 2 | Untreated (DSB peaks) |
| 42736 | + | 651 | promoter-TSS (NM_001033194) | Gtf3c3 | 2 | Untreated (DSB peaks) |
| 1-487 | + | 541 | exon (NM_175751, exon 2 of 10) | Zfp608 | 2 | Untreated (DSB peaks) |
| 1-676 | + | 565 | intron (NM_172867, intron 7 of 12) | Zfp462 | 2 | Untreated (DSB peaks) |
| 1-523 | + | 556 | intron (NM_181415, intron 26 of 28) | Atrnl1 | 2 | Untreated (DSB peaks) |
| 1-1016 | + | 689 | promoter-TSS (NM_019927) | Arih1 | 2 | Untreated (DSB peaks) |
| 1-283 | + | 600 | intron (NM_001289926, intron 13 of 15) | 2010111I01Rik | 2 | Untreated (DSB peaks) |
| 1-411 | + | 579 | promoter-TSS (NM_010140) | Epha3 | 2 | Untreated (DSB peaks) |
| 1-103 | + | 683 | intron (NM_178678, intron 2 of 2) | Lrrtm3 | 2 | Untreated (DSB peaks) |
| 1-686 | + | 557 | intron (NM_207109, intron 20 of 22) | Trim32 | 2 | Untreated (DSB peaks) |
| 1-412 | + | 666 | promoter-TSS (NM_028572) | Vgll3 | 2 | Untreated (DSB peaks) |
| 1-1026 | + | 736 | promoter-TSS (NM_001170907) | Trip4 | 2 | Untreated (DSB peaks) |
| 1-163 | + | 573 | promoter-TSS (NM_001301710) | Mpdu1 | 2 | Untreated (DSB peaks) |
| 1-944 | + | 583 | exon (NM_010592, exon 1 of 1) | Jund | 2 | Untreated (DSB peaks) |
| 1-945 | + | 555 | promoter-TSS (NM_145624) | Zfp709 | 2 | Untreated (DSB peaks) |
| 1-820 | + | 541 | intron (NM_001079822, intron 3 of 12) | Gm15401 | 2 | Untreated (DSB peaks) |
| 1-457 | + | 642 | promoter-TSS (NM_026417) | Yipf4 | 2 | Untreated (DSB peaks) |
| 1-458 | + | 642 | exon (NM_026417, exon 1 of 6) | Yipf4 | 2 | Untreated (DSB peaks) |
| 1-496 | + | 775 | promoter-TSS (NM_001042660) | Gm20544 | 2 | Untreated (DSB peaks) |
| 1-167 | + | 599 | intron (NM_008750, intron 1 of 7) | Mrm3 | 2 | Untreated (DSB peaks) |
| 1-502 | + | 671 | intron (NM_207651, intron 2 of 21) | Slc14a2 | 2 | Untreated (DSB peaks) |
| 1-221 | + | 563 | intron (NM_009014, intron 7 of 10) | Rad51b | 2 | Untreated (DSB peaks) |
| 1-222 | + | 560 | intron (NM_009014, intron 7 of 10) | Rad51b | 2 | Untreated (DSB peaks) |
| 1-224 | + | 815 | intron (NM_009014, intron 7 of 10) | Rad51b | 2 | Untreated (DSB peaks) |
| 1-227 | + | 568 | intron (NM_177267, intron 6 of 8) | Mir1843a | 2 | Untreated (DSB peaks) |
| 1-692 | + | 556 | intron (NM_026821, intron 1 of 1) | Lurap1l | 2 | Untreated (DSB peaks) |
| 1-627 | + | 545 | intron (NM_019971, intron 1 of 5) | Pdgfc | 2 | Untreated (DSB peaks) |
| 1-373 | + | 679 | intron (NM_001082536, intron 1 of 13) | Mkl1 | 2 | Untreated (DSB peaks) |
| 1-629 | + | 551 | promoter-TSS (NM_177260) | Tmem154 | 2 | Untreated (DSB peaks) |
| 1-413 | + | 626 | promoter-TSS (NM_017404) | Mrpl39 | 2 | Untreated (DSB peaks) |
| 1-1038 | + | 773 | promoter-TSS (NM_024195) | Cyb5r4 | 2 | Untreated (DSB peaks) |
| 1-375 | + | 674 | exon (NM_027081, exon 1 of 20) | Dennd6b | 2 | Untreated (DSB peaks) |
| 1-182 | + | 587 | promoter-TSS (NM_008711) | Nog | 2 | Untreated (DSB peaks) |
| 14246 | + | 590 | promoter-TSS (NM_133805) | Cops8 | 2 | Untreated (DSB peaks) |
| 1-420 | + | 762 | intron (NM_027627, intron 4 of 6) | Eva1c | 2 | Untreated (DSB peaks) |
| 1-293 | + | 658 | promoter-TSS (NM_001048267) | Tnpo1 | 2 | Untreated (DSB peaks) |
| 1-900 | + | 548 | 5' UTR (NM_001362166, exon 1 of 19) | Gdpd5 | 2 | Untreated (DSB peaks) |
| 1-233 | + | 544 | intron (NM_183186, intron 1 of 5) | Foxn3 | 2 | Untreated (DSB peaks) |
| 1-236 | + | 723 | intron (NM_028446, intron 16 of 20) | Fbln5 | 2 | Untreated (DSB peaks) |
| 1-193 | + | 565 | promoter-TSS (NM_027658) | Hexim2 | 2 | Untreated (DSB peaks) |
| 1-566 | + | 571 | intron (NM_178890, intron 1 of 16) | Abtb2 | 2 | Untreated (DSB peaks) |
| 1-296 | + | 774 | promoter-TSS (NM_026072) | Srek1ip1 | 2 | Untreated (DSB peaks) |
| 1-297 | + | 594 | promoter-TSS (NM_026072) | Srek1ip1 | 2 | Untreated (DSB peaks) |
| 1-1044 | + | 755 | exon (NM_053159, exon 1 of 10) | Mrpl3 | 2 | Untreated (DSB peaks) |

| | | | | | | |
|---|---|---|---|---|---|---|
| 1-966 | + | 558 | exon (NM_001271915, exon 1 of 1) | B3gnt9 | 2 | Untreated (DSB peaks) |
| 1-701 | + | 603 | promoter-TSS (NM_028209) | Ttc4 | 2 | Untreated (DSB peaks) |
| 1-906 | + | 569 | promoter-TSS (NM_177249) | Usp47 | 2 | Untreated (DSB peaks) |
| 1-836 | + | 552 | intron (NM_080448, intron 1 of 21) | Srgap3 | 2 | Untreated (DSB peaks) |
| 1-766 | + | 648 | promoter-TSS (NM_001033202) | Usp30 | 2 | Untreated (DSB peaks) |
| 1-837 | + | 647 | intron (NM_001253718, intron 18 of 19) | Atg7 | 2 | Untreated (DSB peaks) |
| 1-641 | + | 550 | promoter-TSS (NM_011693) | Vcam1 | 2 | Untreated (DSB peaks) |
| 1-241 | + | 641 | promoter-TSS (NM_028731) | Esyt2 | 2 | Untreated (DSB peaks) |
| 1-704 | + | 548 | promoter-TSS (NM_001355214) | Ppcs | 2 | Untreated (DSB peaks) |
| 1-705 | + | 884 | promoter-TSS (NM_172699) | Foxj3 | 2 | Untreated (DSB peaks) |
| 1-202 | + | 593 | promoter-TSS (NM_027745) | Ccdc57 | 2 | Untreated (DSB peaks) |
| 1-333 | + | 555 | intron (NM_177393, intron 7 of 43) | Nalcn | 2 | Untreated (DSB peaks) |
| 1-773 | + | 661 | 5' UTR (NM_019639, exon 1 of 2) | Ubc | 2 | Untreated (DSB peaks) |
| 1-570 | + | 570 | intron (NM_001284410, intron 1 of 3) | Bcl2l11 | 2 | Untreated (DSB peaks) |
| 1-914 | + | 686 | promoter-TSS (NM_172255) | Wdr11 | 2 | Untreated (DSB peaks) |
| 1-916 | + | 560 | intron (NM_019564, intron 3 of 8) | Htra1 | 2 | Untreated (DSB peaks) |
| 1-778 | + | 548 | intron (NM_177047, intron 5 of 19) | Gm29681 | 2 | Untreated (DSB peaks) |
| 1-645 | + | 600 | promoter-TSS (NM_027927) | Ints12 | 2 | Untreated (DSB peaks) |
| 1-715 | + | 570 | intron (NM_008546, intron 1 of 8) | Mfap2 | 2 | Untreated (DSB peaks) |
| 1-786 | + | 595 | intron (NM_177879, intron 1 of 44) | Sdk1 | 2 | Untreated (DSB peaks) |
| 1-650 | + | 706 | promoter-TSS (NM_001282042) | Sh3glb1 | 2 | Untreated (DSB peaks) |
| 1-578 | + | 687 | intron (NM_001356497, intron 1 of 16) | Slc24a3 | 2 | Untreated (DSB peaks) |
| 1-720 | + | 584 | promoter-TSS (NM_027873) | Ubiad1 | 2 | Untreated (DSB peaks) |
| 21186 | + | 643 | intron (NM_001038619, intron 18 of 20) | 2810442N19Rik | 2 | Untreated (DSB peaks) |
| 23377 | + | 608 | intron (NM_001305643, intron 19 of 19) | Scyl3 | 2 | Untreated (DSB peaks) |
| 24838 | + | 735 | intron (NM_019445, intron 6 of 17) | Fmn2 | 2 | Untreated (DSB peaks) |
| 1-655 | + | 614 | Intergenic | Adgrl2 | 2 | Untreated (DSB peaks) |
| 20455 | + | 616 | Intergenic | Stx6 | 2 | Untreated (DSB peaks) |
| 1-304 | + | 685 | Intergenic | Mnd1-ps | 1 | Untreated (DSB peaks) |
| 1 | + | 649 | Intergenic | Mir6341 | 1 | Untreated (DSB peaks) |
| 1-428 | + | 634 | Intergenic | Mir99b | 1 | Untreated (DSB peaks) |
| 1-533 | + | 618 | Intergenic | Gm3363 | 1 | Untreated (DSB peaks) |
| 1-599 | + | 585 | Intergenic | Tbl1xr1 | 1 | Untreated (DSB peaks) |
| 1-256 | + | 564 | Intergenic | C230035I16Rik | 1 | Untreated (DSB peaks) |
| 1-257 | + | 734 | Intergenic | Btn1a1 | 1 | Untreated (DSB peaks) |
| 1-436 | + | 647 | non-coding (NR_151779, exon 1 of 11) | Capn15 | 1 | Untreated (DSB peaks) |
| 1-440 | + | 583 | Intergenic | Fgd2 | 1 | Untreated (DSB peaks) |
| 1-403 | + | 575 | non-coding (NR_152226, exon 3 of 3) | Gm15638 | 1 | Untreated (DSB peaks) |
| 1-404 | + | 575 | non-coding (NR_152226, exon 3 of 3) | Gm15638 | 1 | Untreated (DSB peaks) |
| 41640 | + | 583 | Intergenic | 9330175M20Rik | 1 | Untreated (DSB peaks) |
| 1-277 | + | 809 | Intergenic | Gprin1 | 1 | Untreated (DSB peaks) |
| 1-489 | + | 615 | Intergenic | Zfp608 | 1 | Untreated (DSB peaks) |
| 1-492 | + | 596 | Intergenic | Slc27a6 | 1 | Untreated (DSB peaks) |
| 1-281 | + | 1000 | Intergenic | Dapk1 | 1 | Untreated (DSB peaks) |
| 1-1024 | + | 569 | Intergenic | Smad6 | 1 | Untreated (DSB peaks) |
| 1-161 | + | 607 | Intergenic | Borcs6 | 1 | Untreated (DSB peaks) |
| 1-819 | + | 541 | Intergenic | Capg | 1 | Untreated (DSB peaks) |

| | | | | | | |
|---|---|---|---|---|---|---|
| 1-951 | + | 542 | Intergenic | Isx | 1 | Untreated (DSB peaks) |
| 1-760 | + | 629 | Intergenic | Cdc7 | 1 | Untreated (DSB peaks) |
| 1-300 | + | 651 | Intergenic | 4921509O07Rik | 1 | Untreated (DSB peaks) |
| 1-132 | + | 573 | intron (NM_001159284, intron 3 of 21) | Smtn | 1 | Untreated (DSB peaks) |
| 1-133 | + | 573 | intron (NM_001159284, intron 3 of 21) | Smtn | 1 | Untreated (DSB peaks) |
| 1-380 | + | 556 | intron (NM_030205, intron 9 of 27) | Vasn | 1 | Untreated (DSB peaks) |
| 1-382 | + | 603 | promoter-TSS (NM_001271586) | Anks3 | 1 | Untreated (DSB peaks) |
| 1-337 | + | 663 | promoter-TSS (NM_011019) | Osmr | 1 | Untreated (DSB peaks) |
| 28856 | + | 693 | promoter-TSS (NM_001359742) | Lrp11 | 1 | Untreated (DSB peaks) |
| 1-509 | + | 660 | promoter-TSS (NR_028418) | Tmem179b | 1 | Untreated (DSB peaks) |
| 1-473 | + | 542 | exon (NM_008058, exon 1 of 1) | Fzd8 | 1 | Untreated (DSB peaks) |
| 1-474 | + | 641 | intron (NM_022021, intron 2 of 8) | Mir1901 | 1 | Untreated (DSB peaks) |
| 1-924 | + | 779 | intron (NM_001270553, intron 6 of 9) | Gm21119 | 1 | Untreated (DSB peaks) |
| 1-391 | + | 1000 | intron (NM_010143, intron 1 of 15) | Ephb3 | 1 | Untreated (DSB peaks) |
| 1-392 | + | 586 | promoter-TSS (NM_001001881) | 2510009E07Rik | 1 | Untreated (DSB peaks) |
| 1-731 | + | 653 | promoter-TSS (NM_001359963) | Napepld | 1 | Untreated (DSB peaks) |
| 18994 | + | 652 | Intergenic | Mir181b-1 | 1 | Untreated (DSB peaks) |
| 1-397 | + | 811 | intron (NM_178665, intron 5 of 12) | Morf4l1-ps1 | 1 | Untreated (DSB peaks) |
| 1-605 | + | 636 | intron (NM_001361034, intron 1 of 16) | Mecom | 1 | Untreated (DSB peaks) |
| 1-734 | + | 740 | promoter-TSS (NM_009206) | Slc4a1ap | 1 | Untreated (DSB peaks) |
| 1-139 | + | 552 | exon (NM_010822, exon 1 of 4) | Mpg | 1 | Untreated (DSB peaks) |
| 1-212 | + | 574 | promoter-TSS (NM_011658) | Twist1 | 1 | Untreated (DSB peaks) |
| 43252 | + | 660 | promoter-TSS (NM_001310513) | Kansl3 | 1 | Untreated (DSB peaks) |
| 43313 | + | 643 | promoter-TSS (NM_001252201) | Map4k4 | 1 | Untreated (DSB peaks) |
| 1-996 | + | 620 | intron (NR_131928, intron 3 of 4) | 3110039I08Rik | 1 | Untreated (DSB peaks) |
| 1-997 | + | 621 | intron (NR_131928, intron 3 of 4) | 3110039I08Rik | 1 | Untreated (DSB peaks) |
| 1-270 | + | 693 | promoter-TSS (NM_025935) | Tbc1d7 | 1 | Untreated (DSB peaks) |
| 1-543 | + | 588 | intron (NM_001301831, intron 6 of 14) | Arhgap15os | 1 | Untreated (DSB peaks) |
| 1-450 | + | 720 | intron (NM_172621, intron 4 of 5) | Clic5 | 1 | Untreated (DSB peaks) |
| 1-316 | + | 543 | intron (NM_008102, intron 1 of 5) | Gch1 | 1 | Untreated (DSB peaks) |
| 1-812 | + | 613 | 5' UTR (NM_027477, exon 1 of 6) | Zfp398 | 1 | Untreated (DSB peaks) |
| 1-151 | + | 540 | promoter-TSS (NM_011694) | Vdac1 | 1 | Untreated (DSB peaks) |
| 1-522 | + | 622 | promoter-TSS (NM_001033573) | Vti1a | 1 | Untreated (DSB peaks) |
| 1-491 | + | 554 | intron (NM_177115, intron 1 of 4) | March3 | 1 | Untreated (DSB peaks) |
| 1-155 | + | 615 | promoter-TSS (NM_022423) | Rnf187 | 1 | Untreated (DSB peaks) |
| 1-101 | + | 551 | 5' UTR (NM_001359273, exon 1 of 16) | Unc5b | 1 | Untreated (DSB peaks) |
| 1-1019 | + | 542 | promoter-TSS (NM_010193) | Fem1b | 1 | Untreated (DSB peaks) |
| 1-946 | + | 541 | intron (NM_001001491, intron 1 of 7) | Tpm4 | 1 | Untreated (DSB peaks) |
| 1-497 | + | 587 | promoter-TSS (NM_001042660) | Gm20544 | 1 | Untreated (DSB peaks) |
| 1-689 | + | 581 | intron (NM_011211, intron 27 of 39) | Ptprd | 1 | Untreated (DSB peaks) |
| 1-690 | + | 581 | intron (NM_011211, intron 27 of 39) | Ptprd | 1 | Untreated (DSB peaks) |
| 11324 | + | 669 | promoter-TSS (NM_001081210) | Mrpl44 | 1 | Untreated (DSB peaks) |
| 11689 | + | 669 | exon (NM_001081210, exon 1 of 4) | Mrpl44 | 1 | Untreated (DSB peaks) |
| 1-286 | + | 844 | promoter-TSS (NR_015587) | 9330111N05Rik | 1 | Untreated (DSB peaks) |
| 1-693 | + | 544 | intron (NM_001305286, intron 27 of 45) | Mpdz | 1 | Untreated (DSB peaks) |
| 1-1035 | + | 601 | intron (NM_199195, intron 8 of 10) | Bckdhb | 1 | Untreated (DSB peaks) |
| 1-750 | + | 546 | promoter-TSS (NM_023054) | Utp3 | 1 | Untreated (DSB peaks) |

| | | | | | | |
|---|---|---|---|---|---|---|
| 1-889 | + | 543 | intron (NM_181407, intron 3 of 13) | Ccdc81 | 1 | Untreated (DSB peaks) |
| 15342 | + | 628 | promoter-TSS (NM_001159718) | Sept2 | 1 | Untreated (DSB peaks) |
| 1-424 | + | 623 | 5' UTR (NM_001081684, exon 1 of 2) | Zbtb21 | 1 | Untreated (DSB peaks) |
| 1-376 | + | 612 | intron (NM_001310622, intron 1 of 25) | Fmnl3 | 1 | Untreated (DSB peaks) |
| 1-757 | + | 721 | promoter-TSS (NM_001081107) | Helq | 1 | Untreated (DSB peaks) |
| 1-235 | + | 643 | promoter-TSS (NM_211355) | Ppp4r3a | 1 | Untreated (DSB peaks) |
| 1-237 | + | 558 | TTS (NM_001199004) | Golga5 | 1 | Untreated (DSB peaks) |
| 1-192 | + | 758 | promoter-TSS (NM_001159492) | Gpatch8 | 1 | Untreated (DSB peaks) |
| 1-1043 | + | 755 | promoter-TSS (NM_053159) | Mrpl3 | 1 | Untreated (DSB peaks) |
| 1-703 | + | 680 | promoter-TSS (NM_023178) | Dmap1 | 1 | Untreated (DSB peaks) |
| 1-198 | + | 627 | promoter-TSS (NM_011594) | Timp2 | 1 | Untreated (DSB peaks) |
| 1-910 | + | 560 | promoter-TSS (NM_019580) | Gde1 | 1 | Untreated (DSB peaks) |
| 1-847 | + | 842 | promoter-TSS (NM_013535) | Grcc10 | 1 | Untreated (DSB peaks) |
| 1-709 | + | 554 | intron (NM_025297, intron 1 of 9) | Mecr | 1 | Untreated (DSB peaks) |
| 1-853 | + | 746 | promoter-TSS (NM_007961) | Etv6 | 1 | Untreated (DSB peaks) |
| 1-713 | + | 798 | promoter-TSS (NM_198248) | Zbtb40 | 1 | Untreated (DSB peaks) |
| 1-714 | + | 570 | intron (NM_008546, intron 1 of 8) | Mfap2 | 1 | Untreated (DSB peaks) |
| 1-648 | + | 553 | promoter-TSS (NM_009472) | Unc5c | 1 | Untreated (DSB peaks) |
| 1-651 | + | 706 | promoter-TSS (NM_001282042) | Sh3glb1 | 1 | Untreated (DSB peaks) |
| 1-724 | + | 720 | promoter-TSS (NM_007859) | Dffb | 1 | Untreated (DSB peaks) |
| 23012 | + | 680 | intron (NM_001305643, intron 19 of 19) | Scyl3 | 1 | Untreated (DSB peaks) |
| 25204 | + | 735 | intron (NM_019445, intron 6 of 17) | Fmn2 | 1 | Untreated (DSB peaks) |
| 27395 | + | 565 | Intergenic | A730004F24Rik | 1 | Untreated (DSB peaks) |
| 25934 | + | 699 | promoter-TSS (NM_145943) | Sde2 | 1 | Untreated (DSB peaks) |
| 1-1107 | + | 551 | promoter-TSS (NM_001253758) | Anp32e | 39 | tRA treated (DSB peaks) |
| 1-971 | + | 617 | promoter-TSS (NM_007615) | Ctnnd1 | 33 | tRA treated (DSB peaks) |
| 1-1189 | + | 702 | promoter-TSS (NM_001037913) | Rmrp | 30 | tRA treated (DSB peaks) |
| 1-935 | + | 560 | promoter-TSS (NM_146116) | Tubb4b | 27 | tRA treated (DSB peaks) |
| 1-826 | + | 1000 | promoter-TSS (NM_009860) | Cdc25c | 26 | tRA treated (DSB peaks) |
| 1-120 | + | 615 | Intergenic | Iyd | 2 | tRA treated (DSB peaks) |
| 1-934 | + | 560 | promoter-TSS (NM_146116) | Tubb4b | 25 | tRA treated (DSB peaks) |
| 1-1161 | + | 696 | 5' UTR (NM_026147, exon 1 of 4) | Rps20 | 4 | tRA treated (DSB peaks) |
| 1-781 | + | 670 | promoter-TSS (NM_001304785) | Ubxn6 | 23 | tRA treated (DSB peaks) |
| 1-417 | + | 632 | 5' UTR (NM_001047159, exon 1 of 10) | Net1 | 2 | tRA treated (DSB peaks) |
| 1-656 | + | 708 | promoter-TSS (NM_011653) | Tuba1a | 23 | tRA treated (DSB peaks) |
| 1-169 | + | 690 | promoter-TSS (NM_001164151) | Tcf3 | 22 | tRA treated (DSB peaks) |
| 1-297 | + | 895 | promoter-TSS (NM_026982) | Tmem256 | 21 | tRA treated (DSB peaks) |
| 1-665 | + | 607 | Intergenic | Slx4 | 5 | tRA treated (DSB peaks) |
| 1-666 | + | 607 | Intergenic | Slx4 | 4 | tRA treated (DSB peaks) |
| 1-644 | + | 642 | promoter-TSS (NM_008495) | Lgals1 | 21 | tRA treated (DSB peaks) |
| 1-861 | + | 776 | exon (NM_134148, exon 9 of 10) | Carns1 | 3 | tRA treated (DSB peaks) |
| 1-733 | + | 830 | Intergenic | 4930548J01Rik | 2 | tRA treated (DSB peaks) |
| 1-668 | + | 588 | Intergenic | Gm5766 | 7 | tRA treated (DSB peaks) |
| 1-862 | + | 690 | intron (NM_001001984, intron 12 of 20) | Grk2 | 10 | tRA treated (DSB peaks) |
| 1-916 | + | 561 | intron (NM_001177843, intron 2 of 23) | Frmd4a | 4 | tRA treated (DSB peaks) |
| 1-917 | + | 562 | intron (NM_001177843, intron 2 of 23) | Frmd4a | 9 | tRA treated (DSB peaks) |
| 1-370 | + | 633 | Intergenic | Itsn2 | 9 | tRA treated (DSB peaks) |

| | | | | | | |
|---|---|---|---|---|---|---|
| 1-121 | + | 660 | intron (NM_001302531, intron 3 of 8) | Esr1 | 3 | tRA treated (DSB peaks) |
| 1-758 | + | 593 | promoter-TSS (NM_008997) | Rab11b | 20 | tRA treated (DSB peaks) |
| 1-1212 | + | 669 | promoter-TSS (NR_151607) | Gm12602 | 20 | tRA treated (DSB peaks) |
| 1-655 | + | 708 | promoter-TSS (NM_011653) | Tuba1a | 20 | tRA treated (DSB peaks) |
| 1-864 | + | 546 | intron (NM_153553, intron 1 of 7) | Npas4 | 3 | tRA treated (DSB peaks) |
| 1-361 | + | 932 | promoter-TSS (NM_007997) | Fdxr | 20 | tRA treated (DSB peaks) |
| 1-814 | + | 574 | non-coding (NR_153824, exon 3 of 3) | Gm26682 | 1 | tRA treated (DSB peaks) |
| 1-876 | + | 666 | promoter-TSS (NM_001167799) | Stx5a | 19 | tRA treated (DSB peaks) |
| 1-418 | + | 578 | Intergenic | 1700016G22Rik | 4 | tRA treated (DSB peaks) |
| 1-395 | + | 641 | promoter-TSS (NR_073563) | Numb | 19 | tRA treated (DSB peaks) |
| 1-767 | + | 584 | promoter-TSS (NM_001010833) | Mdc1 | 18 | tRA treated (DSB peaks) |
| 1-1661 | + | 567 | promoter-TSS (NM_173015) | Mon1b | 18 | tRA treated (DSB peaks) |
| 1-1519 | + | 590 | promoter-TSS (NM_011304) | Gys1 | 17 | tRA treated (DSB peaks) |
| 1-567 | + | 568 | promoter-TSS (NM_001110162) | Kctd9 | 17 | tRA treated (DSB peaks) |
| 1-220 | + | 550 | exon (NM_001146308, exon 1 of 13) | Dbnl | 2 | tRA treated (DSB peaks) |
| 1-221 | + | 550 | intron (NM_001146309, intron 1 of 12) | Dbnl | 3 | tRA treated (DSB peaks) |
| 1-866 | + | 692 | Intergenic | Malat1 | 9 | tRA treated (DSB peaks) |
| 1-867 | + | 583 | Intergenic | Malat1 | 7 | tRA treated (DSB peaks) |
| 1-337 | + | 579 | promoter-TSS (NM_001080964) | Sp2 | 17 | tRA treated (DSB peaks) |
| 1-863 | + | 604 | promoter-TSS (NM_133803) | Dpp3 | 16 | tRA treated (DSB peaks) |
| 1-868 | + | 570 | non-coding (NR_003513, exon 1 of 1) | Neat1 | 36 | tRA treated (DSB peaks) |
| 1-869 | + | 740 | TTS (NM_026169) | Neat1 | 2 | tRA treated (DSB peaks) |
| 1-870 | + | 740 | TTS (NM_026169) | Neat1 | 4 | tRA treated (DSB peaks) |
| 1-1190 | + | 702 | promoter-TSS (NR_001460) | Rmrp | 16 | tRA treated (DSB peaks) |
| 1-904 | + | 648 | promoter-TSS (NM_001177812) | Wbp1l | 16 | tRA treated (DSB peaks) |
| 1-268 | + | 605 | promoter-TSS (NM_013716) | G3bp1 | 16 | tRA treated (DSB peaks) |
| 1-648 | + | 784 | promoter-TSS (NM_001199024) | Pla2g6 | 16 | tRA treated (DSB peaks) |
| 1-1164 | + | 732 | intron (NM_026534, intron 1 of 7) | Ubxn2b | 3 | tRA treated (DSB peaks) |
| 1-223 | + | 605 | 5' UTR (NM_173748, exon 1 of 6) | Nudcd3 | 3 | tRA treated (DSB peaks) |
| 1-1785 | + | 603 | promoter-TSS (NM_178660) | Rbms3 | 16 | tRA treated (DSB peaks) |
| 1-873 | + | 607 | intron (NM_010119, intron 1 of 4) | Ehd1 | 6 | tRA treated (DSB peaks) |
| 1-1473 | + | 580 | promoter-TSS (NM_010749) | M6pr | 16 | tRA treated (DSB peaks) |
| 1-1352 | + | 547 | promoter-TSS (NM_001042421) | Rsrc2 | 16 | tRA treated (DSB peaks) |
| 1-1251 | + | 679 | promoter-TSS (NR_002865) | Rnu11 | 16 | tRA treated (DSB peaks) |
| 1-1494 | + | 878 | Intergenic | Smim17 | 1 | tRA treated (DSB peaks) |
| 34700 | + | 875 | promoter-TSS (NM_023284) | Nuf2 | 16 | tRA treated (DSB peaks) |
| 1-267 | + | 605 | promoter-TSS (NM_013716) | G3bp1 | 15 | tRA treated (DSB peaks) |
| 1-188 | + | 565 | promoter-TSS (NM_011631) | Hsp90b1 | 15 | tRA treated (DSB peaks) |
| 1-357 | + | 667 | promoter-TSS (NR_030703) | Snord104 | 15 | tRA treated (DSB peaks) |
| 1-919 | + | 566 | intron (NM_001110232, intron 2 of 15) | Celf2 | 1 | tRA treated (DSB peaks) |
| 1-920 | + | 620 | intron (NM_001110232, intron 2 of 15) | Celf2 | 8 | tRA treated (DSB peaks) |
| 1-875 | + | 1000 | TTS (NR_105957) | Mir6991 | 1 | tRA treated (DSB peaks) |
| 1-1580 | + | 586 | promoter-TSS (NM_007672) | Mfsd13b | 15 | tRA treated (DSB peaks) |
| 1-524 | + | 565 | Intergenic | Flnb | 3 | tRA treated (DSB peaks) |
| 1-525 | + | 592 | intron (NM_001081427, intron 1 of 45) | Flnb | 11 | tRA treated (DSB peaks) |
| 1-1679 | + | 570 | intron (NM_009534, intron 4 of 7) | Yap1 | 4 | tRA treated (DSB peaks) |
| 1-1680 | + | 570 | intron (NM_009534, intron 4 of 7) | Yap1 | 5 | tRA treated (DSB peaks) |

| | | | | | | |
|---|---|---|---|---|---|---|
| 1-1147 | + | 1000 | promoter-TSS (NM_010516) | Cyr61 | 15 | tRA treated (DSB peaks) |
| 1-1396 | + | 551 | intron (NM_001037865, intron 3 of 34) | Col28a1 | 10 | tRA treated (DSB peaks) |
| 1-1397 | + | 551 | intron (NM_001037865, intron 3 of 34) | Col28a1 | 10 | tRA treated (DSB peaks) |
| 1-830 | + | 567 | promoter-TSS (NR_027885) | Vaultrc5 | 14 | tRA treated (DSB peaks) |
| 1-373 | + | 594 | intron (NM_021429, intron 1 of 6) | Hs1bp3 | 1 | tRA treated (DSB peaks) |
| 1-670 | + | 592 | intron (NM_025821, intron 1 of 3) | Carhsp1 | 1 | tRA treated (DSB peaks) |
| 1-597 | + | 718 | intron (NM_148938, intron 3 of 9) | Slc1a3 | 2 | tRA treated (DSB peaks) |
| 1-952 | + | 561 | promoter-TSS (NR_040356) | 4930555B11Rik | 14 | tRA treated (DSB peaks) |
| 1-921 | + | 671 | Intergenic | 1700061F12Rik | 10 | tRA treated (DSB peaks) |
| 1-647 | + | 785 | promoter-TSS (NM_001199024) | Pla2g6 | 14 | tRA treated (DSB peaks) |
| 1-1060 | + | 560 | exon (NM_025434, exon 1 of 3) | Mrps28 | 2 | tRA treated (DSB peaks) |
| 1-343 | + | 868 | promoter-TSS (NM_011486) | Stat3 | 14 | tRA treated (DSB peaks) |
| 1-986 | + | 604 | promoter-TSS (NM_026271) | Fibin | 14 | tRA treated (DSB peaks) |
| 1-374 | + | 549 | Intergenic | 1700022H16Rik | 8 | tRA treated (DSB peaks) |
| 1-1501 | + | 811 | promoter-TSS (NM_007948) | Ercc1 | 13 | tRA treated (DSB peaks) |
| 1-766 | + | 611 | promoter-TSS (NM_133662) | Ier3 | 13 | tRA treated (DSB peaks) |
| 1-1062 | + | 541 | intron (NM_053182, intron 2 of 7) | Zfp704 | 1 | tRA treated (DSB peaks) |
| 1-880 | + | 629 | intron (NM_019699, intron 1 of 11) | Fads2 | 2 | tRA treated (DSB peaks) |
| 1-881 | + | 629 | intron (NM_019699, intron 1 of 11) | Fads2 | 2 | tRA treated (DSB peaks) |
| 1-953 | + | 545 | promoter-TSS (NR_040356) | 4930555B11Rik | 13 | tRA treated (DSB peaks) |
| 1-651 | + | 627 | promoter-TSS (NM_001168672) | Gtse1 | 13 | tRA treated (DSB peaks) |
| 1-976 | + | 822 | promoter-TSS (NM_007387) | Acp2 | 13 | tRA treated (DSB peaks) |
| 1-1666 | + | 1000 | promoter-TSS (NM_178856) | Gins2 | 13 | tRA treated (DSB peaks) |
| 1-368 | + | 550 | promoter-TSS (NM_178664) | B3gntl1 | 13 | tRA treated (DSB peaks) |
| 1-598 | + | 652 | intron (NR_045063, intron 2 of 6) | 4930556M19Rik | 5 | tRA treated (DSB peaks) |
| 1-599 | + | 652 | intron (NR_045063, intron 2 of 6) | 4930556M19Rik | 4 | tRA treated (DSB peaks) |
| 1-600 | + | 641 | Intergenic | Gm19276 | 4 | tRA treated (DSB peaks) |
| 1-1167 | + | 557 | Intergenic | Ccne2 | 1 | tRA treated (DSB peaks) |
| 1-1379 | + | 783 | promoter-TSS (NM_021392) | Mcm7 | 13 | tRA treated (DSB peaks) |
| 1-769 | + | 812 | promoter-TSS (NM_029632) | Ppp1r11 | 12 | tRA treated (DSB peaks) |
| 1-832 | + | 664 | promoter-TSS (NR_110344) | Tcerg1 | 12 | tRA treated (DSB peaks) |
| 1-601 | + | 578 | Intergenic | 6030458C11Rik | 3 | tRA treated (DSB peaks) |
| 1-734 | + | 597 | intron (NR_027784, intron 2 of 2) | Igf2r | 5 | tRA treated (DSB peaks) |
| 1-1169 | + | 581 | intron (NM_001171801, intron 1 of 4) | Triqk | 4 | tRA treated (DSB peaks) |
| 1-1290 | + | 690 | Intergenic | Sema3a | 5 | tRA treated (DSB peaks) |
| 1-124 | + | 729 | non-coding (NR_038027, exon 1 of 4) | B230208H11Rik | 3 | tRA treated (DSB peaks) |
| 1-1495 | + | 740 | exon (NM_175558, exon 1 of 6) | Zfp446 | 6 | tRA treated (DSB peaks) |
| 1-1535 | + | 552 | promoter-TSS (NM_173445) | Rccd1 | 12 | tRA treated (DSB peaks) |
| 1-175 | + | 674 | promoter-TSS (NM_001358898) | Slc39a3 | 12 | tRA treated (DSB peaks) |
| 1-1681 | + | 763 | Intergenic | Ccdc82 | 7 | tRA treated (DSB peaks) |
| 1-1536 | + | 558 | promoter-TSS (NM_007755) | Cpeb1 | 12 | tRA treated (DSB peaks) |
| 1-1606 | + | 578 | intron (NM_009025, intron 1 of 23) | Rasa3 | 1 | tRA treated (DSB peaks) |
| 1-736 | + | 620 | Intergenic | 4930474M22Rik | 9 | tRA treated (DSB peaks) |
| 1-1682 | + | 720 | intron (NM_001081395, intron 3 of 12) | Amotl1 | 3 | tRA treated (DSB peaks) |
| 1-737 | + | 630 | 5' UTR (NM_011581, exon 1 of 22) | Thbs2 | 8 | tRA treated (DSB peaks) |
| 1-1034 | + | 787 | promoter-TSS (NM_001291114) | Rbm39 | 12 | tRA treated (DSB peaks) |
| 1-1170 | + | 560 | 5' UTR (NM_152812, exon 1 of 7) | Otud6b | 4 | tRA treated (DSB peaks) |

| | | | | | | |
|---|---|---|---|---|---|---|
| 1-527 | + | 665 | intron (NM_001195229, intron 4 of 34) | Nek10 | 6 | tRA treated (DSB peaks) |
| 1-672 | + | 701 | Intergenic | Efcab1 | 4 | tRA treated (DSB peaks) |
| 1-915 | + | 707 | promoter-TSS (NM_025626) | Fam107b | 11 | tRA treated (DSB peaks) |
| 1-123 | + | 638 | promoter-TSS (NM_175374) | Mtrf1l | 11 | tRA treated (DSB peaks) |
| 1-1690 | + | 654 | promoter-TSS (NR_033729) | Fbxl12os | 11 | tRA treated (DSB peaks) |
| 1-768 | + | 883 | promoter-TSS (NM_026987) | Dhx16 | 11 | tRA treated (DSB peaks) |
| 1-248 | + | 624 | promoter-TSS (NM_007897) | Ebf1 | 11 | tRA treated (DSB peaks) |
| 1-740 | + | 654 | Intergenic | Prdm9 | 12 | tRA treated (DSB peaks) |
| 1-419 | + | 815 | Intergenic | 4933412O06Rik | 4 | tRA treated (DSB peaks) |
| 1-1171 | + | 619 | intron (NM_013752, intron 1 of 15) | Nbn | 1 | tRA treated (DSB peaks) |
| 1-1497 | + | 541 | Intergenic | Ccdc9 | 7 | tRA treated (DSB peaks) |
| 1-1619 | + | 619 | promoter-TSS (NM_027973) | Cenpu | 11 | tRA treated (DSB peaks) |
| 1-225 | + | 582 | Intergenic | Sec61g | 2 | tRA treated (DSB peaks) |
| 1-226 | + | 583 | Intergenic | Sec61g | 6 | tRA treated (DSB peaks) |
| 1-528 | + | 558 | intron (NM_001289761, intron 2 of 8) | Rarb | 4 | tRA treated (DSB peaks) |
| 1-227 | + | 702 | Intergenic | Sec61g | 5 | tRA treated (DSB peaks) |
| 1-819 | + | 547 | intron (NM_007664, intron 2 of 15) | Cdh2 | 6 | tRA treated (DSB peaks) |
| 1-1421 | + | 667 | promoter-TSS (NM_172728) | Creb5 | 11 | tRA treated (DSB peaks) |
| 1-1628 | + | 607 | promoter-TSS (NM_001039965) | Zfp869 | 11 | tRA treated (DSB peaks) |
| 1-126 | + | 608 | intron (NR_038020, intron 1 of 8) | Gm20125 | 5 | tRA treated (DSB peaks) |
| 1-127 | + | 819 | intron (NR_038020, intron 1 of 8) | Gm20125 | 10 | tRA treated (DSB peaks) |
| 1-228 | + | 580 | Intergenic | Cnrip1 | 2 | tRA treated (DSB peaks) |
| 1-229 | + | 635 | Intergenic | C1d | 7 | tRA treated (DSB peaks) |
| 1-1291 | + | 756 | Intergenic | Sema3c | 4 | tRA treated (DSB peaks) |
| 1-128 | + | 606 | exon (NM_010828, exon 2 of 2) | Cited2 | 1 | tRA treated (DSB peaks) |
| 1-1531 | + | 967 | promoter-TSS (NM_017462) | Polg | 11 | tRA treated (DSB peaks) |
| 1-1398 | + | 598 | intron (NM_022332, intron 1 of 15) | St7 | 16 | tRA treated (DSB peaks) |
| 1-674 | + | 807 | exon (NM_153790, exon 11 of 11) | Scarf2 | 2 | tRA treated (DSB peaks) |
| 1-1532 | + | 621 | promoter-TSS (NM_017462) | Polg | 11 | tRA treated (DSB peaks) |
| 1-741 | + | 727 | intron (NM_001162909, intron 6 of 8) | Spaca6 | 11 | tRA treated (DSB peaks) |
| 1-922 | + | 570 | non-coding (NR_028376, exon 2 of 2) | Gm17762 | 1 | tRA treated (DSB peaks) |
| 1-1537 | + | 558 | promoter-TSS (NM_007755) | Cpeb1 | 11 | tRA treated (DSB peaks) |
| 1-1326 | + | 785 | promoter-TSS (NM_017367) | 2010109A12Rik | 11 | tRA treated (DSB peaks) |
| 1-1063 | + | 860 | Intergenic | 4930433B08Rik | 7 | tRA treated (DSB peaks) |
| 1-1064 | + | 638 | Intergenic | 4930433B08Rik | 6 | tRA treated (DSB peaks) |
| 1-923 | + | 646 | Intergenic | Commd3 | 2 | tRA treated (DSB peaks) |
| 1-924 | + | 646 | Intergenic | Commd3 | 3 | tRA treated (DSB peaks) |
| 1-1607 | + | 601 | intron (NM_173189, intron 3 of 13) | Mcph1 | 3 | tRA treated (DSB peaks) |
| 1-925 | + | 824 | Intergenic | Gm3363 | 2 | tRA treated (DSB peaks) |
| 1-405 | + | 708 | promoter-TSS (NM_001330702) | Rps6ka5 | 11 | tRA treated (DSB peaks) |
| 1-202 | + | 877 | promoter-TSS (NM_178610) | Krr1 | 11 | tRA treated (DSB peaks) |
| 1-1065 | + | 831 | Intergenic | Trim55 | 10 | tRA treated (DSB peaks) |
| 1-1346 | + | 727 | promoter-TSS (NM_007475) | Rplp0 | 11 | tRA treated (DSB peaks) |
| 1-1608 | + | 1000 | intron (NM_001270553, intron 6 of 9) | 4930467E23Rik | 1 | tRA treated (DSB peaks) |
| 1-230 | + | 597 | Intergenic | Spred2 | 4 | tRA treated (DSB peaks) |
| 1-211 | + | 557 | promoter-TSS (NM_007837) | Ddit3 | 11 | tRA treated (DSB peaks) |
| 1-1609 | + | 1000 | intron (NM_001270553, intron 6 of 9) | 4930467E23Rik | 1 | tRA treated (DSB peaks) |

| | | | | | | |
|---|---|---|---|---|---|---|
| 1-529 | + | 565 | exon (NM_026209, exon 1 of 2) | Saysd1 | 5 | tRA treated (DSB peaks) |
| 1-1008 | + | 742 | promoter-TSS (NM_178404) | Zc3h6 | 11 | tRA treated (DSB peaks) |
| 1-1033 | + | 787 | promoter-TSS (NM_001291114) | Rbm39 | 11 | tRA treated (DSB peaks) |
| 1-886 | + | 955 | Intergenic | Anxa1 | 3 | tRA treated (DSB peaks) |
| 1-592 | + | 813 | promoter-TSS (NM_145930) | Gm2093 | 10 | tRA treated (DSB peaks) |
| 1-593 | + | 812 | promoter-TSS (NM_145930) | Gm2093 | 10 | tRA treated (DSB peaks) |
| 1-878 | + | 679 | promoter-TSS (NM_026007) | Eef1g | 10 | tRA treated (DSB peaks) |
| 1-759 | + | 813 | promoter-TSS (NM_001195298) | Kifc1 | 10 | tRA treated (DSB peaks) |
| 1-765 | + | 612 | promoter-TSS (NM_133662) | Ier3 | 10 | tRA treated (DSB peaks) |
| 1-1691 | + | 654 | non-coding (NR_033729, exon 1 of 3) | Fbxl12os | 8 | tRA treated (DSB peaks) |
| 1-677 | + | 542 | 5' UTR (NM_009900, exon 1 of 24) | Clcn2 | 1 | tRA treated (DSB peaks) |
| 1-555 | + | 638 | promoter-TSS (NM_013768) | Prmt5 | 10 | tRA treated (DSB peaks) |
| 1-1722 | + | 629 | promoter-TSS (NM_025812) | Peak1 | 10 | tRA treated (DSB peaks) |
| 1-233 | + | 579 | Intergenic | Lgalsl | 1 | tRA treated (DSB peaks) |
| 1-1692 | + | 617 | exon (NM_145154, exon 2 of 6) | Angptl6 | 1 | tRA treated (DSB peaks) |
| 1-381 | + | 586 | promoter-TSS (NM_001037746) | Mipol1 | 10 | tRA treated (DSB peaks) |
| 1-498 | + | 639 | promoter-TSS (NM_146231) | Zfp825 | 10 | tRA treated (DSB peaks) |
| 1-420 | + | 631 | Intergenic | Olfr1368 | 4 | tRA treated (DSB peaks) |
| 1-1693 | + | 600 | intron (NM_183408, intron 1 of 14) | Pde4a | 1 | tRA treated (DSB peaks) |
| 1-1292 | + | 655 | Intergenic | Gsap | 8 | tRA treated (DSB peaks) |
| 1-1694 | + | 600 | intron (NM_183408, intron 1 of 14) | Pde4a | 2 | tRA treated (DSB peaks) |
| 1-421 | + | 609 | Intergenic | Trim27 | 8 | tRA treated (DSB peaks) |
| 1-422 | + | 642 | Intergenic | Trim27 | 6 | tRA treated (DSB peaks) |
| 1-423 | + | 557 | Intergenic | Gpx5 | 4 | tRA treated (DSB peaks) |
| 1-424 | + | 850 | Intergenic | Gpx5 | 9 | tRA treated (DSB peaks) |
| 1-165 | + | 836 | promoter-TSS (NM_001347201) | Gucd1 | 10 | tRA treated (DSB peaks) |
| 1-678 | + | 555 | Intergenic | Magef1 | 4 | tRA treated (DSB peaks) |
| 1-679 | + | 570 | Intergenic | Magef1 | 1 | tRA treated (DSB peaks) |
| 1-680 | + | 606 | Intergenic | Magef1 | 3 | tRA treated (DSB peaks) |
| 1-637 | + | 572 | promoter-TSS (NM_001110830) | Rbfox2 | 10 | tRA treated (DSB peaks) |
| 1-354 | + | 730 | promoter-TSS (NM_011947) | Map3k3 | 10 | tRA treated (DSB peaks) |
| 1-355 | + | 729 | promoter-TSS (NM_011947) | Map3k3 | 10 | tRA treated (DSB peaks) |
| 1-210 | + | 558 | promoter-TSS (NM_007837) | Ddit3 | 10 | tRA treated (DSB peaks) |
| 1-1294 | + | 704 | 5' UTR (NM_011188, exon 2 of 13) | Psmc2 | 9 | tRA treated (DSB peaks) |
| 1-213 | + | 581 | promoter-TSS (NR_104288) | Rnf41 | 10 | tRA treated (DSB peaks) |
| 1-426 | + | 588 | Intergenic | Mir1983 | 9 | tRA treated (DSB peaks) |
| 1-427 | + | 656 | Intergenic | Mir1983 | 4 | tRA treated (DSB peaks) |
| 1-428 | + | 656 | Intergenic | Mir1983 | 6 | tRA treated (DSB peaks) |
| 1-1066 | + | 1000 | Intergenic | Tbl1xr1 | 3 | tRA treated (DSB peaks) |
| 1-1067 | + | 562 | Intergenic | Tbl1xr1 | 1 | tRA treated (DSB peaks) |
| 1-429 | + | 828 | Intergenic | Zfp184 | 10 | tRA treated (DSB peaks) |
| 1-430 | + | 603 | Intergenic | Zfp184 | 2 | tRA treated (DSB peaks) |
| 1-1030 | + | 601 | promoter-TSS (NM_001141976) | Tpx2 | 10 | tRA treated (DSB peaks) |
| 1-432 | + | 890 | Intergenic | Pom121l2 | 14 | tRA treated (DSB peaks) |
| 1-534 | + | 574 | exon (NM_145459, exon 3 of 3) | Zfp503 | 6 | tRA treated (DSB peaks) |
| 1-433 | + | 606 | Intergenic | Prss16 | 1 | tRA treated (DSB peaks) |
| 1-434 | + | 616 | Intergenic | Prss16 | 10 | tRA treated (DSB peaks) |

| | | | | | | |
|---|---|---|---|---|---|---|
| 1-435 | + | 610 | Intergenic | Hist1h2ah | 1 | tRA treated (DSB peaks) |
| 1-530 | + | 1000 | promoter-TSS (NM_001360365) | Fam149b | 9 | tRA treated (DSB peaks) |
| 1-744 | + | 874 | promoter-TSS (NM_027008) | Kctd5 | 9 | tRA treated (DSB peaks) |
| 1-927 | + | 714 | intron (NM_019456, intron 1 of 13) | Apbb1ip | 8 | tRA treated (DSB peaks) |
| 1-928 | + | 714 | intron (NM_019456, intron 1 of 13) | Apbb1ip | 10 | tRA treated (DSB peaks) |
| 1-748 | + | 615 | promoter-TSS (NR_027452) | Ube2i | 9 | tRA treated (DSB peaks) |
| 1-235 | + | 637 | exon (NM_009837, exon 1 of 14) | Cct4 | 5 | tRA treated (DSB peaks) |
| 1-236 | + | 637 | intron (NM_009837, intron 1 of 13) | Cct4 | 2 | tRA treated (DSB peaks) |
| 1-1504 | + | 627 | promoter-TSS (NM_001360789) | Hnrnpul1 | 9 | tRA treated (DSB peaks) |
| 1-237 | + | 555 | intron (NM_001035226, intron 2 of 25) | Xpo1 | 10 | tRA treated (DSB peaks) |
| 1-1507 | + | 889 | promoter-TSS (NM_001083912) | Plekhg2 | 9 | tRA treated (DSB peaks) |
| 1-436 | + | 663 | intron (NR_126444, intron 1 of 1) | 4930557F10Rik | 4 | tRA treated (DSB peaks) |
| 1-437 | + | 843 | intron (NR_131052, intron 2 of 3) | 4930586N03Rik | 6 | tRA treated (DSB peaks) |
| 1-438 | + | 549 | Intergenic | Abt1 | 3 | tRA treated (DSB peaks) |
| 1-439 | + | 983 | Intergenic | Abt1 | 6 | tRA treated (DSB peaks) |
| 1-440 | + | 561 | Intergenic | Abt1 | 4 | tRA treated (DSB peaks) |
| 1-441 | + | 581 | Intergenic | C230035I16Rik | 8 | tRA treated (DSB peaks) |
| 1-1295 | + | 676 | non-coding (NR_040469, exon 1 of 1) | 5031425E22Rik | 4 | tRA treated (DSB peaks) |
| 1-442 | + | 651 | Intergenic | C230035I16Rik | 5 | tRA treated (DSB peaks) |
| 1-443 | + | 651 | Intergenic | C230035I16Rik | 7 | tRA treated (DSB peaks) |
| 1-444 | + | 552 | Intergenic | C230035I16Rik | 3 | tRA treated (DSB peaks) |
| 1-445 | + | 575 | Intergenic | C230035I16Rik | 9 | tRA treated (DSB peaks) |
| 1-446 | + | 547 | Intergenic | Btn1a1 | 4 | tRA treated (DSB peaks) |
| 1-447 | + | 596 | Intergenic | Btn1a1 | 5 | tRA treated (DSB peaks) |
| 1-448 | + | 994 | Intergenic | Btn1a1 | 5 | tRA treated (DSB peaks) |
| 1-449 | + | 769 | Intergenic | Btn2a2 | 1 | tRA treated (DSB peaks) |
| 1-743 | + | 818 | Intergenic | 6330415G19Rik | 6 | tRA treated (DSB peaks) |
| 1-930 | + | 636 | intron (NM_029773, intron 1 of 10) | Spopl | 1 | tRA treated (DSB peaks) |
| 1-702 | + | 626 | promoter-TSS (NM_027342) | Ccdc58 | 9 | tRA treated (DSB peaks) |
| 1-1521 | + | 963 | promoter-TSS (NM_175318) | Spty2d1 | 9 | tRA treated (DSB peaks) |
| 1-1697 | + | 625 | Intergenic | Npsr1 | 7 | tRA treated (DSB peaks) |
| 1-1426 | + | 569 | promoter-TSS (NM_019499) | Mad2l1 | 9 | tRA treated (DSB peaks) |
| 1-958 | + | 656 | promoter-TSS (NM_029280) | Mettl5 | 9 | tRA treated (DSB peaks) |
| 1-1429 | + | 687 | promoter-TSS (NM_173001) | Kdm3a | 9 | tRA treated (DSB peaks) |
| 1-931 | + | 856 | intron (NM_011040, intron 2 of 11) | Pax8 | 11 | tRA treated (DSB peaks) |
| 1-932 | + | 600 | intron (NM_011040, intron 2 of 11) | Pax8 | 6 | tRA treated (DSB peaks) |
| 1-859 | + | 644 | promoter-TSS (NM_025797) | Cyb5a | 9 | tRA treated (DSB peaks) |
| 1-746 | + | 564 | intron (NM_001252463, intron 1 of 8) | Mlst8 | 5 | tRA treated (DSB peaks) |
| 1-1444 | + | 806 | promoter-TSS (NM_133934) | Isy1 | 9 | tRA treated (DSB peaks) |
| 1-683 | + | 549 | intron (NM_178665, intron 3 of 12) | Morf4l1-ps1 | 16 | tRA treated (DSB peaks) |
| 1-1447 | + | 591 | promoter-TSS (NM_030251) | Abtb1 | 9 | tRA treated (DSB peaks) |
| 43160 | + | 550 | intron (NM_007733, intron 4 of 50) | Col19a1 | 6 | tRA treated (DSB peaks) |
| 1-1296 | + | 688 | Intergenic | Smarcd3 | 2 | tRA treated (DSB peaks) |
| 1-1217 | + | 705 | promoter-TSS (NM_145549) | Itgb3bp | 9 | tRA treated (DSB peaks) |
| 1-352 | + | 824 | promoter-TSS (NM_027346) | Taco1 | 9 | tRA treated (DSB peaks) |
| 1-129 | + | 665 | Intergenic | Enpp1 | 4 | tRA treated (DSB peaks) |
| 1-984 | + | 599 | promoter-TSS (NM_172671) | Lgr4 | 9 | tRA treated (DSB peaks) |

| | | | | | | |
|---|---|---|---|---|---|---|
| 1-747 | + | 671 | intron (NM_134126, intron 18 of 29) | Tmem204 | 5 | tRA treated (DSB peaks) |
| 1-1776 | + | 541 | promoter-TSS (NM_008633) | Map4 | 9 | tRA treated (DSB peaks) |
| 1-1463 | + | 580 | promoter-TSS (NM_001282126) | Brpf1 | 9 | tRA treated (DSB peaks) |
| 1-1783 | + | 553 | promoter-TSS (NM_153585) | Cnot10 | 9 | tRA treated (DSB peaks) |
| 1-989 | + | 730 | promoter-TSS (NM_026620) | Fam98b | 9 | tRA treated (DSB peaks) |
| 1-131 | + | 549 | intron (NM_018747, intron 3 of 7) | Akap7 | 9 | tRA treated (DSB peaks) |
| 1-132 | + | 549 | intron (NM_018747, intron 3 of 7) | Akap7 | 9 | tRA treated (DSB peaks) |
| 1-1664 | + | 625 | promoter-TSS (NM_028883) | Tldc1 | 9 | tRA treated (DSB peaks) |
| 1-1351 | + | 1000 | promoter-TSS (NM_021505) | Anapc5 | 9 | tRA treated (DSB peaks) |
| 1-133 | + | 548 | intron (NM_018747, intron 1 of 7) | Akap7 | 2 | tRA treated (DSB peaks) |
| 1-450 | + | 1000 | Intergenic | Nrsn1 | 2 | tRA treated (DSB peaks) |
| 1-451 | + | 1000 | Intergenic | Nrsn1 | 1 | tRA treated (DSB peaks) |
| 1-1611 | + | 592 | Intergenic | Fgfr1 | 13 | tRA treated (DSB peaks) |
| 1-134 | + | 796 | intron (NM_001358754, intron 16 of 16) | Smlr1 | 8 | tRA treated (DSB peaks) |
| 1-684 | + | 543 | Intergenic | Trp63 | 5 | tRA treated (DSB peaks) |
| 1-1502 | + | 673 | TTS (NM_001362154) | Tgfb1 | 1 | tRA treated (DSB peaks) |
| 1-1503 | + | 554 | 3' UTR (NM_028771, exon 5 of 5) | Ccdc97 | 3 | tRA treated (DSB peaks) |
| 1-1792 | + | 701 | promoter-TSS (NM_030692) | Sacm1l | 9 | tRA treated (DSB peaks) |
| 1-602 | + | 542 | intron (NM_001353142, intron 1 of 20) | Myo10 | 5 | tRA treated (DSB peaks) |
| 1-1481 | + | 779 | promoter-TSS (NM_009496) | Vamp1 | 9 | tRA treated (DSB peaks) |
| 1-1378 | + | 743 | promoter-TSS (NM_011757) | Zscan21 | 9 | tRA treated (DSB peaks) |
| 1-1175 | + | 579 | Intergenic | Manea | 6 | tRA treated (DSB peaks) |
| 1-135 | + | 668 | intron (NM_176837, intron 12 of 14) | Arhgap18 | 7 | tRA treated (DSB peaks) |
| 1-136 | + | 668 | intron (NM_176837, intron 12 of 14) | Arhgap18 | 8 | tRA treated (DSB peaks) |
| 1-1394 | + | 605 | promoter-TSS (NM_201369) | N4bp2l2 | 9 | tRA treated (DSB peaks) |
| 1-1612 | + | 978 | exon (NM_001101502, exon 2 of 2) | Zfp703 | 1 | tRA treated (DSB peaks) |
| 1-112 | + | 555 | promoter-TSS (NM_026041) | Rrp15 | 9 | tRA treated (DSB peaks) |
| 1-115 | + | 642 | promoter-TSS (NM_198025) | Flvcr1 | 9 | tRA treated (DSB peaks) |
| 1-938 | + | 829 | 5' UTR (NM_001360790, exon 1 of 1) | Brd3os | 2 | tRA treated (DSB peaks) |
| 1-1505 | + | 607 | intron (NM_133210, intron 1 of 1) | Sertad3 | 18 | tRA treated (DSB peaks) |
| 1-939 | + | 619 | Intergenic | Wdr5 | 10 | tRA treated (DSB peaks) |
| 1-603 | + | 869 | intron (NM_001081302, intron 1 of 56) | Trio | 4 | tRA treated (DSB peaks) |
| 1-1299 | + | 647 | intron (NM_153526, intron 1 of 4) | Insig1 | 9 | tRA treated (DSB peaks) |
| 1-751 | + | 588 | Intergenic | Tcp11 | 3 | tRA treated (DSB peaks) |
| 1-752 | + | 588 | Intergenic | Tcp11 | 4 | tRA treated (DSB peaks) |
| 1-219 | + | 646 | promoter-TSS (NM_022889) | Pes1 | 8 | tRA treated (DSB peaks) |
| 1-1165 | + | 776 | promoter-TSS (NR_040645) | 8430436N08Rik | 8 | tRA treated (DSB peaks) |
| 1-879 | + | 578 | promoter-TSS (NM_001286518) | Ahnak | 8 | tRA treated (DSB peaks) |
| 1-671 | + | 590 | promoter-TSS (NM_021428) | Dexi | 8 | tRA treated (DSB peaks) |
| 1-1508 | + | 709 | Intergenic | Zfp36 | 11 | tRA treated (DSB peaks) |
| 1-452 | + | 919 | Intergenic | Prl5a1 | 10 | tRA treated (DSB peaks) |
| 1-453 | + | 917 | Intergenic | Prl5a1 | 11 | tRA treated (DSB peaks) |
| 43132 | + | 715 | promoter-TSS (NM_172294) | Sulf1 | 8 | tRA treated (DSB peaks) |
| 1-1401 | + | 1000 | Intergenic | Ccdc136 | 4 | tRA treated (DSB peaks) |
| 1-755 | + | 864 | Intergenic | Pim1 | 6 | tRA treated (DSB peaks) |
| 1-1687 | + | 760 | promoter-TSS (NM_025844) | Chordc1 | 8 | tRA treated (DSB peaks) |
| 1-1610 | + | 724 | promoter-TSS (NM_020560) | Mrps31 | 8 | tRA treated (DSB peaks) |

| | | | | | | |
|---|---|---|---|---|---|---|
| 1-756 | + | 763 | 5' UTR (NM_028791, exon 1 of 24) | Cmtr1 | 2 | tRA treated (DSB peaks) |
| 1-536 | + | 635 | intron (NM_019979, intron 1 of 4) | Selenok | 1 | tRA treated (DSB peaks) |
| 1-686 | + | 627 | 3' UTR (NM_008235, exon 4 of 4) | Hes1 | 5 | tRA treated (DSB peaks) |
| 1-687 | + | 673 | TTS (NM_008235) | Hes1 | 1 | tRA treated (DSB peaks) |
| 1-688 | + | 673 | Intergenic | Hes1 | 2 | tRA treated (DSB peaks) |
| 1-941 | + | 589 | Intergenic | Pkn3 | 1 | tRA treated (DSB peaks) |
| 1-689 | + | 616 | Intergenic | Cpn2 | 1 | tRA treated (DSB peaks) |
| 1-1301 | + | 613 | promoter-TSS (NM_001294262) | Agbl5 | 8 | tRA treated (DSB peaks) |
| 1-700 | + | 610 | promoter-TSS (NM_028295) | Pdia5 | 8 | tRA treated (DSB peaks) |
| 1-757 | + | 560 | intron (NM_172618, intron 6 of 11) | Gm6402 | 10 | tRA treated (DSB peaks) |
| 1-1184 | + | 615 | promoter-TSS (NM_009736) | Bag1 | 8 | tRA treated (DSB peaks) |
| 1-1069 | + | 961 | Intergenic | Mecom | 1 | tRA treated (DSB peaks) |
| 1-240 | + | 592 | intron (NM_029344, intron 1 of 3) | Acyp2 | 2 | tRA treated (DSB peaks) |
| 1-690 | + | 814 | Intergenic | Fam43a | 3 | tRA treated (DSB peaks) |
| 1-691 | + | 602 | Intergenic | Fam43a | 6 | tRA treated (DSB peaks) |
| 1-944 | + | 549 | intron (NM_009116, intron 1 of 3) | Prrx2 | 4 | tRA treated (DSB peaks) |
| 1-713 | + | 647 | promoter-TSS (NM_001347240) | Bbx | 8 | tRA treated (DSB peaks) |
| 1-798 | + | 572 | promoter-TSS (NM_001331237) | Ltbp1 | 8 | tRA treated (DSB peaks) |
| 1-631 | + | 572 | promoter-TSS (NM_175399) | Exosc4 | 8 | tRA treated (DSB peaks) |
| 1-388 | + | 610 | promoter-TSS (NM_145927) | Fntb | 8 | tRA treated (DSB peaks) |
| 1-177 | + | 575 | promoter-TSS (NM_008844) | Pip5k1c | 8 | tRA treated (DSB peaks) |
| 1-506 | + | 576 | promoter-TSS (NM_025335) | Tmem167 | 8 | tRA treated (DSB peaks) |
| 1-1402 | + | 653 | Intergenic | Mir29b-1 | 11 | tRA treated (DSB peaks) |
| 1-889 | + | 800 | 5' UTR (NM_031249, exon 1 of 1) | Cstf2t | 3 | tRA treated (DSB peaks) |
| 1-1403 | + | 571 | Intergenic | Mir29b-1 | 4 | tRA treated (DSB peaks) |
| 1-692 | + | 592 | Intergenic | Acap2 | 2 | tRA treated (DSB peaks) |
| 1-1699 | + | 705 | Intergenic | Gm7244 | 2 | tRA treated (DSB peaks) |
| 1-1700 | + | 705 | Intergenic | Gm7244 | 1 | tRA treated (DSB peaks) |
| 19360 | + | 602 | promoter-TSS (NM_001355761) | Ube2f | 8 | tRA treated (DSB peaks) |
| 1-344 | + | 585 | promoter-TSS (NM_011289) | Rpl27 | 8 | tRA treated (DSB peaks) |
| 1-693 | + | 801 | Intergenic | Gm536 | 3 | tRA treated (DSB peaks) |
| 1-1305 | + | 714 | Intergenic | Mrpl33 | 6 | tRA treated (DSB peaks) |
| 1-539 | + | 635 | exon (NM_025295, exon 2 of 4) | Btd | 15 | tRA treated (DSB peaks) |
| 1-981 | + | 641 | promoter-TSS (NM_153126) | Nat10 | 8 | tRA treated (DSB peaks) |
| 1-946 | + | 558 | intron (NM_001283047, intron 2 of 11) | Abl1 | 6 | tRA treated (DSB peaks) |
| 1-1306 | + | 599 | intron (NM_144541, intron 6 of 11) | Babam2 | 6 | tRA treated (DSB peaks) |
| 1-1307 | + | 599 | intron (NM_144541, intron 6 of 11) | Babam2 | 8 | tRA treated (DSB peaks) |
| 1-358 | + | 564 | promoter-TSS (NR_040680) | Gm45916 | 8 | tRA treated (DSB peaks) |
| 1-521 | + | 558 | promoter-TSS (NM_172595) | Arl15 | 8 | tRA treated (DSB peaks) |
| 1-1178 | + | 986 | Intergenic | Map3k7 | 12 | tRA treated (DSB peaks) |
| 1-1179 | + | 561 | Intergenic | Bach2 | 9 | tRA treated (DSB peaks) |
| 1-241 | + | 631 | Intergenic | Ubtd2 | 9 | tRA treated (DSB peaks) |
| 1-1578 | + | 557 | promoter-TSS (NR_029433) | 1110004F10Rik | 8 | tRA treated (DSB peaks) |
| 1-375 | + | 766 | Intergenic | Ccdc71l | 5 | tRA treated (DSB peaks) |
| 1-695 | + | 1000 | Intergenic | Tnk2 | 8 | tRA treated (DSB peaks) |
| 1-822 | + | 596 | intron (NR_045381, intron 4 of 4) | Gypc | 9 | tRA treated (DSB peaks) |
| 1-206 | + | 590 | promoter-TSS (NM_010441) | Hmga2 | 8 | tRA treated (DSB peaks) |

| | | | | | | |
|---|---|---|---|---|---|---|
| 1-890 | + | 605 | Intergenic | Pten | 6 | tRA treated (DSB peaks) |
| 1-1236 | + | 805 | promoter-TSS (NM_172700) | Zmpste24 | 8 | tRA treated (DSB peaks) |
| 1-1254 | + | 547 | promoter-TSS (NM_010948) | Nudc | 8 | tRA treated (DSB peaks) |
| 1-1260 | + | 562 | promoter-TSS (NM_013736) | Eloa | 8 | tRA treated (DSB peaks) |
| 1-696 | + | 642 | intron (NM_172615, intron 1 of 18) | Rubcn | 9 | tRA treated (DSB peaks) |
| 1-947 | + | 573 | intron (NM_146119, intron 5 of 13) | Fam129b | 6 | tRA treated (DSB peaks) |
| 1-1264 | + | 664 | promoter-TSS (NM_007838) | Ddost | 8 | tRA treated (DSB peaks) |
| 1-1145 | + | 573 | promoter-TSS (NM_011828) | Selenof | 8 | tRA treated (DSB peaks) |
| 1-242 | + | 542 | intron (NM_001190885, intron 1 of 8) | Gabrp | 4 | tRA treated (DSB peaks) |
| 32509 | + | 550 | promoter-TSS (NR_040562) | 2810442N19Rik | 8 | tRA treated (DSB peaks) |
| 1-114 | + | 643 | promoter-TSS (NM_198025) | Flvcr1 | 8 | tRA treated (DSB peaks) |
| 1-118 | + | 708 | promoter-TSS (NM_029766) | Ints7 | 8 | tRA treated (DSB peaks) |
| 1-860 | + | 826 | promoter-TSS (NM_009212) | Ighmbp2 | 7 | tRA treated (DSB peaks) |
| 1-1162 | + | 697 | promoter-TSS (NM_026147) | Rps20 | 7 | tRA treated (DSB peaks) |
| 1-243 | + | 893 | intron (NM_001190886, intron 1 of 7) | Kcnip1 | 11 | tRA treated (DSB peaks) |
| 1-918 | + | 565 | promoter-TSS (NM_181399) | Usp6nl | 7 | tRA treated (DSB peaks) |
| 1-1613 | + | 641 | intron (NM_001042675, intron 1 of 6) | Rbpms | 8 | tRA treated (DSB peaks) |
| 1-698 | + | 580 | intron (NM_001145884, intron 10 of 14) | Umps | 6 | tRA treated (DSB peaks) |
| 1-699 | + | 580 | intron (NM_001145884, intron 10 of 14) | Umps | 7 | tRA treated (DSB peaks) |
| 1-877 | + | 595 | promoter-TSS (NM_001362044) | Ubxn1 | 7 | tRA treated (DSB peaks) |
| 1-125 | + | 938 | promoter-TSS (NM_001347361) | Pex3 | 7 | tRA treated (DSB peaks) |
| 1-1684 | + | 589 | promoter-TSS (NM_144933) | Med17 | 7 | tRA treated (DSB peaks) |
| 1-231 | + | 778 | promoter-TSS (NM_033523) | Spred2 | 7 | tRA treated (DSB peaks) |
| 1-377 | + | 652 | Intergenic | Twist1 | 5 | tRA treated (DSB peaks) |
| 1-1695 | + | 735 | promoter-TSS (NR_029585) | Mir199a-1 | 7 | tRA treated (DSB peaks) |
| 1-763 | + | 680 | non-coding (NR_037970, exon 1 of 13) | Brd2 | 5 | tRA treated (DSB peaks) |
| 1-1509 | + | 627 | Intergenic | Gpi1 | 6 | tRA treated (DSB peaks) |
| 1-1510 | + | 627 | Intergenic | Gpi1 | 8 | tRA treated (DSB peaks) |
| 1-1696 | + | 735 | promoter-TSS (NR_029585) | Mir199a-1 | 7 | tRA treated (DSB peaks) |
| 1-1511 | + | 627 | Intergenic | 4931406P16Rik | 3 | tRA treated (DSB peaks) |
| 1-750 | + | 750 | promoter-TSS (NM_001285982) | Fam173a | 7 | tRA treated (DSB peaks) |
| 1-1512 | + | 660 | Intergenic | Lsm14a | 11 | tRA treated (DSB peaks) |
| 1-137 | + | 591 | exon (NM_176968, exon 1 of 12) | Nt5dc1 | 3 | tRA treated (DSB peaks) |
| 1-892 | + | 622 | Intergenic | Ch25h | 7 | tRA treated (DSB peaks) |
| 1-1506 | + | 887 | promoter-TSS (NM_001083912) | Plekhg2 | 7 | tRA treated (DSB peaks) |
| 1-942 | + | 649 | promoter-TSS (NM_001281829) | Phyhd1 | 7 | tRA treated (DSB peaks) |
| 1-454 | + | 546 | promoter-TSS (NM_009256) | Serpinb9 | 7 | tRA treated (DSB peaks) |
| 1-608 | + | 543 | promoter-TSS (NM_175009) | Eny2 | 7 | tRA treated (DSB peaks) |
| 1-609 | + | 543 | promoter-TSS (NM_175009) | Eny2 | 7 | tRA treated (DSB peaks) |
| 1-1405 | + | 639 | intron (NM_145575, intron 1 of 11) | Cald1 | 19 | tRA treated (DSB peaks) |
| 1-456 | + | 747 | Intergenic | Fam50b | 10 | tRA treated (DSB peaks) |
| 1-708 | + | 567 | promoter-TSS (NM_001252442) | Phldb2 | 7 | tRA treated (DSB peaks) |
| 1-552 | + | 923 | promoter-TSS (NM_021472) | Ang | 7 | tRA treated (DSB peaks) |
| 1-827 | + | 600 | Intergenic | Egr1 | 3 | tRA treated (DSB peaks) |
| 1-1182 | + | 748 | Intergenic | Zfp292 | 3 | tRA treated (DSB peaks) |
| 1-1183 | + | 749 | Intergenic | Zfp292 | 1 | tRA treated (DSB peaks) |
| 1-244 | + | 989 | Intergenic | Slit3 | 3 | tRA treated (DSB peaks) |

| | | | | | | |
|---|---|---|---|---|---|---|
| 1-245 | + | 991 | Intergenic | Slit3 | 3 | tRA treated (DSB peaks) |
| 1-553 | + | 547 | promoter-TSS (NM_001113358) | Abhd4 | 7 | tRA treated (DSB peaks) |
| 1-1702 | + | 657 | Intergenic | 4933422A05Rik | 3 | tRA treated (DSB peaks) |
| 1-277 | + | 1000 | promoter-TSS (NM_021354) | Drg2 | 7 | tRA treated (DSB peaks) |
| 1-384 | + | 545 | promoter-TSS (NM_009093) | Rps29 | 7 | tRA treated (DSB peaks) |
| 1-1631 | + | 655 | promoter-TSS (NM_001142323) | Haus8 | 7 | tRA treated (DSB peaks) |
| 1-701 | + | 640 | Intergenic | 1600019K03Rik | 2 | tRA treated (DSB peaks) |
| 1-1633 | + | 547 | promoter-TSS (NM_023126) | Rab8a | 7 | tRA treated (DSB peaks) |
| 1-963 | + | 655 | promoter-TSS (NM_001190297) | Gpr155 | 7 | tRA treated (DSB peaks) |
| 1-499 | + | 628 | promoter-TSS (NM_138953) | Ell2 | 7 | tRA treated (DSB peaks) |
| 1-1754 | + | 631 | promoter-TSS (NM_134255) | Elovl5 | 7 | tRA treated (DSB peaks) |
| 1-168 | + | 712 | promoter-TSS (NM_029272) | Ndufs7 | 7 | tRA treated (DSB peaks) |
| 1-1309 | + | 571 | intron (NM_027373, intron 1 of 16) | Afap1 | 2 | tRA treated (DSB peaks) |
| 43221 | + | 680 | Intergenic | Hs6st1 | 3 | tRA treated (DSB peaks) |
| 1-650 | + | 636 | promoter-TSS (NM_020507) | Tob2 | 7 | tRA treated (DSB peaks) |
| 1-1075 | + | 578 | intron (NM_009673, intron 12 of 12) | Mir7009 | 4 | tRA treated (DSB peaks) |
| 1-186 | + | 737 | promoter-TSS (NM_027423) | Polr3b | 7 | tRA treated (DSB peaks) |
| 1-396 | + | 611 | promoter-TSS (NM_023409) | Npc2 | 7 | tRA treated (DSB peaks) |
| 1-727 | + | 548 | promoter-TSS (NM_011434) | Sod1 | 7 | tRA treated (DSB peaks) |
| 1-893 | + | 731 | intron (NR_033510, intron 1 of 4) | Hectd2os | 4 | tRA treated (DSB peaks) |
| 1-894 | + | 731 | intron (NR_033510, intron 1 of 4) | Hectd2os | 3 | tRA treated (DSB peaks) |
| 1-1109 | + | 717 | promoter-TSS (NM_011342) | Sec22b | 7 | tRA treated (DSB peaks) |
| 1-1770 | + | 860 | promoter-TSS (NM_001324534) | Rpl29 | 7 | tRA treated (DSB peaks) |
| 1-1773 | + | 662 | promoter-TSS (NM_001012236) | Trex1 | 7 | tRA treated (DSB peaks) |
| 1-1786 | + | 540 | promoter-TSS (NM_178660) | Rbms3 | 7 | tRA treated (DSB peaks) |
| 1-897 | + | 761 | exon (NM_028263, exon 2 of 2) | Fgfbp3 | 1 | tRA treated (DSB peaks) |
| 1-996 | + | 905 | promoter-TSS (NM_001356467) | Ndufaf1 | 7 | tRA treated (DSB peaks) |
| 1-138 | + | 643 | exon (NM_008538, exon 2 of 2) | Marcks | 7 | tRA treated (DSB peaks) |
| 1-139 | + | 686 | exon (NM_008538, exon 2 of 2) | Marcks | 4 | tRA treated (DSB peaks) |
| 1-140 | + | 681 | Intergenic | Marcks | 4 | tRA treated (DSB peaks) |
| 1-141 | + | 594 | Intergenic | Marcks | 6 | tRA treated (DSB peaks) |
| 1-1406 | + | 687 | Intergenic | Akr1d1 | 5 | tRA treated (DSB peaks) |
| 1-1407 | + | 687 | Intergenic | Akr1d1 | 4 | tRA treated (DSB peaks) |
| 1-703 | + | 566 | intron (NM_008047, intron 2 of 10) | Fstl1 | 3 | tRA treated (DSB peaks) |
| 1-457 | + | 560 | intron (NM_001039188, intron 1 of 9) | Rreb1 | 1 | tRA treated (DSB peaks) |
| 1-458 | + | 560 | intron (NM_001039188, intron 1 of 9) | Rreb1 | 2 | tRA treated (DSB peaks) |
| 1-589 | + | 802 | promoter-TSS (NR_131000) | 1810041H14Rik | 7 | tRA treated (DSB peaks) |
| 1-1353 | + | 631 | promoter-TSS (NM_030241) | Kmt5a | 7 | tRA treated (DSB peaks) |
| 1-1514 | + | 543 | Intergenic | Ccne1 | 1 | tRA treated (DSB peaks) |
| 1-1244 | + | 688 | promoter-TSS (NM_029157) | Sf3a3 | 7 | tRA treated (DSB peaks) |
| 24838 | + | 568 | promoter-TSS (NM_001081756) | Nckap5 | 7 | tRA treated (DSB peaks) |
| 1-1015 | + | 568 | promoter-TSS (NM_011045) | Pcna | 7 | tRA treated (DSB peaks) |
| 1-1390 | + | 572 | promoter-TSS (NM_181730) | Lnx2 | 7 | tRA treated (DSB peaks) |
| 1-1285 | + | 825 | promoter-TSS (NM_001039157) | Ube2j2 | 7 | tRA treated (DSB peaks) |
| 1-1409 | + | 692 | Intergenic | Fmc1 | 13 | tRA treated (DSB peaks) |
| 1-1410 | + | 568 | intron (NM_001170848, intron 1 of 8) | Luc7l2 | 4 | tRA treated (DSB peaks) |
| 1-1286 | + | 585 | promoter-TSS (NM_145557) | 9430015G10Rik | 7 | tRA treated (DSB peaks) |

| | | | | | | |
|---|---|---|---|---|---|---|
| 1-142 | + | 854 | Intergenic | Lama4 | 6 | tRA treated (DSB peaks) |
| 1-143 | + | 629 | intron (NM_010681, intron 1 of 37) | Lama4 | 1 | tRA treated (DSB peaks) |
| 1-1604 | + | 704 | promoter-TSS (NM_015801) | Pnpla6 | 6 | tRA treated (DSB peaks) |
| 1-460 | + | 611 | Intergenic | Slc35b3 | 5 | tRA treated (DSB peaks) |
| 1 | + | 822 | promoter-TSS (NM_011541) | Tcea1 | 6 | tRA treated (DSB peaks) |
| 1-144 | + | 718 | intron (NR_038037, intron 2 of 2) | Gm16364 | 4 | tRA treated (DSB peaks) |
| 1-145 | + | 718 | intron (NR_038037, intron 2 of 2) | Gm16364 | 2 | tRA treated (DSB peaks) |
| 43313 | + | 607 | intron (NM_001145660, intron 2 of 14) | Rfx8 | 5 | tRA treated (DSB peaks) |
| 43344 | + | 633 | intron (NM_001025602, intron 6 of 10) | Il1rl1 | 13 | tRA treated (DSB peaks) |
| 43374 | + | 633 | intron (NM_001025602, intron 6 of 10) | Il1rl1 | 2 | tRA treated (DSB peaks) |
| 1-865 | + | 696 | promoter-TSS (NM_027236) | Eif1ad | 6 | tRA treated (DSB peaks) |
| 1-1058 | + | 1000 | promoter-TSS (NM_001163301) | Pex2 | 6 | tRA treated (DSB peaks) |
| 1-247 | + | 611 | exon (NM_001199274, exon 1 of 7) | Mat2b | 5 | tRA treated (DSB peaks) |
| 1-1703 | + | 615 | intron (NM_133733, intron 1 of 6) | Clmp | 1 | tRA treated (DSB peaks) |
| 1-122 | + | 638 | promoter-TSS (NM_175374) | Mtrf1l | 6 | tRA treated (DSB peaks) |
| 1-1076 | + | 540 | Intergenic | Pgrmc2 | 3 | tRA treated (DSB peaks) |
| 1-146 | + | 635 | intron (NM_133999, intron 1 of 22) | Fig4 | 4 | tRA treated (DSB peaks) |
| 1-147 | + | 635 | intron (NM_133999, intron 1 of 22) | Fig4 | 3 | tRA treated (DSB peaks) |
| 1-874 | + | 585 | promoter-TSS (NM_001360286) | Fkbp2 | 6 | tRA treated (DSB peaks) |
| 1-883 | + | 930 | promoter-TSS (NM_028411) | Cyb561a3 | 6 | tRA treated (DSB peaks) |
| 1-1704 | + | 791 | intron (NR_131930, intron 2 of 3) | Mir100 | 5 | tRA treated (DSB peaks) |
| 1-1705 | + | 691 | Intergenic | Mir125b-1 | 4 | tRA treated (DSB peaks) |
| 1-1706 | + | 571 | Intergenic | Mir125b-1 | 3 | tRA treated (DSB peaks) |
| 1-1707 | + | 574 | Intergenic | Sorl1 | 10 | tRA treated (DSB peaks) |
| 1-1708 | + | 640 | Intergenic | Sorl1 | 4 | tRA treated (DSB peaks) |
| 1-1709 | + | 1000 | Intergenic | Sorl1 | 8 | tRA treated (DSB peaks) |
| 1-1710 | + | 633 | Intergenic | Sorl1 | 6 | tRA treated (DSB peaks) |
| 1-705 | + | 664 | Intergenic | Mir6540 | 4 | tRA treated (DSB peaks) |
| 1-706 | + | 664 | Intergenic | Mir6540 | 3 | tRA treated (DSB peaks) |
| 1-148 | + | 738 | Intergenic | Foxo3 | 1 | tRA treated (DSB peaks) |
| 1-1168 | + | 1000 | promoter-TSS (NM_001081201) | Dpy19l4 | 6 | tRA treated (DSB peaks) |
| 1-738 | + | 630 | promoter-TSS (NM_011581) | Thbs2 | 6 | tRA treated (DSB peaks) |
| 1-461 | + | 568 | Intergenic | Phactr1 | 6 | tRA treated (DSB peaks) |
| 1-462 | + | 723 | Intergenic | Phactr1 | 2 | tRA treated (DSB peaks) |
| 1-1499 | + | 667 | promoter-TSS (NM_010320) | Gng8 | 6 | tRA treated (DSB peaks) |
| 1-1688 | + | 760 | promoter-TSS (NM_025844) | Chordc1 | 6 | tRA treated (DSB peaks) |
| 1-887 | + | 566 | promoter-TSS (NM_153808) | Smc5 | 6 | tRA treated (DSB peaks) |
| 1-1186 | + | 611 | intron (NM_001037709, intron 1 of 11) | Rusc2 | 3 | tRA treated (DSB peaks) |
| 1-1187 | + | 611 | intron (NM_001037709, intron 1 of 11) | Rusc2 | 6 | tRA treated (DSB peaks) |
| 1-1068 | + | 559 | promoter-TSS (NM_178772) | Nceh1 | 6 | tRA treated (DSB peaks) |
| 1-753 | + | 578 | promoter-TSS (NM_011145) | Ppard | 6 | tRA treated (DSB peaks) |
| 1-754 | + | 579 | promoter-TSS (NM_001163820) | Fance | 6 | tRA treated (DSB peaks) |
| 1-1300 | + | 613 | promoter-TSS (NM_001294262) | Agbl5 | 6 | tRA treated (DSB peaks) |
| 1-1311 | + | 575 | Intergenic | Fgfbp1 | 3 | tRA treated (DSB peaks) |
| 1-1303 | + | 997 | promoter-TSS (NM_009206) | Slc4a1ap | 6 | tRA treated (DSB peaks) |
| 1-764 | + | 598 | promoter-TSS (NM_001252468) | Bag6 | 6 | tRA treated (DSB peaks) |
| 1-949 | + | 553 | intron (NM_001301831, intron 10 of 14) | Arhgap15os | 2 | tRA treated (DSB peaks) |

| | | | | | | |
|---|---|---|---|---|---|---|
| 1-246 | + | 560 | promoter-TSS (NM_145962) | Pank3 | 6 | tRA treated (DSB peaks) |
| 1-895 | + | 609 | promoter-TSS (NM_016854) | Ppp1r3c | 6 | tRA treated (DSB peaks) |
| 1-771 | + | 682 | intron (NM_172621, intron 4 of 5) | Clic5 | 3 | tRA treated (DSB peaks) |
| 1-772 | + | 681 | intron (NM_172621, intron 4 of 5) | Clic5 | 1 | tRA treated (DSB peaks) |
| 1-901 | + | 698 | 5' UTR (NM_009128, exon 1 of 6) | Scd2 | 11 | tRA treated (DSB peaks) |
| 1-1185 | + | 659 | promoter-TSS (NM_026893) | Dcaf12 | 6 | tRA treated (DSB peaks) |
| 1-1188 | + | 576 | promoter-TSS (NM_199057) | Rusc2 | 6 | tRA treated (DSB peaks) |
| 43405 | + | 616 | promoter-TSS (NM_011729) | Ercc5 | 6 | tRA treated (DSB peaks) |
| 1-1617 | + | 551 | promoter-TSS (NM_007450) | Slc25a4 | 6 | tRA treated (DSB peaks) |
| 1-773 | + | 650 | intron (NR_073425, intron 5 of 5) | Runx2 | 5 | tRA treated (DSB peaks) |
| 1-913 | + | 571 | promoter-TSS (NM_001355242) | Shoc2 | 6 | tRA treated (DSB peaks) |
| 1-707 | + | 541 | exon (NM_001356392, exon 1 of 5) | Gtpbp8 | 7 | tRA treated (DSB peaks) |
| 1-1423 | + | 738 | promoter-TSS (NM_026637) | Ggct | 6 | tRA treated (DSB peaks) |
| 43435 | + | 675 | Intergenic | Col3a1 | 3 | tRA treated (DSB peaks) |
| 41275 | + | 675 | Intergenic | Col3a1 | 1 | tRA treated (DSB peaks) |
| 1-1518 | + | 592 | intron (NM_001163684, intron 1 of 8) | Nosip | 4 | tRA treated (DSB peaks) |
| 1-249 | + | 682 | Intergenic | Gm12159 | 5 | tRA treated (DSB peaks) |
| 1-774 | + | 594 | Intergenic | Cdc5l | 6 | tRA treated (DSB peaks) |
| 1-478 | + | 604 | promoter-TSS (NM_134059) | Ddx41 | 6 | tRA treated (DSB peaks) |
| 1-782 | + | 582 | promoter-TSS (NR_029827) | A230051N06Rik | 6 | tRA treated (DSB peaks) |
| 1-717 | + | 645 | promoter-TSS (NM_001358433) | Tbc1d23 | 6 | tRA treated (DSB peaks) |
| 1-1196 | + | 585 | promoter-TSS (NM_022814) | Svep1 | 6 | tRA treated (DSB peaks) |
| 41640 | + | 596 | 5' UTR (NM_007737, exon 1 of 54) | Col5a2 | 26 | tRA treated (DSB peaks) |
| 1-621 | + | 642 | promoter-TSS (NM_144549) | Trib1 | 6 | tRA treated (DSB peaks) |
| 42005 | + | 595 | Intergenic | Wdr75 | 9 | tRA treated (DSB peaks) |
| 1-486 | + | 585 | promoter-TSS (NM_008086) | Gas1 | 6 | tRA treated (DSB peaks) |
| 1-466 | + | 624 | intron (NM_001199304, intron 4 of 6) | Atxn1 | 12 | tRA treated (DSB peaks) |
| 42370 | + | 589 | Intergenic | Wdr75 | 12 | tRA treated (DSB peaks) |
| 1-1201 | + | 540 | promoter-TSS (NM_177607) | 4933430I17Rik | 6 | tRA treated (DSB peaks) |
| 1-709 | + | 577 | Intergenic | Phldb2 | 5 | tRA treated (DSB peaks) |
| 1-710 | + | 577 | Intergenic | Phldb2 | 10 | tRA treated (DSB peaks) |
| 1-250 | + | 584 | Intergenic | Lsm11 | 2 | tRA treated (DSB peaks) |
| 1-721 | + | 617 | promoter-TSS (NM_001347245) | Zfp654 | 6 | tRA treated (DSB peaks) |
| 1-568 | + | 568 | promoter-TSS (NM_001110162) | Kctd9 | 6 | tRA treated (DSB peaks) |
| 1-903 | + | 638 | 5' UTR (NM_176785, exon 1 of 1) | Hps6 | 1 | tRA treated (DSB peaks) |
| 1-501 | + | 577 | promoter-TSS (NM_178916) | Rfesd | 6 | tRA treated (DSB peaks) |
| 1-1618 | + | 739 | Intergenic | Acsl1 | 6 | tRA treated (DSB peaks) |
| 1-542 | + | 731 | Intergenic | Bmp4 | 3 | tRA treated (DSB peaks) |
| 1-543 | + | 732 | Intergenic | Bmp4 | 2 | tRA treated (DSB peaks) |
| 1-968 | + | 580 | promoter-TSS (NM_011871) | Prkra | 6 | tRA treated (DSB peaks) |
| 1-315 | + | 553 | promoter-TSS (NM_134025) | Pex12 | 6 | tRA treated (DSB peaks) |
| 1-1209 | + | 568 | promoter-TSS (NR_152448) | Snapc3 | 6 | tRA treated (DSB peaks) |
| 1-905 | + | 691 | exon (NM_025563, exon 1 of 5) | Borcs7 | 5 | tRA treated (DSB peaks) |
| 1-1312 | + | 804 | Intergenic | 4930405L22Rik | 10 | tRA treated (DSB peaks) |
| 1-1096 | + | 556 | promoter-TSS (NM_033573) | Prcc | 6 | tRA treated (DSB peaks) |
| 1-728 | + | 568 | promoter-TSS (NM_026502) | 1110004E09Rik | 6 | tRA treated (DSB peaks) |
| 1-328 | + | 688 | promoter-TSS (NM_146024) | Ankrd40 | 6 | tRA treated (DSB peaks) |

| | | | | | | |
|---|---|---|---|---|---|---|
| 1-1620 | + | 618 | intron (NM_177304, intron 7 of 7) | Enpp6 | 6 | tRA treated (DSB peaks) |
| 1-906 | + | 646 | Intergenic | Neurl1a | 4 | tRA treated (DSB peaks) |
| 1-907 | + | 623 | intron (NM_008018, intron 6 of 14) | Mir6995 | 5 | tRA treated (DSB peaks) |
| 1-908 | + | 623 | intron (NM_008018, intron 6 of 14) | Mir6995 | 6 | tRA treated (DSB peaks) |
| 1-1192 | + | 789 | Intergenic | Sec61b | 11 | tRA treated (DSB peaks) |
| 1-1414 | + | 689 | Intergenic | Rn4.5s | 7 | tRA treated (DSB peaks) |
| 1-510 | + | 564 | promoter-TSS (NM_001271464) | Gfm2 | 6 | tRA treated (DSB peaks) |
| 20455 | + | 609 | promoter-TSS (NM_173760) | Ppip5k2 | 6 | tRA treated (DSB peaks) |
| 1-1415 | + | 566 | Intergenic | Mir704 | 18 | tRA treated (DSB peaks) |
| 1-1416 | + | 545 | 5' UTR (NM_001145881, exon 1 of 5) | Zfp212 | 1 | tRA treated (DSB peaks) |
| 1-1621 | + | 590 | intron (NM_133791, intron 1 of 22) | Wwc2 | 3 | tRA treated (DSB peaks) |
| 1-1313 | + | 641 | intron (NR_045500, intron 3 of 3) | 5730480H06Rik | 5 | tRA treated (DSB peaks) |
| 1-664 | + | 667 | promoter-TSS (NM_026468) | Atp5g2 | 6 | tRA treated (DSB peaks) |
| 1-546 | + | 591 | exon (NM_172600, exon 1 of 16) | Tmem260 | 1 | tRA treated (DSB peaks) |
| 1-410 | + | 655 | promoter-TSS (NM_181328) | Slc25a29 | 6 | tRA treated (DSB peaks) |
| 1-253 | + | 797 | Intergenic | Trim41 | 9 | tRA treated (DSB peaks) |
| 1-254 | + | 560 | Intergenic | Trim41 | 6 | tRA treated (DSB peaks) |
| 1-255 | + | 735 | Intergenic | Trim7 | 12 | tRA treated (DSB peaks) |
| 1-256 | + | 580 | Intergenic | Trim7 | 13 | tRA treated (DSB peaks) |
| 1-257 | + | 594 | Intergenic | Irgm1 | 3 | tRA treated (DSB peaks) |
| 1-258 | + | 595 | Intergenic | Irgm1 | 6 | tRA treated (DSB peaks) |
| 1-1576 | + | 557 | promoter-TSS (NM_011965) | Psma1 | 6 | tRA treated (DSB peaks) |
| 1-1228 | + | 545 | promoter-TSS (NM_177670) | Tmem69 | 6 | tRA treated (DSB peaks) |
| 1-995 | + | 618 | promoter-TSS (NM_019769) | Exd1 | 6 | tRA treated (DSB peaks) |
| 1-1003 | + | 541 | promoter-TSS (NM_001081091) | Cep152 | 6 | tRA treated (DSB peaks) |
| 1-1248 | + | 591 | promoter-TSS (NM_011970) | Psmb2 | 6 | tRA treated (DSB peaks) |
| 1-1713 | + | 589 | intron (NM_010875, intron 1 of 15) | Gm11149 | 2 | tRA treated (DSB peaks) |
| 1-1714 | + | 589 | intron (NM_010875, intron 1 of 15) | Gm11149 | 4 | tRA treated (DSB peaks) |
| 1-1077 | + | 838 | Intergenic | Pcdh18 | 14 | tRA treated (DSB peaks) |
| 1-1078 | + | 839 | Intergenic | Pcdh18 | 10 | tRA treated (DSB peaks) |
| 1-1314 | + | 610 | Intergenic | Adgra3 | 1 | tRA treated (DSB peaks) |
| 1-1584 | + | 717 | promoter-TSS (NM_029420) | Bola2 | 6 | tRA treated (DSB peaks) |
| 1-1716 | + | 581 | Intergenic | Ncam1 | 10 | tRA treated (DSB peaks) |
| 1-1362 | + | 626 | promoter-TSS (NM_133895) | Slc15a4 | 6 | tRA treated (DSB peaks) |
| 1-1717 | + | 815 | Intergenic | Ncam1 | 5 | tRA treated (DSB peaks) |
| 1-1585 | + | 581 | promoter-TSS (NM_172749) | Zfp668 | 6 | tRA treated (DSB peaks) |
| 1-1016 | + | 569 | promoter-TSS (NM_011045) | Pcna | 6 | tRA treated (DSB peaks) |
| 1-1368 | + | 628 | promoter-TSS (NM_010717) | Limk1 | 6 | tRA treated (DSB peaks) |
| 1-1418 | + | 652 | intron (NM_001163645, intron 2 of 21) | Osbpl3 | 5 | tRA treated (DSB peaks) |
| 1-1419 | + | 610 | intron (NM_001163645, intron 2 of 21) | Osbpl3 | 2 | tRA treated (DSB peaks) |
| 1-1376 | + | 728 | promoter-TSS (NM_010312) | Gnb2 | 6 | tRA treated (DSB peaks) |
| 1-1493 | + | 713 | promoter-TSS (NM_001113424) | Amn1 | 6 | tRA treated (DSB peaks) |
| 1-101 | + | 593 | promoter-TSS (NM_153064) | Ndufs2 | 6 | tRA treated (DSB peaks) |
| 1-1719 | + | 582 | exon (NM_133981, exon 1 of 15) | Alg9 | 2 | tRA treated (DSB peaks) |
| 1-549 | + | 999 | Intergenic | Olfr750 | 6 | tRA treated (DSB peaks) |
| 1-550 | + | 632 | Intergenic | Ang | 10 | tRA treated (DSB peaks) |
| 1-551 | + | 825 | Intergenic | Ang | 17 | tRA treated (DSB peaks) |

| | | | | | | |
|---|---|---|---|---|---|---|
| 1-1056 | + | 787 | promoter-TSS (NM_016775) | Dnajc5 | 6 | tRA treated (DSB peaks) |
| 1-951 | + | 568 | intron (NR_040636, intron 1 of 2) | Gm13497 | 8 | tRA treated (DSB peaks) |
| 1-1079 | + | 729 | Intergenic | Elf2 | 7 | tRA treated (DSB peaks) |
| 1-1080 | + | 631 | Intergenic | Elf2 | 10 | tRA treated (DSB peaks) |
| 1-1081 | + | 711 | 5' UTR (NM_016858, exon 1 of 2) | Rab33b | 1 | tRA treated (DSB peaks) |
| 1-263 | + | 633 | intron (NM_001048061, intron 2 of 7) | Hnrnpab | 5 | tRA treated (DSB peaks) |
| 1-667 | + | 558 | promoter-TSS (NM_026508) | Trap1 | 5 | tRA treated (DSB peaks) |
| 42736 | + | 602 | Intergenic | Gm17767 | 6 | tRA treated (DSB peaks) |
| 1-610 | + | 944 | intron (NM_080635, intron 3 of 7) | Eif3h | 10 | tRA treated (DSB peaks) |
| 1-469 | + | 603 | intron (NM_011817, intron 1 of 3) | Gadd45g | 10 | tRA treated (DSB peaks) |
| 1-594 | + | 579 | promoter-TSS (NM_026069) | Rpl37 | 5 | tRA treated (DSB peaks) |
| 1-470 | + | 748 | Intergenic | Gadd45g | 6 | tRA treated (DSB peaks) |
| 1-471 | + | 674 | Intergenic | 4921525O09Rik | 7 | tRA treated (DSB peaks) |
| 1-472 | + | 675 | Intergenic | 4921525O09Rik | 8 | tRA treated (DSB peaks) |
| 1-1420 | + | 597 | Intergenic | Hoxa7 | 7 | tRA treated (DSB peaks) |
| 1-1059 | + | 602 | promoter-TSS (NM_001163301) | Pex2 | 5 | tRA treated (DSB peaks) |
| 1-378 | + | 623 | Intergenic | Gm35135 | 3 | tRA treated (DSB peaks) |
| 1-379 | + | 921 | Intergenic | Arhgap5 | 11 | tRA treated (DSB peaks) |
| 1-833 | + | 656 | 3' UTR (NM_025749, exon 2 of 2) | 1700034E13Rik | 15 | tRA treated (DSB peaks) |
| 1-882 | + | 621 | promoter-TSS (NM_019699) | Fads2 | 5 | tRA treated (DSB peaks) |
| 1-1683 | + | 589 | promoter-TSS (NM_144933) | Med17 | 5 | tRA treated (DSB peaks) |
| 1-1082 | + | 541 | intron (NM_175386, intron 2 of 3) | Lhfp | 3 | tRA treated (DSB peaks) |
| 1-1083 | + | 541 | intron (NM_175386, intron 2 of 3) | Lhfp | 4 | tRA treated (DSB peaks) |
| 1-611 | + | 582 | intron (NM_010162, intron 1 of 10) | Ext1 | 4 | tRA treated (DSB peaks) |
| 1-1193 | + | 636 | intron (NM_013454, intron 1 of 49) | Abca1 | 2 | tRA treated (DSB peaks) |
| 1-612 | + | 540 | intron (NM_010162, intron 1 of 10) | Ext1 | 4 | tRA treated (DSB peaks) |
| 1-613 | + | 540 | intron (NM_010162, intron 1 of 10) | Ext1 | 9 | tRA treated (DSB peaks) |
| 1-909 | + | 571 | Intergenic | Dusp5 | 7 | tRA treated (DSB peaks) |
| 1-910 | + | 571 | Intergenic | Dusp5 | 10 | tRA treated (DSB peaks) |
| 1-1297 | + | 782 | promoter-TSS (NM_016736) | Nub1 | 5 | tRA treated (DSB peaks) |
| 1-130 | + | 654 | promoter-TSS (NM_027347) | Med23 | 5 | tRA treated (DSB peaks) |
| 1-911 | + | 674 | intron (NM_001170847, intron 1 of 13) | Rbm20 | 2 | tRA treated (DSB peaks) |
| 1-749 | + | 580 | promoter-TSS (NM_026238) | Narfl | 5 | tRA treated (DSB peaks) |
| 1-685 | + | 863 | promoter-TSS (NM_001199177) | Opa1 | 5 | tRA treated (DSB peaks) |
| 1-760 | + | 765 | promoter-TSS (NM_009059) | Rgl2 | 5 | tRA treated (DSB peaks) |
| 1-1308 | + | 771 | promoter-TSS (NM_010414) | Htt | 5 | tRA treated (DSB peaks) |
| 1-266 | + | 636 | intron (NM_019417, intron 2 of 6) | Pdlim4 | 5 | tRA treated (DSB peaks) |
| 1-829 | + | 706 | promoter-TSS (NM_001305627) | Hars2 | 5 | tRA treated (DSB peaks) |
| 43282 | + | 870 | promoter-TSS (NM_026913) | Mrpl30 | 5 | tRA treated (DSB peaks) |
| 1-835 | + | 576 | Intergenic | Redrum | 6 | tRA treated (DSB peaks) |
| 1-836 | + | 575 | Intergenic | Redrum | 4 | tRA treated (DSB peaks) |
| 1-837 | + | 619 | Intergenic | Redrum | 14 | tRA treated (DSB peaks) |
| 1-1408 | + | 633 | promoter-TSS (NM_153600) | Ttc26 | 5 | tRA treated (DSB peaks) |
| 1-554 | + | 578 | Intergenic | Mmp14 | 9 | tRA treated (DSB peaks) |
| 1-1310 | + | 672 | promoter-TSS (NM_010474) | Hs3st1 | 5 | tRA treated (DSB peaks) |
| 1-259 | + | 582 | promoter-TSS (NM_008326) | Irgm1 | 5 | tRA treated (DSB peaks) |
| 1-950 | + | 588 | promoter-TSS (NM_001348198) | Mmadhc | 5 | tRA treated (DSB peaks) |

| | | | | | | |
|---|---|---|---|---|---|---|
| 1-149 | + | 707 | promoter-TSS (NM_030250) | Nus1 | 5 | tRA treated (DSB peaks) |
| 1-1422 | + | 737 | promoter-TSS (NM_026637) | Ggct | 5 | tRA treated (DSB peaks) |
| 1-614 | + | 547 | promoter-TSS (NM_134092) | Mrpl13 | 5 | tRA treated (DSB peaks) |
| 1-475 | + | 594 | Intergenic | Gprin1 | 5 | tRA treated (DSB peaks) |
| 1-838 | + | 542 | intron (NM_001360787, intron 2 of 8) | Zfp608 | 7 | tRA treated (DSB peaks) |
| 1-839 | + | 777 | intron (NM_175751, intron 2 of 9) | Zfp608 | 6 | tRA treated (DSB peaks) |
| 1-840 | + | 676 | intron (NM_175751, intron 2 of 9) | Zfp608 | 5 | tRA treated (DSB peaks) |
| 1-780 | + | 703 | promoter-TSS (NM_001252471) | Sh3gl1 | 5 | tRA treated (DSB peaks) |
| 1-784 | + | 647 | promoter-TSS (NM_018730) | Rpl36 | 5 | tRA treated (DSB peaks) |
| 1-841 | + | 904 | Intergenic | Zfp608 | 8 | tRA treated (DSB peaks) |
| 1-842 | + | 751 | Intergenic | Zfp608 | 4 | tRA treated (DSB peaks) |
| 1-843 | + | 788 | Intergenic | Zfp608 | 4 | tRA treated (DSB peaks) |
| 1-620 | + | 599 | promoter-TSS (NM_026746) | Nsmce2 | 5 | tRA treated (DSB peaks) |
| 1-1198 | + | 574 | promoter-TSS (NM_001013577) | Inip | 5 | tRA treated (DSB peaks) |
| 45658 | + | 650 | promoter-TSS (NM_001160039) | Ndufs1 | 5 | tRA treated (DSB peaks) |
| 1-380 | + | 561 | exon (NM_010907, exon 1 of 6) | Nfkbia | 1 | tRA treated (DSB peaks) |
| 1-476 | + | 647 | intron (NM_001114088, intron 2 of 12) | Pdlim7 | 9 | tRA treated (DSB peaks) |
| 1-477 | + | 647 | intron (NM_001114088, intron 2 of 12) | Pdlim7 | 8 | tRA treated (DSB peaks) |
| 1-159 | + | 698 | promoter-TSS (NM_177612) | Ctnna3 | 5 | tRA treated (DSB peaks) |
| 1-493 | + | 583 | promoter-TSS (NM_023507) | Ercc6l2 | 5 | tRA treated (DSB peaks) |
| 1-1741 | + | 974 | promoter-TSS (NM_001170907) | Trip4 | 5 | tRA treated (DSB peaks) |
| 1-1195 | + | 653 | Intergenic | Klf4 | 6 | tRA treated (DSB peaks) |
| 1-714 | + | 623 | Intergenic | Nfkbiz | 9 | tRA treated (DSB peaks) |
| 1-715 | + | 622 | Intergenic | Nfkbiz | 3 | tRA treated (DSB peaks) |
| 1-559 | + | 613 | Intergenic | Nfatc4 | 2 | tRA treated (DSB peaks) |
| 1-779 | + | 606 | intron (NM_145490, intron 1 of 3) | Zfp959 | 2 | tRA treated (DSB peaks) |
| 1-1744 | + | 722 | promoter-TSS (NM_001311101) | Rps27l | 5 | tRA treated (DSB peaks) |
| 1-386 | + | 835 | promoter-TSS (NM_027117) | Klhdc2 | 5 | tRA treated (DSB peaks) |
| 1-1720 | + | 610 | intron (NM_001278274, intron 1 of 1) | Rcn2 | 2 | tRA treated (DSB peaks) |
| 1-495 | + | 570 | promoter-TSS (NM_144837) | Ice1 | 5 | tRA treated (DSB peaks) |
| 1-629 | + | 742 | promoter-TSS (NM_180662) | Trappc9 | 5 | tRA treated (DSB peaks) |
| 11689 | + | 680 | promoter-TSS (NM_001310668) | Zfp142 | 5 | tRA treated (DSB peaks) |
| 1-500 | + | 577 | promoter-TSS (NM_178916) | Rfesd | 5 | tRA treated (DSB peaks) |
| 1-1723 | + | 616 | intron (NM_025812, intron 1 of 9) | Hmg20a | 10 | tRA treated (DSB peaks) |
| 1-783 | + | 677 | Intergenic | Znrf4 | 2 | tRA treated (DSB peaks) |
| 1-575 | + | 643 | promoter-TSS (NM_026177) | Nufip1 | 5 | tRA treated (DSB peaks) |
| 1-151 | + | 553 | Intergenic | Gm36595 | 16 | tRA treated (DSB peaks) |
| 1-152 | + | 546 | Intergenic | Gm36595 | 13 | tRA treated (DSB peaks) |
| 1-632 | + | 855 | promoter-TSS (NM_001164173) | Cpsf1 | 5 | tRA treated (DSB peaks) |
| 1-1640 | + | 1000 | promoter-TSS (NM_178267) | Zfp827 | 5 | tRA treated (DSB peaks) |
| 1-1641 | + | 681 | promoter-TSS (NR_045822) | Mmaa | 5 | tRA treated (DSB peaks) |
| 1-1533 | + | 1000 | promoter-TSS (NM_009682) | Ap3s2 | 5 | tRA treated (DSB peaks) |
| 1-313 | + | 714 | promoter-TSS (NM_183275) | Tefm | 5 | tRA treated (DSB peaks) |
| 1-649 | + | 692 | promoter-TSS (NM_175109) | Rps19bp1 | 5 | tRA treated (DSB peaks) |
| 1-718 | + | 784 | intron (NM_001040397, intron 1 of 4) | Filip1l | 4 | tRA treated (DSB peaks) |
| 1-1446 | + | 645 | promoter-TSS (NM_019685) | Ruvbl1 | 5 | tRA treated (DSB peaks) |
| 1-1214 | + | 799 | promoter-TSS (NM_007670) | Cdkn2b | 5 | tRA treated (DSB peaks) |

| | | | | | | |
|---|---|---|---|---|---|---|
| 1-615 | + | 619 | Intergenic | Zhx2 | 5 | tRA treated (DSB peaks) |
| 1-269 | + | 752 | intron (NM_134189, intron 1 of 11) | 4933424L21Rik | 11 | tRA treated (DSB peaks) |
| 1-270 | + | 752 | intron (NM_134189, intron 1 of 11) | 4933424L21Rik | 4 | tRA treated (DSB peaks) |
| 1-1318 | + | 610 | intron (NM_001310608, intron 1 of 3) | Pcdh7 | 6 | tRA treated (DSB peaks) |
| 1-729 | + | 670 | promoter-TSS (NM_026110) | Paxbp1 | 5 | tRA treated (DSB peaks) |
| 1-1725 | + | 566 | intron (NM_019689, intron 1 of 7) | Arid3b | 6 | tRA treated (DSB peaks) |
| 1-616 | + | 644 | Intergenic | Fam83a | 6 | tRA treated (DSB peaks) |
| 20090 | + | 551 | promoter-TSS (NM_001159718) | Hdlbp | 5 | tRA treated (DSB peaks) |
| 1-190 | + | 556 | promoter-TSS (NM_011629) | Nr2c1 | 5 | tRA treated (DSB peaks) |
| 1-955 | + | 618 | Intergenic | Cytip | 3 | tRA treated (DSB peaks) |
| 1-978 | + | 721 | promoter-TSS (NM_001349003) | Api5 | 5 | tRA treated (DSB peaks) |
| 1-1726 | + | 575 | Intergenic | Mir6385 | 7 | tRA treated (DSB peaks) |
| 1-271 | + | 578 | Intergenic | Zfp692 | 9 | tRA treated (DSB peaks) |
| 1-1084 | + | 634 | Intergenic | Tsc22d2 | 5 | tRA treated (DSB peaks) |
| 1-1103 | + | 727 | promoter-TSS (NM_009336) | Vps72 | 5 | tRA treated (DSB peaks) |
| 1-1085 | + | 601 | exon (NM_001081229, exon 1 of 4) | Tsc22d2 | 2 | tRA treated (DSB peaks) |
| 1-1104 | + | 598 | promoter-TSS (NM_172512) | Gabpb2 | 5 | tRA treated (DSB peaks) |
| 1-1759 | + | 639 | promoter-TSS (NM_001114977) | U2surp | 5 | tRA treated (DSB peaks) |
| 1-617 | + | 751 | Intergenic | Fam91a1 | 5 | tRA treated (DSB peaks) |
| 1-1086 | + | 546 | intron (NM_001040396, intron 1 of 5) | Selenot | 2 | tRA treated (DSB peaks) |
| 1-1653 | + | 567 | promoter-TSS (NM_030198) | Gins3 | 5 | tRA treated (DSB peaks) |
| 1-1106 | + | 649 | promoter-TSS (NM_008562) | Mcl1 | 5 | tRA treated (DSB peaks) |
| 1-509 | + | 583 | promoter-TSS (NM_008255) | Hmgcr | 5 | tRA treated (DSB peaks) |
| 1-1424 | + | 595 | 5' UTR (NM_001081145, exon 1 of 2) | Tigd2 | 1 | tRA treated (DSB peaks) |
| 1-338 | + | 616 | promoter-TSS (NM_008379) | Kpnb1 | 5 | tRA treated (DSB peaks) |
| 1-1197 | + | 766 | intron (NM_001163288, intron 15 of 16) | Gm12596 | 4 | tRA treated (DSB peaks) |
| 1-653 | + | 573 | promoter-TSS (NM_001163487) | Pfkm | 5 | tRA treated (DSB peaks) |
| 1-273 | + | 763 | TTS (NR_037992) | Jmjd4 | 8 | tRA treated (DSB peaks) |
| 1-274 | + | 762 | TTS (NR_037992) | Jmjd4 | 6 | tRA treated (DSB peaks) |
| 1-1550 | + | 741 | promoter-TSS (NM_009825) | Serpinh1 | 5 | tRA treated (DSB peaks) |
| 1-659 | + | 725 | promoter-TSS (NM_031842) | Smarcd1 | 5 | tRA treated (DSB peaks) |
| 1-622 | + | 587 | Intergenic | Trib1 | 5 | tRA treated (DSB peaks) |
| 1-1762 | + | 719 | promoter-TSS (NM_010878) | Nck1 | 5 | tRA treated (DSB peaks) |
| 1-623 | + | 593 | Intergenic | Gm19510 | 8 | tRA treated (DSB peaks) |
| 1-624 | + | 593 | Intergenic | Gm19510 | 7 | tRA treated (DSB peaks) |
| 1-342 | + | 579 | promoter-TSS (NM_024456) | Rab5c | 5 | tRA treated (DSB peaks) |
| 1-484 | + | 764 | Intergenic | A530065N20Rik | 12 | tRA treated (DSB peaks) |
| 1-485 | + | 548 | Intergenic | A530065N20Rik | 8 | tRA treated (DSB peaks) |
| 1-345 | + | 1000 | promoter-TSS (NM_029649) | Tmem101 | 5 | tRA treated (DSB peaks) |
| 1-349 | + | 692 | promoter-TSS (NM_001159492) | Gpatch8 | 5 | tRA treated (DSB peaks) |
| 1-1763 | + | 634 | promoter-TSS (NM_001301689) | Anapc13 | 5 | tRA treated (DSB peaks) |
| 1-584 | + | 548 | promoter-TSS (NR_039592) | Kctd12 | 5 | tRA treated (DSB peaks) |
| 1-487 | + | 1000 | Intergenic | Dapk1 | 5 | tRA treated (DSB peaks) |
| 1-356 | + | 632 | promoter-TSS (NM_001130187) | Smarcd2 | 5 | tRA treated (DSB peaks) |
| 1-488 | + | 616 | Intergenic | Dapk1 | 6 | tRA treated (DSB peaks) |
| 1-1221 | + | 635 | promoter-TSS (NM_028209) | Ttc4 | 5 | tRA treated (DSB peaks) |
| 1-844 | + | 754 | Intergenic | G630071F17Rik | 7 | tRA treated (DSB peaks) |

| | | | | | | |
|---|---|---|---|---|---|---|
| 1-956 | + | 842 | intron (NM_001159564, intron 1 of 17) | Itgb6 | 7 | tRA treated (DSB peaks) |
| 1-625 | + | 553 | 5' UTR (NM_001162926, exon 1 of 2) | Fam84b | 4 | tRA treated (DSB peaks) |
| 44927 | + | 547 | Intergenic | Cd28 | 6 | tRA treated (DSB peaks) |
| 1-153 | + | 661 | intron (NM_001316674, intron 1 of 14) | Sgpl1 | 1 | tRA treated (DSB peaks) |
| 1-278 | + | 851 | Intergenic | Slc47a1 | 3 | tRA treated (DSB peaks) |
| 1-562 | + | 601 | intron (NR_028264, intron 1 of 6) | Dleu2 | 2 | tRA treated (DSB peaks) |
| 1-154 | + | 681 | intron (NM_207000, intron 1 of 8) | H2afy2 | 1 | tRA treated (DSB peaks) |
| 1-1556 | + | 840 | promoter-TSS (NM_025344) | Eif3f | 5 | tRA treated (DSB peaks) |
| 1-1774 | + | 662 | promoter-TSS (NM_001012236) | Trex1 | 5 | tRA treated (DSB peaks) |
| 1-1775 | + | 672 | promoter-TSS (NM_018757) | Nme6 | 5 | tRA treated (DSB peaks) |
| 1-1575 | + | 557 | promoter-TSS (NM_011965) | Psma1 | 5 | tRA treated (DSB peaks) |
| 1-1227 | + | 782 | promoter-TSS (NM_028142) | Nsun4 | 5 | tRA treated (DSB peaks) |
| 1-365 | + | 726 | promoter-TSS (NM_001033528) | Usp36 | 5 | tRA treated (DSB peaks) |
| 23743 | + | 563 | promoter-TSS (NM_001081125) | Gli2 | 5 | tRA treated (DSB peaks) |
| 1-1665 | + | 625 | promoter-TSS (NM_028883) | Tldc1 | 5 | tRA treated (DSB peaks) |
| 1-1730 | + | 656 | Intergenic | Coro2b | 16 | tRA treated (DSB peaks) |
| 1-1237 | + | 804 | promoter-TSS (NM_172700) | Zmpste24 | 5 | tRA treated (DSB peaks) |
| 24473 | + | 623 | promoter-TSS (NM_025860) | Ddx18 | 5 | tRA treated (DSB peaks) |
| 1-1667 | + | 558 | promoter-TSS (NM_145605) | Klhdc4 | 5 | tRA treated (DSB peaks) |
| 45292 | + | 567 | 5' UTR (NM_010939, exon 1 of 16) | Nrp2 | 1 | tRA treated (DSB peaks) |
| 1-284 | + | 567 | Intergenic | Zfp287 | 4 | tRA treated (DSB peaks) |
| 1-590 | + | 597 | promoter-TSS (NM_144844) | Pcca | 5 | tRA treated (DSB peaks) |
| 1-285 | + | 926 | Intergenic | Cdrt4 | 6 | tRA treated (DSB peaks) |
| 1-1242 | + | 638 | promoter-TSS (NM_017475) | Rragc | 5 | tRA treated (DSB peaks) |
| 1-1354 | + | 562 | promoter-TSS (NM_181410) | Eif2b1 | 5 | tRA treated (DSB peaks) |
| 1-489 | + | 774 | intron (NM_001289926, intron 3 of 15) | 2010111I01Rik | 1 | tRA treated (DSB peaks) |
| 1-1732 | + | 664 | Intergenic | Skor1 | 3 | tRA treated (DSB peaks) |
| 1-1733 | + | 559 | Intergenic | Skor1 | 8 | tRA treated (DSB peaks) |
| 1-563 | + | 720 | intron (NM_007798, intron 1 of 9) | Ctsb | 7 | tRA treated (DSB peaks) |
| 1-564 | + | 720 | intron (NM_007798, intron 1 of 9) | Ctsb | 7 | tRA treated (DSB peaks) |
| 1-1734 | + | 572 | Intergenic | Skor1 | 4 | tRA treated (DSB peaks) |
| 1-1477 | + | 647 | promoter-TSS (NM_013535) | Grcc10 | 5 | tRA treated (DSB peaks) |
| 1-1011 | + | 753 | promoter-TSS (NM_025922) | Itpa | 5 | tRA treated (DSB peaks) |
| 46388 | + | 561 | TTS (NM_018796) | Snora41 | 5 | tRA treated (DSB peaks) |
| 1-490 | + | 607 | intron (NM_001289926, intron 13 of 15) | 2010111I01Rik | 12 | tRA treated (DSB peaks) |
| 1-626 | + | 885 | Intergenic | Gm20740 | 4 | tRA treated (DSB peaks) |
| 1-627 | + | 795 | Intergenic | Gm20740 | 2 | tRA treated (DSB peaks) |
| 1-1259 | + | 616 | promoter-TSS (NM_013885) | Clic4 | 5 | tRA treated (DSB peaks) |
| 1-491 | + | 686 | intron (NM_001328514, intron 1 of 23) | Ptch1 | 1 | tRA treated (DSB peaks) |
| 1-492 | + | 686 | intron (NM_001328514, intron 1 of 23) | Ptch1 | 4 | tRA treated (DSB peaks) |
| 1-785 | + | 677 | Intergenic | Fbxl17 | 2 | tRA treated (DSB peaks) |
| 1-786 | + | 621 | Intergenic | Fer | 10 | tRA treated (DSB peaks) |
| 1-1735 | + | 548 | intron (NM_016769, intron 1 of 8) | Smad3 | 5 | tRA treated (DSB peaks) |
| 1-1380 | + | 777 | promoter-TSS (NM_172412) | Stag3 | 5 | tRA treated (DSB peaks) |
| 1-720 | + | 602 | intron (NM_010140, intron 1 of 16) | Epha3 | 3 | tRA treated (DSB peaks) |
| 1-1736 | + | 618 | Intergenic | Smad6 | 5 | tRA treated (DSB peaks) |
| 1-1737 | + | 618 | Intergenic | Smad6 | 5 | tRA treated (DSB peaks) |

| | | | | | | |
|---|---|---|---|---|---|---|
| 46753 | + | 663 | Intergenic | Klf7 | 13 | tRA treated (DSB peaks) |
| 47119 | + | 699 | Intergenic | Klf7 | 7 | tRA treated (DSB peaks) |
| 1-1383 | + | 542 | promoter-TSS (NM_026748) | Ints1 | 5 | tRA treated (DSB peaks) |
| 1-160 | + | 594 | intron (NM_001164376, intron 9 of 17) | Lrrtm3 | 5 | tRA treated (DSB peaks) |
| 1-845 | + | 562 | Intergenic | Atp8b1 | 11 | tRA treated (DSB peaks) |
| 1-1150 | + | 552 | promoter-TSS (NM_009740) | Bcl10 | 5 | tRA treated (DSB peaks) |
| 1-1202 | + | 568 | Intergenic | Pappa | 7 | tRA treated (DSB peaks) |
| 1-1492 | + | 540 | promoter-TSS (NM_010656) | Sspn | 5 | tRA treated (DSB peaks) |
| 1-1738 | + | 658 | Intergenic | Parp16 | 8 | tRA treated (DSB peaks) |
| 1-1739 | + | 658 | Intergenic | Parp16 | 5 | tRA treated (DSB peaks) |
| 1-494 | + | 549 | Intergenic | Nlrp4f | 6 | tRA treated (DSB peaks) |
| 1-1203 | + | 540 | intron (NM_207109, intron 20 of 22) | Trim32 | 4 | tRA treated (DSB peaks) |
| 1-1391 | + | 900 | promoter-TSS (NM_011908) | Ubl3 | 5 | tRA treated (DSB peaks) |
| 1-1425 | + | 821 | exon (NM_027547, exon 1 of 15) | Prdm5 | 1 | tRA treated (DSB peaks) |
| 1-1392 | + | 899 | promoter-TSS (NM_011908) | Ubl3 | 5 | tRA treated (DSB peaks) |
| 1-1057 | + | 565 | promoter-TSS (NM_030708) | Zfhx4 | 4 | tRA treated (DSB peaks) |
| 1-565 | + | 544 | Intergenic | Scara3 | 4 | tRA treated (DSB peaks) |
| 1-846 | + | 909 | exon (NM_013833, exon 3 of 3) | Rax | 1 | tRA treated (DSB peaks) |
| 1-872 | + | 744 | promoter-TSS (NM_026912) | Snx15 | 4 | tRA treated (DSB peaks) |
| 1-818 | + | 679 | promoter-TSS (NM_144860) | Mib1 | 4 | tRA treated (DSB peaks) |
| 1-847 | + | 620 | intron (NM_178793, intron 2 of 10) | Mir694 | 3 | tRA treated (DSB peaks) |
| 1-848 | + | 573 | intron (NM_178793, intron 2 of 10) | Mir694 | 3 | tRA treated (DSB peaks) |
| 1-1319 | + | 587 | Intergenic | 1700126H18Rik | 5 | tRA treated (DSB peaks) |
| 1-1088 | + | 569 | Intergenic | Veph1 | 6 | tRA treated (DSB peaks) |
| 1-1496 | + | 786 | promoter-TSS (NM_001024699) | Zbtb45 | 4 | tRA treated (DSB peaks) |
| 1-739 | + | 557 | promoter-TSS (NM_013684) | Tbp | 4 | tRA treated (DSB peaks) |
| 1-1743 | + | 679 | exon (NM_173413, exon 1 of 8) | Rab8b | 4 | tRA treated (DSB peaks) |
| 1-885 | + | 587 | promoter-TSS (NM_001190483) | Pcsk5 | 4 | tRA treated (DSB peaks) |
| 1-1172 | + | 579 | promoter-TSS (NM_025476) | Rmdn1 | 4 | tRA treated (DSB peaks) |
| 1-926 | + | 571 | promoter-TSS (NM_027715) | Gm3230 | 4 | tRA treated (DSB peaks) |
| 1-161 | + | 734 | Intergenic | Gm31763 | 7 | tRA treated (DSB peaks) |
| 1-742 | + | 754 | promoter-TSS (NM_016891) | Ppp2r1a | 4 | tRA treated (DSB peaks) |
| 1-425 | + | 628 | promoter-TSS (NM_001251833) | Zkscan8 | 4 | tRA treated (DSB peaks) |
| 1-1746 | + | 575 | 5' UTR (NM_001164254, exon 1 of 9) | Tpm1 | 3 | tRA treated (DSB peaks) |
| 1-929 | + | 609 | promoter-TSS (NM_019501) | Pdss1 | 4 | tRA treated (DSB peaks) |
| 1-1428 | + | 598 | Intergenic | Gadd45a | 5 | tRA treated (DSB peaks) |
| 1-238 | + | 685 | promoter-TSS (NM_023651) | Pus10 | 4 | tRA treated (DSB peaks) |
| 1-1748 | + | 693 | intron (NM_001081242, intron 54 of 57) | Mir190a | 6 | tRA treated (DSB peaks) |
| 1-1749 | + | 694 | exon (NM_001081242, exon 54 of 58) | Mir190a | 5 | tRA treated (DSB peaks) |
| 1-286 | + | 576 | Intergenic | Gas7 | 7 | tRA treated (DSB peaks) |
| 1-682 | + | 653 | promoter-TSS (NM_009744) | Bcl6 | 4 | tRA treated (DSB peaks) |
| 1-1298 | + | 782 | promoter-TSS (NM_016736) | Nub1 | 4 | tRA treated (DSB peaks) |
| 1-933 | + | 548 | promoter-TSS (NM_146115) | Tor4a | 4 | tRA treated (DSB peaks) |
| 1-162 | + | 750 | intron (NM_023598, intron 4 of 9) | Arid5b | 6 | tRA treated (DSB peaks) |
| 1-163 | + | 716 | Intergenic | 4930545H06Rik | 11 | tRA treated (DSB peaks) |
| 1-1623 | + | 758 | intron (NM_172753, intron 4 of 9) | 1700125H03Rik | 4 | tRA treated (DSB peaks) |
| 1-1624 | + | 759 | intron (NM_172753, intron 4 of 9) | 1700125H03Rik | 7 | tRA treated (DSB peaks) |

| | | | | | | |
|---|---|---|---|---|---|---|
| 1-164 | + | 570 | Intergenic | Cabcoco1 | 9 | tRA treated (DSB peaks) |
| 1-1625 | + | 590 | intron (NM_172753, intron 1 of 9) | Csgalnact1 | 8 | tRA treated (DSB peaks) |
| 1-1626 | + | 590 | intron (NM_172753, intron 1 of 9) | Csgalnact1 | 7 | tRA treated (DSB peaks) |
| 1-957 | + | 810 | intron (NM_001347161, intron 3 of 10) | Cers6 | 7 | tRA treated (DSB peaks) |
| 1-287 | + | 780 | Intergenic | Aurkb | 5 | tRA treated (DSB peaks) |
| 1-288 | + | 582 | Intergenic | Aurkb | 5 | tRA treated (DSB peaks) |
| 1-289 | + | 888 | Intergenic | Borcs6 | 13 | tRA treated (DSB peaks) |
| 1-290 | + | 908 | TTS (NR_028566) | Snord118 | 18 | tRA treated (DSB peaks) |
| 1-291 | + | 906 | TTS (NR_028566) | Snord118 | 13 | tRA treated (DSB peaks) |
| 1-292 | + | 793 | Intergenic | Hes7 | 8 | tRA treated (DSB peaks) |
| 1-293 | + | 859 | Intergenic | Aloxe3 | 4 | tRA treated (DSB peaks) |
| 1-294 | + | 563 | Intergenic | Aloxe3 | 13 | tRA treated (DSB peaks) |
| 1-383 | + | 545 | exon (NM_009093, exon 1 of 3) | Rps29 | 3 | tRA treated (DSB peaks) |
| 1-937 | + | 828 | promoter-TSS (NM_001360790) | Brd3os | 4 | tRA treated (DSB peaks) |
| 1-1701 | + | 544 | promoter-TSS (NM_011808) | Ets1 | 4 | tRA treated (DSB peaks) |
| 1-891 | + | 605 | promoter-TSS (NM_008960) | Pten | 4 | tRA treated (DSB peaks) |
| 1-295 | + | 650 | 3' UTR (NM_010599, exon 14 of 14) | Kcnab3 | 7 | tRA treated (DSB peaks) |
| 1-789 | + | 599 | non-coding (NR_026848, exon 1 of 1) | C030034I22Rik | 3 | tRA treated (DSB peaks) |
| 1-296 | + | 666 | Intergenic | Kdm6b | 9 | tRA treated (DSB peaks) |
| 1-1627 | + | 607 | 5' UTR (NM_001359086, exon 1 of 6) | Zfp869 | 2 | tRA treated (DSB peaks) |
| 1-376 | + | 552 | promoter-TSS (NM_021524) | Nampt | 4 | tRA treated (DSB peaks) |
| 1-823 | + | 543 | promoter-TSS (NM_144863) | Wdr36 | 4 | tRA treated (DSB peaks) |
| 1-697 | + | 720 | promoter-TSS (NM_011749) | 1700007L15Rik | 4 | tRA treated (DSB peaks) |
| 43191 | + | 545 | promoter-TSS (NM_001359275) | Zfp451 | 4 | tRA treated (DSB peaks) |
| 1-762 | + | 712 | promoter-TSS (NM_011296) | Vps52 | 4 | tRA treated (DSB peaks) |
| 1-1522 | + | 1000 | intron (NR_131145, intron 1 of 3) | Gm29683 | 10 | tRA treated (DSB peaks) |
| 1-722 | + | 947 | intron (NM_028803, intron 1 of 15) | Gbe1 | 1 | tRA treated (DSB peaks) |
| 1-1523 | + | 641 | Intergenic | Gm29683 | 5 | tRA treated (DSB peaks) |
| 1-1524 | + | 551 | exon (NM_009697, exon 1 of 3) | Nr2f2 | 5 | tRA treated (DSB peaks) |
| 1-1525 | + | 551 | exon (NM_009697, exon 1 of 3) | Nr2f2 | 3 | tRA treated (DSB peaks) |
| 1-604 | + | 613 | promoter-TSS (NR_028101) | Rpl30 | 4 | tRA treated (DSB peaks) |
| 1-1181 | + | 661 | promoter-TSS (NM_015824) | Rars2 | 4 | tRA treated (DSB peaks) |
| 1-605 | + | 703 | promoter-TSS (NM_177151) | Vps13b | 4 | tRA treated (DSB peaks) |
| 1-1527 | + | 680 | Intergenic | Nr2f2 | 4 | tRA treated (DSB peaks) |
| 1-948 | + | 544 | promoter-TSS (NM_001145821) | Ggta1 | 4 | tRA treated (DSB peaks) |
| 1-569 | + | 541 | Intergenic | Npm2 | 4 | tRA treated (DSB peaks) |
| 1-828 | + | 710 | promoter-TSS (NM_001357512) | Apbb3 | 4 | tRA treated (DSB peaks) |
| 1-790 | + | 550 | intron (NM_001360665, intron 9 of 14) | Dlgap1 | 4 | tRA treated (DSB peaks) |
| 1-1630 | + | 592 | 5' UTR (NM_010592, exon 1 of 1) | Jund | 5 | tRA treated (DSB peaks) |
| 1-791 | + | 881 | intron (NM_001347411, intron 9 of 14) | Tgif1 | 14 | tRA treated (DSB peaks) |
| 1-792 | + | 655 | intron (NM_001347411, intron 9 of 14) | Tgif1 | 16 | tRA treated (DSB peaks) |
| 1-387 | + | 588 | Intergenic | Frmd6 | 3 | tRA treated (DSB peaks) |
| 1-298 | + | 580 | exon (NM_007573, exon 1 of 6) | C1qbp | 1 | tRA treated (DSB peaks) |
| 1-896 | + | 618 | promoter-TSS (NM_016854) | Ppp1r3c | 4 | tRA treated (DSB peaks) |
| 1-899 | + | 754 | promoter-TSS (NM_021315) | Noc3l | 4 | tRA treated (DSB peaks) |
| 1-1412 | + | 794 | promoter-TSS (NM_001286664) | Ssbp1 | 4 | tRA treated (DSB peaks) |
| 1-960 | + | 1000 | intron (NR_039557, intron 2 of 3).2 | Platr26 | 4 | tRA treated (DSB peaks) |

| | | | | | | |
|---|---|---|---|---|---|---|
| 1-496 | + | 559 | exon (NM_010573, exon 2 of 4) | Irx1 | 1 | tRA treated (DSB peaks) |
| 1-1413 | + | 969 | promoter-TSS (NM_011777) | Zyx | 4 | tRA treated (DSB peaks) |
| 1-902 | + | 626 | promoter-TSS (NM_009771) | Btrc | 4 | tRA treated (DSB peaks) |
| 1-251 | + | 702 | promoter-TSS (NM_001290737) | Thg1l | 4 | tRA treated (DSB peaks) |
| 1-1430 | + | 593 | exon (NM_138592, exon 1 of 13) | Usp39 | 6 | tRA treated (DSB peaks) |
| 1-544 | + | 575 | promoter-TSS (NM_001360455) | Cgrrf1 | 4 | tRA treated (DSB peaks) |
| 1-497 | + | 653 | Intergenic | Irx2 | 6 | tRA treated (DSB peaks) |
| 1-467 | + | 562 | promoter-TSS (NM_010617) | Kif13a | 4 | tRA treated (DSB peaks) |
| 1-724 | + | 542 | 5' UTR (NM_019413, exon 1 of 29) | Robo1 | 2 | tRA treated (DSB peaks) |
| 1-547 | + | 554 | promoter-TSS (NM_144535) | Ap5m1 | 4 | tRA treated (DSB peaks) |
| 1-300 | + | 624 | intron (NM_009671, intron 1 of 24) | Ankfy1 | 3 | tRA treated (DSB peaks) |
| 1-1206 | + | 546 | Intergenic | Aldoart1 | 4 | tRA treated (DSB peaks) |
| 1-1715 | + | 543 | promoter-TSS (NM_001113204) | Ncam1 | 4 | tRA treated (DSB peaks) |
| 1-1432 | + | 555 | Intergenic | Tmsb10 | 6 | tRA treated (DSB peaks) |
| 1-570 | + | 659 | Intergenic | Gm9199 | 3 | tRA treated (DSB peaks) |
| 1-962 | + | 613 | Intergenic | Sp3 | 6 | tRA treated (DSB peaks) |
| 1-262 | + | 564 | promoter-TSS (NM_175334) | Maml1 | 4 | tRA treated (DSB peaks) |
| 1-571 | + | 715 | intron (NR_033185, intron 1 of 15) | Rcbtb2 | 1 | tRA treated (DSB peaks) |
| 1-572 | + | 587 | intron (NM_009029, intron 4 of 26) | Rb1 | 19 | tRA treated (DSB peaks) |
| 1-264 | + | 614 | promoter-TSS (NM_025319) | 0610009B22Rik | 4 | tRA treated (DSB peaks) |
| 1-573 | + | 712 | intron (NM_172527, intron 3 of 4) | Nudt15 | 2 | tRA treated (DSB peaks) |
| 1-1528 | + | 834 | intron (NM_001081345, intron 2 of 38) | Chd2 | 8 | tRA treated (DSB peaks) |
| 1-1529 | + | 558 | Intergenic | 1810026B05Rik | 1 | tRA treated (DSB peaks) |
| 1-1317 | + | 774 | promoter-TSS (NM_144517) | Tbc1d19 | 4 | tRA treated (DSB peaks) |
| 1-1751 | + | 589 | intron (NM_001081153, intron 1 of 30) | Unc13c | 10 | tRA treated (DSB peaks) |
| 1-1752 | + | 589 | intron (NM_001081153, intron 1 of 30) | Unc13c | 8 | tRA treated (DSB peaks) |
| 1-793 | + | 771 | Intergenic | Xdh | 5 | tRA treated (DSB peaks) |
| 1-794 | + | 769 | Intergenic | Xdh | 4 | tRA treated (DSB peaks) |
| 1-1530 | + | 582 | Intergenic | St8sia2 | 5 | tRA treated (DSB peaks) |
| 1-574 | + | 637 | Intergenic | 4933402J15Rik | 5 | tRA treated (DSB peaks) |
| 11324 | + | 981 | TTS (NM_013612) | Ctdsp1 | 9 | tRA treated (DSB peaks) |
| 1-912 | + | 571 | promoter-TSS (NM_001355242) | Shoc2 | 4 | tRA treated (DSB peaks) |
| 1-479 | + | 716 | promoter-TSS (NM_001311137) | B4galt7 | 4 | tRA treated (DSB peaks) |
| 1-1724 | + | 552 | promoter-TSS (NM_146223) | Cplx3 | 4 | tRA treated (DSB peaks) |
| 1-1727 | + | 615 | promoter-TSS (NM_028121) | Adpgk | 4 | tRA treated (DSB peaks) |
| 1-964 | + | 556 | Intergenic | Hoxd12 | 1 | tRA treated (DSB peaks) |
| 1-965 | + | 556 | Intergenic | Hoxd12 | 1 | tRA treated (DSB peaks) |
| 1-1320 | + | 624 | Intergenic | Lnx1 | 4 | tRA treated (DSB peaks) |
| 1-1321 | + | 817 | Intergenic | Gm6116 | 8 | tRA treated (DSB peaks) |
| 1-1322 | + | 634 | Intergenic | Gm6116 | 5 | tRA treated (DSB peaks) |
| 1-1728 | + | 654 | promoter-TSS (NM_019927) | Arih1 | 4 | tRA treated (DSB peaks) |
| 1-1636 | + | 694 | intron (NM_178017, intron 1 of 10) | Hmgxb4 | 2 | tRA treated (DSB peaks) |
| 1-302 | + | 630 | intron (NM_001002764, intron 10 of 18) | Mir684-1 | 5 | tRA treated (DSB peaks) |
| 1-303 | + | 630 | intron (NM_001002764, intron 10 of 18) | Mir684-1 | 4 | tRA treated (DSB peaks) |
| 1-1637 | + | 761 | Intergenic | Hmox1 | 11 | tRA treated (DSB peaks) |
| 1-796 | + | 565 | intron (NM_019919, intron 3 of 33) | Ltbp1 | 8 | tRA treated (DSB peaks) |
| 1-797 | + | 566 | intron (NM_019919, intron 3 of 33) | Ltbp1 | 6 | tRA treated (DSB peaks) |

| | | | | | | |
|---|---|---|---|---|---|---|
| 1-275 | + | 583 | promoter-TSS (NM_001290012) | Pemt | 4 | tRA treated (DSB peaks) |
| 1-276 | + | 596 | promoter-TSS (NM_145427) | Gid4 | 4 | tRA treated (DSB peaks) |
| 1-1200 | + | 746 | promoter-TSS (NM_001276446) | Alad | 4 | tRA treated (DSB peaks) |
| 1-281 | + | 667 | promoter-TSS (NR_002875) | Gm1821 | 4 | tRA treated (DSB peaks) |
| 1-155 | + | 621 | promoter-TSS (NM_053183) | Ddx50 | 4 | tRA treated (DSB peaks) |
| 1-158 | + | 841 | promoter-TSS (NM_001079824) | Pbld2 | 4 | tRA treated (DSB peaks) |
| 1-1740 | + | 629 | promoter-TSS (NM_172536) | Zfp609 | 4 | tRA treated (DSB peaks) |
| 1-966 | + | 707 | intron (NM_010902, intron 1 of 4) | Nfe2l2 | 9 | tRA treated (DSB peaks) |
| 1-967 | + | 707 | intron (NM_010902, intron 1 of 4) | Nfe2l2 | 5 | tRA treated (DSB peaks) |
| 1-1742 | + | 876 | promoter-TSS (NM_019727) | Snx1 | 4 | tRA treated (DSB peaks) |
| 1-1090 | + | 604 | Intergenic | Golim4 | 5 | tRA treated (DSB peaks) |
| 1-1745 | + | 721 | promoter-TSS (NM_001361106) | Rps27l | 4 | tRA treated (DSB peaks) |
| 1-566 | + | 713 | promoter-TSS (NM_009761) | Bnip3l | 4 | tRA treated (DSB peaks) |
| 1-1427 | + | 569 | promoter-TSS (NM_007836) | Gadd45a | 4 | tRA treated (DSB peaks) |
| 1-1204 | + | 621 | promoter-TSS (NM_145990) | Cdk5rap2 | 4 | tRA treated (DSB peaks) |
| 1-1323 | + | 800 | intron (NM_020611, intron 1 of 5) | Srd5a3 | 2 | tRA treated (DSB peaks) |
| 1-308 | + | 581 | intron (NM_008750, intron 1 of 7) | Mrm3 | 7 | tRA treated (DSB peaks) |
| 1-1205 | + | 622 | promoter-TSS (NM_145990) | Cdk5rap2 | 4 | tRA treated (DSB peaks) |
| 1-1207 | + | 624 | intron (NM_011211, intron 10 of 39) | Ptprd | 1 | tRA treated (DSB peaks) |
| 1-576 | + | 633 | 5' UTR (NM_009366, exon 1 of 3) | Tsc22d1 | 8 | tRA treated (DSB peaks) |
| 1-959 | + | 775 | promoter-TSS (NM_027352) | Gorasp2 | 4 | tRA treated (DSB peaks) |
| 1-1632 | + | 761 | promoter-TSS (NM_023126) | Rab8a | 4 | tRA treated (DSB peaks) |
| 1-577 | + | 779 | intron (NR_015485, intron 4 of 4) | 1700108F19Rik | 3 | tRA treated (DSB peaks) |
| 1-309 | + | 613 | intron (NM_001356321, intron 1 of 8) | Gosr1 | 7 | tRA treated (DSB peaks) |
| 1-630 | + | 777 | promoter-TSS (NM_030199) | Zfp623 | 4 | tRA treated (DSB peaks) |
| 1-166 | + | 548 | intron (NM_001347216, intron 1 of 15) | Pcbp3 | 8 | tRA treated (DSB peaks) |
| 1-633 | + | 577 | promoter-TSS (NM_012053) | Rpl8 | 4 | tRA treated (DSB peaks) |
| 1-634 | + | 867 | exon (NM_001347540, exon 1 of 5) | 1110038F14Rik | 8 | tRA treated (DSB peaks) |
| 1-635 | + | 695 | exon (NM_001347540, exon 2 of 5) | 1110038F14Rik | 6 | tRA treated (DSB peaks) |
| 1-636 | + | 578 | Intergenic | 1700109K24Rik | 9 | tRA treated (DSB peaks) |
| 1-310 | + | 601 | promoter-TSS (NM_001024205) | Nufip2 | 4 | tRA treated (DSB peaks) |
| 1-638 | + | 548 | Intergenic | Rbfox2 | 7 | tRA treated (DSB peaks) |
| 1-639 | + | 553 | Intergenic | Apol7a | 4 | tRA treated (DSB peaks) |
| 1-640 | + | 655 | Intergenic | Apol7a | 2 | tRA treated (DSB peaks) |
| 1-1639 | + | 1000 | promoter-TSS (NM_178267) | Zfp827 | 4 | tRA treated (DSB peaks) |
| 1-1638 | + | 663 | intron (NM_010332, intron 1 of 7) | Ednra | 4 | tRA treated (DSB peaks) |
| 1-641 | + | 659 | intron (NM_022410, intron 3 of 40) | Apol8 | 7 | tRA treated (DSB peaks) |
| 1-1753 | + | 670 | Intergenic | Gclc | 8 | tRA treated (DSB peaks) |
| 1-642 | + | 604 | Intergenic | Myh9 | 1 | tRA treated (DSB peaks) |
| 1-174 | + | 644 | promoter-TSS (NM_025349) | Sppl2b | 4 | tRA treated (DSB peaks) |
| 1-799 | + | 548 | intron (NM_015800, intron 7 of 16) | Fez2 | 9 | tRA treated (DSB peaks) |
| 12055 | + | 556 | intron (NM_001004173, intron 1 of 4) | Sgpp2 | 5 | tRA treated (DSB peaks) |
| 1-643 | + | 572 | intron (NM_001355601, intron 10 of 17) | Tmprss6 | 4 | tRA treated (DSB peaks) |
| 1-852 | + | 648 | intron (NM_207651, intron 2 of 21) | Slc14a2 | 2 | tRA treated (DSB peaks) |
| 1-853 | + | 634 | intron (NM_207651, intron 2 of 21) | Slc14a2 | 2 | tRA treated (DSB peaks) |
| 1-578 | + | 602 | exon (NM_173758, exon 1 of 26) | Vwa8 | 2 | tRA treated (DSB peaks) |
| 1-579 | + | 654 | intron (NM_027906, intron 2 of 44) | Vwa8 | 7 | tRA treated (DSB peaks) |

| | | | | | | |
|---|---|---|---|---|---|---|
| 1-181 | + | 639 | promoter-TSS (NM_010914) | Nfyb | 4 | tRA treated (DSB peaks) |
| 1-1434 | + | 755 | promoter-TSS (NM_007517) | Htra2 | 4 | tRA treated (DSB peaks) |
| 1-1435 | + | 566 | promoter-TSS (NM_145571) | Mob1a | 4 | tRA treated (DSB peaks) |
| 1-1438 | + | 704 | promoter-TSS (NM_011585) | Tia1 | 4 | tRA treated (DSB peaks) |
| 1-646 | + | 552 | 5' UTR (NM_008197, exon 1 of 1) | H1f0 | 4 | tRA treated (DSB peaks) |
| 1-854 | + | 574 | Intergenic | Setbp1 | 4 | tRA treated (DSB peaks) |
| 1-855 | + | 574 | Intergenic | Setbp1 | 4 | tRA treated (DSB peaks) |
| 1-311 | + | 560 | Intergenic | Wsb1 | 4 | tRA treated (DSB peaks) |
| 1-334 | + | 627 | promoter-TSS (NM_025287) | Spop | 4 | tRA treated (DSB peaks) |
| 1-508 | + | 596 | promoter-TSS (NM_010169) | F2r | 4 | tRA treated (DSB peaks) |
| 1-1327 | + | 933 | promoter-TSS (NM_053158) | Mrpl1 | 4 | tRA treated (DSB peaks) |
| 1-800 | + | 550 | Intergenic | 4930429F11Rik | 1 | tRA treated (DSB peaks) |
| 1-580 | + | 590 | 5' UTR (NM_001024135, exon 1 of 1) | Kbtbd7 | 2 | tRA treated (DSB peaks) |
| 1-341 | + | 578 | promoter-TSS (NM_024456) | Rab5c | 4 | tRA treated (DSB peaks) |
| 1-1766 | + | 569 | promoter-TSS (NM_172460) | Nphp3 | 4 | tRA treated (DSB peaks) |
| 1-856 | + | 838 | Intergenic | Setbp1 | 18 | tRA treated (DSB peaks) |
| 1-1091 | + | 573 | intron (NM_001162999, intron 1 of 16) | Fnip2 | 11 | tRA treated (DSB peaks) |
| 1-1755 | + | 543 | 5' UTR (NM_009945, exon 1 of 4) | Cox7a2 | 12 | tRA treated (DSB peaks) |
| 1-1219 | + | 604 | promoter-TSS (NM_080555) | Plpp3 | 4 | tRA treated (DSB peaks) |
| 1-312 | + | 678 | Intergenic | Mir365-2 | 10 | tRA treated (DSB peaks) |
| 1-1777 | + | 545 | promoter-TSS (NM_008633) | Map4 | 4 | tRA treated (DSB peaks) |
| 1-1568 | + | 559 | promoter-TSS (NR_015573) | 1700012D14Rik | 4 | tRA treated (DSB peaks) |
| 1-1092 | + | 690 | Intergenic | Fam198b | 9 | tRA treated (DSB peaks) |
| 1-1093 | + | 630 | Intergenic | Fam198b | 4 | tRA treated (DSB peaks) |
| 1-969 | + | 554 | intron (NM_001355141, intron 2 of 15) | Pde1a | 9 | tRA treated (DSB peaks) |
| 1-389 | + | 622 | exon (NM_007564, exon 2 of 2) | Zfp36l1 | 1 | tRA treated (DSB peaks) |
| 1-1660 | + | 820 | promoter-TSS (NR_105819) | | 4 | tRA treated (DSB peaks) |
| 1-1341 | + | 550 | promoter-TSS (NM_018783) | Tfip11 | 4 | tRA treated (DSB peaks) |
| 1-390 | + | 958 | intron (NM_134156, intron 1 of 20) | Actn1 | 5 | tRA treated (DSB peaks) |
| 1-391 | + | 957 | intron (NM_134156, intron 1 of 20) | Actn1 | 4 | tRA treated (DSB peaks) |
| 1-1464 | + | 565 | promoter-TSS (NM_133932) | Tada3 | 4 | tRA treated (DSB peaks) |
| 1-987 | + | 579 | promoter-TSS (NM_009702) | Aqr | 4 | tRA treated (DSB peaks) |
| 1-1534 | + | 551 | TTS (NM_133952) | Rccd1 | 12 | tRA treated (DSB peaks) |
| 1-1230 | + | 563 | promoter-TSS (NM_023178) | Dmap1 | 4 | tRA treated (DSB peaks) |
| 1-392 | + | 620 | intron (NM_177267, intron 6 of 8) | Mir1843a | 9 | tRA treated (DSB peaks) |
| 1-993 | + | 901 | promoter-TSS (NM_029617) | Knl1 | 4 | tRA treated (DSB peaks) |
| 1-998 | + | 603 | promoter-TSS (NM_011941) | Mapkbp1 | 4 | tRA treated (DSB peaks) |
| 1-802 | + | 635 | Intergenic | Sos1 | 2 | tRA treated (DSB peaks) |
| 1-393 | + | 661 | Intergenic | 2310002D06Rik | 2 | tRA treated (DSB peaks) |
| 1-502 | + | 645 | Intergenic | Arrdc3 | 12 | tRA treated (DSB peaks) |
| 1-369 | + | 726 | promoter-TSS (NM_144797) | Metrnl | 4 | tRA treated (DSB peaks) |
| 1-1476 | + | 560 | promoter-TSS (NM_013536) | Emg1 | 4 | tRA treated (DSB peaks) |
| 1-171 | + | 739 | Intergenic | Dot1l | 12 | tRA treated (DSB peaks) |
| 1-172 | + | 603 | Intergenic | Dot1l | 7 | tRA treated (DSB peaks) |
| 1-173 | + | 578 | intron (NM_199322, intron 7 of 27) | Dot1l | 4 | tRA treated (DSB peaks) |
| 1-803 | + | 669 | Intergenic | Map4k3 | 3 | tRA treated (DSB peaks) |
| 1-1479 | + | 596 | promoter-TSS (NM_175557) | Zfp384 | 4 | tRA treated (DSB peaks) |

| | | | | | | |
|---|---|---|---|---|---|---|
| 1-1356 | + | 757 | promoter-TSS (NM_019639) | Ubc | 4 | tRA treated (DSB peaks) |
| 1-1581 | + | 592 | promoter-TSS (NM_026330) | Nsmce1 | 4 | tRA treated (DSB peaks) |
| 1-1208 | + | 586 | intron (NM_026821, intron 1 of 1) | Lurap1l | 6 | tRA treated (DSB peaks) |
| 1-1582 | + | 592 | promoter-TSS (NM_026330) | Nsmce1 | 4 | tRA treated (DSB peaks) |
| 1-1005 | + | 621 | promoter-TSS (NM_029455) | Dtwd1 | 4 | tRA treated (DSB peaks) |
| 1-1094 | + | 663 | intron (NM_019971, intron 1 of 5) | Pdgfc | 5 | tRA treated (DSB peaks) |
| 1-1247 | + | 710 | promoter-TSS (NM_133883) | Adprhl2 | 4 | tRA treated (DSB peaks) |
| 1-1673 | + | 606 | promoter-TSS (NM_001164598) | Irf2bp2 | 4 | tRA treated (DSB peaks) |
| 1-178 | + | 575 | exon (NM_008844, exon 1 of 18) | Pip5k1c | 6 | tRA treated (DSB peaks) |
| 1-1324 | + | 593 | intron (NM_198702, intron 5 of 27) | Adgrl3 | 7 | tRA treated (DSB peaks) |
| 1-1364 | + | 598 | promoter-TSS (NM_008095) | Nipsnap2 | 4 | tRA treated (DSB peaks) |
| 1-1591 | + | 809 | promoter-TSS (NM_026655) | 2310057M21Rik | 4 | tRA treated (DSB peaks) |
| 1-804 | + | 623 | intron (NR_104580, intron 2 of 8) | Slc8a1 | 10 | tRA treated (DSB peaks) |
| 1-1253 | + | 920 | promoter-TSS (NR_110978) | Tmem222 | 4 | tRA treated (DSB peaks) |
| 1-179 | + | 608 | Intergenic | AU041133 | 1 | tRA treated (DSB peaks) |
| 1-394 | + | 549 | intron (NM_001167983, intron 2 of 23) | Sipa1l1 | 8 | tRA treated (DSB peaks) |
| 12785 | + | 549 | intron (NM_010570, intron 1 of 1) | Irs1 | 1 | tRA treated (DSB peaks) |
| 13150 | + | 1000 | 5' UTR (NM_010570, exon 1 of 2) | Irs1 | 2 | tRA treated (DSB peaks) |
| 1-857 | + | 562 | Intergenic | Mbp | 2 | tRA treated (DSB peaks) |
| 1-858 | + | 584 | intron (NM_001025245, intron 1 of 3) | Mbp | 2 | tRA treated (DSB peaks) |
| 1-1369 | + | 628 | promoter-TSS (NM_010717) | Limk1 | 4 | tRA treated (DSB peaks) |
| 1-1262 | + | 552 | promoter-TSS (NM_008321) | Id3 | 4 | tRA treated (DSB peaks) |
| 1-1375 | + | 728 | promoter-TSS (NM_010312) | Gnb2 | 4 | tRA treated (DSB peaks) |
| 1-1642 | + | 675 | intron (NM_177378, intron 1 of 6) | Rnf150 | 7 | tRA treated (DSB peaks) |
| 1-1643 | + | 671 | intron (NM_177378, intron 1 of 6) | Rnf150 | 6 | tRA treated (DSB peaks) |
| 1-1433 | + | 622 | intron (NM_033079, intron 4 of 9) | Dok1 | 11 | tRA treated (DSB peaks) |
| 1-1382 | + | 620 | promoter-TSS (NM_197980) | Cox19 | 4 | tRA treated (DSB peaks) |
| 27760 | + | 581 | promoter-TSS (NM_009888) | Cfh | 4 | tRA treated (DSB peaks) |
| 1-1388 | + | 644 | promoter-TSS (NM_001033313) | Bud31 | 4 | tRA treated (DSB peaks) |
| 1-1393 | + | 916 | promoter-TSS (NM_001286825) | Uspl1 | 4 | tRA treated (DSB peaks) |
| 33239 | + | 700 | promoter-TSS (NM_201364) | Mettl18 | 4 | tRA treated (DSB peaks) |
| 1-1042 | + | 596 | promoter-TSS (NM_133924) | Snx21 | 4 | tRA treated (DSB peaks) |
| 1-182 | + | 696 | Intergenic | 1500009L16Rik | 2 | tRA treated (DSB peaks) |
| 1-183 | + | 696 | Intergenic | 1500009L16Rik | 3 | tRA treated (DSB peaks) |
| 1-109 | + | 942 | promoter-TSS (NM_145943) | Sde2 | 4 | tRA treated (DSB peaks) |
| 1-805 | + | 576 | Intergenic | C430042M11Rik | 6 | tRA treated (DSB peaks) |
| 1-813 | + | 748 | promoter-TSS (NM_001110852) | Crem | 3 | tRA treated (DSB peaks) |
| 1-1539 | + | 598 | exon (NM_133722, exon 1 of 3) | Abhd17c | 2 | tRA treated (DSB peaks) |
| 1-1437 | + | 619 | intron (NM_001077694, intron 41 of 54) | Dysf | 2 | tRA treated (DSB peaks) |
| 1-806 | + | 1000 | exon (NM_001001806, exon 2 of 2) | Zfp36l2 | 1 | tRA treated (DSB peaks) |
| 13881 | + | 563 | TTS (NR_105771) | Mir6353 | 5 | tRA treated (DSB peaks) |
| 1-1540 | + | 551 | intron (NM_007488, intron 5 of 18) | Arnt2 | 7 | tRA treated (DSB peaks) |
| 1-1757 | + | 542 | Intergenic | 4930554C24Rik | 11 | tRA treated (DSB peaks) |
| 1-184 | + | 672 | intron (NM_001004363, intron 4 of 6) | Nuak1 | 6 | tRA treated (DSB peaks) |
| 1-185 | + | 671 | intron (NM_001004363, intron 4 of 6) | Nuak1 | 5 | tRA treated (DSB peaks) |
| 1-807 | + | 668 | Intergenic | Plekhh2 | 10 | tRA treated (DSB peaks) |
| 1-1289 | + | 656 | promoter-TSS (NM_027399) | Steap1 | 3 | tRA treated (DSB peaks) |

| | | | | | | |
|---|---|---|---|---|---|---|
| 1-871 | + | 634 | promoter-TSS (NM_001081041) | Vps51 | 3 | tRA treated (DSB peaks) |
| 1-1644 | + | 584 | intron (NM_001297601, intron 6 of 8) | Lyl1 | 4 | tRA treated (DSB peaks) |
| 1-224 | + | 915 | promoter-TSS (NM_026538) | Ddx56 | 3 | tRA treated (DSB peaks) |
| 1-972 | + | 580 | Intergenic | Mir130a | 7 | tRA treated (DSB peaks) |
| 1-1645 | + | 660 | intron (NM_001081982, intron 2 of 9) | Nfix | 8 | tRA treated (DSB peaks) |
| 1-1646 | + | 659 | intron (NM_001081982, intron 2 of 9) | Nfix | 4 | tRA treated (DSB peaks) |
| 1-595 | + | 666 | promoter-TSS (NM_023118) | Dab2 | 3 | tRA treated (DSB peaks) |
| 1-596 | + | 666 | promoter-TSS (NM_023118) | Dab2 | 3 | tRA treated (DSB peaks) |
| 1-815 | + | 554 | promoter-TSS (NM_007935) | Epc1 | 3 | tRA treated (DSB peaks) |
| 1-526 | + | 542 | promoter-TSS (NM_178279) | Pxk | 3 | tRA treated (DSB peaks) |
| 1-1166 | + | 827 | promoter-TSS (NM_023066) | Asph | 3 | tRA treated (DSB peaks) |
| 1-1095 | + | 702 | 5' UTR (NM_080428, exon 1 of 11) | Fbxw7 | 6 | tRA treated (DSB peaks) |
| 1-973 | + | 639 | Intergenic | Prg2 | 5 | tRA treated (DSB peaks) |
| 1-1647 | + | 661 | TTS (NM_008416) | Junb | 2 | tRA treated (DSB peaks) |
| 1-317 | + | 550 | intron (NM_138681, intron 8 of 25) | Bcas3 | 3 | tRA treated (DSB peaks) |
| 1-232 | + | 576 | promoter-TSS (NM_146243) | Actr2 | 3 | tRA treated (DSB peaks) |
| 1-675 | + | 574 | promoter-TSS (NM_001347176) | Dvl3 | 3 | tRA treated (DSB peaks) |
| 1-318 | + | 599 | Intergenic | Tbx4 | 5 | tRA treated (DSB peaks) |
| 1-533 | + | 584 | promoter-TSS (NM_010811) | Ndst2 | 3 | tRA treated (DSB peaks) |
| 1-1439 | + | 650 | non-coding (NR_131125, exon 5 of 5) | A430078I02Rik | 1 | tRA treated (DSB peaks) |
| 1-1440 | + | 650 | non-coding (NR_131125, exon 5 of 5) | A430078I02Rik | 2 | tRA treated (DSB peaks) |
| 14977 | + | 725 | Intergenic | Ptma | 2 | tRA treated (DSB peaks) |
| 15342 | + | 686 | Intergenic | Ptma | 5 | tRA treated (DSB peaks) |
| 15707 | + | 685 | Intergenic | Ptma | 4 | tRA treated (DSB peaks) |
| 16072 | + | 556 | Intergenic | Ptma | 14 | tRA treated (DSB peaks) |
| 16438 | + | 556 | Intergenic | Ptma | 15 | tRA treated (DSB peaks) |
| 1-319 | + | 628 | Intergenic | Mir21a | 6 | tRA treated (DSB peaks) |
| 1-320 | + | 628 | Intergenic | Mir21a | 6 | tRA treated (DSB peaks) |
| 1-321 | + | 778 | Intergenic | Mir21a | 8 | tRA treated (DSB peaks) |
| 1-322 | + | 555 | Intergenic | Mir21a | 8 | tRA treated (DSB peaks) |
| 1-1441 | + | 554 | Intergenic | Asprv1 | 2 | tRA treated (DSB peaks) |
| 16803 | + | 880 | 5' UTR (NM_172974, exon 2 of 8) | Cops7b | 3 | tRA treated (DSB peaks) |
| 1-323 | + | 545 | intron (NM_001356531, intron 9 of 11) | Mir21a | 2 | tRA treated (DSB peaks) |
| 1-324 | + | 659 | intron (NM_001356531, intron 5 of 11) | Vmp1 | 5 | tRA treated (DSB peaks) |
| 1-820 | + | 566 | promoter-TSS (NM_001033445) | Garem1 | 3 | tRA treated (DSB peaks) |
| 1-397 | + | 733 | Intergenic | Lrrc74a | 9 | tRA treated (DSB peaks) |
| 1-398 | + | 716 | Intergenic | Irf2bpl | 2 | tRA treated (DSB peaks) |
| 1-399 | + | 542 | Intergenic | Irf2bpl | 7 | tRA treated (DSB peaks) |
| 1-400 | + | 542 | Intergenic | Irf2bpl | 4 | tRA treated (DSB peaks) |
| 1-681 | + | 613 | promoter-TSS (NM_001123037) | Eif4a2 | 3 | tRA treated (DSB peaks) |
| 1-239 | + | 685 | promoter-TSS (NM_023651) | Pus10 | 3 | tRA treated (DSB peaks) |
| 1-821 | + | 562 | promoter-TSS (NM_153058) | Mapre2 | 3 | tRA treated (DSB peaks) |
| 1-808 | + | 556 | Intergenic | C330024C12Rik | 5 | tRA treated (DSB peaks) |
| 1-809 | + | 573 | Intergenic | C330024C12Rik | 4 | tRA treated (DSB peaks) |
| 1-810 | + | 573 | Intergenic | C330024C12Rik | 2 | tRA treated (DSB peaks) |
| 1-811 | + | 684 | intron (NM_028639, intron 5 of 19) | Ttc7 | 4 | tRA treated (DSB peaks) |
| 1-745 | + | 949 | promoter-TSS (NM_025954) | Pgp | 3 | tRA treated (DSB peaks) |

| | | | | | | |
|---|---|---|---|---|---|---|
| 1-1400 | + | 744 | promoter-TSS (NM_080847) | Ndufa5 | 3 | tRA treated (DSB peaks) |
| 1-1698 | + | 681 | promoter-TSS (NM_001205367) | Sept7 | 3 | tRA treated (DSB peaks) |
| 1-1210 | + | 694 | 5' UTR (NM_027326, exon 1 of 11) | Mllt3 | 2 | tRA treated (DSB peaks) |
| 1-936 | + | 648 | promoter-TSS (NM_207234) | Rexo4 | 3 | tRA treated (DSB peaks) |
| 1-1541 | + | 1000 | Intergenic | Ctsc | 13 | tRA treated (DSB peaks) |
| 1-943 | + | 545 | promoter-TSS (NM_138748) | Crat | 3 | tRA treated (DSB peaks) |
| 1-1302 | + | 801 | promoter-TSS (NM_133918) | Emilin1 | 3 | tRA treated (DSB peaks) |
| 1-538 | + | 658 | promoter-TSS (NM_029626) | Glt8d1 | 3 | tRA treated (DSB peaks) |
| 1-1448 | + | 576 | Intergenic | Tpra1 | 2 | tRA treated (DSB peaks) |
| 1-1449 | + | 576 | Intergenic | Tpra1 | 2 | tRA treated (DSB peaks) |
| 1-1304 | + | 719 | promoter-TSS (NM_028150) | Slc4a1ap | 3 | tRA treated (DSB peaks) |
| 1-325 | + | 628 | Intergenic | Gm525 | 5 | tRA treated (DSB peaks) |
| 1-824 | + | 617 | promoter-TSS (NR_038151) | Epb41l4aos | 3 | tRA treated (DSB peaks) |
| 1-1072 | + | 709 | promoter-TSS (NM_025978) | Ttc14 | 3 | tRA treated (DSB peaks) |
| 1-1097 | + | 651 | intron (NM_011368, intron 1 of 12) | Shc1 | 3 | tRA treated (DSB peaks) |
| 1-1098 | + | 651 | intron (NM_011368, intron 1 of 12) | Shc1 | 3 | tRA treated (DSB peaks) |
| 1-1542 | + | 606 | Intergenic | Prss23 | 8 | tRA treated (DSB peaks) |
| 1-1543 | + | 554 | intron (NM_181407, intron 3 of 13) | Ccdc81 | 4 | tRA treated (DSB peaks) |
| 1-1404 | + | 684 | promoter-TSS (NM_009658) | Akr1b3 | 3 | tRA treated (DSB peaks) |
| 17168 | + | 749 | intron (NM_029122, intron 7 of 18) | Iqca | 4 | tRA treated (DSB peaks) |
| 17533 | + | 749 | intron (NM_029122, intron 7 of 18) | Iqca | 4 | tRA treated (DSB peaks) |
| 1-726 | + | 594 | non-coding (NR_033476, exon 1 of 1) | Gm10789 | 5 | tRA treated (DSB peaks) |
| 1-1180 | + | 660 | promoter-TSS (NM_015824) | Orc3 | 3 | tRA treated (DSB peaks) |
| 17899 | + | 752 | Intergenic | Ackr3 | 6 | tRA treated (DSB peaks) |
| 1-1513 | + | 767 | promoter-TSS (NM_001360818) | Kctd15 | 3 | tRA treated (DSB peaks) |
| 1-1099 | + | 654 | 5' UTR (NM_011313, exon 1 of 3) | S100a6 | 7 | tRA treated (DSB peaks) |
| 43252 | + | 558 | promoter-TSS (NM_033570) | Cnnm4 | 3 | tRA treated (DSB peaks) |
| 1-459 | + | 603 | promoter-TSS (NM_026382) | Snrnp48 | 3 | tRA treated (DSB peaks) |
| 18629 | + | 583 | Intergenic | Rab17 | 7 | tRA treated (DSB peaks) |
| 18994 | + | 583 | Intergenic | Rab17 | 5 | tRA treated (DSB peaks) |
| 1-898 | + | 1000 | promoter-TSS (NM_001081075) | Fra10ac1 | 3 | tRA treated (DSB peaks) |
| 1-975 | + | 643 | non-coding (NR_110981, exon 8 of 8) | Psmc3 | 10 | tRA treated (DSB peaks) |
| 1-463 | + | 747 | promoter-TSS (NM_025935) | Tbc1d7 | 3 | tRA treated (DSB peaks) |
| 1-1649 | + | 573 | exon (NM_001282000, exon 1 of 21) | Rbl2 | 3 | tRA treated (DSB peaks) |
| 1-507 | + | 1000 | intron (NM_025770, intron 2 of 7) | Atg10 | 6 | tRA treated (DSB peaks) |
| 1-189 | + | 661 | Intergenic | Tmpo | 6 | tRA treated (DSB peaks) |
| 1-464 | + | 747 | promoter-TSS (NM_025935) | Tbc1d7 | 3 | tRA treated (DSB peaks) |
| 1-1515 | + | 546 | promoter-TSS (NM_145582) | Ctu1 | 3 | tRA treated (DSB peaks) |
| 1-1758 | + | 615 | intron (NM_011636, intron 1 of 8) | Plscr1 | 9 | tRA treated (DSB peaks) |
| 1-1325 | + | 650 | 5' UTR (NM_007644, exon 1 of 12) | Scarb2 | 3 | tRA treated (DSB peaks) |
| 1-1650 | + | 576 | Intergenic | Irx6 | 7 | tRA treated (DSB peaks) |
| 1-1450 | + | 735 | intron (NM_175314, intron 18 of 29) | Adamts9 | 2 | tRA treated (DSB peaks) |
| 1-1712 | + | 572 | promoter-TSS (NM_030256) | Bcl9l | 3 | tRA treated (DSB peaks) |
| 1-1517 | + | 546 | promoter-TSS (NM_030693) | Atf5 | 3 | tRA treated (DSB peaks) |
| 1-465 | + | 808 | promoter-TSS (NM_153789) | Mylip | 3 | tRA treated (DSB peaks) |
| 1-1100 | + | 676 | Intergenic | S100a10 | 14 | tRA treated (DSB peaks) |
| 1-1520 | + | 652 | promoter-TSS (NM_203280) | Rpl18 | 3 | tRA treated (DSB peaks) |

| | | | | | | |
|---|---|---|---|---|---|---|
| 1-326 | + | 643 | 5' UTR (NM_008704, exon 1 of 5) | Nme1 | 16 | tRA treated (DSB peaks) |
| 1-776 | + | 884 | promoter-TSS (NM_001161723) | Tfeb | 3 | tRA treated (DSB peaks) |
| 1-545 | + | 591 | promoter-TSS (NM_172600) | Tmem260 | 3 | tRA treated (DSB peaks) |
| 1-191 | + | 556 | intron (NM_011629, intron 1 of 13) | Nr2c1 | 2 | tRA treated (DSB peaks) |
| 1-327 | + | 556 | Intergenic | Wfikkn2 | 3 | tRA treated (DSB peaks) |
| 1-1718 | + | 700 | promoter-TSS (NM_028300) | Pih1d2 | 3 | tRA treated (DSB peaks) |
| 1-1102 | + | 851 | intron (NM_029431, intron 1 of 5) | Them4 | 4 | tRA treated (DSB peaks) |
| 1-834 | + | 688 | promoter-TSS (NM_178686) | Cep120 | 3 | tRA treated (DSB peaks) |
| 1-150 | + | 657 | promoter-TSS (NR_038016) | Man1a | 3 | tRA treated (DSB peaks) |
| 1-402 | + | 611 | Intergenic | Mir8099-2 | 4 | tRA treated (DSB peaks) |
| 43101 | + | 732 | promoter-TSS (NM_001033194) | Gtf3c3 | 3 | tRA treated (DSB peaks) |
| 1-558 | + | 588 | promoter-TSS (NM_010590) | Ajuba | 3 | tRA treated (DSB peaks) |
| 1-192 | + | 676 | intron (NM_018797, intron 21 of 30) | Plxnc1 | 6 | tRA treated (DSB peaks) |
| 1-331 | + | 545 | Intergenic | Gm11544 | 3 | tRA treated (DSB peaks) |
| 1-1651 | + | 689 | intron (NM_001033533, intron 1 of 8) | Ccdc102a | 3 | tRA treated (DSB peaks) |
| 1-332 | + | 661 | intron (NM_007742, intron 42 of 50) | Col1a1 | 45 | tRA treated (DSB peaks) |
| 1-333 | + | 711 | Intergenic | Ppp1r9b | 8 | tRA treated (DSB peaks) |
| 1-1194 | + | 597 | promoter-TSS (NM_009011) | Rad23b | 3 | tRA treated (DSB peaks) |
| 1-716 | + | 626 | promoter-TSS (NM_029092) | Trmt10c | 3 | tRA treated (DSB peaks) |
| 1-1721 | + | 629 | promoter-TSS (NM_025812) | Peak1 | 3 | tRA treated (DSB peaks) |
| 1-560 | + | 547 | promoter-TSS (NM_198605) | Mrpl57 | 3 | tRA treated (DSB peaks) |
| 1-482 | + | 871 | promoter-TSS (NM_001301343) | Rmi1 | 3 | tRA treated (DSB peaks) |
| 43831 | + | 582 | promoter-TSS (NM_027351) | Ppil3 | 3 | tRA treated (DSB peaks) |
| 44197 | + | 582 | promoter-TSS (NM_027351) | Ppil3 | 3 | tRA treated (DSB peaks) |
| 1-483 | + | 600 | promoter-TSS (NR_015349) | Etohd2 | 3 | tRA treated (DSB peaks) |
| 1-561 | + | 608 | promoter-TSS (NM_144843) | Mtmr6 | 3 | tRA treated (DSB peaks) |
| 1-1729 | + | 568 | promoter-TSS (NM_018853) | Rplp1 | 3 | tRA treated (DSB peaks) |
| 1-283 | + | 601 | promoter-TSS (NR_030738) | 2410006H16Rik | 3 | tRA treated (DSB peaks) |
| 1-652 | + | 549 | intron (NM_175344, intron 9 of 19) | LOC102634389 | 5 | tRA treated (DSB peaks) |
| 1-1731 | + | 697 | promoter-TSS (NM_010193) | Fem1b | 3 | tRA treated (DSB peaks) |
| 1-157 | + | 840 | promoter-TSS (NM_001079824) | Hnrnph3 | 3 | tRA treated (DSB peaks) |
| 46023 | + | 650 | promoter-TSS (NM_001160039) | Eef1b2 | 3 | tRA treated (DSB peaks) |
| 1-193 | + | 833 | Intergenic | 4930459C07Rik | 2 | tRA treated (DSB peaks) |
| 1-849 | + | 569 | promoter-TSS (NM_134138) | Psmg2 | 3 | tRA treated (DSB peaks) |
| 1-385 | + | 633 | promoter-TSS (NM_025589) | Rpl36al | 3 | tRA treated (DSB peaks) |
| 1-1526 | + | 648 | promoter-TSS (NM_009697) | Nr2f2 | 3 | tRA treated (DSB peaks) |
| 1-1634 | + | 703 | promoter-TSS (NM_008452) | Klf2 | 3 | tRA treated (DSB peaks) |
| 1-336 | + | 569 | intron (NM_001177898, intron 10 of 10) | Snx11 | 4 | tRA treated (DSB peaks) |
| 1-299 | + | 624 | promoter-TSS (NM_009671) | Ankfy1 | 3 | tRA treated (DSB peaks) |
| 1-961 | + | 617 | promoter-TSS (NM_001018042) | Sp3 | 3 | tRA treated (DSB peaks) |
| 1-1760 | + | 710 | intron (NM_138756, intron 1 of 6) | Slc25a36 | 1 | tRA treated (DSB peaks) |
| 1-1635 | + | 693 | promoter-TSS (NM_001316753) | Hmgxb4 | 3 | tRA treated (DSB peaks) |
| 1-305 | + | 722 | promoter-TSS (NM_010430) | Hic1 | 3 | tRA treated (DSB peaks) |
| 1-307 | + | 736 | promoter-TSS (NM_008916) | Inpp5k | 3 | tRA treated (DSB peaks) |
| 1-503 | + | 772 | promoter-TSS (NR_153410) | Arrdc3 | 3 | tRA treated (DSB peaks) |
| 1-504 | + | 773 | promoter-TSS (NM_001042591) | Arrdc3 | 3 | tRA treated (DSB peaks) |
| 1-176 | + | 940 | promoter-TSS (NM_001282096) | Tjp3 | 3 | tRA treated (DSB peaks) |

| | | | | | | |
|---|---|---|---|---|---|---|
| 1-732 | + | 683 | 5' UTR (NM_001081684, exon 1 of 2) | Zbtb21 | 4 | tRA treated (DSB peaks) |
| 21186 | + | 614 | 5' UTR (NM_013626, exon 2 of 26) | B230216N24Rik | 2 | tRA treated (DSB peaks) |
| 21551 | + | 614 | intron (NM_001357127, intron 1 of 25) | B230216N24Rik | 3 | tRA treated (DSB peaks) |
| 1-340 | + | 734 | Intergenic | Plxdc1 | 10 | tRA treated (DSB peaks) |
| 13516 | + | 574 | promoter-TSS (NM_010472) | Agfg1 | 3 | tRA treated (DSB peaks) |
| 1-1544 | + | 672 | intron (NM_008663, intron 2 of 48) | Myo7a | 5 | tRA treated (DSB peaks) |
| 1-1329 | + | 554 | intron (NM_001024614, intron 1 of 5) | 1700007G11Rik | 2 | tRA treated (DSB peaks) |
| 1-725 | + | 808 | promoter-TSS (NM_017404) | Mrpl39 | 3 | tRA treated (DSB peaks) |
| 1-1215 | + | 561 | intron (NM_001355177, intron 18 of 40) | Patj | 3 | tRA treated (DSB peaks) |
| 1-1216 | + | 561 | intron (NM_001355177, intron 18 of 40) | Patj | 3 | tRA treated (DSB peaks) |
| 1-1546 | + | 593 | Intergenic | Lrrc32 | 4 | tRA treated (DSB peaks) |
| 1-1547 | + | 543 | Intergenic | Lrrc32 | 5 | tRA treated (DSB peaks) |
| 1-187 | + | 657 | promoter-TSS (NM_001302886) | Prdm4 | 3 | tRA treated (DSB peaks) |
| 1-1548 | + | 770 | Intergenic | Wnt11 | 8 | tRA treated (DSB peaks) |
| 1-1549 | + | 769 | Intergenic | Wnt11 | 7 | tRA treated (DSB peaks) |
| 1-1213 | + | 649 | promoter-TSS (NM_009877) | Cdkn2a | 3 | tRA treated (DSB peaks) |
| 1-974 | + | 680 | promoter-TSS (NM_001311116) | Ndufs3 | 3 | tRA treated (DSB peaks) |
| 19725 | + | 648 | promoter-TSS (NM_178051) | Gm28535 | 3 | tRA treated (DSB peaks) |
| 1-1110 | + | 669 | Intergenic | Gm12440 | 5 | tRA treated (DSB peaks) |
| 1-730 | + | 678 | promoter-TSS (NM_145482) | Setd4 | 3 | tRA treated (DSB peaks) |
| 1-1551 | + | 725 | Intergenic | Serpinh1 | 15 | tRA treated (DSB peaks) |
| 1-1552 | + | 725 | Intergenic | Serpinh1 | 18 | tRA treated (DSB peaks) |
| 1-657 | + | 564 | intron (NM_001310622, intron 1 of 25) | Fmnl3 | 1 | tRA treated (DSB peaks) |
| 1-512 | + | 734 | intron (NM_173757, intron 4 of 10) | 6430562O15Rik | 3 | tRA treated (DSB peaks) |
| 1-1451 | + | 632 | Intergenic | Foxp1 | 2 | tRA treated (DSB peaks) |
| 1-1452 | + | 632 | Intergenic | Foxp1 | 1 | tRA treated (DSB peaks) |
| 1-658 | + | 602 | Intergenic | Aqp6 | 6 | tRA treated (DSB peaks) |
| 1-1101 | + | 849 | promoter-TSS (NM_029431) | Them4 | 3 | tRA treated (DSB peaks) |
| 1-1652 | + | 645 | promoter-TSS (NM_008187) | Cfap20 | 3 | tRA treated (DSB peaks) |
| 1-660 | + | 726 | non-coding (NR_028124, exon 1 of 1) | Larp4 | 5 | tRA treated (DSB peaks) |
| 1-1654 | + | 635 | promoter-TSS (NM_010325) | Got2 | 3 | tRA treated (DSB peaks) |
| 1-1761 | + | 713 | Intergenic | Il20rb | 1 | tRA treated (DSB peaks) |
| 1-731 | + | 595 | promoter-TSS (NM_207301) | Wrb | 3 | tRA treated (DSB peaks) |
| 1-661 | + | 896 | intron (NM_001355497, intron 1 of 15) | Tfcp2 | 8 | tRA treated (DSB peaks) |
| 1-196 | + | 688 | Intergenic | 1700017N19Rik | 2 | tRA treated (DSB peaks) |
| 1-194 | + | 781 | promoter-TSS (NM_007569) | Btg1 | 3 | tRA treated (DSB peaks) |
| 1-195 | + | 569 | promoter-TSS (NM_007569) | Btg1 | 3 | tRA treated (DSB peaks) |
| 20821 | + | 609 | promoter-TSS (NM_173760) | Gin1 | 3 | tRA treated (DSB peaks) |
| 1-511 | + | 847 | promoter-TSS (NM_001048267) | Tnpo1 | 3 | tRA treated (DSB peaks) |
| 1-404 | + | 708 | promoter-TSS (NM_001330702) | Rps6ka5 | 3 | tRA treated (DSB peaks) |
| 1-1767 | + | 569 | promoter-TSS (NM_172460) | Nphp3 | 3 | tRA treated (DSB peaks) |
| 1-406 | + | 802 | promoter-TSS (NM_148948) | Dicer1 | 3 | tRA treated (DSB peaks) |
| 1-1218 | + | 837 | intron (NM_146145, intron 1 of 24) | Jak1 | 5 | tRA treated (DSB peaks) |
| 1-1453 | + | 715 | intron (NM_018884, intron 3 of 9) | Pdzrn3 | 2 | tRA treated (DSB peaks) |
| 1-1454 | + | 715 | intron (NM_018884, intron 3 of 9) | Pdzrn3 | 3 | tRA treated (DSB peaks) |
| 1-1554 | + | 762 | promoter-TSS (NM_025301) | Mrpl17 | 3 | tRA treated (DSB peaks) |
| 1-581 | + | 636 | intron (NM_201529, intron 1 of 30) | Lmo7 | 4 | tRA treated (DSB peaks) |

| | | | | | | |
|---|---|---|---|---|---|---|
| 1-582 | + | 637 | intron (NM_201529, intron 2 of 30) | Lmo7 | 5 | tRA treated (DSB peaks) |
| 1-583 | + | 554 | intron (NM_201529, intron 2 of 30) | Lmo7 | 7 | tRA treated (DSB peaks) |
| 1-1223 | + | 606 | promoter-TSS (NM_001301368) | Cdkn2c | 3 | tRA treated (DSB peaks) |
| 1-513 | + | 549 | Intergenic | Cd180 | 3 | tRA treated (DSB peaks) |
| 1-346 | + | 832 | Intergenic | BC030867 | 7 | tRA treated (DSB peaks) |
| 1-347 | + | 588 | Intergenic | BC030867 | 5 | tRA treated (DSB peaks) |
| 1-662 | + | 542 | intron (NM_011244, intron 3 of 9) | Rarg | 1 | tRA treated (DSB peaks) |
| 1-979 | + | 753 | exon (NM_020267, exon 1 of 5) | Trim44 | 15 | tRA treated (DSB peaks) |
| 1-348 | + | 585 | intron (NM_010575, intron 4 of 29) | Itga2b | 7 | tRA treated (DSB peaks) |
| 1-1566 | + | 862 | promoter-TSS (NM_001040131) | Eif4g2 | 3 | tRA treated (DSB peaks) |
| 1-1343 | + | 741 | promoter-TSS (NM_175403) | Mlec | 3 | tRA treated (DSB peaks) |
| 1-1229 | + | 612 | promoter-TSS (NM_011034) | Prdx1 | 3 | tRA treated (DSB peaks) |
| 1-1788 | + | 767 | promoter-TSS (NM_001048146) | Azi2 | 3 | tRA treated (DSB peaks) |
| 1-1764 | + | 927 | intron (NM_019764, intron 5 of 9) | Amotl2 | 5 | tRA treated (DSB peaks) |
| 1-1765 | + | 656 | Intergenic | Gm5627 | 2 | tRA treated (DSB peaks) |
| 1-980 | + | 703 | Intergenic | Cd44 | 4 | tRA treated (DSB peaks) |
| 22647 | + | 670 | promoter-TSS (NR_153836) | Tsn | 3 | tRA treated (DSB peaks) |
| 1-350 | + | 552 | intron (NM_026551, intron 1 of 4) | Mir6931 | 6 | tRA treated (DSB peaks) |
| 1-351 | + | 553 | intron (NM_008707, intron 3 of 11) | Nmt1 | 5 | tRA treated (DSB peaks) |
| 1-1655 | + | 612 | Intergenic | 1600027J07Rik | 4 | tRA treated (DSB peaks) |
| 1-197 | + | 1000 | Intergenic | Slc6a15 | 6 | tRA treated (DSB peaks) |
| 1-994 | + | 627 | promoter-TSS (NM_026574) | Ino80 | 3 | tRA treated (DSB peaks) |
| 1-1656 | + | 706 | Intergenic | Cdh5 | 3 | tRA treated (DSB peaks) |
| 1-1579 | + | 600 | promoter-TSS (NM_025298) | Polr3e | 3 | tRA treated (DSB peaks) |
| 1-366 | + | 561 | promoter-TSS (NM_027745) | Ccdc57 | 3 | tRA treated (DSB peaks) |
| 1-1657 | + | 540 | Intergenic | Cdh5 | 9 | tRA treated (DSB peaks) |
| 1-514 | + | 768 | intron (NM_029447, intron 12 of 12) | Nln | 10 | tRA treated (DSB peaks) |
| 1-515 | + | 636 | intron (NM_029447, intron 12 of 12) | Nln | 5 | tRA treated (DSB peaks) |
| 1-1128 | + | 584 | promoter-TSS (NR_028574) | Snhg8 | 3 | tRA treated (DSB peaks) |
| 1-1480 | + | 596 | promoter-TSS (NM_175557) | Zfp384 | 3 | tRA treated (DSB peaks) |
| 1-1245 | + | 581 | promoter-TSS (NM_025544) | Mrps15 | 3 | tRA treated (DSB peaks) |
| 1-209 | + | 568 | promoter-TSS (NM_009870) | Cdk4 | 3 | tRA treated (DSB peaks) |
| 1-982 | + | 645 | exon (NM_009037, exon 1 of 6) | Rcn1 | 4 | tRA treated (DSB peaks) |
| 1-1483 | + | 772 | promoter-TSS (NM_001346512) | Parp11 | 3 | tRA treated (DSB peaks) |
| 1-215 | + | 674 | promoter-TSS (NR_106183) | Mir8105 | 3 | tRA treated (DSB peaks) |
| 1-1331 | + | 567 | intron (NM_001122768, intron 2 of 2) | Lrrc8d | 22 | tRA treated (DSB peaks) |
| 1-1133 | + | 577 | promoter-TSS (NM_130450) | Elovl6 | 3 | tRA treated (DSB peaks) |
| 1-1134 | + | 577 | promoter-TSS (NR_153369) | Gm35986 | 3 | tRA treated (DSB peaks) |
| 1-1220 | + | 622 | Intergenic | Gm12724 | 2 | tRA treated (DSB peaks) |
| 1-983 | + | 577 | non-coding (NR_033993, exon 1 of 4) | Dnajc24 | 1 | tRA treated (DSB peaks) |
| 1-585 | + | 690 | Intergenic | Trim52 | 12 | tRA treated (DSB peaks) |
| 1-1261 | + | 649 | promoter-TSS (NM_025919) | Rpl11 | 3 | tRA treated (DSB peaks) |
| 1-353 | + | 573 | exon (NM_027346, exon 1 of 5) | Taco1 | 8 | tRA treated (DSB peaks) |
| 1-1377 | + | 571 | promoter-TSS (NM_008788) | Pcolce | 3 | tRA treated (DSB peaks) |
| 1-1597 | + | 550 | promoter-TSS (NM_027201) | Zfp511 | 3 | tRA treated (DSB peaks) |
| 1-1598 | + | 695 | promoter-TSS (NM_053119) | Echs1 | 3 | tRA treated (DSB peaks) |
| 1-1769 | + | 559 | Intergenic | Rpl29 | 3 | tRA treated (DSB peaks) |

| | | | | | | |
|---|---|---|---|---|---|---|
| 1-1599 | + | 544 | promoter-TSS (NM_199301) | Mtg1 | 3 | tRA treated (DSB peaks) |
| 1-1387 | + | 619 | promoter-TSS (NM_177682) | Ccz1 | 3 | tRA treated (DSB peaks) |
| 1-1146 | + | 557 | promoter-TSS (NM_001282042) | Sh3glb1 | 3 | tRA treated (DSB peaks) |
| 1-1389 | + | 697 | promoter-TSS (NM_001256086) | Rnf6 | 3 | tRA treated (DSB peaks) |
| 21916 | + | 673 | intron (NM_027534, intron 1 of 9) | Kdsr | 3 | tRA treated (DSB peaks) |
| 1-1332 | + | 665 | Intergenic | Cdc7 | 9 | tRA treated (DSB peaks) |
| 1-1333 | + | 665 | Intergenic | Cdc7 | 6 | tRA treated (DSB peaks) |
| 1-1334 | + | 594 | Intergenic | Cdc7 | 2 | tRA treated (DSB peaks) |
| 1-198 | + | 587 | Intergenic | Acss3 | 5 | tRA treated (DSB peaks) |
| 1-1277 | + | 586 | promoter-TSS (NM_029035) | Spsb1 | 3 | tRA treated (DSB peaks) |
| 1-1027 | + | 647 | promoter-TSS (NM_172117) | Entpd6 | 3 | tRA treated (DSB peaks) |
| 1-1771 | + | 663 | 5' UTR (NM_009895, exon 1 of 3) | Cish | 8 | tRA treated (DSB peaks) |
| 1-1032 | + | 608 | promoter-TSS (NM_001163452) | Trpc4ap | 3 | tRA treated (DSB peaks) |
| 1-516 | + | 692 | intron (NM_145456, intron 2 of 14) | Zswim6 | 3 | tRA treated (DSB peaks) |
| 1-1338 | + | 609 | intron (NM_016980, intron 1 of 7) | Rpl5 | 2 | tRA treated (DSB peaks) |
| 1-408 | + | 591 | intron (NM_001286343, intron 1 of 1) | Bcl11b | 5 | tRA treated (DSB peaks) |
| 1-409 | + | 766 | intron (NM_001286343, intron 1 of 1) | Bcl11b | 8 | tRA treated (DSB peaks) |
| 31413 | + | 577 | promoter-TSS (NM_001160182) | Tor1aip2 | 3 | tRA treated (DSB peaks) |
| 1-1114 | + | 679 | exon (NM_001289545, exon 1 of 11) | Atxn7l2 | 1 | tRA treated (DSB peaks) |
| 1-1115 | + | 641 | Intergenic | Psrc1 | 3 | tRA treated (DSB peaks) |
| 1-1159 | + | 565 | promoter-TSS (NM_001289524) | Srsf11 | 3 | tRA treated (DSB peaks) |
| 1-1038 | + | 625 | promoter-TSS (NM_001291138) | Ralgapb | 3 | tRA treated (DSB peaks) |
| 1-1772 | + | 623 | 5' UTR (NM_023247, exon 1 of 5) | Ndufaf3 | 1 | tRA treated (DSB peaks) |
| 1-1040 | + | 576 | promoter-TSS (NM_025575) | Sys1 | 3 | tRA treated (DSB peaks) |
| 1-517 | + | 598 | Intergenic | Pde4d | 7 | tRA treated (DSB peaks) |
| 1-1455 | + | 796 | Intergenic | 0610040F04Rik | 3 | tRA treated (DSB peaks) |
| 1-1118 | + | 553 | intron (NM_177859, intron 6 of 11) | Aknad1 | 5 | tRA treated (DSB peaks) |
| 1-1041 | + | 559 | promoter-TSS (NM_133779) | Pigt | 3 | tRA treated (DSB peaks) |
| 1-1288 | + | 563 | promoter-TSS (NM_001042501) | Fam133b | 2 | tRA treated (DSB peaks) |
| 1-371 | + | 798 | promoter-TSS (NM_001099628) | Atad2b | 2 | tRA treated (DSB peaks) |
| 1-1557 | + | 805 | Intergenic | Tub | 4 | tRA treated (DSB peaks) |
| 1-817 | + | 575 | promoter-TSS (NM_177595) | Mkx | 2 | tRA treated (DSB peaks) |
| 1-372 | + | 745 | promoter-TSS (NM_021429) | Hs1bp3 | 2 | tRA treated (DSB peaks) |
| 1-884 | + | 726 | promoter-TSS (NM_015735) | Ddb1 | 2 | tRA treated (DSB peaks) |
| 1-1120 | + | 550 | intron (NM_172685, intron 1 of 9) | Slc25a24 | 3 | tRA treated (DSB peaks) |
| 1-1685 | + | 621 | promoter-TSS (NM_001199484) | 4931406C07Rik | 2 | tRA treated (DSB peaks) |
| 1-1498 | + | 713 | promoter-TSS (NM_198631) | Zc3h4 | 2 | tRA treated (DSB peaks) |
| 1-1500 | + | 1000 | promoter-TSS (NM_007590) | Calm3 | 2 | tRA treated (DSB peaks) |
| 1-1558 | + | 640 | Intergenic | Akip1 | 5 | tRA treated (DSB peaks) |
| 1-1559 | + | 640 | Intergenic | Akip1 | 6 | tRA treated (DSB peaks) |
| 1-1689 | + | 687 | promoter-TSS (NM_001004190) | Zfp560 | 2 | tRA treated (DSB peaks) |
| 1-676 | + | 574 | promoter-TSS (NM_001347176) | Dvl3 | 2 | tRA treated (DSB peaks) |
| 1-1293 | + | 643 | promoter-TSS (NM_001359963) | Napepld | 2 | tRA treated (DSB peaks) |
| 1-985 | + | 600 | 5' UTR (NM_172671, exon 1 of 18) | Lgr4 | 4 | tRA treated (DSB peaks) |
| 1-1173 | + | 729 | promoter-TSS (NM_001356348) | Coq3 | 2 | tRA treated (DSB peaks) |
| 1-940 | + | 583 | promoter-TSS (NM_001166031) | Mrps2 | 2 | tRA treated (DSB peaks) |
| 1-1340 | + | 614 | intron (NM_145147, intron 1 of 9) | Gtpbp6 | 7 | tRA treated (DSB peaks) |

| | | | | | | |
|---|---|---|---|---|---|---|
| 1-412 | + | 632 | Intergenic | 3110009F21Rik | 1 | tRA treated (DSB peaks) |
| 1-1560 | + | 596 | Intergenic | Wee1 | 10 | tRA treated (DSB peaks) |
| 1-1561 | + | 546 | Intergenic | Wee1 | 8 | tRA treated (DSB peaks) |
| 1-1562 | + | 742 | Intergenic | Swap70 | 3 | tRA treated (DSB peaks) |
| 1-888 | + | 913 | promoter-TSS (NM_001081213) | Ermp1 | 2 | tRA treated (DSB peaks) |
| 1-1778 | + | 775 | intron (NM_001360306, intron 3 of 9) | Klhl18 | 4 | tRA treated (DSB peaks) |
| 1-945 | + | 743 | promoter-TSS (NM_144885) | Usp20 | 2 | tRA treated (DSB peaks) |
| 1-413 | + | 684 | Intergenic | 1700001K19Rik | 6 | tRA treated (DSB peaks) |
| 1-1563 | + | 653 | Intergenic | Ampd3 | 9 | tRA treated (DSB peaks) |
| 1-1564 | + | 602 | Intergenic | Ampd3 | 11 | tRA treated (DSB peaks) |
| 1-414 | + | 951 | Intergenic | Hsp90aa1 | 12 | tRA treated (DSB peaks) |
| 1-415 | + | 585 | Intergenic | Hsp90aa1 | 11 | tRA treated (DSB peaks) |
| 1-1565 | + | 717 | Intergenic | Ampd3 | 4 | tRA treated (DSB peaks) |
| 1-1780 | + | 829 | Intergenic | Prss42 | 1 | tRA treated (DSB peaks) |
| 1-1781 | + | 730 | Intergenic | Prss42 | 9 | tRA treated (DSB peaks) |
| 1-518 | + | 546 | non-coding (NR_131193, exon 5 of 5) | 4930526H09Rik | 5 | tRA treated (DSB peaks) |
| 1-1659 | + | 736 | intron (NM_009179, intron 1 of 6) | St3gal2 | 10 | tRA treated (DSB peaks) |
| 1-1177 | + | 572 | promoter-TSS (NM_172688) | Map3k7 | 2 | tRA treated (DSB peaks) |
| 1-1567 | + | 561 | intron (NR_015573, intron 1 of 2) | 1700012D14Rik | 3 | tRA treated (DSB peaks) |
| 1-1071 | + | 585 | promoter-TSS (NM_144519) | Zfp639 | 2 | tRA treated (DSB peaks) |
| 1-199 | + | 779 | Intergenic | Nap1l1 | 2 | tRA treated (DSB peaks) |
| 1-1073 | + | 566 | promoter-TSS (NM_025978) | Ttc14 | 2 | tRA treated (DSB peaks) |
| 1-1074 | + | 784 | promoter-TSS (NM_001286972) | Dnajc19 | 2 | tRA treated (DSB peaks) |
| 1-200 | + | 544 | exon (NM_009344, exon 1 of 2) | Phlda1 | 2 | tRA treated (DSB peaks) |
| 1-201 | + | 544 | exon (NM_009344, exon 1 of 2) | Phlda1 | 3 | tRA treated (DSB peaks) |
| 1-1411 | + | 542 | promoter-TSS (NM_172893) | 4930599N23Rik | 2 | tRA treated (DSB peaks) |
| 1-1191 | + | 745 | promoter-TSS (NM_001190414) | Gne | 2 | tRA treated (DSB peaks) |
| 1-770 | + | 567 | promoter-TSS (NM_199016) | Enpp4 | 2 | tRA treated (DSB peaks) |
| 1-203 | + | 662 | intron (NM_178610, intron 1 of 9) | Krr1 | 7 | tRA treated (DSB peaks) |
| 1-1711 | + | 572 | promoter-TSS (NM_030256) | Bcl9l | 2 | tRA treated (DSB peaks) |
| 1-775 | + | 626 | promoter-TSS (NM_198608) | Aars2 | 2 | tRA treated (DSB peaks) |
| 1-1570 | + | 733 | intron (NM_027587, intron 2 of 9) | Micalcl | 6 | tRA treated (DSB peaks) |
| 1-1571 | + | 733 | intron (NM_027587, intron 2 of 9) | Micalcl | 6 | tRA treated (DSB peaks) |
| 1-1572 | + | 788 | intron (NM_020606, intron 1 of 12) | Parva | 13 | tRA treated (DSB peaks) |
| 1-359 | + | 585 | Intergenic | BC006965 | 5 | tRA treated (DSB peaks) |
| 1-1573 | + | 663 | intron (NM_001166584, intron 1 of 13) | Tead1 | 20 | tRA treated (DSB peaks) |
| 1-1456 | + | 553 | intron (NM_080448, intron 1 of 21) | Srgap3 | 19 | tRA treated (DSB peaks) |
| 1-1457 | + | 741 | 5' UTR (NM_080448, exon 1 of 22) | Srgap3 | 7 | tRA treated (DSB peaks) |
| 1-1782 | + | 699 | Intergenic | AU023762 | 3 | tRA treated (DSB peaks) |
| 1-1458 | + | 579 | intron (NR_027009, intron 1 of 1) | Gt(ROSA)26Sor | 3 | tRA treated (DSB peaks) |
| 1-1616 | + | 711 | promoter-TSS (NM_001039562) | Ankrd37 | 2 | tRA treated (DSB peaks) |
| 1-1460 | + | 564 | intron (NM_028385, intron 1 of 21) | Setd5 | 8 | tRA treated (DSB peaks) |
| 1-1461 | + | 555 | intron (NM_028385, intron 1 of 21) | Setd5 | 5 | tRA treated (DSB peaks) |
| 1-1574 | + | 598 | intron (NR_045978, intron 2 of 5) | 4930543E12Rik | 7 | tRA treated (DSB peaks) |
| 1-520 | + | 573 | Intergenic | Esm1 | 9 | tRA treated (DSB peaks) |
| 1-1417 | + | 563 | promoter-TSS (NM_175098) | Ccdc126 | 2 | tRA treated (DSB peaks) |
| 1-712 | + | 910 | promoter-TSS (NM_010581) | Cd47 | 2 | tRA treated (DSB peaks) |

| | | | | | | |
|---|---|---|---|---|---|---|
| 1-1316 | + | 710 | promoter-TSS (NM_024213) | Anapc4 | 2 | tRA treated (DSB peaks) |
| 1-557 | + | 588 | promoter-TSS (NM_010590) | Ajuba | 2 | tRA treated (DSB peaks) |
| 1-1121 | + | 848 | exon (NM_026043, exon 1 of 15) | Rnpc3 | 3 | tRA treated (DSB peaks) |
| 1-954 | + | 555 | promoter-TSS (NM_010274) | Gpd2 | 2 | tRA treated (DSB peaks) |
| 1-719 | + | 605 | promoter-TSS (NM_025599) | Cmss1 | 2 | tRA treated (DSB peaks) |
| 1-481 | + | 645 | promoter-TSS (NM_029872) | Hnrnpa0 | 2 | tRA treated (DSB peaks) |
| 1-522 | + | 558 | intron (NM_172595, intron 1 of 4) | Arl15 | 3 | tRA treated (DSB peaks) |
| 1-1662 | + | 593 | intron (NM_172466, intron 3 of 22) | Adamts18 | 5 | tRA treated (DSB peaks) |
| 1-272 | + | 699 | promoter-TSS (NR_134309) | Hist3h2a | 2 | tRA treated (DSB peaks) |
| 1-988 | + | 608 | intron (NM_175466, intron 1 of 1) | Zfp770 | 1 | tRA treated (DSB peaks) |
| 1-280 | + | 547 | promoter-TSS (NM_027198) | Ttc19 | 2 | tRA treated (DSB peaks) |
| 1-788 | + | 552 | promoter-TSS (NM_013933) | Vapa | 2 | tRA treated (DSB peaks) |
| 1-1342 | + | 579 | intron (NM_001159941, intron 2 of 6) | Kctd10 | 10 | tRA treated (DSB peaks) |
| 1-1089 | + | 646 | promoter-TSS (NM_001356274) | Rsrc1 | 2 | tRA treated (DSB peaks) |
| 1-1577 | + | 816 | exon (NM_177382, exon 1 of 5) | Cyp2r1 | 3 | tRA treated (DSB peaks) |
| 1-1747 | + | 626 | promoter-TSS (NM_001164254) | Tpm1 | 2 | tRA treated (DSB peaks) |
| 1-1465 | + | 809 | intron (NM_001356371, intron 3 of 5) | Vgll4 | 9 | tRA treated (DSB peaks) |
| 1-1123 | + | 583 | intron (NM_001286750, intron 1 of 5) | Olfm3 | 4 | tRA treated (DSB peaks) |
| 1-628 | + | 607 | promoter-TSS (NM_009177) | St3gal1 | 2 | tRA treated (DSB peaks) |
| 1-1629 | + | 670 | promoter-TSS (NM_001200023) | Zfp963 | 2 | tRA treated (DSB peaks) |
| 1-362 | + | 551 | Intergenic | Atp5h | 8 | tRA treated (DSB peaks) |
| 1-1344 | + | 548 | Intergenic | Sirt4 | 23 | tRA treated (DSB peaks) |
| 1-1345 | + | 548 | Intergenic | Sirt4 | 17 | tRA treated (DSB peaks) |
| 1-1431 | + | 595 | promoter-TSS (NM_145569) | Mat2a | 2 | tRA treated (DSB peaks) |
| 1-795 | + | 617 | promoter-TSS (NM_007566) | Birc6 | 2 | tRA treated (DSB peaks) |
| 1-1225 | + | 683 | exon (NM_181040, exon 1 of 9) | Atpaf1 | 2 | tRA treated (DSB peaks) |
| 1-363 | + | 695 | intron (NM_008211, intron 1 of 3) | H3f3b | 2 | tRA treated (DSB peaks) |
| 1-1226 | + | 783 | exon (NM_028142, exon 1 of 6) | Nsun4 | 4 | tRA treated (DSB peaks) |
| 1-304 | + | 722 | promoter-TSS (NM_010430) | Hic1 | 2 | tRA treated (DSB peaks) |
| 1-645 | + | 550 | promoter-TSS (NM_133800) | Nol12 | 2 | tRA treated (DSB peaks) |
| 1-1784 | + | 668 | Intergenic | Gm4668 | 10 | tRA treated (DSB peaks) |
| 1-970 | + | 579 | promoter-TSS (NM_027091) | Nup35 | 2 | tRA treated (DSB peaks) |
| 1-170 | + | 600 | promoter-TSS (NM_027829) | Mob3a | 2 | tRA treated (DSB peaks) |
| 1-523 | + | 558 | intron (NR_153406, intron 3 of 3) | 4930435F18Rik | 9 | tRA treated (DSB peaks) |
| 1-505 | + | 721 | promoter-TSS (NR_015587) | 9330111N05Rik | 2 | tRA treated (DSB peaks) |
| 1-1467 | + | 753 | exon (NM_021704, exon 1 of 3) | Cxcl12 | 1 | tRA treated (DSB peaks) |
| 1-180 | + | 639 | promoter-TSS (NM_010914) | Nfyb | 2 | tRA treated (DSB peaks) |
| 1-990 | + | 566 | intron (NM_026620, intron 1 of 7) | Fam98b | 4 | tRA treated (DSB peaks) |
| 1-1436 | + | 670 | promoter-TSS (NM_001162521) | Dguok | 2 | tRA treated (DSB peaks) |
| 1-1538 | + | 585 | promoter-TSS (NM_030705) | Tlnrd1 | 2 | tRA treated (DSB peaks) |
| 1-316 | + | 598 | promoter-TSS (NM_145433) | Mrm1 | 2 | tRA treated (DSB peaks) |
| 14246 | + | 639 | promoter-TSS (NM_025386) | Trip12 | 2 | tRA treated (DSB peaks) |
| 14611 | + | 639 | promoter-TSS (NM_025386) | Fbxo36 | 2 | tRA treated (DSB peaks) |
| 1-1468 | + | 669 | Intergenic | Zfp239 | 2 | tRA treated (DSB peaks) |
| 1-991 | + | 563 | intron (NR_040631, intron 3 of 5) | 4930412B13Rik | 4 | tRA treated (DSB peaks) |
| 1-1231 | + | 548 | intron (NM_001285520, intron 5 of 11) | Artn | 6 | tRA treated (DSB peaks) |
| 1-1443 | + | 657 | promoter-TSS (NM_013528) | Gfpt1 | 2 | tRA treated (DSB peaks) |

| | | | | | | |
|---|---|---|---|---|---|---|
| 1-812 | + | 582 | promoter-TSS (NM_007589) | Calm2 | 2 | tRA treated (DSB peaks) |
| 1-1211 | + | 693 | promoter-TSS (NM_027326) | Mllt3 | 2 | tRA treated (DSB peaks) |
| 1-1445 | + | 573 | promoter-TSS (NM_023060) | Eefsec | 2 | tRA treated (DSB peaks) |
| 1-1105 | + | 552 | promoter-TSS (NM_172395) | Cdc42se1 | 2 | tRA treated (DSB peaks) |
| 1-1328 | + | 544 | promoter-TSS (NM_080708) | Bmp2k | 2 | tRA treated (DSB peaks) |
| 1-339 | + | 543 | promoter-TSS (NM_138657) | Socs7 | 2 | tRA treated (DSB peaks) |
| 1-1124 | + | 682 | Intergenic | Mir137 | 4 | tRA treated (DSB peaks) |
| 1-1347 | + | 631 | Intergenic | Med13l | 6 | tRA treated (DSB peaks) |
| 1-1348 | + | 632 | Intergenic | Med13l | 4 | tRA treated (DSB peaks) |
| 1-1349 | + | 793 | Intergenic | Med13l | 4 | tRA treated (DSB peaks) |
| 1-992 | + | 543 | intron (NM_001331221, intron 3 of 3) | Bmf | 2 | tRA treated (DSB peaks) |
| 1-586 | + | 599 | intron (NM_001163676, intron 18 of 29) | Mir6391 | 3 | tRA treated (DSB peaks) |
| 23012 | + | 582 | intron (NM_001081125, intron 2 of 13) | Mir6346 | 11 | tRA treated (DSB peaks) |
| 23377 | + | 570 | intron (NM_001081125, intron 2 of 13) | Mir6346 | 2 | tRA treated (DSB peaks) |
| 1-654 | + | 547 | promoter-TSS (NM_001080981) | Ddx23 | 2 | tRA treated (DSB peaks) |
| 1-1330 | + | 675 | promoter-TSS (NM_001081107) | Helq | 2 | tRA treated (DSB peaks) |
| 1-1234 | + | 612 | Intergenic | Slc2a1 | 5 | tRA treated (DSB peaks) |
| 1-1789 | + | 649 | exon (NM_010851, exon 1 of 6) | Myd88 | 4 | tRA treated (DSB peaks) |
| 1-1553 | + | 861 | promoter-TSS (NM_001146010) | Fchsd2 | 2 | tRA treated (DSB peaks) |
| 1-1111 | + | 564 | promoter-TSS (NM_001253702) | Capza1 | 2 | tRA treated (DSB peaks) |
| 1-205 | + | 617 | intron (NM_133442, intron 1 of 21) | Grip1os2 | 2 | tRA treated (DSB peaks) |
| 1-1768 | + | 553 | promoter-TSS (NM_053159) | Mrpl3 | 2 | tRA treated (DSB peaks) |
| 1-1339 | + | 1000 | promoter-TSS (NM_001253877) | Mtf2 | 2 | tRA treated (DSB peaks) |
| 1-1116 | + | 572 | promoter-TSS (NM_026198) | Tmem167b | 2 | tRA treated (DSB peaks) |
| 1-1222 | + | 544 | promoter-TSS (NM_177045) | Cc2d1b | 2 | tRA treated (DSB peaks) |
| 1-997 | + | 786 | intron (NM_013720, intron 1 of 23) | Mga | 3 | tRA treated (DSB peaks) |
| 1-1472 | + | 546 | non-coding (NR_149727, exon 1 of 11) | Rad52 | 1 | tRA treated (DSB peaks) |
| 1-411 | + | 872 | promoter-TSS (NM_177602) | Wdr25 | 2 | tRA treated (DSB peaks) |
| 1-587 | + | 645 | Intergenic | Oxgr1 | 7 | tRA treated (DSB peaks) |
| 1-588 | + | 644 | Intergenic | Oxgr1 | 7 | tRA treated (DSB peaks) |
| 1-1235 | + | 557 | Intergenic | Scmh1 | 1 | tRA treated (DSB peaks) |
| 1-519 | + | 683 | promoter-TSS (NM_001122963) | Gpbp1 | 2 | tRA treated (DSB peaks) |
| 1-416 | + | 595 | promoter-TSS (NM_173363) | Eif5 | 2 | tRA treated (DSB peaks) |
| 1-1790 | + | 626 | Intergenic | 5830454E08Rik | 2 | tRA treated (DSB peaks) |
| 24108 | + | 549 | Intergenic | En1 | 9 | tRA treated (DSB peaks) |
| 1-1569 | + | 674 | promoter-TSS (NM_177249) | Usp47 | 2 | tRA treated (DSB peaks) |
| 1-1462 | + | 723 | promoter-TSS (NM_026849) | Mtmr14 | 2 | tRA treated (DSB peaks) |
| 1-360 | + | 551 | promoter-TSS (NM_001166503) | Slc39a11 | 2 | tRA treated (DSB peaks) |
| 1-367 | + | 1000 | 5' UTR (NM_139059, exon 1 of 9) | Csnk1d | 2 | tRA treated (DSB peaks) |
| 1-1663 | + | 586 | promoter-TSS (NM_019573) | Wwox | 2 | tRA treated (DSB peaks) |
| 1-1466 | + | 775 | promoter-TSS (NM_001356334) | Raf1 | 2 | tRA treated (DSB peaks) |
| 1-1787 | + | 540 | promoter-TSS (NM_178660) | Rbms3 | 2 | tRA treated (DSB peaks) |
| 1-1125 | + | 758 | Intergenic | Slc44a3 | 7 | tRA treated (DSB peaks) |
| 1-204 | + | 553 | promoter-TSS (NM_024457) | Rap1b | 2 | tRA treated (DSB peaks) |
| 1-1233 | + | 555 | promoter-TSS (NM_001161774) | St3gal3 | 2 | tRA treated (DSB peaks) |
| 1-1469 | + | 552 | promoter-TSS (NM_194339) | Bms1 | 2 | tRA treated (DSB peaks) |
| 1-1470 | + | 552 | promoter-TSS (NM_194339) | Bms1 | 2 | tRA treated (DSB peaks) |

| | | | | | | |
|---|---|---|---|---|---|---|
| 1-207 | + | 689 | intron (NM_153059, intron 1 of 5) | Tmem5 | 6 | tRA treated (DSB peaks) |
| 1-1126 | + | 610 | intron (NM_007378, intron 19 of 49) | Abca4 | 9 | tRA treated (DSB peaks) |
| 1-1471 | + | 898 | promoter-TSS (NM_053204) | 3110021A11Rik | 2 | tRA treated (DSB peaks) |
| 1-1127 | + | 662 | TTS (NR_030439) | Mir760 | 8 | tRA treated (DSB peaks) |
| 1-1791 | + | 719 | promoter-TSS (NM_025339) | Tmem42 | 2 | tRA treated (DSB peaks) |
| 1-999 | + | 1000 | Intergenic | Shf | 12 | tRA treated (DSB peaks) |
| 1-1000 | + | 1000 | Intergenic | Shf | 11 | tRA treated (DSB peaks) |
| 1-1669 | + | 637 | TTS (NR_015497) | 9330133O14Rik | 11 | tRA treated (DSB peaks) |
| 1-1478 | + | 591 | promoter-TSS (NM_007881) | Atn1 | 2 | tRA treated (DSB peaks) |
| 1-1350 | + | 833 | Intergenic | Ift81 | 6 | tRA treated (DSB peaks) |
| 1-1670 | + | 1000 | intron (NM_009824, intron 1 of 11) | Cbfa2t3 | 5 | tRA treated (DSB peaks) |
| 1-1671 | + | 653 | intron (NM_009824, intron 1 of 11) | Cbfa2t3 | 4 | tRA treated (DSB peaks) |
| 1-1474 | + | 559 | Intergenic | Nanog | 5 | tRA treated (DSB peaks) |
| 1-1004 | + | 845 | promoter-TSS (NM_026981) | Dtwd1 | 2 | tRA treated (DSB peaks) |
| 1-1238 | + | 577 | TTS (NM_008917) | Ppt1 | 13 | tRA treated (DSB peaks) |
| 1-1239 | + | 577 | TTS (NM_008917) | Ppt1 | 7 | tRA treated (DSB peaks) |
| 1-1475 | + | 615 | Intergenic | Necap1 | 4 | tRA treated (DSB peaks) |
| 1-1240 | + | 598 | 5' UTR (NM_001358036, exon 1 of 13) | Cap1 | 10 | tRA treated (DSB peaks) |
| 1-208 | + | 755 | intron (NM_001110218, intron 8 of 9) | Mirlet7i | 5 | tRA treated (DSB peaks) |
| 1-1583 | + | 803 | promoter-TSS (NM_183020) | Atxn2l | 2 | tRA treated (DSB peaks) |
| 1-591 | + | 546 | Intergenic | AA536875 | 4 | tRA treated (DSB peaks) |
| 1-1006 | + | 677 | promoter-TSS (NM_025296) | Tmem127 | 2 | tRA treated (DSB peaks) |
| 1-1363 | + | 937 | promoter-TSS (NM_172462) | Zfp11 | 2 | tRA treated (DSB peaks) |
| 1-1241 | + | 584 | intron (NM_001199137, intron 5 of 92) | Mir6398 | 13 | tRA treated (DSB peaks) |
| 1-1592 | + | 540 | promoter-TSS (NM_001039534) | Pstk | 2 | tRA treated (DSB peaks) |
| 1-1136 | + | 855 | promoter-TSS (NM_001163455) | Aimp1 | 2 | tRA treated (DSB peaks) |
| 1-1129 | + | 858 | 5' UTR (NM_146140, exon 1 of 1) | Tram1l1 | 1 | tRA treated (DSB peaks) |
| 1-1017 | + | 546 | promoter-TSS (NM_029148) | Tmx4 | 2 | tRA treated (DSB peaks) |
| 1-1138 | + | 622 | promoter-TSS (NM_025356) | Ube2d3 | 2 | tRA treated (DSB peaks) |
| 1-1672 | + | 594 | intron (NM_175149, intron 1 of 3) | 2310022B05Rik | 5 | tRA treated (DSB peaks) |
| 1-1243 | + | 819 | TTS (NM_010213) | Sf3a3 | 4 | tRA treated (DSB peaks) |
| 1-1600 | + | 541 | promoter-TSS (NM_013782) | Ptdss2 | 2 | tRA treated (DSB peaks) |
| 1-1276 | + | 586 | promoter-TSS (NM_029035) | Spsb1 | 2 | tRA treated (DSB peaks) |
| 29221 | + | 677 | promoter-TSS (NM_007842) | Dhx9 | 2 | tRA treated (DSB peaks) |
| 1-1282 | + | 563 | promoter-TSS (NM_001163019) | A430005L14Rik | 2 | tRA treated (DSB peaks) |
| 1-1287 | + | 585 | promoter-TSS (NM_145557) | 9430015G10Rik | 2 | tRA treated (DSB peaks) |
| 32874 | + | 700 | promoter-TSS (NM_201364) | BC055324 | 2 | tRA treated (DSB peaks) |
| 1-116 | + | 929 | promoter-TSS (NM_001163421) | Tatdn3 | 2 | tRA treated (DSB peaks) |
| 1-1001 | + | 603 | Intergenic | Dut | 6 | tRA treated (DSB peaks) |
| 1-1355 | + | 756 | intron (NM_019639, intron 1 of 1) | Ubc | 2 | tRA treated (DSB peaks) |
| 1-119 | + | 584 | promoter-TSS (NM_198247) | Sertad4 | 2 | tRA treated (DSB peaks) |
| 1-1357 | + | 613 | Intergenic | Ubc | 8 | tRA treated (DSB peaks) |
| 1-1358 | + | 543 | Intergenic | Ubc | 5 | tRA treated (DSB peaks) |
| 1-1359 | + | 790 | Intergenic | Ubc | 4 | tRA treated (DSB peaks) |
| 1-1360 | + | 567 | Intergenic | Ubc | 6 | tRA treated (DSB peaks) |
| 1-1361 | + | 567 | Intergenic | Ubc | 5 | tRA treated (DSB peaks) |
| 1-1002 | + | 556 | intron (NM_007993, intron 2 of 65) | Fbn1 | 6 | tRA treated (DSB peaks) |

| | | | | | | |
|---|---|---|---|---|---|---|
| 1-914 | + | 589 | promoter-TSS (NM_001290370) | Nmt2 | 1 | tRA treated (DSB peaks) |
| 1-669 | + | 691 | promoter-TSS (NM_027446) | Eef2kmt | 1 | tRA treated (DSB peaks) |
| 1-222 | + | 616 | promoter-TSS (NM_001361942) | Pold2 | 1 | tRA treated (DSB peaks) |
| 1-1163 | + | 732 | promoter-TSS (NM_026534) | Ubxn2b | 1 | tRA treated (DSB peaks) |
| 1-1246 | + | 547 | TTS (NM_172145) | Eva1b | 8 | tRA treated (DSB peaks) |
| 1-816 | + | 575 | promoter-TSS (NM_177595) | Mkx | 1 | tRA treated (DSB peaks) |
| 1-1061 | + | 570 | promoter-TSS (NM_133218) | Zfp704 | 1 | tRA treated (DSB peaks) |
| 1-1605 | + | 541 | promoter-TSS (NM_001113518) | Arhgef7 | 1 | tRA treated (DSB peaks) |
| 1-735 | + | 546 | promoter-TSS (NM_175394) | Wtap | 1 | tRA treated (DSB peaks) |
| 1-1130 | + | 545 | Intergenic | Camk2d | 4 | tRA treated (DSB peaks) |
| 1-1686 | + | 621 | promoter-TSS (NM_001199484) | 4931406C07Rik | 1 | tRA treated (DSB peaks) |
| 1-1131 | + | 564 | 5' UTR (NM_001293665, exon 1 of 20) | Camk2d | 4 | tRA treated (DSB peaks) |
| 1-1674 | + | 544 | Intergenic | Irf2bp2 | 9 | tRA treated (DSB peaks) |
| 1-1675 | + | 706 | Intergenic | Irf2bp2 | 11 | tRA treated (DSB peaks) |
| 1-673 | + | 631 | promoter-TSS (NM_145479) | Klhl22 | 1 | tRA treated (DSB peaks) |
| 1-531 | + | 547 | promoter-TSS (NM_134081) | Dnajc9 | 1 | tRA treated (DSB peaks) |
| 1-532 | + | 584 | promoter-TSS (NM_010811) | Ndst2 | 1 | tRA treated (DSB peaks) |
| 1-1132 | + | 771 | intron (NM_001025438, intron 14 of 18) | Ank2 | 7 | tRA treated (DSB peaks) |
| 1-234 | + | 865 | promoter-TSS (NM_023324) | Peli1 | 1 | tRA treated (DSB peaks) |
| 1-431 | + | 541 | promoter-TSS (NM_183014) | Zfp184 | 1 | tRA treated (DSB peaks) |
| 1-1399 | + | 717 | promoter-TSS (NM_138587) | Fam3c | 1 | tRA treated (DSB peaks) |
| 1-1482 | + | 562 | Intergenic | Gm38404 | 6 | tRA treated (DSB peaks) |
| 1-535 | + | 557 | promoter-TSS (NM_001081247) | Polr3a | 1 | tRA treated (DSB peaks) |
| 1-1174 | + | 669 | promoter-TSS (NM_001355512) | Ufl1 | 1 | tRA treated (DSB peaks) |
| 1-537 | + | 595 | promoter-TSS (NM_133761) | Dcp1a | 1 | tRA treated (DSB peaks) |
| 1-1070 | + | 563 | promoter-TSS (NM_008857) | Prkci | 1 | tRA treated (DSB peaks) |
| 1-1676 | + | 1000 | Intergenic | Pard3 | 5 | tRA treated (DSB peaks) |
| 1-1007 | + | 668 | intron (NM_145532, intron 1 of 3) | Mall | 8 | tRA treated (DSB peaks) |
| 1-694 | + | 791 | promoter-TSS (NM_001252434) | Dlg1 | 1 | tRA treated (DSB peaks) |
| 1-1677 | + | 601 | Intergenic | Mir21c | 4 | tRA treated (DSB peaks) |
| 1-1678 | + | 678 | Intergenic | Mir21c | 10 | tRA treated (DSB peaks) |
| 1-212 | + | 823 | intron (NM_054078, intron 2 of 29) | Baz2a | 1 | tRA treated (DSB peaks) |
| 1-1176 | + | 572 | promoter-TSS (NM_172688) | Map3k7 | 1 | tRA treated (DSB peaks) |
| 1-214 | + | 581 | intron (NM_026259, intron 1 of 7) | Rnf41 | 2 | tRA treated (DSB peaks) |
| 25204 | + | 609 | intron (NM_145507, intron 1 of 3) | Dars | 1 | tRA treated (DSB peaks) |
| 1-761 | + | 765 | promoter-TSS (NM_009059) | Rgl2 | 1 | tRA treated (DSB peaks) |
| 1-1484 | + | 601 | Intergenic | Klrb1c | 6 | tRA treated (DSB peaks) |
| 1-1485 | + | 636 | Intergenic | Klrb1c | 5 | tRA treated (DSB peaks) |
| 1-455 | + | 614 | promoter-TSS (NM_009068) | Ripk1 | 1 | tRA treated (DSB peaks) |
| 1-217 | + | 657 | intron (NM_015740, intron 1 of 3) | Bloc1s1 | 7 | tRA treated (DSB peaks) |
| 1-218 | + | 657 | exon (NM_015740, exon 1 of 4) | Bloc1s1 | 7 | tRA treated (DSB peaks) |
| 1-825 | + | 723 | promoter-TSS (NM_025527) | Srp19 | 1 | tRA treated (DSB peaks) |
| 1-540 | + | 642 | promoter-TSS (NM_001301330) | Wapl | 1 | tRA treated (DSB peaks) |
| 1-1010 | + | 640 | exon (NM_009086, exon 1 of 15) | Polr1b | 1 | tRA treated (DSB peaks) |
| 1-1614 | + | 609 | promoter-TSS (NM_015802) | Dlc1 | 1 | tRA treated (DSB peaks) |
| 1-704 | + | 554 | promoter-TSS (NM_177093) | Lrrc58 | 1 | tRA treated (DSB peaks) |
| 1-831 | + | 601 | promoter-TSS (NM_020012) | Rnf14 | 1 | tRA treated (DSB peaks) |

| | | | | | | |
|---|---|---|---|---|---|---|
| 1-1587 | + | 651 | exon (NM_172255, exon 1 of 29) | Wdr11 | 1 | tRA treated (DSB peaks) |
| 1-606 | + | 596 | promoter-TSS (NM_001102458) | Azin1 | 1 | tRA treated (DSB peaks) |
| 1-900 | + | 556 | promoter-TSS (NM_009166) | Sorbs1 | 1 | tRA treated (DSB peaks) |
| 1-1250 | + | 541 | intron (NM_172702, intron 1 of 9) | Serinc2 | 8 | tRA treated (DSB peaks) |
| 1-1588 | + | 586 | intron (NM_001162855, intron 1 of 10) | Nsmce4a | 4 | tRA treated (DSB peaks) |
| 1-1615 | + | 613 | promoter-TSS (NM_019734) | Asah1 | 1 | tRA treated (DSB peaks) |
| 1-1589 | + | 544 | intron (NM_019564, intron 3 of 8) | Htra1 | 6 | tRA treated (DSB peaks) |
| 1-1590 | + | 544 | intron (NM_019564, intron 3 of 8) | Htra1 | 2 | tRA treated (DSB peaks) |
| 1-607 | + | 612 | promoter-TSS (NM_172815) | Rspo2 | 1 | tRA treated (DSB peaks) |
| 1-1516 | + | 550 | promoter-TSS (NM_025880) | 2410002F23Rik | 1 | tRA treated (DSB peaks) |
| 1-1012 | + | 868 | Intergenic | Rnf24 | 7 | tRA treated (DSB peaks) |
| 1-1135 | + | 591 | Intergenic | Papss1 | 1 | tRA treated (DSB peaks) |
| 1-1013 | + | 593 | intron (NM_013460, intron 1 of 1) | Adra1d | 6 | tRA treated (DSB peaks) |
| 1-1365 | + | 564 | intron (NM_177047, intron 5 of 19) | 4930563F08Rik | 6 | tRA treated (DSB peaks) |
| 1-1014 | + | 614 | intron (NM_175445, intron 4 of 10) | Rassf2 | 8 | tRA treated (DSB peaks) |
| 1-541 | + | 587 | promoter-TSS (NM_019425) | Gnpnat1 | 1 | tRA treated (DSB peaks) |
| 1-711 | + | 605 | promoter-TSS (NM_021497) | Nectin3 | 1 | tRA treated (DSB peaks) |
| 1-777 | + | 563 | promoter-TSS (NM_001161723) | Tfeb | 1 | tRA treated (DSB peaks) |
| 1-252 | + | 653 | promoter-TSS (NM_145377) | Trim41 | 1 | tRA treated (DSB peaks) |
| 1-1593 | + | 738 | Intergenic | Chst15 | 18 | tRA treated (DSB peaks) |
| 1-1252 | + | 599 | intron (NM_144907, intron 1 of 9) | Sesn2 | 8 | tRA treated (DSB peaks) |
| 1-548 | + | 597 | promoter-TSS (NM_027492) | Naa30 | 1 | tRA treated (DSB peaks) |
| 25934 | + | 691 | Intergenic | Mdm4 | 1 | tRA treated (DSB peaks) |
| 1-1366 | + | 736 | Intergenic | Auts2 | 6 | tRA treated (DSB peaks) |
| 1-260 | + | 608 | promoter-TSS (NM_001110150) | Mgat1 | 1 | tRA treated (DSB peaks) |
| 1-261 | + | 705 | promoter-TSS (NM_026543) | Mrnip | 1 | tRA treated (DSB peaks) |
| 1-1255 | + | 559 | intron (NM_001080819, intron 1 of 19) | Mir7227 | 11 | tRA treated (DSB peaks) |
| 1-468 | + | 684 | promoter-TSS (NM_175401) | Fbxw17 | 1 | tRA treated (DSB peaks) |
| 1-1257 | + | 590 | Intergenic | Arid1a | 1 | tRA treated (DSB peaks) |
| 1-1137 | + | 646 | Intergenic | Cxxc4 | 3 | tRA treated (DSB peaks) |
| 1-265 | + | 727 | promoter-TSS (NM_029976) | Cdkn2aipnl | 1 | tRA treated (DSB peaks) |
| 1-1258 | + | 624 | intron (NM_144527, intron 1 of 13) | Cep85 | 7 | tRA treated (DSB peaks) |
| 1-1367 | + | 695 | intron (NM_001359066, intron 1 of 32) | Gtf2i | 1 | tRA treated (DSB peaks) |
| 1-1594 | + | 540 | Intergenic | D7Ertd443e | 20 | tRA treated (DSB peaks) |
| 26299 | + | 724 | Intergenic | Kdm5b | 5 | tRA treated (DSB peaks) |
| 1-1315 | + | 677 | promoter-TSS (NM_001308144) | Zcchc4 | 1 | tRA treated (DSB peaks) |
| 1-778 | + | 559 | promoter-TSS (NM_028232) | Sgo1 | 1 | tRA treated (DSB peaks) |
| 1-556 | + | 541 | promoter-TSS (NM_145462) | Haus4 | 1 | tRA treated (DSB peaks) |
| 1-1018 | + | 567 | 5' UTR (NM_019677, exon 1 of 33) | Plcb1 | 1 | tRA treated (DSB peaks) |
| 1-1370 | + | 571 | exon (NM_009745, exon 1 of 6) | Bcl7b | 4 | tRA treated (DSB peaks) |
| 1-1371 | + | 571 | intron (NM_009745, intron 1 of 5) | Bcl7b | 8 | tRA treated (DSB peaks) |
| 1-473 | + | 672 | promoter-TSS (NR_027395) | Rnf44 | 1 | tRA treated (DSB peaks) |
| 1-474 | + | 671 | promoter-TSS (NR_027395) | Rnf44 | 1 | tRA treated (DSB peaks) |
| 43466 | + | 843 | promoter-TSS (NM_001302963) | Maip1 | 1 | tRA treated (DSB peaks) |
| 26665 | + | 544 | Intergenic | Gm4793 | 6 | tRA treated (DSB peaks) |
| 27030 | + | 544 | Intergenic | Gm4793 | 7 | tRA treated (DSB peaks) |
| 27395 | + | 580 | Intergenic | Phlda3 | 4 | tRA treated (DSB peaks) |

| | | | | | | |
|---|---|---|---|---|---|---|
| 1-480 | + | 646 | promoter-TSS (NM_029872) | Hnrnpa0 | 1 | tRA treated (DSB peaks) |
| 1-618 | + | 699 | promoter-TSS (NM_175212) | Tmem65 | 1 | tRA treated (DSB peaks) |
| 1-619 | + | 699 | promoter-TSS (NM_175212) | Tmem65 | 1 | tRA treated (DSB peaks) |
| 44562 | + | 559 | promoter-TSS (NM_173444) | Nbeal1 | 1 | tRA treated (DSB peaks) |
| 1-1140 | + | 557 | intron (NM_001033350, intron 7 of 16) | Bank1 | 5 | tRA treated (DSB peaks) |
| 1-1373 | + | 701 | intron (NM_001291238, intron 3 of 22) | Mir721 | 7 | tRA treated (DSB peaks) |
| 1-1374 | + | 603 | intron (NM_001291238, intron 3 of 22) | Mir721 | 7 | tRA treated (DSB peaks) |
| 1-1487 | + | 652 | intron (NM_019426, intron 1 of 14) | Atf7ip | 7 | tRA treated (DSB peaks) |
| 1-1488 | + | 616 | intron (NM_019426, intron 1 of 14) | Atf7ip | 7 | tRA treated (DSB peaks) |
| 1-1141 | + | 593 | intron (NM_008913, intron 1 of 13) | Ppp3ca | 15 | tRA treated (DSB peaks) |
| 1-1263 | + | 604 | intron (NM_010142, intron 1 of 15) | Ephb2 | 6 | tRA treated (DSB peaks) |
| 1-1019 | + | 548 | exon (NM_028201, exon 6 of 6) | Jag1 | 4 | tRA treated (DSB peaks) |
| 1-1020 | + | 694 | intron (NM_013822, intron 2 of 25) | Jag1 | 1 | tRA treated (DSB peaks) |
| 1-1595 | + | 549 | intron (NM_029886, intron 1 of 3) | 9430038I01Rik | 6 | tRA treated (DSB peaks) |
| 1-1489 | + | 566 | TTS (NM_001164401) | Eps8 | 3 | tRA treated (DSB peaks) |
| 1-279 | + | 846 | promoter-TSS (NM_019921) | Akap10 | 1 | tRA treated (DSB peaks) |
| 1-1199 | + | 756 | promoter-TSS (NM_001290521) | Wdr31 | 1 | tRA treated (DSB peaks) |
| 1-1490 | + | 541 | intron (NM_001271588, intron 1 of 20) | Eps8 | 2 | tRA treated (DSB peaks) |
| 1-282 | + | 667 | promoter-TSS (NM_001313984) | Ubb | 1 | tRA treated (DSB peaks) |
| 1-156 | + | 617 | promoter-TSS (NM_053183) | Ddx50 | 1 | tRA treated (DSB peaks) |
| 1-787 | + | 953 | promoter-TSS (NM_144859) | Pja2 | 1 | tRA treated (DSB peaks) |
| 1-382 | + | 561 | promoter-TSS (NM_172578) | Mis18bp1 | 1 | tRA treated (DSB peaks) |
| 1-1087 | + | 637 | promoter-TSS (NM_001355432) | Ccnl1 | 1 | tRA treated (DSB peaks) |
| 1-1622 | + | 546 | promoter-TSS (NM_025465) | Tma16 | 1 | tRA treated (DSB peaks) |
| 1-1142 | + | 553 | intron (NR_121195, intron 1 of 7) | Tspan5 | 6 | tRA treated (DSB peaks) |
| 1-1143 | + | 553 | intron (NR_121195, intron 1 of 7) | Tspan5 | 11 | tRA treated (DSB peaks) |
| 1-1750 | + | 751 | promoter-TSS (NM_016787) | Bnip2 | 1 | tRA treated (DSB peaks) |
| 10959 | + | 697 | promoter-TSS (NM_026195) | Atic | 1 | tRA treated (DSB peaks) |
| 1-1596 | + | 597 | TTS (NM_001347613) | 6430531B16Rik | 3 | tRA treated (DSB peaks) |
| 1-723 | + | 544 | promoter-TSS (NM_019413) | Robo1 | 1 | tRA treated (DSB peaks) |
| 1-850 | + | 618 | promoter-TSS (NM_008540) | 4930578L24Rik | 1 | tRA treated (DSB peaks) |
| 1-301 | + | 852 | promoter-TSS (NM_001081158) | Cluh | 1 | tRA treated (DSB peaks) |
| 1-851 | + | 551 | promoter-TSS (NM_001042660) | Gm20544 | 1 | tRA treated (DSB peaks) |
| 1-1265 | + | 742 | Intergenic | Rcc2 | 9 | tRA treated (DSB peaks) |
| 1-306 | + | 781 | promoter-TSS (NM_138950) | Wdr81 | 1 | tRA treated (DSB peaks) |
| 12420 | + | 1000 | promoter-TSS (NM_001081210) | Mrpl44 | 1 | tRA treated (DSB peaks) |
| 1-801 | + | 597 | promoter-TSS (NM_001286647) | Atl2 | 1 | tRA treated (DSB peaks) |
| 1-1267 | + | 638 | TTS (NM_013868) | Hspb7 | 8 | tRA treated (DSB peaks) |
| 1-167 | + | 584 | promoter-TSS (NM_007705) | Cirbp | 1 | tRA treated (DSB peaks) |
| 1-1268 | + | 684 | intron (NM_010329, intron 1 of 5) | Pdpn | 5 | tRA treated (DSB peaks) |
| 1-1385 | + | 682 | intron (NM_011182, intron 1 of 12) | Cyth3 | 10 | tRA treated (DSB peaks) |
| 1-1386 | + | 682 | intron (NM_011182, intron 1 of 12) | Cyth3 | 7 | tRA treated (DSB peaks) |
| 1-1601 | + | 631 | intron (NR_131763, intron 1 of 5) | Tnfrsf23 | 10 | tRA treated (DSB peaks) |
| 1-314 | + | 730 | promoter-TSS (NM_027472) | 5730455P16Rik | 1 | tRA treated (DSB peaks) |
| 1-1756 | + | 574 | promoter-TSS (NM_172507) | Sh3bgrl2 | 1 | tRA treated (DSB peaks) |
| 1-1021 | + | 857 | Intergenic | Gm5535 | 4 | tRA treated (DSB peaks) |
| 1-1022 | + | 799 | Intergenic | Gm5535 | 7 | tRA treated (DSB peaks) |

| | | | | | | |
|---|---|---|---|---|---|---|
| 1-401 | + | 795 | promoter-TSS (NM_001289432) | Cipc | 1 | tRA treated (DSB peaks) |
| 1-1442 | + | 658 | promoter-TSS (NM_013528) | Gfpt1 | 1 | tRA treated (DSB peaks) |
| 1-1603 | + | 591 | Intergenic | Ccnd1 | 5 | tRA treated (DSB peaks) |
| 1-1269 | + | 746 | Intergenic | Dhrs3 | 5 | tRA treated (DSB peaks) |
| 1-1270 | + | 600 | Intergenic | Dhrs3 | 3 | tRA treated (DSB peaks) |
| 1-1271 | + | 562 | intron (NM_001276465, intron 67 of 69) | Dhrs3 | 4 | tRA treated (DSB peaks) |
| 1-1272 | + | 1000 | intron (NM_001276465, intron 67 of 69) | Dhrs3 | 2 | tRA treated (DSB peaks) |
| 1-1273 | + | 935 | intron (NM_001276465, intron 67 of 69) | Dhrs3 | 3 | tRA treated (DSB peaks) |
| 18264 | + | 581 | promoter-TSS (NM_133805) | Cops8 | 1 | tRA treated (DSB peaks) |
| 1-1274 | + | 691 | intron (NM_011610, intron 1 of 9) | Tnfrsf1b | 4 | tRA treated (DSB peaks) |
| 1-1648 | + | 573 | promoter-TSS (NM_001282000) | Rbl2 | 1 | tRA treated (DSB peaks) |
| 1-1148 | + | 750 | exon (NM_026993, exon 1 of 6) | Ddah1 | 2 | tRA treated (DSB peaks) |
| 1-1023 | + | 625 | Intergenic | Rin2 | 7 | tRA treated (DSB peaks) |
| 1-1024 | + | 625 | Intergenic | Rin2 | 5 | tRA treated (DSB peaks) |
| 1-1491 | + | 562 | Intergenic | Rassf8 | 5 | tRA treated (DSB peaks) |
| 1-1149 | + | 575 | intron (NM_026993, intron 4 of 5) | Bcl10 | 12 | tRA treated (DSB peaks) |
| 1-1025 | + | 554 | TTS (NM_028724) | Naa20 | 11 | tRA treated (DSB peaks) |
| 1-977 | + | 626 | promoter-TSS (NM_026944) | Alkbh3 | 1 | tRA treated (DSB peaks) |
| 1-329 | + | 836 | promoter-TSS (NM_001013381) | Rsad1 | 1 | tRA treated (DSB peaks) |
| 1-330 | + | 835 | promoter-TSS (NM_001013381) | Rsad1 | 1 | tRA treated (DSB peaks) |
| 1-1152 | + | 665 | exon (NM_001166064, exon 1 of 7) | Syde2 | 2 | tRA treated (DSB peaks) |
| 1-335 | + | 548 | promoter-TSS (NM_172300) | Ube2z | 1 | tRA treated (DSB peaks) |
| 1-1026 | + | 566 | intron (NM_001033298, intron 12 of 13) | Xrn2 | 2 | tRA treated (DSB peaks) |
| 1-1108 | + | 634 | promoter-TSS (NM_001162388) | Pex11b | 1 | tRA treated (DSB peaks) |
| 1-1153 | + | 807 | Intergenic | Ttll7 | 10 | tRA treated (DSB peaks) |
| 1-1154 | + | 924 | Intergenic | Ttll7 | 3 | tRA treated (DSB peaks) |
| 1-1275 | + | 569 | 5' UTR (NM_027873, exon 1 of 2) | Ubiad1 | 6 | tRA treated (DSB peaks) |
| 1-1545 | + | 614 | promoter-TSS (NM_001168539) | Tsku | 1 | tRA treated (DSB peaks) |
| 1-403 | + | 605 | promoter-TSS (NM_001313934) | Calm1 | 1 | tRA treated (DSB peaks) |
| 1-663 | + | 676 | promoter-TSS (NM_001174073) | Pcbp2 | 1 | tRA treated (DSB peaks) |
| 1-407 | + | 559 | promoter-TSS (NM_178613) | Atg2b | 1 | tRA treated (DSB peaks) |
| 1-1112 | + | 583 | promoter-TSS (NM_017397) | Ddx20 | 1 | tRA treated (DSB peaks) |
| 1-1555 | + | 762 | promoter-TSS (NM_025301) | Mrpl17 | 1 | tRA treated (DSB peaks) |
| 1-1278 | + | 662 | intron (NM_173371, intron 1 of 4) | H6pd | 1 | tRA treated (DSB peaks) |
| 1-1279 | + | 662 | 5' UTR (NM_173371, exon 1 of 5) | H6pd | 2 | tRA treated (DSB peaks) |
| 1-1155 | + | 611 | Intergenic | Mir3963 | 6 | tRA treated (DSB peaks) |
| 1-1156 | + | 611 | Intergenic | Mir3963 | 9 | tRA treated (DSB peaks) |
| 1-1113 | + | 1000 | promoter-TSS (NM_026013) | Cept1 | 1 | tRA treated (DSB peaks) |
| 1-1335 | + | 894 | promoter-TSS (NM_011578) | Tgfbr3 | 1 | tRA treated (DSB peaks) |
| 1-1395 | + | 712 | intron (NR_045700, intron 1 of 3) | 1700028E10Rik | 9 | tRA treated (DSB peaks) |
| 1-1280 | + | 614 | intron (NM_001081557, intron 5 of 23) | Gm13090 | 11 | tRA treated (DSB peaks) |
| 1-1336 | + | 551 | promoter-TSS (NM_011578) | Tgfbr3 | 1 | tRA treated (DSB peaks) |
| 1-1281 | + | 567 | exon (NM_178406, exon 6 of 6) | Hes3 | 3 | tRA treated (DSB peaks) |
| 1-1337 | + | 709 | promoter-TSS (NM_001255991) | Btbd8 | 1 | tRA treated (DSB peaks) |
| 28126 | + | 794 | intron (NM_016846, intron 5 of 17) | Rgl1 | 7 | tRA treated (DSB peaks) |
| 1-1117 | + | 572 | promoter-TSS (NM_026198) | Tmem167b | 1 | tRA treated (DSB peaks) |
| 28491 | + | 553 | intron (NM_008485, intron 19 of 23) | Lamc2 | 1 | tRA treated (DSB peaks) |

| | | | | | | |
|---|---|---|---|---|---|---|
| 28856 | + | 554 | intron (NM_008485, intron 19 of 23) | Lamc2 | 5 | tRA treated (DSB peaks) |
| 1-1119 | + | 550 | promoter-TSS (NM_172685) | Slc25a24 | 1 | tRA treated (DSB peaks) |
| 29587 | + | 821 | Intergenic | Npl | 10 | tRA treated (DSB peaks) |
| 1-1157 | + | 674 | 5' UTR (NM_007382, exon 1 of 12) | Acadm | 1 | tRA treated (DSB peaks) |
| 1-1224 | + | 606 | promoter-TSS (NM_001301368) | Cdkn2c | 1 | tRA treated (DSB peaks) |
| 1-1658 | + | 1000 | promoter-TSS (NM_009677) | Ap1g1 | 1 | tRA treated (DSB peaks) |
| 1-1779 | + | 733 | promoter-TSS (NM_010628) | Klhl18 | 1 | tRA treated (DSB peaks) |
| 1-1158 | + | 758 | Intergenic | Erich3 | 5 | tRA treated (DSB peaks) |
| 29952 | + | 608 | Intergenic | Gm9530 | 10 | tRA treated (DSB peaks) |
| 30317 | + | 608 | Intergenic | Gm9530 | 8 | tRA treated (DSB peaks) |
| 30682 | + | 587 | Intergenic | Ier5 | 9 | tRA treated (DSB peaks) |
| 31048 | + | 640 | Intergenic | Ier5 | 2 | tRA treated (DSB peaks) |
| 22282 | + | 768 | promoter-TSS (NM_001081316) | Dsel | 1 | tRA treated (DSB peaks) |
| 1-1284 | + | 814 | 5' UTR (NM_025570, exon 2 of 5) | Mrpl20 | 3 | tRA treated (DSB peaks) |
| 1-1459 | + | 817 | promoter-TSS (NR_027008) | Gt(ROSA)26Sor | 1 | tRA treated (DSB peaks) |
| 1-1122 | + | 848 | promoter-TSS (NM_026043) | Rnpc3 | 1 | tRA treated (DSB peaks) |
| 1-364 | + | 629 | promoter-TSS (NM_011358) | Srsf2 | 1 | tRA treated (DSB peaks) |
| 1-1232 | + | 555 | promoter-TSS (NM_001161774) | St3gal3 | 1 | tRA treated (DSB peaks) |
| 1-1668 | + | 571 | promoter-TSS (NM_009569) | Zfpm1 | 1 | tRA treated (DSB peaks) |
| 1-1249 | + | 679 | promoter-TSS (NM_030252) | Smim12 | 1 | tRA treated (DSB peaks) |
| 1-216 | + | 597 | promoter-TSS (NM_007653) | Cd63 | 1 | tRA treated (DSB peaks) |
| 1-1009 | + | 640 | promoter-TSS (NM_009086) | Polr1b | 1 | tRA treated (DSB peaks) |
| 1-1586 | + | 651 | promoter-TSS (NM_172255) | Wdr11 | 1 | tRA treated (DSB peaks) |
| 25569 | + | 902 | promoter-TSS (NM_172516) | Dstyk | 1 | tRA treated (DSB peaks) |
| 1-1037 | + | 792 | intron (NM_027434, intron 6 of 6) | 2010009K17Rik | 8 | tRA treated (DSB peaks) |
| 1-1256 | + | 589 | promoter-TSS (NM_001080819) | Arid1a | 1 | tRA treated (DSB peaks) |
| 32143 | + | 574 | intron (NM_001085376, intron 20 of 21) | Pappa2 | 5 | tRA treated (DSB peaks) |
| 1-1160 | + | 586 | intron (NM_001356349, intron 2 of 12) | Wls | 6 | tRA treated (DSB peaks) |
| 1-1486 | + | 622 | promoter-TSS (NM_007961) | Etv6 | 1 | tRA treated (DSB peaks) |
| 1-1139 | + | 623 | promoter-TSS (NM_025356) | Ube2d3 | 1 | tRA treated (DSB peaks) |
| 1-1372 | + | 630 | promoter-TSS (NM_011293) | Polr2j | 1 | tRA treated (DSB peaks) |
| 1-1039 | + | 615 | intron (NM_011521, intron 1 of 4) | Sdc4 | 2 | tRA treated (DSB peaks) |
| 1-1381 | + | 587 | promoter-TSS (NM_030565) | Fam20c | 1 | tRA treated (DSB peaks) |
| 1-1266 | + | 648 | promoter-TSS (NM_001145958) | Crocc | 1 | tRA treated (DSB peaks) |
| 1-1384 | + | 625 | promoter-TSS (NM_177879) | Sdk1 | 1 | tRA treated (DSB peaks) |
| 33604 | + | 607 | Intergenic | Mpzl1 | 2 | tRA treated (DSB peaks) |
| 1-1043 | + | 644 | Intergenic | Zmynd8 | 2 | tRA treated (DSB peaks) |
| 1-1044 | + | 603 | Intergenic | 5031425F14Rik | 5 | tRA treated (DSB peaks) |
| 1-1045 | + | 602 | Intergenic | 5031425F14Rik | 6 | tRA treated (DSB peaks) |
| 1-1046 | + | 764 | Intergenic | 5031425F14Rik | 9 | tRA treated (DSB peaks) |
| 1-1144 | + | 568 | promoter-TSS (NM_001190855) | Pdlim5 | 1 | tRA treated (DSB peaks) |
| 1-1048 | + | 563 | Intergenic | A530013C23Rik | 3 | tRA treated (DSB peaks) |
| 33970 | + | 698 | Intergenic | Mir6354 | 7 | tRA treated (DSB peaks) |
| 34335 | + | 599 | Intergenic | Mir6354 | 6 | tRA treated (DSB peaks) |
| 1-1602 | + | 803 | promoter-TSS (NM_001199227) | Osbpl5 | 1 | tRA treated (DSB peaks) |
| 35065 | + | 620 | Intergenic | Ddr2 | 13 | tRA treated (DSB peaks) |
| 35431 | + | 620 | Intergenic | Ddr2 | 7 | tRA treated (DSB peaks) |

| | | | | | | |
|---|---|---|---|---|---|---|
| 1-1049 | + | 757 | Intergenic | Bcas1os2 | 1 | tRA treated (DSB peaks) |
| 35796 | + | 811 | Intergenic | Fcgr4 | 4 | tRA treated (DSB peaks) |
| 36161 | + | 773 | Intergenic | Cfap126 | 10 | tRA treated (DSB peaks) |
| 1-100 | + | 773 | Intergenic | Cfap126 | 8 | tRA treated (DSB peaks) |
| 1-1151 | + | 555 | promoter-TSS (NM_001356284) | 2410004B18Rik | 1 | tRA treated (DSB peaks) |
| 1-102 | + | 666 | intron (NM_001329871, intron 1 of 3) | Pea15a | 6 | tRA treated (DSB peaks) |
| 1-103 | + | 666 | 5' UTR (NM_001329869, exon 1 of 4) | Pea15a | 4 | tRA treated (DSB peaks) |
| 1-1050 | + | 545 | Intergenic | Pmepa1 | 2 | tRA treated (DSB peaks) |
| 1-1051 | + | 685 | Intergenic | Pmepa1 | 1 | tRA treated (DSB peaks) |
| 1-1052 | + | 685 | Intergenic | Pmepa1 | 1 | tRA treated (DSB peaks) |
| 1-1028 | + | 587 | promoter-TSS (NM_029688) | Srxn1 | 1 | tRA treated (DSB peaks) |
| 1-1054 | + | 545 | intron (NM_213733, intron 1 of 11) | Npepl1 | 1 | tRA treated (DSB peaks) |
| 1-1055 | + | 592 | Intergenic | Gm14393 | 4 | tRA treated (DSB peaks) |
| 1-104 | + | 718 | Intergenic | 2310043L19Rik | 4 | tRA treated (DSB peaks) |
| 1-105 | + | 759 | Intergenic | 1700016C15Rik | 3 | tRA treated (DSB peaks) |
| 1-106 | + | 759 | Intergenic | 1700016C15Rik | 4 | tRA treated (DSB peaks) |
| 1-107 | + | 701 | intron (NM_001081175, intron 2 of 7) | Gm5069 | 4 | tRA treated (DSB peaks) |
| 1-108 | + | 700 | intron (NM_001081175, intron 2 of 7) | Gm5069 | 6 | tRA treated (DSB peaks) |
| 1-1029 | + | 608 | promoter-TSS (NM_001083921) | Rbck1 | 1 | tRA treated (DSB peaks) |
| 1-1283 | + | 826 | promoter-TSS (NM_007859) | Dffb | 1 | tRA treated (DSB peaks) |
| 1-110 | + | 770 | intron (NM_012058, intron 1 of 2) | Srp9 | 7 | tRA treated (DSB peaks) |
| 1-111 | + | 697 | Intergenic | Tlr5 | 2 | tRA treated (DSB peaks) |
| 1-1031 | + | 687 | promoter-TSS (NM_199304) | Zfp341 | 1 | tRA treated (DSB peaks) |
| 1-113 | + | 564 | Intergenic | Esrrg | 5 | tRA treated (DSB peaks) |
| 31778 | + | 634 | promoter-TSS (NM_001160018) | Tor1aip1 | 1 | tRA treated (DSB peaks) |
| 1-1035 | + | 660 | promoter-TSS (NM_001139516) | Rbl1 | 1 | tRA treated (DSB peaks) |
| 1-1036 | + | 556 | promoter-TSS (NM_001355165) | Rpn2 | 1 | tRA treated (DSB peaks) |
| 1-1047 | + | 1000 | promoter-TSS (NM_023815) | Trp53rkb | 1 | tRA treated (DSB peaks) |
| 1-1053 | + | 1000 | promoter-TSS (NM_172675) | Stx16 | 1 | tRA treated (DSB peaks) |
| 1-117 | + | 562 | promoter-TSS (NM_025424) | Nenf | 1 | tRA treated (DSB peaks) |
