## Supplementary Table S6 for "ERCC1-XPF Interacts with Topoisomerase IIβ to Facilitate the Repair of Activity-induced DNA Breaks"

### Supplementary Table S6. RNA-Seq and BLISS data profiles of bXPF-bound genes in untreated and tRA-treated MEFs

| Gene name | (0=no XPF recruitment, 1=XPF recruited, IDR-filtered ChIPseq peaks) | DSBs (average DSB count per gene) | mRNA levels (average DESeq normalized counts) | Treatment (0=untreated, 1=tRA-treated, 10mM, 16h) |
|---|---|---|---|---|
| 0610043K17Rik | 0 | 10.5 | 3 | 0 |
| 0610043K17Rik | 0 | 6.5 | 13.6667 | 1 |
| 1010001N08Rik | 0 | 110.5 | 9.66666667 | 0 |
| 1010001N08Rik | 0 | 142 | 26.6666667 | 1 |
| 1600010M07Rik | 0 | 25.5 | 159.333333 | 0 |
| 1600010M07Rik | 0 | 18.5 | 278.666667 | 1 |
| 1700056E22Rik | 0 | 2.5 | 41 | 0 |
| 1700056E22Rik | 0 | 1 | 74 | 1 |
| 2310002F09Rik | 0 | 19 | 61.6666667 | 0 |
| 2310002F09Rik | 0 | 25.5 | 117 | 1 |
| 2900026A02Rik | 0 | 132 | 4813.33333 | 0 |
| 2900026A02Rik | 0 | 117.5 | 1969.66667 | 1 |
| 3300005D01Rik | 0 | 240.5 | 479.666667 | 0 |
| 3300005D01Rik | 0 | 232 | 170 | 1 |
| 4833422C13Rik | 0 | 129 | 63.3333333 | 0 |
| 4833422C13Rik | 0 | 86 | 26.6666667 | 1 |
| 4930447F24Rik | 0 | 104.5 | 28.3333333 | 0 |
| 4930447F24Rik | 0 | 76 | 9.66666667 | 1 |
| 4930556M19Rik | 0 | 109.5 | 66 | 0 |
| 4930556M19Rik | 1 | 93.5 | 108 | 1 |
| 4931406P16Rik | 0 | 79.5 | 3506.33333 | 0 |
| 4931406P16Rik | 0 | 88.5 | 6060.33333 | 1 |
| 5830408C22Rik | 0 | 12 | 17.3333333 | 0 |
| 5830408C22Rik | 0 | 15.5 | 50 | 1 |
| 9030624G23Rik | 0 | 6 | 100.666667 | 0 |
| 9030624G23Rik | 0 | 3 | 167 | 1 |
| 9330188P03Rik | 0 | 14 | 58 | 0 |
| 9330188P03Rik | 0 | 12.5 | 21.3333333 | 1 |
| 9430078K24Rik | 0 | 473.5 | 18.6666667 | 0 |
| 9430078K24Rik | 0 | 507.5 | 2.00003333 | 1 |
| A2m | 0 | 58.5 | 10.6666667 | 0 |
| A2m | 0 | 71.5 | 27.0000333 | 1 |
| A4galt | 0 | 71 | 154.333333 | 0 |
| A4galt | 0 | 67.5 | 77.3333333 | 1 |
| A530016L24Rik | 0 | 24.5 | 18 | 0 |
| A530016L24Rik | 0 | 17 | 46 | 1 |
| A630072M18Rik | 0 | 75 | 105.666667 | 0 |
| A630072M18Rik | 0 | 83.5 | 165.666667 | 1 |
| A730056A06Rik | 0 | 295 | 20 | 0 |
| A730056A06Rik | 0 | 400 | 48.3333333 | 1 |
| A830029E22Rik | 0 | 5.5 | 8.66666667 | 0 |

| | | | | |
|---|---|---|---|---|
| A830029E22Rik | 0 | 7 | 0.0001 | 1 |
| Abca1 | 0 | 150.5 | 1215.66667 | 0 |
| Abca1 | 1 | 177.5 | 2823 | 1 |
| Abca6 | 0 | 158.5 | 22.3333333 | 0 |
| Abca6 | 0 | 172 | 49.3333333 | 1 |
| Abca9 | 0 | 123.5 | 111.333333 | 0 |
| Abca9 | 0 | 182.5 | 175.666667 | 1 |
| Abcb1b | 0 | 99.5 | 888.666667 | 0 |
| Abcb1b | 0 | 95.5 | 638.333333 | 1 |
| Abcc10 | 0 | 36 | 407.666667 | 0 |
| Abcc10 | 0 | 40 | 729 | 1 |
| Abcc3 | 0 | 120.5 | 173.333333 | 0 |
| Abcc3 | 0 | 122.5 | 289.333333 | 1 |
| Abcd2 | 0 | 582 | 38 | 0 |
| Abcd2 | 0 | 565.5 | 69.3333333 | 1 |
| Abcg1 | 0 | 74.5 | 161 | 0 |
| Abcg1 | 0 | 75.5 | 301.666667 | 1 |
| Abcg2 | 0 | 188 | 1878.33333 | 0 |
| Abcg2 | 0 | 205 | 3132.33333 | 1 |
| Abi1 | 0 | 359 | 3248 | 0 |
| Abi1 | 0 | 405 | 4566.33333 | 1 |
| Abi3bp | 0 | 438 | 1705.66667 | 0 |
| Abi3bp | 0 | 486 | 1073.66667 | 1 |
| Ablim1 | 0 | 644.5 | 1398.66667 | 0 |
| Ablim1 | 0 | 489 | 2856.33333 | 1 |
| Acan | 0 | 152 | 1775.33333 | 0 |
| Acan | 0 | 106.5 | 660.333333 | 1 |
| Ackr4 | 0 | 157 | 52.6666667 | 0 |
| Ackr4 | 0 | 160.5 | 22 | 1 |
| Acot11 | 0 | 74 | 447.666667 | 0 |
| Acot11 | 0 | 62.5 | 185.666667 | 1 |
| Acp5 | 0 | 26.5 | 73.3333333 | 0 |
| Acp5 | 0 | 23 | 252 | 1 |
| Acp7 | 0 | 44.5 | 13.6666667 | 0 |
| Acp7 | 0 | 36.5 | 136.666667 | 1 |
| Acsbg1 | 0 | 79 | 78.6666667 | 0 |
| Acsbg1 | 0 | 82 | 141.666667 | 1 |
| Acsl4 | 0 | 141.5 | 3248.33333 | 0 |
| Acsl4 | 0 | 209 | 5447.33333 | 1 |
| Actg2 | 0 | 45.5 | 15640.6667 | 0 |
| Actg2 | 0 | 61.5 | 11420.3333 | 1 |
| Acvr2a | 0 | 141.5 | 2864 | 0 |
| Acvr2a | 0 | 137 | 2063.66667 | 1 |
| Acvrl1 | 0 | 57 | 1114.66667 | 0 |
| Acvrl1 | 0 | 53.5 | 733.333333 | 1 |
| Adam19 | 1 | 240.5 | 17388.3333 | 0 |
| Adam19 | 0 | 286.5 | 25636.3333 | 1 |

| | | | | |
|---|---|---|---|---|
| Adam22 | 0 | 365 | 98 | 0 |
| Adam22 | 0 | 342 | 189 | 1 |
| Adam23 | 0 | 294 | 2583.66667 | 0 |
| Adam23 | 0 | 351.5 | 1864.66667 | 1 |
| Adam33 | 0 | 47.5 | 1390.66667 | 0 |
| Adam33 | 0 | 36 | 975.333333 | 1 |
| Adamts14 | 0 | 108 | 1097 | 0 |
| Adamts14 | 0 | 87 | 2345 | 1 |
| Adamts16 | 0 | 315.5 | 50.3333333 | 0 |
| Adamts16 | 0 | 478 | 10.6666667 | 1 |
| Adamts19 | 0 | 182.5 | 1.6667 | 0 |
| Adamts19 | 0 | 192.5 | 19.3333333 | 1 |
| Adamts4 | 0 | 44 | 1090.66667 | 0 |
| Adamts4 | 0 | 35.5 | 646.666667 | 1 |
| Adamts6 | 0 | 456.5 | 2711.33333 | 0 |
| Adamts6 | 0 | 858 | 3709.33333 | 1 |
| Adamts8 | 0 | 69.5 | 13 | 0 |
| Adamts8 | 0 | 46.5 | 108.333333 | 1 |
| Adamts9 | 0 | 204 | 2848 | 0 |
| Adamts9 | 1 | 240.5 | 4123.33333 | 1 |
| Adamtsl1 | 0 | 503.5 | 1360.66667 | 0 |
| Adamtsl1 | 0 | 595 | 3876.66667 | 1 |
| Adamtsl2 | 0 | 70 | 72.6666667 | 0 |
| Adamtsl2 | 0 | 65 | 31.3333333 | 1 |
| Adamtsl3 | 0 | 465 | 1718.33333 | 0 |
| Adamtsl3 | 0 | 412.5 | 525.666667 | 1 |
| Adamtsl4 | 0 | 41 | 1941.66667 | 0 |
| Adamtsl4 | 0 | 38.5 | 796.666667 | 1 |
| Adarb1 | 0 | 358 | 1037.33333 | 0 |
| Adarb1 | 0 | 424 | 1614.33333 | 1 |
| Adcy4 | 0 | 46 | 361 | 0 |
| Adcy4 | 0 | 36.5 | 237 | 1 |
| Adcyap1r1 | 0 | 105 | 1064 | 0 |
| Adcyap1r1 | 0 | 121.5 | 355 | 1 |
| Add3 | 0 | 168 | 1226 | 0 |
| Add3 | 0 | 159 | 2142.66667 | 1 |
| Adgra2 | 0 | 70.5 | 15749.3333 | 0 |
| Adgra2 | 0 | 66 | 33768.3333 | 1 |
| Adgra3 | 0 | 216 | 2386.33333 | 0 |
| Adgra3 | 0 | 250 | 3272.66667 | 1 |
| Adgrf5 | 0 | 152 | 116.666667 | 0 |
| Adgrf5 | 0 | 210 | 217.666667 | 1 |
| Adgrg1 | 0 | 56.5 | 127 | 0 |
| Adgrg1 | 0 | 45 | 77.3333333 | 1 |
| Adgrg2 | 0 | 214 | 436.333333 | 0 |
| Adgrg2 | 0 | 192 | 232.666667 | 1 |
| Adgrg5 | 0 | 38.5 | 72 | 0 |

| | | | | |
|---|---|---|---|---|
| Adgrg5 | 0 | 49.5 | 154.333333 | 1 |
| Adgrg6 | 0 | 300 | 1828.33333 | 0 |
| Adgrg6 | 0 | 317 | 2939 | 1 |
| Adm | 1 | 22 | 3744.66667 | 0 |
| Adm | 0 | 25 | 8014.66667 | 1 |
| Adora1 | 0 | 49.5 | 46 | 0 |
| Adora1 | 0 | 44.5 | 7 | 1 |
| Adora2a | 0 | 138.5 | 22.3333333 | 0 |
| Adora2a | 0 | 123 | 47.6666667 | 1 |
| Adora2b | 0 | 155 | 725.333333 | 0 |
| Adora2b | 0 | 199 | 1745.66667 | 1 |
| Adra1b | 0 | 263.5 | 316 | 0 |
| Adra1b | 0 | 317 | 477.666667 | 1 |
| Adrb1 | 0 | 4 | 162.333333 | 0 |
| Adrb1 | 0 | 1.5 | 256.333333 | 1 |
| Adrb2 | 0 | 5 | 161.333333 | 0 |
| Adrb2 | 0 | 3.5 | 105.333333 | 1 |
| Afap1l2 | 0 | 437 | 645.333333 | 0 |
| Afap1l2 | 0 | 477 | 3080.66667 | 1 |
| Agtr1a | 0 | 94.5 | 74.3333333 | 0 |
| Agtr1a | 0 | 115.5 | 166.666667 | 1 |
| Agtr2 | 0 | 9.5 | 562.666667 | 0 |
| Agtr2 | 0 | 11.5 | 143 | 1 |
| Ahr | 0 | 145 | 859 | 0 |
| Ahr | 0 | 128.5 | 510.333333 | 1 |
| Ahrr | 0 | 106.5 | 348.666667 | 0 |
| Ahrr | 0 | 166.5 | 155.333333 | 1 |
| AI427809 | 0 | 23 | 38.3333333 | 0 |
| AI427809 | 0 | 19.5 | 87.3333333 | 1 |
| Aif1 | 0 | 24.5 | 57.6666667 | 0 |
| Aif1 | 0 | 35 | 25 | 1 |
| Aifm2 | 0 | 77.5 | 440.333333 | 0 |
| Aifm2 | 0 | 58.5 | 1093.66667 | 1 |
| Ajap1 | 0 | 165 | 0.0001 | 0 |
| Ajap1 | 0 | 138 | 15.3333333 | 1 |
| Ak5 | 0 | 54 | 2047.66667 | 0 |
| Ak5 | 0 | 23 | 1495 | 1 |
| Akap7 | 1 | 257 | 255 | 0 |
| Akap7 | 1 | 284.5 | 420.333333 | 1 |
| Akr1b3 | 0 | 4.5 | 3127.66667 | 0 |
| Akr1b3 | 1 | 2.5 | 5367.33333 | 1 |
| Aldh1a3 | 0 | 42 | 558.666667 | 0 |
| Aldh1a3 | 0 | 25.5 | 94.3333333 | 1 |
| Alkal1 | 0 | 75.5 | 119.666667 | 0 |
| Alkal1 | 0 | 98 | 47 | 1 |
| Alpl | 0 | 101.5 | 29.6666667 | 0 |
| Alpl | 0 | 85.5 | 150.333333 | 1 |

| | | | | |
|---|---|---|---|---|
| Als2cl | 0 | 34.5 | 979.666667 | 0 |
| Als2cl | 0 | 28.5 | 614 | 1 |
| Alx3 | 0 | 34 | 155.666667 | 0 |
| Alx3 | 0 | 27.5 | 98.6666667 | 1 |
| Amhr2 | 0 | 37.5 | 13 | 0 |
| Amhr2 | 0 | 56.5 | 2.6667 | 1 |
| Ammecr1 | 0 | 192 | 2736.33333 | 0 |
| Ammecr1 | 0 | 257.5 | 4423.33333 | 1 |
| Amotl1 | 0 | 145 | 9275.66667 | 0 |
| Amotl1 | 1 | 161.5 | 21407.6667 | 1 |
| Angpt2 | 0 | 258 | 390 | 0 |
| Angpt2 | 0 | 288 | 821.333333 | 1 |
| Angpt4 | 0 | 60 | 58 | 0 |
| Angpt4 | 0 | 57.5 | 221.333333 | 1 |
| Angptl1 | 0 | 137 | 133.666667 | 0 |
| Angptl1 | 0 | 152 | 69.3333333 | 1 |
| Angptl2 | 0 | 288.5 | 11023 | 0 |
| Angptl2 | 0 | 250.5 | 5155 | 1 |
| Angptl4 | 0 | 22.5 | 322 | 0 |
| Angptl4 | 0 | 20.5 | 1021 | 1 |
| Angptl7 | 0 | 137 | 737.666667 | 0 |
| Angptl7 | 0 | 142.5 | 186.333333 | 1 |
| Ank | 0 | 108 | 2431.66667 | 0 |
| Ank | 0 | 92 | 1616.66667 | 1 |
| Ank1 | 0 | 738.5 | 393.333333 | 0 |
| Ank1 | 0 | 715 | 216.666667 | 1 |
| Ank3 | 0 | 1834.5 | 1069.66667 | 0 |
| Ank3 | 0 | 2171 | 596.666667 | 1 |
| Ankk1 | 0 | 102 | 65.6666667 | 0 |
| Ankk1 | 0 | 99 | 149.333333 | 1 |
| Ankrd13b | 0 | 37.5 | 1323.33333 | 0 |
| Ankrd13b | 0 | 28.5 | 2159.66667 | 1 |
| Ankrd28 | 0 | 247 | 5522 | 0 |
| Ankrd28 | 0 | 245 | 3612.33333 | 1 |
| Ankrd44 | 0 | 359 | 750.666667 | 0 |
| Ankrd44 | 0 | 425 | 1990.33333 | 1 |
| Ankrd55 | 0 | 219.5 | 71.3333333 | 0 |
| Ankrd55 | 0 | 224 | 135.666667 | 1 |
| Ano1 | 0 | 231 | 260.333333 | 0 |
| Ano1 | 0 | 255.5 | 109 | 1 |
| Ano5 | 0 | 185.5 | 50 | 0 |
| Ano5 | 0 | 261 | 12.6666667 | 1 |
| Anpep | 0 | 63 | 880.666667 | 0 |
| Anpep | 0 | 47 | 583 | 1 |
| Antxr2 | 1 | 239.5 | 2239 | 0 |
| Antxr2 | 0 | 281.5 | 3059.66667 | 1 |
| Anxa3 | 0 | 569 | 12270.3333 | 0 |

| | | | | |
|---|---|---|---|---|
| Anxa3 | 0 | 721.5 | 16714 | 1 |
| Anxa8 | 0 | 77 | 2057 | 0 |
| Anxa8 | 0 | 114 | 956 | 1 |
| Apba1 | 0 | 261 | 716 | 0 |
| Apba1 | 0 | 269.5 | 293.666667 | 1 |
| Apba2 | 0 | 643.5 | 19.6666667 | 0 |
| Apba2 | 0 | 431.5 | 6.33333333 | 1 |
| Apcdd1 | 0 | 46 | 2461.33333 | 0 |
| Apcdd1 | 0 | 54.5 | 1417.33333 | 1 |
| Apln | 0 | 23.5 | 1215 | 0 |
| Apln | 0 | 19 | 402.666667 | 1 |
| Apod | 0 | 73.5 | 362.333333 | 0 |
| Apod | 0 | 49.5 | 243.666667 | 1 |
| Apol6 | 0 | 39 | 159 | 0 |
| Apol6 | 0 | 26.5 | 78.3333333 | 1 |
| Apol8 | 0 | 1977.5 | 181.333333 | 0 |
| Apol8 | 0 | 2180.5 | 283.666667 | 1 |
| Aqp5 | 0 | 10 | 578 | 0 |
| Aqp5 | 0 | 8 | 958 | 1 |
| Arc | 0 | 13.5 | 476.333333 | 0 |
| Arc | 0 | 11 | 102.333333 | 1 |
| Arg1 | 0 | 18 | 10 | 0 |
| Arg1 | 0 | 24.5 | 74.6666667 | 1 |
| Arhgap12 | 0 | 351 | 758.666667 | 0 |
| Arhgap12 | 0 | 528 | 1137 | 1 |
| Arhgap18 | 0 | 509.5 | 350.333333 | 0 |
| Arhgap18 | 1 | 586 | 1154 | 1 |
| Arhgap20 | 0 | 76.5 | 404.333333 | 0 |
| Arhgap20 | 0 | 108.5 | 223.333333 | 1 |
| Arhgap22 | 1 | 191.5 | 660 | 0 |
| Arhgap22 | 0 | 179.5 | 407 | 1 |
| Arhgap27 | 0 | 121.5 | 149.666667 | 0 |
| Arhgap27 | 0 | 78.5 | 266 | 1 |
| Arhgap42 | 0 | 57.5 | 2595.33333 | 0 |
| Arhgap42 | 0 | 50 | 1850 | 1 |
| Arhgap8 | 0 | 33.5 | 22.3333333 | 0 |
| Arhgap8 | 0 | 40 | 5.00003333 | 1 |
| Arhgdib | 0 | 25.5 | 6974.33333 | 0 |
| Arhgdib | 0 | 28 | 4729.33333 | 1 |
| Arhgef28 | 0 | 471 | 1790 | 0 |
| Arhgef28 | 0 | 456.5 | 1237.66667 | 1 |
| Arhgef37 | 0 | 63.5 | 37.3333333 | 0 |
| Arhgef37 | 0 | 47.5 | 16.3333333 | 1 |
| Arhgef40 | 1 | 3 | 7948.66667 | 0 |
| Arhgef40 | 0 | 3 | 5615 | 1 |
| Arid5b | 1 | 490 | 7516.66667 | 0 |
| Arid5b | 1 | 520 | 4746.66667 | 1 |

| | | | | |
|---|---|---|---|---|
| Arl14epl | 0 | 51.5 | 11.3333333 | 0 |
| Arl14epl | 0 | 69 | 53 | 1 |
| Arl4c | 0 | 19 | 3334 | 0 |
| Arl4c | 0 | 19.5 | 4668 | 1 |
| Arl4d | 0 | 9.5 | 266.333333 | 0 |
| Arl4d | 0 | 7 | 113.333333 | 1 |
| Arl6ip5 | 0 | 74.5 | 5283.33333 | 0 |
| Arl6ip5 | 0 | 66 | 3461.66667 | 1 |
| Armcx6 | 0 | 6 | 246.333333 | 0 |
| Armcx6 | 0 | 4.5 | 364.333333 | 1 |
| Arntl | 0 | 254.5 | 1204.66667 | 0 |
| Arntl | 0 | 230.5 | 692 | 1 |
| Arsi | 0 | 9.5 | 764.666667 | 0 |
| Arsi | 0 | 8.5 | 1599 | 1 |
| Arsj | 0 | 164.5 | 462 | 0 |
| Arsj | 0 | 155.5 | 282.333333 | 1 |
| Asb5 | 0 | 45.5 | 29 | 0 |
| Asb5 | 0 | 52 | 60 | 1 |
| Ascl1 | 0 | 5.5 | 0.0001 | 0 |
| Ascl1 | 0 | 3.5 | 7.33333333 | 1 |
| Asic1 | 0 | 53 | 58 | 0 |
| Asic1 | 0 | 58 | 100 | 1 |
| Aspn | 0 | 123.5 | 8088.33333 | 0 |
| Aspn | 0 | 134 | 2936.66667 | 1 |
| Ass1 | 0 | 1.5 | 5058.66667 | 0 |
| Ass1 | 0 | 3 | 2485.66667 | 1 |
| Astn2 | 1 | 1028 | 202.666667 | 0 |
| Astn2 | 0 | 1241.5 | 307.333333 | 1 |
| Atf3 | 0 | 38.5 | 243 | 0 |
| Atf3 | 0 | 29.5 | 502.333333 | 1 |
| Atg16l2 | 0 | 298.5 | 425.666667 | 0 |
| Atg16l2 | 0 | 513 | 1539 | 1 |
| Atg4a | 0 | 191.5 | 0.3334 | 0 |
| Atg4a | 0 | 208.5 | 7 | 1 |
| Atoh8 | 0 | 73.5 | 2285.66667 | 0 |
| Atoh8 | 0 | 65.5 | 1636.66667 | 1 |
| Atp2b4 | 0 | 29.5 | 3634.66667 | 0 |
| Atp2b4 | 0 | 44 | 1813.33333 | 1 |
| Atp8b2 | 0 | 85 | 10083.3333 | 0 |
| Atp8b2 | 0 | 72 | 6801.33333 | 1 |
| Atp8b4 | 0 | 407 | 106.666667 | 0 |
| Atp8b4 | 0 | 469 | 63.6666667 | 1 |
| Avpr1a | 0 | 13 | 1219 | 0 |
| Avpr1a | 0 | 13 | 604 | 1 |
| B3galt1 | 0 | 203 | 148 | 0 |
| B3galt1 | 0 | 231.5 | 233 | 1 |
| B3gnt8 | 0 | 38 | 29.3333333 | 0 |

| | | | | |
|---|---|---|---|---|
| B3gnt8 | 0 | 49 | 7 | 1 |
| B3gnt9 | 1 | 66.5 | 1515 | 0 |
| B3gnt9 | 0 | 92 | 836 | 1 |
| B4galnt4 | 0 | 7.5 | 56.6666667 | 0 |
| B4galnt4 | 0 | 9 | 31 | 1 |
| Bach2 | 0 | 199.5 | 699 | 0 |
| Bach2 | 0 | 261 | 2705.33333 | 1 |
| Bambi | 0 | 18.5 | 601 | 0 |
| Bambi | 0 | 14.5 | 1238 | 1 |
| BB218582 | 0 | 18 | 10.3333333 | 0 |
| BB218582 | 0 | 18.5 | 25.6666667 | 1 |
| BC106179 | 0 | 1.5 | 5.66666667 | 0 |
| BC106179 | 0 | 1 | 20.6666667 | 1 |
| Bcar1 | 0 | 192.5 | 7895.33333 | 0 |
| Bcar1 | 0 | 197 | 14832 | 1 |
| Bcar3 | 0 | 234.5 | 526.333333 | 0 |
| Bcar3 | 0 | 209 | 775 | 1 |
| Bcas1 | 0 | 88 | 1.3334 | 0 |
| Bcas1 | 0 | 163 | 15.6666667 | 1 |
| Bcl11b | 0 | 436 | 1105.33333 | 0 |
| Bcl11b | 1 | 360.5 | 747.333333 | 1 |
| Bcl2 | 0 | 301.5 | 598.333333 | 0 |
| Bcl2 | 0 | 542 | 855 | 1 |
| Bcl2l1 | 0 | 150 | 3958.33333 | 0 |
| Bcl2l1 | 0 | 188 | 5579.33333 | 1 |
| Bcl2l11 | 1 | 359.5 | 862.666667 | 0 |
| Bcl2l11 | 0 | 522 | 1261.33333 | 1 |
| Bcl7a | 0 | 58 | 1599.33333 | 0 |
| Bcl7a | 0 | 54 | 1049 | 1 |
| Bdkrb1 | 0 | 77.5 | 33.6666667 | 0 |
| Bdkrb1 | 0 | 82 | 6 | 1 |
| Bdkrb2 | 0 | 38.5 | 199 | 0 |
| Bdkrb2 | 0 | 37 | 51.6666667 | 1 |
| Bend6 | 0 | 64.5 | 96.3333333 | 0 |
| Bend6 | 0 | 83.5 | 57.6666667 | 1 |
| Bex3 | 0 | 8.5 | 6288 | 0 |
| Bex3 | 0 | 9 | 13210.6667 | 1 |
| Bhlhe22 | 0 | 143 | 116.666667 | 0 |
| Bhlhe22 | 0 | 155 | 225.666667 | 1 |
| Bicd1 | 0 | 130 | 117.333333 | 0 |
| Bicd1 | 0 | 181.5 | 205.666667 | 1 |
| Bid | 0 | 83 | 1471 | 0 |
| Bid | 0 | 94 | 1045 | 1 |
| Bmp3 | 0 | 581 | 54 | 0 |
| Bmp3 | 0 | 842 | 159.333333 | 1 |
| Bmp6 | 0 | 136 | 139.333333 | 0 |
| Bmp6 | 0 | 127.5 | 266.666667 | 1 |

| | | | | |
|---|---|---|---|---|
| Bmp7 | 0 | 137 | 6 | 0 |
| Bmp7 | 0 | 102 | 39 | 1 |
| Bmp8b | 0 | 7.5 | 94 | 0 |
| Bmp8b | 0 | 7 | 34.3333333 | 1 |
| Bpgm | 0 | 84 | 848.666667 | 0 |
| Bpgm | 0 | 74.5 | 533.333333 | 1 |
| Brinp1 | 0 | 404.5 | 859.666667 | 0 |
| Brinp1 | 0 | 498.5 | 1237.66667 | 1 |
| Btn1a1 | 0 | 34 | 1.6667 | 0 |
| Btn1a1 | 0 | 30.5 | 19.6666667 | 1 |
| C1qtnf1 | 0 | 42 | 616.666667 | 0 |
| C1qtnf1 | 0 | 38 | 1227.33333 | 1 |
| C1qtnf3 | 0 | 38 | 6201.33333 | 0 |
| C1qtnf3 | 0 | 49 | 3649.66667 | 1 |
| C2cd4a | 0 | 4.5 | 23 | 0 |
| C2cd4a | 0 | 4.5 | 6 | 1 |
| C2cd4b | 0 | 2.5 | 5.00006667 | 0 |
| C2cd4b | 0 | 1 | 0.0001 | 1 |
| C2cd4c | 0 | 31 | 11.3333333 | 0 |
| C2cd4c | 0 | 29 | 1.6667 | 1 |
| C3 | 0 | 16 | 204.333333 | 0 |
| C3 | 0 | 14.5 | 132 | 1 |
| C430002N11Rik | 0 | 82.5 | 259 | 0 |
| C430002N11Rik | 0 | 63 | 140.333333 | 1 |
| C530044C16Rik | 0 | 94 | 24 | 0 |
| C530044C16Rik | 0 | 79.5 | 7.66666667 | 1 |
| C77080 | 0 | 119 | 775 | 0 |
| C77080 | 0 | 107 | 444 | 1 |
| Cables1 | 0 | 77 | 269.333333 | 0 |
| Cables1 | 0 | 45.5 | 113 | 1 |
| Cacna1g | 0 | 255 | 3600 | 0 |
| Cacna1g | 0 | 186.5 | 2159.33333 | 1 |
| Cacnb2 | 0 | 460 | 157 | 0 |
| Cacnb2 | 0 | 563 | 449 | 1 |
| Calcb | 0 | 13.5 | 3.33333333 | 0 |
| Calcb | 0 | 13.5 | 35.3333333 | 1 |
| Camk1d | 0 | 407.5 | 3045.33333 | 0 |
| Camk1d | 0 | 410 | 1671.66667 | 1 |
| Camk2n1 | 0 | 7 | 381.333333 | 0 |
| Camk2n1 | 0 | 15 | 2058 | 1 |
| Camkk2 | 0 | 156.5 | 2473 | 0 |
| Camkk2 | 0 | 153 | 1749 | 1 |
| Cap2 | 0 | 171.5 | 544.666667 | 0 |
| Cap2 | 0 | 178.5 | 282 | 1 |
| Capg | 0 | 59.5 | 4138.66667 | 0 |
| Capg | 0 | 60.5 | 2870.66667 | 1 |
| Car2 | 0 | 25.5 | 43.6666667 | 0 |

| | | | | |
|---|---|---|---|---|
| Car2 | 0 | 24.5 | 124.333333 | 1 |
| Car3 | 0 | 12.5 | 6.33336667 | 0 |
| Car3 | 0 | 23 | 21.6666667 | 1 |
| Car5b | 0 | 65.5 | 147.666667 | 0 |
| Car5b | 0 | 94.5 | 79.3333333 | 1 |
| Card10 | 0 | 37 | 334.333333 | 0 |
| Card10 | 0 | 21.5 | 489.666667 | 1 |
| Casp14 | 0 | 17 | 92.6666667 | 0 |
| Casp14 | 0 | 12 | 194.666667 | 1 |
| Casq2 | 0 | 163.5 | 24.3333333 | 0 |
| Casq2 | 0 | 223 | 9.33333333 | 1 |
| Cav1 | 0 | 128 | 33109.6667 | 0 |
| Cav1 | 0 | 100.5 | 13397.3333 | 1 |
| Cav2 | 0 | 27 | 2149 | 0 |
| Cav2 | 0 | 20 | 935 | 1 |
| Cav3 | 0 | 38.5 | 131 | 0 |
| Cav3 | 0 | 37 | 30.3333333 | 1 |
| Cavin2 | 0 | 365 | 11562 | 0 |
| Cavin2 | 0 | 404.5 | 18226.6667 | 1 |
| Cblb | 1 | 405.5 | 1614 | 0 |
| Cblb | 0 | 410.5 | 943.666667 | 1 |
| Cbln1 | 0 | 9.5 | 68 | 0 |
| Cbln1 | 0 | 6 | 7.6667 | 1 |
| Cbr2 | 0 | 6 | 531 | 0 |
| Cbr2 | 0 | 5.5 | 176 | 1 |
| Ccbe1 | 0 | 60 | 506 | 0 |
| Ccbe1 | 0 | 88.5 | 1130 | 1 |
| Ccdc116 | 0 | 80.5 | 110.333333 | 0 |
| Ccdc116 | 0 | 86.5 | 64.3333333 | 1 |
| Ccdc162 | 0 | 150 | 57.3333333 | 0 |
| Ccdc162 | 0 | 171 | 30 | 1 |
| Ccdc18 | 0 | 153.5 | 279 | 0 |
| Ccdc18 | 0 | 184.5 | 403 | 1 |
| Ccdc3 | 0 | 183 | 205.666667 | 0 |
| Ccdc3 | 0 | 213 | 94.6666667 | 1 |
| Ccdc80 | 0 | 172.5 | 24839.3333 | 0 |
| Ccdc80 | 0 | 200.5 | 16269 | 1 |
| Ccdc81 | 1 | 266 | 24.6666667 | 0 |
| Ccdc81 | 1 | 289.5 | 83.6666667 | 1 |
| Cck | 0 | 95 | 274.666667 | 0 |
| Cck | 0 | 111 | 165.333333 | 1 |
| Ccl11 | 0 | 14.5 | 15.3333333 | 0 |
| Ccl11 | 0 | 16.5 | 2 | 1 |
| Ccl2 | 1 | 12 | 2304.33333 | 0 |
| Ccl2 | 0 | 9 | 1104.66667 | 1 |
| Ccl4 | 0 | 5 | 28 | 0 |
| Ccl4 | 0 | 3 | 56.3333333 | 1 |

| | | | | |
|---|---|---|---|---|
| Ccl6 | 0 | 100 | 6.00006667 | 0 |
| Ccl6 | 0 | 125.5 | 67 | 1 |
| Ccl7 | 1 | 10.5 | 2052.33333 | 0 |
| Ccl7 | 0 | 6.5 | 944.333333 | 1 |
| Cd109 | 0 | 224 | 11907.3333 | 0 |
| Cd109 | 0 | 240 | 8492.66667 | 1 |
| Cd300a | 0 | 63 | 33.6666667 | 0 |
| Cd300a | 0 | 47 | 11.6667 | 1 |
| Cd33 | 0 | 41.5 | 80.6666667 | 0 |
| Cd33 | 0 | 34.5 | 32.0000333 | 1 |
| Cd34 | 0 | 50.5 | 281.333333 | 0 |
| Cd34 | 0 | 42 | 159 | 1 |
| Cd36 | 0 | 301.5 | 53 | 0 |
| Cd36 | 0 | 347.5 | 129.333333 | 1 |
| Cd38 | 0 | 87.5 | 59.6666667 | 0 |
| Cd38 | 0 | 92.5 | 153.666667 | 1 |
| Cd44 | 1 | 729.5 | 28196 | 0 |
| Cd44 | 0 | 472 | 16317.3333 | 1 |
| Cd55 | 0 | 31.5 | 559 | 0 |
| Cd55 | 0 | 47.5 | 198 | 1 |
| Cd72 | 0 | 30 | 146.666667 | 0 |
| Cd72 | 0 | 24 | 59.3333333 | 1 |
| Cd82 | 0 | 57.5 | 903.666667 | 0 |
| Cd82 | 0 | 61 | 584 | 1 |
| Cd9 | 0 | 114 | 6742.33333 | 0 |
| Cd9 | 0 | 98.5 | 4286.33333 | 1 |
| Cdh1 | 0 | 95.5 | 15 | 0 |
| Cdh1 | 0 | 88.5 | 4 | 1 |
| Cdh4 | 0 | 781.5 | 320.333333 | 0 |
| Cdh4 | 0 | 679 | 158 | 1 |
| Cdh6 | 0 | 261.5 | 382.333333 | 0 |
| Cdh6 | 0 | 284 | 251 | 1 |
| Cdh8 | 0 | 617 | 182.666667 | 0 |
| Cdh8 | 0 | 714 | 78 | 1 |
| Cdhr1 | 0 | 35 | 45.6666667 | 0 |
| Cdhr1 | 0 | 36.5 | 143.333333 | 1 |
| Cdk6 | 0 | 131 | 1798 | 0 |
| Cdk6 | 0 | 157 | 2746 | 1 |
| Cdk8 | 0 | 9448 | 10286.6667 | 0 |
| Cdk8 | 0 | 6595 | 6699 | 1 |
| Cdkl1 | 0 | 60.5 | 190.666667 | 0 |
| Cdkl1 | 0 | 57.5 | 113 | 1 |
| Cdkn1c | 0 | 133.5 | 4993.66667 | 0 |
| Cdkn1c | 0 | 145.5 | 3286.66667 | 1 |
| Cdo1 | 0 | 42 | 1945.33333 | 0 |
| Cdo1 | 0 | 47.5 | 2731 | 1 |
| Cdsn | 0 | 39 | 454 | 0 |

| | | | | |
|---|---|---|---|---|
| Cdsn | 0 | 27.5 | 111.333333 | 1 |
| Cebpd | 0 | 263 | 1156 | 0 |
| Cebpd | 0 | 329 | 833.333333 | 1 |
| Cemip | 0 | 279.5 | 10882.3333 | 0 |
| Cemip | 0 | 455 | 15183.3333 | 1 |
| Cenpm | 0 | 33 | 412.333333 | 0 |
| Cenpm | 0 | 24 | 645.333333 | 1 |
| Cep85l | 0 | 43.5 | 268.333333 | 0 |
| Cep85l | 0 | 51.5 | 937 | 1 |
| Cfap100 | 0 | 53.5 | 47.6666667 | 0 |
| Cfap100 | 0 | 64 | 104.333333 | 1 |
| Cfap43 | 0 | 81 | 5449.66667 | 0 |
| Cfap43 | 0 | 67 | 3195.66667 | 1 |
| Cfap45 | 0 | 50.5 | 28.6666667 | 0 |
| Cfap45 | 0 | 38.5 | 12 | 1 |
| Cfh | 0 | 102 | 215 | 0 |
| Cfh | 1 | 123 | 1212.66667 | 1 |
| Cfl2 | 0 | 94 | 4287 | 0 |
| Cfl2 | 0 | 123 | 7861.33333 | 1 |
| Cftr | 0 | 268 | 21.3333333 | 0 |
| Cftr | 0 | 345 | 55.3333333 | 1 |
| Cgnl1 | 0 | 400 | 2625.33333 | 0 |
| Cgnl1 | 0 | 236 | 4733.33333 | 1 |
| Cgref1 | 0 | 35.5 | 775.666667 | 0 |
| Cgref1 | 0 | 28 | 513 | 1 |
| Chchd10 | 0 | 7.5 | 171 | 0 |
| Chchd10 | 0 | 5 | 257.333333 | 1 |
| Chga | 0 | 116.5 | 0.0001 | 0 |
| Chga | 0 | 105.5 | 4.66666667 | 1 |
| Chka | 0 | 223 | 1321 | 0 |
| Chka | 0 | 144 | 2279.66667 | 1 |
| Chl1 | 0 | 484 | 176.666667 | 0 |
| Chl1 | 0 | 466.5 | 48 | 1 |
| Chmp4c | 0 | 46 | 111.333333 | 0 |
| Chmp4c | 0 | 55 | 49.3333333 | 1 |
| Chn2 | 0 | 501 | 70 | 0 |
| Chn2 | 0 | 520 | 116.333333 | 1 |
| Chodl | 0 | 101.5 | 92.6666667 | 0 |
| Chodl | 0 | 99.5 | 15.6666667 | 1 |
| Chrd | 0 | 2 | 383 | 0 |
| Chrd | 0 | 1 | 242 | 1 |
| Chrdl1 | 0 | 378 | 148.333333 | 0 |
| Chrdl1 | 0 | 447.5 | 270.666667 | 1 |
| Chrm3 | 0 | 489 | 36.3333333 | 0 |
| Chrm3 | 0 | 565 | 15.6666667 | 1 |
| Chrna1 | 1 | 48.5 | 24.6666667 | 0 |
| Chrna1 | 0 | 43.5 | 52.3333333 | 1 |

| | | | | |
|---|---|---|---|---|
| Chst1 | 0 | 43.5 | 456 | 0 |
| Chst1 | 0 | 50 | 232 | 1 |
| Chst11 | 0 | 376 | 2028.66667 | 0 |
| Chst11 | 0 | 439 | 6660.66667 | 1 |
| Chst13 | 0 | 17.5 | 8 | 0 |
| Chst13 | 0 | 14 | 38 | 1 |
| Chst7 | 0 | 193 | 535.666667 | 0 |
| Chst7 | 0 | 275.5 | 378.333333 | 1 |
| Ckb | 0 | 88 | 6195.66667 | 0 |
| Ckb | 0 | 112.5 | 8568.66667 | 1 |
| Clca3a1 | 0 | 21.5 | 1926.33333 | 0 |
| Clca3a1 | 0 | 18 | 1183.33333 | 1 |
| Cldn1 | 0 | 29.5 | 110.666667 | 0 |
| Cldn1 | 0 | 35 | 23 | 1 |
| Cldn19 | 0 | 39.5 | 14.6666667 | 0 |
| Cldn19 | 0 | 46 | 0.3334 | 1 |
| Clec11a | 0 | 14 | 652.666667 | 0 |
| Clec11a | 0 | 13 | 383.333333 | 1 |
| Clec14a | 0 | 12.5 | 279.666667 | 0 |
| Clec14a | 0 | 10.5 | 66 | 1 |
| Clec4a1 | 0 | 26.5 | 9.66666667 | 0 |
| Clec4a1 | 0 | 26 | 1.00003333 | 1 |
| Clec4a2 | 0 | 27.5 | 21 | 0 |
| Clec4a2 | 0 | 29.5 | 4.00003333 | 1 |
| Clec7a | 0 | 39.5 | 4.33336667 | 0 |
| Clec7a | 0 | 47 | 44.3333333 | 1 |
| Clu | 0 | 44 | 752.333333 | 0 |
| Clu | 0 | 41 | 1545 | 1 |
| Cma2 | 0 | 10.5 | 10.3333333 | 0 |
| Cma2 | 0 | 17.5 | 34 | 1 |
| Cmah | 0 | 277.5 | 186 | 0 |
| Cmah | 0 | 655 | 296 | 1 |
| Cmbl | 0 | 26.5 | 7.00003333 | 0 |
| Cmbl | 0 | 26 | 80.3333333 | 1 |
| Cmklr1 | 0 | 91.5 | 1692.66667 | 0 |
| Cmklr1 | 0 | 63.5 | 398 | 1 |
| Cmss1 | 0 | 298.5 | 48302.3333 | 0 |
| Cmss1 | 1 | 317.5 | 33201 | 1 |
| Cmya5 | 0 | 82.5 | 55 | 0 |
| Cmya5 | 0 | 106 | 19 | 1 |
| Cnr1 | 0 | 53 | 80 | 0 |
| Cnr1 | 0 | 36.5 | 129 | 1 |
| Cntfr | 0 | 40 | 312 | 0 |
| Cntfr | 0 | 27.5 | 487 | 1 |
| Cobl | 0 | 504.5 | 422.333333 | 0 |
| Cobl | 0 | 607.5 | 253 | 1 |
| Col10a1 | 0 | 103 | 744.666667 | 0 |

| | | | | |
|---|---|---|---|---|
| Col10a1 | 0 | 115.5 | 4659.66667 | 1 |
| Col15a1 | 0 | 175.5 | 2388.66667 | 0 |
| Col15a1 | 0 | 144.5 | 527.666667 | 1 |
| Col18a1 | 0 | 218.5 | 8828.66667 | 0 |
| Col18a1 | 0 | 189 | 5865 | 1 |
| Col23a1 | 0 | 277.5 | 244.666667 | 0 |
| Col23a1 | 0 | 282 | 162.333333 | 1 |
| Col24a1 | 0 | 464.5 | 599.666667 | 0 |
| Col24a1 | 0 | 469 | 374.666667 | 1 |
| Col25a1 | 0 | 652 | 77 | 0 |
| Col25a1 | 0 | 706 | 14 | 1 |
| Col26a1 | 0 | 204 | 116.666667 | 0 |
| Col26a1 | 0 | 416 | 57.6667 | 1 |
| Col2a1 | 0 | 110.5 | 6603 | 0 |
| Col2a1 | 0 | 113 | 4054.66667 | 1 |
| Col3a1 | 0 | 1485.5 | 176922 | 0 |
| Col3a1 | 0 | 2341 | 273352.333 | 1 |
| Col5a1 | 0 | 684 | 198393.667 | 0 |
| Col5a1 | 0 | 568 | 127893.333 | 1 |
| Col5a3 | 0 | 190.5 | 7782.33333 | 0 |
| Col5a3 | 0 | 169.5 | 5383.66667 | 1 |
| Col6a1 | 0 | 313 | 73536.6667 | 0 |
| Col6a1 | 0 | 222.5 | 43965.3333 | 1 |
| Col6a2 | 0 | 402.5 | 51626 | 0 |
| Col6a2 | 0 | 272.5 | 31505 | 1 |
| Col6a3 | 0 | 499.5 | 82821 | 0 |
| Col6a3 | 0 | 513.5 | 42218 | 1 |
| Col6a5 | 0 | 239.5 | 49 | 0 |
| Col6a5 | 0 | 221 | 110 | 1 |
| Col6a6 | 0 | 303.5 | 82 | 0 |
| Col6a6 | 0 | 265 | 46.6666667 | 1 |
| Col7a1 | 0 | 50 | 5046.66667 | 0 |
| Col7a1 | 0 | 70 | 9672 | 1 |
| Colec12 | 0 | 288 | 6417.66667 | 0 |
| Colec12 | 0 | 404.5 | 10829.6667 | 1 |
| Coro6 | 0 | 293.5 | 120.333333 | 0 |
| Coro6 | 0 | 408 | 232.333333 | 1 |
| Cp | 0 | 97 | 75.6666667 | 0 |
| Cp | 0 | 135 | 151.333333 | 1 |
| Cpa4 | 0 | 71.5 | 82 | 0 |
| Cpa4 | 0 | 68.5 | 10 | 1 |
| Cpeb1 | 0 | 255.5 | 981.666667 | 0 |
| Cpeb1 | 1 | 287.5 | 633.666667 | 1 |
| Cpeb4 | 0 | 198 | 1491.33333 | 0 |
| Cpeb4 | 0 | 305 | 2136.33333 | 1 |
| Cped1 | 0 | 332.5 | 2130.33333 | 0 |
| Cped1 | 0 | 385.5 | 1115 | 1 |

| | | | | |
|---|---|---|---|---|
| Cplx2 | 0 | 30 | 1395.33333 | 0 |
| Cplx2 | 0 | 27.5 | 2683.33333 | 1 |
| Cpm | 0 | 85.5 | 108.333333 | 0 |
| Cpm | 0 | 79.5 | 535.666667 | 1 |
| Cpn1 | 0 | 57.5 | 6.6667 | 0 |
| Cpn1 | 0 | 52 | 25.6666667 | 1 |
| Cpne8 | 0 | 637 | 479.333333 | 0 |
| Cpne8 | 0 | 806 | 2205.33333 | 1 |
| Cpz | 0 | 33 | 136.333333 | 0 |
| Cpz | 0 | 36.5 | 232.666667 | 1 |
| Crabp2 | 0 | 12.5 | 616.666667 | 0 |
| Crabp2 | 0 | 17 | 4431 | 1 |
| Crim1 | 0 | 316.5 | 14125 | 0 |
| Crim1 | 0 | 542.5 | 23522.6667 | 1 |
| Crip1 | 0 | 9.5 | 1055 | 0 |
| Crip1 | 0 | 8 | 693.666667 | 1 |
| Crip2 | 0 | 66.5 | 4912.33333 | 0 |
| Crip2 | 0 | 56 | 3020.33333 | 1 |
| Cryab | 0 | 56.5 | 4800 | 0 |
| Cryab | 0 | 25 | 3147.66667 | 1 |
| Crybg1 | 0 | 97 | 302 | 0 |
| Crybg1 | 0 | 79.5 | 132 | 1 |
| Csf3 | 0 | 47.5 | 18.6666667 | 0 |
| Csf3 | 0 | 45 | 41 | 1 |
| Csgalnact1 | 0 | 431.5 | 1085.33333 | 0 |
| Csgalnact1 | 1 | 666 | 658 | 1 |
| Csn3 | 0 | 104.5 | 0.0001 | 0 |
| Csn3 | 0 | 112 | 261.333333 | 1 |
| Cspg4 | 0 | 120 | 13890.6667 | 0 |
| Cspg4 | 0 | 61 | 5372.33333 | 1 |
| Csrp2 | 0 | 110.5 | 1180.33333 | 0 |
| Csrp2 | 0 | 121.5 | 780.333333 | 1 |
| Ctnna2 | 0 | 1041 | 283.666667 | 0 |
| Ctnna2 | 0 | 694.5 | 151.666667 | 1 |
| Ctsb | 0 | 102 | 23138 | 0 |
| Ctsb | 1 | 89.5 | 31727.3333 | 1 |
| Ctsc | 0 | 721 | 2196 | 0 |
| Ctsc | 0 | 655 | 3640.33333 | 1 |
| Ctsw | 0 | 6 | 20 | 0 |
| Ctsw | 0 | 4 | 5.33333333 | 1 |
| Cttnbp2 | 0 | 385.5 | 72.3333333 | 0 |
| Cttnbp2 | 0 | 306.5 | 30.3333333 | 1 |
| Cx3cl1 | 0 | 23 | 2506 | 0 |
| Cx3cl1 | 0 | 17.5 | 1156.33333 | 1 |
| Cx3cr1 | 0 | 37.5 | 769 | 0 |
| Cx3cr1 | 0 | 40 | 450.666667 | 1 |
| Cxcl1 | 0 | 7 | 415.666667 | 0 |

| | | | | |
|---|---|---|---|---|
| Cxcl1 | 0 | 5 | 120 | 1 |
| Cxcl12 | 0 | 57 | 10335.3333 | 0 |
| Cxcl12 | 1 | 115 | 14979.6667 | 1 |
| Cxcl14 | 0 | 19 | 5894.33333 | 0 |
| Cxcl14 | 0 | 11 | 1929.66667 | 1 |
| Cxcl15 | 0 | 24 | 51.3333333 | 0 |
| Cxcl15 | 0 | 15 | 12.6666667 | 1 |
| Cxcl16 | 0 | 11 | 295 | 0 |
| Cxcl16 | 0 | 35 | 143.333333 | 1 |
| Cxcl5 | 0 | 7 | 884 | 0 |
| Cxcl5 | 0 | 11 | 278.333333 | 1 |
| Cxcr6 | 0 | 91 | 62 | 0 |
| Cxcr6 | 0 | 93.5 | 27 | 1 |
| Cxxc4 | 0 | 8.5 | 104.666667 | 0 |
| Cxxc4 | 0 | 3.5 | 55.3333333 | 1 |
| Cyb5r3 | 1 | 55.5 | 20026.3333 | 0 |
| Cyb5r3 | 0 | 37.5 | 14557 | 1 |
| Cyb5r4 | 1 | 79 | 895.333333 | 0 |
| Cyb5r4 | 0 | 148.5 | 1241.33333 | 1 |
| Cyp26a1 | 0 | 6.5 | 3.00003333 | 0 |
| Cyp26a1 | 0 | 7 | 41 | 1 |
| Cyp26b1 | 0 | 400 | 1573.33333 | 0 |
| Cyp26b1 | 0 | 555 | 16541.6667 | 1 |
| Cyp2d22 | 0 | 48 | 88.6666667 | 0 |
| Cyp2d22 | 0 | 24 | 154 | 1 |
| Cyp2f2 | 0 | 37 | 11.0000667 | 0 |
| Cyp2f2 | 0 | 39 | 0.0001 | 1 |
| Cyp4b1 | 0 | 31.5 | 13 | 0 |
| Cyp4b1 | 0 | 35.5 | 1.33336667 | 1 |
| Cyp7b1 | 1 | 156 | 104.333333 | 0 |
| Cyp7b1 | 0 | 190.5 | 406.666667 | 1 |
| Cyth3 | 1 | 225.5 | 10556.3333 | 0 |
| Cyth3 | 1 | 276 | 7061.66667 | 1 |
| D030068K23Rik | 0 | 200.5 | 9 | 0 |
| D030068K23Rik | 0 | 206 | 1.3334 | 1 |
| D430019H16Rik | 0 | 56.5 | 62.3333333 | 0 |
| D430019H16Rik | 0 | 55.5 | 33.3333333 | 1 |
| Dact1 | 0 | 114 | 721.666667 | 0 |
| Dact1 | 0 | 84 | 493.333333 | 1 |
| Dapk1 | 0 | 179 | 1072 | 0 |
| Dapk1 | 0 | 188 | 2106.33333 | 1 |
| Dapk2 | 0 | 200 | 2.66666667 | 0 |
| Dapk2 | 0 | 196.5 | 14 | 1 |
| Dbpht2 | 0 | 31.5 | 20 | 0 |
| Dbpht2 | 0 | 32 | 4 | 1 |
| Dcaf12l1 | 0 | 4.5 | 158 | 0 |
| Dcaf12l1 | 0 | 5.5 | 313 | 1 |

| | | | | |
|---|---|---|---|---|
| Dcbld2 | 0 | 217.5 | 5472 | 0 |
| Dcbld2 | 0 | 390 | 11438.6667 | 1 |
| Dchs2 | 0 | 5.5 | 58 | 0 |
| Dchs2 | 0 | 5 | 15.3333333 | 1 |
| Dclk1 | 0 | 984.5 | 2109.66667 | 0 |
| Dclk1 | 0 | 1497 | 1358.66667 | 1 |
| Dclk2 | 0 | 551.5 | 310.333333 | 0 |
| Dclk2 | 0 | 787.5 | 541.333333 | 1 |
| Ddah2 | 0 | 30.5 | 2823 | 0 |
| Ddah2 | 0 | 26.5 | 3976 | 1 |
| Ddc | 1 | 227 | 99 | 0 |
| Ddc | 0 | 306.5 | 202.666667 | 1 |
| Ddit4l | 0 | 12.5 | 477 | 0 |
| Ddit4l | 0 | 9.5 | 305.333333 | 1 |
| Ddx4 | 0 | 117 | 115.666667 | 0 |
| Ddx4 | 0 | 52.5 | 36.6666667 | 1 |
| Dennd5b | 0 | 157 | 399.666667 | 0 |
| Dennd5b | 0 | 180 | 565 | 1 |
| Depdc5 | 0 | 362 | 691.666667 | 0 |
| Depdc5 | 0 | 210 | 1023 | 1 |
| Dhrs3 | 0 | 428.5 | 837.333333 | 0 |
| Dhrs3 | 1 | 808 | 9502.66667 | 1 |
| Dhrs9 | 0 | 198 | 832.333333 | 0 |
| Dhrs9 | 0 | 197.5 | 595.333333 | 1 |
| Dio2 | 0 | 122.5 | 38.6666667 | 0 |
| Dio2 | 0 | 147.5 | 103.333333 | 1 |
| Dio3 | 0 | 3 | 308.666667 | 0 |
| Dio3 | 0 | 2 | 491 | 1 |
| Dio3os | 0 | 3 | 33.6666667 | 0 |
| Dio3os | 0 | 0 | 71.3333333 | 1 |
| Diras2 | 0 | 44 | 38 | 0 |
| Diras2 | 0 | 68.5 | 12 | 1 |
| Dkk1 | 0 | 359 | 34.6666667 | 0 |
| Dkk1 | 0 | 416 | 15.6666667 | 1 |
| Dkk2 | 0 | 190 | 1401.33333 | 0 |
| Dkk2 | 0 | 206 | 576.333333 | 1 |
| Dlg2 | 0 | 1991.5 | 62.6666667 | 0 |
| Dlg2 | 0 | 2361 | 26.3333333 | 1 |
| Dlk2 | 0 | 25 | 326.333333 | 0 |
| Dlk2 | 0 | 28 | 808 | 1 |
| Dmd | 0 | 2497 | 1645.66667 | 0 |
| Dmd | 0 | 4024.5 | 2564.66667 | 1 |
| Dmrt2 | 0 | 18 | 146.333333 | 0 |
| Dmrt2 | 0 | 19.5 | 87 | 1 |
| Dmtn | 0 | 85 | 43.6666667 | 0 |
| Dmtn | 0 | 62.5 | 83.3333333 | 1 |
| Dnah7b | 0 | 156 | 20.3333333 | 0 |

| | | | | |
|---|---|---|---|---|
| Dnah7b | 0 | 165 | 54.3333333 | 1 |
| Dner | 0 | 113.5 | 375 | 0 |
| Dner | 0 | 146 | 758.666667 | 1 |
| Dnm1 | 0 | 98 | 2222.66667 | 0 |
| Dnm1 | 0 | 48.5 | 1065.33333 | 1 |
| Dock8 | 0 | 247 | 250.666667 | 0 |
| Dock8 | 0 | 235 | 607.333333 | 1 |
| Dpp4 | 0 | 645 | 35.6666667 | 0 |
| Dpp4 | 0 | 621.5 | 167.333333 | 1 |
| Dpt | 0 | 58.5 | 969 | 0 |
| Dpt | 0 | 69 | 602.666667 | 1 |
| Dpysl3 | 0 | 393 | 17700 | 0 |
| Dpysl3 | 0 | 448.5 | 27539.6667 | 1 |
| Dpysl5 | 0 | 168 | 34.3333333 | 0 |
| Dpysl5 | 0 | 201 | 12.3333333 | 1 |
| Dram1 | 0 | 68.5 | 653.666667 | 0 |
| Dram1 | 0 | 60 | 357.333333 | 1 |
| Dtx4 | 0 | 51.5 | 608 | 0 |
| Dtx4 | 0 | 41.5 | 1973 | 1 |
| Dusp1 | 0 | 21 | 5124.33333 | 0 |
| Dusp1 | 0 | 26 | 10685.3333 | 1 |
| Dusp14 | 0 | 175 | 717 | 0 |
| Dusp14 | 0 | 170.5 | 1254.33333 | 1 |
| Dusp2 | 0 | 2.5 | 21 | 0 |
| Dusp2 | 0 | 3 | 6.00003333 | 1 |
| Dusp27 | 0 | 70.5 | 116.333333 | 0 |
| Dusp27 | 0 | 73 | 297.333333 | 1 |
| Dusp4 | 0 | 146.5 | 2413.33333 | 0 |
| Dusp4 | 0 | 154 | 1503.66667 | 1 |
| Dusp6 | 0 | 54.5 | 2985 | 0 |
| Dusp6 | 0 | 48.5 | 2134 | 1 |
| Dusp7 | 0 | 75.5 | 2670.66667 | 0 |
| Dusp7 | 0 | 66.5 | 1718.66667 | 1 |
| Dusp9 | 0 | 13 | 440.666667 | 0 |
| Dusp9 | 0 | 10.5 | 834.333333 | 1 |
| Dynap | 0 | 9.5 | 56.3333333 | 0 |
| Dynap | 0 | 16 | 21.6666667 | 1 |
| E030030I06Rik | 0 | 148 | 34.3333333 | 0 |
| E030030I06Rik | 0 | 120 | 62.6666667 | 1 |
| Ece1 | 0 | 172.5 | 12653.6667 | 0 |
| Ece1 | 0 | 159 | 8609.33333 | 1 |
| Ecm1 | 0 | 33 | 7613 | 0 |
| Ecm1 | 0 | 45.5 | 5615 | 1 |
| Edn1 | 0 | 39 | 518 | 0 |
| Edn1 | 0 | 34.5 | 1022.66667 | 1 |
| Ednrb | 0 | 117.5 | 109.333333 | 0 |
| Ednrb | 0 | 114.5 | 55.6666667 | 1 |

| | | | | |
|---|---|---|---|---|
| Efemp1 | 0 | 125 | 1649.33333 | 0 |
| Efemp1 | 0 | 151 | 4143.66667 | 1 |
| Efna1 | 0 | 9.5 | 157.333333 | 0 |
| Efna1 | 0 | 10.5 | 84.3333333 | 1 |
| Efna3 | 0 | 19.5 | 155.666667 | 0 |
| Efna3 | 0 | 15.5 | 91 | 1 |
| Efnb1 | 0 | 41.5 | 4298 | 0 |
| Efnb1 | 0 | 30.5 | 6760.66667 | 1 |
| Egfl6 | 0 | 82.5 | 996 | 0 |
| Egfl6 | 0 | 83.5 | 652.333333 | 1 |
| Egfr | 0 | 425 | 3874.66667 | 0 |
| Egfr | 0 | 562 | 9648.66667 | 1 |
| Egr1 | 0 | 132 | 10126.6667 | 0 |
| Egr1 | 0 | 96.5 | 15748.6667 | 1 |
| Egr3 | 0 | 307.5 | 388.666667 | 0 |
| Egr3 | 0 | 255 | 171 | 1 |
| Ehd3 | 0 | 52.5 | 1019 | 0 |
| Ehd3 | 0 | 53 | 713.333333 | 1 |
| Ehd4 | 0 | 216 | 2389 | 0 |
| Ehd4 | 0 | 246.5 | 3334 | 1 |
| Elavl2 | 0 | 354 | 43.6666667 | 0 |
| Elavl2 | 0 | 592.5 | 23.6666667 | 1 |
| Elfn1 | 1 | 227 | 474.666667 | 0 |
| Elfn1 | 0 | 185 | 251 | 1 |
| Eln | 0 | 131.5 | 31085.6667 | 0 |
| Eln | 0 | 111 | 15977.3333 | 1 |
| Elobl | 0 | 10.5 | 12.6666667 | 0 |
| Elobl | 0 | 14 | 48 | 1 |
| Elovl4 | 0 | 46.5 | 178.666667 | 0 |
| Elovl4 | 0 | 58.5 | 288 | 1 |
| Eml1 | 0 | 341.5 | 2237.33333 | 0 |
| Eml1 | 0 | 303 | 1618.66667 | 1 |
| Eml2 | 0 | 11.5 | 641.666667 | 0 |
| Eml2 | 0 | 15 | 974.333333 | 1 |
| Eml4 | 0 | 352.5 | 1669 | 0 |
| Eml4 | 0 | 278.5 | 2296.66667 | 1 |
| Emp2 | 0 | 76.5 | 6308.33333 | 0 |
| Emp2 | 0 | 54 | 4649.33333 | 1 |
| Emp3 | 0 | 43.5 | 7539 | 0 |
| Emp3 | 0 | 41.5 | 4763.33333 | 1 |
| Enho | 0 | 108 | 108.666667 | 0 |
| Enho | 0 | 108 | 195.666667 | 1 |
| Enox1 | 0 | 584.5 | 382 | 0 |
| Enox1 | 0 | 852 | 171.666667 | 1 |
| Enpp2 | 0 | 364 | 86.6666667 | 0 |
| Enpp2 | 0 | 349 | 186.666667 | 1 |
| Enpp3 | 0 | 164 | 54.3333333 | 0 |

| | | | | |
|---|---|---|---|---|
| Enpp3 | 0 | 153 | 176 | 1 |
| Epb41l1 | 0 | 289 | 4280.66667 | 0 |
| Epb41l1 | 0 | 291 | 6554.66667 | 1 |
| Epb41l3 | 0 | 475 | 9685.66667 | 0 |
| Epb41l3 | 0 | 468 | 5963.66667 | 1 |
| Epb41l4a | 0 | 223.5 | 602 | 0 |
| Epb41l4a | 0 | 209.5 | 402 | 1 |
| Epb41l4b | 0 | 243.5 | 385.666667 | 0 |
| Epb41l4b | 0 | 141 | 231.333333 | 1 |
| Epha1 | 0 | 15.5 | 545 | 0 |
| Epha1 | 0 | 18 | 263.333333 | 1 |
| Epha4 | 0 | 36 | 1698 | 0 |
| Epha4 | 0 | 21 | 747 | 1 |
| Ephx1 | 0 | 137 | 7073 | 0 |
| Ephx1 | 0 | 128 | 4752.33333 | 1 |
| Ephx2 | 0 | 81 | 91.3333333 | 0 |
| Ephx2 | 0 | 96 | 171.333333 | 1 |
| Epn3 | 0 | 66 | 109 | 0 |
| Epn3 | 0 | 49.5 | 54.3333333 | 1 |
| Epop | 0 | 17 | 948.333333 | 0 |
| Epop | 0 | 19.5 | 1329 | 1 |
| Ermp1 | 0 | 197 | 2324.33333 | 0 |
| Ermp1 | 1 | 203.5 | 5110.33333 | 1 |
| Errfi1 | 1 | 97 | 6618 | 0 |
| Errfi1 | 0 | 110.5 | 15472 | 1 |
| Esd | 0 | 436.5 | 7270.66667 | 0 |
| Esd | 0 | 420 | 11461.6667 | 1 |
| Esm1 | 0 | 20.5 | 19 | 0 |
| Esm1 | 0 | 19 | 224.666667 | 1 |
| Esrp2 | 0 | 6 | 162 | 0 |
| Esrp2 | 0 | 4 | 244 | 1 |
| Etfbkmt | 0 | 69 | 46 | 0 |
| Etfbkmt | 0 | 40 | 109 | 1 |
| Etv1 | 0 | 186 | 211.666667 | 0 |
| Etv1 | 0 | 210 | 82.3333333 | 1 |
| Etv4 | 0 | 152 | 2344.33333 | 0 |
| Etv4 | 0 | 102 | 1603 | 1 |
| Eva1b | 1 | 48 | 10267.6667 | 0 |
| Eva1b | 1 | 48 | 7134.66667 | 1 |
| Eva1c | 1 | 174.5 | 71.6666667 | 0 |
| Eva1c | 0 | 205 | 30.3333333 | 1 |
| Eya2 | 0 | 306.5 | 64.6666667 | 0 |
| Eya2 | 0 | 289 | 198.666667 | 1 |
| Eya4 | 0 | 522 | 924 | 0 |
| Eya4 | 0 | 582 | 427 | 1 |
| F11r | 0 | 80 | 505.666667 | 0 |
| F11r | 0 | 86 | 1649.66667 | 1 |

| | | | | |
|---|---|---|---|---|
| F2rl1 | 0 | 32.5 | 1222.66667 | 0 |
| F2rl1 | 0 | 23 | 664.333333 | 1 |
| F730043M19Rik | 0 | 58.5 | 197.333333 | 0 |
| F730043M19Rik | 0 | 53 | 430.666667 | 1 |
| Fabp3 | 0 | 41 | 361 | 0 |
| Fabp3 | 0 | 33 | 250 | 1 |
| Fads2 | 0 | 77 | 6017.33333 | 0 |
| Fads2 | 1 | 74.5 | 8604.66667 | 1 |
| Fads3 | 0 | 93 | 9020 | 0 |
| Fads3 | 0 | 96.5 | 6432 | 1 |
| Fam107b | 1 | 201 | 3670.66667 | 0 |
| Fam107b | 1 | 318.5 | 8281.66667 | 1 |
| Fam110b | 0 | 352 | 455 | 0 |
| Fam110b | 0 | 359.5 | 303 | 1 |
| Fam129a | 0 | 245.5 | 3967 | 0 |
| Fam129a | 0 | 290 | 6879 | 1 |
| Fam129b | 1 | 139 | 25269.3333 | 0 |
| Fam129b | 1 | 132.5 | 15370.6667 | 1 |
| Fam131a | 0 | 43.5 | 166.666667 | 0 |
| Fam131a | 0 | 40.5 | 105.666667 | 1 |
| Fam131b | 0 | 72 | 623 | 0 |
| Fam131b | 0 | 58 | 424.666667 | 1 |
| Fam13b | 0 | 128 | 2455.66667 | 0 |
| Fam13b | 0 | 192.5 | 5076.66667 | 1 |
| Fam13c | 0 | 300 | 623 | 0 |
| Fam13c | 0 | 148.5 | 357.666667 | 1 |
| Fam167a | 0 | 64.5 | 40.3333333 | 0 |
| Fam167a | 0 | 48.5 | 9.66666667 | 1 |
| Fam169b | 0 | 8.5 | 117.333333 | 0 |
| Fam169b | 0 | 5.5 | 58 | 1 |
| Fam184a | 0 | 114.5 | 45.3333333 | 0 |
| Fam184a | 0 | 138 | 112.666667 | 1 |
| Fam186b | 0 | 40 | 0.66673333 | 0 |
| Fam186b | 0 | 37.5 | 7.33333333 | 1 |
| Fam198b | 0 | 151.5 | 14741.3333 | 0 |
| Fam198b | 0 | 254.5 | 21618 | 1 |
| Fam227a | 0 | 54 | 28.6666667 | 0 |
| Fam227a | 0 | 52 | 53.6666667 | 1 |
| Fam227b | 0 | 87.5 | 16 | 0 |
| Fam227b | 0 | 91 | 4.33336667 | 1 |
| Fam43a | 0 | 11.5 | 1557.33333 | 0 |
| Fam43a | 0 | 5.5 | 2365.33333 | 1 |
| Fam49a | 0 | 172.5 | 331 | 0 |
| Fam49a | 0 | 200 | 195.666667 | 1 |
| Fam78b | 0 | 360 | 54 | 0 |
| Fam78b | 0 | 345 | 97 | 1 |
| Fap | 0 | 134.5 | 511.333333 | 0 |

| | | | | |
|---|---|---|---|---|
| Fap | 0 | 200 | 3059 | 1 |
| Fblim1 | 0 | 69 | 4692.33333 | 0 |
| Fblim1 | 0 | 80 | 6759 | 1 |
| Fbln5 | 1 | 181.5 | 35064 | 0 |
| Fbln5 | 0 | 159 | 19026 | 1 |
| Fbln7 | 0 | 72 | 106 | 0 |
| Fbln7 | 0 | 94.5 | 1030.33333 | 1 |
| Fbn2 | 0 | 421 | 15587 | 0 |
| Fbn2 | 0 | 474 | 9679.66667 | 1 |
| Fbxl7 | 0 | 586 | 696.666667 | 0 |
| Fbxl7 | 0 | 914.5 | 1347.33333 | 1 |
| Fbxl8 | 0 | 10 | 227.333333 | 0 |
| Fbxl8 | 0 | 6 | 337.333333 | 1 |
| Fcgr2b | 0 | 31 | 71.3333333 | 0 |
| Fcgr2b | 0 | 35.5 | 33.3333333 | 1 |
| Fcrls | 0 | 18 | 653 | 0 |
| Fcrls | 0 | 15.5 | 274.666667 | 1 |
| Fez2 | 0 | 336 | 3881.33333 | 0 |
| Fez2 | 1 | 902 | 6005.33333 | 1 |
| Fgf10 | 0 | 138 | 1085.66667 | 0 |
| Fgf10 | 0 | 178 | 332.666667 | 1 |
| Fgf12 | 0 | 1362 | 31.6666667 | 0 |
| Fgf12 | 0 | 1286.5 | 11.6666667 | 1 |
| Fgf18 | 0 | 134 | 93.3333333 | 0 |
| Fgf18 | 0 | 164 | 310 | 1 |
| Fgf5 | 0 | 56.5 | 111.333333 | 0 |
| Fgf5 | 0 | 60.5 | 186.333333 | 1 |
| Fgf7 | 0 | 280.5 | 1380 | 0 |
| Fgf7 | 0 | 344 | 510.666667 | 1 |
| Fgf9 | 0 | 87.5 | 105 | 0 |
| Fgf9 | 0 | 88 | 16 | 1 |
| Fgfr4 | 0 | 42 | 6.33336667 | 0 |
| Fgfr4 | 0 | 35 | 23 | 1 |
| Fgl2 | 0 | 210 | 346 | 0 |
| Fgl2 | 0 | 204 | 1022.66667 | 1 |
| Fhdc1 | 0 | 188.5 | 2491.66667 | 0 |
| Fhdc1 | 0 | 169.5 | 1206 | 1 |
| Fhl1 | 0 | 139.5 | 13852.3333 | 0 |
| Fhl1 | 0 | 178 | 19486 | 1 |
| Fhl5 | 0 | 109 | 59.3333333 | 0 |
| Fhl5 | 0 | 141.5 | 194.333333 | 1 |
| Fhod3 | 0 | 976 | 185.333333 | 0 |
| Fhod3 | 0 | 1090 | 1127.33333 | 1 |
| Fibcd1 | 0 | 31 | 63.3333333 | 0 |
| Fibcd1 | 0 | 25.5 | 19 | 1 |
| Fibin | 0 | 14 | 1576.33333 | 0 |
| Fibin | 1 | 25 | 3804.66667 | 1 |

| | | | | |
|---|---|---|---|---|
| Filip1 | 0 | 414.5 | 37.3333333 | 0 |
| Filip1 | 0 | 379.5 | 13.6666667 | 1 |
| Filip1l | 0 | 1157 | 7660 | 0 |
| Filip1l | 1 | 1136 | 10787 | 1 |
| Fjx1 | 0 | 5 | 1223.66667 | 0 |
| Fjx1 | 0 | 3 | 2557.33333 | 1 |
| Flrt1 | 0 | 152 | 158.666667 | 0 |
| Flrt1 | 0 | 113 | 316.666667 | 1 |
| Flrt2 | 0 | 242 | 10263.6667 | 0 |
| Flrt2 | 0 | 263 | 5523.66667 | 1 |
| Flt1 | 0 | 379.5 | 11555 | 0 |
| Flt1 | 0 | 360.5 | 7962 | 1 |
| Flvcr2 | 0 | 110.5 | 90 | 0 |
| Flvcr2 | 0 | 108.5 | 146 | 1 |
| Fmnl1 | 0 | 50.5 | 360 | 0 |
| Fmnl1 | 0 | 60 | 246 | 1 |
| Fmo2 | 0 | 49 | 13 | 0 |
| Fmo2 | 0 | 68 | 134.333333 | 1 |
| Fmo3 | 0 | 54.5 | 3 | 0 |
| Fmo3 | 0 | 81.5 | 39.3333333 | 1 |
| Fmo6 | 0 | 64.5 | 0.66673333 | 0 |
| Fmo6 | 0 | 107 | 20.3333333 | 1 |
| Fmod | 0 | 34.5 | 6568.66667 | 0 |
| Fmod | 0 | 24 | 3921.33333 | 1 |
| Fndc1 | 0 | 188 | 4267.33333 | 0 |
| Fndc1 | 0 | 189.5 | 2826.33333 | 1 |
| Fos | 0 | 57.5 | 2685 | 0 |
| Fos | 0 | 49.5 | 7371 | 1 |
| Fosb | 0 | 300.5 | 960 | 0 |
| Fosb | 0 | 155.5 | 3487.33333 | 1 |
| Fosl1 | 0 | 19.5 | 2094 | 0 |
| Fosl1 | 0 | 12 | 1382.33333 | 1 |
| Fosl2 | 0 | 443 | 12334 | 0 |
| Fosl2 | 0 | 504.5 | 8723.33333 | 1 |
| Foxc1 | 0 | 496 | 1109 | 0 |
| Foxc1 | 0 | 550.5 | 1561.66667 | 1 |
| Foxe3 | 0 | 4.5 | 11.3334 | 0 |
| Foxe3 | 0 | 8.5 | 0.0001 | 1 |
| Frem1 | 0 | 161 | 340.666667 | 0 |
| Frem1 | 0 | 216 | 91.3333333 | 1 |
| Frk | 0 | 458 | 405 | 0 |
| Frk | 0 | 442.5 | 574.333333 | 1 |
| Frmpd3 | 0 | 124.5 | 53.6666667 | 0 |
| Frmpd3 | 0 | 123 | 25 | 1 |
| Fscn1 | 0 | 162.5 | 42074.3333 | 0 |
| Fscn1 | 0 | 172 | 57628 | 1 |
| Fst | 0 | 114.5 | 10117.6667 | 0 |

| | | | | |
|---|---|---|---|---|
| Fst | 0 | 42 | 2755.33333 | 1 |
| Fxyd5 | 0 | 10 | 1877 | 0 |
| Fxyd5 | 0 | 23.5 | 1325.33333 | 1 |
| Fxyd6 | 0 | 48.5 | 653.666667 | 0 |
| Fxyd6 | 0 | 36 | 395.666667 | 1 |
| Fxyd7 | 0 | 5 | 23.3333333 | 0 |
| Fxyd7 | 0 | 8 | 3.00003333 | 1 |
| Fyb | 0 | 97 | 125.666667 | 0 |
| Fyb | 0 | 148 | 60 | 1 |
| Fzd7 | 0 | 24 | 8067.33333 | 0 |
| Fzd7 | 0 | 37.5 | 17022.3333 | 1 |
| G630016G05Rik | 0 | 4 | 84.6666667 | 0 |
| G630016G05Rik | 0 | 3.5 | 165.666667 | 1 |
| Gab2 | 0 | 390 | 668.333333 | 0 |
| Gab2 | 0 | 673 | 1039.66667 | 1 |
| Gabbr2 | 0 | 369.5 | 25 | 0 |
| Gabbr2 | 0 | 361 | 4 | 1 |
| Gabra1 | 0 | 64.5 | 54.3333333 | 0 |
| Gabra1 | 0 | 80 | 94.3333333 | 1 |
| Gabra3 | 0 | 96.5 | 12 | 0 |
| Gabra3 | 0 | 125 | 37 | 1 |
| Gabrb2 | 0 | 372.5 | 71.6666667 | 0 |
| Gabrb2 | 0 | 469 | 135 | 1 |
| Gabre | 0 | 11 | 93 | 0 |
| Gabre | 0 | 9.5 | 160.666667 | 1 |
| Gadd45a | 0 | 93 | 521 | 0 |
| Gadd45a | 1 | 103 | 983.666667 | 1 |
| Galnt14 | 0 | 315 | 221.666667 | 0 |
| Galnt14 | 0 | 361 | 413.333333 | 1 |
| Galnt9 | 0 | 237.5 | 192.333333 | 0 |
| Galnt9 | 0 | 204 | 31.3333333 | 1 |
| Gap43 | 0 | 44.5 | 785 | 0 |
| Gap43 | 0 | 58.5 | 316 | 1 |
| Garem2 | 0 | 3 | 296.333333 | 0 |
| Garem2 | 0 | 1.5 | 165.333333 | 1 |
| Gas1 | 0 | 45 | 8910.33333 | 0 |
| Gas1 | 1 | 48.5 | 15129.3333 | 1 |
| Gas2 | 0 | 156.5 | 1046.66667 | 0 |
| Gas2 | 0 | 152.5 | 440.666667 | 1 |
| Gas6 | 0 | 106.5 | 21690.3333 | 0 |
| Gas6 | 0 | 76.5 | 9263.33333 | 1 |
| Gas7 | 0 | 329 | 1745.66667 | 0 |
| Gas7 | 0 | 433.5 | 3189.66667 | 1 |
| Gata6 | 0 | 24 | 291.333333 | 0 |
| Gata6 | 0 | 20.5 | 425 | 1 |
| Gba2 | 0 | 4.5 | 582.333333 | 0 |
| Gba2 | 0 | 8 | 869.333333 | 1 |

| | | | | |
|---|---|---|---|---|
| Gbx2 | 0 | 200 | 11.6666667 | 0 |
| Gbx2 | 0 | 227.5 | 29.3333333 | 1 |
| Gcnt2 | 1 | 200 | 1073.33333 | 0 |
| Gcnt2 | 0 | 231.5 | 698.333333 | 1 |
| Gcnt3 | 0 | 23 | 4.66666667 | 0 |
| Gcnt3 | 0 | 16.5 | 29.3333333 | 1 |
| Gda | 0 | 111 | 108.666667 | 0 |
| Gda | 0 | 159 | 547 | 1 |
| Gdf10 | 0 | 15.5 | 242.333333 | 0 |
| Gdf10 | 0 | 13.5 | 691.666667 | 1 |
| Gdf6 | 0 | 63.5 | 307 | 0 |
| Gdf6 | 0 | 73 | 889.666667 | 1 |
| Gdnf | 0 | 185 | 199 | 0 |
| Gdnf | 0 | 204 | 746 | 1 |
| Gdpd5 | 1 | 154.5 | 762 | 0 |
| Gdpd5 | 0 | 165 | 2310.66667 | 1 |
| Gem | 0 | 24 | 313.666667 | 0 |
| Gem | 0 | 21 | 181.333333 | 1 |
| Gfpt2 | 0 | 66 | 855.333333 | 0 |
| Gfpt2 | 0 | 78 | 1611.33333 | 1 |
| Gfra1 | 0 | 331 | 3636.33333 | 0 |
| Gfra1 | 0 | 355 | 2670 | 1 |
| Gfra4 | 0 | 16 | 216.333333 | 0 |
| Gfra4 | 0 | 22 | 320 | 1 |
| Ggct | 0 | 15.5 | 124.666667 | 0 |
| Ggct | 1 | 19 | 65.3333333 | 1 |
| Ghr | 0 | 326.5 | 3772.66667 | 0 |
| Ghr | 0 | 519 | 6950.33333 | 1 |
| Gimap6 | 0 | 19.5 | 5.66666667 | 0 |
| Gimap6 | 0 | 24.5 | 40 | 1 |
| Git1 | 0 | 40.5 | 4691 | 0 |
| Git1 | 0 | 36.5 | 7075.33333 | 1 |
| Gja3 | 0 | 47.5 | 42.6666667 | 0 |
| Gja3 | 0 | 49 | 17.6666667 | 1 |
| Gja4 | 0 | 6 | 8.6667 | 0 |
| Gja4 | 0 | 2 | 27.6666667 | 1 |
| Gja5 | 0 | 122 | 35.3333333 | 0 |
| Gja5 | 0 | 107 | 6.00003333 | 1 |
| Gjb2 | 0 | 17.5 | 7928 | 0 |
| Gjb2 | 0 | 11 | 2072 | 1 |
| Gjb3 | 0 | 8.5 | 404.333333 | 0 |
| Gjb3 | 0 | 14 | 191.666667 | 1 |
| Gjb4 | 0 | 20 | 110.666667 | 0 |
| Gjb4 | 0 | 17.5 | 55 | 1 |
| Gjb5 | 0 | 7 | 92.3333333 | 0 |
| Gjb5 | 0 | 4.5 | 44.3333333 | 1 |
| Gldn | 0 | 26.5 | 235.666667 | 0 |

| | | | | |
|---|---|---|---|---|
| Gldn | 0 | 30 | 121.666667 | 1 |
| Gli1 | 0 | 30.5 | 21.6666667 | 0 |
| Gli1 | 0 | 23.5 | 3.33333333 | 1 |
| Glis1 | 0 | 315 | 488.666667 | 0 |
| Glis1 | 0 | 282.5 | 271.333333 | 1 |
| Glrx | 0 | 76 | 791.666667 | 0 |
| Glrx | 0 | 59.5 | 375 | 1 |
| Glul | 0 | 21.5 | 1196.66667 | 0 |
| Glul | 0 | 23 | 2153 | 1 |
| Gm10190 | 0 | 145 | 17.6667 | 0 |
| Gm10190 | 1 | 148 | 0.0001 | 1 |
| Gm10516 | 0 | 57.5 | 95.3333333 | 0 |
| Gm10516 | 0 | 63 | 185.333333 | 1 |
| Gm10575 | 0 | 9 | 357.333333 | 0 |
| Gm10575 | 0 | 6.5 | 239.666667 | 1 |
| Gm10941 | 0 | 5 | 17.3333333 | 0 |
| Gm10941 | 0 | 2.5 | 4.66666667 | 1 |
| Gm13986 | 0 | 352 | 525 | 0 |
| Gm13986 | 0 | 420 | 314 | 1 |
| Gm2115 | 0 | 96.5 | 6054.66667 | 0 |
| Gm2115 | 0 | 113 | 3816.66667 | 1 |
| Gm26682 | 0 | 106.5 | 53 | 0 |
| Gm26682 | 0 | 110.5 | 99.6666667 | 1 |
| Gm26705 | 0 | 10 | 111.666667 | 0 |
| Gm26705 | 0 | 9.5 | 57.6666667 | 1 |
| Gm2694 | 0 | 83 | 65.6666667 | 0 |
| Gm2694 | 0 | 72 | 22.6666667 | 1 |
| Gm3219 | 0 | 43.5 | 42 | 0 |
| Gm3219 | 0 | 31.5 | 14.3333333 | 1 |
| Gm3417 | 0 | 1 | 0.0001 | 0 |
| Gm3417 | 0 | 0.5 | 4.6667 | 1 |
| Gm4070 | 0 | 0 | 7.33333333 | 0 |
| Gm4070 | 0 | 1 | 56.6666667 | 1 |
| Gm5424 | 0 | 95.5 | 2224.33333 | 0 |
| Gm5424 | 0 | 104 | 1090.66667 | 1 |
| Gm6093 | 0 | 10.5 | 64.6666667 | 0 |
| Gm6093 | 0 | 5.5 | 110.666667 | 1 |
| Gm6644 | 0 | 30 | 25 | 0 |
| Gm6644 | 0 | 18.5 | 54 | 1 |
| Gnai1 | 0 | 135.5 | 694.333333 | 0 |
| Gnai1 | 0 | 149 | 350.666667 | 1 |
| Gnao1 | 0 | 0 | 945 | 0 |
| Gnao1 | 0 | 0 | 598.666667 | 1 |
| Gnb4 | 0 | 87.5 | 1851 | 0 |
| Gnb4 | 0 | 88.5 | 998.666667 | 1 |
| Gorab | 0 | 68.5 | 1204.33333 | 0 |
| Gorab | 0 | 85.5 | 818 | 1 |

| | | | | |
|---|---|---|---|---|
| Gpat3 | 0 | 110 | 65.6666667 | 0 |
| Gpat3 | 0 | 108.5 | 109.666667 | 1 |
| Gpbar1 | 0 | 42 | 123.333333 | 0 |
| Gpbar1 | 0 | 51 | 32.6666667 | 1 |
| Gper1 | 0 | 44 | 198.333333 | 0 |
| Gper1 | 0 | 41 | 14.3333333 | 1 |
| Gphn | 0 | 534 | 9490.66667 | 0 |
| Gphn | 0 | 399 | 6795.33333 | 1 |
| Gpi1 | 0 | 106.5 | 14756.3333 | 0 |
| Gpi1 | 0 | 178.5 | 23966.6667 | 1 |
| Gpnmb | 0 | 49.5 | 1843 | 0 |
| Gpnmb | 0 | 35 | 1140.66667 | 1 |
| Gpr20 | 0 | 39 | 6.66666667 | 0 |
| Gpr20 | 0 | 28 | 0.0001 | 1 |
| Gpr39 | 0 | 224.5 | 171.666667 | 0 |
| Gpr39 | 0 | 270.5 | 438 | 1 |
| Gpr88 | 0 | 76 | 96.3333333 | 0 |
| Gpr88 | 0 | 75 | 50.6666667 | 1 |
| Gprc5a | 0 | 69 | 2260.66667 | 0 |
| Gprc5a | 0 | 66 | 3870.66667 | 1 |
| Gprc5b | 0 | 200.5 | 860.333333 | 0 |
| Gprc5b | 0 | 209 | 4345.33333 | 1 |
| Gprin3 | 1 | 146.5 | 83 | 0 |
| Gprin3 | 0 | 163 | 19.6666667 | 1 |
| Gpx3 | 0 | 60 | 793 | 0 |
| Gpx3 | 0 | 117 | 1458.33333 | 1 |
| Gramd2 | 0 | 24.5 | 68.6666667 | 0 |
| Gramd2 | 0 | 17 | 17 | 1 |
| Greb1l | 0 | 239 | 119 | 0 |
| Greb1l | 0 | 296 | 271.666667 | 1 |
| Grem1 | 0 | 66 | 6086 | 0 |
| Grem1 | 0 | 66 | 2217.33333 | 1 |
| Grem2 | 0 | 706.5 | 10842 | 0 |
| Grem2 | 0 | 798.5 | 6027.66667 | 1 |
| Gria1 | 0 | 557.5 | 258.666667 | 0 |
| Gria1 | 0 | 726.5 | 172 | 1 |
| Gria4 | 0 | 468.5 | 18 | 0 |
| Gria4 | 0 | 797 | 48.6666667 | 1 |
| Grid2 | 0 | 1348.5 | 178.666667 | 0 |
| Grid2 | 0 | 1604.5 | 109 | 1 |
| Grik1 | 0 | 511.5 | 165 | 0 |
| Grik1 | 0 | 430.5 | 95.3333333 | 1 |
| Grik5 | 0 | 179 | 1682 | 0 |
| Grik5 | 0 | 161 | 1062 | 1 |
| Grin2b | 0 | 460.5 | 23.6666667 | 0 |
| Grin2b | 0 | 522.5 | 9.33333333 | 1 |
| Grpr | 0 | 49.5 | 14.3333333 | 0 |

| | | | | |
|---|---|---|---|---|
| Grpr | 0 | 82 | 110.666667 | 1 |
| Gsdmd | 0 | 17.5 | 790.333333 | 0 |
| Gsdmd | 0 | 9 | 556.333333 | 1 |
| Gsg1 | 1 | 93.5 | 25 | 0 |
| Gsg1 | 0 | 78.5 | 8.33333333 | 1 |
| Gsn | 0 | 191.5 | 26101 | 0 |
| Gsn | 0 | 172.5 | 19217.3333 | 1 |
| Gsta1 | 0 | 1 | 7.66666667 | 0 |
| Gsta1 | 0 | 0.5 | 1.6667 | 1 |
| Gsta4 | 0 | 55.5 | 791 | 0 |
| Gsta4 | 0 | 52 | 443 | 1 |
| Gsto1 | 0 | 29.5 | 11283.3333 | 0 |
| Gsto1 | 0 | 16.5 | 7092.33333 | 1 |
| Gsto2 | 0 | 70 | 87.6666667 | 0 |
| Gsto2 | 0 | 58.5 | 39.6666667 | 1 |
| Gstt1 | 0 | 21 | 105.666667 | 0 |
| Gstt1 | 0 | 23 | 60.3333333 | 1 |
| Gucy1a1 | 0 | 35.5 | 103 | 0 |
| Gucy1a1 | 0 | 33 | 363.666667 | 1 |
| Gucy1b1 | 0 | 107 | 164 | 0 |
| Gucy1b1 | 0 | 110 | 313.333333 | 1 |
| Gulp1 | 0 | 227 | 1379.66667 | 0 |
| Gulp1 | 0 | 294.5 | 3145 | 1 |
| Gvin1 | 0 | 0 | 19.6666667 | 0 |
| Gvin1 | 0 | 0 | 87.3333333 | 1 |
| Gzmb | 0 | 8.5 | 3.33333333 | 0 |
| Gzmb | 0 | 14 | 18.3333333 | 1 |
| Gzmd | 0 | 10.5 | 29.3333333 | 0 |
| Gzmd | 0 | 8.5 | 65.6666667 | 1 |
| Gzme | 0 | 7 | 127.333333 | 0 |
| Gzme | 0 | 6.5 | 386.333333 | 1 |
| H2-M3 | 0 | 9.5 | 132 | 0 |
| H2-M3 | 0 | 11 | 80 | 1 |
| Hal | 0 | 94 | 47 | 0 |
| Hal | 0 | 72.5 | 20.6666667 | 1 |
| Hapln1 | 0 | 349.5 | 77 | 0 |
| Hapln1 | 0 | 373 | 28.6666667 | 1 |
| Has1 | 0 | 35.5 | 18.3333333 | 0 |
| Has1 | 0 | 31.5 | 6.66666667 | 1 |
| Hcn1 | 0 | 97 | 235 | 0 |
| Hcn1 | 0 | 93.5 | 91 | 1 |
| Hcn2 | 0 | 47.5 | 181.666667 | 0 |
| Hcn2 | 0 | 50.5 | 275.333333 | 1 |
| Hcn4 | 0 | 176.5 | 6.66666667 | 0 |
| Hcn4 | 0 | 190.5 | 88.6666667 | 1 |
| Hdac5 | 0 | 8 | 3238.33333 | 0 |
| Hdac5 | 0 | 4.5 | 2258 | 1 |

| | | | | |
|---|---|---|---|---|
| Hecw1 | 0 | 389.5 | 60 | 0 |
| Hecw1 | 0 | 458 | 23.3333333 | 1 |
| Heg1 | 0 | 312 | 5030 | 0 |
| Heg1 | 0 | 342 | 13505.6667 | 1 |
| Helb | 0 | 145 | 818.666667 | 0 |
| Helb | 0 | 140.5 | 572.333333 | 1 |
| Helz | 0 | 94.5 | 2089.33333 | 0 |
| Helz | 0 | 99 | 3112 | 1 |
| Herc3 | 0 | 67.5 | 536.333333 | 0 |
| Herc3 | 0 | 102.5 | 1422.33333 | 1 |
| Hes3 | 0 | 21.5 | 0.0001 | 0 |
| Hes3 | 1 | 23.5 | 7.6667 | 1 |
| Hey1 | 0 | 16.5 | 371.333333 | 0 |
| Hey1 | 0 | 12 | 192 | 1 |
| Hey2 | 0 | 20.5 | 175.333333 | 0 |
| Hey2 | 0 | 24 | 81.3333333 | 1 |
| Heyl | 0 | 33 | 342.666667 | 0 |
| Heyl | 0 | 27.5 | 101.333333 | 1 |
| Hfe | 0 | 36.5 | 164.666667 | 0 |
| Hfe | 0 | 27.5 | 95.6666667 | 1 |
| Hgf | 0 | 564 | 634.666667 | 0 |
| Hgf | 0 | 730.5 | 224 | 1 |
| Hic1 | 1 | 147.5 | 11484 | 0 |
| Hic1 | 1 | 182.5 | 33526.6667 | 1 |
| Hipk2 | 0 | 323 | 2336.66667 | 0 |
| Hipk2 | 0 | 376.5 | 3406.66667 | 1 |
| Hipk4 | 0 | 70.5 | 125.666667 | 0 |
| Hipk4 | 0 | 64 | 248.666667 | 1 |
| Hist1h2ah | 0 | 4 | 18.6666667 | 0 |
| Hist1h2ah | 0 | 6 | 3.00006667 | 1 |
| Hist1h2al | 0 | 5 | 196.333333 | 0 |
| Hist1h2al | 0 | 2.5 | 112.666667 | 1 |
| Hist1h2an | 0 | 4 | 9.00003333 | 0 |
| Hist1h2an | 0 | 1.5 | 0.0001 | 1 |
| Hist1h2bc | 0 | 26.5 | 577.333333 | 0 |
| Hist1h2bc | 0 | 23.5 | 408 | 1 |
| Hist1h2bl | 0 | 3 | 10.6666667 | 0 |
| Hist1h2bl | 0 | 4.5 | 0.0001 | 1 |
| Hist2h2aa2 | 0 | 5.5 | 23.3333333 | 0 |
| Hist2h2aa2 | 0 | 5 | 3.00006667 | 1 |
| Hivep3 | 0 | 370.5 | 810.666667 | 0 |
| Hivep3 | 0 | 346.5 | 373.333333 | 1 |
| Hk2 | 0 | 160 | 3371 | 0 |
| Hk2 | 0 | 182.5 | 6982.66667 | 1 |
| Hlf | 0 | 170.5 | 28 | 0 |
| Hlf | 0 | 160.5 | 125 | 1 |
| Hmga2 | 0 | 84.5 | 29089.3333 | 0 |

| | | | | |
|---|---|---|---|---|
| Hmga2 | 1 | 72.5 | 17459 | 1 |
| Hmx3 | 0 | 0 | 3.66673333 | 0 |
| Hmx3 | 0 | 0 | 16.6666667 | 1 |
| Hnrnpu | 0 | 138 | 285.333333 | 0 |
| Hnrnpu | 0 | 123 | 163 | 1 |
| Homer2 | 0 | 187 | 336.333333 | 0 |
| Homer2 | 0 | 141.5 | 215.333333 | 1 |
| Hopx | 0 | 91.5 | 95 | 0 |
| Hopx | 0 | 137 | 41.3333333 | 1 |
| Hoxaas3 | 0 | 17 | 18 | 0 |
| Hoxaas3 | 0 | 18.5 | 57.3333333 | 1 |
| Hoxb3 | 0 | 55.5 | 826.333333 | 0 |
| Hoxb3 | 0 | 77 | 1366.33333 | 1 |
| Hoxb4 | 0 | 4.5 | 306 | 0 |
| Hoxb4 | 0 | 3.5 | 450.333333 | 1 |
| Hoxc4 | 0 | 5 | 425.666667 | 0 |
| Hoxc4 | 0 | 5.5 | 643.666667 | 1 |
| Hoxc5 | 0 | 6 | 486 | 0 |
| Hoxc5 | 0 | 4.5 | 1013.33333 | 1 |
| Hoxd12 | 0 | 4 | 63 | 0 |
| Hoxd12 | 0 | 4 | 180.666667 | 1 |
| Hpgd | 0 | 60 | 139.333333 | 0 |
| Hpgd | 0 | 70.5 | 77.3333333 | 1 |
| Hr | 0 | 31.5 | 113.333333 | 0 |
| Hr | 0 | 24 | 21.6666667 | 1 |
| Hrh1 | 0 | 234 | 33.3333333 | 0 |
| Hrh1 | 0 | 245.5 | 64.3333333 | 1 |
| Hs3st1 | 0 | 42.5 | 69 | 0 |
| Hs3st1 | 1 | 52 | 781 | 1 |
| Hs3st5 | 0 | 740 | 42.6666667 | 0 |
| Hs3st5 | 0 | 925.5 | 21.6666667 | 1 |
| Hs6st1 | 0 | 95.5 | 4756.66667 | 0 |
| Hs6st1 | 0 | 100 | 7102.33333 | 1 |
| Hsd17b1 | 0 | 18 | 36.6666667 | 0 |
| Hsd17b1 | 0 | 6 | 16.3333333 | 1 |
| Hspa12a | 0 | 234.5 | 789.666667 | 0 |
| Hspa12a | 0 | 223.5 | 1226.66667 | 1 |
| Hspa4l | 0 | 58 | 22.3333333 | 0 |
| Hspa4l | 0 | 73 | 4 | 1 |
| Hspb6 | 0 | 1 | 1802.66667 | 0 |
| Hspb6 | 0 | 3 | 1088.33333 | 1 |
| Hspb7 | 0 | 12 | 445.666667 | 0 |
| Hspb7 | 1 | 7.5 | 164.333333 | 1 |
| Htr1d | 0 | 46 | 0.66673333 | 0 |
| Htr1d | 0 | 48 | 8.33333333 | 1 |
| Htr5b | 0 | 80.5 | 0.66673333 | 0 |
| Htr5b | 0 | 76.5 | 10.3333333 | 1 |

| | | | | |
|---|---|---|---|---|
| Htr7 | 0 | 194 | 39 | 0 |
| Htr7 | 0 | 266.5 | 16 | 1 |
| Htra1 | 1 | 142.5 | 5738.33333 | 0 |
| Htra1 | 1 | 150 | 16732.3333 | 1 |
| Icmt | 0 | 41.5 | 4991.66667 | 0 |
| Icmt | 0 | 23 | 7636.66667 | 1 |
| Id2 | 0 | 62.5 | 2200.33333 | 0 |
| Id2 | 0 | 85.5 | 3232.66667 | 1 |
| Id3 | 0 | 61.5 | 12918.3333 | 0 |
| Id3 | 1 | 70.5 | 17922.6667 | 1 |
| Ifi202b | 0 | 39 | 299.333333 | 0 |
| Ifi202b | 0 | 34.5 | 175.666667 | 1 |
| Ifi204 | 0 | 5 | 342 | 0 |
| Ifi204 | 0 | 4.5 | 200.333333 | 1 |
| Ifi205 | 0 | 15 | 16 | 0 |
| Ifi205 | 0 | 10 | 39.3333333 | 1 |
| Ifi27l2b | 0 | 5 | 156 | 0 |
| Ifi27l2b | 0 | 5.5 | 247.666667 | 1 |
| Ifitm1 | 0 | 4.5 | 53.6666667 | 0 |
| Ifitm1 | 0 | 3.5 | 26 | 1 |
| Ifitm10 | 0 | 93.5 | 238.666667 | 0 |
| Ifitm10 | 0 | 98.5 | 122.333333 | 1 |
| Ifitm3 | 0 | 6.5 | 7907.33333 | 0 |
| Ifitm3 | 0 | 9.5 | 5660.33333 | 1 |
| Ifitm6 | 0 | 5 | 3.33333333 | 0 |
| Ifitm6 | 0 | 4 | 16.6666667 | 1 |
| Ifngr1 | 0 | 48 | 1754 | 0 |
| Ifngr1 | 0 | 39 | 3205.33333 | 1 |
| Igfbp2 | 0 | 111 | 13510.3333 | 0 |
| Igfbp2 | 0 | 76 | 6624 | 1 |
| Igfbp3 | 0 | 16.5 | 8826.33333 | 0 |
| Igfbp3 | 0 | 37 | 42212.6667 | 1 |
| Igfbp5 | 0 | 80.5 | 16335 | 0 |
| Igfbp5 | 0 | 57.5 | 5914.66667 | 1 |
| Igfbp6 | 0 | 23 | 1720 | 0 |
| Igfbp6 | 0 | 23 | 3757.66667 | 1 |
| Igfbpl1 | 0 | 42 | 4.66666667 | 0 |
| Igfbpl1 | 0 | 24.5 | 21 | 1 |
| Igip | 0 | 7 | 153 | 0 |
| Igip | 0 | 12.5 | 261 | 1 |
| Igsf10 | 0 | 87 | 3481.33333 | 0 |
| Igsf10 | 0 | 88.5 | 753.333333 | 1 |
| Il11 | 0 | 16 | 279.666667 | 0 |
| Il11 | 0 | 11 | 89.3333333 | 1 |
| Il13ra1 | 0 | 212.5 | 1529 | 0 |
| Il13ra1 | 0 | 255 | 864 | 1 |
| Il15 | 0 | 1035 | 10 | 0 |

| | | | | |
|---|---|---|---|---|
| Il15 | 0 | 825.5 | 28.6666667 | 1 |
| Il15ra | 0 | 186 | 335 | 0 |
| Il15ra | 0 | 244 | 193 | 1 |
| Il16 | 0 | 187 | 36 | 0 |
| Il16 | 0 | 171 | 12.3333333 | 1 |
| Il17ra | 0 | 108 | 2485 | 0 |
| Il17ra | 0 | 120.5 | 1722 | 1 |
| Il1r1 | 0 | 225 | 3348.33333 | 0 |
| Il1r1 | 0 | 222.5 | 1720.33333 | 1 |
| Il1rap | 0 | 255 | 1342 | 0 |
| Il1rap | 0 | 278 | 850.666667 | 1 |
| Il1rl1 | 0 | 97 | 17617.3333 | 0 |
| Il1rl1 | 1 | 82.5 | 7569 | 1 |
| Il1rl2 | 0 | 242 | 1483 | 0 |
| Il1rl2 | 0 | 122.5 | 904 | 1 |
| Il31ra | 0 | 182 | 9920 | 0 |
| Il31ra | 0 | 200 | 5234 | 1 |
| Il33 | 0 | 111 | 126 | 0 |
| Il33 | 0 | 152 | 201.333333 | 1 |
| Il34 | 0 | 147 | 77.6666667 | 0 |
| Il34 | 0 | 268 | 45.6666667 | 1 |
| Il6 | 0 | 23.5 | 56.6666667 | 0 |
| Il6 | 0 | 16 | 21 | 1 |
| Il6ra | 0 | 141.5 | 426.666667 | 0 |
| Il6ra | 0 | 85.5 | 273 | 1 |
| Ilk | 0 | 3 | 19810.3333 | 0 |
| Ilk | 0 | 0.5 | 27807.3333 | 1 |
| Inhbb | 0 | 11.5 | 3838.33333 | 0 |
| Inhbb | 0 | 12.5 | 1619.66667 | 1 |
| Inmt | 0 | 5 | 0.0001 | 0 |
| Inmt | 0 | 5 | 88.3333333 | 1 |
| Inpp5d | 0 | 284 | 335.333333 | 0 |
| Inpp5d | 0 | 289 | 228.333333 | 1 |
| Iqck | 0 | 0.5 | 42 | 0 |
| Iqck | 0 | 0 | 94.3333333 | 1 |
| Iqsec2 | 0 | 144 | 900.333333 | 0 |
| Iqsec2 | 0 | 152 | 1286 | 1 |
| Irak2 | 0 | 194 | 1264.33333 | 0 |
| Irak2 | 0 | 139 | 2284.33333 | 1 |
| Irf2 | 0 | 366 | 772.666667 | 0 |
| Irf2 | 0 | 256 | 503 | 1 |
| Irf2bpl | 0 | 7 | 7229 | 0 |
| Irf2bpl | 0 | 10 | 10532 | 1 |
| Isg15 | 0 | 7.5 | 319.666667 | 0 |
| Isg15 | 0 | 4.5 | 178.333333 | 1 |
| Itga2 | 0 | 195.5 | 128.666667 | 0 |
| Itga2 | 0 | 216.5 | 219 | 1 |

| | | | | |
|---|---|---|---|---|
| Itga4 | 0 | 125 | 1046 | 0 |
| Itga4 | 0 | 152.5 | 1868.33333 | 1 |
| Itgad | 0 | 30.5 | 7.66666667 | 0 |
| Itgad | 0 | 22 | 0.66673333 | 1 |
| Itgal | 0 | 119 | 221 | 0 |
| Itgal | 0 | 118 | 105.666667 | 1 |
| Itgax | 0 | 98 | 8.33336667 | 0 |
| Itgax | 0 | 86.5 | 0.66673333 | 1 |
| Itgb2 | 0 | 54 | 273.333333 | 0 |
| Itgb2 | 0 | 65.5 | 491.333333 | 1 |
| Itgb3bp | 0 | 90.5 | 8.33333333 | 0 |
| Itgb3bp | 1 | 101.5 | 0.3334 | 1 |
| Itgb7 | 0 | 45.5 | 44.6666667 | 0 |
| Itgb7 | 0 | 54 | 111.333333 | 1 |
| Itgbl1 | 0 | 182.5 | 1203 | 0 |
| Itgbl1 | 0 | 178 | 826 | 1 |
| Itih2 | 0 | 43.5 | 239.333333 | 0 |
| Itih2 | 0 | 59 | 1315.33333 | 1 |
| Itih5 | 0 | 185.5 | 244.666667 | 0 |
| Itih5 | 0 | 186.5 | 1135 | 1 |
| Itpr3 | 0 | 45.5 | 6471.33333 | 0 |
| Itpr3 | 0 | 39.5 | 4190.66667 | 1 |
| Jade2 | 0 | 87.5 | 1044 | 0 |
| Jade2 | 0 | 83 | 621.666667 | 1 |
| Jag1 | 0 | 221.5 | 8803.33333 | 0 |
| Jag1 | 1 | 218.5 | 5132 | 1 |
| Jak2 | 0 | 89.5 | 2263.66667 | 0 |
| Jak2 | 0 | 188 | 1468.33333 | 1 |
| Jcad | 0 | 93 | 5227.33333 | 0 |
| Jcad | 0 | 94 | 3587.33333 | 1 |
| Jmjd4 | 0 | 16 | 728 | 0 |
| Jmjd4 | 1 | 22 | 1233.66667 | 1 |
| Jun | 0 | 73.5 | 9256.33333 | 0 |
| Jun | 0 | 68.5 | 12529 | 1 |
| Katna1 | 0 | 111.5 | 21 | 0 |
| Katna1 | 0 | 107 | 3.00003333 | 1 |
| Kazald1 | 0 | 26 | 295.333333 | 0 |
| Kazald1 | 0 | 17.5 | 146 | 1 |
| Kbtbd11 | 0 | 124.5 | 38.6666667 | 0 |
| Kbtbd11 | 0 | 99 | 95.3333333 | 1 |
| Kcna1 | 0 | 23 | 16.3333333 | 0 |
| Kcna1 | 0 | 22 | 43 | 1 |
| Kcna2 | 0 | 19.5 | 6.66666667 | 0 |
| Kcna2 | 0 | 25 | 30.3333333 | 1 |
| Kcna3 | 0 | 4.5 | 28.6666667 | 0 |
| Kcna3 | 0 | 2.5 | 9.00003333 | 1 |
| Kcne4 | 0 | 350 | 569 | 0 |

| | | | | |
|---|---|---|---|---|
| Kcne4 | 0 | 410.5 | 149.666667 | 1 |
| Kcnf1 | 0 | 14.5 | 417 | 0 |
| Kcnf1 | 0 | 5.5 | 116 | 1 |
| Kcnh1 | 0 | 713 | 45.3333333 | 0 |
| Kcnh1 | 0 | 540 | 287 | 1 |
| Kcnj15 | 0 | 273 | 407.333333 | 0 |
| Kcnj15 | 0 | 227.5 | 259 | 1 |
| Kcnj8 | 0 | 37 | 69.3333333 | 0 |
| Kcnj8 | 0 | 41.5 | 36.6666667 | 1 |
| Kcnk10 | 0 | 86 | 59.3333333 | 0 |
| Kcnk10 | 0 | 97 | 12.6666667 | 1 |
| Kcnk5 | 0 | 102.5 | 119 | 0 |
| Kcnk5 | 0 | 82 | 221.333333 | 1 |
| Kcnn2 | 0 | 917.5 | 31 | 0 |
| Kcnn2 | 0 | 1163.5 | 64.3333333 | 1 |
| Kif11 | 0 | 92.5 | 20 | 0 |
| Kif11 | 0 | 96.5 | 57 | 1 |
| Kif1a | 0 | 147.5 | 6758 | 0 |
| Kif1a | 0 | 152.5 | 4560.66667 | 1 |
| Kif26a | 0 | 56.5 | 90.3333333 | 0 |
| Kif26a | 0 | 45 | 49.3333333 | 1 |
| Kif3c | 0 | 70 | 1143 | 0 |
| Kif3c | 0 | 78.5 | 1755.33333 | 1 |
| Kif5c | 0 | 222 | 683.333333 | 0 |
| Kif5c | 0 | 223.5 | 400.333333 | 1 |
| Kit | 0 | 167.5 | 16.6666667 | 0 |
| Kit | 0 | 147.5 | 73.6666667 | 1 |
| Klf12 | 0 | 469.5 | 215.666667 | 0 |
| Klf12 | 0 | 612.5 | 123.666667 | 1 |
| Klf13 | 0 | 32.5 | 5754.66667 | 0 |
| Klf13 | 0 | 26 | 2730 | 1 |
| Klf14 | 0 | 5 | 6.6667 | 0 |
| Klf14 | 0 | 5 | 0.66673333 | 1 |
| Klf4 | 0 | 21 | 4059.33333 | 0 |
| Klf4 | 0 | 13 | 2462 | 1 |
| Klf9 | 0 | 105.5 | 1716 | 0 |
| Klf9 | 0 | 108.5 | 2771.33333 | 1 |
| Klhdc8a | 0 | 29 | 4585.66667 | 0 |
| Klhdc8a | 0 | 21 | 1971.66667 | 1 |
| Klhl13 | 0 | 225.5 | 755 | 0 |
| Klhl13 | 0 | 245.5 | 1195.33333 | 1 |
| Klk8 | 0 | 37 | 475.666667 | 0 |
| Klk8 | 0 | 21.5 | 1104 | 1 |
| Kng1 | 0 | 57 | 3.3334 | 0 |
| Kng1 | 0 | 41.5 | 28.0000333 | 1 |
| Kng2 | 0 | 37 | 16.3333333 | 0 |
| Kng2 | 0 | 49 | 77.6666667 | 1 |

| | | | | |
|---|---|---|---|---|
| Krt13 | 0 | 8.5 | 5.66666667 | 0 |
| Krt13 | 0 | 11.5 | 64 | 1 |
| Krt14 | 0 | 10 | 50.6666667 | 0 |
| Krt14 | 0 | 11 | 17.6666667 | 1 |
| Krt19 | 0 | 11.5 | 839.666667 | 0 |
| Krt19 | 0 | 10 | 1371 | 1 |
| Krt20 | 0 | 16.5 | 83.6666667 | 0 |
| Krt20 | 0 | 14.5 | 364 | 1 |
| L1cam | 0 | 70.5 | 214.333333 | 0 |
| L1cam | 0 | 65.5 | 131 | 1 |
| L3mbtl3 | 0 | 213.5 | 732.333333 | 0 |
| L3mbtl3 | 0 | 254 | 1025 | 1 |
| Lactb | 0 | 42.5 | 704.666667 | 0 |
| Lactb | 0 | 40 | 1236.33333 | 1 |
| Lama1 | 0 | 246 | 240.333333 | 0 |
| Lama1 | 0 | 242 | 368.333333 | 1 |
| Lars2 | 0 | 172.5 | 25777.6667 | 0 |
| Lars2 | 0 | 119 | 16947 | 1 |
| Layn | 0 | 75 | 1428 | 0 |
| Layn | 0 | 67.5 | 930.333333 | 1 |
| Lce1g | 0 | 2.5 | 4.33336667 | 0 |
| Lce1g | 0 | 2.5 | 0.0001 | 1 |
| Ldb2 | 1 | 1162 | 368.666667 | 0 |
| Ldb2 | 0 | 1079 | 185.666667 | 1 |
| Ldhb | 0 | 50 | 733.666667 | 0 |
| Ldhb | 0 | 74 | 1084.33333 | 1 |
| Ldlrad4 | 0 | 938.5 | 1285 | 0 |
| Ldlrad4 | 0 | 961 | 774.333333 | 1 |
| Lef1 | 0 | 169.5 | 69.3333333 | 0 |
| Lef1 | 0 | 163 | 116.333333 | 1 |
| Leng9 | 0 | 26 | 218 | 0 |
| Leng9 | 0 | 23 | 137.666667 | 1 |
| Lepr | 0 | 176 | 88.6666667 | 0 |
| Lepr | 0 | 187 | 145.333333 | 1 |
| Lfng | 0 | 43.5 | 846.333333 | 0 |
| Lfng | 0 | 44.5 | 544 | 1 |
| Lgalsl | 0 | 20.5 | 3853.66667 | 0 |
| Lgalsl | 0 | 26 | 2571.66667 | 1 |
| Lgi3 | 0 | 19 | 17.6666667 | 0 |
| Lgi3 | 0 | 14 | 47.3333333 | 1 |
| Lgr5 | 0 | 20.5 | 3203 | 0 |
| Lgr5 | 0 | 13.5 | 296 | 1 |
| Lgr6 | 0 | 194 | 176.666667 | 0 |
| Lgr6 | 0 | 159 | 3689.33333 | 1 |
| Lhx8 | 0 | 264 | 96.3333333 | 0 |
| Lhx8 | 0 | 297 | 46.6666667 | 1 |
| Lhx9 | 0 | 75 | 1003 | 0 |

| | | | | |
|---|---|---|---|---|
| Lhx9 | 0 | 42.5 | 525.666667 | 1 |
| Limk2 | 0 | 195.5 | 3832.66667 | 0 |
| Limk2 | 0 | 254 | 2774.33333 | 1 |
| Lims2 | 0 | 27 | 1381.33333 | 0 |
| Lims2 | 0 | 21.5 | 2368.66667 | 1 |
| Lingo2 | 0 | 1719.5 | 284.666667 | 0 |
| Lingo2 | 0 | 2501 | 191.333333 | 1 |
| Lingo4 | 0 | 25 | 5 | 0 |
| Lingo4 | 0 | 20 | 0.0001 | 1 |
| Lipg | 0 | 51 | 3701.66667 | 0 |
| Lipg | 0 | 42.5 | 1685 | 1 |
| Litaf | 0 | 151.5 | 6090.66667 | 0 |
| Litaf | 0 | 148 | 4099.33333 | 1 |
| Lix1 | 0 | 120 | 15.6666667 | 0 |
| Lix1 | 0 | 125 | 4.33333333 | 1 |
| Lmna | 0 | 413.5 | 43831 | 0 |
| Lmna | 0 | 179.5 | 22275.6667 | 1 |
| Lmo4 | 0 | 43 | 2865 | 0 |
| Lmo4 | 0 | 45.5 | 4157 | 1 |
| Loxl4 | 0 | 144 | 2780 | 0 |
| Loxl4 | 0 | 189 | 4766 | 1 |
| Lpar1 | 0 | 234.5 | 1847.66667 | 0 |
| Lpar1 | 0 | 233 | 1008.66667 | 1 |
| Lpar6 | 0 | 110.5 | 864.333333 | 0 |
| Lpar6 | 0 | 119.5 | 1192.33333 | 1 |
| Lrch2 | 0 | 92.5 | 1051 | 0 |
| Lrch2 | 0 | 118.5 | 4264.66667 | 1 |
| Lrp2 | 0 | 201 | 1032.66667 | 0 |
| Lrp2 | 0 | 232 | 730 | 1 |
| Lrrc10b | 0 | 63 | 48 | 0 |
| Lrrc10b | 0 | 53.5 | 303.333333 | 1 |
| Lrrc17 | 1 | 286.5 | 6137 | 0 |
| Lrrc17 | 0 | 271.5 | 2130.66667 | 1 |
| Lrrc32 | 0 | 67 | 12623 | 0 |
| Lrrc32 | 0 | 120.5 | 31525.3333 | 1 |
| Lrrc4 | 0 | 526 | 16.6666667 | 0 |
| Lrrc4 | 0 | 586.5 | 4.33333333 | 1 |
| Lrrc4c | 0 | 2473 | 262 | 0 |
| Lrrc4c | 0 | 3710 | 93.3333333 | 1 |
| Lrrc55 | 0 | 11.5 | 19 | 0 |
| Lrrc55 | 0 | 21 | 6.00003333 | 1 |
| Lrrc75b | 0 | 26 | 193 | 0 |
| Lrrc75b | 0 | 21 | 67 | 1 |
| Lrrc8e | 0 | 18.5 | 221 | 0 |
| Lrrc8e | 0 | 17 | 125 | 1 |
| Lrrn2 | 0 | 106.5 | 269.666667 | 0 |
| Lrrn2 | 0 | 93 | 126.333333 | 1 |

| | | | | |
|---|---|---|---|---|
| Lrrn3 | 0 | 1524.5 | 349.666667 | 0 |
| Lrrn3 | 0 | 2203.5 | 172 | 1 |
| Lrrn4cl | 0 | 38 | 400.333333 | 0 |
| Lrrn4cl | 0 | 17 | 874.666667 | 1 |
| Lrrtm1 | 0 | 1528 | 80.3333333 | 0 |
| Lrrtm1 | 0 | 1882 | 39 | 1 |
| Lrrtm2 | 0 | 368 | 126.666667 | 0 |
| Lrrtm2 | 0 | 442 | 26.6666667 | 1 |
| Lrrtm3 | 1 | 1249.5 | 161 | 0 |
| Lrrtm3 | 1 | 1513 | 72.3333333 | 1 |
| Ltbp3 | 0 | 59 | 12378.6667 | 0 |
| Ltbp3 | 0 | 65.5 | 20469.6667 | 1 |
| Lum | 0 | 17 | 2626 | 0 |
| Lum | 0 | 19 | 1185 | 1 |
| Luzp2 | 0 | 617 | 61.3333333 | 0 |
| Luzp2 | 0 | 717 | 100.666667 | 1 |
| Lvrn | 0 | 89.5 | 283.333333 | 0 |
| Lvrn | 0 | 78.5 | 688.333333 | 1 |
| Lxn | 0 | 95 | 2504 | 0 |
| Lxn | 0 | 93 | 1764.33333 | 1 |
| Ly6a | 0 | 10 | 2923.66667 | 0 |
| Ly6a | 0 | 11 | 882.333333 | 1 |
| Ly6c1 | 0 | 4 | 426.666667 | 0 |
| Ly6c1 | 0 | 1.5 | 114.666667 | 1 |
| Ly6i | 0 | 8.5 | 32.3333333 | 0 |
| Ly6i | 0 | 15 | 8 | 1 |
| Lypd1 | 0 | 743 | 65.3333333 | 0 |
| Lypd1 | 0 | 904 | 158.666667 | 1 |
| Mafb | 1 | 6.5 | 562 | 0 |
| Mafb | 0 | 6.5 | 801.333333 | 1 |
| Mageb3 | 0 | 93.5 | 2.6667 | 0 |
| Mageb3 | 0 | 110.5 | 15 | 1 |
| Magi3 | 1 | 339.5 | 1256.33333 | 0 |
| Magi3 | 0 | 409.5 | 1933.66667 | 1 |
| Mamdc2 | 0 | 166.5 | 655 | 0 |
| Mamdc2 | 0 | 221.5 | 1494 | 1 |
| Map1a | 0 | 112.5 | 6352.33333 | 0 |
| Map1a | 0 | 94 | 4109.33333 | 1 |
| Map2k3 | 0 | 61 | 8122.33333 | 0 |
| Map2k3 | 0 | 47 | 4554 | 1 |
| Map2k3os | 0 | 23.5 | 207.666667 | 0 |
| Map2k3os | 0 | 34 | 121.333333 | 1 |
| Map4k3 | 0 | 285.5 | 4850.66667 | 0 |
| Map4k3 | 0 | 289 | 3269.33333 | 1 |
| Map6 | 0 | 111.5 | 6702 | 0 |
| Map6 | 0 | 103 | 4132 | 1 |
| Marcksl1 | 0 | 29 | 19750.6667 | 0 |

| | | | | |
|---|---|---|---|---|
| Marcksl1 | 0 | 30 | 26920.6667 | 1 |
| Mark1 | 0 | 107.5 | 878 | 0 |
| Mark1 | 0 | 118 | 1361.33333 | 1 |
| Masp1 | 0 | 182 | 6598.33333 | 0 |
| Masp1 | 0 | 254 | 13040.3333 | 1 |
| Mbd1 | 0 | 35 | 2401 | 0 |
| Mbd1 | 0 | 76 | 5161.33333 | 1 |
| Mbnl3 | 0 | 596 | 294.666667 | 0 |
| Mbnl3 | 0 | 504 | 445 | 1 |
| Mcam | 0 | 157 | 8576 | 0 |
| Mcam | 0 | 125 | 5917 | 1 |
| Mcf2l | 0 | 144.5 | 351 | 0 |
| Mcf2l | 0 | 143 | 210.333333 | 1 |
| Mchr1 | 0 | 11 | 19.6667 | 0 |
| Mchr1 | 0 | 10.5 | 4.6667 | 1 |
| Mcpt8 | 0 | 6 | 189 | 0 |
| Mcpt8 | 0 | 6 | 501 | 1 |
| Mcpt9 | 0 | 1 | 4 | 0 |
| Mcpt9 | 0 | 0 | 17 | 1 |
| Mdfi | 0 | 78 | 239.333333 | 0 |
| Mdfi | 0 | 56 | 352.666667 | 1 |
| Me1 | 0 | 128.5 | 3887.66667 | 0 |
| Me1 | 0 | 168 | 5279.66667 | 1 |
| Me3 | 0 | 226 | 225.666667 | 0 |
| Me3 | 0 | 236 | 347.333333 | 1 |
| Medag | 0 | 41.5 | 1072.33333 | 0 |
| Medag | 0 | 62 | 2829 | 1 |
| Megf10 | 0 | 229.5 | 440.333333 | 0 |
| Megf10 | 0 | 285.5 | 266 | 1 |
| Megf6 | 0 | 132 | 2699.33333 | 0 |
| Megf6 | 0 | 120.5 | 1046.66667 | 1 |
| Meis2 | 0 | 1035 | 1556.33333 | 0 |
| Meis2 | 0 | 1500.5 | 2491.66667 | 1 |
| Meox1 | 0 | 50.5 | 229.333333 | 0 |
| Meox1 | 0 | 39 | 37 | 1 |
| Mest | 0 | 17.5 | 5631.66667 | 0 |
| Mest | 0 | 16.5 | 2087.66667 | 1 |
| Metrnl | 0 | 39.5 | 3192 | 0 |
| Metrnl | 1 | 59.5 | 8891.33333 | 1 |
| Mfap3l | 0 | 99.5 | 617.666667 | 0 |
| Mfap3l | 0 | 113.5 | 336 | 1 |
| Mgll | 0 | 152 | 5960.33333 | 0 |
| Mgll | 0 | 128.5 | 3578 | 1 |
| Mical2 | 0 | 478 | 23245.6667 | 0 |
| Mical2 | 0 | 234.5 | 16028.6667 | 1 |
| Mical3 | 0 | 165.5 | 1620 | 0 |
| Mical3 | 0 | 176.5 | 2356.66667 | 1 |

| | | | | |
|---|---|---|---|---|
| Mid1 | 0 | 429 | 854 | 0 |
| Mid1 | 0 | 573.5 | 1675.66667 | 1 |
| Mir27a | 0 | 1.5 | 1.00006667 | 0 |
| Mir27a | 0 | 2.5 | 8.66666667 | 1 |
| Mir450b | 0 | 5 | 8.00006667 | 0 |
| Mir450b | 0 | 6 | 0.0001 | 1 |
| Mir762 | 0 | 108.5 | 705.333333 | 0 |
| Mir762 | 0 | 121 | 137.3334 | 1 |
| Mitf | 0 | 427.5 | 174 | 0 |
| Mitf | 0 | 508 | 112.666667 | 1 |
| Mkx | 0 | 259 | 100.666667 | 0 |
| Mkx | 1 | 316 | 182.333333 | 1 |
| Mmp13 | 0 | 15.5 | 93.3333333 | 0 |
| Mmp13 | 0 | 17 | 444.666667 | 1 |
| Mmp17 | 0 | 116.5 | 1650 | 0 |
| Mmp17 | 0 | 111.5 | 2519 | 1 |
| Mmp9 | 0 | 20.5 | 389.666667 | 0 |
| Mmp9 | 0 | 13 | 1001.33333 | 1 |
| Mn1 | 0 | 124 | 2240.33333 | 0 |
| Mn1 | 0 | 106.5 | 1623 | 1 |
| Mocos | 0 | 53 | 123.333333 | 0 |
| Mocos | 0 | 55.5 | 225.666667 | 1 |
| Morc4 | 0 | 221 | 1091.66667 | 0 |
| Morc4 | 0 | 194.5 | 1597 | 1 |
| Mov10 | 0 | 84.5 | 1289.33333 | 0 |
| Mov10 | 0 | 81.5 | 1969 | 1 |
| Mpp1 | 0 | 38.5 | 2375.66667 | 0 |
| Mpp1 | 0 | 50.5 | 3484.33333 | 1 |
| Mpped2 | 0 | 326 | 174 | 0 |
| Mpped2 | 0 | 375 | 324.666667 | 1 |
| Mpzl2 | 0 | 19 | 79 | 0 |
| Mpzl2 | 0 | 11 | 150 | 1 |
| Mras | 0 | 85.5 | 2164.66667 | 0 |
| Mras | 0 | 99.5 | 4571 | 1 |
| Mrc1 | 0 | 81 | 379 | 0 |
| Mrc1 | 0 | 84 | 208.333333 | 1 |
| Mrc2 | 0 | 335 | 17906.6667 | 0 |
| Mrc2 | 0 | 465 | 36462.3333 | 1 |
| Mrgprf | 0 | 28 | 1406.33333 | 0 |
| Mrgprf | 0 | 19 | 536.333333 | 1 |
| Mrvi1 | 0 | 123.5 | 774.666667 | 0 |
| Mrvi1 | 0 | 137.5 | 1180.33333 | 1 |
| Ms4a6b | 0 | 27 | 43 | 0 |
| Ms4a6b | 0 | 37.5 | 14 | 1 |
| Ms4a6c | 0 | 57 | 88 | 0 |
| Ms4a6c | 0 | 61.5 | 33 | 1 |
| Ms4a6d | 0 | 24 | 94.3333333 | 0 |

| | | | | |
|---|---|---|---|---|
| Ms4a6d | 0 | 27 | 48.3333333 | 1 |
| Ms4a7 | 0 | 33 | 109.333333 | 0 |
| Ms4a7 | 0 | 43.5 | 27 | 1 |
| Msx1 | 0 | 2.5 | 956 | 0 |
| Msx1 | 1 | 3 | 507 | 1 |
| Mvp | 0 | 49 | 3418.33333 | 0 |
| Mvp | 0 | 47 | 2321 | 1 |
| Mxd4 | 0 | 128.5 | 3220.66667 | 0 |
| Mxd4 | 0 | 109.5 | 2313.66667 | 1 |
| Mxra7 | 0 | 99.5 | 3917.66667 | 0 |
| Mxra7 | 0 | 71 | 2499 | 1 |
| Mybl1 | 0 | 112 | 1844 | 0 |
| Mybl1 | 0 | 156 | 3280.33333 | 1 |
| Mycl | 0 | 24.5 | 332.333333 | 0 |
| Mycl | 0 | 34 | 553 | 1 |
| Mycn | 0 | 15.5 | 95 | 0 |
| Mycn | 0 | 11 | 50.3333333 | 1 |
| Myh1 | 0 | 52.5 | 130 | 0 |
| Myh1 | 0 | 41.5 | 32.3333333 | 1 |
| Myh11 | 0 | 215 | 2804 | 0 |
| Myh11 | 0 | 198 | 1592.66667 | 1 |
| Myh2 | 0 | 29 | 38 | 0 |
| Myh2 | 0 | 24 | 7 | 1 |
| Myl12b | 0 | 50.5 | 34.3333333 | 0 |
| Myl12b | 0 | 66 | 70.6666667 | 1 |
| Myo18b | 0 | 192.5 | 4.66666667 | 0 |
| Myo18b | 0 | 155 | 0.0001 | 1 |
| Myo1e | 0 | 292.5 | 4256 | 0 |
| Myo1e | 0 | 287 | 3050.33333 | 1 |
| Myo1f | 0 | 118 | 221 | 0 |
| Myo1f | 0 | 96.5 | 132.666667 | 1 |
| Myo7a | 0 | 234.5 | 1790.33333 | 0 |
| Myo7a | 1 | 331.5 | 1165.66667 | 1 |
| Myocd | 0 | 228 | 206.333333 | 0 |
| Myocd | 0 | 271 | 80.6666667 | 1 |
| Nab2 | 0 | 96 | 10101.3333 | 0 |
| Nab2 | 0 | 47 | 6878.66667 | 1 |
| Naip1 | 0 | 83.5 | 0.0001 | 0 |
| Naip1 | 0 | 96 | 18.3333333 | 1 |
| Nap1l5 | 0 | 142 | 81.6666667 | 0 |
| Nap1l5 | 0 | 164.5 | 33 | 1 |
| Nav2 | 0 | 838 | 3109.66667 | 0 |
| Nav2 | 0 | 894.5 | 4307.66667 | 1 |
| Ncam2 | 0 | 950.5 | 48.3333333 | 0 |
| Ncam2 | 0 | 777 | 23.6666667 | 1 |
| Ncoa3 | 0 | 199.5 | 1677 | 0 |
| Ncoa3 | 0 | 238 | 2981 | 1 |

| | | | | |
|---|---|---|---|---|
| Ncor2 | 0 | 487.5 | 19300.3333 | 0 |
| Ncor2 | 0 | 496 | 13016.3333 | 1 |
| Ndnf | 0 | 76.5 | 996.666667 | 0 |
| Ndnf | 0 | 86.5 | 508 | 1 |
| Ndrg1 | 0 | 102.5 | 1471 | 0 |
| Ndrg1 | 0 | 89.5 | 411 | 1 |
| Ndrg4 | 0 | 26 | 2267.66667 | 0 |
| Ndrg4 | 0 | 19 | 1312 | 1 |
| Ndst3 | 0 | 237.5 | 158.666667 | 0 |
| Ndst3 | 0 | 248.5 | 100.333333 | 1 |
| Neat1 | 0 | 32.5 | 4898.66667 | 0 |
| Neat1 | 1 | 48.5 | 3290.33333 | 1 |
| Necab1 | 0 | 176 | 4.33333333 | 0 |
| Necab1 | 0 | 197.5 | 0.0001 | 1 |
| Nectin2 | 0 | 110 | 5228.33333 | 0 |
| Nectin2 | 0 | 111.5 | 7271.66667 | 1 |
| Nefl | 0 | 14 | 22.3333333 | 0 |
| Nefl | 0 | 11.5 | 53 | 1 |
| Nefm | 0 | 10 | 42 | 0 |
| Nefm | 0 | 6 | 88.3333333 | 1 |
| Nek2 | 0 | 88.5 | 43 | 0 |
| Nek2 | 0 | 80 | 74 | 1 |
| Nek7 | 0 | 489 | 4281 | 0 |
| Nek7 | 0 | 345.5 | 2609.33333 | 1 |
| Nell1 | 0 | 493 | 14.6666667 | 0 |
| Nell1 | 0 | 565 | 4 | 1 |
| Nell2 | 0 | 363.5 | 381 | 0 |
| Nell2 | 0 | 425 | 771.666667 | 1 |
| Nes | 0 | 86 | 9007.66667 | 0 |
| Nes | 0 | 128.5 | 17261.6667 | 1 |
| Neurl1b | 0 | 52.5 | 169.666667 | 0 |
| Neurl1b | 0 | 58 | 99.3333333 | 1 |
| Neurl3 | 0 | 15 | 59.3333333 | 0 |
| Neurl3 | 0 | 15 | 120.666667 | 1 |
| Nfatc2 | 0 | 647 | 110.333333 | 0 |
| Nfatc2 | 0 | 358 | 66.3333333 | 1 |
| Nfatc4 | 0 | 49 | 5017.66667 | 0 |
| Nfatc4 | 0 | 27.5 | 2616 | 1 |
| Nfkbiz | 0 | 203 | 1566 | 0 |
| Nfkbiz | 0 | 221 | 2144.33333 | 1 |
| Ngef | 0 | 153.5 | 99.6666667 | 0 |
| Ngef | 0 | 136.5 | 19.6666667 | 1 |
| Ngf | 0 | 249.5 | 923.333333 | 0 |
| Ngf | 0 | 137 | 429 | 1 |
| Ngfr | 0 | 41.5 | 364.666667 | 0 |
| Ngfr | 0 | 38.5 | 632.666667 | 1 |
| Nhs | 0 | 701 | 299.333333 | 0 |

| | | | | |
|---|---|---|---|---|
| Nhs | 0 | 610 | 197.666667 | 1 |
| Nhsl1 | 0 | 224 | 970 | 0 |
| Nhsl1 | 0 | 219.5 | 1334.33333 | 1 |
| Nid1 | 0 | 203 | 51642.3333 | 0 |
| Nid1 | 0 | 342 | 83552 | 1 |
| Nid2 | 0 | 93 | 9049.33333 | 0 |
| Nid2 | 0 | 94 | 12754 | 1 |
| Nipal1 | 0 | 53 | 1053.66667 | 0 |
| Nipal1 | 0 | 64 | 593.666667 | 1 |
| Nkain1 | 0 | 84 | 39.6666667 | 0 |
| Nkain1 | 0 | 87 | 66.3333333 | 1 |
| Nkain4 | 0 | 50.5 | 102 | 0 |
| Nkain4 | 0 | 28.5 | 62 | 1 |
| Nkd2 | 0 | 125.5 | 1058.33333 | 0 |
| Nkd2 | 0 | 123.5 | 601 | 1 |
| Nlrc5 | 0 | 29.5 | 72.6666667 | 0 |
| Nlrc5 | 0 | 26.5 | 31.3333333 | 1 |
| Nmnat2 | 0 | 275.5 | 334 | 0 |
| Nmnat2 | 0 | 280 | 213.333333 | 1 |
| Noct | 0 | 71.5 | 2416 | 0 |
| Noct | 0 | 75.5 | 3621.66667 | 1 |
| Nod1 | 0 | 85.5 | 795.666667 | 0 |
| Nod1 | 0 | 88 | 276.333333 | 1 |
| Nog | 1 | 3.5 | 703 | 0 |
| Nog | 0 | 1 | 345.333333 | 1 |
| Nos1 | 0 | 325 | 26.6666667 | 0 |
| Nos1 | 0 | 317.5 | 3.3334 | 1 |
| Notch1 | 1 | 41.5 | 4543.66667 | 0 |
| Notch1 | 0 | 39.5 | 2801.33333 | 1 |
| Notch4 | 0 | 27 | 11 | 0 |
| Notch4 | 0 | 32.5 | 32 | 1 |
| Nov | 0 | 25.5 | 7561.66667 | 0 |
| Nov | 0 | 22 | 5417.66667 | 1 |
| Npnt | 0 | 199.5 | 3416 | 0 |
| Npnt | 0 | 214.5 | 1603.66667 | 1 |
| Nppb | 0 | 11 | 150.333333 | 0 |
| Nppb | 0 | 2 | 84 | 1 |
| Npr1 | 0 | 31.5 | 23 | 0 |
| Npr1 | 0 | 27.5 | 63.3333333 | 1 |
| Npr3 | 0 | 256.5 | 1442.66667 | 0 |
| Npr3 | 0 | 319 | 4628 | 1 |
| Nptx1 | 0 | 18.5 | 117.333333 | 0 |
| Nptx1 | 0 | 19 | 31.3333333 | 1 |
| Npy1r | 0 | 33 | 159 | 0 |
| Npy1r | 0 | 24.5 | 88.6666667 | 1 |
| Npy2r | 0 | 37 | 9.6667 | 0 |
| Npy2r | 0 | 25 | 47.6666667 | 1 |

| | | | | |
|---|---|---|---|---|
| Nr4a1 | 0 | 109.5 | 5354 | 0 |
| Nr4a1 | 0 | 52.5 | 2514 | 1 |
| Nr4a3 | 0 | 90.5 | 155.666667 | 0 |
| Nr4a3 | 0 | 90 | 74 | 1 |
| Nrarp | 0 | 6.5 | 147 | 0 |
| Nrarp | 0 | 4.5 | 31.3333333 | 1 |
| Nrcam | 0 | 774 | 131.333333 | 0 |
| Nrcam | 0 | 974 | 388 | 1 |
| Nrep | 0 | 88 | 1069.33333 | 0 |
| Nrep | 1 | 120 | 1501 | 1 |
| Nrk | 0 | 606 | 1223 | 0 |
| Nrk | 0 | 748 | 2531.66667 | 1 |
| Nrxn1 | 0 | 1500 | 8 | 0 |
| Nrxn1 | 0 | 2235 | 30.3333333 | 1 |
| Nt5e | 0 | 119 | 294.333333 | 0 |
| Nt5e | 0 | 114 | 428.666667 | 1 |
| Ntn4 | 0 | 29 | 1632.33333 | 0 |
| Ntn4 | 0 | 28 | 3654.33333 | 1 |
| Ntng2 | 0 | 198 | 43 | 0 |
| Ntng2 | 0 | 153 | 76.3333333 | 1 |
| Ntrk3 | 0 | 1627 | 235 | 0 |
| Ntrk3 | 0 | 1384 | 111 | 1 |
| Nupr1 | 1 | 17 | 2066.33333 | 0 |
| Nupr1 | 0 | 19.5 | 4001.66667 | 1 |
| Ogn | 0 | 75.5 | 4577.66667 | 0 |
| Ogn | 0 | 79.5 | 28618 | 1 |
| Olfm1 | 0 | 67 | 1161.33333 | 0 |
| Olfm1 | 0 | 88 | 761.333333 | 1 |
| Olfm2 | 0 | 168.5 | 380 | 0 |
| Olfm2 | 0 | 142 | 868 | 1 |
| Olfml2a | 0 | 59.5 | 263 | 0 |
| Olfml2a | 0 | 46.5 | 143.333333 | 1 |
| Olfml2b | 0 | 195 | 4700 | 0 |
| Olfml2b | 0 | 193 | 2531 | 1 |
| Olfr1033 | 0 | 14 | 46.3333333 | 0 |
| Olfr1033 | 0 | 17 | 205.666667 | 1 |
| Olr1 | 0 | 42 | 154.666667 | 0 |
| Olr1 | 0 | 128 | 690.333333 | 1 |
| Ophn1 | 0 | 887.5 | 288.333333 | 0 |
| Ophn1 | 0 | 751 | 440 | 1 |
| Orai2 | 0 | 41 | 739.333333 | 0 |
| Orai2 | 0 | 37.5 | 437.333333 | 1 |
| Otogl | 0 | 174 | 6.66666667 | 0 |
| Otogl | 0 | 184.5 | 25.6666667 | 1 |
| Otud7a | 0 | 458.5 | 104.666667 | 0 |
| Otud7a | 0 | 524 | 53.6666667 | 1 |
| Otud7b | 0 | 97 | 2373 | 0 |

| | | | | |
|---|---|---|---|---|
| Otud7b | 0 | 139.5 | 3340.66667 | 1 |
| P2rx7 | 0 | 60 | 134.666667 | 0 |
| P2rx7 | 0 | 65.5 | 80 | 1 |
| P2ry1 | 0 | 27 | 36.3333333 | 0 |
| P2ry1 | 0 | 27 | 127 | 1 |
| P2ry10b | 0 | 36 | 623.333333 | 0 |
| P2ry10b | 0 | 49.5 | 312 | 1 |
| P3h2 | 0 | 308 | 175 | 0 |
| P3h2 | 0 | 389.5 | 352.333333 | 1 |
| P4ha1 | 0 | 136 | 7099 | 0 |
| P4ha1 | 0 | 169 | 9694.66667 | 1 |
| P4ha3 | 0 | 103 | 5911 | 0 |
| P4ha3 | 0 | 106.5 | 2903.66667 | 1 |
| Pacs2 | 0 | 124.5 | 5440 | 0 |
| Pacs2 | 0 | 164 | 3844.33333 | 1 |
| Pag1 | 0 | 219 | 1053 | 0 |
| Pag1 | 0 | 481 | 2719.66667 | 1 |
| Pak6 | 0 | 88 | 6 | 0 |
| Pak6 | 0 | 80 | 36.6666667 | 1 |
| Papss2 | 0 | 210.5 | 1814.33333 | 0 |
| Papss2 | 0 | 269.5 | 1186.66667 | 1 |
| Paqr8 | 0 | 90.5 | 671 | 0 |
| Paqr8 | 0 | 95 | 1041.66667 | 1 |
| Parp3 | 0 | 63 | 3972.66667 | 0 |
| Parp3 | 0 | 39 | 2504.66667 | 1 |
| Parp8 | 0 | 208.5 | 1457.33333 | 0 |
| Parp8 | 0 | 346.5 | 2492.66667 | 1 |
| Pbx1 | 0 | 1735 | 3775.33333 | 0 |
| Pbx1 | 0 | 1824 | 5506.33333 | 1 |
| Pcdh10 | 0 | 77 | 609 | 0 |
| Pcdh10 | 0 | 87 | 1583 | 1 |
| Pcdh11x | 0 | 39 | 540.333333 | 0 |
| Pcdh11x | 0 | 41 | 923.666667 | 1 |
| Pcdh17 | 0 | 31.5 | 100.666667 | 0 |
| Pcdh17 | 0 | 33 | 192.666667 | 1 |
| Pcdh20 | 0 | 13.5 | 152.333333 | 0 |
| Pcdh20 | 0 | 13 | 62 | 1 |
| Pcdh8 | 0 | 2 | 10 | 0 |
| Pcdh8 | 0 | 5 | 1.6667 | 1 |
| Pcdh9 | 0 | 2969 | 2923.33333 | 0 |
| Pcdh9 | 0 | 2792 | 4369.66667 | 1 |
| Pcolce2 | 0 | 129 | 888.666667 | 0 |
| Pcolce2 | 0 | 131.5 | 566 | 1 |
| Pcp4l1 | 0 | 52 | 500.333333 | 0 |
| Pcp4l1 | 0 | 44.5 | 203.666667 | 1 |
| Pcx | 0 | 268.5 | 1018.33333 | 0 |
| Pcx | 0 | 283.5 | 1560 | 1 |

| | | | | |
|---|---|---|---|---|
| Pcyt1b | 0 | 168.5 | 291 | 0 |
| Pcyt1b | 0 | 179.5 | 198 | 1 |
| Pde4b | 0 | 718 | 1109.66667 | 0 |
| Pde4b | 0 | 831.5 | 714.333333 | 1 |
| Pde4d | 0 | 625 | 1077.66667 | 0 |
| Pde4d | 0 | 744 | 567 | 1 |
| Pde5a | 0 | 87.5 | 638.333333 | 0 |
| Pde5a | 0 | 90.5 | 1008.66667 | 1 |
| Pde8b | 0 | 322 | 1215.33333 | 0 |
| Pde8b | 0 | 409.5 | 2570.33333 | 1 |
| Pdgfb | 0 | 15.5 | 144.333333 | 0 |
| Pdgfb | 0 | 13.5 | 223.666667 | 1 |
| Pdgfd | 0 | 233 | 1049 | 0 |
| Pdgfd | 0 | 149 | 1446.33333 | 1 |
| Pdgfra | 0 | 75 | 11567.6667 | 0 |
| Pdgfra | 0 | 91.5 | 19818.6667 | 1 |
| Pdgfrl | 0 | 147.5 | 1663 | 0 |
| Pdgfrl | 0 | 159 | 3272 | 1 |
| Pdlim7 | 0 | 35 | 12647.3333 | 0 |
| Pdlim7 | 1 | 48.5 | 21007 | 1 |
| Pdzd2 | 0 | 241.5 | 538.666667 | 0 |
| Pdzd2 | 0 | 289 | 312.666667 | 1 |
| Pdzrn3 | 1 | 765 | 5862.33333 | 0 |
| Pdzrn3 | 1 | 841 | 4157.33333 | 1 |
| Pdzrn4 | 0 | 655.5 | 4.00003333 | 0 |
| Pdzrn4 | 0 | 745 | 18.3333333 | 1 |
| Peli2 | 0 | 81 | 730 | 0 |
| Peli2 | 0 | 76.5 | 261 | 1 |
| Penk | 0 | 18 | 2248.66667 | 0 |
| Penk | 0 | 10.5 | 494.666667 | 1 |
| Per1 | 0 | 51 | 1922 | 0 |
| Per1 | 0 | 49 | 1393.66667 | 1 |
| Per2 | 0 | 68 | 480 | 0 |
| Per2 | 0 | 51.5 | 800.333333 | 1 |
| Pex26 | 0 | 46 | 361 | 0 |
| Pex26 | 0 | 42.5 | 536 | 1 |
| Pfkfb3 | 0 | 346 | 1194.33333 | 0 |
| Pfkfb3 | 0 | 277 | 1661.33333 | 1 |
| Pgam2 | 0 | 21 | 29.6666667 | 0 |
| Pgam2 | 0 | 13.5 | 3.66673333 | 1 |
| Pgap1 | 0 | 286 | 418.666667 | 0 |
| Pgap1 | 0 | 330.5 | 773.333333 | 1 |
| Pgpep1 | 0 | 53.5 | 1135.66667 | 0 |
| Pgpep1 | 0 | 56.5 | 2355.66667 | 1 |
| Phactr1 | 0 | 873.5 | 84.6666667 | 0 |
| Phactr1 | 0 | 936.5 | 44.6666667 | 1 |
| Pi15 | 0 | 51.5 | 858.666667 | 0 |

| | | | | |
|---|---|---|---|---|
| Pi15 | 0 | 60.5 | 582.333333 | 1 |
| Pid1 | 0 | 264 | 623.666667 | 0 |
| Pid1 | 0 | 315.5 | 1023 | 1 |
| Piezo2 | 0 | 398.5 | 2304.33333 | 0 |
| Piezo2 | 0 | 581.5 | 6005.33333 | 1 |
| Pik3cb | 0 | 123 | 913 | 0 |
| Pik3cb | 0 | 137 | 1458.33333 | 1 |
| Pik3ip1 | 0 | 61 | 550.666667 | 0 |
| Pik3ip1 | 0 | 62 | 326.333333 | 1 |
| Pik3r5 | 0 | 103.5 | 169.666667 | 0 |
| Pik3r5 | 0 | 106 | 350.333333 | 1 |
| Pim1 | 0 | 24 | 1553 | 0 |
| Pim1 | 0 | 25 | 6635 | 1 |
| Pim3 | 0 | 10.5 | 3701.33333 | 0 |
| Pim3 | 0 | 10 | 5262 | 1 |
| Pir | 0 | 159 | 121.666667 | 0 |
| Pir | 0 | 102 | 76.6666667 | 1 |
| Pitpnc1 | 0 | 361.5 | 918.666667 | 0 |
| Pitpnc1 | 0 | 764 | 3711.66667 | 1 |
| Pknox2 | 0 | 569 | 476.333333 | 0 |
| Pknox2 | 1 | 389 | 336.666667 | 1 |
| Pkp1 | 0 | 160.5 | 158 | 0 |
| Pkp1 | 0 | 133.5 | 19.3333333 | 1 |
| Pkp2 | 0 | 88 | 817.333333 | 0 |
| Pkp2 | 0 | 68.5 | 1573.33333 | 1 |
| Pla2g16 | 0 | 87 | 642 | 0 |
| Pla2g16 | 0 | 103.5 | 404 | 1 |
| Pla2g7 | 0 | 28 | 919.333333 | 0 |
| Pla2g7 | 0 | 21.5 | 524.666667 | 1 |
| Plau | 0 | 24.5 | 5490.33333 | 0 |
| Plau | 0 | 26.5 | 10254.3333 | 1 |
| Plb1 | 0 | 297.5 | 27 | 0 |
| Plb1 | 0 | 276 | 8 | 1 |
| Plcd3 | 0 | 100.5 | 733.333333 | 0 |
| Plcd3 | 0 | 78 | 1152 | 1 |
| Plce1 | 0 | 306 | 717.666667 | 0 |
| Plce1 | 0 | 354 | 465.333333 | 1 |
| Plch1 | 0 | 617 | 35.3333333 | 0 |
| Plch1 | 0 | 600 | 67.6666667 | 1 |
| Plcxd1 | 0 | 6 | 38 | 0 |
| Plcxd1 | 0 | 3.5 | 14 | 1 |
| Plcxd2 | 1 | 124.5 | 962 | 0 |
| Plcxd2 | 0 | 133 | 684.666667 | 1 |
| Pld6 | 0 | 74 | 0.0001 | 0 |
| Pld6 | 0 | 103 | 6 | 1 |
| Plec | 0 | 546.5 | 46450 | 0 |
| Plec | 0 | 452.5 | 27272 | 1 |

| | | | | |
|---|---|---|---|---|
| Plek2 | 0 | 38 | 5.66666667 | 0 |
| Plek2 | 0 | 32 | 0.66673333 | 1 |
| Plekha2 | 0 | 93.5 | 1202.33333 | 0 |
| Plekha2 | 0 | 78 | 2333 | 1 |
| Plekha4 | 0 | 36 | 236 | 0 |
| Plekha4 | 0 | 32 | 97.6666667 | 1 |
| Plekha6 | 0 | 23 | 122.666667 | 0 |
| Plekha6 | 0 | 36.5 | 410.333333 | 1 |
| Plekha7 | 0 | 349.5 | 223.666667 | 0 |
| Plekha7 | 0 | 333.5 | 677.333333 | 1 |
| Plekhb1 | 0 | 66.5 | 22.6666667 | 0 |
| Plekhb1 | 0 | 113 | 49 | 1 |
| Plekhg1 | 0 | 324.5 | 159.333333 | 0 |
| Plekhg1 | 1 | 394 | 685.333333 | 1 |
| Plekhg4 | 0 | 69 | 165.333333 | 0 |
| Plekhg4 | 0 | 55 | 98 | 1 |
| Plin2 | 0 | 27 | 2282 | 0 |
| Plin2 | 0 | 25.5 | 3430.33333 | 1 |
| Plin4 | 0 | 51 | 72.6666667 | 0 |
| Plin4 | 0 | 48.5 | 12.6666667 | 1 |
| Plk2 | 0 | 35.5 | 9100 | 0 |
| Plk2 | 0 | 75.5 | 14029 | 1 |
| Plpp1 | 1 | 302 | 8282.33333 | 0 |
| Plpp1 | 0 | 359 | 4102 | 1 |
| Plppr4 | 0 | 73 | 165.666667 | 0 |
| Plppr4 | 0 | 74.5 | 76.6666667 | 1 |
| Plscr1 | 0 | 16.5 | 355.666667 | 0 |
| Plscr1 | 1 | 19 | 715 | 1 |
| Plscr2 | 0 | 85 | 171.666667 | 0 |
| Plscr2 | 0 | 131 | 695.666667 | 1 |
| Plxna2 | 0 | 237 | 1583.33333 | 0 |
| Plxna2 | 0 | 264.5 | 913.666667 | 1 |
| Plxnc1 | 0 | 251.5 | 826.333333 | 0 |
| Plxnc1 | 1 | 269 | 310.666667 | 1 |
| Pmaip1 | 0 | 19.5 | 761.333333 | 0 |
| Pmaip1 | 0 | 20.5 | 1198.33333 | 1 |
| Pmepa1 | 0 | 115 | 12192.6667 | 0 |
| Pmepa1 | 0 | 176 | 22887 | 1 |
| Pml | 0 | 49 | 3680 | 0 |
| Pml | 0 | 54.5 | 7104.33333 | 1 |
| Pmp22 | 1 | 108.5 | 6174 | 0 |
| Pmp22 | 0 | 153 | 12119.3333 | 1 |
| Podnl1 | 0 | 25 | 544.666667 | 0 |
| Podnl1 | 0 | 26.5 | 345 | 1 |
| Pop1 | 0 | 152.5 | 1570.33333 | 0 |
| Pop1 | 0 | 148 | 2430.66667 | 1 |
| Popdc3 | 0 | 53 | 16.6666667 | 0 |

| | | | | |
|---|---|---|---|---|
| Popdc3 | 0 | 49.5 | 4.66666667 | 1 |
| Postn | 1 | 464 | 82636.3333 | 0 |
| Postn | 0 | 416.5 | 58845.3333 | 1 |
| Ppbp | 0 | 5 | 465 | 0 |
| Ppbp | 0 | 2.5 | 1176.66667 | 1 |
| Ppl | 0 | 75 | 166 | 0 |
| Ppl | 0 | 67.5 | 106 | 1 |
| Ppm1h | 0 | 2 | 199.333333 | 0 |
| Ppm1h | 0 | 1.5 | 545.333333 | 1 |
| Ppm1j | 0 | 14.5 | 136.666667 | 0 |
| Ppm1j | 0 | 14 | 230.666667 | 1 |
| Ppp1r12b | 0 | 454.5 | 1367.66667 | 0 |
| Ppp1r12b | 0 | 454.5 | 3470.33333 | 1 |
| Ppp1r3c | 0 | 18.5 | 39.6666667 | 0 |
| Ppp1r3c | 1 | 17 | 9 | 1 |
| Ppp1r3d | 0 | 4 | 331 | 0 |
| Ppp1r3d | 0 | 4 | 226.333333 | 1 |
| Prag1 | 0 | 70 | 1133.66667 | 0 |
| Prag1 | 0 | 79 | 2571.33333 | 1 |
| Prdm1 | 0 | 45 | 305.333333 | 0 |
| Prdm1 | 0 | 45 | 115.333333 | 1 |
| Prdm16 | 0 | 0 | 553.666667 | 0 |
| Prdm16 | 0 | 1 | 380 | 1 |
| Prelp | 0 | 29.5 | 4315 | 0 |
| Prelp | 0 | 13 | 1528.33333 | 1 |
| Prickle1 | 0 | 238 | 1285 | 0 |
| Prickle1 | 0 | 320 | 2569.66667 | 1 |
| Prickle2 | 0 | 767 | 733.333333 | 0 |
| Prickle2 | 0 | 903.5 | 1210.66667 | 1 |
| Prkag2 | 0 | 397.5 | 1675.66667 | 0 |
| Prkag2 | 0 | 426.5 | 2432 | 1 |
| Prkar2b | 0 | 123.5 | 2566.33333 | 0 |
| Prkar2b | 0 | 115.5 | 1414.66667 | 1 |
| Prkca | 0 | 419.5 | 2671.33333 | 0 |
| Prkca | 0 | 441 | 1805.66667 | 1 |
| Prkd1 | 0 | 559 | 1039.33333 | 0 |
| Prkd1 | 0 | 747.5 | 2199.66667 | 1 |
| Prkg2 | 0 | 195 | 1080.33333 | 0 |
| Prkg2 | 0 | 197.5 | 378 | 1 |
| Prl3d1 | 0 | 9.5 | 0.0001 | 0 |
| Prl3d1 | 0 | 12.5 | 6.33336667 | 1 |
| Prl3d3 | 0 | 13 | 2.66673333 | 0 |
| Prl3d3 | 0 | 12 | 15.6667 | 1 |
| Prl8a9 | 0 | 8 | 8.33336667 | 0 |
| Prl8a9 | 0 | 22 | 33 | 1 |
| Prokr2 | 0 | 33 | 29 | 0 |
| Prokr2 | 0 | 32 | 8.66666667 | 1 |

| | | | | |
|---|---|---|---|---|
| Prom1 | 0 | 266 | 56 | 0 |
| Prom1 | 0 | 258.5 | 29.6666667 | 1 |
| Prr5l | 0 | 398.5 | 818.666667 | 0 |
| Prr5l | 0 | 474.5 | 1229 | 1 |
| Prrt4 | 0 | 17.5 | 33.6666667 | 0 |
| Prrt4 | 0 | 11.5 | 11 | 1 |
| Prss22 | 0 | 13 | 12.6666667 | 0 |
| Prss22 | 0 | 8 | 41.3333333 | 1 |
| Prss23 | 0 | 78.5 | 47751 | 0 |
| Prss23 | 0 | 120.5 | 78107.6667 | 1 |
| Prss35 | 0 | 130.5 | 559 | 0 |
| Prss35 | 0 | 125.5 | 4893 | 1 |
| Prss46 | 0 | 20 | 28 | 0 |
| Prss46 | 0 | 20 | 6.6667 | 1 |
| Prune2 | 0 | 385.5 | 704 | 0 |
| Prune2 | 0 | 395.5 | 439.333333 | 1 |
| Prx | 0 | 67 | 1200 | 0 |
| Prx | 0 | 36.5 | 1695 | 1 |
| Psd4 | 0 | 101 | 219.666667 | 0 |
| Psd4 | 0 | 82 | 140.666667 | 1 |
| Pstpip1 | 0 | 78 | 201.333333 | 0 |
| Pstpip1 | 0 | 63.5 | 313.666667 | 1 |
| Ptchd4 | 0 | 289.5 | 451.666667 | 0 |
| Ptchd4 | 0 | 382 | 642 | 1 |
| Ptgdr2 | 0 | 18 | 1.00003333 | 0 |
| Ptgdr2 | 0 | 9.5 | 40.3333333 | 1 |
| Ptger1 | 0 | 52.5 | 139 | 0 |
| Ptger1 | 0 | 66.5 | 59.3333333 | 1 |
| Ptger4 | 0 | 72 | 660.333333 | 0 |
| Ptger4 | 0 | 56 | 372.666667 | 1 |
| Ptgir | 0 | 13.5 | 224.666667 | 0 |
| Ptgir | 0 | 8.5 | 74.3333333 | 1 |
| Ptgis | 0 | 66.5 | 1095.33333 | 0 |
| Ptgis | 0 | 51 | 674.333333 | 1 |
| Ptgs1 | 0 | 65.5 | 1498.66667 | 0 |
| Ptgs1 | 0 | 72 | 1080.66667 | 1 |
| Ptgs2os2 | 0 | 14.5 | 174.666667 | 0 |
| Ptgs2os2 | 0 | 18.5 | 67.6666667 | 1 |
| Ptk7 | 0 | 122.5 | 24349.3333 | 0 |
| Ptk7 | 0 | 136.5 | 17406.3333 | 1 |
| Ptn | 0 | 732.5 | 11881.6667 | 0 |
| Ptn | 0 | 834 | 7346.33333 | 1 |
| Ptpn22 | 0 | 75.5 | 16.3333333 | 0 |
| Ptpn22 | 0 | 93 | 46 | 1 |
| Ptpn3 | 0 | 142 | 392.333333 | 0 |
| Ptpn3 | 0 | 129.5 | 247 | 1 |
| Ptprb | 0 | 214.5 | 525.333333 | 0 |

| | | | | |
|---|---|---|---|---|
| Ptprb | 0 | 201 | 272 | 1 |
| Ptprc | 0 | 179.5 | 251 | 0 |
| Ptprc | 0 | 231.5 | 159 | 1 |
| Ptprj | 0 | 277.5 | 660.333333 | 0 |
| Ptprj | 0 | 316.5 | 1007.33333 | 1 |
| Ptprk | 0 | 801 | 2074 | 0 |
| Ptprk | 0 | 763 | 4280.66667 | 1 |
| Ptprq | 0 | 287.5 | 279.666667 | 0 |
| Ptprq | 0 | 340 | 570.666667 | 1 |
| Ptprv | 0 | 29.5 | 1244.66667 | 0 |
| Ptprv | 0 | 30.5 | 894 | 1 |
| Pttg1ip | 0 | 63 | 8765.66667 | 0 |
| Pttg1ip | 0 | 57.5 | 14811 | 1 |
| Ptx3 | 0 | 311.5 | 7234 | 0 |
| Ptx3 | 0 | 339 | 2321.33333 | 1 |
| Pxn | 0 | 151 | 2699.33333 | 0 |
| Pxn | 0 | 77 | 4555.33333 | 1 |
| Pygl | 0 | 59.5 | 89 | 0 |
| Pygl | 0 | 52.5 | 177.333333 | 1 |
| Rab11fip1 | 0 | 47 | 232.333333 | 0 |
| Rab11fip1 | 0 | 54.5 | 357 | 1 |
| Rab3c | 0 | 356 | 165.666667 | 0 |
| Rab3c | 0 | 429 | 17 | 1 |
| Rab3il1 | 0 | 102.5 | 494.333333 | 0 |
| Rab3il1 | 0 | 72.5 | 325.666667 | 1 |
| Rad9a | 0 | 14 | 11.6666667 | 0 |
| Rad9a | 0 | 15 | 105.666667 | 1 |
| Rai14 | 0 | 150.5 | 9276.66667 | 0 |
| Rai14 | 0 | 251 | 16373.6667 | 1 |
| Ralgds | 0 | 123 | 1726.66667 | 0 |
| Ralgds | 0 | 116 | 1109 | 1 |
| Ralgps2 | 0 | 138 | 389 | 0 |
| Ralgps2 | 0 | 89.5 | 596.666667 | 1 |
| Ramp3 | 0 | 13.5 | 578.333333 | 0 |
| Ramp3 | 0 | 13.5 | 152 | 1 |
| Rap1gap | 0 | 198.5 | 164 | 0 |
| Rap1gap | 0 | 257.5 | 279 | 1 |
| Rap1gap2 | 0 | 232 | 1624 | 0 |
| Rap1gap2 | 0 | 220 | 955.666667 | 1 |
| Rarb | 1 | 1516.5 | 62.3333333 | 0 |
| Rarb | 1 | 1744.5 | 2906.33333 | 1 |
| Rarg | 0 | 68 | 4444.66667 | 0 |
| Rarg | 1 | 67 | 9277 | 1 |
| Rarres1 | 0 | 51 | 371.333333 | 0 |
| Rarres1 | 0 | 48 | 225 | 1 |
| Rarres2 | 0 | 11.5 | 54 | 0 |
| Rarres2 | 0 | 19 | 98.6666667 | 1 |

| | | | | |
|---|---|---|---|---|
| Rasa1 | 0 | 99.5 | 12044 | 0 |
| Rasa1 | 0 | 101.5 | 8607.33333 | 1 |
| Rasa2 | 0 | 143.5 | 1258.33333 | 0 |
| Rasa2 | 0 | 187 | 910 | 1 |
| Rasa3 | 0 | 170.5 | 5884.66667 | 0 |
| Rasa3 | 1 | 149.5 | 3890.66667 | 1 |
| Rasgrp1 | 0 | 98.5 | 59.3333333 | 0 |
| Rasgrp1 | 0 | 100.5 | 24 | 1 |
| Rasgrp3 | 0 | 470 | 1033.66667 | 0 |
| Rasgrp3 | 0 | 604 | 1557.66667 | 1 |
| Rasl10b | 0 | 119.5 | 31.6666667 | 0 |
| Rasl10b | 0 | 129.5 | 320.666667 | 1 |
| Rasl11a | 0 | 12 | 276.333333 | 0 |
| Rasl11a | 0 | 3.5 | 450 | 1 |
| Rassf2 | 0 | 178.5 | 816.333333 | 0 |
| Rassf2 | 1 | 180.5 | 1128 | 1 |
| Rassf4 | 0 | 37 | 168 | 0 |
| Rassf4 | 0 | 24.5 | 89.3333333 | 1 |
| Rassf6 | 0 | 132.5 | 580.666667 | 0 |
| Rassf6 | 0 | 86.5 | 361.333333 | 1 |
| Rassf7 | 0 | 14 | 646.666667 | 0 |
| Rassf7 | 0 | 7.5 | 446 | 1 |
| Rassf9 | 0 | 66.5 | 16.6666667 | 0 |
| Rassf9 | 0 | 77.5 | 2.6667 | 1 |
| Rb1cc1 | 0 | 141.5 | 6227 | 0 |
| Rb1cc1 | 0 | 147 | 4504.33333 | 1 |
| Rbm20 | 0 | 208.5 | 204.333333 | 0 |
| Rbm20 | 1 | 178 | 136 | 1 |
| Rbm24 | 0 | 24 | 33.6666667 | 0 |
| Rbm24 | 0 | 21.5 | 11.6666667 | 1 |
| Rbp1 | 0 | 186 | 879.666667 | 0 |
| Rbp1 | 0 | 182 | 1699.66667 | 1 |
| Rbpms | 1 | 338.5 | 1719.33333 | 0 |
| Rbpms | 1 | 367.5 | 3371.66667 | 1 |
| Rcan2 | 0 | 472 | 1691.33333 | 0 |
| Rcan2 | 0 | 520.5 | 951 | 1 |
| Rcor3 | 0 | 33 | 374.333333 | 0 |
| Rcor3 | 0 | 27 | 587.666667 | 1 |
| Relb | 0 | 90 | 762.333333 | 0 |
| Relb | 0 | 138 | 539 | 1 |
| Rem1 | 0 | 17 | 165.333333 | 0 |
| Rem1 | 0 | 15.5 | 71.6666667 | 1 |
| Rerg | 0 | 361.5 | 805.333333 | 0 |
| Rerg | 0 | 341.5 | 186.333333 | 1 |
| Ret | 0 | 130 | 128.666667 | 0 |
| Ret | 0 | 56 | 70.6666667 | 1 |
| Rev3l | 0 | 183.5 | 1474.66667 | 0 |

| | | | | |
|---|---|---|---|---|
| Rev3l | 0 | 223.5 | 2287.66667 | 1 |
| Rflnb | 0 | 166 | 2730.66667 | 0 |
| Rflnb | 0 | 149.5 | 1420.66667 | 1 |
| Rgl1 | 1 | 485.5 | 3024.33333 | 0 |
| Rgl1 | 1 | 505 | 5614.66667 | 1 |
| Rgmb | 0 | 63 | 1387.66667 | 0 |
| Rgmb | 0 | 65.5 | 2468.66667 | 1 |
| Rgs16 | 0 | 19.5 | 3905.66667 | 0 |
| Rgs16 | 0 | 22.5 | 6699.33333 | 1 |
| Rgs17 | 0 | 115.5 | 306.666667 | 0 |
| Rgs17 | 0 | 120 | 169.666667 | 1 |
| Rgs2 | 0 | 12.5 | 417.333333 | 0 |
| Rgs2 | 0 | 14.5 | 166 | 1 |
| Rgs21 | 0 | 124 | 24 | 0 |
| Rgs21 | 0 | 140.5 | 9.33333333 | 1 |
| Rgs4 | 0 | 20 | 2252.66667 | 0 |
| Rgs4 | 0 | 16.5 | 554.666667 | 1 |
| Rgs7 | 0 | 680 | 5.66666667 | 0 |
| Rgs7 | 0 | 1039.5 | 19.3333333 | 1 |
| Rhobtb2 | 0 | 48 | 612.666667 | 0 |
| Rhobtb2 | 0 | 44 | 882.666667 | 1 |
| Rhoc | 0 | 91 | 11878 | 0 |
| Rhoc | 0 | 85.5 | 17535 | 1 |
| Rida | 0 | 23 | 194.666667 | 0 |
| Rida | 0 | 23.5 | 997 | 1 |
| Rin2 | 0 | 156 | 1071.33333 | 0 |
| Rin2 | 0 | 219 | 1899 | 1 |
| Ripk2 | 0 | 103 | 837 | 0 |
| Ripk2 | 0 | 99.5 | 484 | 1 |
| Ripk4 | 0 | 70.5 | 24.6666667 | 0 |
| Ripk4 | 0 | 60.5 | 9.6667 | 1 |
| Rnd1 | 0 | 21 | 711.666667 | 0 |
| Rnd1 | 0 | 19 | 2190.33333 | 1 |
| Rnf128 | 0 | 219 | 1215 | 0 |
| Rnf128 | 0 | 252 | 2080.66667 | 1 |
| Rnf144b | 0 | 171 | 173 | 0 |
| Rnf144b | 0 | 180 | 110 | 1 |
| Rnf150 | 0 | 353.5 | 1911.66667 | 0 |
| Rnf150 | 1 | 422 | 1118.33333 | 1 |
| Rnf39 | 0 | 15 | 99.6666667 | 0 |
| Rnf39 | 0 | 10.5 | 48 | 1 |
| Robo2 | 0 | 1505.5 | 744.333333 | 0 |
| Robo2 | 0 | 2386.5 | 437.333333 | 1 |
| Ror1 | 0 | 638 | 644.666667 | 0 |
| Ror1 | 0 | 984 | 410 | 1 |
| Rprm | 0 | 7 | 182.333333 | 0 |
| Rprm | 0 | 2.5 | 96.3333333 | 1 |

| | | | | |
|---|---|---|---|---|
| Rspo2 | 0 | 549 | 690.333333 | 0 |
| Rspo2 | 1 | 500 | 161.666667 | 1 |
| Rtl3 | 0 | 13.5 | 2359.33333 | 0 |
| Rtl3 | 0 | 13.5 | 1074.33333 | 1 |
| Rtn4rl1 | 0 | 167.5 | 287.666667 | 0 |
| Rtn4rl1 | 0 | 156.5 | 423.333333 | 1 |
| Rufy1 | 0 | 66 | 1256.33333 | 0 |
| Rufy1 | 0 | 63.5 | 803.666667 | 1 |
| Runx3 | 0 | 118 | 411 | 0 |
| Runx3 | 0 | 105.5 | 266.666667 | 1 |
| Rusc2 | 1 | 199.5 | 4947 | 0 |
| Rusc2 | 1 | 192.5 | 8690.33333 | 1 |
| Rxra | 0 | 204.5 | 4064 | 0 |
| Rxra | 0 | 192.5 | 2924.33333 | 1 |
| S100a10 | 0 | 59.5 | 9583.66667 | 0 |
| S100a10 | 0 | 41.5 | 6915.33333 | 1 |
| S100a16 | 0 | 27 | 1021 | 0 |
| S100a16 | 0 | 14 | 1957 | 1 |
| S100a4 | 0 | 17 | 4408.66667 | 0 |
| S100a4 | 0 | 19 | 2648.33333 | 1 |
| S100a7a | 0 | 9 | 304 | 0 |
| S100a7a | 0 | 15 | 159 | 1 |
| S1pr1 | 0 | 19.5 | 2421.66667 | 0 |
| S1pr1 | 0 | 16 | 5508.33333 | 1 |
| S1pr3 | 0 | 147.5 | 11747.3333 | 0 |
| S1pr3 | 0 | 182 | 16368.3333 | 1 |
| Samd1 | 0 | 34.5 | 4745.66667 | 0 |
| Samd1 | 0 | 30.5 | 3228.33333 | 1 |
| Sap18 | 0 | 5 | 4937 | 0 |
| Sap18 | 0 | 4.5 | 3485.33333 | 1 |
| Sap18b | 0 | 0.5 | 1796 | 0 |
| Sap18b | 0 | 0 | 1259.33333 | 1 |
| Sardh | 0 | 247 | 1027 | 0 |
| Sardh | 0 | 208 | 701.666667 | 1 |
| Scara3 | 0 | 66.5 | 1842 | 0 |
| Scara3 | 0 | 69.5 | 2647 | 1 |
| Scara5 | 0 | 287.5 | 139 | 0 |
| Scara5 | 0 | 188.5 | 25.6666667 | 1 |
| Scarna2 | 0 | 36 | 13 | 0 |
| Scarna2 | 0 | 42.5 | 0.0001 | 1 |
| Scd1 | 0 | 44.5 | 11690 | 0 |
| Scd1 | 0 | 42 | 17838.3333 | 1 |
| Scd2 | 0 | 33.5 | 56002.3333 | 0 |
| Scd2 | 1 | 27.5 | 77839 | 1 |
| Scml4 | 0 | 389 | 10.6666667 | 0 |
| Scml4 | 0 | 309 | 2.66666667 | 1 |
| Scn3a | 0 | 262 | 18 | 0 |

| | | | | |
|---|---|---|---|---|
| Scn3a | 0 | 449 | 4.33336667 | 1 |
| Scn5a | 0 | 230 | 3.33333333 | 0 |
| Scn5a | 0 | 199.5 | 13.6666667 | 1 |
| Scn7a | 0 | 112.5 | 69.6666667 | 0 |
| Scn7a | 0 | 113 | 152 | 1 |
| Scn9a | 0 | 387.5 | 0.66673333 | 0 |
| Scn9a | 0 | 293.5 | 11.3333333 | 1 |
| Scrn1 | 0 | 107 | 990 | 0 |
| Scrn1 | 0 | 130.5 | 619 | 1 |
| Scrt1 | 0 | 5.5 | 3.66673333 | 0 |
| Scrt1 | 0 | 7 | 18.6666667 | 1 |
| Scube1 | 0 | 182 | 119.333333 | 0 |
| Scube1 | 0 | 207 | 253 | 1 |
| Scube3 | 0 | 67.5 | 296 | 0 |
| Scube3 | 0 | 61.5 | 490 | 1 |
| Scx | 0 | 43.5 | 413 | 0 |
| Scx | 0 | 48 | 206 | 1 |
| Sele | 0 | 27.5 | 33.6666667 | 0 |
| Sele | 0 | 29 | 15.6666667 | 1 |
| Sema3a | 0 | 347 | 5917.66667 | 0 |
| Sema3a | 0 | 464.5 | 10781 | 1 |
| Sema3c | 1 | 247.5 | 3467 | 0 |
| Sema3c | 0 | 299.5 | 5974 | 1 |
| Sema3e | 0 | 428 | 1834.66667 | 0 |
| Sema3e | 0 | 519 | 2945 | 1 |
| Sema3g | 0 | 22.5 | 30.6666667 | 0 |
| Sema3g | 0 | 16.5 | 11.3333333 | 1 |
| Sema5a | 0 | 551 | 2447.66667 | 0 |
| Sema5a | 0 | 767.5 | 3351 | 1 |
| Sema5b | 0 | 256.5 | 389.666667 | 0 |
| Sema5b | 0 | 226.5 | 177 | 1 |
| Sema6a | 0 | 292 | 920.666667 | 0 |
| Sema6a | 0 | 347.5 | 1758 | 1 |
| Sema7a | 0 | 48.5 | 909 | 0 |
| Sema7a | 0 | 56 | 373.333333 | 1 |
| Serpina3f | 0 | 13 | 44.3333333 | 0 |
| Serpina3f | 0 | 12 | 8.33333333 | 1 |
| Serpina3g | 0 | 42.5 | 627 | 0 |
| Serpina3g | 0 | 19.5 | 166.666667 | 1 |
| Serpina3h | 0 | 16 | 1509 | 0 |
| Serpina3h | 0 | 11 | 249 | 1 |
| Serpina3i | 0 | 10 | 410.666667 | 0 |
| Serpina3i | 0 | 11.5 | 55 | 1 |
| Serpina3m | 0 | 12 | 10.6666667 | 0 |
| Serpina3m | 0 | 11.5 | 2.00003333 | 1 |
| Serpina3n | 0 | 22.5 | 694 | 0 |
| Serpina3n | 0 | 13.5 | 312 | 1 |

| | | | | |
|---|---|---|---|---|
| Serpinb1a | 0 | 21 | 76.3333333 | 0 |
| Serpinb1a | 0 | 27 | 27.6666667 | 1 |
| Serpinb8 | 0 | 35.5 | 302.333333 | 0 |
| Serpinb8 | 0 | 77 | 170.666667 | 1 |
| Serping1 | 0 | 28 | 620.333333 | 0 |
| Serping1 | 0 | 24.5 | 1197.66667 | 1 |
| Serpini1 | 0 | 152.5 | 46 | 0 |
| Serpini1 | 0 | 153 | 94.6666667 | 1 |
| Sez6l | 0 | 197 | 45 | 0 |
| Sez6l | 0 | 185.5 | 20 | 1 |
| Sfn | 0 | 29 | 704 | 0 |
| Sfn | 0 | 15.5 | 400.333333 | 1 |
| Sfrp2 | 0 | 19.5 | 4234.66667 | 0 |
| Sfrp2 | 0 | 15.5 | 711.333333 | 1 |
| Sfrp4 | 0 | 54.5 | 12.3333333 | 0 |
| Sfrp4 | 0 | 51 | 56.3333333 | 1 |
| Sfxn3 | 0 | 57 | 5699 | 0 |
| Sfxn3 | 0 | 49.5 | 3637.33333 | 1 |
| Sgcd | 0 | 516.5 | 2800.33333 | 0 |
| Sgcd | 0 | 636 | 1730 | 1 |
| Sgk1 | 0 | 266 | 4043.66667 | 0 |
| Sgk1 | 0 | 298.5 | 2837 | 1 |
| Sgpl1 | 0 | 150 | 6691.66667 | 0 |
| Sgpl1 | 1 | 142.5 | 9986 | 1 |
| Sgpp1 | 0 | 53.5 | 1862.33333 | 0 |
| Sgpp1 | 0 | 44 | 1332.33333 | 1 |
| Sh2d1b1 | 0 | 14 | 33 | 0 |
| Sh2d1b1 | 0 | 31 | 12.6666667 | 1 |
| Sh3d21 | 0 | 74 | 182.666667 | 0 |
| Sh3d21 | 0 | 94 | 114.333333 | 1 |
| Sh3gl3 | 0 | 146 | 75.3333333 | 0 |
| Sh3gl3 | 0 | 242 | 203 | 1 |
| Sh3kbp1 | 0 | 911 | 6295.66667 | 0 |
| Sh3kbp1 | 0 | 1139 | 10069.6667 | 1 |
| Sh3rf2 | 0 | 266 | 27.3333333 | 0 |
| Sh3rf2 | 0 | 170.5 | 67.3333333 | 1 |
| Sh3rf3 | 0 | 430 | 1092.66667 | 0 |
| Sh3rf3 | 0 | 418.5 | 623 | 1 |
| Shank2 | 0 | 395.5 | 773.666667 | 0 |
| Shank2 | 0 | 360.5 | 363.333333 | 1 |
| Shc4 | 0 | 86.5 | 143.333333 | 0 |
| Shc4 | 0 | 113.5 | 272.333333 | 1 |
| Shisa2 | 0 | 430.5 | 73.3333333 | 0 |
| Shisa2 | 0 | 417.5 | 34.3333333 | 1 |
| Shox2 | 0 | 27 | 1020.66667 | 0 |
| Shox2 | 0 | 17 | 1672 | 1 |
| Shroom2 | 0 | 567 | 423 | 0 |

| | | | | |
|---|---|---|---|---|
| Shroom2 | 0 | 716.5 | 794.666667 | 1 |
| Siglecg | 0 | 27.5 | 2384 | 0 |
| Siglecg | 0 | 26 | 1540.66667 | 1 |
| Sik1 | 0 | 38.5 | 1819 | 0 |
| Sik1 | 0 | 46.5 | 5053.33333 | 1 |
| Sim2 | 0 | 201 | 780.666667 | 0 |
| Sim2 | 0 | 182.5 | 1728.33333 | 1 |
| Sipa1l1 | 0 | 669 | 6397 | 0 |
| Sipa1l1 | 1 | 886 | 11453.3333 | 1 |
| Sipa1l2 | 0 | 323.5 | 3430.66667 | 0 |
| Sipa1l2 | 0 | 300.5 | 1692.33333 | 1 |
| Sirpa | 0 | 79.5 | 2995.33333 | 0 |
| Sirpa | 0 | 72 | 1895.66667 | 1 |
| Six1 | 0 | 10.5 | 935 | 0 |
| Six1 | 0 | 8.5 | 1391.66667 | 1 |
| Skil | 0 | 155 | 2906 | 0 |
| Skil | 0 | 192.5 | 4006.33333 | 1 |
| Sla | 0 | 334.5 | 88 | 0 |
| Sla | 0 | 240 | 34.3333333 | 1 |
| Slc10a6 | 1 | 198.5 | 9 | 0 |
| Slc10a6 | 0 | 204 | 0.0001 | 1 |
| Slc12a6 | 0 | 113.5 | 48.6666667 | 0 |
| Slc12a6 | 0 | 119 | 84 | 1 |
| Slc13a5 | 0 | 47 | 115 | 0 |
| Slc13a5 | 0 | 51.5 | 12.3333333 | 1 |
| Slc14a2 | 1 | 1863 | 26 | 0 |
| Slc14a2 | 1 | 1730 | 10 | 1 |
| Slc15a3 | 0 | 32.5 | 261.666667 | 0 |
| Slc15a3 | 0 | 31.5 | 410 | 1 |
| Slc16a12 | 0 | 111.5 | 14.6666667 | 0 |
| Slc16a12 | 0 | 153 | 61 | 1 |
| Slc16a2 | 0 | 194.5 | 526.666667 | 0 |
| Slc16a2 | 0 | 196 | 278 | 1 |
| Slc17a6 | 0 | 160.5 | 7.66666667 | 0 |
| Slc17a6 | 0 | 164 | 1.00003333 | 1 |
| Slc1a3 | 1 | 173 | 815 | 0 |
| Slc1a3 | 1 | 174 | 186.666667 | 1 |
| Slc20a1 | 0 | 91 | 8706.33333 | 0 |
| Slc20a1 | 0 | 52.5 | 6401.33333 | 1 |
| Slc20a2 | 0 | 36 | 4036.66667 | 0 |
| Slc20a2 | 0 | 38 | 5980 | 1 |
| Slc22a23 | 0 | 204 | 276 | 0 |
| Slc22a23 | 0 | 225 | 421 | 1 |
| Slc24a3 | 1 | 467 | 498 | 0 |
| Slc24a3 | 0 | 597 | 1165 | 1 |
| Slc25a24 | 0 | 114.5 | 3816 | 0 |
| Slc25a24 | 1 | 134 | 5379.33333 | 1 |

| | | | | |
|---|---|---|---|---|
| Slc25a51 | 0 | 124 | 4034.66667 | 0 |
| Slc25a51 | 0 | 141 | 6255 | 1 |
| Slc26a10 | 0 | 30 | 30 | 0 |
| Slc26a10 | 0 | 27.5 | 12.6666667 | 1 |
| Slc27a1 | 0 | 44.5 | 627 | 0 |
| Slc27a1 | 0 | 47 | 401.666667 | 1 |
| Slc29a1 | 0 | 35.5 | 3090.33333 | 0 |
| Slc29a1 | 0 | 25.5 | 1502.66667 | 1 |
| Slc29a2 | 0 | 17 | 517.333333 | 0 |
| Slc29a2 | 0 | 13.5 | 320.333333 | 1 |
| Slc2a13 | 0 | 360 | 326.666667 | 0 |
| Slc2a13 | 0 | 425 | 566.333333 | 1 |
| Slc2a5 | 0 | 36 | 21 | 0 |
| Slc2a5 | 0 | 32 | 7.66666667 | 1 |
| Slc38a4 | 0 | 145 | 4551 | 0 |
| Slc38a4 | 0 | 219 | 8348.66667 | 1 |
| Slc38a6 | 0 | 122 | 315 | 0 |
| Slc38a6 | 0 | 150 | 492.666667 | 1 |
| Slc4a10 | 0 | 619.5 | 10.3333667 | 0 |
| Slc4a10 | 0 | 765 | 69 | 1 |
| Slc4a4 | 0 | 993 | 1031.33333 | 0 |
| Slc4a4 | 0 | 1316 | 2274.33333 | 1 |
| Slc52a3 | 0 | 20.5 | 18 | 0 |
| Slc52a3 | 0 | 39.5 | 68.6666667 | 1 |
| Slc5a7 | 0 | 31 | 473 | 0 |
| Slc5a7 | 0 | 41.5 | 45 | 1 |
| Slc6a17 | 0 | 87.5 | 2311.33333 | 0 |
| Slc6a17 | 0 | 98 | 763 | 1 |
| Slc6a6 | 1 | 201.5 | 19695.3333 | 0 |
| Slc6a6 | 0 | 182 | 13599.6667 | 1 |
| Slc7a2 | 0 | 70 | 783 | 0 |
| Slc7a2 | 0 | 94.5 | 548.333333 | 1 |
| Slc8a1 | 0 | 1336 | 1115 | 0 |
| Slc8a1 | 1 | 1378 | 2331.66667 | 1 |
| Slc9a3r1 | 0 | 43 | 1725.66667 | 0 |
| Slc9a3r1 | 0 | 38.5 | 1215 | 1 |
| Slco2a1 | 0 | 121 | 3058 | 0 |
| Slco2a1 | 0 | 126.5 | 5840.33333 | 1 |
| Slco2b1 | 0 | 87 | 26.6666667 | 0 |
| Slco2b1 | 0 | 87.5 | 68 | 1 |
| Slf2 | 0 | 117.5 | 2445.33333 | 0 |
| Slf2 | 0 | 193.5 | 6052.66667 | 1 |
| Slfn5 | 0 | 39 | 34.0000333 | 0 |
| Slfn5 | 0 | 34.5 | 10.3333667 | 1 |
| Slitrk2 | 0 | 15.5 | 16.6666667 | 0 |
| Slitrk2 | 0 | 17.5 | 5 | 1 |
| Slpi | 0 | 5 | 105.666667 | 0 |

| | | | | |
|---|---|---|---|---|
| Slpi | 0 | 4 | 60.6666667 | 1 |
| Smad1 | 0 | 131 | 1606.66667 | 0 |
| Smad1 | 0 | 164 | 2333.66667 | 1 |
| Smad3 | 0 | 213 | 4985.66667 | 0 |
| Smad3 | 1 | 280 | 7666 | 1 |
| Smad6 | 0 | 203 | 1523 | 0 |
| Smad6 | 0 | 274.5 | 2319.66667 | 1 |
| Smg6 | 0 | 231 | 1223 | 0 |
| Smg6 | 0 | 229 | 1758.33333 | 1 |
| Smim3 | 0 | 75 | 856 | 0 |
| Smim3 | 0 | 62.5 | 552.333333 | 1 |
| Smoc1 | 0 | 275 | 1349.66667 | 0 |
| Smoc1 | 0 | 273 | 860.333333 | 1 |
| Smoc2 | 0 | 163.5 | 5190 | 0 |
| Smoc2 | 0 | 144 | 3476.33333 | 1 |
| Snap47 | 0 | 88 | 2529 | 0 |
| Snap47 | 0 | 138.5 | 3710 | 1 |
| Snap91 | 0 | 195.5 | 50.3333333 | 0 |
| Snap91 | 0 | 264 | 172.333333 | 1 |
| Sncaip | 0 | 319 | 138.666667 | 0 |
| Sncaip | 0 | 320 | 242.333333 | 1 |
| Sncg | 0 | 7 | 50 | 0 |
| Sncg | 0 | 3.5 | 23 | 1 |
| Snph | 0 | 135 | 34 | 0 |
| Snph | 0 | 137 | 62.6666667 | 1 |
| Snrnp40 | 0 | 72 | 11.3333333 | 0 |
| Snrnp40 | 0 | 63 | 100 | 1 |
| Snta1 | 0 | 51.5 | 1228.33333 | 0 |
| Snta1 | 0 | 49 | 1961.66667 | 1 |
| Snx9 | 0 | 102 | 9487 | 0 |
| Snx9 | 0 | 102.5 | 6282.33333 | 1 |
| Sod3 | 0 | 54.5 | 1206.33333 | 0 |
| Sod3 | 0 | 53.5 | 641.333333 | 1 |
| Sorcs1 | 0 | 1692.5 | 164.333333 | 0 |
| Sorcs1 | 0 | 1602.5 | 91.3333333 | 1 |
| Sord | 0 | 57 | 611.333333 | 0 |
| Sord | 0 | 60.5 | 1229 | 1 |
| Sox6 | 0 | 1016.5 | 180.666667 | 0 |
| Sox6 | 0 | 1200.5 | 115.666667 | 1 |
| Sox9 | 0 | 19 | 802 | 0 |
| Sox9 | 0 | 10 | 278 | 1 |
| Sp7 | 0 | 20.5 | 53.6666667 | 0 |
| Sp7 | 0 | 19.5 | 13.6666667 | 1 |
| Sparcl1 | 0 | 75.5 | 156 | 0 |
| Sparcl1 | 0 | 65 | 410.333333 | 1 |
| Spcs3 | 0 | 20.5 | 4377.66667 | 0 |
| Spcs3 | 0 | 25 | 5980 | 1 |

| | | | | |
|---|---|---|---|---|
| Specc1 | 0 | 479.5 | 4843.33333 | 0 |
| Specc1 | 0 | 638 | 10309 | 1 |
| Spin4 | 0 | 97 | 372 | 0 |
| Spin4 | 0 | 110.5 | 526.666667 | 1 |
| Spink6 | 0 | 33.5 | 0.0001 | 0 |
| Spink6 | 0 | 32.5 | 4.33333333 | 1 |
| Spire1 | 0 | 79 | 2548.33333 | 0 |
| Spire1 | 0 | 92.5 | 1763 | 1 |
| Spock1 | 0 | 1775 | 44.6666667 | 0 |
| Spock1 | 0 | 1825 | 19.3333333 | 1 |
| Spon2 | 0 | 13.5 | 129.666667 | 0 |
| Spon2 | 0 | 27 | 4669.66667 | 1 |
| Spp1 | 0 | 30 | 9879 | 0 |
| Spp1 | 0 | 49 | 16502.3333 | 1 |
| Sprr1a | 0 | 6 | 17.3333333 | 0 |
| Sprr1a | 0 | 4.5 | 649 | 1 |
| Sprr3 | 0 | 8.5 | 37 | 0 |
| Sprr3 | 0 | 29 | 109.333333 | 1 |
| Spsb4 | 0 | 122 | 97.6666667 | 0 |
| Spsb4 | 0 | 133.5 | 234.666667 | 1 |
| Sqor | 0 | 89 | 346.333333 | 0 |
| Sqor | 0 | 79.5 | 238.666667 | 1 |
| Srgap1 | 0 | 364 | 821 | 0 |
| Srgap1 | 0 | 372.5 | 1414 | 1 |
| Srgap3 | 1 | 288.5 | 912 | 0 |
| Srgap3 | 1 | 310.5 | 2225.33333 | 1 |
| Srr | 0 | 35.5 | 519.666667 | 0 |
| Srr | 0 | 39 | 751.333333 | 1 |
| Ssc5d | 0 | 53 | 1759 | 0 |
| Ssc5d | 0 | 45.5 | 1016.66667 | 1 |
| Ssh3 | 0 | 38.5 | 704.333333 | 0 |
| Ssh3 | 0 | 24.5 | 456.333333 | 1 |
| Sstr2 | 0 | 23 | 39.3333333 | 0 |
| Sstr2 | 0 | 11 | 6 | 1 |
| Sstr4 | 0 | 2 | 63.3333333 | 0 |
| Sstr4 | 0 | 1 | 167.333333 | 1 |
| Ssx2ip | 0 | 88.5 | 736.333333 | 0 |
| Ssx2ip | 0 | 91 | 1182 | 1 |
| St3gal4 | 0 | 49.5 | 1090.66667 | 0 |
| St3gal4 | 1 | 51 | 748 | 1 |
| St6gal1 | 0 | 298 | 411.666667 | 0 |
| St6gal1 | 0 | 314.5 | 668.666667 | 1 |
| Stac | 0 | 322 | 56.3333333 | 0 |
| Stac | 0 | 319 | 13.3333333 | 1 |
| Stard10 | 0 | 46.5 | 133 | 0 |
| Stard10 | 0 | 41.5 | 522.333333 | 1 |
| Stat5a | 0 | 99 | 877.666667 | 0 |

| | | | | |
|---|---|---|---|---|
| Stat5a | 0 | 89.5 | 630.333333 | 1 |
| Stc1 | 0 | 39.5 | 472.666667 | 0 |
| Stc1 | 0 | 28 | 244.333333 | 1 |
| Steap3 | 0 | 84.5 | 1676.33333 | 0 |
| Steap3 | 0 | 75.5 | 901.666667 | 1 |
| Stk40 | 0 | 129 | 4504.33333 | 0 |
| Stk40 | 0 | 101 | 3003.66667 | 1 |
| Stmn2 | 0 | 77 | 198.333333 | 0 |
| Stmn2 | 0 | 84.5 | 132 | 1 |
| Stmn4 | 0 | 59.5 | 9.33333333 | 0 |
| Stmn4 | 0 | 57 | 141.666667 | 1 |
| Stoml1 | 0 | 37 | 754.666667 | 0 |
| Stoml1 | 0 | 17 | 1049.33333 | 1 |
| Ston1 | 1 | 90.5 | 8709 | 0 |
| Ston1 | 0 | 132 | 13684 | 1 |
| Stra6 | 0 | 43 | 4 | 0 |
| Stra6 | 0 | 36.5 | 252.333333 | 1 |
| Stx1a | 0 | 119 | 794.666667 | 0 |
| Stx1a | 0 | 77 | 1479 | 1 |
| Styk1 | 0 | 25.5 | 374 | 0 |
| Styk1 | 0 | 35 | 539.666667 | 1 |
| Susd2 | 0 | 192.5 | 316.666667 | 0 |
| Susd2 | 0 | 208 | 166 | 1 |
| Susd5 | 0 | 75 | 23.6667 | 0 |
| Susd5 | 0 | 76.5 | 7.33336667 | 1 |
| Sv2c | 0 | 216 | 1219 | 0 |
| Sv2c | 0 | 244 | 270 | 1 |
| Svep1 | 0 | 282 | 11205 | 0 |
| Svep1 | 1 | 323.5 | 6098.66667 | 1 |
| Swap70 | 0 | 417 | 2827 | 0 |
| Swap70 | 0 | 460 | 4187 | 1 |
| Sybu | 0 | 260 | 15.6666667 | 0 |
| Sybu | 0 | 306 | 4.33333333 | 1 |
| Syde2 | 0 | 103 | 1132 | 0 |
| Syde2 | 1 | 112.5 | 2180.66667 | 1 |
| Syk | 0 | 147.5 | 548 | 0 |
| Syk | 0 | 133 | 342.333333 | 1 |
| Syn3 | 1 | 253 | 73.6666667 | 0 |
| Syn3 | 0 | 402 | 32.3333333 | 1 |
| Syna | 0 | 9.5 | 12.3333333 | 0 |
| Syna | 0 | 5 | 3.33336667 | 1 |
| Syne3 | 1 | 227 | 2413.33333 | 0 |
| Syne3 | 0 | 160.5 | 1088.33333 | 1 |
| Synj2 | 0 | 467.5 | 2627.66667 | 0 |
| Synj2 | 0 | 407 | 1506 | 1 |
| Synm | 0 | 185.5 | 387 | 0 |
| Synm | 0 | 104.5 | 269.333333 | 1 |

| | | | | |
|---|---|---|---|---|
| Syt13 | 0 | 78.5 | 6966 | 0 |
| Syt13 | 0 | 72 | 3126 | 1 |
| Syt5 | 0 | 82 | 41.6666667 | 0 |
| Syt5 | 0 | 115 | 15 | 1 |
| Syt7 | 0 | 138 | 31 | 0 |
| Syt7 | 0 | 54.5 | 125.333333 | 1 |
| Syt8 | 0 | 9.5 | 127 | 0 |
| Syt8 | 0 | 7.5 | 73.3333333 | 1 |
| Tarm1 | 0 | 46 | 0.6667 | 0 |
| Tarm1 | 0 | 58.5 | 9.33333333 | 1 |
| Tbl1xr1 | 0 | 186.5 | 1197 | 0 |
| Tbl1xr1 | 0 | 225.5 | 771 | 1 |
| Tbxa2r | 0 | 32 | 864 | 0 |
| Tbxa2r | 0 | 36.5 | 464.666667 | 1 |
| Tbxas1 | 0 | 191 | 89 | 0 |
| Tbxas1 | 0 | 233 | 286.333333 | 1 |
| Tcaf2 | 0 | 71.5 | 871.666667 | 0 |
| Tcaf2 | 0 | 72.5 | 543.666667 | 1 |
| Tdrkh | 0 | 69 | 312 | 0 |
| Tdrkh | 0 | 65.5 | 138.333333 | 1 |
| Tecta | 0 | 200 | 12 | 0 |
| Tecta | 0 | 174 | 35 | 1 |
| Tenm4 | 1 | 97.5 | 724.666667 | 0 |
| Tenm4 | 0 | 102.5 | 279.666667 | 1 |
| Tes | 0 | 145 | 3682.33333 | 0 |
| Tes | 0 | 164 | 5450.66667 | 1 |
| Tg | 0 | 216 | 105.666667 | 0 |
| Tg | 0 | 270 | 28.3333333 | 1 |
| Tgfa | 0 | 174 | 102.333333 | 0 |
| Tgfa | 0 | 150.5 | 49.6666667 | 1 |
| Tgfb2 | 0 | 297 | 2934.66667 | 0 |
| Tgfb2 | 0 | 404 | 6728.66667 | 1 |
| Tgfbr3 | 1 | 436 | 3014.66667 | 0 |
| Tgfbr3 | 1 | 405.5 | 2034 | 1 |
| Tgm2 | 0 | 61 | 1714 | 0 |
| Tgm2 | 0 | 60.5 | 12128.3333 | 1 |
| Thbd | 0 | 16 | 6447.33333 | 0 |
| Thbd | 0 | 28 | 13140.3333 | 1 |
| Thbs2 | 0 | 114 | 42211.3333 | 0 |
| Thbs2 | 1 | 182 | 58440.3333 | 1 |
| Thbs4 | 0 | 104.5 | 355.666667 | 0 |
| Thbs4 | 0 | 90 | 245.333333 | 1 |
| Thra | 0 | 95 | 2596 | 0 |
| Thra | 0 | 72 | 1783 | 1 |
| Thsd4 | 0 | 972 | 651.333333 | 0 |
| Thsd4 | 0 | 1048 | 960 | 1 |
| Tiam1 | 0 | 374.5 | 1870.33333 | 0 |

| | | | | |
|---|---|---|---|---|
| Tiam1 | 0 | 361.5 | 1346.66667 | 1 |
| Tifab | 0 | 14 | 99 | 0 |
| Tifab | 0 | 31 | 57.3333333 | 1 |
| Timp1 | 0 | 96.5 | 20445.3333 | 0 |
| Timp1 | 0 | 140 | 12745 | 1 |
| Timp3 | 1 | 584.5 | 74154 | 0 |
| Timp3 | 0 | 541 | 39304.6667 | 1 |
| Tinagl1 | 0 | 26 | 4636.66667 | 0 |
| Tinagl1 | 0 | 29.5 | 15279.6667 | 1 |
| Tll1 | 0 | 262.5 | 7218.33333 | 0 |
| Tll1 | 0 | 330.5 | 3996.66667 | 1 |
| Tll2 | 0 | 186 | 4 | 0 |
| Tll2 | 0 | 189 | 19.6666667 | 1 |
| Tlr2 | 0 | 88 | 239 | 0 |
| Tlr2 | 0 | 75.5 | 127 | 1 |
| Tlr5 | 0 | 116.5 | 106.333333 | 0 |
| Tlr5 | 0 | 102 | 55.6666667 | 1 |
| Tlr9 | 0 | 17 | 30 | 0 |
| Tlr9 | 0 | 13.5 | 10 | 1 |
| Tm9sf3 | 0 | 190 | 12022.3333 | 0 |
| Tm9sf3 | 0 | 204 | 19065.3333 | 1 |
| Tmem108 | 0 | 259.5 | 123 | 0 |
| Tmem108 | 0 | 290 | 206 | 1 |
| Tmem119 | 0 | 20 | 5158.66667 | 0 |
| Tmem119 | 0 | 20.5 | 11256.6667 | 1 |
| Tmem125 | 0 | 52.5 | 40.3333333 | 0 |
| Tmem125 | 0 | 51 | 16.3333333 | 1 |
| Tmem151a | 0 | 25 | 200 | 0 |
| Tmem151a | 0 | 15 | 112.333333 | 1 |
| Tmem164 | 0 | 144.5 | 3769.33333 | 0 |
| Tmem164 | 0 | 123.5 | 5147.66667 | 1 |
| Tmem176a | 0 | 13.5 | 1485.66667 | 0 |
| Tmem176a | 0 | 7.5 | 3405.33333 | 1 |
| Tmem176b | 0 | 23.5 | 2767.33333 | 0 |
| Tmem176b | 0 | 20.5 | 5673.66667 | 1 |
| Tmem178 | 0 | 155 | 107.666667 | 0 |
| Tmem178 | 0 | 140.5 | 17.6666667 | 1 |
| Tmem182 | 0 | 115.5 | 1.33336667 | 0 |
| Tmem182 | 0 | 131 | 11.6666667 | 1 |
| Tmem184a | 0 | 17.5 | 19.6666667 | 0 |
| Tmem184a | 0 | 13.5 | 4.33333333 | 1 |
| Tmem200b | 0 | 120 | 1224.66667 | 0 |
| Tmem200b | 0 | 131.5 | 2009.66667 | 1 |
| Tmem204 | 0 | 109 | 229.666667 | 0 |
| Tmem204 | 1 | 82 | 104.666667 | 1 |
| Tmem255a | 0 | 43.5 | 1.00003333 | 0 |
| Tmem255a | 0 | 65 | 23 | 1 |

| | | | | |
|---|---|---|---|---|
| Tmem26 | 0 | 102 | 328 | 0 |
| Tmem26 | 0 | 94 | 77.6666667 | 1 |
| Tmem35a | 0 | 25 | 86.3333333 | 0 |
| Tmem35a | 0 | 15.5 | 48.3333333 | 1 |
| Tmem40 | 0 | 66.5 | 35.3333333 | 0 |
| Tmem40 | 0 | 55 | 7.66666667 | 1 |
| Tmem45a | 1 | 141.5 | 1391 | 0 |
| Tmem45a | 0 | 137.5 | 767.666667 | 1 |
| Tmem56 | 0 | 114.5 | 66.6666667 | 0 |
| Tmem56 | 0 | 99.5 | 36.3333333 | 1 |
| Tmem82 | 0 | 5 | 9.33333333 | 0 |
| Tmem82 | 0 | 5.5 | 0.66673333 | 1 |
| Tmem86a | 0 | 11.5 | 564.333333 | 0 |
| Tmem86a | 0 | 10.5 | 2065 | 1 |
| Tmtc2 | 0 | 746 | 507 | 0 |
| Tmtc2 | 0 | 969 | 790.666667 | 1 |
| Tmtc4 | 0 | 63.5 | 847.666667 | 0 |
| Tmtc4 | 0 | 88 | 1685.33333 | 1 |
| Tnfaip3 | 0 | 29 | 792.333333 | 0 |
| Tnfaip3 | 0 | 32 | 2435 | 1 |
| Tnfaip6 | 0 | 44 | 466 | 0 |
| Tnfaip6 | 0 | 45.5 | 296 | 1 |
| Tnfaip8l3 | 0 | 88 | 70.3333333 | 0 |
| Tnfaip8l3 | 0 | 79.5 | 17 | 1 |
| Tnfrsf11a | 0 | 170 | 1106.33333 | 0 |
| Tnfrsf11a | 0 | 200.5 | 800.666667 | 1 |
| Tnfrsf11b | 0 | 61.5 | 1783.66667 | 0 |
| Tnfrsf11b | 0 | 62 | 862 | 1 |
| Tnfrsf19 | 0 | 175 | 49 | 0 |
| Tnfrsf19 | 0 | 177.5 | 106.666667 | 1 |
| Tnfrsf1b | 0 | 96.5 | 1778.33333 | 0 |
| Tnfrsf1b | 1 | 104 | 4207.66667 | 1 |
| Tnfrsf25 | 0 | 34.5 | 40.6666667 | 0 |
| Tnfrsf25 | 0 | 26 | 13.6666667 | 1 |
| Tnfsf11 | 0 | 72.5 | 588.333333 | 0 |
| Tnfsf11 | 0 | 64 | 230.666667 | 1 |
| Tnfsf18 | 0 | 30 | 54 | 0 |
| Tnfsf18 | 0 | 30.5 | 156 | 1 |
| Tnk2 | 0 | 133.5 | 17607.3333 | 0 |
| Tnk2 | 0 | 165 | 31302.3333 | 1 |
| Tnn | 0 | 64.5 | 4060 | 0 |
| Tnn | 0 | 67.5 | 1272.66667 | 1 |
| Tnnt2 | 0 | 88 | 372 | 0 |
| Tnnt2 | 0 | 64 | 117 | 1 |
| Tnxb | 0 | 70.5 | 164 | 0 |
| Tnxb | 0 | 67 | 315 | 1 |
| Tob1 | 0 | 13.5 | 1064.66667 | 0 |

| | | | | |
|---|---|---|---|---|
| Tob1 | 0 | 7.5 | 717.333333 | 1 |
| Tox | 0 | 500 | 306.333333 | 0 |
| Tox | 0 | 599 | 190 | 1 |
| Tpbg | 0 | 16 | 4940.66667 | 0 |
| Tpbg | 0 | 15.5 | 3390 | 1 |
| Tpm1 | 0 | 420.5 | 122552 | 0 |
| Tpm1 | 1 | 1264.5 | 255247 | 1 |
| Trabd2b | 0 | 410 | 764 | 0 |
| Trabd2b | 0 | 430 | 432.333333 | 1 |
| Traf1 | 0 | 182.5 | 18 | 0 |
| Traf1 | 0 | 104 | 3.66666667 | 1 |
| Trib2 | 0 | 29 | 3256.33333 | 0 |
| Trib2 | 0 | 22.5 | 1410.33333 | 1 |
| Trim16 | 0 | 46.5 | 3436 | 0 |
| Trim16 | 0 | 48.5 | 2275 | 1 |
| Trim47 | 0 | 29.5 | 1913.33333 | 0 |
| Trim47 | 0 | 23 | 1093.66667 | 1 |
| Trim9 | 0 | 229 | 33 | 0 |
| Trim9 | 0 | 307 | 10.6666667 | 1 |
| Triobp | 0 | 164.5 | 2958.33333 | 0 |
| Triobp | 0 | 123.5 | 4439 | 1 |
| Trps1 | 1 | 689.5 | 1441.33333 | 0 |
| Trps1 | 0 | 1132.5 | 901.333333 | 1 |
| Trpv2 | 0 | 27.5 | 728.666667 | 0 |
| Trpv2 | 0 | 37 | 489.333333 | 1 |
| Tslp | 0 | 224 | 232 | 0 |
| Tslp | 0 | 249 | 46 | 1 |
| Tspan11 | 0 | 269 | 916.333333 | 0 |
| Tspan11 | 0 | 252 | 257 | 1 |
| Tspan12 | 0 | 440.5 | 139.333333 | 0 |
| Tspan12 | 0 | 555 | 234.666667 | 1 |
| Tspan18 | 0 | 159 | 717.666667 | 0 |
| Tspan18 | 0 | 166 | 1602.33333 | 1 |
| Tspan7 | 0 | 204 | 421.333333 | 0 |
| Tspan7 | 0 | 198 | 792 | 1 |
| Tspan8 | 0 | 142 | 13.6666667 | 0 |
| Tspan8 | 0 | 176 | 0.6667 | 1 |
| Ttc19 | 0 | 281.5 | 1405.66667 | 0 |
| Ttc19 | 1 | 332 | 1998 | 1 |
| Ttc28 | 0 | 164 | 6554.33333 | 0 |
| Ttc28 | 0 | 179.5 | 9248.33333 | 1 |
| Ttc39b | 0 | 238 | 2806.33333 | 0 |
| Ttc39b | 0 | 212 | 1785.33333 | 1 |
| Ttll11 | 0 | 184 | 504.666667 | 0 |
| Ttll11 | 0 | 174 | 2529.33333 | 1 |
| Ttll7 | 0 | 486 | 472.666667 | 0 |
| Ttll7 | 0 | 409 | 237.666667 | 1 |

| | | | | |
|---|---|---|---|---|
| Ttyh2 | 0 | 67 | 1435.33333 | 0 |
| Ttyh2 | 0 | 70.5 | 2315.33333 | 1 |
| Twist2 | 0 | 265 | 3313.66667 | 0 |
| Twist2 | 0 | 316 | 4506 | 1 |
| Txnrd1 | 0 | 107.5 | 17719 | 0 |
| Txnrd1 | 0 | 87 | 24582 | 1 |
| Txnrd2 | 0 | 20.5 | 2.66666667 | 0 |
| Txnrd2 | 0 | 31 | 11.3334 | 1 |
| U90926 | 0 | 83.5 | 708 | 0 |
| U90926 | 0 | 103.5 | 1495.66667 | 1 |
| Uap1l1 | 0 | 30.5 | 2274.33333 | 0 |
| Uap1l1 | 0 | 15.5 | 1635.33333 | 1 |
| Ubap1l | 0 | 40 | 33 | 0 |
| Ubap1l | 0 | 42.5 | 59.6666667 | 1 |
| Ube2ql1 | 0 | 57 | 118.333333 | 0 |
| Ube2ql1 | 0 | 60 | 444.666667 | 1 |
| Ucn2 | 0 | 42 | 55 | 0 |
| Ucn2 | 0 | 44 | 106.666667 | 1 |
| Ugt2b35 | 0 | 24 | 1.00006667 | 0 |
| Ugt2b35 | 0 | 25.5 | 11.3333333 | 1 |
| Unc13a | 0 | 43.5 | 166.333333 | 0 |
| Unc13a | 0 | 48 | 311.666667 | 1 |
| Unc13c | 0 | 745 | 175.333333 | 0 |
| Unc13c | 1 | 819.5 | 87.3333333 | 1 |
| Unc5d | 0 | 1859 | 60.3333333 | 0 |
| Unc5d | 0 | 1525 | 30.6666667 | 1 |
| Upk1b | 0 | 63 | 459.333333 | 0 |
| Upk1b | 0 | 65 | 296 | 1 |
| Ush1g | 0 | 7.5 | 32 | 0 |
| Ush1g | 0 | 11.5 | 64 | 1 |
| Usp43 | 0 | 213.5 | 14 | 0 |
| Usp43 | 0 | 248.5 | 216.333333 | 1 |
| Usp44 | 0 | 58 | 6.33333333 | 0 |
| Usp44 | 0 | 52.5 | 22.6666667 | 1 |
| Vcan | 0 | 551.5 | 22837.6667 | 0 |
| Vcan | 0 | 810 | 13517 | 1 |
| Vdr | 0 | 80 | 1012.33333 | 0 |
| Vdr | 0 | 70.5 | 705.333333 | 1 |
| Vegfd | 0 | 221 | 1330.33333 | 0 |
| Vegfd | 0 | 221.5 | 565.666667 | 1 |
| Vgf | 0 | 17.5 | 451 | 0 |
| Vgf | 0 | 14 | 276.666667 | 1 |
| Vill | 0 | 40.5 | 220 | 0 |
| Vill | 0 | 58 | 123.333333 | 1 |
| Vip | 0 | 32 | 27 | 0 |
| Vip | 0 | 19.5 | 7 | 1 |
| Vipr1 | 0 | 63.5 | 24.6666667 | 0 |

| | | | | |
|---|---|---|---|---|
| Vipr1 | 0 | 49 | 6 | 1 |
| Vmn1r40 | 0 | 3.5 | 0.0001 | 0 |
| Vmn1r40 | 0 | 3 | 11 | 1 |
| Vsir | 0 | 159 | 314.333333 | 0 |
| Vsir | 0 | 161 | 862.333333 | 1 |
| Wars | 0 | 41 | 4690.33333 | 0 |
| Wars | 0 | 51.5 | 7988 | 1 |
| Wdr95 | 0 | 88.5 | 29.6666667 | 0 |
| Wdr95 | 0 | 76 | 4.33336667 | 1 |
| Wee1 | 0 | 122 | 1408 | 0 |
| Wee1 | 0 | 162 | 2843 | 1 |
| Wipi1 | 0 | 81 | 4574 | 0 |
| Wipi1 | 0 | 73 | 3305 | 1 |
| Wisp1 | 0 | 156.5 | 31640 | 0 |
| Wisp1 | 0 | 126 | 14371.3333 | 1 |
| Wnk3 | 0 | 157.5 | 236.333333 | 0 |
| Wnk3 | 0 | 206.5 | 347.666667 | 1 |
| Wnk4 | 0 | 33 | 110.666667 | 0 |
| Wnk4 | 0 | 19.5 | 172.666667 | 1 |
| Wnt10a | 0 | 18.5 | 161.666667 | 0 |
| Wnt10a | 0 | 16 | 46.6666667 | 1 |
| Wnt10b | 0 | 14.5 | 98.3333333 | 0 |
| Wnt10b | 0 | 9 | 864.666667 | 1 |
| Wnt11 | 0 | 593 | 234.666667 | 0 |
| Wnt11 | 0 | 288.5 | 423 | 1 |
| Wnt2b | 0 | 69.5 | 129.333333 | 0 |
| Wnt2b | 0 | 77 | 1002 | 1 |
| Wnt4 | 0 | 45.5 | 1983 | 0 |
| Wnt4 | 0 | 40.5 | 1091.66667 | 1 |
| Wnt7a | 0 | 95 | 11.3333333 | 0 |
| Wnt7a | 0 | 80 | 3.00003333 | 1 |
| Wnt9a | 0 | 57 | 478.333333 | 0 |
| Wnt9a | 0 | 66 | 3220.66667 | 1 |
| Wt1 | 0 | 97 | 189.666667 | 0 |
| Wt1 | 0 | 92 | 285.666667 | 1 |
| Xaf1 | 0 | 22 | 254 | 0 |
| Xaf1 | 0 | 19 | 158 | 1 |
| Xdh | 0 | 147 | 176 | 0 |
| Xdh | 0 | 172.5 | 113.666667 | 1 |
| Zbtb18 | 0 | 28 | 953 | 0 |
| Zbtb18 | 0 | 35 | 1954.66667 | 1 |
| Zc3h12d | 0 | 35.5 | 22.6666667 | 0 |
| Zc3h12d | 0 | 30 | 5.66666667 | 1 |
| Zc3h8 | 0 | 39.5 | 115.666667 | 0 |
| Zc3h8 | 0 | 33 | 180 | 1 |
| Zdbf2 | 0 | 300.5 | 1046.33333 | 0 |
| Zdbf2 | 0 | 262 | 736.666667 | 1 |

| | | | | |
|---|---|---|---|---|
| Zdhhc14 | 0 | 306 | 379.333333 | 0 |
| Zdhhc14 | 0 | 279.5 | 231 | 1 |
| Zeb2 | 0 | 109.5 | 4258.33333 | 0 |
| Zeb2 | 0 | 121.5 | 2811 | 1 |
| Zeb2os | 0 | 5.5 | 56.6666667 | 0 |
| Zeb2os | 0 | 9.5 | 30.6666667 | 1 |
| Zfhx3 | 0 | 305.5 | 2070.33333 | 0 |
| Zfhx3 | 0 | 318.5 | 1437 | 1 |
| Zfp36 | 0 | 28.5 | 3196.33333 | 0 |
| Zfp36 | 0 | 26 | 4893.33333 | 1 |
| Zfp365 | 0 | 69 | 1115 | 0 |
| Zfp365 | 0 | 60.5 | 722 | 1 |
| Zfp36l2 | 0 | 272 | 15754.3333 | 0 |
| Zfp36l2 | 1 | 297 | 9940.33333 | 1 |
| Zfp521 | 1 | 1069 | 2130.66667 | 0 |
| Zfp521 | 0 | 711 | 1267.33333 | 1 |
| Zfp53 | 0 | 47 | 509 | 0 |
| Zfp53 | 0 | 54 | 763 | 1 |
| Zfp618 | 1 | 221 | 1033.66667 | 0 |
| Zfp618 | 0 | 263.5 | 1790 | 1 |
| Zfp804a | 0 | 248 | 31.6666667 | 0 |
| Zfp804a | 0 | 274.5 | 119 | 1 |
| Zfp820 | 0 | 67 | 228.666667 | 0 |
| Zfp820 | 0 | 67 | 377.333333 | 1 |
| Zfp827 | 1 | 375 | 2547.33333 | 0 |
| Zfp827 | 1 | 619 | 3816 | 1 |
| Zim1 | 0 | 44.5 | 664.333333 | 0 |
| Zim1 | 0 | 51.5 | 1396.33333 | 1 |
| Zswim6 | 1 | 256 | 1629.33333 | 0 |
| Zswim6 | 1 | 388 | 2329.66667 | 1 |
| Zswim7 | 0 | 36 | 248.333333 | 0 |
| Zswim7 | 0 | 37 | 402 | 1 |
